# Analysis of Post-Translational Modifications in the Urinary Proteome of Stroke-Prone Spontaneously Hypertensive Rats (SHRSP)

**DOI:** 10.1101/2025.05.25.655980

**Authors:** Yan Su, Youhe Gao

**Affiliations:** Gene Engineering Drug and Biotechnology Beijing Key Laboratory, College of Life Sciences, Beijing Normal University, Beijing 100875, China

**Keywords:** Hypertension, SHRSP, Urinary proteome, PTMs

## Abstract

Hypertension, one of the most prevalent chronic non-communicable diseases worldwide, is a primary risk factor for stroke and coronary heart disease. This study investigated dynamic changes in protein post-translational modifications (PTMs) and associated biological processes during hypertension progression by comparing urinary proteome PTMs between stroke-prone spontaneously hypertensive rats (SHRSP) and normal rats at 1-, 8-, and 14-15-month-old stages. Results revealed both stage-specific and shared PTM alterations across hypertensive phases. Notably, multiple differentially modified proteins identified here have been previously implicated in hypertension pathogenesis or progression. Randomized group validation demonstrated that more than 95% of differential modifications between the hypertensive and control groups exhibited non-random characteristics. Furthermore, proteins harboring differential PTMs were significantly enriched in hypertension-related biological pathways and KEGG pathways, including regulation of systemic arterial blood pressure, renin-angiotensin system (RAS), and complement and coagulation cascades. These results highlight distinct urinary proteome PTM profiles between SHRSP and normal rats, providing new insights into the mechanisms of hypertension.

## 1 Introduction

Hypertension, one of the most prevalent chronic non-communicable diseases globally, serves as a core risk factor for cardiovascular and cerebrovascular events^[1]^. As a multifactorial disorder associated with genetic and environmental factors, hypertension is a primary risk factor for stroke and coronary heart disease, with complications including heart failure, peripheral vascular disease, renal dysfunction, retinal hemorrhage, and others^[2][3]^. According to the Global Burden of Disease Study published in The Lancet Neurology in 2024, stroke-related deaths worldwide reached 7.3 million in 2021, making it the third leading cause of death after ischemic heart disease and COVID-19. Hypertension is the foremost risk factor for stroke, linked to more than half of global stroke occurrences, and hypertensive patients exhibit a significantly higher stroke risk than the normal population^[4][5]^. Sequelae of stroke, such as limb paralysis and cognitive dysfunction, not only severely impact patients’ quality of life but also impose substantial economic burdens on families and society^[6]^.

Urinary proteomics, as a non-invasive detection technique, can sensitively reflect the pathophysiological changes of the body by analyzing proteins and their modification states in urine, emerging as an important tool for discovering early biomarkers of diseases^[7][8]^. In recent years, with the development of liquid chromatography-mass spectrometry (LC-MS)-based proteomics technologies, high-precision identification of low-abundance modified proteins in urine has been achieved^[9]^. Post-translational modifications (PTMs) of proteins, as key molecular events regulating protein function, play important regulatory roles in pathological processes such as hypertensive vascular endothelial dysfunction, smooth muscle cell proliferation, and oxidative stress injury^[10][11]^. For example, abnormal protein phosphorylation can activate the renin-angiotensin system^[12]^. Histone acetylation modifications influence the proliferation and migration of vascular smooth muscle cells by regulating gene expression and cell functions during vascular remodeling, playing a crucial role in hypertension-related vascular lesions and further promoting the process of vascular remodeling^[13]^. The interaction between advanced glycation end products and their receptors activates cellular signaling pathways, upregulates inflammation- and oxidative stress-related factors, and exacerbates arterial aging and vascular pathological changes^[14]^.

Stroke-prone spontaneously hypertensive rats (SHRSP) are classic animal models for studying hypertensive stroke, characterized by spontaneous hypertension, progressive blood-brain barrier damage, and high stroke susceptibility, with a pathological process highly similar to human hypertensive stroke^[15]^. Studies have found that in the natural growth process of SHRSP rats, blood pressure begins to increase significantly at around 10 weeks, reaches hypertensive levels at 20-25 weeks, and subsequently develops cerebrovascular lesions gradually^[16]^. In this study, urinary samples from SHRSP rats at 1 month, 8 months, and 14-15 months of age were selected. By comparing the differences in urinary proteome PTMs with those of normal rats, we aimed to explore the protein modifications and biological process changes in different stages of hypertension, and preliminarily investigate the overall dynamic characteristics of protein PTMs during the course of hypertension.

## 2 Materials and Methods

### 2.1 Experimental Animals and Model Establishment

Relevant data of hypertensive and control group rats in this study were derived from published works of our laboratory^[17][18]^. Male stroke-prone SHRSP and male Sprague-Dawley (SD) rats were purchased from Beijing Vital River Laboratory Animal Technology Co., Ltd. All rats were housed in a standard environment at room temperature (22 ± 1) °C, with humidity of 65%-70% and a 12-h light-dark cycle. All experimental operations were approved by the Ethics Committee of the Institute of Basic Medicine, Chinese Academy of Medical Sciences, with approval numbers ACUC-A02-2014-007 and ACUC-A02-2015-004.

Urine was collected from SHRSP rats at 1 month, 8 months, and 14 months of age, and from SD rats at 30 days, 240 days, and 450 days of age using metabolic cages. The samples of 1-month-old SHRSP rats and 30-day-old SD rats were set as the experimental and control groups for the 1-month-old group, 8-month-old SHRSP rats and 240-day-old SD rats for the 8-month-old group, and 14-month-old SHRSP rats and 450-day-old SD rats for the 14-15-month-old group. The collected urine samples were centrifuged at 3000 r/min for 30 min and then stored at −80 °C.

### 2.2 Urine Sample Processing

Urinary protein extraction procedure: Urine samples were first thawed, then centrifuged at 12,000×g for 30 min at 4 °C, and the supernatant was transferred to a new EP tube. Four volumes of absolute ethanol were added, mixed, and precipitated at −20 °C for one day. The next day, the mixture was centrifuged at 12,000×g for 30 min at 4 °C, the supernatant was discarded, and the protein precipitate was resuspended in lysis buffer (containing 8 mol/L urea, 2 mol/L thiourea, 25 mmol/L dithiothreitol, 50 mmol/L). After centrifugation at 12,000×g for 30 min at 4 °C, the supernatant was retained, and protein concentration was determined by the Bradford method.

Protease digestion procedure: 100 μg of urinary protein sample was placed in an EP tube, and 25 mmol/L NH4HCO3 solution was added to make the total volume 200 μL as a standby sample. Dithiothreitol (DTT, Sigma) solution was then added to a final concentration of 20 mM, heated in a 95 °C metal bath for 10 min, cooled to room temperature, and iodoacetamide (IAA, Sigma) was added to a concentration of 50 mM, followed by incubation in the dark at room temperature for 40 min. After washing the membrane of a 10 kDa ultrafiltration tube (Pall, Port Washington, NY, USA), the prepared sample was added to the ultrafiltration tube and centrifuged at 14,000×g for 40 min at 18 °C. Then 200 μL of UA solution was added, centrifuged at 14,000×g for 30 min at 18 °C, repeated twice. Subsequently, 25 mmol/L NH4HCO3 solution was added, centrifuged at 14,000×g for 30 min at 18 °C, repeated twice. Finally, trypsin (Trypsin Gold, Promega, Fitchburg, WI, USA) was added at a ratio of 1:50 (trypsin:protein) and incubated overnight at 37 °C. After overnight incubation, the digested filtrate was collected by centrifugation, desalted using an Oasis HLB solid-phase extraction column, vacuum-dried, and the peptide lyophilate was stored at −80 °C.

### 2.3 LC-MS/MS Tandem Mass Spectrometry Analysis

Peptides were reconstituted in 0.1% formic acid in water to adjust the concentration to 0.5 μg/μL. Then, the peptides were separated by a Thermo EASY-Nlc1200 liquid chromatography system with the following parameters: elution time controlled at 90 min, and the elution gradient consisted of phase A (0.1% formic acid) and phase B (80% acetonitrile). Finally, an Orbitrap Fusion Lumos Tribird mass spectrometer was used to perform multiple data-dependent mass spectrometry data acquisitions on the separated peptides in a data-independent acquisition mode.

### 2.4 Open-pFind Unrestricted Modification Search

Using pFind Studio software (version 3.2.1, Institute of Computing Technology, Chinese Academy of Sciences), we performed unrestricted modification searches on the raw files of mass spectrometry results for each sample using default parameters. The database used was the Rattus norvegicus database downloaded from the UniProt website (https://www.uniprot.org), updated to September 2024. The instrument type was set to HCD-FTMS, the enzyme used was trypsin with full specificity, allowing up to 2 missed cleavages. The precursor mass tolerance was set to ±20 ppm, the fragment mass tolerance was also ±20 ppm, and an open search mode was selected. The screening criteria specified that the false discovery rate (FDR) at the peptide level should be less than 1%.

### 2.5 Bioinformatics Analysis of Protein PTMs

After performing unrestricted modification searches, post-translational modification PROTEIN files for each sample were obtained. Subsequently, a Python script named pFind_protein_contrast_script was downloaded from the GitHub platform (https://github.com/daheitu/scripts_for_pFind3_protocol.io) to summarize the post-translational modification identification results (i.e., PROTEIN files) of different samples. Group comparisons between the hypertensive and control groups were performed to screen for differential modifications. The screening criteria for differential modifications were: fold change (FC) ≥ 1.5 or ≤ 0.67, and two-tailed paired t-test P < 0.05. The proteins containing the screened differential modifications were queried for names and functions via the Uniprot website. Functional analysis of differential modifications was conducted by searching published literature in the PubMed database (https://pubmed.ncbi.nlm.nih.gov).

## 3 Results Analysis

### 3.1 Multivariate Statistical Analysis of Grouped Data

#### 3.1.1 Random Grouping

The control group samples (n=7) and experimental group samples (n=6) were randomly divided into two groups, yielding a total of 1,716 grouping types. For all random combination types, the average number of differential modifications across all random iterations was calculated using the same screening criteria. The ratio of the average number of differential modifications from random groupings to that from normal grouping was defined as the proportion of randomly generated differential modifications, as listed in Table 1. These results showed that randomly generated differential modifications were minimal, indicating high reliability in screening differential modifications. This suggests that most of the identified differential modifications were not randomly generated but represented actual differences between SHRSP rats and normal rats at different ages.

**Table 1.**
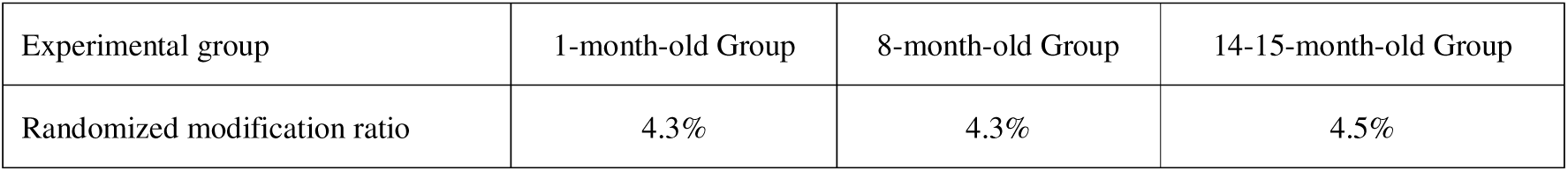
Proportion of Randomly Generated Differential Modifications from Random Grouping.

#### 3.1.2 Unsupervised Cluster Analysis and Principal Component Analysis

A total of 20,769 PTMs were identified in the 1-month-old group, 26,313 in the 8-month-old group, and 29,564 in the 14-15-month-old group (FDR<1%). Unsupervised cluster analysis (HCA) and principal component analysis (PCA) were performed on the total modifications of the three groups, with results shown in Figures 1–3. The results indicated significant differences in urinary protein PTMs between the experimental and control groups in all three age groups. PCA results showed that sample points of the experimental groups in the three groups were concentrated, indicating small within-group variation and similar sample data; in contrast, sample points of the control groups were more dispersed. This reflects that hypertension has a substantial impact on the rat body, and the effect of hypertension on urinary protein PTMs is evident before significant blood pressure elevation in rats.

**Figure 1.**
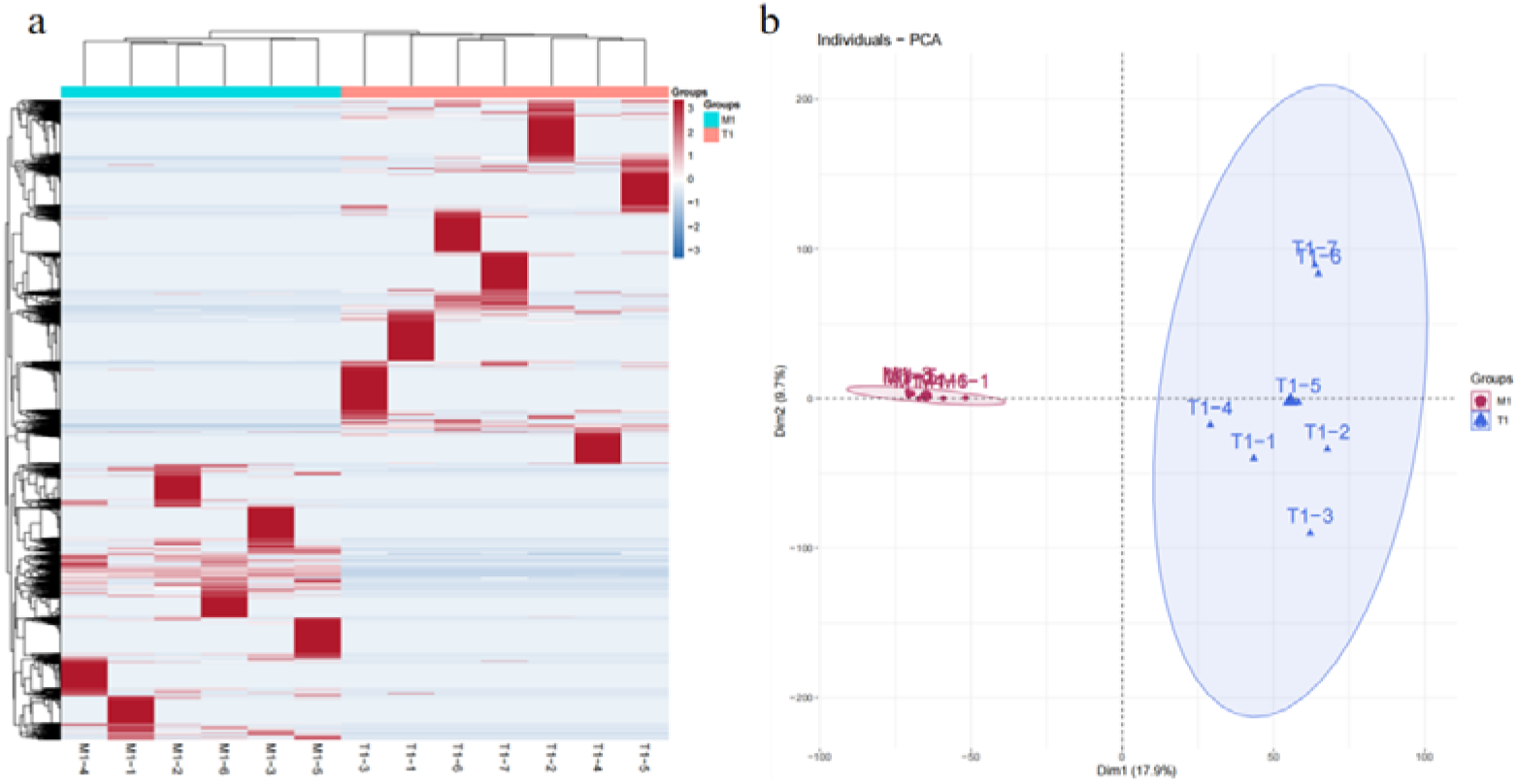
Unsupervised cluster analysis (Panel a) and principal component analysis (Panel b) of urinary protein post-translational modifications (PTMs) in 1-month-old rats. Group T1 is the control group, and M1 is the experimental group.

**Figure 2.**
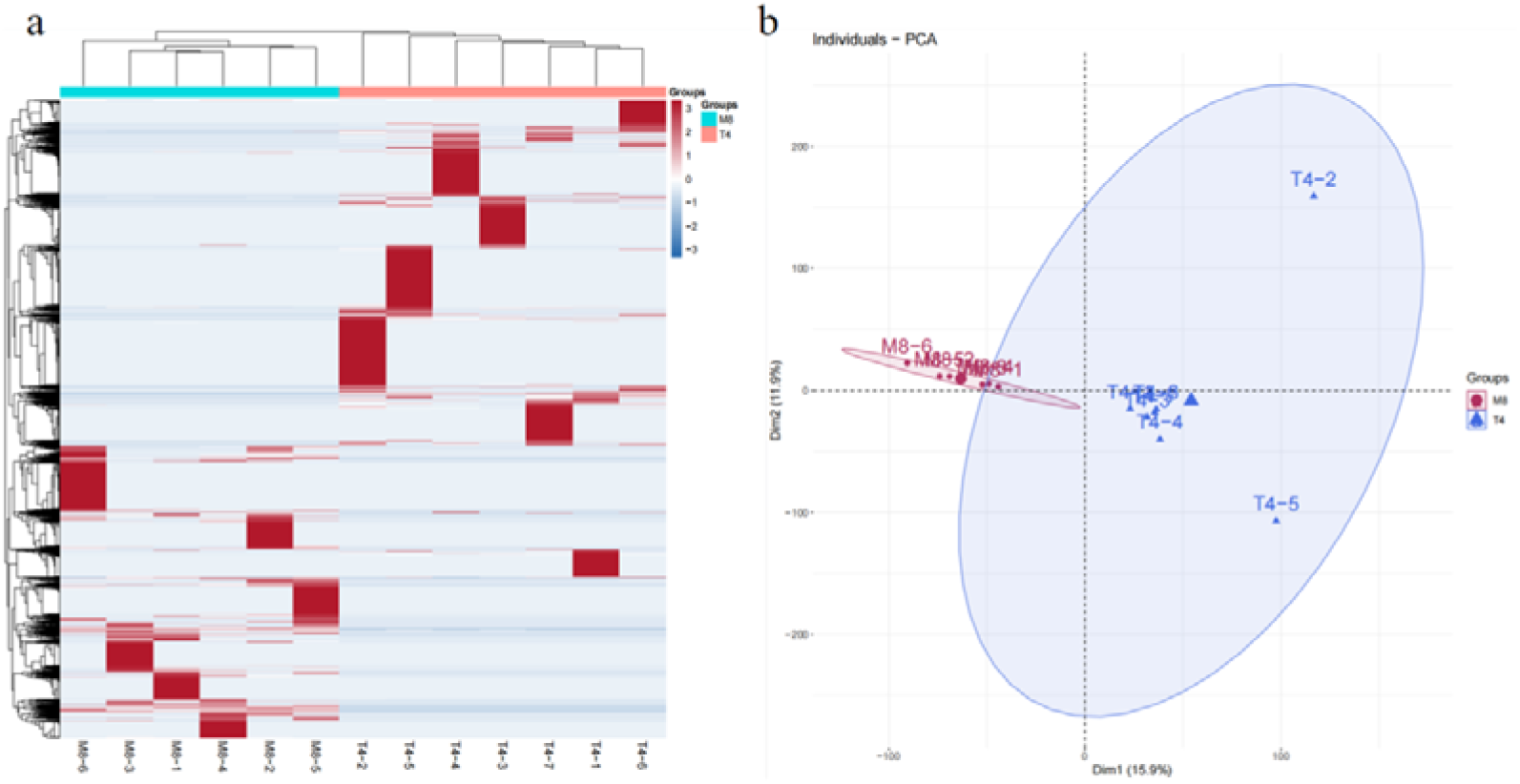
Unsupervised cluster analysis (Panel a) and principal component analysis (Panel b) of urinary protein PTMs in 8-month-old rats. Group T4 is the control group, and M8 is the experimental group.

**Figure 3.**
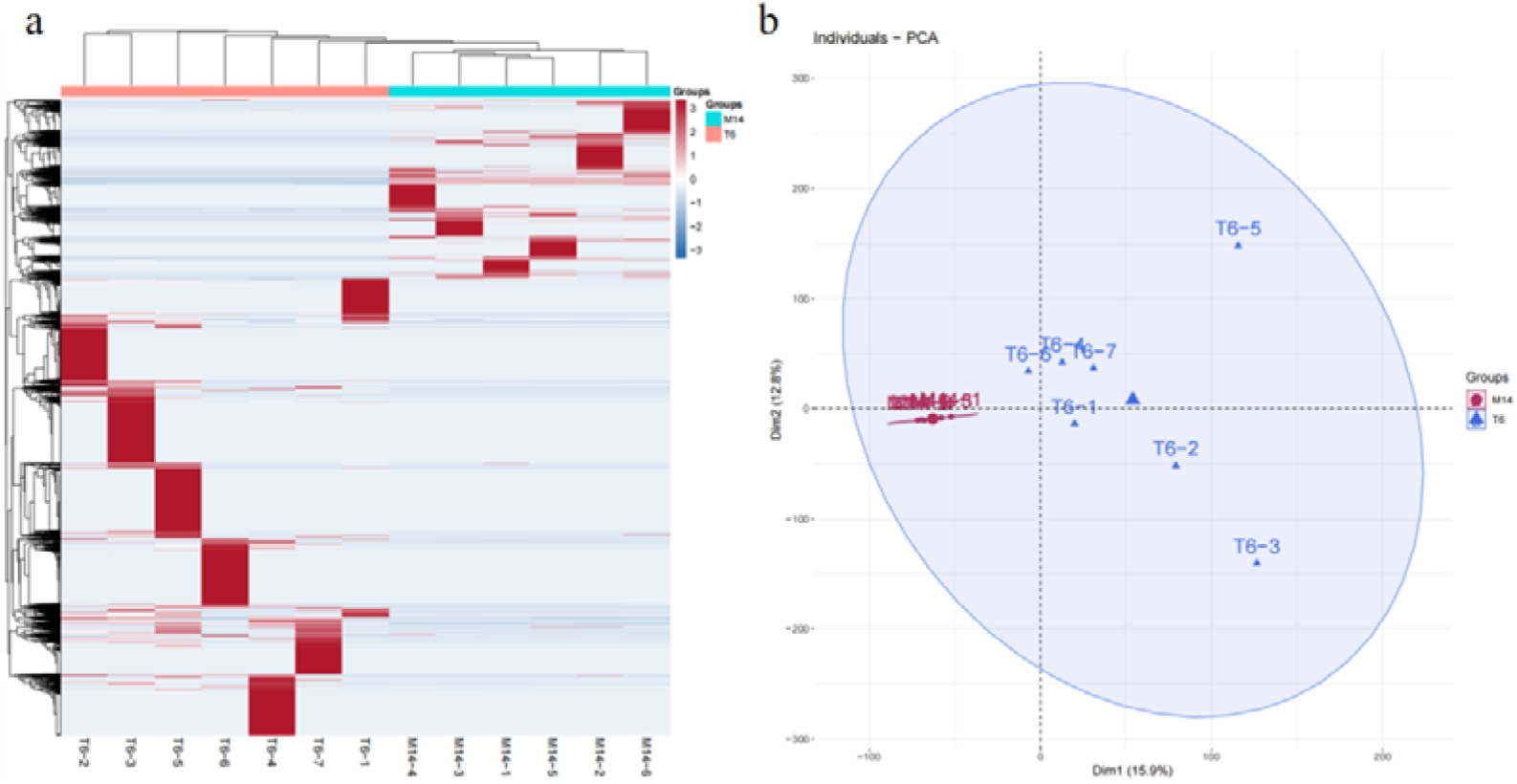
Unsupervised cluster analysis (Panel a) and principal component analysis (Panel b) of urinary protein PTMs in 14-15-month-old rats. Group T6 is the control group, and M14 is the experimental group.

### 3.2 Analysis of Urinary Proteome PTMs in Different Stages of Hypertension

PTMs in the experimental and control groups of the 1-month-old, 8-month-old, and 14-15-month-old groups were compared respectively, with the screening criteria for differential modifications set as: FC ≥ 1.5 or ≤ 0.67, and two-tailed paired t-test P < 0.05. The results showed that a total of 1,590 differential modifications were identified in the 1-month-old group, involving 204 proteins with differential modifications, and detailed information is listed in Supplementary Table 1. The 8-month-old group identified 1,297 differential modifications involving 126 proteins, with details in Supplementary Table 2. The 14-15-month-old group identified 1,596 differential modifications involving 127 proteins, as listed in Supplementary Table 3. The types of differential modifications identified in the three groups were 318, 316, and 349, respectively, with detailed information in Supplementary Table 4.

We constructed Venn diagrams for the differential modifications, proteins with differential modifications, and types of differential modifications identified in the 1-month-old, 8-month-old, and 14-15-month-old groups, as shown in Figure 4. The results indicated that the three groups had numerous unique differential modifications and corresponding proteins, suggesting that different stages of hypertension development are associated with specific modification changes. Additionally, common differential modifications and corresponding proteins existed among the three groups, implying that certain shared modification mechanisms might underlie different hypertensive stages. The specific and shared differential modifications of the three groups are discussed separately below.

**Figure 4.**
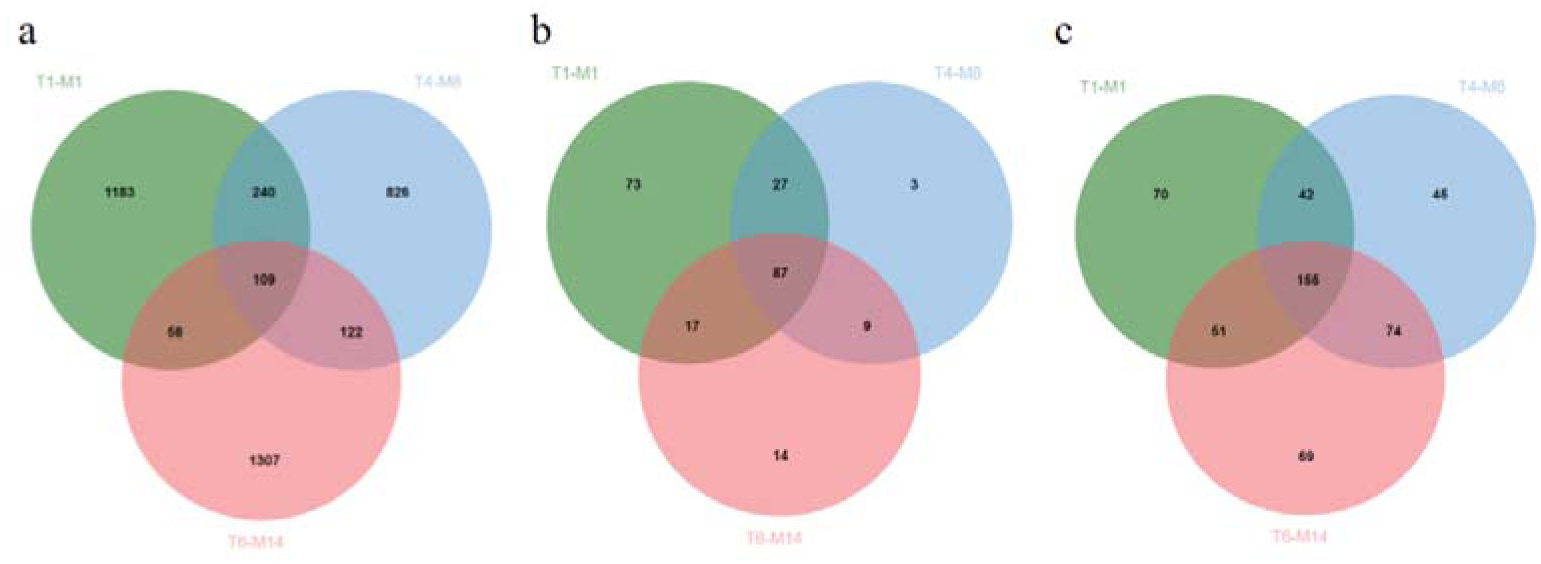
Venn diagrams of urinary protein PTMs in 1-month-old (T1-M1), 8-month-old (T4-M8), and 14-15-month-old (T6-M14) rats. a. Venn diagram of differential modifications; b. Venn diagram of proteins with differential modifications; c. Venn diagram of types of differential PTMs.

#### 3.2.1 Analysis of Unique Differential PTMs in Urinary Proteome of 1-month-old Rats

Literature searches in the PubMed database for proteins containing identified PTMs showed that some proteins have been reported to be associated with hypertension in previous studies.

For proteins with PTMs showing “from present to absent” changes:

Q64230, Meprin A subunit alpha (P=3.72E-05). Its binding to heparan sulfate is crucial for protecting endothelial cells from inflammatory cell extravasation. In patients with idiopathic pulmonary arterial hypertension, reduced binding of Meprin A subunit alpha occurs due to decreased endothelial heparan sulfate levels, potentially leading to pathological vascular remodeling^[19]^.

P07861, Neprilysin (P=1.67E-04). This protein converts angiotensin-1 to angiotensin-(1-7), a process that activates MAS-related G protein-coupled receptors. MAS receptors antagonize angiotensin type 1 receptors, reducing reactive oxygen species and inflammation to improve renal injury^[20]^.

Q9ESG3, Collectrin (P=2.52E-03). Studies have linked Collectrin mRNA and protein expression in renal tissue to increased blood pressure. Global Collectrin deficiency leads to endothelial dysfunction, enhanced salt sensitivity, and hypertension^[21][22]^.

O55006, Protein RoBo-1 (P=5.56E-03). Robo receptors 1 and 4 and their ligand Slit2 are novel regulators of endothelial cell function. Compared with human umbilical vein endothelial cells, pulmonary artery endothelial cells from patients with pulmonary arterial hypertension exhibit increased Robo receptor 1 expression, decreased Robo receptor 4 expression, and resistance to the anti-inflammatory effect of Slit2, which may be associated with pulmonary arterial hypertension^[23]^.

Q63691, Monocyte differentiation antigen CD14 (P=1.05E-02). Studies show increased numbers of monocytes and macrophages around cerebral blood vessels in hypertensive and aged rats, which may enhance endothelial cell responsiveness to mediators, shifting from an anticoagulant to a procoagulant surface^[24]^. As a monocyte marker, CD14 may be indirectly associated with hypertension and vascular diseases.

P53790, Sodium/glucose cotransporter 1 (P=1.11E-02), potentially involved in sodium transport disorders observed in hypertension. This protein is transcriptionally regulated in the intestine of hypertensive rats, leading to reduced molecular weight^[25]^.

P04642, L-lactate dehydrogenase A chain (P=1.11E-02). Its mediated lactate production promotes pulmonary vascular remodeling in pulmonary arterial hypertension by activating the Akt signaling pathway, indicating the protein’s potential role in metabolic reprogramming and vascular remodeling for regulating pulmonary arterial hypertension^[26]^.

P50123, Glutamyl aminopeptidase (P=2.05E-02), participating in the catabolic pathway of the renin-angiotensin system to form angiotensin Ⅲ, which is involved in blood pressure regulation and angiogenesis^[27]^.

Q9Z0W7, Chloride intracellular channel protein 4 (P=3.33E-02), overexpressed in the renal proximal tubules of SHRSP. This suggests significant changes in renal parenchyma during the early stage of hypertension development, with this protein potentially serving as an early marker of tubule alterations caused by increased blood pressure^[28]^.

P0DP31, Calmodulin-3 (P=4.40E-02), which regulates eNOS and iNOS activities and participates in related signaling pathway modulation in vitro vessels of portal hypertensive rats, influencing NO production and thereby leading to hyporeactivity of vessels to vasoconstrictors^[29]^.

P29534, Vascular cell adhesion protein 1 (P=4.65E-02), associated with the occurrence and development of hypertension. Studies have shown increased levels of this protein in patients with essential hypertension, which correlates with norepinephrine concentration^[30]^.

P11232, Thioredoxin (P=4.72E-02), playing an important role in age-related hypertension. In mice with higher human thioredoxin levels, age-related hypertension is reversed and maintained at lower levels for a prolonged period. Injection of human recombinant thioredoxin reduces hypertension in aged wild-type mice^[31]^.

For proteins with PTMs that changed from absent to present:

P01015, Angiotensinogen (P=8.04E-04), related to hypertension and vascular diseases. Studies have shown that angiotensinogen gene knockout delays and alleviates cold-induced hypertension^[32]^.

P04937, Fibronectin (P=2.27E-03). Research has found lower plasma fibronectin levels in patients with portal hypertension, possibly due to increased consumption by an enlarged spleen^[33]^.

P04903, Glutathione S-transferase alpha-2 (P=3.35E-02). Studies have indicated that glutathione S-transferase variants are risk factors for essential hypertension in Italian patients, with the Glutathione S-transferase T1-null phenotype significantly more frequent in hypertensive patients than in normotensive participants^[34]^.

For proteins with PTMs showing more than 10-fold upregulation/downregulation, P12020, Cysteine-rich secretory protein 1 (FC=10.89, P=2.30E-02). Some members of the cysteine-rich secretory protein family exhibit anti-angiogenic activity. For example, cysteine-rich oral gland proteins have anti-angiogenic activity, which can induce apoptosis of vascular endothelial cells by affecting mitochondrial-dependent pathways and inhibit proliferation and adhesion of vascular endothelial cells^[35]^.

#### 3.2.2 Analysis of Unique Differential PTMs in Urinary Proteome of 8-Month-Old Rats

Literature searches for proteins containing identified PTMs were conducted in the Pubmed database, and some proteins showed correlation with hypertension in existing studies.

For proteins with PTMs that changed from present to absent:

Q99PS8, Histidine-rich glycoprotein (P=1.04E-02). As a plasma protein, it has been reported to regulate coagulation, fibrinolysis, and angiogenesis by binding to heparin, fibrinogen, thrombospondin, plasminogen, etc., and interacting with various cell types such as endothelial cells, red blood cells, neutrophils, and platelets^[36]^.

Q63678, Zinc-alpha-2-glycoprotein (P=1.62E-02). It may serve as an early biomarker in diabetic nephropathy. Urinary and serum Zinc-alpha-2-glycoprotein helps detect diabetic nephropathy in type 2 diabetic patients earlier than microalbuminuria^[37]^.

P04916, Retinol-binding protein 4 (P=2.78E-02). Studies have found that changes in the binding saturation of this protein with retinol may link renal dysfunction and insulin resistance to atherosclerosis^[38]^.

P23785, Progranulin (P=4.97E-02). Circulating Progranulin regulates vascular tone and blood pressure by activating EphrinA2 and Sortilin1 receptors as well as endothelial nitric oxide synthase. Mice deficient in Progranulin exhibit increased blood pressure, which can be restored by recombinant Progranulin supplementation^[39]^.

Q63691, Monocyte differentiation antigen CD14 (P=4.65E-02); P11232, Thioredoxin (P=3.83E-02); and P0DP31, Calmodulin-3 (P=2.78E-02) are proteins with PTMs shared with the 1-month-old group, and detailed analysis is provided in section 3.2.1.

For proteins with PTMs that changed from absent to present:

P01026, Complement C3 (P=1.66E-03), is an important factor in the pathogenesis of hypertension. In SHRSP, Complement C3 is highly expressed in mesenchymal tissues, inducing the synthetic phenotype and overgrowth of mesenchymal cells through maintaining dedifferentiated cells. Increased C3 expression induces salt-sensitive hypertension and activates the renin-angiotensin system. C3 also participates in epithelial-mesenchymal transition and epithelial cell dedifferentiation in the unilateral ureteral obstruction kidney, leading to increased blood pressure^[40]^.

P02767, Transthyretin (P=3.16E-02). SHRSP exhibit more Transthyretin immunoreactive accumulation in the choroid plexus, with lower Transthyretin monomer expression in cerebrospinal fluid and higher expression in blood. This indicates that hypertension alters the in vivo distribution and expression form of Transthyretin, suggesting a mutual influence between Transthyretin and hypertension^[41]^.

#### 3.2.3 Analysis of Unique Differential PTMs in Urinary Proteome of 14-15-Month-Old Rats

Literature searches for proteins containing identified PTMs were conducted in the Pubmed database, and some proteins showed correlation with hypertension in existing studies.

For proteins with PTMs that changed from present to absent:

P08721, Osteopontin (P=2.78E-02). Its expression is a powerful predictor of cardiovascular diseases, independent of traditional risk factors. Acute increases in Osteopontin in cardiovascular diseases are protective, alleviating vascular calcification and promoting post-ischemic neovascularization; chronic increases in Osteopontin are clinically associated with an increased risk of major adverse cardiovascular events^[42]^.

P08649, Complement C4 (P=3.75E-02), and P01026, Complement C3 (P=2.03E-02, FC=8.56), both play roles in the complement activation pathway. After ischemic stroke, Complement C3 exhibits acute elevation, confirming complement activation, which may be one of the mechanisms leading to further brain tissue damage. Complement C3 and Complement C4 are unique predictors of poor outcomes in diabetic stroke, associated with adverse clinical outcomes in stroke patients^[43][44]^.

For proteins with PTMs that changed from absent to present:

P04639, Apolipoprotein A-I (P=6.72E-04), has anti-inflammatory effects, inhibiting the production of inflammatory factors and activation of inflammatory cells, thereby reducing inflammatory damage to vascular endothelial cells^[45]^. Studies have shown that apoA-I infusion promotes cholesterol metabolism and excretion, reducing the risk of atherosclerosis^[46]^.

P08934, Kininogen-1 (P=1.89E-03), exerts anti-angiogenic properties to inhibit tumor cell growth. Overexpression of this protein suppresses cell viability and angiogenesis in glioma cells^[47]^.

For proteins with PTMs showing more than 10-fold upregulation/downregulation, P02680, Fibrinogen gamma chain (FC=12.25, P=2.15E-02), plays a key role in fibrin clot formation, platelet aggregation, and wound healing. These binding interactions may also contribute to pathophysiological processes such as inflammation and thrombosis^[48]^.

Additionally, P04642, L-lactate dehydrogenase A chain (P=1.34E-03); P01015, Angiotensinogen (P=1.99E-02); and P50123, Glutamyl aminopeptidase (P=2.90E-02) are proteins with PTMs shared with the 1-month-old group, with detailed analysis provided in section 3.2.1. P02767, Transthyretin (P=6.16E-06); Q63678, Zinc-alpha-2-glycoprotein (P=6.97E-05); and Q99PS8, Histidine-rich glycoprotein (P=5.30E-03) are proteins with PTMs shared with the 8-month-old group, with detailed analysis provided in section 3.2.2.

#### 3.2.4 Analysis of Common Differential PTMs in Urinary Proteome of 1-Month-Old, 8-Month-Old, and 14-15-Month-Old Rats

By comparing the FC of common differential PTMs among the three groups, specific information is listed in Supplementary Table 5. PTMs of certain proteins showed FC changes in the 1-month-old, 8-month-old, and 14-15-month-old groups, which may reflect the dynamic regulation of protein functions during hypertension progression and the pathophysiological mechanisms at different stages of hypertension development.

For modifications with gradually increasing FC in the 1-month-old, 8-month-old, and 14-15-month-old groups:

Cystatin-related protein 2 had a modification of “4,Carbamidomethyl[C]” on the sequence “LVNCPFEEQTEQLKR”, with FC changes of 134.4, 179.1, and #DIV/0! (where #DIV/0! indicates a “from absent to present” change). Studies have found that serum cystatin C, urinary microalbumin, and β2-microglobulin can effectively reflect renal injury, and combined detection has high accuracy for diagnosing early renal injury in gestational hypertension syndrome. Cystatin-related protein 2, belonging to the cystatin family like serum cystatin C, may share structural and functional similarities^[49]^.

Hemopexin had a modification of “6,Carbamidomethyl[C]” on the sequence “DYFISCPGR”, with FC changes of 1.7, 5.6, and 5.9. When hemolysis occurs, hemoglobin degradation releases free heme, which has strong oxidative activity, triggering oxidative stress, damaging vascular endothelial cells, promoting vasoconstriction, and thus increasing blood pressure. Hemopexin tightly binds to free heme to form a complex,reducing the adverse effects of free heme and indirectly stabilizing blood pressure^[50]^.

Serotransferrin had a modification of “6,Carbamidomethyl[C]” on the sequence “TSYQDCIK”, with FC changes of 0.02, 5.0, and 5.4. Serotransferrin, the main iron-containing protein in plasma, is responsible for transporting iron absorbed by the digestive tract and released from red blood cell degradation. Studies have shown that Serotransferrin levels are positively correlated with blood pressure and hypertension^[51]^.

Albumin had a modification of “18,Carbamidomethyl[C]” on the sequence “DDNPNLPPFQRPEAEAMCTSFQENPTSFLGHYLHEVAR”, with FC changes of 4.6, 6.7, and 9.3. In this study, multiple modification sites of Albumin were identified: 84 in the 1-month-old group, 392 in the 8-month-old group, and 218 in the 14-15-month-old group. In the 14-15-month-old group, the modification “2,Hex(3)[N]” on the sequence “DNCFATEGPNLVAR” showed a fold change of 525. Existing studies have shown that Albumin has a nonlinear relationship with hypertension incidence risk, and moderately increasing and highly stable serum albumin concentration trajectories are associated with reduced hypertension risk in specific populations^[52][53]^. Additionally, studies suggest that hypertensive patients should routinely detect urinary albumin/creatinine ratio to identify high-risk individuals early, strengthen treatment, and reduce the risk of cardio-renal events^[54]^.

Clusterin had a modification of “0,Succinyl_2H(4)[AnyN-term]” on the sequence “TQQYNELLHSLQSK”, with FC changes of 2.8, 6.2, and 18.7. Plasminogen had a modification of “7,Carbamidomethyl[C]” on the sequence “TPENFPCK”, with FC changes of 2.0, 3.4, and 3.5. These two modifications have not yet been shown to be associated with hypertension in existing studies.

For modifications with gradually decreasing FC in the 1-month-old, 8-month-old, and 14-15-month-old groups: Major urinary protein had a modification of “2,CarbamidomethylDTT[C]” on the sequence “LCEAHGITR”, with FC changes of 73.0, 25.9, and 6.0. Alpha-1-acid glycoprotein had a modification of “0,Gln->pyro-Glu[AnyN-termQ]” on the sequence “QQLELEKETK”, with FC changes of 3.5, 2.6, and 2.1. Prosaposin had modifications of “4,Carbamidomethyl[C];7,Carbamidomethyl[C]” on the sequence “SLPCDICK”, with FC changes of 1.7, 0.1, and 0. These three modifications have not yet been shown to be associated with hypertension in existing studies.

Furthermore, the 1-month-old, 8-month-old, and 14-15-month-old groups shared differentially modified proteins, which underwent PTMs at different sites or types in different groups, indicating that these proteins may serve as hub molecules continuously regulated during hypertension development. Some proteins have been shown to be associated with hypertension in existing studies:

Fetuin-B, a cytokine regulating lipid metabolism, was found to have significantly elevated serum levels in patients with essential hypertension, serving as an independent risk factor for essential hypertension^[55]^.

C-reactive protein, an inflammatory marker associated with increased cardiovascular risk, is typically elevated in hypertensive patients. Regular, sustained aerobic exercise and other training can significantly reduce C-reactive protein levels in hypertensive patients, alleviating inflammation to manage hypertension^[56]^.

Meprin A subunit beta, where Meprin A plays an important role in vascular remodeling in pulmonary arterial hypertension, may represent a novel molecule targetable for pulmonary arterial hypertension. Remodeled pulmonary arteries from explanted lungs of patients with idiopathic pulmonary arterial hypertension exhibit significant Meprin A expression^[57]^.

Serum amyloid P-component prevents cardiac remodeling by inhibiting the recruitment of profibrotic macrophages in hypertensive heart disease, and its reduced level is associated with the progression of diastolic dysfunction^[58]^.

Alpha-1-antiproteinase may serve as a potential biomarker for monitoring hypertension. Studies have found altered expression of serum Alpha-1-antiproteinase in SHRSP, which is eliminated by captopril treatment^[59]^.

Dipeptidyl peptidase 4: In a rat model of portal hypertension, Dipeptidyl peptidase 4 inhibitors significantly reduce portal pressure, upregulate nitric oxide synthase expression, and exert protective effects in reversing vascular aging^[60]^.

T-kininogen 2, Submandibular glandular kallikrein-9, Prostatic glandular kallikrein-6, Glandular kallikrein-7, and Kallikrein-1 are related to the kallikrein-kinin system. Studies have shown that when the normal function of the kallikrein-kinin system is inhibited, blood pressure increases, indicating its important role in blood pressure regulation^[61]^.

Cystatin-C has predictive value for hypertension control. Hypertensive patients with uncontrolled blood pressure have higher Cystatin-C levels than those with controlled blood pressure, and elevated serum cystatin C levels increase the likelihood of uncontrolled blood pressure in hypertensive patients, suggesting that serum cystatin C concentration can detect mild renal dysfunction in patients with poorly controlled blood pressure and normal serum creatinine levels^[62]^.

Prostatic spermine-binding protein: Polyamines such as spermine are significantly elevated in the plasma of patients with idiopathic pulmonary arterial hypertension and correlate with disease severity. Exogenous spermine promotes proliferation and migration of pulmonary arterial smooth muscle cells, exacerbating pulmonary vascular remodeling. Binding of prostatic polyamine-binding protein to spermine may play a role in this process^[63]^.

Prothrombin has been less studied in relation to hypertension, but existing studies have shown that prothrombin fragment F1+2 can serve as a sensitive marker for thrombosis, and hypertensive patients have an increased risk of thrombosis. Lifestyle factors in hypertensive patients, such as lack of exercise and smoking, may affect prothrombin fragment F1+2 levels^[64]^.

### 3.3 Biological Pathway Analysis of Urinary Proteome PTMs in Different Stages of Hypertension

The DAVID database was used to perform enrichment analysis of biological processes (BP) and KEGG pathways for proteins with differential modifications identified in the 1-month-old, 8-month-old, and 14-15-month-old groups. The results showed that 246 BP pathways were significantly enriched in the 1-month-old group, 182 in the 8-month-old group, and 155 in the 14-15-month-old group, with detailed information listed in Supplementary Tables 6, 7, and 8, respectively.

Among these, four biological pathways—zymogen activation, proteolysis, regulation of systemic arterial blood pressure, and acute-phase response—were highly significantly enriched in all three groups, ranking among the top six with the smallest P-values in each group’s enrichment results. This suggests that these pathways play key regulatory roles as core biological processes throughout the continuous pathological process of hypertension.

KEGG pathway enrichment analysis showed that the Renin-angiotensin system (RAS) and Complement and coagulation cascades were highly significantly enriched in all three groups, ranking among the top five with the smallest P-values in each group’s enrichment results. Numerous studies have shown that RAS is a key system regulating blood pressure and fluid balance, playing an important role in the occurrence, development, and treatment of hypertension^[65][66]^. The Complement and coagulation cascades also play a role in the hypertensive process, as complement activation fragments such as C3a and C5a exacerbate the pathological process of hypertension by promoting vascular inflammation and tissue damage^[67]^.

## 4 Discussion

This study investigated the dynamic characteristics of protein modifications during the course of hypertension by comparing urinary proteome PTMs between SHRSP rats of different ages and normal SD rats. The results showed significant proteomic PTM differences between the hypertensive and normal groups at 1 month, 8 months, and 14-15 months of age. Each stage exhibited both unique hypertension-related differentially modified proteins and biological pathways, as well as numerous common changes. Combined with the disease development pattern of SHRSP rats, their blood pressure begins to increase significantly at approximately 10 weeks of natural growth and reaches hypertensive levels at 20-25 weeks. Therefore, 1 month of age corresponds to the normotensive compensation period, during which the body may maintain homeostasis via specific protein modifications during early pathological changes before blood pressure elevation; 8 months corresponds to the early-stage vascular remodeling phase of hypertension, where altered protein modifications may participate in regulating the molecular processes of vascular structural remodeling; 14-15 months corresponds to the decompensated hypertension and high-risk stroke period, where related protein modifications may reflect the decline of compensatory mechanisms and increased complication risks^[16][68]^. The correlation between modification characteristics presented by urinary proteomics and the course of hypertension provides important clues for exploring novel biomarkers that can be used to warn different stages of hypertensive diseases.

It should be noted that the functional annotation of most identified PTMs in this study regarding their roles in hypertensive pathology remains incomplete, which objectively restricts the depth of analysis for specific modification events. This indicates that future studies should combine targeted modified proteomics with functional experiments to further clarify the impact of key modification sites on disease risk. Additionally, this study has some limitations. Dynamic blood pressure monitoring was not performed on the experimental and control rats, primarily because blood pressure measurement may induce stress responses in rats, leading to blood pressure fluctuations and compromising the stability of the urinary proteome; thus, grouping was primarily based on existing literature on SHRSP rats. Although the sample size of this study was small, random grouping verification showed that more than 95% of the screened differential modifications exhibited non-random characteristics, which significantly reduced the possibility of small-sample study results being interfered with by accidental factors. Finally, future research could evaluate the reversal effect of antihypertensive drugs on urinary PTM characteristics and attempt to provide monitoring tools for personalized medicine.

## 5 Conclusion

In summary, urinary protein PTMs differ between SHRSP rats and normal rats. This suggests that urinary proteomics can be used to study the mechanisms of hypertension, providing a new perspective for evaluating the course of hypertension.

