## Appendix for "Analysis of Post-Translational Modifications in the Urinary Proteome of Stroke-Prone Spontaneously Hypertensive Rats (SHRSP)"

Appendix Table 1 Differential modifications screened by FC ≥ 1.5 or ≤ 0.67 with P < 0.05 in the 1-month-old group

| Uniprot ID | Peptide | Modification | Control group | Experimental group | Flod change | P value |
| --- | --- | --- | --- | --- | --- | --- |
| P02761 | TPEDGEYFVEYDGGNTFTILK | 10,Cation_Na[E]; | 792.4 | 0.0 | 0.0 | 1.92E-07 |
| P02770 | FPNAEFAEITK | 3,Deamidated[N]; | 102.4 | 0.0 | 0.0 | 4.37E-06 |
| Q63257 | VLGDPTCFSDYIR | 7,Carbamidomethyl[C]; | 45.0 | 0.0 | 0.0 | 1.15E-05 |
| P83121 | CQTYPDEICAWVVVTTR | 1,Carbamidomethyl[C];9,Carbamidomethyl[C]; | 610.7 | 0.0 | 0.0 | 1.58E-05 |
| P81828 | CQTYPDEICAWVVVTTR | 1,Carbamidomethyl[C];9,Carbamidomethyl[C]; | 610.7 | 0.0 | 0.0 | 1.58E-05 |
| P81827 | CQTYPDEICAWVVVTTR | 1,Carbamidomethyl[C];9,Carbamidomethyl[C]; | 610.7 | 0.0 | 0.0 | 1.58E-05 |
| Q64230 | TNICGMIQGTR | 4,Carbamidomethyl[C]; | 71.6 | 0.0 | 0.0 | 3.72E-05 |
| P81828 | TSFDSTGFCHVGR | 9,Carbamidomethyl[C]; | 124.7 | 0.0 | 0.0 | 6.35E-05 |
| P02770 | CCTLPEAQR | 1,Carbamidomethyl[C];2,Carbamidomethyl[C]; | 30.9 | 0.0 | 0.0 | 6.97E-05 |
| Q04807 | LTMLSQAVQK | 3,Oxidation[M]; | 21.0 | 0.0 | 0.0 | 7.00E-05 |
| P12788 | HPEYNKDTLDNDIMLIK | 6,TMT2plex[K]; | 31.7 | 0.0 | 0.0 | 8.50E-05 |
| P07314 | TPALCEVFCR | 5,Carbamidomethyl[C];9,Carbamidomethyl[C]; | 41.1 | 0.0 | 0.0 | 1.38E-04 |
| P02770 | QEPERNECFLQHK | 8,Carbamidomethyl[C]; | 21.9 | 0.0 | 0.0 | 1.53E-04 |
| P07861 | DYYECTGIYK | 5,Carbamidomethyl[C]; | 18.0 | 0.0 | 0.0 | 1.67E-04 |
| P22057 | QGHDTVQPNFQQDK | 0,Gln->pyro-Glu[AnyN-termQ]; | 6.4 | 0.0 | 0.0 | 1.75E-04 |
| P07522 | NLDPASVPPR | 0,Ethylphosphate[AnyN-term]; | 12.4 | 0.0 | 0.0 | 1.84E-04 |
| Q6P7A9 | AVPTQCDVTPNSR | 6,Carbamidomethyl[C]; | 10.3 | 0.0 | 0.0 | 2.03E-04 |
| P02770 | YNEVLTQCCTESDK | 8,Carbamidomethyl[C];9,Carbamidomethyl[C]; | 132.0 | 0.0 | 0.0 | 2.43E-04 |
| P20059 | CNADPGLSALLSDHR | 1,Carbamidomethyl[C]; | 30.4 | 0.0 | 0.0 | 2.71E-04 |
| P83121 | CQTYPDEICAWVVVTTR | 7,Cation_Na[E];9,Carbamidomethyl[C]; | 249.0 | 0.0 | 0.0 | 3.58E-04 |
| P81828 | CQTYPDEICAWVVVTTR | 7,Cation_Na[E];9,Carbamidomethyl[C]; | 249.0 | 0.0 | 0.0 | 3.58E-04 |
| P81827 | CQTYPDEICAWVVVTTR | 7,Cation_Na[E];9,Carbamidomethyl[C]; | 249.0 | 0.0 | 0.0 | 3.58E-04 |
| P63018 | LLQDFFNGK | 7,Deamidated[N]; | 24.9 | 0.0 | 0.0 | 3.60E-04 |
| P14659 | LLQDFFNGK | 7,Deamidated[N]; | 24.9 | 0.0 | 0.0 | 3.60E-04 |
| Q08420 | MGLATSLAGPHSILGR | 1,Oxidation[M]; | 16.7 | 0.0 | 0.0 | 3.86E-04 |
| P83121 | CQTYPDEICAWVVVTTR | 1,maleimide[C]; | 134.6 | 0.0 | 0.0 | 3.95E-04 |
| P81828 | CQTYPDEICAWVVVTTR | 1,maleimide[C]; | 134.6 | 0.0 | 0.0 | 3.95E-04 |
| P81827 | CQTYPDEICAWVVVTTR | 1,maleimide[C]; | 134.6 | 0.0 | 0.0 | 3.95E-04 |
| Q01177 | CTTPPPPPGPTYQCLK | 1,Carbamidomethyl[C];14,Carbamidomethyl[C]; | 8.6 | 0.0 | 0.0 | 4.01E-04 |
| P01836 | RDGVLDSVTDQDSK | 14,Lys->Asp[K]; | 12.9 | 0.0 | 0.0 | 4.01E-04 |
| P01835 | RDGVLDSVTDQDSK | 14,Lys->Asp[K]; | 12.9 | 0.0 | 0.0 | 4.01E-04 |
| P98158 | GFTCACPDDFQTVQLR | 4,Carbamidomethyl[C];6,Carbamidomethyl[C]; | 11.6 | 0.0 | 0.0 | 4.33E-04 |
| P07861 | LIQNMDASAEPCTDFFK | 12,Carbamidomethyl[C]; | 54.4 | 0.0 | 0.0 | 4.51E-04 |
| P02770 | CCSGSLVER | 1,Carbamidomethyl[C];2,Carbamidomethyl[C]; | 26.1 | 0.0 | 0.0 | 5.04E-04 |
| P20760 | SNSMVTLGCLVK | 9,Carbamidomethyl[C]; | 14.6 | 0.0 | 0.0 | 7.37E-04 |
| P20759 | SNSMVTLGCLVK | 9,Carbamidomethyl[C]; | 14.6 | 0.0 | 0.0 | 7.37E-04 |
| P07522 | GQCLYSLSER | 3,Carbamidomethyl[C]; | 45.0 | 0.0 | 0.0 | 7.78E-04 |
| P10960 | GCSFLPDPYQK | 2,Carbamidomethyl[C]; | 16.3 | 0.0 | 0.0 | 7.80E-04 |
| P20611 | FPLGPCPR | 6,Carbamidomethyl[C]; | 16.7 | 0.0 | 0.0 | 7.99E-04 |
| P18292 | EECSIPVCGQEGR | 3,Carbamidomethyl[C];8,Carbamidomethyl[C]; | 12.4 | 0.0 | 0.0 | 8.24E-04 |
| P12346 | EGVCPEGSIDSAPVK | 4,Carbamidomethyl[C]; | 27.0 | 0.0 | 0.0 | 8.99E-04 |
| Q9JJ19 | IVEVNGVCMEGK | 5,Deamidated[N];8,Carbamidomethyl[C];9,Oxidation[M]; | 5.1 | 0.0 | 0.0 | 9.65E-04 |
| P22057 | TEEDIVFLPQPDK | 1,Hex(1)NeuAc(1)[T]; | 2.6 | 0.0 | 0.0 | 9.65E-04 |
| P20767 | NSFTCQVTHEGNTVEK | 5,Carbamidomethyl[C];12,Deamidated[N]; | 7.7 | 0.0 | 0.0 | 9.65E-04 |
| P00689 | VGFMLAHPYGFTR | 0,Unknown_210[AnyN-term]; | 10.3 | 0.0 | 0.0 | 9.65E-04 |
| P14740 | KYPLLIDVYAGPCSQK | 13,Carbamidomethyl[C]; | 37.3 | 0.0 | 0.0 | 1.04E-03 |
| P07647 | AYDHNNDLMLLHLSKPADITGGVK | 6,Deamidated[N]; | 19.3 | 0.0 | 0.0 | 1.05E-03 |
| P07314 | QPLSSMCPSIIVDKDGK | 7,Carbamidomethyl[C]; | 20.6 | 0.0 | 0.0 | 1.06E-03 |
| Q6IE52 | VTASPQSLCGLR | 9,Carbamidomethyl[C]; | 23.1 | 0.0 | 0.0 | 1.12E-03 |
| Q03626 | VTASPQSLCGLR | 9,Carbamidomethyl[C]; | 23.1 | 0.0 | 0.0 | 1.12E-03 |
| P14046 | VTASPQSLCGLR | 9,Carbamidomethyl[C]; | 23.1 | 0.0 | 0.0 | 1.12E-03 |
| P02770 | YNEVLTQCCTESDK | 2,Deamidated[N];8,Carbamidomethyl[C];9,Carbamidomethyl[C]; | 12.0 | 0.0 | 0.0 | 1.16E-03 |
| P12346 | LLEACTFHK | 5,Carbamidomethyl[C]; | 15.9 | 0.0 | 0.0 | 1.20E-03 |
| P02770 | EDQLKTVMGDFAQFVDK | 4,trifluoro[L]; | 40.3 | 0.0 | 0.0 | 1.30E-03 |
| O70244 | QCMNDFLEVR | 2,Carbamidomethyl[C]; | 14.6 | 0.0 | 0.0 | 1.30E-03 |
| P02770 | DNCFATEGPNLVAR | 3,SulfoGMBS[C]; | 82.3 | 0.0 | 0.0 | 1.32E-03 |
| P02770 | CCTLPEAQR | 1,maleimide[C]; | 8.6 | 0.0 | 0.0 | 1.36E-03 |
| P20760 | SNSMVTLGCLVK | 2,Deamidated[N];9,Carbamidomethyl[C]; | 19.3 | 0.0 | 0.0 | 1.54E-03 |
| P20759 | SNSMVTLGCLVK | 2,Deamidated[N];9,Carbamidomethyl[C]; | 19.3 | 0.0 | 0.0 | 1.54E-03 |
| P14562 | YSGTCGAQLVTLK | 5,Carbamidomethyl[C]; | 5.6 | 0.0 | 0.0 | 1.57E-03 |
| Q9R0T4 | DWVIPPINCPENQK | 9,Carbamidomethyl[C]; | 63.0 | 0.0 | 0.0 | 1.59E-03 |
| P12346 | DLKQEDFQLLCPDGTK | 11,Carbamidomethyl[C]; | 27.9 | 0.0 | 0.0 | 1.64E-03 |
| P15083 | SSVTFECDLGR | 7,Carbamidomethyl[C]; | 205.3 | 0.0 | 0.0 | 1.67E-03 |
| P02761 | VFMQHIDVLENSLGFK | 3,Oxidation[M];10,Cation_Na[E];11,Deamidated[N]; | 18.9 | 0.0 | 0.0 | 1.70E-03 |
| P02761 | CEAHGITR | 1,Carbamidomethyl[C]; | 83.6 | 0.0 | 0.0 | 1.75E-03 |
| P48500 | LPADTEVVCAPPTAYIDFAR | 9,Carbamidomethyl[C]; | 7.7 | 0.0 | 0.0 | 1.75E-03 |
| Q01177 | KQLAAGSIADCLAK | 11,Carbamidomethyl[C]; | 6.4 | 0.0 | 0.0 | 1.82E-03 |
| Q5FVF9 | QPCLGSDPGCPQDGR | 3,Carbamidomethyl[C];10,Carbamidomethyl[C]; | 9.4 | 0.0 | 0.0 | 1.86E-03 |
| Q01460 | EDCPCSEASLCRPIR | 3,Carbamidomethyl[C];5,Carbamidomethyl[C];11,Carbamidomethyl[C]; | 9.4 | 0.0 | 0.0 | 1.86E-03 |
| O70513 | VDAECMPVVR | 5,Carbamidomethyl[C]; | 21.0 | 0.0 | 0.0 | 1.88E-03 |
| Q6IUU3 | DTWPPACPPLEPAK | 7,Carbamidomethyl[C]; | 49.3 | 0.0 | 0.0 | 1.93E-03 |
| P07647 | AYDHNNDLMLLHLSKPADITGGVK | 5,Deamidated[N]; | 22.3 | 0.0 | 0.0 | 2.14E-03 |
| P20767 | ANGAPISQGVDTANPTK | 2,Deamidated[N]; | 50.6 | 0.0 | 0.0 | 2.15E-03 |
| P12346 | DCTGNFCLFR | 2,Carbamidomethyl[C];7,Carbamidomethyl[C]; | 36.9 | 0.0 | 0.0 | 2.21E-03 |
| Q4FZV0 | FQSAVQYAECQSK | 10,Carbamidomethyl[C]; | 14.6 | 0.0 | 0.0 | 2.33E-03 |
| P36374 | VGSTCLTSGWGSITPLK | 5,Carbamidomethyl[C]; | 7.3 | 0.0 | 0.0 | 2.33E-03 |
| Q64240 | KGVCEEISGVYQK | 4,Carbamidomethyl[C]; | 50.6 | 0.0 | 0.0 | 2.37E-03 |
| P98158 | NPCASASCSHLCLLSAQAPR | 3,Carbamidomethyl[C];8,Carbamidomethyl[C];12,Carbamidomethyl[C]; | 15.9 | 0.0 | 0.0 | 2.52E-03 |
| Q9ESG3 | ELCRPDAENAFK | 3,Carbamidomethyl[C]; | 11.1 | 0.0 | 0.0 | 2.52E-03 |
| P02781 | LEEFDAPPEAVEANLK | 0,CLIP_TRAQ_2[AnyN-term]; | 41.1 | 0.0 | 0.0 | 2.56E-03 |
| P07522 | VNLDPASVPPR | 2,Asn->Cys[N]; | 14.6 | 0.0 | 0.0 | 2.64E-03 |
| P07647 | LVSQSFQHPDYIPVFMR | 1,Xle->Met[L]; | 16.7 | 0.0 | 0.0 | 2.89E-03 |
| P82450 | QCAVYHTSSVLPAPPFTAR | 2,Carbamidomethyl[C]; | 12.4 | 0.0 | 0.0 | 2.90E-03 |
| Q6DGG1 | EARPGNGQAVR | 6,Deamidated[N]; | 8.6 | 0.0 | 0.0 | 2.99E-03 |
| Q64230 | IAHVTLNEEK | 7,Deamidated[N]; | 4.3 | 0.0 | 0.0 | 2.99E-03 |
| P31977 | ILQLCMGNHELYMR | 5,Carbamidomethyl[C];6,Oxidation[M]; | 4.3 | 0.0 | 0.0 | 2.99E-03 |
| P98158 | DCSDGSDEPDLCPHR | 2,Carbamidomethyl[C];12,Carbamidomethyl[C]; | 11.1 | 0.0 | 0.0 | 3.08E-03 |
| P81828 | LALQCFRCTSFDSTGFCHVGR | 8,Carbamidomethyl[C];17,Carbamidomethyl[C]; | 21.4 | 0.0 | 0.0 | 3.15E-03 |
| P01835 | ADYESHNLYTCEVVHK | 11,Carbamidomethyl[C]; | 1129.3 | 0.0 | 0.0 | 3.22E-03 |
| P01835 | ADYESHNLYTCEVVHK | 4,Cation_Na[E];11,Carbamidomethyl[C]; | 43.3 | 0.0 | 0.0 | 3.37E-03 |
| P83121 | CQTYPDEICAWVVVTTR | 9,maleimide[C]; | 57.4 | 0.0 | 0.0 | 3.37E-03 |
| P81828 | CQTYPDEICAWVVVTTR | 9,maleimide[C]; | 57.4 | 0.0 | 0.0 | 3.37E-03 |
| P81827 | CQTYPDEICAWVVVTTR | 9,maleimide[C]; | 57.4 | 0.0 | 0.0 | 3.37E-03 |
| P83121 | PDEICAWVVVTTR | 5,Carbamidomethyl[C]; | 165.0 | 0.0 | 0.0 | 3.39E-03 |
| P81828 | PDEICAWVVVTTR | 5,Carbamidomethyl[C]; | 165.0 | 0.0 | 0.0 | 3.39E-03 |
| P81827 | PDEICAWVVVTTR | 5,Carbamidomethyl[C]; | 165.0 | 0.0 | 0.0 | 3.39E-03 |
| P12346 | KPVTEFATCHLAQAPNHVVVSR | 9,Carbamidomethyl[C]; | 59.6 | 0.0 | 0.0 | 3.40E-03 |
| Q9EPB1 | DLFLQGAYDTISQNFGTCQSLSSPK | 18,Carbamidomethyl[C]; | 33.0 | 0.0 | 0.0 | 3.54E-03 |
| P27590 | FSIQMFR | 5,Oxidation[M]; | 215.1 | 0.0 | 0.0 | 3.64E-03 |
| P81828 | DSTGFCHVGR | 6,Carbamidomethyl[C]; | 168.4 | 0.0 | 0.0 | 3.64E-03 |
| Q9R0T4 | DWVIPPINCPENQK | 8,Deamidated[N];9,Carbamidomethyl[C]; | 20.6 | 0.0 | 0.0 | 3.84E-03 |
| P19218 | MPETCVNVYR | 5,Carbamidomethyl[C]; | 8.6 | 0.0 | 0.0 | 4.06E-03 |
| Q64319 | KQCYFHQFLK | 3,Carbamidomethyl[C]; | 10.3 | 0.0 | 0.0 | 4.66E-03 |
| Q68FP1 | QHGMDDDGTGQK | 0,Gln->pyro-Glu[AnyN-termQ]; | 3.4 | 0.0 | 0.0 | 4.66E-03 |
| P05545 | QPILSEFQEK | 0,PyMIC[AnyN-term]; | 9.0 | 0.0 | 0.0 | 4.84E-03 |
| P05544 | QPILSEFQEK | 0,PyMIC[AnyN-term]; | 9.0 | 0.0 | 0.0 | 4.84E-03 |
| P02761 | EKIEENGSMR | 0,Glu->pyro-Glu[AnyN-termE]; | 9.0 | 0.0 | 0.0 | 4.84E-03 |
| P02761 | VFMQHIDVLENSLGFK | 3,Oxidation[M];10,Cation_Na[E]; | 97.3 | 0.0 | 0.0 | 4.90E-03 |
| P98158 | AIVLDPCR | 7,Carbamidomethyl[C]; | 14.6 | 0.0 | 0.0 | 4.93E-03 |
| P13635 | DIASGLIGPLILCK | 13,Carbamidomethyl[C]; | 12.0 | 0.0 | 0.0 | 4.98E-03 |
| P05545 | TSMVLVNYLLFK | 3,Oxidation[M]; | 12.9 | 0.0 | 0.0 | 5.01E-03 |
| P05544 | TSMVLVNYLLFK | 3,Oxidation[M]; | 12.9 | 0.0 | 0.0 | 5.01E-03 |
| P05545 | TQGKIAELFSELDER | 4,PyridoxalPhosphate[K]; | 38.6 | 0.0 | 0.0 | 5.20E-03 |
| P27590 | MAETCVPVLR | 5,Carbamidomethyl[C]; | 202.7 | 0.0 | 0.0 | 5.26E-03 |
| Q9JHY1 | KGSVYSPQTAVQVPENDSVK | 16,Deamidated[N]; | 4.7 | 0.0 | 0.0 | 5.32E-03 |
| P07861 | SESQMDITDINAPKPK | 5,Oxidation[M]; | 4.7 | 0.0 | 0.0 | 5.32E-03 |
| P05544 | TQGKIAELFSDLEER | 4,PyridoxalPhosphate[K]; | 7.3 | 0.0 | 0.0 | 5.38E-03 |
| Q6IRK9 | TYPDTDSFNTVAEITGSK | 0,Dimethyl_2H(6)13C(2)[AnyN-term]; | 13.3 | 0.0 | 0.0 | 5.41E-03 |
| Q5XI43 | EVLVVALGAPALMTCVNR | 15,Carbamidomethyl[C]; | 49.3 | 0.0 | 0.0 | 5.42E-03 |
| P02770 | SQCLAEIEHDNIPADLPSIAADFVEDKEVCK | 6,Cation_Na[E];8,Cation_Na[E];30,Carbamidomethyl[C]; | 9.9 | 0.0 | 0.0 | 5.54E-03 |
| O55006 | NVIILNNIK | 6,Deamidated[N]; | 10.3 | 0.0 | 0.0 | 5.56E-03 |
| Q01177 | YSPSTHPSEGLEENYCR | 16,Carbamidomethyl[C]; | 13.7 | 0.0 | 0.0 | 5.67E-03 |
| P20767 | NSFTCQVTHEGNTVEK | 1,Deamidated[N];5,Carbamidomethyl[C]; | 31.3 | 0.0 | 0.0 | 6.04E-03 |
| Q9JHB9 | SGSGCSILDEVIR | 5,CarbamidomethylDTT[C]; | 32.1 | 0.0 | 0.0 | 6.39E-03 |
| P02780 | SGSGCSILDEVIR | 5,CarbamidomethylDTT[C]; | 32.1 | 0.0 | 0.0 | 6.39E-03 |
| Q9QX79 | IFYETVHGQCK | 10,Carbamidomethyl[C]; | 9.4 | 0.0 | 0.0 | 6.46E-03 |
| P27590 | CQLQSLGFMK | 1,Carbamidomethyl[C]; | 57.4 | 0.0 | 0.0 | 6.47E-03 |
| P20059 | CNADPGLSALLSDHR | 1,Carbamidomethyl[C];2,Deamidated[N]; | 10.3 | 0.0 | 0.0 | 6.53E-03 |
| P07861 | QSNSLNYGGIGMVIGHEITHGFDDNGR | 0,Gln->pyro-Glu[AnyN-termQ];25,Deamidated[N]; | 6.9 | 0.0 | 0.0 | 6.78E-03 |
| P20059 | CNADPGLSALLSDHR | 0,Pyro-carbamidomethyl[AnyN-termC]; | 29.6 | 0.0 | 0.0 | 6.85E-03 |
| P02770 | CCAEGDPPACYGTVLAEFQPLVEEPK | 1,CarbamidomethylDTT[C]; | 8.1 | 0.0 | 0.0 | 7.19E-03 |
| O35568 | SGNENGEFYLR | 5,Deamidated[N]; | 9.4 | 0.0 | 0.0 | 7.71E-03 |
| P27590 | STDYGAGYSCDSDMHGWYR | 10,Carbamidomethyl[C];14,Oxidation[M]; | 24.9 | 0.0 | 0.0 | 8.16E-03 |
| P07861 | MDITDINAPKPK | 0,Acetyl[ProteinN-term];1,Oxidation[M]; | 4.3 | 0.0 | 0.0 | 8.24E-03 |
| P07522 | ADLGGMDVK | 6,Oxidation[M]; | 8.6 | 0.0 | 0.0 | 8.24E-03 |
| P02782 | QICELVAHETISFLMK | 3,DHP[C]; | 2.1 | 0.0 | 0.0 | 8.24E-03 |
| P01835 | ADYESHNLYTCEVVHK | 11,Cys->Dha[C]; | 6.4 | 0.0 | 0.0 | 8.24E-03 |
| P00758 | QPGDDYSNDLMLLHLSQPADITDGVK | 0,Formyl[AnyN-term]; | 2.1 | 0.0 | 0.0 | 8.24E-03 |
| O70244 | SCGGYLHADR | 2,Carbamidomethyl[C]; | 6.4 | 0.0 | 0.0 | 8.24E-03 |
| P28826 | MTDDNGSYLWDRPSK | 5,Deamidated[N]; | 23.1 | 0.0 | 0.0 | 8.33E-03 |
| P02761 | NGDWFSIVVASNKR | 0,Pyridylacetyl[AnyN-term](Phenylisocyanate[AnyN-term]); | 14.6 | 0.0 | 0.0 | 8.46E-03 |
| P07861 | QSNSLNYGGIGMVIGHEITHGFDDNGR | 25,Deamidated[N]; | 43.3 | 0.0 | 0.0 | 8.58E-03 |
| P15684 | QLQTDLSVIPVINR | 0,Biotin_Thermo-21328[AnyN-term]; | 15.0 | 0.0 | 0.0 | 8.77E-03 |
| P09005 | MQQVESSLQPETLKK | 1,Oxidation[M]; | 12.9 | 0.0 | 0.0 | 8.97E-03 |
| P05545 | MQQVESSLQPETLKK | 1,Oxidation[M]; | 12.9 | 0.0 | 0.0 | 8.97E-03 |
| P05544 | MQQVESSLQPETLKK | 1,Oxidation[M]; | 12.9 | 0.0 | 0.0 | 8.97E-03 |
| P02761 | EKIEENGSMR | 6,Ammonia-loss[N]; | 17.1 | 0.0 | 0.0 | 9.06E-03 |
| P04797 | LVINGKPITIFQER | 4,Deamidated[N]; | 21.0 | 0.0 | 0.0 | 9.18E-03 |
| P02761 | EDGEYFVEYDGGNTFTILK | 3,Gly->Met[G]; | 10.7 | 0.0 | 0.0 | 9.30E-03 |
| P07647 | AYDHNNDLMLLHLSK | 6,Deamidated[N]; | 27.4 | 0.0 | 0.0 | 9.44E-03 |
| P01836 | GVLDSVTDQDSK | 0,glycidamide[AnyN-term]; | 7.7 | 0.0 | 0.0 | 9.47E-03 |
| P01835 | GVLDSVTDQDSK | 0,glycidamide[AnyN-term]; | 7.7 | 0.0 | 0.0 | 9.47E-03 |
| P02761 | LCEAHGITR | 2,SulfurDioxide[C]; | 12.0 | 0.0 | 0.0 | 9.60E-03 |
| P02761 | EKIEENGSMR | 6,Deamidated[N];9,Oxidation[M]; | 89.6 | 0.0 | 0.0 | 9.80E-03 |
| P02770 | PPACYGTVLAEFQPLVEEPK | 4,Carbamidomethyl[C]; | 21.0 | 0.0 | 0.0 | 1.03E-02 |
| P36373 | EKVTDLMLCAGELEGGK | 9,Carbamidomethyl[C]; | 9.0 | 0.0 | 0.0 | 1.04E-02 |
| P15950 | EKVTDLMLCAGELEGGK | 9,Carbamidomethyl[C]; | 9.0 | 0.0 | 0.0 | 1.04E-02 |
| P12346 | ASDSSINWNNLK | 9,Deamidated[N]; | 16.3 | 0.0 | 0.0 | 1.04E-02 |
| Q64230 | PVENRQAIMTILDQEPDAR | 0,SPITC[AnyN-term]; | 4.7 | 0.0 | 0.0 | 1.05E-02 |
| Q63691 | GLISALCPHKFPTLQVLALR | 7,Carbamidomethyl[C]; | 9.4 | 0.0 | 0.0 | 1.05E-02 |
| P00689 | AVGFMLAHPYGFTR | 0,Iodoacetanilide_13C(6)[AnyN-term]; | 4.7 | 0.0 | 0.0 | 1.05E-02 |
| Q9JJ40 | INGVFVDKEEHAQVVDLVR | 2,Deamidated[N]; | 48.0 | 0.0 | 0.0 | 1.05E-02 |
| P98158 | CDNGHCIEMGR | 1,Carbamidomethyl[C];3,Deamidated[N];6,Carbamidomethyl[C]; | 5.6 | 0.0 | 0.0 | 1.07E-02 |
| P07522 | SIHHIRHQARHDLLTMAIFGDK | 0,Ethylphosphate[AnyN-term]; | 5.6 | 0.0 | 0.0 | 1.07E-02 |
| P07522 | HDLLTMAIFGDK | 2,Dehydrated[D]; | 5.6 | 0.0 | 0.0 | 1.07E-02 |
| P63259 | EYDESGPSIVHR | 0,Ethylphosphate[AnyN-term]; | 16.3 | 0.0 | 0.0 | 1.09E-02 |
| P60711 | EYDESGPSIVHR | 0,Ethylphosphate[AnyN-term]; | 16.3 | 0.0 | 0.0 | 1.09E-02 |
| P07522 | HDLLTMAIFGDK | 5,Dehydrated[T]; | 23.1 | 0.0 | 0.0 | 1.09E-02 |
| Q6P7A9 | AVPTQCDVTPNSR | 6,Carbamidomethyl[C];11,Deamidated[N]; | 5.1 | 0.0 | 0.0 | 1.11E-02 |
| Q64319 | DFENLVAAVHDK | 1,Dehydrated[D]; | 5.1 | 0.0 | 0.0 | 1.11E-02 |
| P53790 | IACVLPSECK | 3,Carbamidomethyl[C];9,Carbamidomethyl[C]; | 5.1 | 0.0 | 0.0 | 1.11E-02 |
| P42123 | VIGSGCNLDSAR | 6,Carbamidomethyl[C]; | 5.1 | 0.0 | 0.0 | 1.11E-02 |
| P36374 | EVTDVMLCAGEMDGGK | 8,Carbamidomethyl[C]; | 15.4 | 0.0 | 0.0 | 1.11E-02 |
| P19629 | VIGSGCNLDSAR | 6,Carbamidomethyl[C]; | 5.1 | 0.0 | 0.0 | 1.11E-02 |
| P04642 | VIGSGCNLDSAR | 6,Carbamidomethyl[C]; | 5.1 | 0.0 | 0.0 | 1.11E-02 |
| P19468 | GYVSDIDCR | 8,Carbamidomethyl[C]; | 6.4 | 0.0 | 0.0 | 1.13E-02 |
| P26644 | ITCPPPPIPK | 3,Carbamidomethyl[C]; | 9.9 | 0.0 | 0.0 | 1.17E-02 |
| Q64230 | PEFNAIIGQR | 0,NIC[AnyN-term](ICPL[AnyN-term]); | 3.9 | 0.0 | 0.0 | 1.17E-02 |
| P07861 | LPIDENQLSLEMNK | 6,Deamidated[N]; | 3.9 | 0.0 | 0.0 | 1.17E-02 |
| P02782 | QICELVAHETISFLMK | 3,HNE-Delta_H(2)O[C]; | 17.1 | 0.0 | 0.0 | 1.17E-02 |
| P01836 | TSSSPVVKSFNR | 0,Carbamyl[AnyN-term]; | 8.6 | 0.0 | 0.0 | 1.17E-02 |
| P01835 | TSSSPVVKSFNR | 0,Carbamyl[AnyN-term]; | 8.6 | 0.0 | 0.0 | 1.17E-02 |
| Q9JJ19 | LVEVNGENVEK | 5,Deamidated[N]; | 23.6 | 0.0 | 0.0 | 1.20E-02 |
| P05544 | SISTDYSLK | 0,CLIP_TRAQ_2[AnyN-term]; | 6.0 | 0.0 | 0.0 | 1.34E-02 |
| O70244 | LSTQPSCNSEHLIVFNGIR | 7,Carbamidomethyl[C];16,Deamidated[N]; | 9.0 | 0.0 | 0.0 | 1.39E-02 |
| P12346 | DSAFGLLR | 1,Dehydrated[D]; | 14.1 | 0.0 | 0.0 | 1.48E-02 |
| P01836 | SKDSTYSMSSTLSLTK | 1,Glucuronyl[S]; | 17.6 | 0.0 | 0.0 | 1.50E-02 |
| P01835 | SKDSTYSMSSTLSLTK | 1,Glucuronyl[S]; | 17.6 | 0.0 | 0.0 | 1.50E-02 |
| P02761 | GNLDVAKLNGDWFSIVVASNKR | 7,3-phosphoglyceryl[K]; | 61.3 | 0.0 | 0.0 | 1.51E-02 |
| P13635 | DLYSGLIGPLIVCR | 13,Carbamidomethyl[C]; | 10.3 | 0.0 | 0.0 | 1.52E-02 |
| P02770 | LGEYGFQNAVLVR | 3,Cation_Na[E]; | 3.4 | 0.0 | 0.0 | 1.52E-02 |
| P36373 | VTDLMLCAGELEGGK | 7,Carbamidomethyl[C]; | 3.4 | 0.0 | 0.0 | 1.52E-02 |
| P15950 | VTDLMLCAGELEGGK | 7,Carbamidomethyl[C]; | 3.4 | 0.0 | 0.0 | 1.52E-02 |
| P00758 | LYEDEPFAQHR | 0,NP40[AnyN-term]; | 3.4 | 0.0 | 0.0 | 1.52E-02 |
| O70244 | NGDSPGSPVIGR | 1,Deamidated[N]; | 8.1 | 0.0 | 0.0 | 1.53E-02 |
| P07522 | LIFWSSVVTGNLHR | 3,Phe->CamCys[F]; | 9.4 | 0.0 | 0.0 | 1.53E-02 |
| P27590 | VCQDPCNVYETLTEYWR | 6,4-ONE[C]; | 15.4 | 0.0 | 0.0 | 1.54E-02 |
| P01836 | DGVLDSVTDQDSK | 5,Cation_Na[D]; | 12.0 | 0.0 | 0.0 | 1.55E-02 |
| P01835 | DGVLDSVTDQDSK | 5,Cation_Na[D]; | 12.0 | 0.0 | 0.0 | 1.55E-02 |
| P22282 | LDNCPFEEQTEQLKR | 4,Carbamidomethyl[C]; | 231.0 | 0.0 | 0.0 | 1.55E-02 |
| P28826 | LSQVLSGPENDHSNMGQCK | 18,Carbamidomethyl[C]; | 14.6 | 0.0 | 0.0 | 1.56E-02 |
| P98158 | CQTTNICVPR | 1,Carbamidomethyl[C];7,Carbamidomethyl[C]; | 12.4 | 0.0 | 0.0 | 1.57E-02 |
| Q63041 | ATPLSLCALTAVDQSVLLLKPEAK | 7,Carbamidomethyl[C]; | 4.3 | 0.0 | 0.0 | 1.57E-02 |
| Q01177 | PWCFTTDPNKR | 3,Carbamidomethyl[C]; | 4.3 | 0.0 | 0.0 | 1.57E-02 |
| P07647 | AYDHNNDLMLLHLSK | 5,Deamidated[N]; | 17.6 | 0.0 | 0.0 | 1.60E-02 |
| P06760 | VPSSFNDITQEAELR | 0,Methylthio[AnyN-term]; | 21.0 | 0.0 | 0.0 | 1.62E-02 |
| Q64230 | GAILNAFEMFR | 5,Deamidated[N]; | 16.7 | 0.0 | 0.0 | 1.64E-02 |
| Q4QQW8 | TNTKPSVGSGSCSALIK | 12,Carbamidomethyl[C]; | 2.6 | 0.0 | 0.0 | 1.67E-02 |
| P36376 | VIDLPTEEPK | 1,Val->Pro[V]; | 5.1 | 0.0 | 0.0 | 1.67E-02 |
| P36375 | VIDLPTEEPK | 1,Val->Pro[V]; | 5.1 | 0.0 | 0.0 | 1.67E-02 |
| P36374 | VIDLPTEEPK | 1,Val->Pro[V]; | 5.1 | 0.0 | 0.0 | 1.67E-02 |
| P36373 | VIDLPTEEPK | 1,Val->Pro[V]; | 5.1 | 0.0 | 0.0 | 1.67E-02 |
| P21704 | LIPDSADTTATSTHCAYDR | 15,Carbamidomethyl[C]; | 5.1 | 0.0 | 0.0 | 1.67E-02 |
| P15083 | QGANGYCATLISSNGYLSK | 0,Gln->pyro-Glu[AnyN-termQ];7,Carbamidomethyl[C]; | 7.7 | 0.0 | 0.0 | 1.67E-02 |
| P07647 | VIDLPTEEPK | 1,Val->Pro[V]; | 5.1 | 0.0 | 0.0 | 1.67E-02 |
| P02781 | TLAGQVCQALQDVTITFLLNPEEELKR | 7,Bromobimane[C]; | 2.6 | 0.0 | 0.0 | 1.67E-02 |
| O35568 | QTSPVSAMLVLVK | 0,Gln->pyro-Glu[AnyN-termQ]; | 2.6 | 0.0 | 0.0 | 1.67E-02 |
| P27590 | CDSDMHGWYR | 1,ICAT-C[C]; | 13.7 | 0.0 | 0.0 | 1.69E-02 |
| P22282 | LDNCPFEEQTEQLKR | 3,Deamidated[N];4,Carbamidomethyl[C]; | 9.4 | 0.0 | 0.0 | 1.71E-02 |
| P22057 | DQGLTEEDIVFLPQPDK | 0,Amidine[AnyN-term]; | 4.7 | 0.0 | 0.0 | 1.71E-02 |
| P22282 | LDNCPFEEQTEQLK | 3,Deamidated[N];4,Carbamidomethyl[C]; | 11.6 | 0.0 | 0.0 | 1.73E-02 |
| Q5XI43 | IDDLEPADEGIYSCHLHHHYCGLHER | 14,Carbamidomethyl[C];21,Carbamidomethyl[C]; | 56.6 | 0.0 | 0.0 | 1.73E-02 |
| Q9JHB9 | SGCSILDEVIR | 3,Carbamidomethyl[C]; | 36.0 | 0.0 | 0.0 | 1.74E-02 |
| P02780 | SGCSILDEVIR | 3,Carbamidomethyl[C]; | 36.0 | 0.0 | 0.0 | 1.74E-02 |
| P81828 | QKCQTYPDEICAWVVVTTR | 3,Cys->Ala[C]; | 50.1 | 0.0 | 0.0 | 1.76E-02 |
| Q9QZA2 | CSDIVFAR | 1,Carbamidomethyl[C]; | 3.0 | 0.0 | 0.0 | 1.77E-02 |
| P98158 | NDCGDYSDER | 3,Carbamidomethyl[C]; | 3.0 | 0.0 | 0.0 | 1.77E-02 |
| P98158 | QIISNLNNPR | 0,Gln->pyro-Glu[AnyN-termQ]; | 3.0 | 0.0 | 0.0 | 1.77E-02 |
| P98158 | CDGVYDCR | 1,Carbamidomethyl[C];7,Carbamidomethyl[C]; | 3.0 | 0.0 | 0.0 | 1.77E-02 |
| P28826 | LSQVLSGPENDHSNMGQCK | 10,Deamidated[N];18,Carbamidomethyl[C]; | 12.0 | 0.0 | 0.0 | 1.77E-02 |
| P22283 | VNCPFEEQTEQLKR | 2,Hep[N]; | 6.0 | 0.0 | 0.0 | 1.77E-02 |
| P15083 | TIELQVAEATK | 0,Ethylphosphate[AnyN-term]; | 3.0 | 0.0 | 0.0 | 1.77E-02 |
| P08592 | EVCSEQAETGPCR | 3,Carbamidomethyl[C];12,Carbamidomethyl[C]; | 6.0 | 0.0 | 0.0 | 1.77E-02 |
| P02781 | LEEFDAPPEAVEANLK | 0,CarboxymethylDMAP[AnyN-term]; | 3.0 | 0.0 | 0.0 | 1.77E-02 |
| P02770 | AADKDNCFATEGPNLVAR | 7,Ethanolyl[C]; | 3.0 | 0.0 | 0.0 | 1.77E-02 |
| P02761 | NGDWFSIVVASNK | 0,Iodoacetanilide_13C(6)[AnyN-term]; | 3.0 | 0.0 | 0.0 | 1.77E-02 |
| P01836 | HNLYTCEVVHK | 2,Deamidated[N];6,Carbamidomethyl[C]; | 3.0 | 0.0 | 0.0 | 1.77E-02 |
| P01835 | HNLYTCEVVHK | 2,Deamidated[N];6,Carbamidomethyl[C]; | 3.0 | 0.0 | 0.0 | 1.77E-02 |
| G3V928 | NDCGDYSDER | 3,Carbamidomethyl[C]; | 3.0 | 0.0 | 0.0 | 1.77E-02 |
| P02761 | NGDWFSIVVASNK | 0,mTRAQ[AnyN-term]; | 26.1 | 0.0 | 0.0 | 1.81E-02 |
| P29598 | RPWCYVQIGLK | 4,Carbamidomethyl[C]; | 42.0 | 0.0 | 0.0 | 1.82E-02 |
| P02761 | VFMQHIDVLENSLGFK | 0,Delta_H(2)C(2)[AnyN-term]; | 6.4 | 0.0 | 0.0 | 1.86E-02 |
| P01836 | RDGVLDSVTDQDSK | 2,Dehydrated[D]; | 6.4 | 0.0 | 0.0 | 1.86E-02 |
| P01835 | RDGVLDSVTDQDSK | 2,Dehydrated[D]; | 6.4 | 0.0 | 0.0 | 1.86E-02 |
| P27590 | CDSDMHGWYR | 0,C8-QAT[AnyN-term]; | 62.1 | 0.0 | 0.0 | 1.89E-02 |
| P02770 | VSTPTLVEAAR | 0,Unknown_302[AnyN-term]; | 20.1 | 0.0 | 0.0 | 1.91E-02 |
| P61589 | IGAFGYMECSAK | 9,Carbamidomethyl[C]; | 11.1 | 0.0 | 0.0 | 1.92E-02 |
| Q64230 | THTLLDHCAFEK | 8,Carbamidomethyl[C]; | 21.4 | 0.0 | 0.0 | 1.92E-02 |
| P07522 | QARHDLLTMAIFGDK | 0,LG-pyrrole[AnyN-term]; | 18.9 | 0.0 | 0.0 | 1.94E-02 |
| P48508 | LFIVGSNSSSSTR | 7,Deamidated[N]; | 7.7 | 0.0 | 0.0 | 1.94E-02 |
| P07314 | QPLSSMCPSIIVDK | 7,Carbamidomethyl[C]; | 15.4 | 0.0 | 0.0 | 1.94E-02 |
| P80067 | AISYCHETMTGWVHDVLGR | 5,Carbamidomethyl[C]; | 11.6 | 0.0 | 0.0 | 1.97E-02 |
| P81828 | LALQCFRCTSFDSTGFCHVGR | 5,Carbamidomethyl[C];17,Carbamidomethyl[C]; | 14.1 | 0.0 | 0.0 | 2.04E-02 |
| Q9JJ19 | DPDSPAEASGLR | 0,CarboxymethylDMAP[AnyN-term]; | 5.6 | 0.0 | 0.0 | 2.05E-02 |
| P50123 | MNFAEEEPSKK | 0,Acetyl[ProteinN-term]; | 5.6 | 0.0 | 0.0 | 2.05E-02 |
| P20767 | TPTLTVFPPSTEELQGNK | 1,Dimethylphosphothione[T]; | 5.6 | 0.0 | 0.0 | 2.05E-02 |
| Q63041 | TEVNTNHVLIYIEK | 4,Deamidated[N]; | 7.3 | 0.0 | 0.0 | 2.10E-02 |
| P01048 | QEEGAQELNCNDETVFQAVDTALK | 0,Gln->pyro-Glu[AnyN-termQ];10,Carbamidomethyl[C]; | 16.7 | 0.0 | 0.0 | 2.11E-02 |
| Q63041 | TEVNTNHVLIYIEK | 6,Deamidated[N]; | 12.0 | 0.0 | 0.0 | 2.12E-02 |
| P02770 | RPCFSALTVDETYVPK | 3,Carbamidomethyl[C];11,Cation_Na[E]; | 12.0 | 0.0 | 0.0 | 2.12E-02 |
| P20767 | TPTLTVFPPSTEELQGNK | 0,GIST-Quat[AnyN-term]; | 25.3 | 0.0 | 0.0 | 2.13E-02 |
| P29598 | TDSCSGDSGGPLICNIDGR | 4,Carbamidomethyl[C];14,Carbamidomethyl[C]; | 19.3 | 0.0 | 0.0 | 2.14E-02 |
| Q08420 | GGNQASVQNGNAGR | 9,Deamidated[N]; | 15.9 | 0.0 | 0.0 | 2.23E-02 |
| P01836 | RDGVLDSVTDQDSK | 0,C+12[AnyN-term]; | 3.9 | 0.0 | 0.0 | 2.23E-02 |
| P01835 | RDGVLDSVTDQDSK | 0,C+12[AnyN-term]; | 3.9 | 0.0 | 0.0 | 2.23E-02 |
| Q01177 | CEGETDFICR | 1,Carbamidomethyl[C];9,Carbamidomethyl[C]; | 6.0 | 0.0 | 0.0 | 2.24E-02 |
| P29598 | GEHCEIDTSK | 4,Carbamidomethyl[C]; | 6.0 | 0.0 | 0.0 | 2.24E-02 |
| P19218 | STENTVSSEECDSHLR | 11,Carbamidomethyl[C]; | 6.0 | 0.0 | 0.0 | 2.24E-02 |
| Q62867 | VVPIRLDLNDAQYETLFR | 0,Unknown_250[AnyN-term]; | 8.6 | 0.0 | 0.0 | 2.25E-02 |
| P50123 | GCQIYLENFK | 2,Carbamidomethyl[C]; | 10.7 | 0.0 | 0.0 | 2.26E-02 |
| P20059 | GATYAFSGSHYWR | 3,Dehydrated[T]; | 10.7 | 0.0 | 0.0 | 2.26E-02 |
| P06760 | TSHYPYSEEVLQLCDR | 14,Carbamidomethyl[C]; | 74.1 | 0.0 | 0.0 | 2.27E-02 |
| P00758 | NLYEDEPFAQHR | 3,monomethylphosphothione[Y]; | 6.4 | 0.0 | 0.0 | 2.28E-02 |
| P27590 | GDGWYCECSPGFCEPGLDCLPQGPSGK | 8,Carbamidomethyl[C];13,Carbamidomethyl[C];19,Carbamidomethyl[C]; | 5.1 | 0.0 | 0.0 | 2.30E-02 |
| P22057 | DQGLTEEDIVFLPQPDK | 1,Cation_Ca[II][D]; | 5.1 | 0.0 | 0.0 | 2.30E-02 |
| Q68FP1 | AQHVQVEEGSEPDGFWEALGGK | 12,Pro->Ser[P]; | 12.9 | 0.0 | 0.0 | 2.39E-02 |
| Q9EQV6 | LLNGRPPLGFLNPR | 3,Deamidated[N]; | 10.7 | 0.0 | 0.0 | 2.42E-02 |
| P83121 | HICQTYPDEICAWVVVTTR | 11,Cys->Ala[C]; | 20.6 | 0.0 | 0.0 | 2.43E-02 |
| P02761 | IKENGECRELYLVAYK | 7,Carbamidomethyl[C]; | 47.6 | 0.0 | 0.0 | 2.46E-02 |
| P54311 | ELAGHTGYLSCCR | 11,Carbamidomethyl[C];12,Carbamidomethyl[C]; | 4.7 | 0.0 | 0.0 | 2.48E-02 |
| Q9JJ40 | EGNSFGFSLK | 3,Deamidated[N]; | 12.9 | 0.0 | 0.0 | 2.49E-02 |
| P22282 | RTPGETMYYISLPGSVR | 7,Met->His[M]; | 4.3 | 0.0 | 0.0 | 2.49E-02 |
| P19468 | CNQIANELCECPELLGSGFR | 1,Carbamidomethyl[C];9,Carbamidomethyl[C];11,Carbamidomethyl[C]; | 4.3 | 0.0 | 0.0 | 2.49E-02 |
| P07154 | MIQLHNGEYSNGK | 6,Deamidated[N];11,Deamidated[N]; | 8.6 | 0.0 | 0.0 | 2.49E-02 |
| P01836 | RDGVLDSVTDQDSK | 6,Dehydrated[D]; | 4.3 | 0.0 | 0.0 | 2.49E-02 |
| P01835 | RDGVLDSVTDQDSK | 6,Dehydrated[D]; | 4.3 | 0.0 | 0.0 | 2.49E-02 |
| P01836 | RDGVLDSVTDQDSK | 12,Dehydrated[D]; | 9.9 | 0.0 | 0.0 | 2.50E-02 |
| P01835 | RDGVLDSVTDQDSK | 12,Dehydrated[D]; | 9.9 | 0.0 | 0.0 | 2.50E-02 |
| P27590 | CDSDMHGWYR | 0,Brij58[AnyN-term]; | 18.4 | 0.0 | 0.0 | 2.58E-02 |
| P81827 | RICQTYPDEICAWVVVTTR | 3,Carbamidomethyl[C];11,Carbamidomethyl[C]; | 6.9 | 0.0 | 0.0 | 2.60E-02 |
| P02761 | NGSMRVFMQHIDVLENSLGFK | 0,Acetyl[ProteinN-term]; | 5.6 | 0.0 | 0.0 | 2.62E-02 |
| P22283 | CSFQWEFCNIK | 1,Carbamidomethyl[C];8,Carbamidomethyl[C]; | 29.1 | 0.0 | 0.0 | 2.63E-02 |
| P05544 | MPKFSISTDYSLK | 3,Xlink_BS2G[217][K]; | 28.3 | 0.0 | 0.0 | 2.64E-02 |
| P02782 | LEMYNAPPAAVEAK | 0,Iodoacetanilide_13C(6)[AnyN-term]; | 10.3 | 0.0 | 0.0 | 2.65E-02 |
| P83121 | DEICAWVVVTTR | 2,Cation_Na[E];4,Carbamidomethyl[C]; | 6.4 | 0.0 | 0.0 | 2.72E-02 |
| P81828 | DEICAWVVVTTR | 2,Cation_Na[E];4,Carbamidomethyl[C]; | 6.4 | 0.0 | 0.0 | 2.72E-02 |
| P81827 | DEICAWVVVTTR | 2,Cation_Na[E];4,Carbamidomethyl[C]; | 6.4 | 0.0 | 0.0 | 2.72E-02 |
| P20767 | STPTLTVFPPSTEELQGNK | 2,Dehydrated[T]; | 27.4 | 0.0 | 0.0 | 2.78E-02 |
| P20767 | NSFTCQVTHEGNTVEK | 5,Carbamidomethyl[C]; | 80.6 | 0.0 | 0.0 | 2.78E-02 |
| Q01177 | CQEWAAQEPHSHR | 1,Dioxidation[C]; | 7.3 | 0.0 | 0.0 | 2.78E-02 |
| P02770 | ADKDNCFATEGPNLVAR | 6,DNCB_hapten[C]; | 10.3 | 0.0 | 0.0 | 2.82E-02 |
| P02761 | LNGDWFSIVVASNKR | 0,Delta_H(2)C(2)[AnyN-term]; | 10.3 | 0.0 | 0.0 | 2.82E-02 |
| P20059 | GECQSEGVLFFQGNR | 3,Carbamyl[C]; | 16.3 | 0.0 | 0.0 | 2.82E-02 |
| Q64230 | IPYILADNLDLNAK | 0,SPITC[AnyN-term]; | 7.7 | 0.0 | 0.0 | 2.84E-02 |
| P02780 | VKPYVQDHFTEK | 0,Ethylphosphate[AnyN-term]; | 7.7 | 0.0 | 0.0 | 2.84E-02 |
| P05545 | TEGAAATAVTAALK | 0,NDA[AnyN-term]; | 81.4 | 0.0 | 0.0 | 2.85E-02 |
| P83121 | HICQTYPDEICAWVVVTTR | 3,Carbamidomethyl[C];11,Carbamidomethyl[C]; | 202.3 | 0.0 | 0.0 | 2.93E-02 |
| P83121 | HICQTYPDEICAWVVVTTR | 3,Cys->Ala[C]; | 16.7 | 0.0 | 0.0 | 2.96E-02 |
| Q9Z1P2 | EGLLLWCQR | 7,Carbamidomethyl[C]; | 3.4 | 0.0 | 0.0 | 3.00E-02 |
| Q9QXQ0 | EGLLLWCQR | 7,Carbamidomethyl[C]; | 3.4 | 0.0 | 0.0 | 3.00E-02 |
| Q63772 | LCDKDVNECSQK | 2,Carbamidomethyl[C];9,Carbamidomethyl[C]; | 1.7 | 0.0 | 0.0 | 3.00E-02 |
| Q63621 | GCTEIVNFHNVQPK | 2,Carbamidomethyl[C]; | 1.7 | 0.0 | 0.0 | 3.00E-02 |
| P97840 | NTQINNSWGPEER | 5,Deamidated[N]; | 12.0 | 0.0 | 0.0 | 3.00E-02 |
| P81827 | ALQCFRCESLDSTGLCR | 0,Acetyl[ProteinN-term];16,Carbamidomethyl[C]; | 5.1 | 0.0 | 0.0 | 3.00E-02 |
| P63018 | VEIIANDQGNR | 6,Deamidated[N]; | 3.4 | 0.0 | 0.0 | 3.00E-02 |
| P55063 | VEIIANDQGNR | 6,Deamidated[N]; | 3.4 | 0.0 | 0.0 | 3.00E-02 |
| P50399 | GGKIYKVPSTEAEALASSLMGLFEK | 10,Thr->Pro[T]; | 1.7 | 0.0 | 0.0 | 3.00E-02 |
| P47967 | NTQINNSWGPEER | 5,Deamidated[N]; | 12.0 | 0.0 | 0.0 | 3.00E-02 |
| P36374 | NNLLEDEPFAQHR | 0,Formyl[AnyN-term]; | 1.7 | 0.0 | 0.0 | 3.00E-02 |
| P36373 | NNLLEDEPFAQHR | 0,Formyl[AnyN-term]; | 1.7 | 0.0 | 0.0 | 3.00E-02 |
| P29598 | TCYHGNGQSYR | 2,Carbamidomethyl[C];6,Deamidated[N]; | 1.7 | 0.0 | 0.0 | 3.00E-02 |
| P28826 | GSGCWSSVGNIHAGK | 4,Carbamidomethyl[C]; | 3.4 | 0.0 | 0.0 | 3.00E-02 |
| P25093 | VGFGQCAGK | 6,Carbamidomethyl[C]; | 1.7 | 0.0 | 0.0 | 3.00E-02 |
| P22283 | CPFEEQTEQLKR | 1,Carbamidomethyl[C]; | 3.4 | 0.0 | 0.0 | 3.00E-02 |
| P22282 | CPFEEQTEQLKR | 1,Carbamidomethyl[C]; | 3.4 | 0.0 | 0.0 | 3.00E-02 |
| P22282 | LKDVYDVFK | 0,Formyl[AnyN-term]; | 3.4 | 0.0 | 0.0 | 3.00E-02 |
| P20767 | SRNSFTCQVTHEGNTVEK | 3,Deamidated[N];7,Carbamidomethyl[C]; | 1.7 | 0.0 | 0.0 | 3.00E-02 |
| P19468 | LGCPGFTLPEHRPNPEEGGASK | 3,Carbamidomethyl[C]; | 6.9 | 0.0 | 0.0 | 3.00E-02 |
| P14659 | VEIIANDQGNR | 6,Deamidated[N]; | 3.4 | 0.0 | 0.0 | 3.00E-02 |
| P10252 | MLHETEDQWK | 1,Oxidation[M]; | 1.7 | 0.0 | 0.0 | 3.00E-02 |
| P0DMW1 | VEIIANDQGNR | 6,Deamidated[N]; | 3.4 | 0.0 | 0.0 | 3.00E-02 |
| P0DMW0 | VEIIANDQGNR | 6,Deamidated[N]; | 3.4 | 0.0 | 0.0 | 3.00E-02 |
| P06761 | VEIIANDQGNR | 6,Deamidated[N]; | 3.4 | 0.0 | 0.0 | 3.00E-02 |
| P02781 | LEEFDAPPEAVEANLK | 0,Iodoacetanilide_13C(6)[AnyN-term]; | 1.7 | 0.0 | 0.0 | 3.00E-02 |
| P02780 | PYVQDHFTEK | 8,Pent(1)HexNAc(1)[T]; | 1.7 | 0.0 | 0.0 | 3.00E-02 |
| P01836 | VLDSVTDQDSKDSTYSMSSTLSLTK | 4,pyrophospho[S]; | 1.7 | 0.0 | 0.0 | 3.00E-02 |
| P01836 | RDGVLDSVTDQDSK | 9,Thr->Asn[T]; | 1.7 | 0.0 | 0.0 | 3.00E-02 |
| P01835 | VLDSVTDQDSKDSTYSMSSTLSLTK | 4,pyrophospho[S]; | 1.7 | 0.0 | 0.0 | 3.00E-02 |
| P01835 | RDGVLDSVTDQDSK | 9,Thr->Asn[T]; | 1.7 | 0.0 | 0.0 | 3.00E-02 |
| O70244 | NGGSSTSPLLDK | 1,Deamidated[N]; | 5.1 | 0.0 | 0.0 | 3.00E-02 |
| G3V686 | TLSCTSVK | 4,Carbamidomethyl[C]; | 1.7 | 0.0 | 0.0 | 3.00E-02 |
| G3V686 | DGTTCHCQCAVMDWATAR | 0,AccQTag[AnyN-term]; | 1.7 | 0.0 | 0.0 | 3.00E-02 |
| P20059 | GECQSEGVLFFQGNR | 3,Carbamidomethyl[C]; | 22.7 | 0.0 | 0.0 | 3.09E-02 |
| Q62687 | AQASGMDPLVDIEDERPK | 0,Acetyl[ProteinN-term];6,Oxidation[M]; | 7.3 | 0.0 | 0.0 | 3.14E-02 |
| P02781 | ELEEFDAPPEAVEANLK | 6,Asp->Lys[D]; | 36.0 | 0.0 | 0.0 | 3.15E-02 |
| P05545 | IAELFSELDER | 0,C+12[AnyN-term]; | 6.4 | 0.0 | 0.0 | 3.18E-02 |
| Q9JJ40 | GSNGYGFYLR | 3,Deamidated[N]; | 16.3 | 0.0 | 0.0 | 3.18E-02 |
| P42854 | WRENNCISELPYVCK | 4,Deamidated[N];6,Carbamidomethyl[C];14,Carbamidomethyl[C]; | 10.3 | 0.0 | 0.0 | 3.18E-02 |
| P10252 | EDQGVDYTWYEDSGPFPQR | 1,Glu->Xle[E]; | 11.1 | 0.0 | 0.0 | 3.24E-02 |
| P07647 | AYDHNNDLMLLHLSKPADITGGVK | 6,Ammonia-loss[N]; | 5.6 | 0.0 | 0.0 | 3.24E-02 |
| P07522 | ISVLILDILDKR | 1,Xle->Pro[I]; | 5.6 | 0.0 | 0.0 | 3.24E-02 |
| P02770 | CCAEGDPPACYGTVLAEFQPLVEEPK | 2,CarbamidomethylDTT[C]; | 5.6 | 0.0 | 0.0 | 3.24E-02 |
| Q5XI43 | GDEGVYTCNLHHHYCHLYESLAVR | 8,Carbamidomethyl[C];15,Carbamidomethyl[C]; | 33.0 | 0.0 | 0.0 | 3.26E-02 |
| Q9Z0W7 | DEFTNTCPSDK | 7,Carbamidomethyl[C]; | 4.7 | 0.0 | 0.0 | 3.33E-02 |
| Q6P7S1 | PGLIGSIPGPFGEEMR | 0,Diethylphosphate[AnyN-term]; | 4.7 | 0.0 | 0.0 | 3.33E-02 |
| Q63257 | TSTCEWQLDSTVDCSSQLLLDYR | 4,Carbamidomethyl[C];14,Carbamidomethyl[C]; | 4.7 | 0.0 | 0.0 | 3.33E-02 |
| P63018 | TVTNAVVTVPAYFNDSQR | 4,Deamidated[N]; | 4.7 | 0.0 | 0.0 | 3.33E-02 |
| P54313 | ELPGHTGYLSCCR | 11,Carbamidomethyl[C];12,Carbamidomethyl[C]; | 4.7 | 0.0 | 0.0 | 3.33E-02 |
| P31044 | GNDISSGTVLSEYVGSGPPK | 3,Cation_Na[D]; | 4.7 | 0.0 | 0.0 | 3.33E-02 |
| P21704 | LTNQMAEAISDHYPVEVTLR | 5,Met->Gln[M]; | 4.7 | 0.0 | 0.0 | 3.33E-02 |
| P09005 | MQQVESSLQPETLKK | 0,ISD_z+2_ion[AnyN-term]; | 4.7 | 0.0 | 0.0 | 3.33E-02 |
| P05545 | MQQVESSLQPETLKK | 0,ISD_z+2_ion[AnyN-term]; | 4.7 | 0.0 | 0.0 | 3.33E-02 |
| P05544 | MQQVESSLQPETLKK | 0,ISD_z+2_ion[AnyN-term]; | 4.7 | 0.0 | 0.0 | 3.33E-02 |
| O35353 | ELPGHTGYLSCCR | 11,Carbamidomethyl[C];12,Carbamidomethyl[C]; | 4.7 | 0.0 | 0.0 | 3.33E-02 |
| Q63041 | MMGASEVAQEVEVR | 0,C+12[AnyN-term]; | 10.3 | 0.0 | 0.0 | 3.37E-02 |
| Q64230 | SIDWGWGQAISHQMLMR | 0,Methyl[AnyN-term]; | 13.3 | 0.0 | 0.0 | 3.37E-02 |
| P98158 | ECSESEFR | 2,Carbamidomethyl[C]; | 6.9 | 0.0 | 0.0 | 3.42E-02 |
| P06760 | FLINGKPFYFQGVNKHEDSDIR | 4,Deamidated[N]; | 6.9 | 0.0 | 0.0 | 3.42E-02 |
| Q9JJ40 | EAPAPISAPLEAPGSATTEDVGDHKPK | 0,Glu->pyro-Glu[AnyN-termE]; | 3.9 | 0.0 | 0.0 | 3.49E-02 |
| P05545 | QPILSEFQEK | 0,Iodoacetanilide[AnyN-term]; | 3.9 | 0.0 | 0.0 | 3.49E-02 |
| P05544 | QPILSEFQEK | 0,Iodoacetanilide[AnyN-term]; | 3.9 | 0.0 | 0.0 | 3.49E-02 |
| P01835 | ADYESHNLYTCEVVHK | 11,Propionamide_2H(3)[C]; | 3.9 | 0.0 | 0.0 | 3.49E-02 |
| P21704 | LTNQMAEAISDHYPVEVTLR | 7,Glu->Xle[E]; | 3.9 | 0.0 | 0.0 | 3.49E-02 |
| P02780 | VKPYVQDHFTEK | 2,Xlink_DSSO[104][K]; | 3.9 | 0.0 | 0.0 | 3.49E-02 |
| Q64319 | TNNGFVQNEDIQEQDPDSR | 2,Deamidated[N]; | 4.3 | 0.0 | 0.0 | 3.53E-02 |
| Q64230 | QAIMTILDQEPDAR | 10,Cation_Na[E]; | 4.3 | 0.0 | 0.0 | 3.53E-02 |
| P21704 | AEAISDHYPVEVTLR | 2,Cation_Li[E]; | 4.3 | 0.0 | 0.0 | 3.53E-02 |
| P07522 | HDLLTMAIFGDK | 6,Oxidation[M]; | 4.3 | 0.0 | 0.0 | 3.53E-02 |
| P05544 | IAELFSDLEER | 1,Xle->Cys[I]; | 4.3 | 0.0 | 0.0 | 3.53E-02 |
| P02770 | TNCELYEKLGEYGFQNAVLVR | 2,HexNAc(1)dHex(2)[N]; | 6.4 | 0.0 | 0.0 | 3.66E-02 |
| P01836 | HNLYTCEVVHK | 6,Carbamidomethyl[C]; | 209.1 | 0.0 | 0.0 | 3.68E-02 |
| P01835 | HNLYTCEVVHK | 6,Carbamidomethyl[C]; | 209.1 | 0.0 | 0.0 | 3.68E-02 |
| P00758 | EEVTDLMLCAGEMDGGKDTCK | 9,Carbamidomethyl[C];20,Carbamidomethyl[C]; | 11.1 | 0.0 | 0.0 | 3.71E-02 |
| P02625 | SMTDLLSAEDIKK | 0,Acetyl[ProteinN-term]; | 5.1 | 0.0 | 0.0 | 3.75E-02 |
| P30120 | GYQSDHFACLPR | 9,Carbamidomethyl[C]; | 5.1 | 0.0 | 0.0 | 3.75E-02 |
| P18292 | RRANSGFLEELRK | 5,Hex(1)NeuAc(1)[S]; | 5.1 | 0.0 | 0.0 | 3.75E-02 |
| P02761 | LNGDWFSIVVASNK | 0,Delta_H(2)C(2)[AnyN-term]; | 5.1 | 0.0 | 0.0 | 3.75E-02 |
| P09005 | TEATAATGVATVIR | 0,PyMIC[AnyN-term]; | 8.6 | 0.0 | 0.0 | 3.81E-02 |
| P05544 | TEATAATGVATVIR | 0,PyMIC[AnyN-term]; | 8.6 | 0.0 | 0.0 | 3.81E-02 |
| P27590 | INFECSYPLDMK | 5,Carboxymethyl_13C(2)[C]; | 12.4 | 0.0 | 0.0 | 3.82E-02 |
| Q64230 | NLFQGDILLPR | 1,NeuAc[N]; | 3.0 | 0.0 | 0.0 | 3.82E-02 |
| Q01177 | GCVANPHSWPWQISLR | 0,LG-pyrrole[AnyN-term]; | 3.0 | 0.0 | 0.0 | 3.82E-02 |
| P98158 | IASANMDGTSLK | 5,Deamidated[N]; | 3.0 | 0.0 | 0.0 | 3.82E-02 |
| P50123 | MLQDWITPEK | 2,Xle->Cys[L]; | 3.0 | 0.0 | 0.0 | 3.82E-02 |
| P26051 | YAGVFHVEKNGR | 10,Deamidated[N]; | 3.0 | 0.0 | 0.0 | 3.82E-02 |
| P19218 | YCTDPSTAPK | 2,Carbamidomethyl[C]; | 3.0 | 0.0 | 0.0 | 3.82E-02 |
| P02782 | ANASQICELVAHETISFLMK | 7,SulfoGMBS[C]; | 6.0 | 0.0 | 0.0 | 3.82E-02 |
| Q6IUU3 | SYVQSFFGCR | 9,Carbamidomethyl[C]; | 13.3 | 0.0 | 0.0 | 3.83E-02 |
| P50399 | FLMANGQLVK | 5,Deamidated[N]; | 7.7 | 0.0 | 0.0 | 3.83E-02 |
| P50398 | FLMANGQLVK | 5,Deamidated[N]; | 7.7 | 0.0 | 0.0 | 3.83E-02 |
| Q68FP1 | PKAGALNSNDAFVLK | 0,Unknown_420[AnyN-term]; | 11.1 | 0.0 | 0.0 | 3.88E-02 |
| Q62740 | ESGDPSTCAFQR | 8,Carbamidomethyl[C]; | 5.6 | 0.0 | 0.0 | 3.88E-02 |
| P83121 | EICAWVVVTTR | 0,Acetyl[ProteinN-term];3,Carbamidomethyl[C]; | 5.6 | 0.0 | 0.0 | 3.88E-02 |
| P81828 | EICAWVVVTTR | 0,Acetyl[ProteinN-term];3,Carbamidomethyl[C]; | 5.6 | 0.0 | 0.0 | 3.88E-02 |
| P81827 | EICAWVVVTTR | 0,Acetyl[ProteinN-term];3,Carbamidomethyl[C]; | 5.6 | 0.0 | 0.0 | 3.88E-02 |
| P01836 | RDGVLDSVTDQDSK | 14,Lys->AminoadipicAcid[K]; | 10.7 | 0.0 | 0.0 | 3.93E-02 |
| P01835 | RDGVLDSVTDQDSK | 14,Lys->AminoadipicAcid[K]; | 10.7 | 0.0 | 0.0 | 3.93E-02 |
| P02761 | GETFQLMVLYGR | 0,Ethylphosphate[AnyN-term]; | 18.0 | 0.0 | 0.0 | 4.08E-02 |
| P02770 | AADKDNCFATEGPNLVAR | 8,Phe->Trp[F]; | 8.6 | 0.0 | 0.0 | 4.09E-02 |
| P02761 | NGETFQLMVLYGR | 1,Deamidated[N];8,Oxidation[M]; | 15.4 | 0.0 | 0.0 | 4.09E-02 |
| P13432 | NPPTQLHSTEQANTK | 7,His->Xle[H]; | 12.9 | 0.0 | 0.0 | 4.15E-02 |
| P02781 | ELEEFDAPPEAVEANLK | 6,Asp->Gln[D]; | 6.4 | 0.0 | 0.0 | 4.15E-02 |
| P02761 | VFMQHIDVLENSLGFK | 5,His->Xle[H]; | 13.3 | 0.0 | 0.0 | 4.17E-02 |
| P63018 | VCNPIITK | 2,Carbamidomethyl[C]; | 4.7 | 0.0 | 0.0 | 4.24E-02 |
| Q9R0T4 | TVNRDTGVISVVTSGLDR | 0,Unknown_250[AnyN-term]; | 24.0 | 0.0 | 0.0 | 4.25E-02 |
| P27590 | STDYGAGYSCDSDMHGWYR | 10,Cys->Dha[C]; | 12.0 | 0.0 | 0.0 | 4.25E-02 |
| O70244 | FCTSLSTPPLK | 2,Carbamidomethyl[C]; | 7.3 | 0.0 | 0.0 | 4.27E-02 |
| P23680 | VGQYSLYIGNSK | 10,Deamidated[N]; | 12.4 | 0.0 | 0.0 | 4.35E-02 |
| Q68FP1 | PSAAYLWVGTGASDAEK | 2,Pentose[S]; | 8.6 | 0.0 | 0.0 | 4.37E-02 |
| P24090 | QQAEHAVEGDCDFHILK | 11,Carbamidomethyl[C]; | 8.6 | 0.0 | 0.0 | 4.37E-02 |
| P02761 | TPEDGEYFVEYDGGNTFTILK | 12,Cation_Na[D]; | 25.7 | 0.0 | 0.0 | 4.37E-02 |
| P05545 | IKDLTTPYIRDEELSCSVLELK | 16,Carbamidomethyl[C]; | 178.7 | 0.0 | 0.0 | 4.37E-02 |
| P02761 | NGETFQLMVLYGR | 8,Oxidation[M]; | 44.6 | 0.0 | 0.0 | 4.39E-02 |
| P27590 | CLWSTEIQVK | 1,Carbamyl[C]; | 18.0 | 0.0 | 0.0 | 4.40E-02 |
| P0DP31 | VFDKDGNGYISAAELR | 7,Deamidated[N]; | 6.0 | 0.0 | 0.0 | 4.40E-02 |
| P0DP30 | VFDKDGNGYISAAELR | 7,Deamidated[N]; | 6.0 | 0.0 | 0.0 | 4.40E-02 |
| P0DP29 | VFDKDGNGYISAAELR | 7,Deamidated[N]; | 6.0 | 0.0 | 0.0 | 4.40E-02 |
| Q63041 | QNNICFDNWWVDEYHTQADHSAAR | 5,Carbamidomethyl[C]; | 21.4 | 0.0 | 0.0 | 4.42E-02 |
| Q9JHB9 | SGCSILDEVIR | 3,Cys->Dha[C]; | 2.6 | 0.0 | 0.0 | 4.53E-02 |
| Q62687 | SGMDPLVDIEDERPK | 1,Hex(2)[S]; | 2.6 | 0.0 | 0.0 | 4.53E-02 |
| P83121 | PDEICAWVVVTTR | 0,Acetyl[ProteinN-term];3,Cation_Na[E];5,Carbamidomethyl[C]; | 2.6 | 0.0 | 0.0 | 4.53E-02 |
| P81828 | PDEICAWVVVTTR | 0,Acetyl[ProteinN-term];3,Cation_Na[E];5,Carbamidomethyl[C]; | 2.6 | 0.0 | 0.0 | 4.53E-02 |
| P81827 | PDEICAWVVVTTR | 0,Acetyl[ProteinN-term];3,Cation_Na[E];5,Carbamidomethyl[C]; | 2.6 | 0.0 | 0.0 | 4.53E-02 |
| P36953 | RPCFEHLK | 3,Carbamidomethyl[C]; | 2.6 | 0.0 | 0.0 | 4.53E-02 |
| P36953 | SLAMVQQECK | 4,Oxidation[M];9,Carbamidomethyl[C]; | 2.6 | 0.0 | 0.0 | 4.53E-02 |
| P22282 | GETMYYISLPGSVR | 0,Triton[AnyN-term]; | 2.6 | 0.0 | 0.0 | 4.53E-02 |
| P15083 | ETCDVIINTLGK | 0,Propyl_2H(6)[AnyN-term]; | 2.6 | 0.0 | 0.0 | 4.53E-02 |
| P13432 | TQLHSTEQANTK | 4,His->Xle[H]; | 2.6 | 0.0 | 0.0 | 4.53E-02 |
| P07861 | QSNSLNYGGIGMVIGHEITHGFDDNGR | 3,Deamidated[N]; | 2.6 | 0.0 | 0.0 | 4.53E-02 |
| P05545 | KIFSQQADLSR | 1,NHS-fluorescein[K]; | 2.6 | 0.0 | 0.0 | 4.53E-02 |
| P02780 | SGCSILDEVIR | 3,Cys->Dha[C]; | 2.6 | 0.0 | 0.0 | 4.53E-02 |
| P02761 | TKDLSSDIKEK | 0,Carbamyl[AnyN-term]; | 2.6 | 0.0 | 0.0 | 4.53E-02 |
| P81828 | QKCQTYPDEICAWVVVTTR | 11,Nethylmaleimide[C]; | 9.9 | 0.0 | 0.0 | 4.53E-02 |
| P12788 | HPEYNKDTLDNDIMLIK | 6,TMT[K]; | 5.6 | 0.0 | 0.0 | 4.54E-02 |
| P02761 | HIDVLENSLGFK | 1,Dimethylphosphothione[H]; | 5.6 | 0.0 | 0.0 | 4.54E-02 |
| P00758 | QPGDDYSNDLMLLHLSQPADITDGVK | 4,Cation_Na[D]; | 5.6 | 0.0 | 0.0 | 4.54E-02 |
| Q99041 | TINFQIECTPVK | 8,Carbamidomethyl[C]; | 8.1 | 0.0 | 0.0 | 4.64E-02 |
| P01835 | ADYESHNLYTCEVVHK | 11,Carbamidomethyl[C];12,Cation_Na[E]; | 16.3 | 0.0 | 0.0 | 4.64E-02 |
| Q9QZA2 | LLDEEEATDNDLR | 1,Xle->Thr[L]; | 2.1 | 0.0 | 0.0 | 4.65E-02 |
| Q9JJ19 | IVEVNGVCMEGK | 8,Carbamidomethyl[C];9,Oxidation[M]; | 2.1 | 0.0 | 0.0 | 4.65E-02 |
| Q63041 | DAVNEEESVHWQRPK | 0,Formyl[AnyN-term]; | 2.1 | 0.0 | 0.0 | 4.65E-02 |
| Q4FZV0 | QHHGNGNNEMLHQVQLHFQLPQR | 0,Gln->pyro-Glu[AnyN-termQ];5,Deamidated[N]; | 2.1 | 0.0 | 0.0 | 4.65E-02 |
| P81828 | LLMSSLLALQCFRCTSFDSTGFCHVGR | 11,GG[C](Dicarbamidomethyl[C]);14,Carbamidomethyl[C];23,Carbamidomethyl[C]; | 2.1 | 0.0 | 0.0 | 4.65E-02 |
| P63259 | IVTNWDDMEK | 0,Propionyl_13C(3)[AnyN-term]; | 2.1 | 0.0 | 0.0 | 4.65E-02 |
| P60711 | IVTNWDDMEK | 0,Propionyl_13C(3)[AnyN-term]; | 2.1 | 0.0 | 0.0 | 4.65E-02 |
| P51635 | TASSVLLHTGQK | 0,Methylthio[AnyN-term]; | 2.1 | 0.0 | 0.0 | 4.65E-02 |
| P36953 | TINPTVDHCCR | 9,Carbamidomethyl[C];10,Carbamidomethyl[C]; | 4.3 | 0.0 | 0.0 | 4.65E-02 |
| P29534 | NTEISVHPSTR | 1,Asn->Cys[N]; | 2.1 | 0.0 | 0.0 | 4.65E-02 |
| P27590 | HTEDTTIQVTENGESSQAR | 0,AccQTag[AnyN-term]; | 2.1 | 0.0 | 0.0 | 4.65E-02 |
| P24090 | QQAEHAVEGDCDFHILK | 0,Gln->pyro-Glu[AnyN-termQ];11,Carbamidomethyl[C]; | 2.1 | 0.0 | 0.0 | 4.65E-02 |
| P20760 | KETWQQGNTFTCSVLHEGLHNHHTEK | 12,Carbamidomethyl[C]; | 2.1 | 0.0 | 0.0 | 4.65E-02 |
| P15684 | VVATTQMQAADAR | 7,Oxidation[M]; | 2.1 | 0.0 | 0.0 | 4.65E-02 |
| P15083 | YWCHWEADENGR | 3,Carbamidomethyl[C]; | 2.1 | 0.0 | 0.0 | 4.65E-02 |
| P10959 | PPEPAEPWSFVK | 0,Iminobiotin[AnyN-term]; | 2.1 | 0.0 | 0.0 | 4.65E-02 |
| P07522 | NCQSGPLER | 1,Deamidated[N];2,Carbamidomethyl[C]; | 2.1 | 0.0 | 0.0 | 4.65E-02 |
| P02782 | SQICELVAHETISFLMK | 5,Carboxy[E]; | 2.1 | 0.0 | 0.0 | 4.65E-02 |
| P02761 | VFMQHIDVLENSLGFK | 12,Formyl[S](Ser->Asp[S]); | 2.1 | 0.0 | 0.0 | 4.65E-02 |
| P01836 | RDGVLDSVTDQDSK | 14,Lys->Xle[K]; | 2.1 | 0.0 | 0.0 | 4.65E-02 |
| P01835 | RDGVLDSVTDQDSK | 14,Lys->Xle[K]; | 2.1 | 0.0 | 0.0 | 4.65E-02 |
| P00758 | NNLYEDEPFAQHR | 4,Sulfo[Y]; | 6.4 | 0.0 | 0.0 | 4.65E-02 |
| O88766 | CGVPDSGDFLLTPGSPK | 1,Carbamidomethyl[C]; | 2.1 | 0.0 | 0.0 | 4.65E-02 |
| P14841 | APQEADASEEGVQR | 1,MurNAc[A]; | 3.4 | 0.0 | 0.0 | 4.72E-02 |
| P11232 | MIKPFFHSLCDK | 10,Carbamidomethyl[C]; | 13.7 | 0.0 | 0.0 | 4.72E-02 |
| P07861 | SESQMDITDINAPKPK | 2,Cation_Na[E]; | 3.4 | 0.0 | 0.0 | 4.72E-02 |
| P02780 | VKPYVQDHFTEK | 0,Succinyl_2H(4)[AnyN-term]; | 3.4 | 0.0 | 0.0 | 4.72E-02 |
| P02770 | PEAEAMCTSFQENPTSFLGHYLHEVAR | 0,NHS-LC-Biotin[AnyN-term]; | 3.4 | 0.0 | 0.0 | 4.72E-02 |
| P48199 | YETHGDVFIKPQLWPLTDCCES | 19,Carbamidomethyl[C];20,Carbamidomethyl[C]; | 15.0 | 0.0 | 0.0 | 4.84E-02 |
| Q64230 | LADNLDLNAK | 0,Ethylphosphate[AnyN-term]; | 3.9 | 0.0 | 0.0 | 4.88E-02 |
| Q63772 | MQCFSVTER | 3,Carbamidomethyl[C]; | 3.9 | 0.0 | 0.0 | 4.88E-02 |
| Q63691 | VPLQALDLSHNSLR | 11,Deamidated[N]; | 3.9 | 0.0 | 0.0 | 4.88E-02 |
| Q63041 | THITNAFNWLSMK | 12,Oxidation[M]; | 3.9 | 0.0 | 0.0 | 4.88E-02 |
| P80067 | GTDECAIESIAMAAIPIPK | 5,Carbamidomethyl[C]; | 3.9 | 0.0 | 0.0 | 4.88E-02 |
| P22283 | CPFEEQTEQLKR | 0,Pyro-carbamidomethyl[AnyN-termC]; | 3.9 | 0.0 | 0.0 | 4.88E-02 |
| P22282 | CPFEEQTEQLKR | 0,Pyro-carbamidomethyl[AnyN-termC]; | 3.9 | 0.0 | 0.0 | 4.88E-02 |
| P19218 | CLLGGLGFK | 1,Carbamidomethyl[C]; | 3.9 | 0.0 | 0.0 | 4.88E-02 |
| P08723 | LCLTSLSFTTNK | 2,Trioxidation[C]; | 3.9 | 0.0 | 0.0 | 4.88E-02 |
| P07647 | DHNNDLMLLHLSKPADITGGVK | 13,Xlink_BS2G[217][K]; | 3.9 | 0.0 | 0.0 | 4.88E-02 |
| P05545 | IFSQQADLSR | 0,C+12[AnyN-term]; | 3.9 | 0.0 | 0.0 | 4.88E-02 |
| Q68FP1 | QVEEGSEPDGFWEALGGK | 6,Hex(1)Pent(2)[S]; | 12.4 | 0.0 | 0.0 | 4.89E-02 |
| P98158 | GSYECFCVDGFK | 5,Carbamidomethyl[C];7,Carbamidomethyl[C]; | 19.7 | 0.0 | 0.0 | 4.91E-02 |
| P02780 | LVKPYVQDHFTEK | 2,Val->His[V]; | 17.1 | 0.0 | 0.0 | 4.94E-02 |
| P27590 | GDGWYCECSPGFCEPGLDCLPQGPSGK | 6,Carbamidomethyl[C];8,Carbamidomethyl[C];19,Carbamidomethyl[C]; | 8.1 | 0.0 | 0.0 | 4.96E-02 |
| P00758 | CATDNYQVWLGR | 0,Propionamide[AnyN-term]; | 9.9 | 0.0 | 0.0 | 4.97E-02 |
| P00689 | CGSGNSAGTHSTCGSYFNPNNR | 0,Pyro-carbamidomethyl[AnyN-termC]; | 6.0 | 0.0 | 0.0 | 4.98E-02 |
| P81828 | QKCQTYPDEICAWVVVTTR | 3,ethylsulfonylethyl[C](BEMAD_C[C]); | 138.9 | 1.3 | 0.0 | 3.72E-02 |
| P02770 | CFATEGPNLVAR | 0,Methyl[AnyN-term]; | 15.0 | 0.2 | 0.0 | 2.81E-02 |
| P97546 | AEDSGEYHCVYHFVSAPK | 9,Carbamidomethyl[C]; | 14.6 | 0.2 | 0.0 | 8.88E-03 |
| P20059 | ECQSEGVLFFQGNR | 2,maleimide[C]; | 9.9 | 0.2 | 0.0 | 1.41E-02 |
| Q63556 | DEELSCSVLELK | 6,Carbamidomethyl[C]; | 54.0 | 1.0 | 0.0 | 1.59E-03 |
| P09005 | DEELSCSVLELK | 6,Carbamidomethyl[C]; | 54.0 | 1.0 | 0.0 | 1.59E-03 |
| P05545 | DEELSCSVLELK | 6,Carbamidomethyl[C]; | 54.0 | 1.0 | 0.0 | 1.59E-03 |
| P05544 | DEELSCSVLELK | 6,Carbamidomethyl[C]; | 54.0 | 1.0 | 0.0 | 1.59E-03 |
| P12346 | TSYQDCIK | 6,Carbamidomethyl[C]; | 7.3 | 0.2 | 0.0 | 5.94E-04 |
| P98158 | VCNHVDDCSDNSDEK | 2,Carbamidomethyl[C];8,Carbamidomethyl[C]; | 7.3 | 0.2 | 0.0 | 3.41E-02 |
| P22282 | LDNCPFEEQTEQLK | 4,Carbamidomethyl[C]; | 103.7 | 5.0 | 0.0 | 1.64E-02 |
| P15684 | VLRMLSSFLTEDLFK | 6,Hex(1)HexA(1)[S]; | 14.6 | 1.0 | 0.1 | 1.50E-03 |
| P68136 | DLTDYLMK | 7,Oxidation[M]; | 4.7 | 0.3 | 0.1 | 4.32E-02 |
| P68035 | DLTDYLMK | 7,Oxidation[M]; | 4.7 | 0.3 | 0.1 | 4.32E-02 |
| P63269 | DLTDYLMK | 7,Oxidation[M]; | 4.7 | 0.3 | 0.1 | 4.32E-02 |
| P63259 | DLTDYLMK | 7,Oxidation[M]; | 4.7 | 0.3 | 0.1 | 4.32E-02 |
| P62738 | DLTDYLMK | 7,Oxidation[M]; | 4.7 | 0.3 | 0.1 | 4.32E-02 |
| P60711 | DLTDYLMK | 7,Oxidation[M]; | 4.7 | 0.3 | 0.1 | 4.32E-02 |
| P36953 | DLLEDCCNR | 6,Carbamidomethyl[C];7,Carbamidomethyl[C]; | 16.3 | 1.3 | 0.1 | 3.83E-03 |
| P10252 | ILEYFPNGK | 7,Deamidated[N]; | 26.1 | 2.3 | 0.1 | 3.00E-02 |
| P27590 | CPHTEDTTIQVTENGESSQAR | 1,Carbamidomethyl[C];14,Deamidated[N]; | 194.6 | 20.3 | 0.1 | 6.19E-03 |
| Q01460 | DDVCAIAK | 4,Carbamidomethyl[C]; | 3.0 | 0.3 | 0.1 | 2.78E-02 |
| P36374 | NNLLEDEPFAQHR | 2,Ammonia-loss[N]; | 12.0 | 1.3 | 0.1 | 3.30E-02 |
| P36373 | NNLLEDEPFAQHR | 2,Ammonia-loss[N]; | 12.0 | 1.3 | 0.1 | 3.30E-02 |
| P36374 | NNLLEDEPFAQHR | 1,Ammonia-loss[N]; | 8.1 | 1.2 | 0.1 | 2.03E-02 |
| P36373 | NNLLEDEPFAQHR | 1,Ammonia-loss[N]; | 8.1 | 1.2 | 0.1 | 2.03E-02 |
| P27590 | DWMSIVTPAR | 3,Oxidation[M]; | 168.9 | 27.5 | 0.2 | 1.53E-02 |
| P82450 | TFEISCCSDHQCK | 6,Carbamidomethyl[C];7,Carbamidomethyl[C];12,Carbamidomethyl[C]; | 9.0 | 1.5 | 0.2 | 2.53E-03 |
| P21704 | LTNQMAEAISDHYPVEVTLR | 7,Cation_Na[E]; | 15.9 | 3.3 | 0.2 | 3.75E-02 |
| P27590 | CPHTEDTTIQVTENGESSQAR | 1,Carbamidomethyl[C]; | 602.6 | 144.8 | 0.2 | 3.62E-02 |
| P18292 | REECSIPVCGQEGR | 4,Carbamidomethyl[C];9,Carbamidomethyl[C]; | 37.3 | 9.2 | 0.2 | 3.07E-02 |
| O55006 | IHAMGCATQTACNLK | 6,Carbamidomethyl[C];12,Carbamidomethyl[C]; | 35.6 | 9.0 | 0.3 | 3.22E-02 |
| P02770 | AETFTFHSDICTLPDKEK | 11,Carbamidomethyl[C]; | 234.9 | 60.7 | 0.3 | 1.02E-02 |
| P02770 | ECCHGDLLECADDR | 2,Carbamidomethyl[C];3,Carbamidomethyl[C];10,Carbamidomethyl[C]; | 49.7 | 13.0 | 0.3 | 1.09E-03 |
| P14046 | ISLCHGNPTFSSETK | 4,Carbamidomethyl[C]; | 18.4 | 5.0 | 0.3 | 4.78E-02 |
| P02761 | EKIEENGSMR | 6,Deamidated[N]; | 155.1 | 42.7 | 0.3 | 1.60E-02 |
| P27590 | MAETCVPVLR | 1,Oxidation[M];5,Carbamidomethyl[C]; | 148.3 | 41.8 | 0.3 | 1.45E-03 |
| P29598 | KPSSTVDQQGFQCGQK | 13,Carbamidomethyl[C]; | 13.3 | 4.0 | 0.3 | 5.25E-03 |
| P00758 | CVEAHKEEVTDLMLCAGEMDGGK | 1,Carbamidomethyl[C];15,Carbamidomethyl[C]; | 51.0 | 16.0 | 0.3 | 3.00E-02 |
| P02761 | VFMQHIDVLENSLGFK | 11,Deamidated[N]; | 49.7 | 18.0 | 0.4 | 2.90E-03 |
| P02761 | LNGDWFSIVVASNKR | 2,Deamidated[N]; | 578.6 | 216.5 | 0.4 | 6.66E-04 |
| P02770 | RPCFSALTVDETYVPK | 3,Cys->Dha[C]; | 9.0 | 3.5 | 0.4 | 2.60E-02 |
| P14841 | SQTNLTNCPFHDQPHLMR | 8,Carbamidomethyl[C]; | 11.6 | 19.2 | 1.7 | 2.59E-02 |
| P20059 | DYFISCPGR | 6,Carbamidomethyl[C]; | 34.3 | 58.5 | 1.7 | 1.31E-02 |
| P10960 | SLPCDICK | 4,Carbamidomethyl[C];7,Carbamidomethyl[C]; | 9.4 | 16.2 | 1.7 | 1.38E-02 |
| Q01177 | TPENFPCK | 7,Carbamidomethyl[C]; | 10.3 | 20.5 | 2.0 | 1.49E-04 |
| P10960 | LGPGVSDICK | 9,Carbamidomethyl[C]; | 10.7 | 21.5 | 2.0 | 1.99E-04 |
| P05371 | TQQYNELLHSLQSK | 0,Succinyl_2H(4)[AnyN-term]; | 4.3 | 12.2 | 2.8 | 1.14E-02 |
| P02761 | QHIDVLENSLGFK | 0,Gln->pyro-Glu[AnyN-termQ]; | 7.3 | 22.7 | 3.1 | 6.48E-03 |
| P02764 | QQLELEKETK | 0,Gln->pyro-Glu[AnyN-termQ]; | 6.9 | 24.2 | 3.5 | 2.63E-03 |
| P16391 | EGPEYWEQQTR | 8,Deamidated[Q]; | 3.9 | 15.2 | 3.9 | 4.88E-03 |
| P02761 | YVMFHLINFK | 3,Oxidation[M]; | 53.6 | 211.2 | 3.9 | 2.79E-06 |
| P00758 | CATDNYQVWLGR | 0,Pyro-carbamidomethyl[AnyN-termC]; | 3.0 | 13.5 | 4.5 | 8.41E-04 |
| P02770 | DDNPNLPPFQRPEAEAMCTSFQENPTSFLGHYLHEVAR | 18,Carbamidomethyl[C]; | 9.9 | 45.7 | 4.6 | 1.92E-03 |
| P02761 | NGDWFSIVVASNKR | 0,Methylthio[AnyN-term]; | 1.7 | 8.7 | 5.1 | 5.85E-03 |
| P02761 | VFMQHIDVLENSLGFK | 7,Cation_Ca[II][D]; | 13.3 | 69.8 | 5.3 | 4.03E-04 |
| P02761 | VFMQHIDVLENSLGFKFR | 0,ISD_z+2_ion[AnyN-term]; | 1.7 | 9.7 | 5.6 | 1.54E-02 |
| O70417 | VDGGFNYIQTR | 2,SulfanilicAcid[D]; | 2.6 | 16.0 | 6.2 | 2.59E-04 |
| P02761 | LCEAHGITR | 0,Methyl_2H(2)[AnyN-term]; | 1.7 | 11.3 | 6.6 | 3.57E-02 |
| P07647 | VSQSFQHPDYIPVFMR | 0,PyMIC[AnyN-term]; | 0.9 | 6.3 | 7.4 | 2.36E-02 |
| P02761 | GNLDVAKLNGDWFSIVVASNKR | 7,monomethylphosphothione[K]; | 1.3 | 10.0 | 7.8 | 2.81E-02 |
| P36953 | ADWCQPLK | 4,Carbamidomethyl[C]; | 0.4 | 3.3 | 7.8 | 5.46E-03 |
| P07647 | NLVKDEPFAQR | 0,Ethylphosphate[AnyN-term]; | 1.7 | 14.2 | 8.3 | 5.03E-03 |
| Q64230 | GDPGNSDGGIYLDDITLTETPCPTGVWTIR | 22,Carbamidomethyl[C]; | 0.9 | 7.3 | 8.6 | 8.64E-05 |
| P05371 | LTQQYNELLHSLQSK | 1,Xle->Cys[L]; | 0.4 | 4.3 | 10.1 | 1.03E-02 |
| P02770 | LPEAQRLPCVEDYLSAILNR | 9,CarbamidomethylDTT[C]; | 0.4 | 4.5 | 10.5 | 4.01E-04 |
| P12020 | CIYNHSPLQHR | 1,Carbamidomethyl[C]; | 1.3 | 14.0 | 10.9 | 2.30E-02 |
| P83121 | ICAWVVVTTR | 2,Trioxidation[C]; | 0.4 | 5.0 | 11.7 | 7.89E-03 |
| P81828 | ICAWVVVTTR | 2,Trioxidation[C]; | 0.4 | 5.0 | 11.7 | 7.89E-03 |
| P81827 | ICAWVVVTTR | 2,Trioxidation[C]; | 0.4 | 5.0 | 11.7 | 7.89E-03 |
| P02761 | ELYLVAYK | 1,Cation_Na[E]; | 9.0 | 107.7 | 12.0 | 4.54E-07 |
| P02782 | CELVAHETISFLMK | 1,Carbamidomethyl[C]; | 1.3 | 16.2 | 12.6 | 1.89E-03 |
| P14841 | QEADASEEGVQR | 0,NIC[AnyN-term](ICPL[AnyN-term]); | 0.4 | 5.5 | 12.8 | 1.11E-02 |
| Q68FP1 | EGSEPDGFWEALGGK | 0,Glu->pyro-Glu+Methyl[AnyN-termE]; | 2.6 | 35.7 | 13.9 | 2.59E-07 |
| P05371 | TQQYNELLHSLQSK | 0,NIC[AnyN-term](ICPL[AnyN-term]); | 0.4 | 6.2 | 14.4 | 3.57E-02 |
| P02761 | FMQHIDVLENSLGFK | 2,Oxidation[M]; | 6.4 | 93.5 | 14.5 | 7.24E-05 |
| P07647 | VSQSFQHPDYIPVFMR | 2,O-Et-N-diMePhospho[S]; | 0.9 | 16.5 | 19.3 | 5.14E-03 |
| P02761 | TDYDRYVMFHLINFK | 8,Oxidation[M]; | 1.7 | 35.7 | 20.8 | 1.06E-06 |
| P02782 | LEMYNAPPAAVEAK | 0,Diethylphosphate[AnyN-term]; | 1.7 | 40.3 | 23.5 | 1.60E-04 |
| P36374 | NLLEDEPFAQHR | 0,Succinyl_2H(4)[AnyN-term]; | 0.4 | 10.5 | 24.5 | 1.44E-02 |
| P36373 | NLLEDEPFAQHR | 0,Succinyl_2H(4)[AnyN-term]; | 0.4 | 10.5 | 24.5 | 1.44E-02 |
| P02782 | LEMYNAPPAAVEAK | 0,CAF[AnyN-term]; | 0.4 | 10.7 | 24.9 | 4.48E-02 |
| P29598 | CGGSLISPCWVASATHCFVNQPK | 1,Carbamidomethyl[C];9,Carbamidomethyl[C];17,Carbamidomethyl[C]; | 0.4 | 11.0 | 25.7 | 6.96E-04 |
| P02761 | LCEAHGITR | 2,Methylthio[C]; | 0.9 | 22.2 | 25.9 | 3.72E-03 |
| P02770 | LPEAQRLPCVEDYLSAILNR | 9,CarboxymethylDTT[C]; | 0.4 | 11.2 | 26.1 | 7.53E-03 |
| Q9JMI9 | YAIAVVLSFSR | 10,HexNAc(1)dHex(1)[S]; | 3.0 | 83.7 | 27.9 | 8.80E-03 |
| P02761 | FVEYDGGNTFTILK | 0,Propionamide[AnyN-term]; | 0.4 | 13.5 | 31.5 | 2.22E-04 |
| P02761 | LCEAHGITR | 0,Methyl_2H(3)13C(1)[AnyN-term]; | 0.9 | 31.7 | 36.9 | 3.24E-02 |
| P02761 | FAKLCEAHGITR | 0,CLIP_TRAQ_2[AnyN-term]; | 0.4 | 18.0 | 42.0 | 3.13E-03 |
| P02782 | SEEELKKELEMYNAPPAAVEAK | 10,Cation_Ca[II][E]; | 0.4 | 19.8 | 46.3 | 2.67E-02 |
| P02761 | LNGDWFSIVVASNKR | 2,Deamidated_18O(1)[N](Delta_H(1)N(-1)18O(1)[N]); | 0.4 | 25.8 | 60.3 | 1.60E-04 |
| P02761 | LCEAHGITR | 2,CarbamidomethylDTT[C]; | 5.1 | 375.3 | 73.0 | 5.79E-03 |
| P22283 | LVNCPFEEQTEQLKR | 4,Carbamidomethyl[C]; | 0.9 | 115.2 | 134.4 | 9.54E-04 |
| P00787 | EQWSNCPTIAQIR | 6,Carbamidomethyl[C]; | 0.0 | 32.7 | #DIV/0! | 2.75E-08 |
| P09005 | MQQVESSLQPETLKK | 5,Cation_Na[E]; | 0.0 | 19.5 | #DIV/0! | 3.12E-08 |
| P05545 | MQQVESSLQPETLKK | 5,Cation_Na[E]; | 0.0 | 19.5 | #DIV/0! | 3.12E-08 |
| P05544 | MQQVESSLQPETLKK | 5,Cation_Na[E]; | 0.0 | 19.5 | #DIV/0! | 3.12E-08 |
| Q9R0T4 | DTANWLEINPETGVISTR | 1,Cation_Na[D]; | 0.0 | 16.3 | #DIV/0! | 2.16E-07 |
| P02770 | AADKDNCFATEGPNLVAR | 7,Carbamidomethyl[C]; | 0.0 | 36.7 | #DIV/0! | 2.69E-07 |
| Q64240 | TIAACNLPIVQGPCR | 5,Carbamidomethyl[C];14,Carbamidomethyl[C]; | 0.0 | 61.7 | #DIV/0! | 4.21E-07 |
| P17475 | VFNNDADLSGITEDAPLK | 7,Cation_Na[D]; | 0.0 | 17.7 | #DIV/0! | 5.89E-07 |
| P02761 | VFMQHIDVLENSLGFK | 3,Oxidation[M]; | 0.0 | 1585.3 | #DIV/0! | 6.22E-07 |
| P63259 | QEYDESGPSIVHR | 0,Gln->pyro-Glu[AnyN-termQ]; | 0.0 | 17.5 | #DIV/0! | 6.45E-07 |
| P60711 | QEYDESGPSIVHR | 0,Gln->pyro-Glu[AnyN-termQ]; | 0.0 | 17.5 | #DIV/0! | 6.45E-07 |
| P08932 | FSVATQICNITPGK | 8,Carbamidomethyl[C]; | 0.0 | 24.5 | #DIV/0! | 7.24E-07 |
| P01048 | FSVATQICNITPGK | 8,Carbamidomethyl[C]; | 0.0 | 24.5 | #DIV/0! | 7.24E-07 |
| Q64240 | TIAACNLPIVQGPCR | 5,Carbamidomethyl[C];6,Deamidated[N];14,Carbamidomethyl[C]; | 0.0 | 26.3 | #DIV/0! | 1.17E-06 |
| P02770 | AADKDNCFATEGPNLVAR | 6,Deamidated[N];7,Carbamidomethyl[C]; | 0.0 | 40.2 | #DIV/0! | 1.33E-06 |
| P01835 | RRDGVLDSVTDQDSK | 8,HexNAc(1)dHex(1)[S]; | 0.0 | 12.0 | #DIV/0! | 1.34E-06 |
| P02770 | APQVSTPTLVEAAR | 0,ISD_z+2_ion[AnyN-term]; | 0.0 | 21.3 | #DIV/0! | 1.80E-06 |
| P14740 | CLSCDLNPER | 1,Carbamidomethyl[C];4,Carbamidomethyl[C]; | 0.0 | 14.8 | #DIV/0! | 2.04E-06 |
| P15083 | WSNDGCHILPSHDEGAR | 6,Carbamidomethyl[C]; | 0.0 | 17.7 | #DIV/0! | 2.74E-06 |
| Q6IE52 | ALGCLEASWETIEQGR | 4,Carbamidomethyl[C]; | 0.0 | 12.5 | #DIV/0! | 3.43E-06 |
| Q03626 | ALGCLEASWETIEQGR | 4,Carbamidomethyl[C]; | 0.0 | 12.5 | #DIV/0! | 3.43E-06 |
| P02761 | CEAHGITR | 1,Glutathione[C]; | 0.0 | 27.7 | #DIV/0! | 3.58E-06 |
| P07522 | LFWTDTGMSPR | 8,Oxidation[M]; | 0.0 | 22.8 | #DIV/0! | 3.75E-06 |
| P81827 | CESLDSTGLCR | 1,Carbamidomethyl[C];10,Carbamidomethyl[C]; | 0.0 | 807.8 | #DIV/0! | 4.24E-06 |
| P02764 | EFQTTDDQCVYNFTHLGVQR | 9,Carbamidomethyl[C]; | 0.0 | 41.3 | #DIV/0! | 4.64E-06 |
| P00787 | GENHCGIESEIVAGIPR | 5,Carbamidomethyl[C]; | 0.0 | 31.8 | #DIV/0! | 4.85E-06 |
| P15083 | ETCDVIINTLGK | 3,Carbamidomethyl[C]; | 0.0 | 24.8 | #DIV/0! | 5.35E-06 |
| O70513 | LASAYGATELQGYCGR | 14,Carbamidomethyl[C]; | 0.0 | 25.0 | #DIV/0! | 5.51E-06 |
| P02761 | VFMQHIDVLENSLGFK | 3,Oxidation[M];11,Deamidated[N]; | 0.0 | 15.8 | #DIV/0! | 5.80E-06 |
| P02782 | ELEMYNAPPAAVEAK | 3,Cation_Ca[II][E]; | 0.0 | 39.5 | #DIV/0! | 8.60E-06 |
| P15083 | NKETCDVIINTLGK | 5,Carbamidomethyl[C]; | 0.0 | 14.7 | #DIV/0! | 9.03E-06 |
| P08426 | ANTFDNDIMLIK | 0,Ethylphosphate[AnyN-term]; | 0.0 | 24.8 | #DIV/0! | 1.14E-05 |
| P01836 | DSTYSMSSTLSLTK | 3,Dehydrated[T]; | 0.0 | 70.7 | #DIV/0! | 1.43E-05 |
| P01835 | DSTYSMSSTLSLTK | 3,Dehydrated[T]; | 0.0 | 70.7 | #DIV/0! | 1.43E-05 |
| P07151 | IPNIEMSDLSFSK | 1,Xle->Pro[I]; | 0.0 | 29.3 | #DIV/0! | 1.51E-05 |
| P20059 | FNPVTGEVPPR | 2,Asn->Met[N]; | 0.0 | 40.2 | #DIV/0! | 1.51E-05 |
| Q01460 | YDSELMCYAHSK | 7,Carbamidomethyl[C]; | 0.0 | 12.2 | #DIV/0! | 1.61E-05 |
| P07632 | QDDLGKGGNEESTK | 0,Gln->pyro-Glu[AnyN-termQ]; | 0.0 | 5.3 | #DIV/0! | 1.74E-05 |
| P01836 | DSTYSMSSTLSLTK | 6,Oxidation[M]; | 0.0 | 291.7 | #DIV/0! | 1.81E-05 |
| P01835 | DSTYSMSSTLSLTK | 6,Oxidation[M]; | 0.0 | 291.7 | #DIV/0! | 1.81E-05 |
| P15684 | QVTPLFAYFK | 0,Gln->pyro-Glu[AnyN-termQ]; | 0.0 | 12.7 | #DIV/0! | 1.86E-05 |
| P04276 | VCQELSTLGKDDFR | 2,Carbamidomethyl[C]; | 0.0 | 14.0 | #DIV/0! | 1.93E-05 |
| P27274 | QVYQQCWR | 6,Carbamidomethyl[C]; | 0.0 | 25.5 | #DIV/0! | 2.17E-05 |
| P02761 | VFMQHIDVLENSLGFK | 0,PEITC[AnyN-term]; | 0.0 | 71.5 | #DIV/0! | 2.23E-05 |
| P06866 | YVMLPVADQEKCELHYEK | 12,Carbamidomethyl[C]; | 0.0 | 21.8 | #DIV/0! | 2.32E-05 |
| P00762 | LGEHNINVLEGDEQFINAAK | 12,Dehydrated[D]; | 0.0 | 14.3 | #DIV/0! | 2.37E-05 |
| P10960 | TCAWIHDSSLSASCK | 2,Carbamidomethyl[C];14,Carbamidomethyl[C]; | 0.0 | 23.3 | #DIV/0! | 2.44E-05 |
| P22057 | GHDTVQPNFQQDK | 0,GIST-Quat[AnyN-term]; | 0.0 | 12.5 | #DIV/0! | 2.58E-05 |
| P02770 | MSQRFPNAEFAEITK | 0,Phenylisocyanate_2H(5)[AnyN-term]; | 0.0 | 48.3 | #DIV/0! | 2.73E-05 |
| P00758 | NNLYEDEPFAQHR | 1,Deamidated[N]; | 0.0 | 18.3 | #DIV/0! | 3.02E-05 |
| P02761 | DLSSDIKEK | 1,Dehydrated[D]; | 0.0 | 17.0 | #DIV/0! | 3.30E-05 |
| Q499T2 | ATASLPQGSSEGAATCK | 16,Carbamidomethyl[C]; | 0.0 | 14.7 | #DIV/0! | 3.45E-05 |
| P12346 | DQYELLCLDNTR | 7,Carbamidomethyl[C]; | 0.0 | 14.5 | #DIV/0! | 3.69E-05 |
| P60711 | DDDIAALVVDNGSGMCK | 0,Acetyl[ProteinN-term];16,Carbamidomethyl[C]; | 0.0 | 10.0 | #DIV/0! | 3.73E-05 |
| P02782 | CVDQMSNGDR | 1,Carbamidomethyl[C];5,Oxidation[M]; | 0.0 | 8.2 | #DIV/0! | 3.87E-05 |
| P83121 | CISFDSTGFCYVGR | 1,Carbamidomethyl[C];10,Carbamidomethyl[C]; | 0.0 | 89.5 | #DIV/0! | 3.91E-05 |
| P20059 | DPGLSALLSDHR | 1,iodoTMT[D]; | 0.0 | 19.3 | #DIV/0! | 3.96E-05 |
| P98158 | CAPGYIREPDGK | 1,Carbamidomethyl[C]; | 0.0 | 11.3 | #DIV/0! | 4.08E-05 |
| P27274 | CYNCLDPVSSCK | 1,Carbamidomethyl[C];4,Carbamidomethyl[C];11,Carbamidomethyl[C]; | 0.0 | 59.7 | #DIV/0! | 4.64E-05 |
| P83121 | STGFCYVGR | 5,Carbamidomethyl[C]; | 0.0 | 11.3 | #DIV/0! | 5.08E-05 |
| P02770 | QTALAELVK | 0,Gln->pyro-Glu[AnyN-termQ]; | 0.0 | 25.2 | #DIV/0! | 5.96E-05 |
| Q68FP1 | AQHVQVEEGSEPDGFWEALGGK | 11,Cation_Ca[II][E]; | 0.0 | 18.0 | #DIV/0! | 6.23E-05 |
| Q64319 | DFTTTQVGMHDLVR | 0,ISD_z+2_ion[AnyN-term]; | 0.0 | 14.7 | #DIV/0! | 6.53E-05 |
| Q499T2 | SLPQGSSEGAATCK | 13,Carbamidomethyl[C]; | 0.0 | 9.7 | #DIV/0! | 6.97E-05 |
| P02761 | NLDVAKLNGDWFSIVVASNKR | 0,NIC[AnyN-term](ICPL[AnyN-term]); | 0.0 | 7.2 | #DIV/0! | 7.31E-05 |
| P00689 | QYDPHTADGR | 0,Gln->pyro-Glu[AnyN-termQ]; | 0.0 | 9.8 | #DIV/0! | 7.69E-05 |
| P02761 | AKLNGDWFSIVVASNK | 3,Xle->Ala[L]; | 0.0 | 8.8 | #DIV/0! | 7.72E-05 |
| P98158 | CAADGSPPLLLLPENVR | 1,Carbamidomethyl[C]; | 0.0 | 18.0 | #DIV/0! | 7.74E-05 |
| Q499T2 | ASLPQGSSEGAATCK | 14,Carbamidomethyl[C]; | 0.0 | 8.0 | #DIV/0! | 7.98E-05 |
| P02761 | EKIEENGSMR | 1,Cation_Ca[II][E]; | 0.0 | 6.5 | #DIV/0! | 8.53E-05 |
| P02761 | CEAHGITR | 0,Pyro-carbamidomethyl[AnyN-termC]; | 0.0 | 21.0 | #DIV/0! | 8.71E-05 |
| P02761 | LCEAHGITR | 0,dNIC[AnyN-term]; | 0.0 | 6.2 | #DIV/0! | 9.23E-05 |
| P02770 | GDFAQFVDK | 0,CLIP_TRAQ_2[AnyN-term]; | 0.0 | 9.8 | #DIV/0! | 9.60E-05 |
| O70417 | AYLISNTPVDGGFNYIQTR | 10,Cation_Na[D]; | 0.0 | 19.0 | #DIV/0! | 9.81E-05 |
| P17475 | FNNDADLSGITEDAPLK | 0,GIST-Quat_2H(3)[AnyN-term]; | 0.0 | 12.5 | #DIV/0! | 9.99E-05 |
| Q01177 | TAVTAAGTPCQEWAAQEPHSHR | 10,Carbamidomethyl[C]; | 0.0 | 30.7 | #DIV/0! | 1.02E-04 |
| P02761 | FMQHIDVLENSLGFK | 0,NDA[AnyN-term]; | 0.0 | 64.2 | #DIV/0! | 1.04E-04 |
| P07647 | VGSICLASGWGMTNPSEMK | 5,Carbamidomethyl[C];12,Oxidation[M]; | 0.0 | 19.5 | #DIV/0! | 1.06E-04 |
| P22282 | LDNCPFEEQTEQLKR | 4,NIPCAM[C]; | 0.0 | 46.3 | #DIV/0! | 1.07E-04 |
| P15978 | VEHEGLPEPLSQR | 0,C+12[AnyN-term]; | 0.0 | 23.5 | #DIV/0! | 1.17E-04 |
| P12346 | WCAVSEHENTK | 2,Carbamidomethyl[C]; | 0.0 | 10.7 | #DIV/0! | 1.19E-04 |
| P07632 | GVANVSIEDR | 0,dNIC[AnyN-term]; | 0.0 | 5.8 | #DIV/0! | 1.21E-04 |
| P22057 | DQGLTEEDIVFLPQPDK | 2,Gln->His[Q]; | 0.0 | 9.0 | #DIV/0! | 1.35E-04 |
| P01836 | RDGVLDSVTDQDSK | 0,Iodoacetanilide[AnyN-term]; | 0.0 | 6.0 | #DIV/0! | 1.42E-04 |
| P01835 | RDGVLDSVTDQDSK | 0,Iodoacetanilide[AnyN-term]; | 0.0 | 6.0 | #DIV/0! | 1.42E-04 |
| P07522 | DQILAGDNADLSKEVASLDNSPK | 12,Hex(2)[S]; | 0.0 | 12.5 | #DIV/0! | 1.59E-04 |
| P02761 | ELYLVAYK | 8,Acetylhypusine[K]; | 0.0 | 18.2 | #DIV/0! | 1.62E-04 |
| P02761 | GETFQLMVLYGR | 0,Phenylisocyanate_2H(5)[AnyN-term]; | 0.0 | 26.8 | #DIV/0! | 1.84E-04 |
| P20611 | CPLQDFLR | 1,Carbamidomethyl[C]; | 0.0 | 33.0 | #DIV/0! | 1.93E-04 |
| P24090 | CNLLAEK | 1,Carbamidomethyl[C]; | 0.0 | 36.0 | #DIV/0! | 1.94E-04 |
| P02761 | FMQHIDVLENSLGFK | 0,Acetyl[ProteinN-term]; | 0.0 | 76.0 | #DIV/0! | 1.96E-04 |
| P02761 | RVFMQHIDVLENSLGFK | 0,Pyridylacetyl[AnyN-term](Phenylisocyanate[AnyN-term]); | 0.0 | 11.7 | #DIV/0! | 1.97E-04 |
| P02781 | EEELKRELEEFDAPPEAVEANLK | 0,Iodoacetanilide[AnyN-term]; | 0.0 | 9.8 | #DIV/0! | 1.98E-04 |
| P14740 | YYQLGCR | 6,Carbamidomethyl[C]; | 0.0 | 9.2 | #DIV/0! | 1.99E-04 |
| Q9R0T4 | EPDTFMEQK | 6,Oxidation[M]; | 0.0 | 15.0 | #DIV/0! | 2.00E-04 |
| P02761 | LNGDWFSIVVASNKR | 2,Asn->Met[N]; | 0.0 | 65.5 | #DIV/0! | 2.07E-04 |
| P02770 | ADKDNCFATEGPNLVAR | 6,Dimethylphosphothione[C]; | 0.0 | 15.5 | #DIV/0! | 2.16E-04 |
| P29598 | TIQTICLPPR | 6,Carbamidomethyl[C]; | 0.0 | 143.5 | #DIV/0! | 2.18E-04 |
| Q62867 | EKLKKFFNILTVNTDGK | 6,Trioxidation[F]; | 0.0 | 17.0 | #DIV/0! | 2.29E-04 |
| P00787 | MCEAGYSTSYKEDK | 2,Carbamidomethyl[C]; | 0.0 | 13.2 | #DIV/0! | 2.30E-04 |
| P07522 | YVCNCVIGYIGER | 3,Carbamidomethyl[C];5,Carbamidomethyl[C]; | 0.0 | 26.8 | #DIV/0! | 2.31E-04 |
| P02761 | VFMQHIDVLENSLGFKFR | 3,Oxidation[M]; | 0.0 | 30.0 | #DIV/0! | 2.34E-04 |
| P04276 | YCSSQIDAEMR | 2,Carbamidomethyl[C]; | 0.0 | 13.3 | #DIV/0! | 2.39E-04 |
| P00758 | ATDNYQVWLGR | 0,CLIP_TRAQ_2[AnyN-term]; | 0.0 | 17.3 | #DIV/0! | 2.48E-04 |
| P02770 | LQACCDKPVLQK | 4,Carbamidomethyl[C];5,Carbamidomethyl[C]; | 0.0 | 32.8 | #DIV/0! | 2.56E-04 |
| P02761 | DLSSDIKEKFAK | 0,Carbamyl[AnyN-term]; | 0.0 | 13.8 | #DIV/0! | 2.64E-04 |
| P02761 | MQHIDVLENSLGFK | 5,Cation_Ca[II][D]; | 0.0 | 16.7 | #DIV/0! | 2.73E-04 |
| P02761 | TKDLSSDIK | 0,Carbamyl[AnyN-term]; | 0.0 | 21.0 | #DIV/0! | 2.85E-04 |
| P08426 | IIRHPSYNANTFDNDIMLIK | 3,Arg->Ser[R]; | 0.0 | 27.0 | #DIV/0! | 2.88E-04 |
| P02761 | RVFMQHIDVLENSLGFK | 2,Val->Cys[V]; | 0.0 | 5.5 | #DIV/0! | 3.01E-04 |
| P20059 | FNPVTGEVPPR | 2,Bacillosamine[N]; | 0.0 | 17.5 | #DIV/0! | 3.07E-04 |
| P13635 | EMGPTYADPVCLSK | 11,Carbamidomethyl[C]; | 0.0 | 6.2 | #DIV/0! | 3.20E-04 |
| Q9R0T4 | HKNMFTVNRDTGVISVVTSGLDR | 11,PhosphoCytidine[T]; | 0.0 | 21.3 | #DIV/0! | 3.27E-04 |
| P02761 | GNLDVAKLNGDWFSIVVASNKR | 9,Deamidated[N]; | 0.0 | 21.0 | #DIV/0! | 3.28E-04 |
| P02761 | GSMRVFMQHIDVLENSLGFK | 0,3sulfo[AnyN-term]; | 0.0 | 5.3 | #DIV/0! | 3.36E-04 |
| P02761 | VFMQHIDVLENSLGFK | 5,Carboxyethyl[H](Ethoxyformyl[H]); | 0.0 | 37.5 | #DIV/0! | 3.52E-04 |
| P02761 | VFMQHIDVLENSLGFK | 0,C+12[AnyN-term]; | 0.0 | 13.2 | #DIV/0! | 3.57E-04 |
| P02761 | MQHIDVLENSLGFK | 0,Triton[AnyN-term]; | 0.0 | 8.7 | #DIV/0! | 3.75E-04 |
| P20059 | ECHRGECQSEGVLFFQGNR | 2,MDCC[C]; | 0.0 | 15.7 | #DIV/0! | 3.83E-04 |
| P02761 | VFMQHIDVLENSLGFK | 4,Xlink_DFDNB[Q]; | 0.0 | 52.7 | #DIV/0! | 3.90E-04 |
| Q64230 | QQCLQFFYK | 3,Carbamidomethyl[C]; | 0.0 | 25.0 | #DIV/0! | 4.00E-04 |
| P02761 | FAKLCEAHGITR | 0,ISD_z+2_ion[AnyN-term]; | 0.0 | 7.7 | #DIV/0! | 4.05E-04 |
| P00762 | LGEHNINVLEGDEQFINAAK | 7,Ammonia-loss[N]; | 0.0 | 17.0 | #DIV/0! | 4.16E-04 |
| P36953 | IANDAIQDMLCDMK | 11,Carbamidomethyl[C]; | 0.0 | 3.5 | #DIV/0! | 4.45E-04 |
| P02761 | NGSMRVFMQHIDVLENSLGFK | 4,Oxidation[M];8,Oxidation[M]; | 0.0 | 16.7 | #DIV/0! | 4.51E-04 |
| P02781 | ELEEFDAPPEAVEANLK | 6,Cation_Ca[II][D]; | 0.0 | 10.2 | #DIV/0! | 4.55E-04 |
| P08932 | KTEEDLCVGCFQPIPMDSSDLKPVLK | 7,Carbamidomethyl[C];10,Carbamidomethyl[C]; | 0.0 | 10.0 | #DIV/0! | 4.72E-04 |
| P01048 | KTEEDLCVGCFQPIPMDSSDLKPVLK | 7,Carbamidomethyl[C];10,Carbamidomethyl[C]; | 0.0 | 10.0 | #DIV/0! | 4.72E-04 |
| P02761 | LNGDWFSIVVASNK | 4,Cation_Na[D]; | 0.0 | 431.2 | #DIV/0! | 4.74E-04 |
| P02761 | MQHIDVLENSLGFK | 0,mTRAQ[AnyN-term]; | 0.0 | 7.7 | #DIV/0! | 4.84E-04 |
| P32038 | NMMCAESNR | 4,Carbamidomethyl[C]; | 0.0 | 9.0 | #DIV/0! | 4.94E-04 |
| P02770 | LQACCDKPVLQK | 4,maleimide[C]; | 0.0 | 4.5 | #DIV/0! | 4.94E-04 |
| P20059 | ECQSEGVLFFQGNRK | 2,Diethylphosphate[C]; | 0.0 | 8.7 | #DIV/0! | 4.97E-04 |
| P00689 | MCGSGNSAGTHSTCGSYFNPNNR | 0,Iodoacetanilide[AnyN-term]; | 0.0 | 14.7 | #DIV/0! | 4.99E-04 |
| P02761 | NIIDLTKTDR | 0,NIC[AnyN-term](ICPL[AnyN-term]); | 0.0 | 27.0 | #DIV/0! | 5.14E-04 |
| P05545 | AVLDVDETGTEGAAATAVTAALK | 6,Cation_Na[D]; | 0.0 | 11.3 | #DIV/0! | 5.15E-04 |
| P02761 | NGDWFSIVVASNKR | 0,Galactosyl[AnyN-term]; | 0.0 | 12.5 | #DIV/0! | 5.29E-04 |
| P02761 | PEDGEYFVEYDGGNTFTILK | 17,PhosphoHexNAc[T]; | 0.0 | 7.0 | #DIV/0! | 5.46E-04 |
| Q9JHB9 | SGSGCSILDEVIR | 5,Carboxymethyl[C]; | 0.0 | 13.5 | #DIV/0! | 5.50E-04 |
| P02780 | SGSGCSILDEVIR | 5,Carboxymethyl[C]; | 0.0 | 13.5 | #DIV/0! | 5.50E-04 |
| P81828 | SLLMSSLLALQCFRCTSFDSTGFCHVGR | 4,Oxidation[M];15,Carbamidomethyl[C];24,Carbamidomethyl[C]; | 0.0 | 8.2 | #DIV/0! | 5.53E-04 |
| P08932 | TELTADCETK | 7,Carbamidomethyl[C]; | 0.0 | 8.2 | #DIV/0! | 5.53E-04 |
| P01048 | TELTADCETK | 7,Carbamidomethyl[C]; | 0.0 | 8.2 | #DIV/0! | 5.53E-04 |
| Q64240 | AFAELWAFDAAQGK | 1,Carbonyl[A]; | 0.0 | 8.7 | #DIV/0! | 5.65E-04 |
| P02761 | NGDWFSIVVASNKR | 0,C+12[AnyN-term]; | 0.0 | 20.7 | #DIV/0! | 5.72E-04 |
| P02761 | LINFKNGETFQLMVLYGR | 0,Diisopropylphosphate[AnyN-term]; | 0.0 | 36.8 | #DIV/0! | 6.02E-04 |
| P02761 | VEYDGGNTFTILK | 0,DiART6plex118[AnyN-term]; | 0.0 | 14.3 | #DIV/0! | 6.08E-04 |
| P02761 | NGDWFSIVVASNK | 0,NIC[AnyN-term](ICPL[AnyN-term]); | 0.0 | 4.7 | #DIV/0! | 6.29E-04 |
| Q64230 | TKIPEFNAIIGQR | 1,PhosphoHex(2)[T]; | 0.0 | 7.5 | #DIV/0! | 6.41E-04 |
| P08937 | LDSECQTHTVVGQK | 5,Carbamidomethyl[C]; | 0.0 | 7.5 | #DIV/0! | 6.41E-04 |
| P15684 | QVTPLFAYFK | 3,Dehydrated[T]; | 0.0 | 6.7 | #DIV/0! | 6.42E-04 |
| P02761 | VLENSLGFK | 0,ISD_z+2_ion[AnyN-term]; | 0.0 | 16.8 | #DIV/0! | 7.05E-04 |
| Q68FP1 | TPSAAYLWVGTGASDAEK | 1,Didehydro[T]; | 0.0 | 9.2 | #DIV/0! | 7.38E-04 |
| P15684 | KVVATTQMQAADAR | 8,Oxidation[M]; | 0.0 | 2.5 | #DIV/0! | 7.46E-04 |
| P81828 | ALQCFRCTSFDSTGFCHVGR | 4,Carbamidomethyl[C];7,Carbamidomethyl[C];16,Carbamidomethyl[C]; | 0.0 | 44.3 | #DIV/0! | 7.57E-04 |
| P22282 | TPGETMYYISLPGSVR | 6,Met->Gln[M]; | 0.0 | 9.8 | #DIV/0! | 7.68E-04 |
| P01015 | RFVQAVTGWK | 0,Cytopiloyne+water[AnyN-term]; | 0.0 | 5.5 | #DIV/0! | 8.04E-04 |
| P36373 | HLSQPADITDGVK | 0,C+12[AnyN-term]; | 0.0 | 8.2 | #DIV/0! | 8.17E-04 |
| P00758 | HLSQPADITDGVK | 0,C+12[AnyN-term]; | 0.0 | 8.2 | #DIV/0! | 8.17E-04 |
| P00758 | VIDLPIEEPK | 1,Val->Ser[V]; | 0.0 | 24.5 | #DIV/0! | 8.27E-04 |
| P27590 | WRSTDYGAGYSCDSDMHGWYR | 4,Dehydrated[T]; | 0.0 | 12.5 | #DIV/0! | 8.70E-04 |
| P27274 | CCQADLCNK | 1,Carbamidomethyl[C];2,Carbamidomethyl[C];7,Carbamidomethyl[C]; | 0.0 | 4.7 | #DIV/0! | 9.17E-04 |
| P02761 | LNGDWFSIVVASNKR | 14,Carbamyl[K]; | 0.0 | 5.2 | #DIV/0! | 9.81E-04 |
| P02761 | FMQHIDVLENSLGFK | 3,Xlink_DFDNB[Q]; | 0.0 | 12.8 | #DIV/0! | 1.00E-03 |
| P02761 | VFMQHIDVLENSLGFK | 0,Cytopiloyne+water[AnyN-term]; | 0.0 | 14.8 | #DIV/0! | 1.05E-03 |
| P20760 | SVSELPIVHR | 2,Val->Cys[V]; | 0.0 | 3.3 | #DIV/0! | 1.09E-03 |
| Q64230 | CQLAKIQTFQGDSDHNWK | 2,Gln->Cys[Q]; | 0.0 | 1.5 | #DIV/0! | 1.11E-03 |
| P12346 | FDEFFSQGCAPGYK | 2,Cation_Na[D];9,Carbamidomethyl[C]; | 0.0 | 15.3 | #DIV/0! | 1.16E-03 |
| P02761 | TKDLSSDIK | 0,Guanidinyl[AnyN-term]; | 0.0 | 3.2 | #DIV/0! | 1.17E-03 |
| P02761 | LCEAHGITR | 2,2-succinyl[C]; | 0.0 | 26.0 | #DIV/0! | 1.18E-03 |
| P02761 | FMQHIDVLENSLGFK | 0,Diisopropylphosphate[AnyN-term]; | 0.0 | 50.0 | #DIV/0! | 1.21E-03 |
| P83121 | EICAWVVVTTR | 3,Trioxidation[C]; | 0.0 | 5.2 | #DIV/0! | 1.27E-03 |
| P81828 | EICAWVVVTTR | 3,Trioxidation[C]; | 0.0 | 5.2 | #DIV/0! | 1.27E-03 |
| P81827 | EICAWVVVTTR | 3,Trioxidation[C]; | 0.0 | 5.2 | #DIV/0! | 1.27E-03 |
| P02770 | KQTALAELVK | 0,Carbamyl[AnyN-term]; | 0.0 | 4.3 | #DIV/0! | 1.29E-03 |
| P02761 | IEENGSMRVFMQHIDVLENSLGFK | 0,Carboxymethyl[AnyN-term]; | 0.0 | 8.3 | #DIV/0! | 1.29E-03 |
| P07522 | VNLDPASVPPR | 1,Val->Thr[V]; | 0.0 | 11.0 | #DIV/0! | 1.29E-03 |
| Q64240 | GECVPGDREVESTSFAR | 3,Carbamidomethyl[C]; | 0.0 | 11.2 | #DIV/0! | 1.43E-03 |
| P12346 | KTSYQDCIK | 7,Carbamidomethyl[C]; | 0.0 | 2.7 | #DIV/0! | 1.46E-03 |
| P02770 | LQACCDKPVLQK | 5,maleimide[C]; | 0.0 | 4.0 | #DIV/0! | 1.46E-03 |
| P02761 | VAKLNGDWFSIVVASNKR | 3,CresylSaligeninPhosphate[K]; | 0.0 | 8.8 | #DIV/0! | 1.47E-03 |
| P02782 | ELEMYNAPPAAVEAK | 6,Deamidated[N]; | 0.0 | 12.3 | #DIV/0! | 1.49E-03 |
| P27590 | DWMSIVTPAR | 1,Cation_Na[D]; | 0.0 | 18.7 | #DIV/0! | 1.51E-03 |
| P02782 | QICELVAHETISFLMK | 3,Cys->SecNEM_2H(5)[C]; | 0.0 | 24.3 | #DIV/0! | 1.51E-03 |
| P22283 | LVNCPFEEQTEQLKR | 4,monomethylphosphothione[C]; | 0.0 | 10.8 | #DIV/0! | 1.56E-03 |
| P02761 | HIDVLENSLGFK | 2,Xle->Thr[I]; | 0.0 | 6.0 | #DIV/0! | 1.58E-03 |
| P02761 | EKIEENGSMR | 1,Glu->Met[E]; | 0.0 | 4.8 | #DIV/0! | 1.71E-03 |
| P01835 | WKIDGTERRDGVLDSVTDQDSK | 15,dHex(1)Hex(2)[S]; | 0.0 | 7.0 | #DIV/0! | 1.76E-03 |
| P17475 | YRKISSNLADFAFSLYR | 0,Ethylphosphate[AnyN-term]; | 0.0 | 4.3 | #DIV/0! | 1.76E-03 |
| Q63041 | AEQGAYLGPLPYK | 0,ISD_z+2_ion[AnyN-term]; | 0.0 | 10.3 | #DIV/0! | 1.77E-03 |
| Q9JHB9 | SGSGCSILDEVIR | 5,Carbamidomethyl[C]; | 0.0 | 273.3 | #DIV/0! | 1.81E-03 |
| P02780 | SGSGCSILDEVIR | 5,Carbamidomethyl[C]; | 0.0 | 273.3 | #DIV/0! | 1.81E-03 |
| P01836 | DGVLDSVTDQDSKDSTYSMSSTLSLTK | 0,Carbamyl[AnyN-term]; | 0.0 | 18.7 | #DIV/0! | 1.89E-03 |
| P01835 | DGVLDSVTDQDSKDSTYSMSSTLSLTK | 0,Carbamyl[AnyN-term]; | 0.0 | 18.7 | #DIV/0! | 1.89E-03 |
| P02761 | LCEAHGITR | 0,ICPL_13C(6)[AnyN-term]; | 0.0 | 13.7 | #DIV/0! | 1.89E-03 |
| P02761 | MQHIDVLENSLGFK | 0,TMAB_2H(9)[AnyN-term]; | 0.0 | 32.7 | #DIV/0! | 1.89E-03 |
| P05545 | ALYQAEAFVADFK | 2,Carbonyl[L]; | 0.0 | 8.0 | #DIV/0! | 1.90E-03 |
| P85971 | DLPAATAPAGPASFAR | 2,Xle->Cys[L]; | 0.0 | 5.7 | #DIV/0! | 1.94E-03 |
| P06760 | ESGPTLDMPVPSSFNDITQEAELR | 7,Ammonium[D]; | 0.0 | 4.5 | #DIV/0! | 2.00E-03 |
| P02761 | FAKLCEAHGITR | 5,2-succinyl[C]; | 0.0 | 9.3 | #DIV/0! | 2.01E-03 |
| P02770 | FKDLGEQHFK | 3,Cation_Na[D]; | 0.0 | 9.7 | #DIV/0! | 2.03E-03 |
| P36373 | PGDDHSNDLMLLHLSQPADITDGVK | 15,pyrophospho[S]; | 0.0 | 7.5 | #DIV/0! | 2.08E-03 |
| P02761 | RVFMQHIDVLENSLGFK | 12,Deamidated[N]; | 0.0 | 8.2 | #DIV/0! | 2.10E-03 |
| P02761 | DLSSDIKEKFAK | 4,Carbamyl[S]; | 0.0 | 6.0 | #DIV/0! | 2.13E-03 |
| P21704 | TNQMAEAISDHYPVEVTLR | 0,PyMIC[AnyN-term]; | 0.0 | 4.8 | #DIV/0! | 2.15E-03 |
| P32038 | TLCDVAGWGVVTHAGR | 3,Cys->Glu[C]; | 0.0 | 11.3 | #DIV/0! | 2.18E-03 |
| P06866 | GAVSPVGVQPILNK | 3,Val->Asn[V]; | 0.0 | 11.7 | #DIV/0! | 2.22E-03 |
| P04937 | QDGHLWCSTTSNYEQDQK | 7,Carbamidomethyl[C]; | 0.0 | 5.7 | #DIV/0! | 2.27E-03 |
| Q9R0T4 | HKNMFTVNRDTGVISVVTSGLDR | 6,PhosphoCytidine[T]; | 0.0 | 13.8 | #DIV/0! | 2.27E-03 |
| P15684 | SQTAIKLPASVSTIMDR | 6,Lys->Cys[K]; | 0.0 | 7.5 | #DIV/0! | 2.28E-03 |
| P02782 | CVDQMSNGDR | 0,Pyro-carbamidomethyl[AnyN-termC]; | 0.0 | 12.5 | #DIV/0! | 2.29E-03 |
| P15978 | PGLGEPRFISVGYVDDTQFVR | 0,Succinyl_2H(4)[AnyN-term]; | 0.0 | 10.5 | #DIV/0! | 2.30E-03 |
| P22282 | TPGETMYYISLPGSVR | 5,Dehydrated[T]; | 0.0 | 12.0 | #DIV/0! | 2.36E-03 |
| P02781 | ELEEFDAPPEAVEANLK | 5,Nitro[F]; | 0.0 | 9.0 | #DIV/0! | 2.40E-03 |
| P01836 | TYSMSSTLSLTK | 2,AEBS[Y]; | 0.0 | 12.8 | #DIV/0! | 2.41E-03 |
| P01835 | TYSMSSTLSLTK | 2,AEBS[Y]; | 0.0 | 12.8 | #DIV/0! | 2.41E-03 |
| P17475 | NNYHSEAFSVNFADSEEAKK | 14,Cation_Na[D]; | 0.0 | 8.3 | #DIV/0! | 2.42E-03 |
| P22283 | SKLVNCPFEEQTEQLKR | 6,Cys->SecNEM_2H(5)[C]; | 0.0 | 10.3 | #DIV/0! | 2.46E-03 |
| P02761 | TPEDGEYFVEYDGGNTFTILK | 0,Succinyl[AnyN-term]; | 0.0 | 7.2 | #DIV/0! | 2.47E-03 |
| P02761 | NGETFQLMVLYGR | 0,C+12[AnyN-term]; | 0.0 | 63.8 | #DIV/0! | 2.48E-03 |
| P07647 | NHTRQRAYDHNNDLMLLHLSKPADITGGVK | 3,Phosphopropargyl[T]; | 0.0 | 7.8 | #DIV/0! | 2.52E-03 |
| P02770 | QRFPNAEFAEITK | 0,Pyridylacetyl[AnyN-term](Phenylisocyanate[AnyN-term]); | 0.0 | 17.2 | #DIV/0! | 2.53E-03 |
| P05545 | RALYQAEAFVADFK | 1,Arg->Lys[R]; | 0.0 | 4.5 | #DIV/0! | 2.54E-03 |
| P02761 | DVAKLNGDWFSIVVASNKR | 0,Diisopropylphosphate[AnyN-term]; | 0.0 | 9.0 | #DIV/0! | 2.54E-03 |
| P02761 | FMQHIDVLENSLGFK | 0,PEITC[AnyN-term]; | 0.0 | 35.7 | #DIV/0! | 2.56E-03 |
| P02761 | NGETFQLMVLYGR | 7,Xle->Ser[L]; | 0.0 | 17.7 | #DIV/0! | 2.56E-03 |
| P15083 | CPVLVGTQALVQEGYEGR | 1,Propionamide_2H(3)[C]; | 0.0 | 6.2 | #DIV/0! | 2.58E-03 |
| P07522 | VNLDPASVPPR | 1,Val->Cys[V]; | 0.0 | 11.5 | #DIV/0! | 2.64E-03 |
| P83121 | ICQTYPDEICAWVVVTTR | 10,maleimide[C]; | 0.0 | 21.2 | #DIV/0! | 2.64E-03 |
| P81827 | ICQTYPDEICAWVVVTTR | 10,maleimide[C]; | 0.0 | 21.2 | #DIV/0! | 2.64E-03 |
| P15978 | PGLGEPRFISVGYVDDTQFVR | 5,Cation_Ag[E]; | 0.0 | 3.7 | #DIV/0! | 2.72E-03 |
| P12346 | HLAQAPNHVVVSR | 0,Propionyl_13C(3)[AnyN-term]; | 0.0 | 4.0 | #DIV/0! | 2.76E-03 |
| P02761 | MQHIDVLENSLGFK | 0,LG-pyrrole[AnyN-term]; | 0.0 | 8.0 | #DIV/0! | 2.76E-03 |
| P02770 | HPDYSVSLLLR | 0,Iodoacetanilide_13C(6)[AnyN-term]; | 0.0 | 8.5 | #DIV/0! | 2.87E-03 |
| P01836 | QDSKDSTYSMSSTLSLTK | 1,Gln->Val[Q]; | 0.0 | 3.3 | #DIV/0! | 2.89E-03 |
| P01835 | QDSKDSTYSMSSTLSLTK | 1,Gln->Val[Q]; | 0.0 | 3.3 | #DIV/0! | 2.89E-03 |
| P05545 | EAKKFINDYVSNQTQGKIAELFSELDER | 5,Trioxidation[F]; | 0.0 | 15.5 | #DIV/0! | 2.90E-03 |
| P98158 | GGQCIPGQWHCDR | 4,Carbamidomethyl[C];11,Carbamidomethyl[C]; | 0.0 | 9.3 | #DIV/0! | 2.92E-03 |
| P02761 | TPEDGEYFVEYDGGNTFTILK | 8,Phe->CamCys[F]; | 0.0 | 24.5 | #DIV/0! | 2.94E-03 |
| P02761 | HIDVLENSLGFK | 12,Lys->Asn[K]; | 0.0 | 2.7 | #DIV/0! | 2.96E-03 |
| P02770 | AADKDNCFATEGPNLVAR | 3,Cation_Na[D];5,Cation_Na[D];7,Carbamidomethyl[C]; | 0.0 | 15.8 | #DIV/0! | 2.97E-03 |
| P07314 | FQPSIQLAR | 0,Ethylphosphate[AnyN-term]; | 0.0 | 11.2 | #DIV/0! | 3.07E-03 |
| P01836 | DSKDSTYSMSSTLSLTK | 3,Acetyldeoxyhypusine[K]; | 0.0 | 14.5 | #DIV/0! | 3.10E-03 |
| P01835 | DSKDSTYSMSSTLSLTK | 3,Acetyldeoxyhypusine[K]; | 0.0 | 14.5 | #DIV/0! | 3.10E-03 |
| P14841 | LLGAPQEADASEEGVQR | 9,Cation_Na[D]; | 0.0 | 7.5 | #DIV/0! | 3.13E-03 |
| P02780 | LVKPYVQDHFTEK | 1,Carbonyl[L]; | 0.0 | 9.0 | #DIV/0! | 3.15E-03 |
| P02761 | LNGDWFSIVVASNKR | 5,Delta_H(-4)O(3)[W]; | 0.0 | 19.5 | #DIV/0! | 3.18E-03 |
| P02761 | FMQHIDVLENSLGFK | 1,Phe->His[F]; | 0.0 | 5.2 | #DIV/0! | 3.27E-03 |
| P04937 | KYSFCTDHAVLVQTR | 2,Tyr->Xle[Y]; | 0.0 | 3.2 | #DIV/0! | 3.27E-03 |
| P02761 | HIDVLENSLGFK | 2,Xle->Tyr[I]; | 0.0 | 3.2 | #DIV/0! | 3.27E-03 |
| P12346 | TTYEEYLGAEYLQAVGNIR | 0,Biotin_Thermo-21328[AnyN-term]; | 0.0 | 11.2 | #DIV/0! | 3.28E-03 |
| P47967 | VAVDGQHICEYSHR | 9,Carbamidomethyl[C]; | 0.0 | 9.7 | #DIV/0! | 3.30E-03 |
| Q9JHB9 | GSGCSILDEVIR | 0,Propyl_2H(6)[AnyN-term]; | 0.0 | 12.3 | #DIV/0! | 3.32E-03 |
| P02780 | GSGCSILDEVIR | 0,Propyl_2H(6)[AnyN-term]; | 0.0 | 12.3 | #DIV/0! | 3.32E-03 |
| P15684 | ALGDTPAPNIDTTELVER | 4,Cation_Ca[II][D]; | 0.0 | 14.2 | #DIV/0! | 3.34E-03 |
| P27590 | CNVYETLTEYWR | 1,Cys->Val[C]; | 0.0 | 7.0 | #DIV/0! | 3.42E-03 |
| Q9JHB9 | SGSGCSILDEVIR | 0,GIST-Quat[AnyN-term]; | 0.0 | 10.2 | #DIV/0! | 3.48E-03 |
| P02780 | SGSGCSILDEVIR | 0,GIST-Quat[AnyN-term]; | 0.0 | 10.2 | #DIV/0! | 3.48E-03 |
| P83121 | ICQTYPDEICAWVVVTTR | 2,maleimide[C]; | 0.0 | 36.5 | #DIV/0! | 3.57E-03 |
| P81827 | ICQTYPDEICAWVVVTTR | 2,maleimide[C]; | 0.0 | 36.5 | #DIV/0! | 3.57E-03 |
| P02761 | FMQHIDVLENSLGFK | 1,AzidoF[F]; | 0.0 | 9.7 | #DIV/0! | 3.59E-03 |
| P00758 | NNLYEDEPFAQHR | 2,Deamidated[N]; | 0.0 | 6.3 | #DIV/0! | 3.61E-03 |
| P02761 | LNGDWFSIVVASNK | 4,Cation_Fe[II][D]; | 0.0 | 10.8 | #DIV/0! | 3.62E-03 |
| Q6IE52 | ESVVFVQTDKPMYKPGQSVK | 3,Val->Ser[V]; | 0.0 | 3.7 | #DIV/0! | 3.68E-03 |
| P27590 | INFECSYPLDMK | 5,Ethanolamine[C]; | 0.0 | 7.3 | #DIV/0! | 3.68E-03 |
| P14046 | ESVVFVQTDKPMYKPGQSVK | 3,Val->Ser[V]; | 0.0 | 3.7 | #DIV/0! | 3.68E-03 |
| P02761 | KIEENGSMR | 8,Oxidation[M]; | 0.0 | 8.3 | #DIV/0! | 3.69E-03 |
| P24268 | LLDIACWVHHK | 6,Carbamidomethyl[C]; | 0.0 | 14.0 | #DIV/0! | 3.70E-03 |
| P02782 | CVDQMSNGDR | 1,Carbamidomethyl[C];5,Oxidation[M];7,Deamidated[N]; | 0.0 | 7.7 | #DIV/0! | 3.72E-03 |
| P02761 | LNGDWFSIVVASNK | 2,Deamidated[N];4,Cation_Na[D]; | 0.0 | 59.8 | #DIV/0! | 3.73E-03 |
| P02761 | LNGDWFSIVVASNKR | 1,Xle->Glu[L]; | 0.0 | 6.2 | #DIV/0! | 3.95E-03 |
| P02761 | DNIIDLTKTDR | 1,Cation_Na[D];2,Deamidated[N]; | 0.0 | 19.0 | #DIV/0! | 3.96E-03 |
| Q63041 | LQDQSNIQR | 1,Carbonyl[L]; | 0.0 | 0.8 | #DIV/0! | 4.10E-03 |
| P02761 | DYDRYVMFHLINFK | 14,Xlink_EGS[115][K]; | 0.0 | 6.7 | #DIV/0! | 4.10E-03 |
| P00884 | LDQGGAPLAGTNK | 1,Xle->Thr[L]; | 0.0 | 0.8 | #DIV/0! | 4.10E-03 |
| P02761 | LCEAHGITR | 2,Trioxidation[C]; | 0.0 | 204.5 | #DIV/0! | 4.15E-03 |
| P83121 | CISFDSTGFCYVGR | 5,Cation_Na[D];10,Carbamidomethyl[C]; | 0.0 | 22.3 | #DIV/0! | 4.18E-03 |
| P07151 | IPNIEMSDLSFSK | 8,Cation_Na[D]; | 0.0 | 10.5 | #DIV/0! | 4.18E-03 |
| P81827 | LQCFRCESLDSTGLCR | 3,Carbamidomethyl[C];15,Carbamidomethyl[C]; | 0.0 | 8.7 | #DIV/0! | 4.22E-03 |
| P14740 | SALDKMLQDVQMPSK | 1,AEC-MAEC_2H(4)[S]; | 0.0 | 7.2 | #DIV/0! | 4.28E-03 |
| P02761 | FAKLCEAHGITR | 5,Trioxidation[C]; | 0.0 | 19.5 | #DIV/0! | 4.28E-03 |
| P02770 | SFLGHYLHEVAR | 1,Phosphopantetheine[S]; | 0.0 | 6.2 | #DIV/0! | 4.34E-03 |
| P02782 | QICELVAHETISFLMK | 0,GIST-Quat_2H(3)[AnyN-term]; | 0.0 | 8.5 | #DIV/0! | 4.39E-03 |
| P01836 | DSTYSMSSTLSLTK | 4,Dehydrated[Y]; | 0.0 | 22.2 | #DIV/0! | 4.40E-03 |
| P01835 | DSTYSMSSTLSLTK | 4,Dehydrated[Y]; | 0.0 | 22.2 | #DIV/0! | 4.40E-03 |
| P02761 | REKIEENGSMRVFMQHIDVLENSLGFK | 7,Asn->Met[N]; | 0.0 | 5.5 | #DIV/0! | 4.40E-03 |
| P02761 | LNGDWFSIVVASNKR | 1,Carbonyl[L]; | 0.0 | 17.3 | #DIV/0! | 4.42E-03 |
| P12346 | CLRDGGGDVAFVK | 4,Cation_Ag[D]; | 0.0 | 4.0 | #DIV/0! | 4.48E-03 |
| P02761 | NLDVAKLNGDWFSIVVASNKR | 0,dNIC[AnyN-term]; | 0.0 | 8.3 | #DIV/0! | 4.54E-03 |
| P22282 | TPGETMYYISLPGSVR | 5,Thr->Pro[T]; | 0.0 | 16.8 | #DIV/0! | 4.60E-03 |
| P02761 | QHIDVLENSLGFK | 0,TMAB_2H(9)[AnyN-term]; | 0.0 | 9.7 | #DIV/0! | 4.69E-03 |
| P12346 | PEGTTYEEYLGAEYLQAVGNIR | 0,NIC[AnyN-term](ICPL[AnyN-term]); | 0.0 | 6.0 | #DIV/0! | 4.69E-03 |
| P07647 | SHDLQCVNIHLLSNEK | 6,Bromobimane[C]; | 0.0 | 9.2 | #DIV/0! | 5.02E-03 |
| P00759 | SHDLQCVNIHLLSNEK | 6,Bromobimane[C]; | 0.0 | 9.2 | #DIV/0! | 5.02E-03 |
| O55006 | IDTSCVSGSPPLR | 5,Carbamidomethyl[C]; | 0.0 | 14.2 | #DIV/0! | 5.10E-03 |
| P83121 | CQTYPDEICAWVVVTTR | 9,Carbamidomethyl[C]; | 0.0 | 52.3 | #DIV/0! | 5.19E-03 |
| P81828 | CQTYPDEICAWVVVTTR | 9,Carbamidomethyl[C]; | 0.0 | 52.3 | #DIV/0! | 5.19E-03 |
| P81827 | CQTYPDEICAWVVVTTR | 9,Carbamidomethyl[C]; | 0.0 | 52.3 | #DIV/0! | 5.19E-03 |
| P20767 | TVFPPSTEELQGNK | 1,HexNAc(1)NeuAc(1)[T]; | 0.0 | 7.0 | #DIV/0! | 5.25E-03 |
| P09005 | QGKMQQVESSLQPETLKK | 3,CresylSaligeninPhosphate[K]; | 0.0 | 2.3 | #DIV/0! | 5.25E-03 |
| P05545 | QGKMQQVESSLQPETLKK | 3,CresylSaligeninPhosphate[K]; | 0.0 | 2.3 | #DIV/0! | 5.25E-03 |
| P05544 | QGKMQQVESSLQPETLKK | 3,CresylSaligeninPhosphate[K]; | 0.0 | 2.3 | #DIV/0! | 5.25E-03 |
| P02770 | RMSQRFPNAEFAEITK | 4,Gln->Pro[Q]; | 0.0 | 2.8 | #DIV/0! | 5.27E-03 |
| P83121 | HICQTYPDEICAWVVVTTR | 8,Cation_Na[D]; | 0.0 | 17.7 | #DIV/0! | 5.27E-03 |
| P02761 | FAKLCEAHGITR | 3,Acetyldeoxyhypusine[K]; | 0.0 | 17.0 | #DIV/0! | 5.32E-03 |
| P07522 | DGCHWVR | 3,Carbamidomethyl[C]; | 0.0 | 7.5 | #DIV/0! | 5.50E-03 |
| P02761 | IDVLENSLGFK | 0,Carboxymethyl[AnyN-term]; | 0.0 | 7.7 | #DIV/0! | 5.55E-03 |
| P02761 | LNGDWFSIVVASNKR | 0,C+12[AnyN-term]; | 0.0 | 15.2 | #DIV/0! | 5.57E-03 |
| P02761 | LNGDWFSIVVASNK | 2,Asn->Thr[N]; | 0.0 | 13.7 | #DIV/0! | 5.59E-03 |
| P22283 | LVNCPFEEQTEQLKR | 4,2-succinyl[C]; | 0.0 | 14.2 | #DIV/0! | 5.60E-03 |
| P05545 | QPEDQAEINTGSALFIDK | 0,NDA[AnyN-term]; | 0.0 | 11.8 | #DIV/0! | 5.62E-03 |
| P02770 | QEPERNECFLQHKDDNPNLPPFQRPEAEAMCTSFQENPTSFLGHYLHEVAR | 8,GG[C](Dicarbamidomethyl[C]); | 0.0 | 7.3 | #DIV/0! | 5.70E-03 |
| P02761 | TDYDRYVMFHLINFK | 11,Dioxidation[L]; | 0.0 | 7.3 | #DIV/0! | 5.70E-03 |
| P98158 | CDNGYCIPASWR | 1,Carbamidomethyl[C];6,Carbamidomethyl[C]; | 0.0 | 9.2 | #DIV/0! | 5.81E-03 |
| P22283 | EICYFQLYPDYIEQNIR | 3,Trioxidation[C]; | 0.0 | 9.3 | #DIV/0! | 5.82E-03 |
| P07171 | LQNLIQELLQAR | 0,CAF[AnyN-term]; | 0.0 | 3.5 | #DIV/0! | 5.93E-03 |
| P00787 | HEAGDVMGGHAIR | 2,Carbonyl[E]; | 0.0 | 3.5 | #DIV/0! | 5.93E-03 |
| P02761 | FMQHIDVLENSLGFK | 2,Oxidation[M];6,Cation_Na[D]; | 0.0 | 14.3 | #DIV/0! | 6.01E-03 |
| P05544 | LGAKDSTMEEILEGLK | 8,Oxidation[M]; | 0.0 | 1.8 | #DIV/0! | 6.02E-03 |
| Q64230 | AIMTILDQEPDAR | 0,Succinyl[AnyN-term]; | 0.0 | 1.8 | #DIV/0! | 6.02E-03 |
| P02761 | DWFSIVVASNKR | 0,SPITC[AnyN-term]; | 0.0 | 35.0 | #DIV/0! | 6.05E-03 |
| P05545 | KTRALYQAEAFVADFK | 2,Pent(1)HexNAc(1)[T]; | 0.0 | 22.7 | #DIV/0! | 6.16E-03 |
| P02761 | DLSSDIKEK | 0,Methyl[AnyN-term]; | 0.0 | 8.7 | #DIV/0! | 6.17E-03 |
| P01836 | DGVLDSVTDQDSK | 1,Cation_Na[D];5,Cation_Na[D]; | 0.0 | 2.2 | #DIV/0! | 6.17E-03 |
| P01835 | DGVLDSVTDQDSK | 1,Cation_Na[D];5,Cation_Na[D]; | 0.0 | 2.2 | #DIV/0! | 6.17E-03 |
| P02761 | DVAKLNGDWFSIVVASNKR | 0,Biotin_Thermo-21328[AnyN-term]; | 0.0 | 7.2 | #DIV/0! | 6.18E-03 |
| P13432 | TTEYQYQWQLTAPDPTPLSNPPTQLHSTEQANTK | 14,Ammonium[D]; | 0.0 | 10.2 | #DIV/0! | 6.20E-03 |
| P02761 | SIVVASNKR | 0,CLIP_TRAQ_2[AnyN-term]; | 0.0 | 14.0 | #DIV/0! | 6.38E-03 |
| P15684 | ALACSNEVWILNR | 0,BITC[AnyN-term]; | 0.0 | 11.0 | #DIV/0! | 6.45E-03 |
| P83121 | RHICQTYPDEICAWVVVTTR | 12,Cys->Dha[C]; | 0.0 | 4.8 | #DIV/0! | 6.49E-03 |
| Q5XI43 | HWDLSGGPGSQGR | 0,Triton[AnyN-term]; | 0.0 | 10.7 | #DIV/0! | 6.62E-03 |
| P02761 | DYDRYVMFHLINFK | 2,Methylphosphonate[Y]; | 0.0 | 8.3 | #DIV/0! | 6.72E-03 |
| P47728 | LLPVQENFLLK | 2,Xle->Glu[L]; | 0.0 | 2.5 | #DIV/0! | 6.75E-03 |
| P07171 | LLPVQENFLLK | 2,Xle->Glu[L]; | 0.0 | 2.5 | #DIV/0! | 6.75E-03 |
| P02761 | KTPEDGEYFVEYDGGNTFTILK | 1,Lys->Thr[K]; | 0.0 | 2.5 | #DIV/0! | 6.75E-03 |
| P81828 | LALQCFRCTSFDSTGFCHVGR | 17,Carbamidomethyl[C]; | 0.0 | 27.8 | #DIV/0! | 6.81E-03 |
| Q63556 | VFSQQADLSR | 1,Oxidation[V]; | 0.0 | 5.2 | #DIV/0! | 6.84E-03 |
| P09005 | VFSQQADLSR | 1,Oxidation[V]; | 0.0 | 5.2 | #DIV/0! | 6.84E-03 |
| P05544 | VFSQQADLSR | 1,Oxidation[V]; | 0.0 | 5.2 | #DIV/0! | 6.84E-03 |
| P00787 | NFYNVDISYLK | 1,Asn->Met[N]; | 0.0 | 7.2 | #DIV/0! | 6.89E-03 |
| P02761 | LNGDWFSIVVASNK | 1,Xle->Thr[L]; | 0.0 | 5.7 | #DIV/0! | 6.90E-03 |
| P07647 | LLHLSKPADITGGVK | 0,SPITC_13C(6)[AnyN-term]; | 0.0 | 4.2 | #DIV/0! | 6.98E-03 |
| P15684 | TAIKLPASVSTIMDR | 4,Xlink_DTSSP[192][K]; | 0.0 | 6.2 | #DIV/0! | 7.00E-03 |
| P50123 | FSSADRSSFIDDAFALAR | 0,Methyl_2H(3)13C(1)[AnyN-term]; | 0.0 | 3.8 | #DIV/0! | 7.06E-03 |
| P02782 | AHETISFLMK | 0,3sulfo[AnyN-term]; | 0.0 | 1.5 | #DIV/0! | 7.08E-03 |
| P02761 | DYDRYVMFHLINFK | 5,Sulfo[Y]; | 0.0 | 1.5 | #DIV/0! | 7.08E-03 |
| P83121 | PDEICAWVVVTTR | 5,Cys->Trp[C]; | 0.0 | 62.0 | #DIV/0! | 7.14E-03 |
| P81828 | PDEICAWVVVTTR | 5,Cys->Trp[C]; | 0.0 | 62.0 | #DIV/0! | 7.14E-03 |
| P81827 | PDEICAWVVVTTR | 5,Cys->Trp[C]; | 0.0 | 62.0 | #DIV/0! | 7.14E-03 |
| P27590 | YGAGYSCDSDMHGWYR | 7,iodoTMT[C]; | 0.0 | 5.5 | #DIV/0! | 7.22E-03 |
| P27590 | ERDWMSIVTPAR | 0,Propyl[AnyN-term]; | 0.0 | 9.8 | #DIV/0! | 7.31E-03 |
| P09005 | ESSLQPETLK | 0,Biotin_Thermo-21330[AnyN-term]; | 0.0 | 7.7 | #DIV/0! | 7.38E-03 |
| P05545 | ESSLQPETLK | 0,Biotin_Thermo-21330[AnyN-term]; | 0.0 | 7.7 | #DIV/0! | 7.38E-03 |
| P05544 | ESSLQPETLK | 0,Biotin_Thermo-21330[AnyN-term]; | 0.0 | 7.7 | #DIV/0! | 7.38E-03 |
| P02770 | VFLGTFLYEYSR | 0,ICPL_13C(6)[AnyN-term]; | 0.0 | 7.5 | #DIV/0! | 7.41E-03 |
| P15684 | ATLVNEADKLR | 0,NIC[AnyN-term](ICPL[AnyN-term]); | 0.0 | 2.7 | #DIV/0! | 7.44E-03 |
| P02761 | DGEYFVEYDGGNTFTILK | 0,ESP[AnyN-term]; | 0.0 | 2.7 | #DIV/0! | 7.44E-03 |
| P02761 | LCEAHGITR | 1,Xle->Cys[L]; | 0.0 | 6.5 | #DIV/0! | 7.48E-03 |
| P22057 | GHDTVQPNFQQDK | 2,His->Ser[H]; | 0.0 | 2.8 | #DIV/0! | 7.49E-03 |
| P08723 | EDGEHVTQVSGTR | 0,Propyl_2H(6)[AnyN-term]; | 0.0 | 5.2 | #DIV/0! | 7.54E-03 |
| P02761 | RVFMQHIDVLENSLGFK | 6,Formylasparagine[H]; | 0.0 | 13.5 | #DIV/0! | 7.60E-03 |
| P83121 | ICQTYPDEICAWVVVTTR | 2,NEMsulfur[C]; | 0.0 | 19.8 | #DIV/0! | 7.73E-03 |
| P81827 | ICQTYPDEICAWVVVTTR | 2,NEMsulfur[C]; | 0.0 | 19.8 | #DIV/0! | 7.73E-03 |
| P02761 | DLSSDIKEK | 9,Carbamyl[K]; | 0.0 | 6.3 | #DIV/0! | 7.78E-03 |
| P97546 | NGVELTATR | 1,Deamidated[N]; | 0.0 | 5.5 | #DIV/0! | 7.85E-03 |
| P02770 | YNEVLTQCCTESDK | 8,Carbamidomethyl[C];9,Carbamidomethyl[C];13,Cation_Na[D]; | 0.0 | 19.7 | #DIV/0! | 7.91E-03 |
| P81827 | LQCFRCESLDSTGLCR | 6,Carbamidomethyl[C];15,Carbamidomethyl[C]; | 0.0 | 8.0 | #DIV/0! | 8.05E-03 |
| P02761 | EKFAKLCEAHGITR | 0,Cytopiloyne+water[AnyN-term]; | 0.0 | 8.0 | #DIV/0! | 8.05E-03 |
| Q64319 | DFTTTQVGMHDLVR | 0,NDA[AnyN-term]; | 0.0 | 7.3 | #DIV/0! | 8.09E-03 |
| P22283 | CPFEEQTEQLKR | 1,MDCC[C]; | 0.0 | 7.5 | #DIV/0! | 8.10E-03 |
| P22282 | CPFEEQTEQLKR | 1,MDCC[C]; | 0.0 | 7.5 | #DIV/0! | 8.10E-03 |
| P02761 | EKIEENGSMR | 0,Dimethyl_2H(6)13C(2)[AnyN-term]; | 0.0 | 10.7 | #DIV/0! | 8.12E-03 |
| P07522 | DKHTGKNVVRVNLDPASVPPR | 7,Asn->Phe[N]; | 0.0 | 3.2 | #DIV/0! | 8.27E-03 |
| P00689 | GFRLDAAKHMWPGDIK | 0,Diethylphosphate[AnyN-term]; | 0.0 | 3.2 | #DIV/0! | 8.27E-03 |
| P01836 | TYSMSSTLSLTK | 0,SPITC_13C(6)[AnyN-term]; | 0.0 | 8.0 | #DIV/0! | 8.36E-03 |
| P01835 | TYSMSSTLSLTK | 0,SPITC_13C(6)[AnyN-term]; | 0.0 | 8.0 | #DIV/0! | 8.36E-03 |
| P02761 | HIDVLENSLGFK | 1,Aspartylurea[H]; | 0.0 | 21.5 | #DIV/0! | 8.43E-03 |
| P02761 | LNGDWFSIVVASNK | 4,Cation_Fe[III][D]; | 0.0 | 5.5 | #DIV/0! | 8.51E-03 |
| P01836 | DSTYSMSSTLSLTK | 1,Ammonium[D]; | 0.0 | 9.0 | #DIV/0! | 8.52E-03 |
| P01835 | DSTYSMSSTLSLTK | 1,Ammonium[D]; | 0.0 | 9.0 | #DIV/0! | 8.52E-03 |
| P81828 | QKCQTYPDEICAWVVVTTR | 8,Ammonium[D]; | 0.0 | 2.3 | #DIV/0! | 8.63E-03 |
| P68136 | AGFAGDDAPR | 0,C+12[AnyN-term]; | 0.0 | 2.3 | #DIV/0! | 8.63E-03 |
| P68035 | AGFAGDDAPR | 0,C+12[AnyN-term]; | 0.0 | 2.3 | #DIV/0! | 8.63E-03 |
| P63269 | AGFAGDDAPR | 0,C+12[AnyN-term]; | 0.0 | 2.3 | #DIV/0! | 8.63E-03 |
| P63259 | AGFAGDDAPR | 0,C+12[AnyN-term]; | 0.0 | 2.3 | #DIV/0! | 8.63E-03 |
| P62738 | AGFAGDDAPR | 0,C+12[AnyN-term]; | 0.0 | 2.3 | #DIV/0! | 8.63E-03 |
| P60711 | AGFAGDDAPR | 0,C+12[AnyN-term]; | 0.0 | 2.3 | #DIV/0! | 8.63E-03 |
| P07522 | LFWTDTGMSPR | 0,Methyl_2H(2)[AnyN-term]; | 0.0 | 12.2 | #DIV/0! | 8.70E-03 |
| P36375 | CVNIHLLSNEK | 1,Carbamidomethyl[C]; | 0.0 | 6.0 | #DIV/0! | 8.71E-03 |
| P36374 | CVNIHLLSNEK | 1,Carbamidomethyl[C]; | 0.0 | 6.0 | #DIV/0! | 8.71E-03 |
| P36373 | CVNIHLLSNEK | 1,Carbamidomethyl[C]; | 0.0 | 6.0 | #DIV/0! | 8.71E-03 |
| P15950 | CVNIHLLSNEK | 1,Carbamidomethyl[C]; | 0.0 | 6.0 | #DIV/0! | 8.71E-03 |
| P07647 | CVNIHLLSNEK | 1,Carbamidomethyl[C]; | 0.0 | 6.0 | #DIV/0! | 8.71E-03 |
| P00759 | CVNIHLLSNEK | 1,Carbamidomethyl[C]; | 0.0 | 6.0 | #DIV/0! | 8.71E-03 |
| P02761 | VFMQHIDVLENSLGFK | 5,DNCB_hapten[H]; | 0.0 | 15.8 | #DIV/0! | 8.75E-03 |
| P02761 | VEYDGGNTFTILK | 0,mTRAQ_13C(6)15N(2)[AnyN-term]; | 0.0 | 4.0 | #DIV/0! | 8.99E-03 |
| Q63041 | NTLPLNFDK | 0,NIC[AnyN-term](ICPL[AnyN-term]); | 0.0 | 3.5 | #DIV/0! | 9.05E-03 |
| P01836 | DGVLDSVTDQDSK | 1,Dehydrated[D]; | 0.0 | 6.2 | #DIV/0! | 9.07E-03 |
| P01835 | DGVLDSVTDQDSK | 1,Dehydrated[D]; | 0.0 | 6.2 | #DIV/0! | 9.07E-03 |
| P20767 | PTLTVFPPSTEELQGNK | 0,3sulfo[AnyN-term]; | 0.0 | 8.3 | #DIV/0! | 9.26E-03 |
| P10959 | HRAISESGVVLTTNLDKK | 0,Phosphogluconoylation[AnyN-term]; | 0.0 | 3.0 | #DIV/0! | 9.28E-03 |
| P02761 | MQHIDVLENSLGFK | 0,Iodoacetanilide_13C(6)[AnyN-term]; | 0.0 | 6.0 | #DIV/0! | 9.28E-03 |
| P22282 | TMYYISLPGSVR | 0,Cytopiloyne+water[AnyN-term]; | 0.0 | 6.7 | #DIV/0! | 9.36E-03 |
| Q63041 | DLNDNDAYSVFQSIGLK | 0,CLIP_TRAQ_2[AnyN-term]; | 0.0 | 6.5 | #DIV/0! | 9.38E-03 |
| P36373 | VGSTCLASGWGSTKPLIWEFPDDLQCVNIHLLSNEK | 5,Carbamidomethyl[C];26,Carbamidomethyl[C]; | 0.0 | 4.7 | #DIV/0! | 9.58E-03 |
| P02770 | LRDNYGELADCCAK | 10,Cation_Na[D];11,Carbamidomethyl[C];12,Carbamidomethyl[C]; | 0.0 | 3.8 | #DIV/0! | 9.79E-03 |
| P02761 | EKIEENGSMRVFMQHIDVLENSLGFK | 0,NDA[AnyN-term]; | 0.0 | 6.7 | #DIV/0! | 9.84E-03 |
| P05545 | IAELFSELDER | 3,Cation_Li[E]; | 0.0 | 6.5 | #DIV/0! | 9.88E-03 |
| P02770 | VQEVTDFAK | 0,NIC[AnyN-term](ICPL[AnyN-term]); | 0.0 | 16.0 | #DIV/0! | 9.90E-03 |
| P36373 | TSWGSVPCAKTNMPAIYTK | 0,ISD_z+2_ion[AnyN-term]; | 0.0 | 2.5 | #DIV/0! | 9.94E-03 |
| P15083 | WSNDGCHILPSHDEGAR | 3,Asn->Tyr[N]; | 0.0 | 2.5 | #DIV/0! | 9.94E-03 |
| P02761 | DLSSDIKEKFAKLCEAHGITR | 9,monomethylphosphothione[K]; | 0.0 | 2.8 | #DIV/0! | 1.00E-02 |
| P07522 | VHLHAQPGTENR | 0,NIC[AnyN-term](ICPL[AnyN-term]); | 0.0 | 4.0 | #DIV/0! | 1.03E-02 |
| P02761 | FAKLCEAHGITR | 1,Trioxidation[F]; | 0.0 | 4.0 | #DIV/0! | 1.03E-02 |
| Q9JHB9 | GSGCSILDEVIR | 4,Thiadiazole[C]; | 0.0 | 1.3 | #DIV/0! | 1.03E-02 |
| Q03626 | VYHKEESSCIHSSCTAER | 0,Methyl_2H(2)[AnyN-term];9,Carbamidomethyl[C];14,Carbamidomethyl[C]; | 0.0 | 1.3 | #DIV/0! | 1.03E-02 |
| P14046 | VYHKEESSCIHSSCTAER | 0,Methyl_2H(2)[AnyN-term];9,Carbamidomethyl[C];14,Carbamidomethyl[C]; | 0.0 | 1.3 | #DIV/0! | 1.03E-02 |
| P02780 | GSGCSILDEVIR | 4,Thiadiazole[C]; | 0.0 | 1.3 | #DIV/0! | 1.03E-02 |
| P02761 | FMQHIDVLENSLGFK | 0,Hex[AnyN-term]; | 0.0 | 3.2 | #DIV/0! | 1.04E-02 |
| P02761 | LCEAHGITR | 0,Phosphogluconoylation[AnyN-term]; | 0.0 | 3.2 | #DIV/0! | 1.04E-02 |
| P02761 | LNGDWFSIVVASNK | 15,Cation_Na[AnyC-term]; | 0.0 | 8.5 | #DIV/0! | 1.06E-02 |
| Q64240 | CGVPGDGYEELTR | 1,4AcAllylGal[C]; | 0.0 | 14.8 | #DIV/0! | 1.06E-02 |
| P14740 | CGIAVAPVSR | 1,Trioxidation[C]; | 0.0 | 5.2 | #DIV/0! | 1.06E-02 |
| P01836 | DGVLDSVTDQDSKDSTYSMSSTLSLTK | 1,Cation_Na[D];5,Cation_Na[D]; | 0.0 | 16.7 | #DIV/0! | 1.07E-02 |
| P01835 | DGVLDSVTDQDSKDSTYSMSSTLSLTK | 1,Cation_Na[D];5,Cation_Na[D]; | 0.0 | 16.7 | #DIV/0! | 1.07E-02 |
| P15978 | PGLGEPRFISVGYVDDTQFVR | 0,NIC[AnyN-term](ICPL[AnyN-term]); | 0.0 | 16.2 | #DIV/0! | 1.08E-02 |
| P02761 | NGETFQLMVLYGR | 0,Methyl[AnyN-term]; | 0.0 | 17.5 | #DIV/0! | 1.08E-02 |
| Q6IE52 | ESVVFVQTDKPMYKPGQSVK | 4,Val->Ser[V]; | 0.0 | 1.7 | #DIV/0! | 1.08E-02 |
| P14046 | ESVVFVQTDKPMYKPGQSVK | 4,Val->Ser[V]; | 0.0 | 1.7 | #DIV/0! | 1.08E-02 |
| P02761 | KLCEAHGITR | 0,Methyl_2H(2)[AnyN-term]; | 0.0 | 13.0 | #DIV/0! | 1.09E-02 |
| P07647 | SFQHPDYIPVFMR | 1,HexNAc(2)[S]; | 0.0 | 6.0 | #DIV/0! | 1.11E-02 |
| Q01177 | TSETDCMYGNGK | 6,Carbamidomethyl[C]; | 0.0 | 4.5 | #DIV/0! | 1.12E-02 |
| P83121 | HICQTYPDEICAWVVVTTR | 7,Pro->Gly[P]; | 0.0 | 30.8 | #DIV/0! | 1.12E-02 |
| P02782 | LEMYNAPPAAVEAK | 0,PyMIC[AnyN-term]; | 0.0 | 6.3 | #DIV/0! | 1.15E-02 |
| P02761 | LCEAHGITR | 2,GG[C](Dicarbamidomethyl[C]); | 0.0 | 22.3 | #DIV/0! | 1.15E-02 |
| P02761 | VEYDGGNTFTILK | 0,BITC[AnyN-term]; | 0.0 | 18.7 | #DIV/0! | 1.15E-02 |
| P07522 | LSVIEMADLDGSK | 6,Oxidation[M]; | 0.0 | 4.7 | #DIV/0! | 1.16E-02 |
| P02761 | VAKLNGDWFSIVVASNK | 3,Xlink_DTSSP[192][K]; | 0.0 | 9.3 | #DIV/0! | 1.16E-02 |
| P02761 | DWFSIVVASNKR | 2,Trp->Kynurenin[W]; | 0.0 | 12.8 | #DIV/0! | 1.17E-02 |
| Q9R0T4 | DTANWLEINPETGVISTR | 1,Cation_Al[III][D]; | 0.0 | 1.0 | #DIV/0! | 1.17E-02 |
| Q64319 | SFKDSDKDGNGDLK | 10,Deamidated[N]; | 0.0 | 1.0 | #DIV/0! | 1.17E-02 |
| P98158 | TGSENPLLVVASR | 1,PET[T]; | 0.0 | 11.0 | #DIV/0! | 1.17E-02 |
| P36375 | LPTEEPKVGSTCLASGWGSTK | 0,ISD_z+2_ion[AnyN-term]; | 0.0 | 2.0 | #DIV/0! | 1.17E-02 |
| P36373 | LPTEEPKVGSTCLASGWGSTK | 0,ISD_z+2_ion[AnyN-term]; | 0.0 | 2.0 | #DIV/0! | 1.17E-02 |
| P15684 | TRALEQALEK | 1,Glucuronyl[T]; | 0.0 | 2.0 | #DIV/0! | 1.17E-02 |
| P07861 | ISGAAVVNAFYSSGR | 2,Glucuronyl[S]; | 0.0 | 1.0 | #DIV/0! | 1.17E-02 |
| P02782 | ASQICELVAHETISFLMK | 4,Xle->Met[I]; | 0.0 | 2.0 | #DIV/0! | 1.17E-02 |
| P02770 | LRDNYGELADCCAK | 11,DTT[C]; | 0.0 | 3.0 | #DIV/0! | 1.17E-02 |
| P00787 | QGSCGSCWAFGAVEAMSDR | 7,EQAT_2H(5)[C]; | 0.0 | 1.0 | #DIV/0! | 1.17E-02 |
| P00786 | NQGACGSCWTFSTTGALESAVAIASGK | 5,Carbamidomethyl[C];8,Carbamidomethyl[C]; | 0.0 | 1.0 | #DIV/0! | 1.17E-02 |
| Q63041 | WWVDEYHTQADHSAAR | 0,Piperidine[AnyN-term]; | 0.0 | 4.2 | #DIV/0! | 1.18E-02 |
| P02761 | RGNLDVAKLNGDWFSIVVASNKR | 10,Asn->His[N]; | 0.0 | 9.0 | #DIV/0! | 1.20E-02 |
| Q9JHB9 | CSILDEVIR | 1,Dimethylphosphothione[C]; | 0.0 | 3.7 | #DIV/0! | 1.21E-02 |
| Q64319 | PIFGTMKDFENLVAAVHDK | 0,Cytopiloyne+water[AnyN-term]; | 0.0 | 1.8 | #DIV/0! | 1.21E-02 |
| Q63041 | YNILPEAEGEAPFTLK | 0,Formyl[AnyN-term]; | 0.0 | 1.8 | #DIV/0! | 1.21E-02 |
| P50123 | TNLLYDPLLSASSNQQR | 1,PET[T]; | 0.0 | 1.8 | #DIV/0! | 1.21E-02 |
| P14841 | PRLLGAPQEADASEEGVQR | 0,Dicarbamidomethyl[AnyN-term]; | 0.0 | 1.8 | #DIV/0! | 1.21E-02 |
| P02780 | CSILDEVIR | 1,Dimethylphosphothione[C]; | 0.0 | 3.7 | #DIV/0! | 1.21E-02 |
| P02761 | ENGSMRVFMQHIDVLENSLGFK | 0,Acetyl[ProteinN-term]; | 0.0 | 1.8 | #DIV/0! | 1.21E-02 |
| P02761 | FMQHIDVLENSLGFK | 4,DNCB_hapten[H]; | 0.0 | 12.0 | #DIV/0! | 1.22E-02 |
| P07522 | LRGSMLKPSSLVVVHPLAK | 0,Unknown_302[AnyN-term]; | 0.0 | 8.2 | #DIV/0! | 1.23E-02 |
| P20059 | NPVTGEVPPR | 0,PEITC[AnyN-term]; | 0.0 | 6.2 | #DIV/0! | 1.26E-02 |
| Q64230 | DSSQPGQVDHTLVGR | 0,GIST-Quat[AnyN-term]; | 0.0 | 1.2 | #DIV/0! | 1.27E-02 |
| P21704 | LTNQMAEAISDHYPVEVTLR | 3,Asn->His[N]; | 0.0 | 1.2 | #DIV/0! | 1.27E-02 |
| P02761 | EENGSMRVFMQHIDVLENSLGFK | 7,Arg->Lys[R]; | 0.0 | 1.2 | #DIV/0! | 1.27E-02 |
| P02761 | CRELYLVAYKTPEDGEYFVEYDGGNTFTILK | 0,dNIC[AnyN-term]; | 0.0 | 1.2 | #DIV/0! | 1.27E-02 |
| P21704 | LLDELNRDIPDNYR | 3,Cation_Na[D]; | 0.0 | 4.3 | #DIV/0! | 1.28E-02 |
| P02761 | FMQHIDVLENSLGFK | 4,Aspartylurea[H]; | 0.0 | 4.3 | #DIV/0! | 1.28E-02 |
| P07522 | SVLILDILDKR | 0,dNIC[AnyN-term]; | 0.0 | 2.8 | #DIV/0! | 1.29E-02 |
| P22283 | TPGVTIYYITLPGSVR | 1,Deoxy[T]; | 0.0 | 3.8 | #DIV/0! | 1.29E-02 |
| P00884 | YTPEQVAMATVTALHR | 3,Pro->Cys[P]; | 0.0 | 3.8 | #DIV/0! | 1.29E-02 |
| P02770 | TVDETYVPK | 0,TNBS[AnyN-term]; | 0.0 | 9.3 | #DIV/0! | 1.29E-02 |
| Q9JHB9 | SGCSILDEVIR | 0,AccQTag[AnyN-term]; | 0.0 | 4.2 | #DIV/0! | 1.32E-02 |
| P02780 | SGCSILDEVIR | 0,AccQTag[AnyN-term]; | 0.0 | 4.2 | #DIV/0! | 1.32E-02 |
| P02761 | PEDGEYFVEYDGGNTFTILK | 6,DiART6plex115[Y]; | 0.0 | 14.3 | #DIV/0! | 1.32E-02 |
| P81828 | QKCQTYPDEICAWVVVTTR | 3,Carbamidomethyl[C];11,Carbamidomethyl[C]; | 0.0 | 119.7 | #DIV/0! | 1.34E-02 |
| P07647 | VGSICLASGWGMTNPSEMK | 5,Propionamide_2H(3)[C]; | 0.0 | 9.3 | #DIV/0! | 1.35E-02 |
| P02761 | VFMQHIDVLENSLGFK | 2,Phe->Asp[F]; | 0.0 | 5.7 | #DIV/0! | 1.36E-02 |
| Q64230 | AIMTILDQEPDAR | 0,PyMIC[AnyN-term]; | 0.0 | 2.5 | #DIV/0! | 1.36E-02 |
| P02782 | SEEELKKELEMYNAPPAAVEAK | 8,Cation_Ca[II][E]; | 0.0 | 2.5 | #DIV/0! | 1.36E-02 |
| P01836 | STYSMSSTLSLTK | 0,Iodoacetanilide[AnyN-term]; | 0.0 | 2.5 | #DIV/0! | 1.36E-02 |
| P01835 | STYSMSSTLSLTK | 0,Iodoacetanilide[AnyN-term]; | 0.0 | 2.5 | #DIV/0! | 1.36E-02 |
| P42854 | RPSGHLVSVLSGSEASFVSSLIK | 14,Cation_Ca[II][E]; | 0.0 | 12.7 | #DIV/0! | 1.37E-02 |
| P27590 | INFECSYPLDMK | 5,Carbamidomethyl[C];10,Cation_Na[D]; | 0.0 | 7.7 | #DIV/0! | 1.37E-02 |
| Q6DGG1 | GYVPVAPICTDKINAADYAR | 9,Carbamidomethyl[C]; | 0.0 | 4.8 | #DIV/0! | 1.38E-02 |
| P15083 | CPVLVGTQALVQEGYEGR | 0,Dimethyl_2H(4)[AnyN-term]; | 0.0 | 6.2 | #DIV/0! | 1.38E-02 |
| P07632 | DGPVQGVIHFEQK | 3,Pro->Phe[P]; | 0.0 | 3.7 | #DIV/0! | 1.39E-02 |
| P02782 | SQICELVAHETISFLMK | 0,Propyl[AnyN-term]; | 0.0 | 3.7 | #DIV/0! | 1.39E-02 |
| P47728 | LLPVQENFLLK | 1,Xle->Glu[L]; | 0.0 | 4.3 | #DIV/0! | 1.41E-02 |
| P07171 | LLPVQENFLLK | 1,Xle->Glu[L]; | 0.0 | 4.3 | #DIV/0! | 1.41E-02 |
| P02782 | EEELKKELEMYNAPPAAVEAK | 10,Oxidation[M]; | 0.0 | 4.3 | #DIV/0! | 1.41E-02 |
| P02761 | NGETFQLMVLYGR | 3,Carbonyl[E]; | 0.0 | 12.0 | #DIV/0! | 1.44E-02 |
| Q9EQV9 | KHGLAFQSGHVLSALPR | 0,SMA[AnyN-term]; | 0.0 | 7.7 | #DIV/0! | 1.45E-02 |
| P02761 | KIEENGSMR | 5,Deamidated[N];8,Oxidation[M]; | 0.0 | 3.8 | #DIV/0! | 1.45E-02 |
| P83121 | CQTYPDEICAWVVVTTR | 1,Carbamidomethyl[C]; | 0.0 | 14.2 | #DIV/0! | 1.47E-02 |
| P81828 | CQTYPDEICAWVVVTTR | 1,Carbamidomethyl[C]; | 0.0 | 14.2 | #DIV/0! | 1.47E-02 |
| P81827 | CQTYPDEICAWVVVTTR | 1,Carbamidomethyl[C]; | 0.0 | 14.2 | #DIV/0! | 1.47E-02 |
| P02761 | QHIDVLENSLGFK | 3,Xle->Asp[I]; | 0.0 | 3.2 | #DIV/0! | 1.51E-02 |
| P00787 | NFYNVDISYLK | 2,Phe->Met[F]; | 0.0 | 5.5 | #DIV/0! | 1.53E-02 |
| P81828 | KCQTYPDEICAWVVVTTR | 2,Cys->Trp[C]; | 0.0 | 2.2 | #DIV/0! | 1.55E-02 |
| P02770 | RFPNAEFAEITK | 0,CLIP_TRAQ_4[AnyN-term]; | 0.0 | 2.2 | #DIV/0! | 1.55E-02 |
| Q9JHB9 | IRGTINSTVTLHDYMK | 0,Unknown_302[AnyN-term]; | 0.0 | 5.7 | #DIV/0! | 1.59E-02 |
| Q5M876 | VAVTGGTHGNEMCGVYLAR | 13,Carbamidomethyl[C]; | 0.0 | 2.8 | #DIV/0! | 1.59E-02 |
| P22282 | EICYFVVYPDYIEQNIHAVR | 1,Propargylamine[E]; | 0.0 | 5.7 | #DIV/0! | 1.59E-02 |
| P02780 | IRGTINSTVTLHDYMK | 0,Unknown_302[AnyN-term]; | 0.0 | 5.7 | #DIV/0! | 1.59E-02 |
| P02770 | PERNECFLQHKDDNPNLPPFQRPEAEAMCTSFQENPTSFLGHYLHEVAR | 29,4AcAllylGal[C]; | 0.0 | 3.8 | #DIV/0! | 1.63E-02 |
| P02761 | DVLENSLGFK | 0,dNIC[AnyN-term]; | 0.0 | 11.5 | #DIV/0! | 1.63E-02 |
| P02761 | GDWFSIVVASNKR | 0,Hex[AnyN-term]; | 0.0 | 8.7 | #DIV/0! | 1.65E-02 |
| P02761 | VFMQHIDVLENSLGFK | 0,Piperidine[AnyN-term]; | 0.0 | 9.2 | #DIV/0! | 1.65E-02 |
| P06866 | LQTEGDGIYTLNSEK | 4,Glu->His[E]; | 0.0 | 8.5 | #DIV/0! | 1.67E-02 |
| P02761 | TPEDGEYFVEYDGGNTFTILK | 10,Hydroxamic_acid[E]; | 0.0 | 3.5 | #DIV/0! | 1.67E-02 |
| P02761 | IDVLENSLGFK | 0,Propionyl_13C(3)[AnyN-term]; | 0.0 | 3.5 | #DIV/0! | 1.67E-02 |
| P00758 | QPGDDYSNDLMLLHLSQPADITDGVK | 0,Gln->pyro-Glu[AnyN-termQ];5,Cation_Na[D]; | 0.0 | 3.5 | #DIV/0! | 1.67E-02 |
| P07647 | QSFQHPDYIPVFMR | 2,CresylSaligeninPhosphate[S]; | 0.0 | 5.7 | #DIV/0! | 1.67E-02 |
| P07522 | DTKLSVIEMADLDGSKR | 2,HexN[T]; | 0.0 | 5.7 | #DIV/0! | 1.67E-02 |
| P02761 | KLCEAHGITR | 0,ISD_z+2_ion[AnyN-term]; | 0.0 | 8.2 | #DIV/0! | 1.68E-02 |
| P07522 | CDTKLSVIEMADLDGSKR | 7,Val->Arg[V]; | 0.0 | 5.5 | #DIV/0! | 1.70E-02 |
| Q64319 | FMGTEVSAESTER | 2,Oxidation[M]; | 0.0 | 5.3 | #DIV/0! | 1.71E-02 |
| P22283 | LNDVYDIFK | 1,Carbonyl[L]; | 0.0 | 2.7 | #DIV/0! | 1.71E-02 |
| P02761 | LNGDWFSIVVASNKR | 8,Xle->Arg[I]; | 0.0 | 2.7 | #DIV/0! | 1.71E-02 |
| Q64230 | QAIMTILDQEPDAR | 4,Met->Cys[M]; | 0.0 | 3.0 | #DIV/0! | 1.72E-02 |
| P31211 | FSGNTKDVPLTLTMVHK | 8,Val->Cys[V]; | 0.0 | 3.0 | #DIV/0! | 1.72E-02 |
| P20059 | PVTSVDAAFR | 0,dNIC[AnyN-term]; | 0.0 | 1.5 | #DIV/0! | 1.72E-02 |
| P05545 | DKEQPILSEFQEK | 0,Galactosyl[AnyN-term]; | 0.0 | 4.5 | #DIV/0! | 1.72E-02 |
| P05544 | DKEQPILSEFQEK | 0,Galactosyl[AnyN-term]; | 0.0 | 4.5 | #DIV/0! | 1.72E-02 |
| P04937 | WAPPPSIELTNLLVR | 6,PhosphoHex[S]; | 0.0 | 1.5 | #DIV/0! | 1.72E-02 |
| P02770 | QENPTSFLGHYLHEVAR | 5,HexNAc(1)NeuAc(1)[T]; | 0.0 | 1.5 | #DIV/0! | 1.72E-02 |
| P02761 | IKEKFAKLCEAHGITR | 0,PyMIC[AnyN-term]; | 0.0 | 1.5 | #DIV/0! | 1.72E-02 |
| P01836 | HNLYTCEVVHKTSSSPVVK | 6,Ub-Br2[C]; | 0.0 | 1.5 | #DIV/0! | 1.72E-02 |
| P01835 | HNLYTCEVVHKTSSSPVVK | 6,Ub-Br2[C]; | 0.0 | 1.5 | #DIV/0! | 1.72E-02 |
| P00787 | NGPVEGAFTVFSDFLTYK | 0,Propyl[AnyN-term]; | 0.0 | 1.5 | #DIV/0! | 1.72E-02 |
| P02761 | FMQHIDVLENSLGFK | 0,Carbamyl[AnyN-term]; | 0.0 | 27.3 | #DIV/0! | 1.72E-02 |
| P98158 | VTGSENPLLVVASR | 5,Cation_Na[E]; | 0.0 | 4.7 | #DIV/0! | 1.73E-02 |
| Q64240 | EYCGVPGDGYEELTR | 3,Carbamidomethyl[C];8,Cation_Na[D]; | 0.0 | 9.3 | #DIV/0! | 1.73E-02 |
| P36373 | PGDDHSNDLMLLHLSQPADITDGVK | 6,pyrophospho[S]; | 0.0 | 2.3 | #DIV/0! | 1.73E-02 |
| P02761 | KIEENGSMRVFMQHIDVLENSLGFK | 0,PyMIC[AnyN-term]; | 0.0 | 2.3 | #DIV/0! | 1.73E-02 |
| P02761 | ETFQLMVLYGR | 0,BITC[AnyN-term]; | 0.0 | 11.5 | #DIV/0! | 1.74E-02 |
| P02761 | TPEDGEYFVEYDGGNTFTILK | 10,Carbonyl[E]; | 0.0 | 30.0 | #DIV/0! | 1.75E-02 |
| P26644 | VCPFAGILENGVVR | 2,Carbamidomethyl[C]; | 0.0 | 13.7 | #DIV/0! | 1.76E-02 |
| P02761 | EKIEENGSMR | 2,Lys->Thr[K]; | 0.0 | 3.2 | #DIV/0! | 1.76E-02 |
| Q64230 | DDADWVHEDSSQPGQVDHTLVGR | 12,Methyl+Deamidated[Q]; | 0.0 | 3.7 | #DIV/0! | 1.77E-02 |
| P07522 | VTGNLHRADLGGMDVK | 0,TMAB_2H(9)[AnyN-term]; | 0.0 | 5.0 | #DIV/0! | 1.77E-02 |
| P02761 | GNLDVAKLNGDWFSIVVASNKR | 20,Deamidated[N]; | 0.0 | 2.5 | #DIV/0! | 1.77E-02 |
| P02761 | VFMQHIDVLENSLGFK | 0,Diisopropylphosphate[AnyN-term]; | 0.0 | 12.3 | #DIV/0! | 1.79E-02 |
| P12346 | VSTVLTAQK | 2,Hex(1)HexNAc(1)Phos(1)[S]; | 0.0 | 2.0 | #DIV/0! | 1.80E-02 |
| P04937 | LGVRPSQGGEAPR | 0,C+12[AnyN-term]; | 0.0 | 2.0 | #DIV/0! | 1.80E-02 |
| P02770 | STPTLVEAAR | 0,Cytopiloyne+water[AnyN-term]; | 0.0 | 2.0 | #DIV/0! | 1.80E-02 |
| P02761 | KLCEAHGITR | 0,Iodoacetanilide[AnyN-term]; | 0.0 | 2.0 | #DIV/0! | 1.80E-02 |
| P02761 | EKIEENGSMR | 1,Propargylamine[E]; | 0.0 | 4.0 | #DIV/0! | 1.80E-02 |
| P01836 | DSTYSMSSTLSLTK | 1,Dehydrated[D]; | 0.0 | 4.0 | #DIV/0! | 1.80E-02 |
| P01835 | DSTYSMSSTLSLTK | 1,Dehydrated[D]; | 0.0 | 4.0 | #DIV/0! | 1.80E-02 |
| P42854 | SSGNSGQNVWIGLHDPTLGQEPNR | 4,Asn->Met[N]; | 0.0 | 5.2 | #DIV/0! | 1.80E-02 |
| P02761 | KIEENGSMRVFMQHIDVLENSLGFK | 0,Iodoacetanilide[AnyN-term]; | 0.0 | 5.2 | #DIV/0! | 1.80E-02 |
| P28826 | VVFEGVGGPGASSGGLSIDDINLSETR | 0,Methyl_2H(2)[AnyN-term]; | 0.0 | 3.8 | #DIV/0! | 1.81E-02 |
| P02761 | AKLCEAHGITR | 0,Diethylphosphate[AnyN-term]; | 0.0 | 8.3 | #DIV/0! | 1.83E-02 |
| P42854 | SSGNSGQNVWIGLHDPTLGQEPNR | 5,Ser->Cys[S]; | 0.0 | 8.2 | #DIV/0! | 1.84E-02 |
| P14841 | EGVQRALDFAVSEYNK | 6,Ala->His[A]; | 0.0 | 4.7 | #DIV/0! | 1.85E-02 |
| P02770 | NPTSFLGHYLHEVAR | 0,Delta_H(2)C(2)[AnyN-term]; | 0.0 | 9.5 | #DIV/0! | 1.85E-02 |
| P02761 | VLENSLGFK | 0,NIC[AnyN-term](ICPL[AnyN-term]); | 0.0 | 4.5 | #DIV/0! | 1.85E-02 |
| Q63691 | NAGMETTSGVCSALAAAR | 11,Carbamidomethyl[C]; | 0.0 | 6.8 | #DIV/0! | 1.86E-02 |
| P02761 | PEDGEYFVEYDGGNTFTILK | 6,DiART6plex118[Y]; | 0.0 | 15.3 | #DIV/0! | 1.86E-02 |
| P27590 | GERDWMSIVTPAR | 0,ISD_z+2_ion[AnyN-term]; | 0.0 | 10.3 | #DIV/0! | 1.88E-02 |
| P07647 | VGSICLASGWGMTNPSEMK | 5,Carbamidomethyl[C];18,Oxidation[M]; | 0.0 | 22.8 | #DIV/0! | 1.89E-02 |
| P02761 | RGNLDVAKLNGDWFSIVVASNKR | 3,Asn->His[N]; | 0.0 | 3.5 | #DIV/0! | 1.89E-02 |
| P02761 | HLINFKNGETFQLMVLYGR | 0,Formyl[AnyN-term]; | 0.0 | 4.2 | #DIV/0! | 1.91E-02 |
| P27590 | STDYGAGYSCDSDMHGWYR | 3,Cation_Na[D];10,Carbamidomethyl[C]; | 0.0 | 14.2 | #DIV/0! | 1.91E-02 |
| P14740 | YMGLPTPEDNLDHYR | 0,Methyl_2H(3)13C(1)[AnyN-term]; | 0.0 | 7.8 | #DIV/0! | 1.92E-02 |
| P22282 | DNCPFEEQTEQLK | 0,Iodoacetanilide_13C(6)[AnyN-term]; | 0.0 | 2.8 | #DIV/0! | 1.93E-02 |
| P06866 | LQTEGDGIYTLNSEK | 2,Gln->His[Q]; | 0.0 | 2.8 | #DIV/0! | 1.93E-02 |
| Q5XI43 | VRLEVTDDPLLSR | 0,Cytopiloyne+water[AnyN-term]; | 0.0 | 4.3 | #DIV/0! | 1.97E-02 |
| P36374 | NNLLEDEPFAQHR | 6,Cation_Ca[II][D]; | 0.0 | 1.8 | #DIV/0! | 1.97E-02 |
| P36373 | NNLLEDEPFAQHR | 6,Cation_Ca[II][D]; | 0.0 | 1.8 | #DIV/0! | 1.97E-02 |
| P21704 | TNQMAEAISDHYPVEVTLR | 0,Diethylphosphate[AnyN-term]; | 0.0 | 7.3 | #DIV/0! | 1.97E-02 |
| P07522 | IESSSLQGSDR | 1,Xle->Thr[I]; | 0.0 | 1.8 | #DIV/0! | 1.97E-02 |
| P00758 | QPGDDYSNDLMLLHLSQPADITDGVK | 0,Gln->pyro-Glu[AnyN-termQ];9,Cation_Na[D]; | 0.0 | 1.8 | #DIV/0! | 1.97E-02 |
| Q63041 | AESPVFVQTDKPIYKPGQTVK | 10,Cation_Na[D]; | 0.0 | 1.7 | #DIV/0! | 1.99E-02 |
| P04937 | LDSPTGFDSSDVTANSFTVHWVAPR | 0,SPITC[AnyN-term]; | 0.0 | 5.0 | #DIV/0! | 1.99E-02 |
| P02761 | KLCEAHGITR | 0,Succinyl[AnyN-term]; | 0.0 | 13.7 | #DIV/0! | 1.99E-02 |
| P02761 | DNIIDLTKTDR | 0,Delta_H(2)C(2)[AnyN-term]; | 0.0 | 3.8 | #DIV/0! | 1.99E-02 |
| Q9R0J8 | HQADACHAYQIIHR | 6,Carbamidomethyl[C]; | 0.0 | 5.2 | #DIV/0! | 2.01E-02 |
| P07647 | HLSKPADITGGVK | 3,Ser->Gly[S]; | 0.0 | 5.7 | #DIV/0! | 2.01E-02 |
| P02761 | MQHIDVLENSLGFK | 0,GIST-Quat_2H(9)[AnyN-term]; | 0.0 | 7.5 | #DIV/0! | 2.01E-02 |
| P07522 | WCDTKLSVIEMADLDGSK | 2,Nitrosyl[C]; | 0.0 | 3.0 | #DIV/0! | 2.02E-02 |
| P02761 | KLCEAHGITR | 1,DNCB_hapten[K]; | 0.0 | 3.0 | #DIV/0! | 2.02E-02 |
| P02761 | FMQHIDVLENSLGFK | 0,AccQTag[AnyN-term]; | 0.0 | 21.2 | #DIV/0! | 2.04E-02 |
| P02770 | CFATEGPNLVAR | 0,Methyl_2H(2)[AnyN-term]; | 0.0 | 12.7 | #DIV/0! | 2.05E-02 |
| P02761 | LCEAHGITRDNIIDLTKTDR | 2,Carbamidomethyl[C]; | 0.0 | 10.3 | #DIV/0! | 2.06E-02 |
| P02761 | VFMQHIDVLENSLGFK | 3,Oxidation[M];7,Cation_Na[D]; | 0.0 | 8.8 | #DIV/0! | 2.07E-02 |
| P83121 | DEICAWVVVTTR | 4,Cys->SecNEM_2H(5)[C]; | 0.0 | 11.3 | #DIV/0! | 2.08E-02 |
| P81828 | DEICAWVVVTTR | 4,Cys->SecNEM_2H(5)[C]; | 0.0 | 11.3 | #DIV/0! | 2.08E-02 |
| P81827 | DEICAWVVVTTR | 4,Cys->SecNEM_2H(5)[C]; | 0.0 | 11.3 | #DIV/0! | 2.08E-02 |
| P02782 | LEMYNAPPAAVEAK | 0,Iodoacetanilide[AnyN-term]; | 0.0 | 12.3 | #DIV/0! | 2.08E-02 |
| P47967 | FDENAVVR | 0,C+12[AnyN-term]; | 0.0 | 2.7 | #DIV/0! | 2.10E-02 |
| P12788 | DTLDNDIMLIK | 1,phosphoRibosyl[D]; | 0.0 | 2.7 | #DIV/0! | 2.10E-02 |
| P02761 | GNLDVAKLNGDWFSIVVASNKR | 0,NDA[AnyN-term]; | 0.0 | 5.3 | #DIV/0! | 2.10E-02 |
| P00787 | QGSCGSCWAFGAVEAMSDR | 4,EQAT_2H(5)[C]; | 0.0 | 2.7 | #DIV/0! | 2.10E-02 |
| P02780 | LVKPYVQDHFTEKAVK | 0,Carbamyl[AnyN-term]; | 0.0 | 7.5 | #DIV/0! | 2.11E-02 |
| Q68FP1 | SAAYLWVGTGASDAEK | 1,Hep[S]; | 0.0 | 6.7 | #DIV/0! | 2.11E-02 |
| P10758 | GYCVSVTSNSGYK | 3,Carbamidomethyl[C]; | 0.0 | 4.3 | #DIV/0! | 2.12E-02 |
| P07647 | AYDHNNDLMLLHLSK | 14,Dehydrated[S]; | 0.0 | 2.2 | #DIV/0! | 2.12E-02 |
| P02761 | LINFKNGETFQLMVLYGR | 0,Hex[AnyN-term]; | 0.0 | 2.2 | #DIV/0! | 2.12E-02 |
| P02761 | TPEDGEYFVEYDGGNTFTILK | 12,Asp->Lys[D]; | 0.0 | 4.5 | #DIV/0! | 2.13E-02 |
| P38918 | QVETELFPCLR | 9,Carbamidomethyl[C]; | 0.0 | 6.2 | #DIV/0! | 2.15E-02 |
| P04937 | FTNIGPDTMR | 9,Oxidation[M]; | 0.0 | 6.5 | #DIV/0! | 2.18E-02 |
| P13432 | LSNPPTQLHSTEQANTK | 0,C+12[AnyN-term]; | 0.0 | 2.5 | #DIV/0! | 2.21E-02 |
| P02761 | VFMQHIDVLENSLGFK | 5,Crotonaldehyde[H]; | 0.0 | 2.5 | #DIV/0! | 2.21E-02 |
| P02761 | VFMQHIDVLENSLGFKFR | 15,Phe->CamCys[F]; | 0.0 | 2.5 | #DIV/0! | 2.21E-02 |
| P02761 | LCEAHGITR | 1,Xle->Met[L]; | 0.0 | 2.5 | #DIV/0! | 2.21E-02 |
| P02761 | QHIDVLENSLGFK | 8,Deamidated[N]; | 0.0 | 6.3 | #DIV/0! | 2.25E-02 |
| P01681 | DVQMTQSPSYLAASPGESVSISCK | 1,Cation_Na[D];4,Oxidation[M];23,Carbamidomethyl[C]; | 0.0 | 31.0 | #DIV/0! | 2.26E-02 |
| P02761 | LNGDWFSIVVASNK | 0,glycidamide[AnyN-term]; | 0.0 | 7.5 | #DIV/0! | 2.26E-02 |
| P68136 | SYELPDGQVITIGNER | 1,azole[S]; | 0.0 | 4.5 | #DIV/0! | 2.27E-02 |
| P68035 | SYELPDGQVITIGNER | 1,azole[S]; | 0.0 | 4.5 | #DIV/0! | 2.27E-02 |
| P63269 | SYELPDGQVITIGNER | 1,azole[S]; | 0.0 | 4.5 | #DIV/0! | 2.27E-02 |
| P63259 | SYELPDGQVITIGNER | 1,azole[S]; | 0.0 | 4.5 | #DIV/0! | 2.27E-02 |
| P62738 | SYELPDGQVITIGNER | 1,azole[S]; | 0.0 | 4.5 | #DIV/0! | 2.27E-02 |
| P60711 | SYELPDGQVITIGNER | 1,azole[S]; | 0.0 | 4.5 | #DIV/0! | 2.27E-02 |
| P98158 | TCKVTGSENPLLVVASR | 3,Dap-DSP[K]; | 0.0 | 8.5 | #DIV/0! | 2.28E-02 |
| P02761 | REKIEENGSMR | 0,Galactosyl[AnyN-term]; | 0.0 | 2.8 | #DIV/0! | 2.28E-02 |
| P02761 | TPEDGEYFVEYDGGNTFTILK | 9,Val->Asn[V]; | 0.0 | 7.7 | #DIV/0! | 2.28E-02 |
| P83121 | ICQTYPDEICAWVVVTTRDGK | 2,Carbamidomethyl[C];10,Carbamidomethyl[C]; | 0.0 | 10.7 | #DIV/0! | 2.32E-02 |
| P81827 | ICQTYPDEICAWVVVTTRDGK | 2,Carbamidomethyl[C];10,Carbamidomethyl[C]; | 0.0 | 10.7 | #DIV/0! | 2.32E-02 |
| P36373 | LSQPADITDGVK | 0,BITC[AnyN-term]; | 0.0 | 5.2 | #DIV/0! | 2.32E-02 |
| P00758 | LSQPADITDGVK | 0,BITC[AnyN-term]; | 0.0 | 5.2 | #DIV/0! | 2.32E-02 |
| P32038 | TLCDVAGWGVVTHAGR | 3,Cys->Met[C]; | 0.0 | 3.0 | #DIV/0! | 2.34E-02 |
| P07522 | SVIEGSDLSGK | 2,Val->Ser[V]; | 0.0 | 3.0 | #DIV/0! | 2.34E-02 |
| P08723 | EHVTQVSGTR | 1,Glu->Met[E]; | 0.0 | 5.8 | #DIV/0! | 2.36E-02 |
| P02761 | VFMQHIDVLENSLGFK | 0,Propionamide[AnyN-term]; | 0.0 | 24.7 | #DIV/0! | 2.37E-02 |
| P08932 | CFQPIPMDSSDLKPVLK | 1,Dimethylphosphothione[C]; | 0.0 | 4.7 | #DIV/0! | 2.37E-02 |
| P01048 | CFQPIPMDSSDLKPVLK | 1,Dimethylphosphothione[C]; | 0.0 | 4.7 | #DIV/0! | 2.37E-02 |
| P02781 | ELEEFDAPPEAVEANLK | 5,Trioxidation[F]; | 0.0 | 3.8 | #DIV/0! | 2.38E-02 |
| P02761 | EENGSMRVFMQHIDVLENSLGFK | 6,Met->Cys[M]; | 0.0 | 3.7 | #DIV/0! | 2.39E-02 |
| P15083 | CPVLVGTQALVQEGYEGR | 1,Dioxidation[C]; | 0.0 | 4.2 | #DIV/0! | 2.40E-02 |
| P01836 | DGVLDSVTDQDSKDSTYSMSSTLSLTK | 1,Carboxy[D]; | 0.0 | 11.3 | #DIV/0! | 2.41E-02 |
| P01835 | DGVLDSVTDQDSKDSTYSMSSTLSLTK | 1,Carboxy[D]; | 0.0 | 11.3 | #DIV/0! | 2.41E-02 |
| Q9JHB9 | ASGSGCSILDEVIR | 6,Carbamidomethyl[C]; | 0.0 | 6.0 | #DIV/0! | 2.42E-02 |
| P02780 | ASGSGCSILDEVIR | 6,Carbamidomethyl[C]; | 0.0 | 6.0 | #DIV/0! | 2.42E-02 |
| P22283 | PFEEQTEQLKR | 0,ISD_z+2_ion[AnyN-term]; | 0.0 | 9.3 | #DIV/0! | 2.44E-02 |
| P22282 | PFEEQTEQLKR | 0,ISD_z+2_ion[AnyN-term]; | 0.0 | 9.3 | #DIV/0! | 2.44E-02 |
| Q6AYD4 | LSNKPGVTLVHSISTR | 0,AccQTag[AnyN-term]; | 0.0 | 1.3 | #DIV/0! | 2.50E-02 |
| Q64319 | YSELYHDFTTTQVGMHDLVR | 5,Tyr->Arg[Y]; | 0.0 | 1.3 | #DIV/0! | 2.50E-02 |
| P83121 | RHICQTYPDEICAWVVVTTR | 4,Cys->Dha[C]; | 0.0 | 0.7 | #DIV/0! | 2.50E-02 |
| P83121 | FRCISFDSTGFCYVGR | 3,Cys->Trp[C]; | 0.0 | 0.7 | #DIV/0! | 2.50E-02 |
| P83121 | VGRHICQTYPDEICAWVVVTTR | 6,Carbamidomethyl[C];14,Carbamidomethyl[C]; | 0.0 | 0.7 | #DIV/0! | 2.50E-02 |
| P83121 | PDEICAWVVVTTR | 5,GG[C](Dicarbamidomethyl[C]); | 0.0 | 0.7 | #DIV/0! | 2.50E-02 |
| P81828 | RQKCQTYPDEICAWVVVTTR | 4,Carbamidomethyl[C];12,Carbamidomethyl[C]; | 0.0 | 1.3 | #DIV/0! | 2.50E-02 |
| P81828 | PDEICAWVVVTTR | 5,GG[C](Dicarbamidomethyl[C]); | 0.0 | 0.7 | #DIV/0! | 2.50E-02 |
| P81827 | PDEICAWVVVTTR | 5,GG[C](Dicarbamidomethyl[C]); | 0.0 | 0.7 | #DIV/0! | 2.50E-02 |
| P48508 | EFPDVLECTMSHAVEK | 8,Carbamidomethyl[C]; | 0.0 | 0.7 | #DIV/0! | 2.50E-02 |
| P22282 | TPGETMYYISLPGSVR | 0,Methylthio[AnyN-term]; | 0.0 | 0.7 | #DIV/0! | 2.50E-02 |
| P17475 | NNYHSEAFSVNFADSEEAKK | 6,Cation_Na[E]; | 0.0 | 2.0 | #DIV/0! | 2.50E-02 |
| P17475 | VFNNDADLSGITEDAPLK | 5,Cation_Na[D]; | 0.0 | 2.0 | #DIV/0! | 2.50E-02 |
| P17475 | VNFADSEEAKKVINDYVEK | 0,Dimethyl_2H(4)[AnyN-term]; | 0.0 | 0.7 | #DIV/0! | 2.50E-02 |
| P17475 | PKLSISGTYNLK | 0,LG-anhyropyrrole[AnyN-term]; | 0.0 | 0.7 | #DIV/0! | 2.50E-02 |
| P10758 | ESGTTAANVWIGLHDPK | 1,Cation_Na[E]; | 0.0 | 1.3 | #DIV/0! | 2.50E-02 |
| P08723 | YKEFLLEDGEHVTQVSGTR | 7,Carboxy[E]; | 0.0 | 1.3 | #DIV/0! | 2.50E-02 |
| P07647 | AYDHNNDLMLLHLSKPADITGGVK | 13,Dioxidation[L]; | 0.0 | 1.3 | #DIV/0! | 2.50E-02 |
| P05545 | AVLDVDETGTEGAAATAVTAALK | 4,Cation_Na[D]; | 0.0 | 1.3 | #DIV/0! | 2.50E-02 |
| P04937 | IPAPTNLKFTQVSPTTLTAQWTAPSVK | 11,Xlink_DFDNB[Q]; | 0.0 | 0.7 | #DIV/0! | 2.50E-02 |
| P02770 | AADKDNCFATEGPNLVAR | 7,NEM_2H(5)+H2O[C]; | 0.0 | 0.7 | #DIV/0! | 2.50E-02 |
| P02761 | PEDGEYFVEYDGGNTFTILK | 5,Unknown_216[E]; | 0.0 | 1.3 | #DIV/0! | 2.50E-02 |
| P02761 | LNGDWFSIVVASNKR | 5,Delta_O(4)[W]; | 0.0 | 0.7 | #DIV/0! | 2.50E-02 |
| P02761 | IEENGSMRVFMQHIDVLENSLGFK | 4,Asn->His[N]; | 0.0 | 1.3 | #DIV/0! | 2.50E-02 |
| P02761 | FAKLCEAHGITR | 2,Carbonyl[A]; | 0.0 | 1.3 | #DIV/0! | 2.50E-02 |
| P02761 | GNLDVAKLNGDWFSIVVASNKR | 2,Deamidated[N];9,Deamidated[N]; | 0.0 | 1.3 | #DIV/0! | 2.50E-02 |
| P02761 | TKDLSSDIKEK | 0,Succinyl[AnyN-term]; | 0.0 | 0.7 | #DIV/0! | 2.50E-02 |
| P01836 | DGVLDSVTDQDSKDSTYSMSSTLSLTK | 9,Cation_Na[D];11,Cation_Na[D]; | 0.0 | 0.7 | #DIV/0! | 2.50E-02 |
| P01835 | DGVLDSVTDQDSKDSTYSMSSTLSLTK | 9,Cation_Na[D];11,Cation_Na[D]; | 0.0 | 0.7 | #DIV/0! | 2.50E-02 |
| P00787 | ENHCGIESEIVAGIPR | 4,maleimide[C]; | 0.0 | 2.7 | #DIV/0! | 2.50E-02 |
| P12346 | KPVTEFATCHLAQAPNHVVVSR | 5,Cation_Fe[III][E]; | 0.0 | 4.5 | #DIV/0! | 2.56E-02 |
| P09006 | LDVAETGTEAAAATGVK | 0,Tween20[AnyN-term]; | 0.0 | 5.0 | #DIV/0! | 2.56E-02 |
| P17475 | TDTSQQDQSPTYR | 0,C+12[AnyN-term]; | 0.0 | 4.3 | #DIV/0! | 2.58E-02 |
| P81828 | KCQTYPDEICAWVVVTTR | 7,Cation_Na[D];10,Carbamidomethyl[C]; | 0.0 | 6.8 | #DIV/0! | 2.59E-02 |
| Q5RKI7 | LQNAFDAEFPEVSR | 1,Xle->Glu[L]; | 0.0 | 3.5 | #DIV/0! | 2.59E-02 |
| P07632 | SLSGEHSIIGR | 0,ISD_z+2_ion[AnyN-term]; | 0.0 | 3.5 | #DIV/0! | 2.59E-02 |
| P02761 | AKLCEAHGITR | 0,PEITC[AnyN-term]; | 0.0 | 9.8 | #DIV/0! | 2.61E-02 |
| P02761 | GDWFSIVVASNKR | 5,Hep[S]; | 0.0 | 3.2 | #DIV/0! | 2.61E-02 |
| P02770 | DNIPADLPSIAADFVEDK | 0,Ethylphosphate[AnyN-term]; | 0.0 | 5.5 | #DIV/0! | 2.64E-02 |
| P07647 | GSICLASGWGMTNPSEMK | 4,QAT[C]; | 0.0 | 4.7 | #DIV/0! | 2.64E-02 |
| P02761 | NGDWFSIVVASNK | 0,Diethylphosphate[AnyN-term]; | 0.0 | 4.7 | #DIV/0! | 2.64E-02 |
| P02761 | LCEAHGITR | 0,Methylthio[AnyN-term]; | 0.0 | 2.8 | #DIV/0! | 2.64E-02 |
| P02770 | MGDFAQFVDK | 0,C+12[AnyN-term]; | 0.0 | 3.0 | #DIV/0! | 2.67E-02 |
| P31211 | FSGNTKDVPLTLTMVHK | 4,Deamidated_18O(1)[N](Delta_H(1)N(-1)18O(1)[N]); | 0.0 | 2.5 | #DIV/0! | 2.68E-02 |
| P06911 | DDNGEALYNFR | 0,Triton[AnyN-term]; | 0.0 | 10.0 | #DIV/0! | 2.71E-02 |
| P68136 | YELPDGQVITIGNER | 1,Furan[Y]; | 0.0 | 2.2 | #DIV/0! | 2.74E-02 |
| P68035 | YELPDGQVITIGNER | 1,Furan[Y]; | 0.0 | 2.2 | #DIV/0! | 2.74E-02 |
| P63269 | YELPDGQVITIGNER | 1,Furan[Y]; | 0.0 | 2.2 | #DIV/0! | 2.74E-02 |
| P63259 | YELPDGQVITIGNER | 1,Furan[Y]; | 0.0 | 2.2 | #DIV/0! | 2.74E-02 |
| P62738 | YELPDGQVITIGNER | 1,Furan[Y]; | 0.0 | 2.2 | #DIV/0! | 2.74E-02 |
| P60711 | YELPDGQVITIGNER | 1,Furan[Y]; | 0.0 | 2.2 | #DIV/0! | 2.74E-02 |
| P17475 | VFNNDADLSGITEDAPLK | 0,Dimethyl_2H(4)[AnyN-term]; | 0.0 | 4.3 | #DIV/0! | 2.74E-02 |
| P12346 | TTIFEVLPQK | 1,Decanoyl[T]; | 0.0 | 2.2 | #DIV/0! | 2.74E-02 |
| P05545 | EKTRALYQAEAFVADFK | 4,Arg->GluSA[R]; | 0.0 | 2.2 | #DIV/0! | 2.74E-02 |
| P02770 | RPCFSALTVDETYVPK | 6,Ala->His[A]; | 0.0 | 2.2 | #DIV/0! | 2.74E-02 |
| P02761 | VMFHLINFK | 0,mTRAQ_13C(6)15N(2)[AnyN-term]; | 0.0 | 4.3 | #DIV/0! | 2.74E-02 |
| Q64319 | MPEGKWPNWMIGGPETSR | 5,BHTOH[K]; | 0.0 | 10.0 | #DIV/0! | 2.74E-02 |
| P22282 | PDYIEQNIHAVR | 0,BITC[AnyN-term]; | 0.0 | 3.3 | #DIV/0! | 2.77E-02 |
| P07647 | AYDHNNDLMLLHLSKPADITGGVK | 9,Met->AspSA[M]; | 0.0 | 3.3 | #DIV/0! | 2.77E-02 |
| P02770 | NCFATEGPNLVAR | 1,Deamidated_18O(1)[N](Delta_H(1)N(-1)18O(1)[N]); | 0.0 | 3.3 | #DIV/0! | 2.77E-02 |
| P02761 | MRVFMQHIDVLENSLGFK | 1,Met->Thr[M]; | 0.0 | 3.3 | #DIV/0! | 2.77E-02 |
| Q63041 | DQSVLLLKPEAK | 0,Unknown_234[AnyN-term]; | 0.0 | 2.3 | #DIV/0! | 2.78E-02 |
| Q0PMD2 | VLPGGDTYMHEGFER | 0,C+12[AnyN-term]; | 0.0 | 2.3 | #DIV/0! | 2.78E-02 |
| P02770 | DDNPNLPPFQRPEAEAMCTSFQENPTSFLGHYLHEVAR | 18,Carbamidomethyl[C];24,Deamidated[N]; | 0.0 | 2.3 | #DIV/0! | 2.78E-02 |
| P02761 | MQHIDVLENSLGFK | 3,His->Phe[H]; | 0.0 | 2.3 | #DIV/0! | 2.78E-02 |
| P02761 | KLCEAHGITR | 0,C+12[AnyN-term]; | 0.0 | 3.8 | #DIV/0! | 2.78E-02 |
| P20059 | ECHRGECQSEGVLFFQGNR | 7,MDCC[C]; | 0.0 | 7.2 | #DIV/0! | 2.82E-02 |
| P02761 | GETFQLMVLYGR | 3,Dimethylphosphothione[T]; | 0.0 | 5.7 | #DIV/0! | 2.83E-02 |
| P02761 | LCEAHGITRDNIIDLTK | 2,Carbamidomethyl[C];14,Cation_Na[D]; | 0.0 | 6.0 | #DIV/0! | 2.84E-02 |
| P07522 | LFWTDTGMSPR | 2,Phe->Lys[F]; | 0.0 | 1.8 | #DIV/0! | 2.84E-02 |
| P02770 | TCVADENAENCDKSIHTLFGDK | 2,Carbamidomethyl[C];11,Carbamidomethyl[C]; | 0.0 | 1.8 | #DIV/0! | 2.84E-02 |
| P02770 | GDFAQFVDK | 0,DiLeu4plex118[AnyN-term]; | 0.0 | 1.8 | #DIV/0! | 2.84E-02 |
| P00787 | FSEDINLPESFDAR | 0,Galactosyl[AnyN-term]; | 0.0 | 1.8 | #DIV/0! | 2.84E-02 |
| P02761 | VAKLNGDWFSIVVASNKR | 5,PhosphoHex(2)[N]; | 0.0 | 3.5 | #DIV/0! | 2.84E-02 |
| P09006 | AVLDVAETGTEAAAATGVK | 4,Cation_Na[D]; | 0.0 | 4.0 | #DIV/0! | 2.88E-02 |
| P02782 | SQICELVAHETISFLMK | 0,Amidine[AnyN-term]; | 0.0 | 7.5 | #DIV/0! | 2.90E-02 |
| P02761 | EYDGGNTFTILK | 0,Succinyl_2H(4)[AnyN-term]; | 0.0 | 4.2 | #DIV/0! | 2.92E-02 |
| P02761 | PEDGEYFVEYDGGNTFTILK | 10,DiART6plex115[Y]; | 0.0 | 5.3 | #DIV/0! | 2.93E-02 |
| P00758 | NLYEDEPFAQHR | 0,Succinyl[AnyN-term]; | 0.0 | 2.7 | #DIV/0! | 2.93E-02 |
| P02761 | VFMQHIDVLENSLGFK | 0,Methyl[AnyN-term]; | 0.0 | 18.5 | #DIV/0! | 2.97E-02 |
| P02761 | KTPEDGEYFVEYDGGNTFTILK | 2,Thr->Ala[T]; | 0.0 | 3.8 | #DIV/0! | 2.99E-02 |
| Q6IRK9 | LQQDGLENVHLEQVR | 4,Cation_Ca[II][D]; | 0.0 | 1.5 | #DIV/0! | 3.01E-02 |
| P15684 | NAGAMENWGLVTYR | 0,mTRAQ[AnyN-term]; | 0.0 | 1.5 | #DIV/0! | 3.01E-02 |
| P13432 | PPTQLHSTEQANTK | 0,Phenylisocyanate_2H(5)[AnyN-term]; | 0.0 | 3.0 | #DIV/0! | 3.01E-02 |
| P02761 | LNGDWFSIVVASNK | 0,ISD_z+2_ion[AnyN-term]; | 0.0 | 1.5 | #DIV/0! | 3.01E-02 |
| P02761 | KDLSSDIKEK | 0,Ethylphosphate[AnyN-term]; | 0.0 | 1.5 | #DIV/0! | 3.01E-02 |
| P02761 | KLCEAHGITR | 2,Carbonyl[L]; | 0.0 | 1.5 | #DIV/0! | 3.01E-02 |
| P02761 | GNLDVAKLNGDWFSIVVASNKR | 2,Deamidated[N];20,Deamidated[N]; | 0.0 | 1.5 | #DIV/0! | 3.01E-02 |
| P00758 | NNLYEDEPFAQHR | 6,Cation_Ca[II][D]; | 0.0 | 3.0 | #DIV/0! | 3.01E-02 |
| P83121 | ICAWVVVTTR | 2,IBTP[C]; | 0.0 | 2.8 | #DIV/0! | 3.02E-02 |
| P81828 | ICAWVVVTTR | 2,IBTP[C]; | 0.0 | 2.8 | #DIV/0! | 3.02E-02 |
| P81827 | ICAWVVVTTR | 2,IBTP[C]; | 0.0 | 2.8 | #DIV/0! | 3.02E-02 |
| P02761 | MQHIDVLENSLGFK | 14,Lys->His[K]; | 0.0 | 2.8 | #DIV/0! | 3.02E-02 |
| P02761 | AKLCEAHGITR | 0,Unknown_306[AnyN-term]; | 0.0 | 2.8 | #DIV/0! | 3.02E-02 |
| P02761 | ITRDNIIDLTK | 0,3sulfo[AnyN-term]; | 0.0 | 5.5 | #DIV/0! | 3.04E-02 |
| Q9JHB9 | GSGCSILDEVIR | 0,Carbamyl[AnyN-term]; | 0.0 | 3.3 | #DIV/0! | 3.05E-02 |
| P24090 | VEFVIAATDCTGQEVTDPAK | 10,Carbamidomethyl[C]; | 0.0 | 1.7 | #DIV/0! | 3.05E-02 |
| P22282 | LDNCPFEEQTEQLK | 4,Propionamide_2H(3)[C]; | 0.0 | 5.0 | #DIV/0! | 3.05E-02 |
| P02780 | GSGCSILDEVIR | 0,Carbamyl[AnyN-term]; | 0.0 | 3.3 | #DIV/0! | 3.05E-02 |
| P02770 | KQTALAELVK | 1,Carbamyl[K]; | 0.0 | 1.7 | #DIV/0! | 3.05E-02 |
| P02761 | KLCEAHGITR | 0,PyMIC[AnyN-term]; | 0.0 | 1.7 | #DIV/0! | 3.05E-02 |
| P02761 | LCEAHGITR | 0,Iodoacetanilide_13C(6)[AnyN-term]; | 0.0 | 1.7 | #DIV/0! | 3.05E-02 |
| P02761 | INFKNGETFQLMVLYGR | 4,CresylSaligeninPhosphate[K]; | 0.0 | 3.3 | #DIV/0! | 3.05E-02 |
| P81827 | RICQTYPDEICAWVVVTTR | 3,Carbamidomethyl[C]; | 0.0 | 3.7 | #DIV/0! | 3.07E-02 |
| Q9JHB9 | GSGCSILDEVIR | 0,Galactosyl[AnyN-term]; | 0.0 | 4.8 | #DIV/0! | 3.07E-02 |
| P02780 | GSGCSILDEVIR | 0,Galactosyl[AnyN-term]; | 0.0 | 4.8 | #DIV/0! | 3.07E-02 |
| P01836 | SKDSTYSMSSTLSLTK | 0,Unknown_216[AnyN-term]; | 0.0 | 3.5 | #DIV/0! | 3.09E-02 |
| P01835 | SKDSTYSMSSTLSLTK | 0,Unknown_216[AnyN-term]; | 0.0 | 3.5 | #DIV/0! | 3.09E-02 |
| P01836 | TDQDSKDSTYSMSSTLSLTK | 6,Methyl_2H(3)+Acetyl_2H(3)[K]; | 0.0 | 7.7 | #DIV/0! | 3.15E-02 |
| P01835 | TDQDSKDSTYSMSSTLSLTK | 6,Methyl_2H(3)+Acetyl_2H(3)[K]; | 0.0 | 7.7 | #DIV/0! | 3.15E-02 |
| P83121 | QTYPDEICAWVVVTTR | 8,NEMsulfur[C]; | 0.0 | 4.5 | #DIV/0! | 3.16E-02 |
| P81828 | QTYPDEICAWVVVTTR | 8,NEMsulfur[C]; | 0.0 | 4.5 | #DIV/0! | 3.16E-02 |
| P81827 | QTYPDEICAWVVVTTR | 8,NEMsulfur[C]; | 0.0 | 4.5 | #DIV/0! | 3.16E-02 |
| P02761 | VFMQHIDVLENSLGFK | 0,Hex[AnyN-term]; | 0.0 | 4.5 | #DIV/0! | 3.16E-02 |
| P07522 | LSVIEMADLDGSK | 1,Xle->Gln[L]; | 0.0 | 17.0 | #DIV/0! | 3.18E-02 |
| Q64240 | CIQFIYGGCK | 0,Methyl_2H(2)[AnyN-term];1,Carbamidomethyl[C];9,Carbamidomethyl[C]; | 0.0 | 2.5 | #DIV/0! | 3.18E-02 |
| P06866 | SPVGVQPILNK | 1,PhosphoHex[S]; | 0.0 | 5.0 | #DIV/0! | 3.18E-02 |
| P02761 | NGDWFSIVVASNKR | 4,Nitro[W]; | 0.0 | 2.5 | #DIV/0! | 3.18E-02 |
| P80067 | ETMTGWVHDVLGR | 0,Brij35[AnyN-term]; | 0.0 | 4.7 | #DIV/0! | 3.21E-02 |
| P00787 | ILRGENHCGIESEIVAGIPR | 8,Carbamidomethyl[C]; | 0.0 | 7.0 | #DIV/0! | 3.22E-02 |
| P36373 | SSNNYQVWLGR | 1,BEMAD_ST_2H(6)[S]; | 0.0 | 3.2 | #DIV/0! | 3.23E-02 |
| P02770 | MCTSFQENPTSFLGHYLHEVAR | 2,Cys->Trp[C]; | 0.0 | 3.2 | #DIV/0! | 3.23E-02 |
| O35568 | SGNENGEFYLR | 1,Ser->Pro[S]; | 0.0 | 4.3 | #DIV/0! | 3.24E-02 |
| P31211 | VPLTLTMVHK | 0,Phenylisocyanate_2H(5)[AnyN-term]; | 0.0 | 2.0 | #DIV/0! | 3.27E-02 |
| P02761 | MQHIDVLENSLGFK | 1,Oxidation[M];5,Cation_Na[D]; | 0.0 | 2.0 | #DIV/0! | 3.27E-02 |
| P00689 | GAGGASILTFWDAR | 0,DiART6plex118[AnyN-term]; | 0.0 | 2.0 | #DIV/0! | 3.27E-02 |
| P01836 | DGVLDSVTDQDSK | 0,Methyl_2H(2)[AnyN-term]; | 0.0 | 26.0 | #DIV/0! | 3.31E-02 |
| P01835 | DGVLDSVTDQDSK | 0,Methyl_2H(2)[AnyN-term]; | 0.0 | 26.0 | #DIV/0! | 3.31E-02 |
| P02761 | KIEENGSMRVFMQHIDVLENSLGFK | 5,Deamidated[N]; | 0.0 | 6.5 | #DIV/0! | 3.33E-02 |
| P07647 | AYDHNNDLMLLHLSK | 5,Ammonia-loss[N]; | 0.0 | 3.3 | #DIV/0! | 3.34E-02 |
| Q6AYD4 | SPRSKPTAQYQWERLAPSSQVFFGPALDTVR | 7,Phosphopropargyl[T]; | 0.0 | 3.5 | #DIV/0! | 3.35E-02 |
| P36373 | KPGDDHSNDLMLLHLSQPADITDGVK | 13,Dioxidation[L]; | 0.0 | 1.2 | #DIV/0! | 3.35E-02 |
| P22283 | NKLNDVYDIFK | 2,Glucosylgalactosyl[K]; | 0.0 | 4.7 | #DIV/0! | 3.35E-02 |
| P16573 | DQQFNPIQTSVQFR | 0,CLIP_TRAQ_4[AnyN-term]; | 0.0 | 1.2 | #DIV/0! | 3.35E-02 |
| P07647 | LHLSKPADITGGVK | 4,Pent(1)HexNAc(1)[S]; | 0.0 | 1.2 | #DIV/0! | 3.35E-02 |
| P07314 | DIQEAGGIMTVEDLNNYR | 3,Deamidated[Q]; | 0.0 | 1.2 | #DIV/0! | 3.35E-02 |
| P06760 | FEKQWYRQPLRESGPTLDMPVPSSFNDITQEAELR | 0,Unknown_250[AnyN-term]; | 0.0 | 1.2 | #DIV/0! | 3.35E-02 |
| P05545 | INTGSALFIDKEQPILSEFQEK | 5,BEMAD_ST_2H(6)[S]; | 0.0 | 1.2 | #DIV/0! | 3.35E-02 |
| P05544 | INTGSALFIDKEQPILSEFQEK | 5,BEMAD_ST_2H(6)[S]; | 0.0 | 1.2 | #DIV/0! | 3.35E-02 |
| P04903 | IQSPEDLEK | 0,CAF[AnyN-term]; | 0.0 | 1.2 | #DIV/0! | 3.35E-02 |
| P02761 | GNLDVAKLNGDWFSIVVASNKR | 5,Val->Cys[V]; | 0.0 | 2.3 | #DIV/0! | 3.35E-02 |
| P02761 | FAKLCEAHGITR | 0,GIST-Quat_2H(9)[AnyN-term]; | 0.0 | 1.2 | #DIV/0! | 3.35E-02 |
| P02761 | GNLDVAKLNGDWFSIVVASNKR | 2,Deamidated[N]; | 0.0 | 1.2 | #DIV/0! | 3.35E-02 |
| P01836 | IDGSEQRDGVLDSVTDQDSK | 0,Carboxymethyl[AnyN-term]; | 0.0 | 2.3 | #DIV/0! | 3.35E-02 |
| P00884 | YTPEQVAMATVTALHR | 4,Glu->His[E]; | 0.0 | 1.2 | #DIV/0! | 3.35E-02 |
| P00502 | IQSPEDLEK | 0,CAF[AnyN-term]; | 0.0 | 1.2 | #DIV/0! | 3.35E-02 |
| P22283 | LVNCPFEEQTEQLKR | 3,Deamidated[N];4,Carbamidomethyl[C]; | 0.0 | 13.3 | #DIV/0! | 3.35E-02 |
| P05371 | LTQQYNELLHSLQSK | 1,Carbonyl[L]; | 0.0 | 3.0 | #DIV/0! | 3.36E-02 |
| P02761 | GNLDVAKLNGDWFSIVVASNKR | 8,trifluoro[L]; | 0.0 | 3.0 | #DIV/0! | 3.36E-02 |
| Q9JJ19 | VEKETHQQVVSR | 0,Succinyl[AnyN-term]; | 0.0 | 2.7 | #DIV/0! | 3.37E-02 |
| Q64319 | EKLDYITALNIK | 2,Phosphoadenosine[K]; | 0.0 | 5.3 | #DIV/0! | 3.37E-02 |
| P83121 | QTYPDEICAWVVVTTR | 8,CarbamidomethylDTT[C]; | 0.0 | 14.5 | #DIV/0! | 3.38E-02 |
| P81828 | QTYPDEICAWVVVTTR | 8,CarbamidomethylDTT[C]; | 0.0 | 14.5 | #DIV/0! | 3.38E-02 |
| P81827 | QTYPDEICAWVVVTTR | 8,CarbamidomethylDTT[C]; | 0.0 | 14.5 | #DIV/0! | 3.38E-02 |
| P42854 | AYGSYCYALFSVSK | 6,Ethanolyl[C]; | 0.0 | 7.8 | #DIV/0! | 3.38E-02 |
| P83121 | DEICAWVVVTTR | 0,Methyl_2H(2)[AnyN-term];4,Carbamidomethyl[C]; | 0.0 | 18.8 | #DIV/0! | 3.40E-02 |
| P81828 | DEICAWVVVTTR | 0,Methyl_2H(2)[AnyN-term];4,Carbamidomethyl[C]; | 0.0 | 18.8 | #DIV/0! | 3.40E-02 |
| P81827 | DEICAWVVVTTR | 0,Methyl_2H(2)[AnyN-term];4,Carbamidomethyl[C]; | 0.0 | 18.8 | #DIV/0! | 3.40E-02 |
| P02761 | RGNLDVAKLNGDWFSIVVASNK | 3,Deamidated[N]; | 0.0 | 2.2 | #DIV/0! | 3.41E-02 |
| P02770 | AEAMCTSFQENPTSFLGHYLHEVAR | 5,BHT[C]; | 0.0 | 8.5 | #DIV/0! | 3.46E-02 |
| Q499T2 | LNKVEACLLDKLEK | 7,Carbamidomethyl[C]; | 0.0 | 9.3 | #DIV/0! | 3.46E-02 |
| P81828 | LQCFRCTSFDSTGFCHVGR | 3,Carbamidomethyl[C];6,Carbamidomethyl[C];15,Carbamidomethyl[C]; | 0.0 | 4.8 | #DIV/0! | 3.48E-02 |
| P07151 | LKNGKKIPNIEMSDLSFSK | 0,Phosphogluconoylation[AnyN-term]; | 0.0 | 10.5 | #DIV/0! | 3.50E-02 |
| P23680 | SIVLGQEQDTYGGGFDK | 0,ESP[AnyN-term]; | 0.0 | 9.2 | #DIV/0! | 3.51E-02 |
| P26051 | FETCRYGFIEGHVVIPR | 4,SulfurDioxide[C]; | 0.0 | 11.7 | #DIV/0! | 3.54E-02 |
| Q64240 | AFAELWAFDAAQGK | 2,Phe->CamCys[F]; | 0.0 | 3.7 | #DIV/0! | 3.54E-02 |
| P20059 | NPVTGEVPPR | 1,Hex[N]; | 0.0 | 12.7 | #DIV/0! | 3.55E-02 |
| P02761 | PEDGEYFVEYDGGNTFTILK | 15,PhosphoHexNAc[T]; | 0.0 | 6.7 | #DIV/0! | 3.55E-02 |
| P02761 | EKFAKLCEAHGITR | 0,Ethylphosphate[AnyN-term]; | 0.0 | 5.5 | #DIV/0! | 3.57E-02 |
| Q9JHB9 | GSGCSILDEVIR | 4,GG[C](Dicarbamidomethyl[C]); | 0.0 | 12.2 | #DIV/0! | 3.58E-02 |
| P02780 | GSGCSILDEVIR | 4,GG[C](Dicarbamidomethyl[C]); | 0.0 | 12.2 | #DIV/0! | 3.58E-02 |
| P24090 | QYGFCK | 5,Carbamidomethyl[C]; | 0.0 | 5.3 | #DIV/0! | 3.60E-02 |
| P02770 | QEPERNECFLQHKDDNPNLPPFQRPEAEAMCTSFQENPTSFLGHYLHEVAR | 31,GG[C](Dicarbamidomethyl[C]); | 0.0 | 10.2 | #DIV/0! | 3.60E-02 |
| P27590 | RDWMSIVTPAR | 0,AccQTag[AnyN-term]; | 0.0 | 3.5 | #DIV/0! | 3.62E-02 |
| P07151 | KKIPNIEMSDLSFSK | 1,Lys->CamCys[K]; | 0.0 | 3.3 | #DIV/0! | 3.63E-02 |
| P07522 | WCDTKLSVIEMADLDGSK | 2,Propionamide_2H(3)[C]; | 0.0 | 10.2 | #DIV/0! | 3.63E-02 |
| P02761 | KLCEAHGITR | 0,Amidine[AnyN-term]; | 0.0 | 7.5 | #DIV/0! | 3.63E-02 |
| P08723 | IFLTVIDLIK | 7,Cation_Na[D]; | 0.0 | 3.8 | #DIV/0! | 3.64E-02 |
| P27590 | PGLDCLPQGPSGK | 5,Carbamidomethyl[C]; | 0.0 | 3.8 | #DIV/0! | 3.64E-02 |
| P02761 | DWFSIVVASNK | 0,Unknown_306[AnyN-term]; | 0.0 | 8.3 | #DIV/0! | 3.65E-02 |
| P02761 | DLSSDIKEK | 0,Carbamyl[AnyN-term]; | 0.0 | 8.7 | #DIV/0! | 3.67E-02 |
| P02761 | KLCEAHGITR | 1,Lys->AminoadipicAcid[K]; | 0.0 | 11.7 | #DIV/0! | 3.69E-02 |
| P02761 | TPEDGEYFVEYDGGNTFTILK | 13,Gly->Arg[G]; | 0.0 | 2.5 | #DIV/0! | 3.69E-02 |
| P02761 | QHIDVLENSLGFK | 0,Gln->pyro-Glu[AnyN-termQ];4,Cation_Na[D]; | 0.0 | 2.5 | #DIV/0! | 3.69E-02 |
| P02761 | DNIIDLTKTDR | 8,Formyl[K]; | 0.0 | 2.5 | #DIV/0! | 3.69E-02 |
| P02761 | LNGDWFSIVVASNKR | 1,Oxidation[L]; | 0.0 | 17.0 | #DIV/0! | 3.73E-02 |
| Q01177 | TPANFPDAGLEMNYCR | 7,Cation_Na[D];15,Carbamidomethyl[C]; | 0.0 | 8.2 | #DIV/0! | 3.74E-02 |
| P01836 | DSTYSMSSTLSLTK | 2,Ser->Cys[S]; | 0.0 | 8.8 | #DIV/0! | 3.76E-02 |
| P01835 | DSTYSMSSTLSLTK | 2,Ser->Cys[S]; | 0.0 | 8.8 | #DIV/0! | 3.76E-02 |
| Q6MG71 | MDSLQKGLCPSFLLPSTPALGR | 0,3sulfo[AnyN-term]; | 0.0 | 1.8 | #DIV/0! | 3.79E-02 |
| P02761 | AKLCEAHGITR | 0,Diisopropylphosphate[AnyN-term]; | 0.0 | 3.7 | #DIV/0! | 3.79E-02 |
| P02761 | NGETFQLMVLYGR | 5,Phe->Arg[F]; | 0.0 | 6.3 | #DIV/0! | 3.79E-02 |
| P07647 | LSHDLQCVNIHLLSNEK | 4,Cation_Na[D];7,Carbamidomethyl[C]; | 0.0 | 6.0 | #DIV/0! | 3.81E-02 |
| P81827 | RICQTYPDEICAWVVVTTR | 10,Xle->Asp[I]; | 0.0 | 2.8 | #DIV/0! | 3.82E-02 |
| P12020 | TVSPSGSDLLR | 1,Thr->Xle[T]; | 0.0 | 2.7 | #DIV/0! | 3.83E-02 |
| P10959 | PEPAEPWSFVK | 0,NIC[AnyN-term](ICPL[AnyN-term]); | 0.0 | 2.7 | #DIV/0! | 3.83E-02 |
| P07522 | WCDTKLSVIEMADLDGSK | 0,Formyl[AnyN-term]; | 0.0 | 2.7 | #DIV/0! | 3.83E-02 |
| P02761 | VFMQHIDVLENSLGFK | 6,Xle->Trp[I]; | 0.0 | 14.8 | #DIV/0! | 3.84E-02 |
| P83121 | ICQTYPDEICAWVVVTTR | 0,Methyl_2H(2)[AnyN-term];7,Cation_Na[D];10,Carbamidomethyl[C]; | 0.0 | 6.7 | #DIV/0! | 3.85E-02 |
| P81827 | ICQTYPDEICAWVVVTTR | 0,Methyl_2H(2)[AnyN-term];7,Cation_Na[D];10,Carbamidomethyl[C]; | 0.0 | 6.7 | #DIV/0! | 3.85E-02 |
| P83121 | CAWVVVTTR | 1,Methylthio[C]; | 0.0 | 25.3 | #DIV/0! | 3.86E-02 |
| P81828 | CAWVVVTTR | 1,Methylthio[C]; | 0.0 | 25.3 | #DIV/0! | 3.86E-02 |
| P81827 | CAWVVVTTR | 1,Methylthio[C]; | 0.0 | 25.3 | #DIV/0! | 3.86E-02 |
| P12020 | VSPSGSDLLR | 0,Ethylphosphate[AnyN-term]; | 0.0 | 3.2 | #DIV/0! | 3.88E-02 |
| P07861 | DGDLVDWWTQQSANNFK | 0,Methyl_2H(2)[AnyN-term]; | 0.0 | 3.2 | #DIV/0! | 3.88E-02 |
| P02761 | LCEAHGITR | 0,CLIP_TRAQ_2[AnyN-term]; | 0.0 | 4.0 | #DIV/0! | 3.88E-02 |
| Q64240 | ELWAFDAAQGK | 0,Unknown_302[AnyN-term]; | 0.0 | 3.3 | #DIV/0! | 3.92E-02 |
| P02761 | DLSSDIKEKFAKLCEAHGITR | 9,Lys->Thr[K]; | 0.0 | 3.3 | #DIV/0! | 3.92E-02 |
| P02780 | VKPYVQDHFTEKAVK | 0,Hex[AnyN-term]; | 0.0 | 4.3 | #DIV/0! | 3.93E-02 |
| Q64240 | AFAELWAFDAAQGK | 0,Methyl[AnyN-term]; | 0.0 | 2.3 | #DIV/0! | 3.95E-02 |
| P48199 | SFSIFSYATK | 2,Phe->His[F]; | 0.0 | 2.3 | #DIV/0! | 3.95E-02 |
| P31977 | KKAPDFVFYAPR | 0,LG-pyrrole[AnyN-term]; | 0.0 | 4.7 | #DIV/0! | 3.95E-02 |
| P22282 | TPGETMYYISLPGSVR | 5,Didehydro[T]; | 0.0 | 2.3 | #DIV/0! | 3.95E-02 |
| P22282 | VYPDYIEQNIHAVR | 1,Val->Cys[V]; | 0.0 | 2.3 | #DIV/0! | 3.95E-02 |
| O35763 | KKAPDFVFYAPR | 0,LG-pyrrole[AnyN-term]; | 0.0 | 4.7 | #DIV/0! | 3.95E-02 |
| Q64240 | YCGVPGDGYEELTR | 2,CarbamidomethylDTT[C]; | 0.0 | 5.0 | #DIV/0! | 3.95E-02 |
| P98158 | YWVDSRYDYIETVTYDGIQR | 4,Cation_Ca[II][D]; | 0.0 | 4.2 | #DIV/0! | 4.03E-02 |
| Q9R0T4 | DINDNAPIFNPSTYQGQVLENEVGAR | 13,Thr->Tyr[T]; | 0.0 | 1.0 | #DIV/0! | 4.09E-02 |
| Q01177 | WSDTSPHVPK | 0,ISD_z+2_ion[AnyN-term]; | 0.0 | 1.0 | #DIV/0! | 4.09E-02 |
| P82450 | VVWYQGENNANYNR | 0,Propyl[AnyN-term]; | 0.0 | 2.0 | #DIV/0! | 4.09E-02 |
| P21704 | LTNQMAEAISDHYPVEVTLR | 3,Deamidated[N];5,Oxidation[M]; | 0.0 | 1.0 | #DIV/0! | 4.09E-02 |
| P20059 | FLRFNPVTGEVPPR | 3,phosphoRibosyl[R]; | 0.0 | 1.0 | #DIV/0! | 4.09E-02 |
| P05371 | LDAMQDSFTRASGIIDTLFQDR | 3,HNE[A]; | 0.0 | 1.0 | #DIV/0! | 4.09E-02 |
| P02770 | CCAEGDPPACYGTVLAEFQPLVEEPK | 10,4-ONE[C]; | 0.0 | 1.0 | #DIV/0! | 4.09E-02 |
| P02770 | DDNPNLPPFQRPEAEAMCTSFQENPTSFLGHYLHEVAR | 5,Deamidated[N];18,Carbamidomethyl[C]; | 0.0 | 1.0 | #DIV/0! | 4.09E-02 |
| P02761 | EENGSMRVFMQHIDVLENSLGFK | 5,PhosphoHex[S]; | 0.0 | 2.0 | #DIV/0! | 4.09E-02 |
| P02761 | FAKLCEAHGITR | 3,Lys->AminoadipicAcid[K]; | 0.0 | 1.0 | #DIV/0! | 4.09E-02 |
| P02761 | KLCEAHGITR | 1,Xlink_DFDNB[K]; | 0.0 | 2.0 | #DIV/0! | 4.09E-02 |
| P02650 | LEEQLGPVAEETR | 0,Iodoacetanilide[AnyN-term]; | 0.0 | 1.0 | #DIV/0! | 4.09E-02 |
| P02650 | SKMEEQTQQIR | 3,Oxidation[M]; | 0.0 | 1.0 | #DIV/0! | 4.09E-02 |
| P01048 | NAELESGNQFVLYR | 0,Galactosyl[AnyN-term]; | 0.0 | 1.0 | #DIV/0! | 4.09E-02 |
| P01015 | DSGALHFLGR | 1,Cation_Li[D]; | 0.0 | 2.0 | #DIV/0! | 4.09E-02 |
| P00787 | NGPVEGAFTVFSDFLTYK | 0,Guanidinyl[AnyN-term]; | 0.0 | 1.0 | #DIV/0! | 4.09E-02 |
| P83121 | EICAWVVVTTR | 3,2-succinyl[C]; | 0.0 | 7.5 | #DIV/0! | 4.10E-02 |
| P81828 | EICAWVVVTTR | 3,2-succinyl[C]; | 0.0 | 7.5 | #DIV/0! | 4.10E-02 |
| P81827 | EICAWVVVTTR | 3,2-succinyl[C]; | 0.0 | 7.5 | #DIV/0! | 4.10E-02 |
| Q9EQV9 | HGLAFQSGHVLSALPR | 1,PhosphoUridine[H]; | 0.0 | 2.2 | #DIV/0! | 4.11E-02 |
| Q68FP1 | QSRVVQGKEPAHLMSLFGGK | 0,Unknown_420[AnyN-term]; | 0.0 | 2.2 | #DIV/0! | 4.11E-02 |
| P15684 | SQTAIKLPASVSTIMDR | 7,Xle->Ser[L]; | 0.0 | 2.2 | #DIV/0! | 4.11E-02 |
| P07151 | IQKTPQIQVYSR | 0,Carbamyl[AnyN-term]; | 0.0 | 2.2 | #DIV/0! | 4.11E-02 |
| P02770 | SQRFPNAEFAEITK | 0,Dimethyl_2H(4)[AnyN-term]; | 0.0 | 2.2 | #DIV/0! | 4.11E-02 |
| P02761 | NGDWFSIVVASNKR | 0,Acetyl_2H(3)[AnyN-term]; | 0.0 | 2.2 | #DIV/0! | 4.11E-02 |
| P07632 | PVQGVIHFEQK | 0,SPITC_13C(6)[AnyN-term]; | 0.0 | 6.2 | #DIV/0! | 4.12E-02 |
| Q5GRG2 | HGLQVVADVCK | 10,Carbamidomethyl[C]; | 0.0 | 5.7 | #DIV/0! | 4.13E-02 |
| P17475 | EDAQETDTSQQDQSPTYR | 1,Cation_Fe[III][E]; | 0.0 | 4.8 | #DIV/0! | 4.19E-02 |
| P02761 | DVLENSLGFK | 0,NIC[AnyN-term](ICPL[AnyN-term]); | 0.0 | 3.2 | #DIV/0! | 4.21E-02 |
| Q9JHB9 | SGCSILDEVIR | 1,O-Et-N-diMePhospho[S]; | 0.0 | 0.8 | #DIV/0! | 4.22E-02 |
| Q68FP1 | HGMDDDGTGQK | 0,Diethylphosphate[AnyN-term]; | 0.0 | 0.8 | #DIV/0! | 4.22E-02 |
| Q63041 | IKEDGTGIELTGTGSCEIANTLSK | 13,Methylphosphonate[T]; | 0.0 | 0.8 | #DIV/0! | 4.22E-02 |
| Q5XI43 | HDRQRVVHWDLSGGPGSQGR | 6,Val->Tyr[V]; | 0.0 | 1.7 | #DIV/0! | 4.22E-02 |
| P97546 | SENKNEGQDAMMYCK | 12,Oxidation[M];14,Carbamidomethyl[C]; | 0.0 | 0.8 | #DIV/0! | 4.22E-02 |
| P36953 | ADWCQPLKEDLQNKR | 4,Trioxidation[C]; | 0.0 | 1.7 | #DIV/0! | 4.22E-02 |
| P20059 | FNPVTGEVPPR | 1,Oxidation[F](Phe->Tyr[F]); | 0.0 | 0.8 | #DIV/0! | 4.22E-02 |
| P20059 | NPVTGEVPPR | 0,Unknown_162[AnyN-term]; | 0.0 | 2.5 | #DIV/0! | 4.22E-02 |
| P07632 | GNVAAGKDGVANVSIEDR | 8,Dehydrated[D]; | 0.0 | 0.8 | #DIV/0! | 4.22E-02 |
| P06866 | NATAKDIAPTLTLYVGK | 5,3-phosphoglyceryl[K]; | 0.0 | 0.8 | #DIV/0! | 4.22E-02 |
| P05371 | SGIIDTLFQDR | 1,AEC-MAEC_2H(4)[S]; | 0.0 | 1.7 | #DIV/0! | 4.22E-02 |
| P02780 | SGCSILDEVIR | 1,O-Et-N-diMePhospho[S]; | 0.0 | 0.8 | #DIV/0! | 4.22E-02 |
| P02780 | LVKPYVQDHFTEK | 1,Xle->Phe[L]; | 0.0 | 0.8 | #DIV/0! | 4.22E-02 |
| P02770 | QEPERNECFLQHKDDNPNLPPFQRPEAEAMCTSFQENPTSFLGHYLHEVAR | 0,ICPL_13C(6)2H(4)[AnyN-term]; | 0.0 | 1.7 | #DIV/0! | 4.22E-02 |
| P02770 | KCCAEGDPPACYGTVLAEFQPLVEEPK | 2,Nitrosyl[C]; | 0.0 | 0.8 | #DIV/0! | 4.22E-02 |
| P02770 | AELAKYMCENQATISSK | 2,Glu->His[E]; | 0.0 | 0.8 | #DIV/0! | 4.22E-02 |
| P02761 | DLSSDIKEKFAK | 9,Carbamyl[K]; | 0.0 | 0.8 | #DIV/0! | 4.22E-02 |
| P02761 | EDGEYFVEYDGGNTFTILK | 0,Phosphogluconoylation[AnyN-term]; | 0.0 | 0.8 | #DIV/0! | 4.22E-02 |
| P02761 | VFMQHIDVLENSLGFK | 5,His->Thr[H]; | 0.0 | 0.8 | #DIV/0! | 4.22E-02 |
| P02761 | ENGSMRVFMQHIDVLENSLGFK | 0,Galactosyl[AnyN-term]; | 0.0 | 0.8 | #DIV/0! | 4.22E-02 |
| P02761 | DLSSDIKEK | 0,ISD_z+2_ion[AnyN-term]; | 0.0 | 0.8 | #DIV/0! | 4.22E-02 |
| P01836 | DSTYSMSSTLSLTK | 0,Methyl_2H(2)[AnyN-term]; | 0.0 | 2.5 | #DIV/0! | 4.22E-02 |
| P01836 | SVKWKIDGSEQR | 0,Dansyl[AnyN-term]; | 0.0 | 0.8 | #DIV/0! | 4.22E-02 |
| P01835 | DSTYSMSSTLSLTK | 0,Methyl_2H(2)[AnyN-term]; | 0.0 | 2.5 | #DIV/0! | 4.22E-02 |
| P00787 | NHCGIESEIVAGIPR | 3,Dipyridyl[C]; | 0.0 | 0.8 | #DIV/0! | 4.22E-02 |
| P00786 | IMGEDSYPYIGK | 1,Xle->Tyr[I]; | 0.0 | 0.8 | #DIV/0! | 4.22E-02 |
| P48199 | VYVGRVFSPNVLNWR | 0,Dimethyl_2H(4)[AnyN-term]; | 0.0 | 2.8 | #DIV/0! | 4.24E-02 |
| P02781 | PPEAVEANLK | 1,OxProBiotinRed[P]; | 0.0 | 2.8 | #DIV/0! | 4.24E-02 |
| P02761 | MRVFMQHIDVLENSLGFK | 3,Val->Ser[V]; | 0.0 | 2.8 | #DIV/0! | 4.24E-02 |
| Q9JMI9 | YAIAVVLSFSR | 8,HexNAc(1)dHex(1)[S]; | 0.0 | 21.8 | #DIV/0! | 4.25E-02 |
| P07522 | LYWVDLER | 5,Cation_Na[D]; | 0.0 | 4.7 | #DIV/0! | 4.26E-02 |
| P02761 | MQHIDVLENSLGFK | 3,His->Glu[H]; | 0.0 | 6.2 | #DIV/0! | 4.29E-02 |
| Q6IE52 | VKDSARAHFSVMGDILSSAIK | 4,Dimethylphosphothione[S]; | 0.0 | 1.3 | #DIV/0! | 4.29E-02 |
| Q03626 | VKDSARAHFSVMGDILSSAIK | 4,Dimethylphosphothione[S]; | 0.0 | 1.3 | #DIV/0! | 4.29E-02 |
| P36374 | LKTPADITDGVK | 0,Diethylphosphate[AnyN-term]; | 0.0 | 1.3 | #DIV/0! | 4.29E-02 |
| P27590 | ERDWMSIVTPAR | 4,Delta_O(4)[W]; | 0.0 | 1.3 | #DIV/0! | 4.29E-02 |
| P22057 | QGHDTVQPNFQQDK | 3,His->Met[H]; | 0.0 | 1.3 | #DIV/0! | 4.29E-02 |
| P20059 | CQSEGVLFFQGNRK | 3,Pent(2)[S]; | 0.0 | 1.3 | #DIV/0! | 4.29E-02 |
| P20059 | NPLPAAHETVAK | 1,Methyl+Deamidated[N](Asn->Glu[N]); | 0.0 | 1.3 | #DIV/0! | 4.29E-02 |
| P14046 | VKDSARAHFSVMGDILSSAIK | 4,Dimethylphosphothione[S]; | 0.0 | 1.3 | #DIV/0! | 4.29E-02 |
| P07522 | YKTLLSRQMGMVFALDYDPVESK | 0,Dimethyl_2H(6)13C(2)[AnyN-term]; | 0.0 | 1.3 | #DIV/0! | 4.29E-02 |
| P05545 | EINTGSALFIDKEQPILSEFQEK | 0,Methyl[AnyN-term]; | 0.0 | 1.3 | #DIV/0! | 4.29E-02 |
| P05545 | IFSQQADLSR | 0,Diisopropylphosphate[AnyN-term]; | 0.0 | 1.3 | #DIV/0! | 4.29E-02 |
| P05544 | EINTGSALFIDKEQPILSEFQEK | 0,Methyl[AnyN-term]; | 0.0 | 1.3 | #DIV/0! | 4.29E-02 |
| P05544 | IAELFSDLEER | 3,Cation_Li[E]; | 0.0 | 1.3 | #DIV/0! | 4.29E-02 |
| P02780 | DYMKLVKPYVQDHFTEK | 2,Fluoro[Y]; | 0.0 | 1.3 | #DIV/0! | 4.29E-02 |
| P02770 | CCAEGDPPACYGTVLAEFQPLVEEPK | 1,HNE+Delta_H(2)[C]; | 0.0 | 1.3 | #DIV/0! | 4.29E-02 |
| P02761 | TPEDGEYFVEYDGGNTFTILK | 15,Asn->Lys[N]; | 0.0 | 1.3 | #DIV/0! | 4.29E-02 |
| P02761 | DNIIDLTKTDRCLQA | 12,NEMsulfur[C]; | 0.0 | 1.3 | #DIV/0! | 4.29E-02 |
| P02761 | WFSIVVASNKR | 0,Carbamidomethyl[AnyN-term]; | 0.0 | 1.3 | #DIV/0! | 4.29E-02 |
| P02761 | EKFAKLCEAHGITR | 0,BITC[AnyN-term]; | 0.0 | 2.7 | #DIV/0! | 4.29E-02 |
| Q01177 | SLDGYVSTQGASLHSLTK | 1,DAET[S]; | 0.0 | 3.8 | #DIV/0! | 4.32E-02 |
| P13432 | PTPLSNPPTQLHSTEQANTK | 4,Xle->Cys[L]; | 0.0 | 4.7 | #DIV/0! | 4.41E-02 |
| P02761 | LDVAKLNGDWFSIVVASNKR | 5,CresylSaligeninPhosphate[K]; | 0.0 | 6.5 | #DIV/0! | 4.42E-02 |
| Q62867 | LNDAQYETLFR | 0,Diethyl[AnyN-term]; | 0.0 | 3.5 | #DIV/0! | 4.43E-02 |
| P11232 | LIESKEAFQEALAAAGDK | 0,Succinyl[AnyN-term]; | 0.0 | 3.5 | #DIV/0! | 4.43E-02 |
| P02761 | NGDWFSIVVASNKR | 1,Methylthio[N]; | 0.0 | 3.5 | #DIV/0! | 4.43E-02 |
| P22283 | LVNCPFEEQTEQLKR | 4,Acetyl[C]; | 0.0 | 8.0 | #DIV/0! | 4.45E-02 |
| Q68FP1 | ALNSNDAFVLK | 0,Iodoacetanilide_13C(6)[AnyN-term]; | 0.0 | 3.0 | #DIV/0! | 4.46E-02 |
| P15684 | IKLPASVSTIMDR | 2,Dap-DSP[K]; | 0.0 | 1.5 | #DIV/0! | 4.46E-02 |
| P02770 | LKTVMGDFAQFVDK | 3,Decanoyl[T]; | 0.0 | 1.5 | #DIV/0! | 4.46E-02 |
| P02761 | KLCEAHGITR | 0,Diethylphosphate[AnyN-term]; | 0.0 | 1.5 | #DIV/0! | 4.46E-02 |
| P02761 | EENGSMRVFMQHIDVLENSLGFK | 7,Arg->Glu[R]; | 0.0 | 1.5 | #DIV/0! | 4.46E-02 |
| P02761 | KEKFAKLCEAHGITR | 8,IDEnT[C]; | 0.0 | 1.5 | #DIV/0! | 4.46E-02 |
| P02782 | EELKKELEMYNAPPAAVEAK | 6,GlycerylPE[E]; | 0.0 | 4.0 | #DIV/0! | 4.51E-02 |
| P01836 | QDSKDSTYSMSSTLSLTK | 1,Gln->Thr[Q]; | 0.0 | 3.3 | #DIV/0! | 4.52E-02 |
| P01835 | QDSKDSTYSMSSTLSLTK | 1,Gln->Thr[Q]; | 0.0 | 3.3 | #DIV/0! | 4.52E-02 |
| P07647 | NDLMLLHLSKPADITGGVK | 4,Met->Cys[M]; | 0.0 | 3.7 | #DIV/0! | 4.53E-02 |
| P02761 | LNGDWFSIVVASNKR | 1,Xle->Gln[L]; | 0.0 | 13.2 | #DIV/0! | 4.54E-02 |
| P02761 | LNGDWFSIVVASNKR | 0,Methyl_2H(2)[AnyN-term];2,Deamidated[N]; | 0.0 | 9.7 | #DIV/0! | 4.55E-02 |
| P02770 | RAELAKYMCENQATISSK | 9,maleimide[C]; | 0.0 | 6.2 | #DIV/0! | 4.56E-02 |
| P02770 | HPDYSVSLLLR | 0,mTRAQ[AnyN-term]; | 0.0 | 2.3 | #DIV/0! | 4.57E-02 |
| P02770 | SIHTLFGDKLCAIPK | 11,Carbamidomethyl[C]; | 0.0 | 5.3 | #DIV/0! | 4.65E-02 |
| P22057 | DQGLTEEDIVFLPQPDK | 1,Cation_Li[D]; | 0.0 | 2.8 | #DIV/0! | 4.66E-02 |
| P07647 | VSQSFQHPDYIPVFMR | 0,Diethylphosphate[AnyN-term]; | 0.0 | 2.8 | #DIV/0! | 4.66E-02 |
| P02770 | RPCFSALTVDETYVPK | 2,Pro->Tyr[P]; | 0.0 | 8.3 | #DIV/0! | 4.71E-02 |
| P02761 | KLCEAHGITR | 0,Piperidine[AnyN-term]; | 0.0 | 6.3 | #DIV/0! | 4.71E-02 |
| P02761 | DYDRYVMFHLINFK | 14,HCysThiolactone[K]; | 0.0 | 4.7 | #DIV/0! | 4.72E-02 |
| O35217 | SSCNLFQDIFLHLDK | 3,Carbamidomethyl[C]; | 0.0 | 6.0 | #DIV/0! | 4.74E-02 |
| Q6IE52 | DSARAHFSVMGDILSSAIK | 8,HexNAc(1)dHex(1)[S]; | 0.0 | 2.5 | #DIV/0! | 4.76E-02 |
| Q03626 | DSARAHFSVMGDILSSAIK | 8,HexNAc(1)dHex(1)[S]; | 0.0 | 2.5 | #DIV/0! | 4.76E-02 |
| P36374 | NNLLEDEPFAQHR | 6,Cation_Na[D]; | 0.0 | 2.5 | #DIV/0! | 4.76E-02 |
| P36373 | NNLLEDEPFAQHR | 6,Cation_Na[D]; | 0.0 | 2.5 | #DIV/0! | 4.76E-02 |
| P14046 | DSARAHFSVMGDILSSAIK | 8,HexNAc(1)dHex(1)[S]; | 0.0 | 2.5 | #DIV/0! | 4.76E-02 |
| P02761 | AKLCEAHGITR | 2,PyridoxalPhosphate[K]; | 0.0 | 2.7 | #DIV/0! | 4.77E-02 |
| P02761 | KEKFAKLCEAHGITR | 2,Cation_Na[E]; | 0.0 | 2.7 | #DIV/0! | 4.77E-02 |
| P02761 | NLDVAKLNGDWFSIVVASNKR | 0,NIC[AnyN-term](ICPL[AnyN-term]);8,Deamidated[N]; | 0.0 | 5.3 | #DIV/0! | 4.77E-02 |
| P83121 | ICQTYPDEICAWVVVTTR | 10,NEMsulfur[C]; | 0.0 | 25.5 | #DIV/0! | 4.78E-02 |
| P81827 | ICQTYPDEICAWVVVTTR | 10,NEMsulfur[C]; | 0.0 | 25.5 | #DIV/0! | 4.78E-02 |
| Q63041 | ATPLSLCALTAVDQSVLLLKPEAK | 12,Val->Glu[V]; | 0.0 | 3.8 | #DIV/0! | 4.78E-02 |
| Q6IUU3 | EQMAAASMHQVK | 0,Carboxymethyl[AnyN-term]; | 0.0 | 1.8 | #DIV/0! | 4.78E-02 |
| Q5XI43 | QRVVHWDLSGGPGSQGR | 0,Biotin_Thermo-21330[AnyN-term]; | 0.0 | 1.8 | #DIV/0! | 4.78E-02 |
| P98158 | CGCAFGTLGNDGK | 1,Carbamidomethyl[C];3,Carbamidomethyl[C]; | 0.0 | 1.8 | #DIV/0! | 4.78E-02 |
| P21704 | TNQMAEAISDHYPVEVTLR | 0,Iodoacetanilide[AnyN-term]; | 0.0 | 1.8 | #DIV/0! | 4.78E-02 |
| P02761 | KDLSSDIKEK | 0,Succinyl[AnyN-term]; | 0.0 | 1.8 | #DIV/0! | 4.78E-02 |
| P02761 | KFAKLCEAHGITR | 1,CresylSaligeninPhosphate[K]; | 0.0 | 1.8 | #DIV/0! | 4.78E-02 |
| P01836 | DGVLDSVTDQDSKDSTYSMSSTLSLTK | 0,Acetyl[ProteinN-term]; | 0.0 | 1.8 | #DIV/0! | 4.78E-02 |
| P01835 | DGVLDSVTDQDSKDSTYSMSSTLSLTK | 0,Acetyl[ProteinN-term]; | 0.0 | 1.8 | #DIV/0! | 4.78E-02 |
| Q9JJ19 | AVDPDSPAEASGLR | 2,Val->Cys[V]; | 0.0 | 2.2 | #DIV/0! | 4.82E-02 |
| P98158 | VFWTDPMQEK | 0,Methyl[AnyN-term]; | 0.0 | 2.2 | #DIV/0! | 4.82E-02 |
| P02770 | CPYEEHIKLVQEVTDFAK | 0,PyMIC[AnyN-term]; | 0.0 | 2.2 | #DIV/0! | 4.82E-02 |
| P02761 | FMQHIDVLENSLGFK | 0,Amidine[AnyN-term]; | 0.0 | 2.2 | #DIV/0! | 4.82E-02 |
| P01836 | DGVLDSVTDQDSKDSTYSMSSTLSLTK | 0,Acetyl_2H(3)[AnyN-term]; | 0.0 | 2.2 | #DIV/0! | 4.82E-02 |
| P01835 | DGVLDSVTDQDSKDSTYSMSSTLSLTK | 0,Acetyl_2H(3)[AnyN-term]; | 0.0 | 2.2 | #DIV/0! | 4.82E-02 |
| Q5XI43 | LSGGPGSQGR | 0,GIST-Quat_2H(3)[AnyN-term]; | 0.0 | 3.0 | #DIV/0! | 4.84E-02 |
| P02761 | KIEENGSMR | 0,NIC[AnyN-term](ICPL[AnyN-term]); | 0.0 | 3.0 | #DIV/0! | 4.84E-02 |
| P83121 | CQTYPDEICAWVVVTTR | 1,CarbamidomethylDTT[C]; | 0.0 | 7.2 | #DIV/0! | 4.85E-02 |
| P81828 | CQTYPDEICAWVVVTTR | 1,CarbamidomethylDTT[C]; | 0.0 | 7.2 | #DIV/0! | 4.85E-02 |
| P81827 | CQTYPDEICAWVVVTTR | 1,CarbamidomethylDTT[C]; | 0.0 | 7.2 | #DIV/0! | 4.85E-02 |
| P36373 | PGDDHSNDLMLLHLSQPADITDGVK | 13,DNCB_hapten[H]; | 0.0 | 2.0 | #DIV/0! | 4.93E-02 |
| P02770 | YTQKAPQVSTPTLVEAAR | 0,Galactosyl[AnyN-term]; | 0.0 | 2.0 | #DIV/0! | 4.93E-02 |
| P02761 | QHIDVLENSLGFK | 0,Diethylphosphate[AnyN-term]; | 0.0 | 4.0 | #DIV/0! | 4.93E-02 |
| P02761 | IEENGSMRVFMQHIDVLENSLGFK | 7,Carbamidomethyl[M]; | 0.0 | 2.0 | #DIV/0! | 4.93E-02 |
| P01836 | DGVLDSVTDQDSK | 5,Cation_Na[D];11,Cation_Na[D]; | 0.0 | 2.0 | #DIV/0! | 4.93E-02 |
| P01835 | DGVLDSVTDQDSK | 5,Cation_Na[D];11,Cation_Na[D]; | 0.0 | 2.0 | #DIV/0! | 4.93E-02 |
| P00787 | MCEAGYSTSYK | 2,Carbamidomethyl[C]; | 0.0 | 2.0 | #DIV/0! | 4.93E-02 |
| P00758 | QPGDDYSNDLMLLHLSQPADITDGVK | 0,Gln->pyro-Glu[AnyN-termQ];4,Cation_Na[D]; | 0.0 | 2.0 | #DIV/0! | 4.93E-02 |
| P02761 | NGETFQLMVLYGR | 3,Cation_Ca[II][E]; | 0.0 | 15.3 | #DIV/0! | 4.97E-02 |

Appendix Table 2 Differential modifications screened by FC ≥ 1.5 or ≤ 0.67 with P < 0.05 in the 8-month-old group

| Uniprot ID | Peptide | Modification | Control group | Experimental group | Flod change | P value |
| --- | --- | --- | --- | --- | --- | --- |
| P81827 | CQTYPDEICAWVVVTTR | 1,Carbamidomethyl[C];9,Carbamidomethyl[C]; | 395.1 | 0.0 | 0.0 | 4.11E-07 |
| P81828 | CQTYPDEICAWVVVTTR | 1,Carbamidomethyl[C];9,Carbamidomethyl[C]; | 395.1 | 0.0 | 0.0 | 4.11E-07 |
| P83121 | CQTYPDEICAWVVVTTR | 1,Carbamidomethyl[C];9,Carbamidomethyl[C]; | 395.1 | 0.0 | 0.0 | 4.11E-07 |
| P14046 | VTASPQSLCGLR | 9,Carbamidomethyl[C]; | 18.4 | 0.0 | 0.0 | 1.73E-05 |
| Q03626 | VTASPQSLCGLR | 9,Carbamidomethyl[C]; | 18.4 | 0.0 | 0.0 | 1.73E-05 |
| Q6IE52 | VTASPQSLCGLR | 9,Carbamidomethyl[C]; | 18.4 | 0.0 | 0.0 | 1.73E-05 |
| P27590 | INFECSYPLDMK | 5,Carbamidomethyl[C]; | 40.1 | 0.0 | 0.0 | 9.21E-05 |
| P05545 | TEGAAATAVTAALK | 0,NDA[AnyN-term]; | 231.6 | 0.0 | 0.0 | 1.92E-04 |
| P36953 | ADWCQPLKEDLQNKR | 4,Carbamidomethyl[C]; | 7.4 | 0.0 | 0.0 | 2.63E-04 |
| P12346 | DLKQEDFQLLCPDGTK | 11,Carbamidomethyl[C]; | 43.4 | 0.0 | 0.0 | 3.28E-04 |
| P20767 | ANGAPISQGVDTANPTK | 2,Deamidated[N]; | 21.9 | 0.0 | 0.0 | 3.73E-04 |
| P02770 | FPNAEFAEITK | 3,Deamidated[N]; | 163.9 | 0.0 | 0.0 | 6.41E-04 |
| Q9R0T4 | DWVIPPINCPENQK | 9,Carbamidomethyl[C]; | 56.7 | 0.0 | 0.0 | 6.54E-04 |
| P10960 | GCSFLPDPYQK | 2,Carbamidomethyl[C]; | 15.3 | 0.0 | 0.0 | 7.79E-04 |
| P12346 | KPVTEFATCHLAQAPNHVVVSR | 9,Carbamidomethyl[C];16,Deamidated[N]; | 51.0 | 0.0 | 0.0 | 9.36E-04 |
| P13635 | DIASGLIGPLILCK | 13,Carbamidomethyl[C]; | 30.0 | 0.0 | 0.0 | 1.03E-03 |
| Q63257 | TSTCEWQLDSTVDCSSQLLLDYR | 4,Carbamidomethyl[C];14,Carbamidomethyl[C]; | 8.0 | 0.0 | 0.0 | 1.08E-03 |
| P20759 | SNSMVTLGCLVK | 2,Deamidated[N];9,Carbamidomethyl[C]; | 38.4 | 0.0 | 0.0 | 1.19E-03 |
| P20760 | SNSMVTLGCLVK | 2,Deamidated[N];9,Carbamidomethyl[C]; | 38.4 | 0.0 | 0.0 | 1.19E-03 |
| P18292 | EECSIPVCGQEGR | 3,Carbamidomethyl[C];8,Carbamidomethyl[C]; | 6.7 | 0.0 | 0.0 | 1.21E-03 |
| P02770 | AADKDNCFATEGPNLVAR | 8,Phe->Trp[F]; | 76.4 | 0.0 | 0.0 | 1.65E-03 |
| P12346 | DCTGNFCLFR | 2,Carbamidomethyl[C];7,Carbamidomethyl[C]; | 57.1 | 0.0 | 0.0 | 2.18E-03 |
| P02770 | QEPERNECFLQHK | 8,Carbamidomethyl[C]; | 21.4 | 0.0 | 0.0 | 2.36E-03 |
| P02761 | EKIEENGSMR | 6,Deamidated[N];9,Oxidation[M]; | 127.7 | 0.0 | 0.0 | 2.52E-03 |
| P02770 | YMCENQATISSK | 2,Oxidation[M];3,Carbamidomethyl[C]; | 68.9 | 0.0 | 0.0 | 2.65E-03 |
| P02770 | CCAEGDPPACYGTVLAEFQPLVEEPK | 2,CarbamidomethylDTT[C]; | 27.0 | 0.0 | 0.0 | 2.80E-03 |
| P07314 | TPALCEVFCR | 5,Carbamidomethyl[C];9,Carbamidomethyl[C]; | 17.4 | 0.0 | 0.0 | 2.95E-03 |
| P20759 | SNSMVTLGCLVK | 9,Carbamidomethyl[C]; | 53.0 | 0.0 | 0.0 | 3.07E-03 |
| P20760 | SNSMVTLGCLVK | 9,Carbamidomethyl[C]; | 53.0 | 0.0 | 0.0 | 3.07E-03 |
| P07314 | QPLSSMCPSIIVDKDGK | 7,Carbamidomethyl[C]; | 17.7 | 0.0 | 0.0 | 3.07E-03 |
| P81828 | TSFDSTGFCHVGR | 9,Carbamidomethyl[C]; | 120.7 | 0.0 | 0.0 | 3.09E-03 |
| P98158 | GFTCACPDDFQTVQLR | 4,Carbamidomethyl[C];6,Carbamidomethyl[C]; | 4.4 | 0.0 | 0.0 | 3.42E-03 |
| P12346 | LLEACTFHK | 5,Carbamidomethyl[C]; | 15.7 | 0.0 | 0.0 | 3.72E-03 |
| Q63257 | VLGDPTCFSDYIR | 7,Carbamidomethyl[C]; | 30.3 | 0.0 | 0.0 | 4.05E-03 |
| P27590 | MAETCVPVLR | 5,Carbamidomethyl[C]; | 195.9 | 0.0 | 0.0 | 4.14E-03 |
| P81827 | LDSTGLCR | 7,Carbamidomethyl[C]; | 20.4 | 0.0 | 0.0 | 4.29E-03 |
| Q6P7A9 | AVPTQCDVTPNSR | 6,Carbamidomethyl[C]; | 4.0 | 0.0 | 0.0 | 4.35E-03 |
| P07522 | VNLDPASVPPR | 2,Asn->Cys[N]; | 9.7 | 0.0 | 0.0 | 4.40E-03 |
| Q64230 | THTLLDHCAFEK | 8,Carbamidomethyl[C]; | 11.3 | 0.0 | 0.0 | 4.46E-03 |
| P02770 | AMCTSFQENPTSFLGHYLHEVAR | 3,4-ONE+Delta_H(-2)O(-1)[C]; | 17.3 | 0.0 | 0.0 | 4.62E-03 |
| P22282 | LKDVYDVFK | 0,Formyl[AnyN-term]; | 15.1 | 0.0 | 0.0 | 4.91E-03 |
| P12346 | KPVTEFATCHLAQAPNHVVVSR | 9,Carbamidomethyl[C]; | 156.7 | 0.0 | 0.0 | 4.91E-03 |
| P10252 | ILEYFPNGK | 7,Deamidated[N]; | 5.6 | 0.0 | 0.0 | 4.96E-03 |
| P05545 | IKDLTTPYIRDEELSCSVLELK | 16,Carbamidomethyl[C]; | 966.9 | 0.0 | 0.0 | 4.98E-03 |
| P05544 | TEATAATGVATVIR | 0,PyMIC[AnyN-term]; | 60.3 | 0.0 | 0.0 | 5.52E-03 |
| P09005 | TEATAATGVATVIR | 0,PyMIC[AnyN-term]; | 60.3 | 0.0 | 0.0 | 5.52E-03 |
| P02761 | CEAHGITR | 1,Carbamidomethyl[C]; | 91.6 | 0.0 | 0.0 | 5.71E-03 |
| P05545 | TGTEGAAATAVTAALK | 4,Hydroxamic_acid[E]; | 10.9 | 0.0 | 0.0 | 6.08E-03 |
| P20760 | CKVNSGAFPAPIEK | 1,Carbamidomethyl[C]; | 6.7 | 0.0 | 0.0 | 6.20E-03 |
| P02781 | LEEFDAPPEAVEANLK | 0,CLIP_TRAQ_2[AnyN-term]; | 25.1 | 0.0 | 0.0 | 6.22E-03 |
| P02770 | ADKDNCFATEGPNLVAR | 6,Nmethylmaleimide[C]; | 35.3 | 0.0 | 0.0 | 6.34E-03 |
| P02761 | DLSSDIKEK | 0,Formyl[AnyN-term]; | 2.0 | 0.0 | 0.0 | 6.37E-03 |
| P05544 | MQQVESSLQPETLKK | 1,Oxidation[M]; | 30.9 | 0.0 | 0.0 | 6.47E-03 |
| P05545 | MQQVESSLQPETLKK | 1,Oxidation[M]; | 30.9 | 0.0 | 0.0 | 6.47E-03 |
| P09005 | MQQVESSLQPETLKK | 1,Oxidation[M]; | 30.9 | 0.0 | 0.0 | 6.47E-03 |
| Q04807 | LTMLSQAVQK | 3,Oxidation[M]; | 12.9 | 0.0 | 0.0 | 6.49E-03 |
| P01835 | GVLDSVTDQDSKDSTYSMSSTLSLTK | 4,Cation_Ag[D]; | 11.0 | 0.0 | 0.0 | 6.64E-03 |
| P01836 | GVLDSVTDQDSKDSTYSMSSTLSLTK | 4,Cation_Ag[D]; | 11.0 | 0.0 | 0.0 | 6.64E-03 |
| P98158 | NPCASASCSHLCLLSAQAPR | 3,Carbamidomethyl[C];8,Carbamidomethyl[C];12,Carbamidomethyl[C]; | 9.7 | 0.0 | 0.0 | 6.71E-03 |
| P12346 | TAGWNIPMGLLFSR | 8,Oxidation[M]; | 24.6 | 0.0 | 0.0 | 6.92E-03 |
| Q01177 | YSPSTHPSEGLEENYCR | 16,Carbamidomethyl[C]; | 11.6 | 0.0 | 0.0 | 7.41E-03 |
| O70513 | VDAECMPVVR | 5,Carbamidomethyl[C]; | 5.7 | 0.0 | 0.0 | 7.45E-03 |
| P81827 | ESLDSTGLCR | 9,Carbamidomethyl[C]; | 5.7 | 0.0 | 0.0 | 7.45E-03 |
| P02770 | CCTLPEAQR | 1,Carbamidomethyl[C];2,Carbamidomethyl[C]; | 48.3 | 0.0 | 0.0 | 7.62E-03 |
| Q9JHB9 | QCFLDQTNK | 2,Carbamidomethyl[C]; | 5.0 | 0.0 | 0.0 | 7.72E-03 |
| P81828 | DSTGFCHVGR | 6,Carbamidomethyl[C]; | 91.3 | 0.0 | 0.0 | 7.89E-03 |
| P01835 | SKDSTYSMSSTLSLTK | 0,NDA[AnyN-term]; | 2.1 | 0.0 | 0.0 | 8.24E-03 |
| P01836 | SKDSTYSMSSTLSLTK | 0,NDA[AnyN-term]; | 2.1 | 0.0 | 0.0 | 8.24E-03 |
| Q64230 | DDADWVHEDSSQPGQVDHTLVGR | 6,Val->Thr[V]; | 2.1 | 0.0 | 0.0 | 8.24E-03 |
| P15083 | SSVTFECDLGR | 7,Carbamidomethyl[C]; | 173.1 | 0.0 | 0.0 | 8.25E-03 |
| P20767 | NSFTCQVTHEGNTVEK | 5,Carbamidomethyl[C];12,Deamidated[N]; | 8.4 | 0.0 | 0.0 | 8.31E-03 |
| P05545 | AVLDVDETGTEGAAATAVTAALK | 10,Didehydro[T]; | 15.7 | 0.0 | 0.0 | 8.34E-03 |
| P02770 | CCAEGDPPACYGTVLAEFQPLVEEPK | 1,CarbamidomethylDTT[C]; | 15.7 | 0.0 | 0.0 | 8.40E-03 |
| Q9JJ40 | INGVFVDKEEHAQVVDLVR | 2,Deamidated[N]; | 11.7 | 0.0 | 0.0 | 8.49E-03 |
| P02780 | SGCSILDEVIR | 3,Carbamidomethyl[C]; | 82.0 | 0.0 | 0.0 | 8.53E-03 |
| Q9JHB9 | SGCSILDEVIR | 3,Carbamidomethyl[C]; | 82.0 | 0.0 | 0.0 | 8.53E-03 |
| Q64240 | QGPCRAFAELWAFDAAQGK | 4,BHT[C]; | 4.1 | 0.0 | 0.0 | 8.55E-03 |
| P36953 | SLAMVQQECK | 4,Oxidation[M];9,Carbamidomethyl[C]; | 3.4 | 0.0 | 0.0 | 8.69E-03 |
| P02770 | ADKDNCFATEGPNLVAR | 6,DNCB_hapten[C]; | 46.4 | 0.0 | 0.0 | 9.13E-03 |
| Q64230 | SIDWGWGQAISHQMLMR | 14,Oxidation[M]; | 5.9 | 0.0 | 0.0 | 9.18E-03 |
| P81828 | LALQCFRCTSFDSTGFCHVGR | 5,Carbamidomethyl[C];17,Carbamidomethyl[C]; | 39.1 | 0.0 | 0.0 | 9.36E-03 |
| P26051 | YAGVFHVEKNGR | 10,Deamidated[N]; | 2.0 | 0.0 | 0.0 | 9.60E-03 |
| Q01177 | CTTPPPPPGPTYQCLK | 1,Carbamidomethyl[C];14,Carbamidomethyl[C]; | 8.4 | 0.0 | 0.0 | 9.66E-03 |
| P22282 | TPGETMYYISLPGSVR | 6,Oxidation[M]; | 14.9 | 0.0 | 0.0 | 9.69E-03 |
| Q01177 | KQLAAGSIADCLAK | 11,Carbamidomethyl[C]; | 16.7 | 0.0 | 0.0 | 9.74E-03 |
| P81828 | ALQCFRCTSFDSTGFCHVGR | 4,GG[C](Dicarbamidomethyl[C]);7,Carbamidomethyl[C];16,Carbamidomethyl[C]; | 4.4 | 0.0 | 0.0 | 9.92E-03 |
| P20059 | CNADPGLSALLSDHR | 0,Pyro-carbamidomethyl[AnyN-termC]; | 20.3 | 0.0 | 0.0 | 1.02E-02 |
| Q99PS8 | TSEVVIGQCK | 9,Carbamidomethyl[C]; | 5.4 | 0.0 | 0.0 | 1.04E-02 |
| P02770 | LQACCDKPVLQK | 4,Carbamidomethyl[C]; | 3.1 | 0.0 | 0.0 | 1.05E-02 |
| P02770 | APQVSTPTLVEAAR | 5,Formyl[S](Ser->Asp[S]); | 11.6 | 0.0 | 0.0 | 1.07E-02 |
| P27590 | FSIQMFR | 5,Oxidation[M]; | 195.9 | 0.0 | 0.0 | 1.12E-02 |
| P13635 | DLYSGLIGPLIVCR | 13,Carbamidomethyl[C]; | 21.9 | 0.0 | 0.0 | 1.12E-02 |
| P02780 | LVKPYVQDHFTEK | 0,Carbamyl[AnyN-term]; | 8.3 | 0.0 | 0.0 | 1.12E-02 |
| P15083 | LVGTQALVQEGYEGR | 0,Cytopiloyne[AnyN-term]; | 2.1 | 0.0 | 0.0 | 1.13E-02 |
| P20760 | KETWQQGNTFTCSVLHEGLHNHHTEK | 12,Carbamidomethyl[C]; | 15.7 | 0.0 | 0.0 | 1.13E-02 |
| P81828 | RCTSFDSTGFCHVGR | 2,Carbamidomethyl[C];11,Carbamidomethyl[C]; | 8.1 | 0.0 | 0.0 | 1.16E-02 |
| P02770 | HPYFYAPELLYYAEK | 0,Formyl[AnyN-term]; | 12.6 | 0.0 | 0.0 | 1.19E-02 |
| P02782 | EMYNAPPAAVEAK | 0,Unknown_250[AnyN-term]; | 20.1 | 0.0 | 0.0 | 1.22E-02 |
| P02770 | TVMGDFAQFVDK | 3,Oxidation[M]; | 295.7 | 0.0 | 0.0 | 1.24E-02 |
| P13635 | TFCSEPEKVDKDNEDFQESNR | 3,Carbamidomethyl[C]; | 4.0 | 0.0 | 0.0 | 1.24E-02 |
| P02761 | GDWFSIVVASNKR | 0,Unknown_234[AnyN-term]; | 5.4 | 0.0 | 0.0 | 1.25E-02 |
| P02761 | IKENGECRELYLVAYK | 7,Carbamidomethyl[C]; | 46.6 | 0.0 | 0.0 | 1.26E-02 |
| P22282 | LDNCPFEEQTEQLKR | 1,trifluoro[L]; | 3.6 | 0.0 | 0.0 | 1.28E-02 |
| P12346 | ASDSSINWNNLK | 9,Deamidated[N]; | 51.6 | 0.0 | 0.0 | 1.29E-02 |
| P20767 | NSFTCQVTHEGNTVEK | 1,Deamidated[N];5,Carbamidomethyl[C]; | 11.4 | 0.0 | 0.0 | 1.33E-02 |
| P02770 | APQVSTPTLVEAAR | 8,Formyl[T](Thr->Glu[T]); | 5.0 | 0.0 | 0.0 | 1.36E-02 |
| O70244 | SCGGYLHADR | 2,Carbamidomethyl[C]; | 11.1 | 0.0 | 0.0 | 1.37E-02 |
| P15083 | YWCHWEADENGR | 3,Carbamidomethyl[C]; | 2.9 | 0.0 | 0.0 | 1.37E-02 |
| P27590 | CQLQSLGFMK | 1,Carbamidomethyl[C]; | 75.0 | 0.0 | 0.0 | 1.37E-02 |
| P06760 | TSHYPYSEEVLQLCDR | 14,Carbamidomethyl[C]; | 37.0 | 0.0 | 0.0 | 1.37E-02 |
| Q64230 | TNICGMIQGTR | 4,Carbamidomethyl[C]; | 43.6 | 0.0 | 0.0 | 1.38E-02 |
| P20767 | NSFTCQVTHEGNTVEK | 5,Carbamidomethyl[C]; | 34.3 | 0.0 | 0.0 | 1.38E-02 |
| P07522 | LSVIEMADLDGSKR | 6,Oxidation[M]; | 13.9 | 0.0 | 0.0 | 1.39E-02 |
| P01835 | VLDSVTDQDSKDSTYSMSSTLSLTK | 0,Diisopropylphosphate[AnyN-term]; | 18.7 | 0.0 | 0.0 | 1.42E-02 |
| P01836 | VLDSVTDQDSKDSTYSMSSTLSLTK | 0,Diisopropylphosphate[AnyN-term]; | 18.7 | 0.0 | 0.0 | 1.42E-02 |
| P81828 | LALQCFRCTSFDSTGFCHVGR | 8,Carbamidomethyl[C];17,Carbamidomethyl[C]; | 46.6 | 0.0 | 0.0 | 1.44E-02 |
| P07861 | LIQNMDASAEPCTDFFK | 12,Carbamidomethyl[C]; | 62.4 | 0.0 | 0.0 | 1.46E-02 |
| P22282 | LDNCPFEEQTEQLKR | 4,Carbamidomethyl[C]; | 210.4 | 0.0 | 0.0 | 1.50E-02 |
| P36953 | LCFFYNKK | 2,Carbamidomethyl[C]; | 2.7 | 0.0 | 0.0 | 1.53E-02 |
| P09006 | MQQVEASLQPETLR | 1,Oxidation[M]; | 7.6 | 0.0 | 0.0 | 1.53E-02 |
| P12346 | NGDGKEDLIWEILK | 0,C+12[AnyN-term]; | 10.6 | 0.0 | 0.0 | 1.54E-02 |
| P01835 | DSKDSTYSMSSTLSLTK | 0,Propionyl_13C(3)[AnyN-term]; | 32.0 | 0.0 | 0.0 | 1.55E-02 |
| P01836 | DSKDSTYSMSSTLSLTK | 0,Propionyl_13C(3)[AnyN-term]; | 32.0 | 0.0 | 0.0 | 1.55E-02 |
| P07151 | KIPNIEMSDLSFSK | 7,Oxidation[M]; | 4.9 | 0.0 | 0.0 | 1.56E-02 |
| P36953 | TINPTVDHCCR | 9,Carbamidomethyl[C];10,Carbamidomethyl[C]; | 6.1 | 0.0 | 0.0 | 1.58E-02 |
| Q63678 | CLAYDFYPQR | 1,Carbamidomethyl[C]; | 9.7 | 0.0 | 0.0 | 1.62E-02 |
| P07861 | MDITDINAPKPK | 0,Acetyl[ProteinN-term];1,Oxidation[M]; | 8.0 | 0.0 | 0.0 | 1.63E-02 |
| Q01177 | CEGETDFICR | 1,Carbamidomethyl[C];9,Carbamidomethyl[C]; | 6.6 | 0.0 | 0.0 | 1.66E-02 |
| P12346 | GTDFQLNQLQGK | 7,Deamidated[N]; | 8.6 | 0.0 | 0.0 | 1.67E-02 |
| P28826 | LSQVLSGPENDHSNMGQCK | 18,Carbamidomethyl[C]; | 13.1 | 0.0 | 0.0 | 1.72E-02 |
| P12346 | EGVCPEGSIDSAPVK | 4,Carbamidomethyl[C]; | 57.7 | 0.0 | 0.0 | 1.76E-02 |
| P15684 | VVATTQMQAADAR | 7,Oxidation[M]; | 3.0 | 0.0 | 0.0 | 1.77E-02 |
| P02770 | YNEVLTQCCTESDK | 8,Carbamidomethyl[C];9,Carbamidomethyl[C]; | 342.0 | 0.0 | 0.0 | 1.80E-02 |
| P02770 | PPACYGTVLAEFQPLVEEPK | 4,Carbamidomethyl[C]; | 108.7 | 0.0 | 0.0 | 1.81E-02 |
| P97546 | AEDSGEYHCVYHFVSAPK | 9,Carbamidomethyl[C]; | 10.7 | 0.0 | 0.0 | 1.82E-02 |
| Q9JJ19 | IVEVNGVCMEGK | 5,Deamidated[N];8,Carbamidomethyl[C];9,Oxidation[M]; | 5.4 | 0.0 | 0.0 | 1.89E-02 |
| P20611 | FPLGPCPR | 6,Carbamidomethyl[C]; | 6.1 | 0.0 | 0.0 | 1.91E-02 |
| P36373 | LVSQSFPHPDYKPFLMR | 0,Formyl[AnyN-term]; | 7.4 | 0.0 | 0.0 | 1.92E-02 |
| P22282 | LDNCPFEEQTEQLKR | 3,Deamidated[N];4,Carbamidomethyl[C]; | 9.1 | 0.0 | 0.0 | 1.93E-02 |
| P22282 | RTPGETMYYISLPGSVR | 7,Met->His[M]; | 7.9 | 0.0 | 0.0 | 1.94E-02 |
| P07522 | HDLLTMAIFGDK | 6,Oxidation[M]; | 15.1 | 0.0 | 0.0 | 1.95E-02 |
| P12346 | FDEFFSQGCAPGYK | 9,Carbamidomethyl[C]; | 14.3 | 0.0 | 0.0 | 1.97E-02 |
| P98158 | AIVLDPCR | 7,Carbamidomethyl[C]; | 5.9 | 0.0 | 0.0 | 1.97E-02 |
| P17475 | AFHHLLQTLNRPDSELQLNTGNGLFVNK | 19,Deamidated[N]; | 4.1 | 0.0 | 0.0 | 1.99E-02 |
| P02770 | TCVADENAENCDK | 11,Carbamidomethyl[C]; | 6.7 | 0.0 | 0.0 | 1.99E-02 |
| P05544 | AVLDVDETGTEATAATGVATVIR | 0,ISD_z+2_ion[AnyN-term]; | 44.0 | 0.0 | 0.0 | 2.02E-02 |
| P02770 | QENPTSFLGHYLHEVAR | 0,Dimethyl_2H(4)[AnyN-term]; | 6.4 | 0.0 | 0.0 | 2.04E-02 |
| P04276 | VCQELSTLGK | 2,Carbamidomethyl[C]; | 7.1 | 0.0 | 0.0 | 2.11E-02 |
| P08723 | FLLEDGEHVTQVSGTR | 0,Phenylisocyanate_2H(5)[AnyN-term]; | 6.3 | 0.0 | 0.0 | 2.13E-02 |
| P02761 | MQHIDVLENSLGFK | 0,C+12[AnyN-term]; | 8.7 | 0.0 | 0.0 | 2.13E-02 |
| Q64240 | CIQFIYGGCK | 1,Carbamidomethyl[C];9,Carbamidomethyl[C]; | 19.3 | 0.0 | 0.0 | 2.20E-02 |
| P07522 | GQCLYSLSER | 3,Carbamidomethyl[C]; | 12.9 | 0.0 | 0.0 | 2.22E-02 |
| P01048 | EALGHSIAQLNAQHNHIFYFK | 0,Glu->pyro-Glu[AnyN-termE]; | 7.4 | 0.0 | 0.0 | 2.23E-02 |
| P08932 | EALGHSIAQLNAQHNHIFYFK | 0,Glu->pyro-Glu[AnyN-termE]; | 7.4 | 0.0 | 0.0 | 2.23E-02 |
| P02761 | LCEAHGITR | 2,maleimide[C]; | 2.9 | 0.0 | 0.0 | 2.25E-02 |
| P02770 | DKDNCFATEGPNLVAR | 5,BHTOH[C]; | 7.7 | 0.0 | 0.0 | 2.26E-02 |
| P02761 | TKDLSSDIK | 1,Dehydrated[T]; | 2.1 | 0.0 | 0.0 | 2.28E-02 |
| P05544 | TSMVLVNYLLFK | 3,Oxidation[M]; | 17.9 | 0.0 | 0.0 | 2.29E-02 |
| P05545 | TSMVLVNYLLFK | 3,Oxidation[M]; | 17.9 | 0.0 | 0.0 | 2.29E-02 |
| Q01177 | NLEENYCR | 7,Carbamidomethyl[C]; | 3.7 | 0.0 | 0.0 | 2.33E-02 |
| P02761 | LCEAHGITR | 2,Cys->Dha[C]; | 28.9 | 0.0 | 0.0 | 2.35E-02 |
| P05545 | GAAATAVTAALK | 5,PhosphoHex(2)[T]; | 42.4 | 0.0 | 0.0 | 2.35E-02 |
| P22282 | DNCPFEEQTEQLKR | 3,4-ONE+Delta_H(-2)O(-1)[C]; | 4.0 | 0.0 | 0.0 | 2.36E-02 |
| P36373 | KPGDDHSNDLMLLHLSQPADITDGVK | 23,Dehydrated[D]; | 5.9 | 0.0 | 0.0 | 2.42E-02 |
| P98158 | GSYECFCVDGFK | 5,Carbamidomethyl[C];7,Carbamidomethyl[C]; | 7.3 | 0.0 | 0.0 | 2.43E-02 |
| P20059 | NPLPAAHETVAK | 1,Deamidated[N]; | 1.6 | 0.0 | 0.0 | 2.48E-02 |
| P02761 | LCEAHGITRDNIIDLTK | 2,Carbamidomethyl[C]; | 11.4 | 0.0 | 0.0 | 2.48E-02 |
| P02761 | VFMQHIDVLENSLGFK | 5,His->Xle[H]; | 8.3 | 0.0 | 0.0 | 2.56E-02 |
| Q68FP1 | AGALNSNDAFVLK | 4,Xle->Cys[L]; | 4.7 | 0.0 | 0.0 | 2.57E-02 |
| P02780 | PYVQDHFTEK | 0,Formyl[AnyN-term]; | 4.6 | 0.0 | 0.0 | 2.60E-02 |
| P02770 | TCVADENAENCDK | 2,Carbamidomethyl[C]; | 3.0 | 0.0 | 0.0 | 2.63E-02 |
| Q9QX79 | IHSMCPDCPHPVDLSAPSVLEAATESLAK | 4,Oxidation[M];5,Carbamidomethyl[C];8,Carbamidomethyl[C]; | 12.3 | 0.0 | 0.0 | 2.65E-02 |
| P02761 | EKIEENGSMR | 0,Glu->pyro-Glu[AnyN-termE]; | 12.6 | 0.0 | 0.0 | 2.67E-02 |
| P12346 | NGDGKEDLIWEILK | 1,Asn->Xle[N]; | 20.1 | 0.0 | 0.0 | 2.68E-02 |
| P02770 | CCSGSLVER | 1,Carbamidomethyl[C];2,Carbamidomethyl[C]; | 118.7 | 0.0 | 0.0 | 2.69E-02 |
| P02770 | AADKDNCFATEGPNLVAR | 7,CarbamidomethylDTT[C]; | 7.1 | 0.0 | 0.0 | 2.69E-02 |
| P12346 | NGDGKEDLIWEILK | 3,Dehydrated[D]; | 5.0 | 0.0 | 0.0 | 2.70E-02 |
| P01835 | IFPPSTEQLATGGASVVCLMNNFYPR | 18,Carbamidomethyl[C]; | 6.4 | 0.0 | 0.0 | 2.72E-02 |
| P05545 | TGTEGAAATAVTAALK | 1,Thr->Asp[T]; | 6.3 | 0.0 | 0.0 | 2.74E-02 |
| P01835 | SKDSTYSMSSTLSLTK | 1,Glucuronyl[S]; | 11.6 | 0.0 | 0.0 | 2.74E-02 |
| P01836 | SKDSTYSMSSTLSLTK | 1,Glucuronyl[S]; | 11.6 | 0.0 | 0.0 | 2.74E-02 |
| P0DP29 | VFDKDGNGYISAAELR | 7,Deamidated[N]; | 9.7 | 0.0 | 0.0 | 2.78E-02 |
| P0DP30 | VFDKDGNGYISAAELR | 7,Deamidated[N]; | 9.7 | 0.0 | 0.0 | 2.78E-02 |
| P0DP31 | VFDKDGNGYISAAELR | 7,Deamidated[N]; | 9.7 | 0.0 | 0.0 | 2.78E-02 |
| P04916 | DPNGLTPETR | 3,Deamidated[N]; | 5.9 | 0.0 | 0.0 | 2.78E-02 |
| Q99041 | FSVESLGLNNMK | 11,Oxidation[M]; | 5.9 | 0.0 | 0.0 | 2.78E-02 |
| P02781 | ELEEFDAPPEAVEANLK | 0,C+12[AnyN-term]; | 23.3 | 0.0 | 0.0 | 2.79E-02 |
| P61589 | IGAFGYMECSAK | 9,Carbamidomethyl[C]; | 7.4 | 0.0 | 0.0 | 2.81E-02 |
| P02770 | VGTKCCTLPEAQR | 5,Carbamidomethyl[C];6,Carbamidomethyl[C]; | 4.1 | 0.0 | 0.0 | 2.92E-02 |
| P07314 | QPLSSMCPSIIVDK | 7,Carbamidomethyl[C]; | 4.3 | 0.0 | 0.0 | 2.94E-02 |
| Q63041 | GGENCIKAEDITHNGIVYTPK | 5,Carbamidomethyl[C];14,Deamidated[N]; | 4.3 | 0.0 | 0.0 | 2.94E-02 |
| P02761 | FAKLCEAHGITR | 5,Carbamidomethyl[C]; | 2.3 | 0.0 | 0.0 | 3.00E-02 |
| P02782 | QICELVAHETISFLMK | 3,HNE-Delta_H(2)O[C]; | 2.3 | 0.0 | 0.0 | 3.00E-02 |
| P05545 | LDVDETGTEGAAATAVTAALK | 0,Tween20[AnyN-term]; | 3.4 | 0.0 | 0.0 | 3.00E-02 |
| Q63257 | SFPVNTLTSGVR | 1,Isopropylphospho[S]; | 1.7 | 0.0 | 0.0 | 3.00E-02 |
| P00758 | QPGDDYSNDLMLLHLSQPADITDGVK | 0,Formyl[AnyN-term]; | 4.1 | 0.0 | 0.0 | 3.05E-02 |
| Q9QX79 | DGYMLTLNR | 4,Oxidation[M]; | 8.3 | 0.0 | 0.0 | 3.05E-02 |
| P15684 | QLQTDLSVIPVINR | 0,Biotin_Thermo-21328[AnyN-term]; | 7.4 | 0.0 | 0.0 | 3.16E-02 |
| P07522 | VNLDPASVPPR | 0,C+12[AnyN-term]; | 6.7 | 0.0 | 0.0 | 3.16E-02 |
| P13635 | GQFPLSIQPMGVR | 10,Oxidation[M]; | 19.9 | 0.0 | 0.0 | 3.18E-02 |
| P32038 | EVKPGTLCDVAGWGVVTHAGR | 8,Delta_H(4)C(3)O(1)[C]; | 3.4 | 0.0 | 0.0 | 3.18E-02 |
| P05544 | AVLDVDETGTEATAATGVATVIR | 7,Glu->Asn[E]; | 99.4 | 0.0 | 0.0 | 3.20E-02 |
| P02782 | SEEELKKELEMYNAPPAAVEAK | 13,Ammonia-loss[N]; | 3.9 | 0.0 | 0.0 | 3.20E-02 |
| P30120 | GYQSDHFACLPR | 9,Carbamidomethyl[C]; | 3.9 | 0.0 | 0.0 | 3.20E-02 |
| P07151 | IPNIEMSDLSFSK | 6,Oxidation[M]; | 9.4 | 0.0 | 0.0 | 3.21E-02 |
| P15083 | DDNGRFSVLITGLR | 3,Deamidated[N]; | 27.0 | 0.0 | 0.0 | 3.23E-02 |
| P22282 | LLSKLDNCPFEEQTEQLKR | 8,Carbamidomethyl[C]; | 7.1 | 0.0 | 0.0 | 3.23E-02 |
| P07647 | AYDHNNDLMLLHLSK | 6,Deamidated[N]; | 12.3 | 0.0 | 0.0 | 3.27E-02 |
| Q08420 | MGLATSLAGPHSILGR | 1,Oxidation[M]; | 4.1 | 0.0 | 0.0 | 3.30E-02 |
| P02770 | PTSFLGHYLHEVAR | 0,Methyl_2H(3)13C(1)[AnyN-term]; | 3.1 | 0.0 | 0.0 | 3.33E-02 |
| P02780 | QCFLDQTDK | 2,Cys->Dha[C]; | 1.6 | 0.0 | 0.0 | 3.33E-02 |
| P06761 | VEIIANDQGNR | 6,Deamidated[N]; | 1.6 | 0.0 | 0.0 | 3.33E-02 |
| P07522 | LIFWSSVVTGNLHR | 1,Xle->Glu[L]; | 1.6 | 0.0 | 0.0 | 3.33E-02 |
| P0DMW0 | VEIIANDQGNR | 6,Deamidated[N]; | 1.6 | 0.0 | 0.0 | 3.33E-02 |
| P0DMW1 | VEIIANDQGNR | 6,Deamidated[N]; | 1.6 | 0.0 | 0.0 | 3.33E-02 |
| P14659 | VEIIANDQGNR | 6,Deamidated[N]; | 1.6 | 0.0 | 0.0 | 3.33E-02 |
| P55063 | VEIIANDQGNR | 6,Deamidated[N]; | 1.6 | 0.0 | 0.0 | 3.33E-02 |
| P63018 | VEIIANDQGNR | 6,Deamidated[N]; | 1.6 | 0.0 | 0.0 | 3.33E-02 |
| P81828 | LQCFRCTSFDSTGFCHVGR | 3,Carbamidomethyl[C];15,Carbamidomethyl[C]; | 1.6 | 0.0 | 0.0 | 3.33E-02 |
| P07522 | ADLGGMDVK | 6,Oxidation[M]; | 5.4 | 0.0 | 0.0 | 3.33E-02 |
| P05545 | IAELFSELDER | 0,C+12[AnyN-term]; | 11.3 | 0.0 | 0.0 | 3.35E-02 |
| P02781 | ELEEFDAPPEAVEANLK | 6,Asp->Lys[D]; | 43.6 | 0.0 | 0.0 | 3.36E-02 |
| P12788 | HPEYNKDTLDNDIMLIK | 6,TMT[K]; | 8.0 | 0.0 | 0.0 | 3.37E-02 |
| P14659 | LLQDFFNGK | 7,Deamidated[N]; | 12.3 | 0.0 | 0.0 | 3.38E-02 |
| P63018 | LLQDFFNGK | 7,Deamidated[N]; | 12.3 | 0.0 | 0.0 | 3.38E-02 |
| P02770 | EDQLKTVMGDFAQFVDK | 4,trifluoro[L]; | 103.6 | 0.0 | 0.0 | 3.40E-02 |
| P82450 | TFEISCCSDHQCK | 6,Carbamidomethyl[C];7,Carbamidomethyl[C];12,Carbamidomethyl[C]; | 2.3 | 0.0 | 0.0 | 3.42E-02 |
| P01835 | DGVLDSVTDQDSKDSTYSMSSTLSLTK | 4,Xle->Thr[L]; | 6.9 | 0.0 | 0.0 | 3.46E-02 |
| P01836 | DGVLDSVTDQDSKDSTYSMSSTLSLTK | 4,Xle->Thr[L]; | 6.9 | 0.0 | 0.0 | 3.46E-02 |
| P02782 | SQICELVAHETISFLMK | 4,Trioxidation[C]; | 5.0 | 0.0 | 0.0 | 3.47E-02 |
| P36373 | NNLLEDEPFAQHR | 1,Ammonia-loss[N]; | 5.0 | 0.0 | 0.0 | 3.47E-02 |
| P36374 | NNLLEDEPFAQHR | 1,Ammonia-loss[N]; | 5.0 | 0.0 | 0.0 | 3.47E-02 |
| P05544 | QPILSEFQEK | 0,PyMIC[AnyN-term]; | 6.6 | 0.0 | 0.0 | 3.48E-02 |
| P05545 | QPILSEFQEK | 0,PyMIC[AnyN-term]; | 6.6 | 0.0 | 0.0 | 3.48E-02 |
| P15083 | CHILPSHDEGAR | 1,Cys->Gly[C]; | 3.9 | 0.0 | 0.0 | 3.49E-02 |
| P02761 | EKIEENGSMR | 6,Ammonia-loss[N]; | 8.0 | 0.0 | 0.0 | 3.51E-02 |
| P02770 | VSTPTLVEAAR | 0,Unknown_302[AnyN-term]; | 12.0 | 0.0 | 0.0 | 3.51E-02 |
| P02770 | FQENPTSFLGHYLHEVAR | 0,SPITC[AnyN-term]; | 19.7 | 0.0 | 0.0 | 3.52E-02 |
| P27590 | GYRGDGWYCECSPGFCEPGLDCLPQGPSGK | 9,Carbamidomethyl[C];11,GG[C](Dicarbamidomethyl[C]);22,Carbamidomethyl[C]; | 1.4 | 0.0 | 0.0 | 3.53E-02 |
| P07522 | RLFWTDTGMSPR | 9,Oxidation[M]; | 3.1 | 0.0 | 0.0 | 3.55E-02 |
| P07522 | HDLLTMAIFGDK | 5,Dehydrated[T]; | 10.6 | 0.0 | 0.0 | 3.56E-02 |
| P07647 | VGSICLASGWGMTNPSEMK | 5,Carbamidomethyl[C]; | 30.7 | 0.0 | 0.0 | 3.61E-02 |
| P12346 | CAVSEHENTK | 0,Biotin[AnyN-term]; | 4.7 | 0.0 | 0.0 | 3.63E-02 |
| P12788 | HPEYNKDTLDNDIMLIK | 6,TMT2plex[K]; | 21.6 | 0.0 | 0.0 | 3.64E-02 |
| P02782 | QICELVAHETISFLMK | 3,4-ONE+Delta_H(-2)O(-1)[C]; | 3.9 | 0.0 | 0.0 | 3.64E-02 |
| P02782 | VAHETISFLMK | 0,Propionyl[AnyN-term]; | 12.0 | 0.0 | 0.0 | 3.65E-02 |
| P01835 | DSKDSTYSMSSTLSLTK | 2,AEC-MAEC_2H(4)[S]; | 2.1 | 0.0 | 0.0 | 3.66E-02 |
| P01836 | DSKDSTYSMSSTLSLTK | 2,AEC-MAEC_2H(4)[S]; | 2.1 | 0.0 | 0.0 | 3.66E-02 |
| P02761 | TPEDGEYFVEYDGGNTFTILK | 3,Cation_Na[E]; | 2.1 | 0.0 | 0.0 | 3.66E-02 |
| P14046 | YMVLVPSQLYTETPEK | 2,Oxidation[M]; | 2.1 | 0.0 | 0.0 | 3.66E-02 |
| Q03626 | YMVLVPSQLYTETPEK | 2,Oxidation[M]; | 2.1 | 0.0 | 0.0 | 3.66E-02 |
| Q63041 | AEDITHNGIVYTPK | 14,Lys->Asp[K]; | 2.1 | 0.0 | 0.0 | 3.66E-02 |
| Q6IE52 | YMVLVPSQLYTETPEK | 2,Oxidation[M]; | 2.1 | 0.0 | 0.0 | 3.66E-02 |
| P05545 | QLHSLTLASINTDFTLSLYK | 10,Xle->Lys[I]; | 115.4 | 0.0 | 0.0 | 3.67E-02 |
| P02770 | MSQRFPNAEFAEITK | 0,Diethylphosphate[AnyN-term]; | 23.7 | 0.0 | 0.0 | 3.69E-02 |
| P22057 | QGHDTVQPNFQQDKFLGR | 0,Gln->pyro-Glu[AnyN-termQ]; | 4.4 | 0.0 | 0.0 | 3.71E-02 |
| P17475 | GTEAAGATVVEAVPMSLPPQVK | 15,Oxidation[M]; | 26.4 | 0.0 | 0.0 | 3.72E-02 |
| P05544 | AVLDVDETGTEATAATGVATVIR | 10,Dehydrated[T]; | 5.6 | 0.0 | 0.0 | 3.73E-02 |
| P02770 | FPNAEFAEITK | 2,Pro->Thr[P]; | 1.7 | 0.0 | 0.0 | 3.75E-02 |
| P02770 | CFATEGPNLVAR | 1,Carbamidomethyl[C]; | 6.7 | 0.0 | 0.0 | 3.75E-02 |
| P02770 | RPCFSALTVDETYVPK | 3,Trioxidation[C]; | 34.9 | 0.0 | 0.0 | 3.76E-02 |
| P07647 | LVSQSFQHPDYIPVFMR | 16,Oxidation[M]; | 20.7 | 0.0 | 0.0 | 3.78E-02 |
| P05544 | QPILSEFQEK | 0,Iodoacetanilide[AnyN-term]; | 2.9 | 0.0 | 0.0 | 3.81E-02 |
| P05545 | QPILSEFQEK | 0,Iodoacetanilide[AnyN-term]; | 2.9 | 0.0 | 0.0 | 3.81E-02 |
| Q68FP1 | AQHVQVEEGSEPDGFWEALGGK | 12,Pro->Ser[P]; | 17.1 | 0.0 | 0.0 | 3.82E-02 |
| P02770 | TNCELYEKLGEYGFQNAVLVR | 1,HexNAc(1)NeuAc(1)[T]; | 3.0 | 0.0 | 0.0 | 3.82E-02 |
| P02770 | APQVSTPTLVEAAR | 6,Formyl[T](Thr->Glu[T]); | 7.7 | 0.0 | 0.0 | 3.83E-02 |
| P11232 | MIKPFFHSLCDK | 10,Carbamidomethyl[C]; | 5.1 | 0.0 | 0.0 | 3.83E-02 |
| P19218 | YCTDPSTAPK | 2,Carbamidomethyl[C]; | 2.6 | 0.0 | 0.0 | 3.83E-02 |
| P02770 | YNEVLTQCCTESDK | 2,Deamidated[N];8,Carbamidomethyl[C];9,Carbamidomethyl[C]; | 6.3 | 0.0 | 0.0 | 3.84E-02 |
| P07522 | GSMLKPSSLVVVHPLAK | 0,Formyl[AnyN-term]; | 6.4 | 0.0 | 0.0 | 3.87E-02 |
| Q63041 | ATPLSLCALTAVDQSVLLLKPEAK | 7,Carbamidomethyl[C]; | 4.9 | 0.0 | 0.0 | 3.89E-02 |
| P02770 | PYFYAPELLYYAEK | 1,Glycosyl[P]; | 3.6 | 0.0 | 0.0 | 3.93E-02 |
| P12346 | HAVVARNGDGKEDLIWEILK | 1,Diethylphosphothione[H]; | 9.4 | 0.0 | 0.0 | 3.96E-02 |
| P17475 | AFHHLLQTLNRPDSELQLNTGNGLFVNK | 22,Deamidated[N]; | 16.4 | 0.0 | 0.0 | 3.96E-02 |
| P36374 | AYNDEVTDVMLCAGEMDGGKDICK | 12,Carbamidomethyl[C];23,Carbamidomethyl[C]; | 44.3 | 0.0 | 0.0 | 4.01E-02 |
| Q01460 | EDCPCSEASLCRPIR | 3,Carbamidomethyl[C];5,Carbamidomethyl[C];11,Carbamidomethyl[C]; | 3.1 | 0.0 | 0.0 | 4.01E-02 |
| P12346 | YLGHSYVTAIR | 0,Pyridylacetyl[AnyN-term](Phenylisocyanate[AnyN-term]); | 2.7 | 0.0 | 0.0 | 4.01E-02 |
| P05544 | GTEATAATGVATVIR | 5,Phospho[T]; | 25.7 | 0.0 | 0.0 | 4.04E-02 |
| P09005 | GTEATAATGVATVIR | 5,Phospho[T]; | 25.7 | 0.0 | 0.0 | 4.04E-02 |
| P05545 | VLDVDETGTEGAAATAVTAALK | 0,Piperidine[AnyN-term]; | 3.0 | 0.0 | 0.0 | 4.08E-02 |
| P02770 | DQLKTVMGDFAQFVDK | 0,Triton[AnyN-term]; | 4.1 | 0.0 | 0.0 | 4.08E-02 |
| Q64240 | AFAELWAFDAAQGK | 4,Glu->Gln[E]; | 3.3 | 0.0 | 0.0 | 4.10E-02 |
| P02781 | RELEEFDAPPEAVEANLK | 13,Val->Gln[V]; | 3.6 | 0.0 | 0.0 | 4.11E-02 |
| P02761 | IEENGSMRVFMQHIDVLENSLGFK | 8,Arg->Gly[R]; | 10.9 | 0.0 | 0.0 | 4.11E-02 |
| P22283 | CSFQWEFCNIK | 1,Carbamidomethyl[C];8,Carbamidomethyl[C]; | 55.4 | 0.0 | 0.0 | 4.13E-02 |
| P02781 | ELEEFDAPPEAVEANLK | 6,Asp->Gln[D]; | 12.4 | 0.0 | 0.0 | 4.14E-02 |
| P02780 | SGSGCSILDEVIR | 5,CarbamidomethylDTT[C]; | 33.9 | 0.0 | 0.0 | 4.16E-02 |
| Q9JHB9 | SGSGCSILDEVIR | 5,CarbamidomethylDTT[C]; | 33.9 | 0.0 | 0.0 | 4.16E-02 |
| P02782 | SEEELKKELEMYNAPPAAVEAK | 8,Glu->Asn[E]; | 1.6 | 0.0 | 0.0 | 4.24E-02 |
| P32038 | EDREVKPGTLCDVAGWGVVTHAGR | 11,Carbamidomethyl[C]; | 5.4 | 0.0 | 0.0 | 4.25E-02 |
| P04276 | YCSSQIDAEMR | 2,Carbamidomethyl[C];10,Oxidation[M]; | 2.4 | 0.0 | 0.0 | 4.27E-02 |
| P07522 | HQARHDLLTMAIFGDK | 0,NDA[AnyN-term]; | 2.3 | 0.0 | 0.0 | 4.28E-02 |
| P01835 | VLDSVTDQDSKDSTYSMSSTLSLTK | 16,Hex[S]; | 5.9 | 0.0 | 0.0 | 4.39E-02 |
| P01836 | VLDSVTDQDSKDSTYSMSSTLSLTK | 16,Hex[S]; | 5.9 | 0.0 | 0.0 | 4.39E-02 |
| Q62867 | VVPIRLDLNDAQYETLFR | 0,Unknown_250[AnyN-term]; | 5.6 | 0.0 | 0.0 | 4.39E-02 |
| P14046 | GMYESLPVVAVK | 2,Oxidation[M]; | 6.0 | 0.0 | 0.0 | 4.40E-02 |
| P05544 | MQQVESSLQPETLKK | 0,ISD_z+2_ion[AnyN-term]; | 4.3 | 0.0 | 0.0 | 4.40E-02 |
| P05545 | MQQVESSLQPETLKK | 0,ISD_z+2_ion[AnyN-term]; | 4.3 | 0.0 | 0.0 | 4.40E-02 |
| P09005 | MQQVESSLQPETLKK | 0,ISD_z+2_ion[AnyN-term]; | 4.3 | 0.0 | 0.0 | 4.40E-02 |
| Q5XI43 | IDDLEPADEGIYSCHLHHHYCGLHER | 14,Carbamidomethyl[C];21,Carbamidomethyl[C]; | 10.4 | 0.0 | 0.0 | 4.43E-02 |
| P02770 | QENPTSFLGHYLHEVAR | 4,Pro->Lys[P]; | 27.6 | 0.0 | 0.0 | 4.47E-02 |
| P36373 | NNLLEDEPFAQHR | 2,Ammonia-loss[N]; | 6.7 | 0.0 | 0.0 | 4.51E-02 |
| P36374 | NNLLEDEPFAQHR | 2,Ammonia-loss[N]; | 6.7 | 0.0 | 0.0 | 4.51E-02 |
| P01835 | ADYESHNLYTCEVVHK | 7,Deamidated[N];11,Carbamidomethyl[C]; | 2.6 | 0.0 | 0.0 | 4.53E-02 |
| P02770 | DKDNCFATEGPNLVAR | 0,Unknown_234[AnyN-term]; | 2.6 | 0.0 | 0.0 | 4.53E-02 |
| P20759 | MNGQPQENYK | 2,Deamidated[N]; | 5.9 | 0.0 | 0.0 | 4.53E-02 |
| P20760 | MNGQPQENYK | 2,Deamidated[N]; | 5.9 | 0.0 | 0.0 | 4.53E-02 |
| P02761 | GNLDVAKLNGDWFSIVVASNKR | 7,DNCB_hapten[K]; | 46.6 | 0.0 | 0.0 | 4.59E-02 |
| P02761 | VFMQHIDVLENSLGFK | 12,Formyl[S](Ser->Asp[S]); | 32.3 | 0.0 | 0.0 | 4.60E-02 |
| Q63691 | GLISALCPHKFPTLQVLALR | 7,Carbamidomethyl[C]; | 10.1 | 0.0 | 0.0 | 4.65E-02 |
| P05544 | AVLDVDETGTEATAATGVATVIR | 7,Cation_Na[E]; | 22.1 | 0.0 | 0.0 | 4.65E-02 |
| P02761 | LNGDWFSIVVASNKR | 12,Formyl[S](Ser->Asp[S]); | 1.4 | 0.0 | 0.0 | 4.65E-02 |
| P02770 | LRDNYGELADCCAK | 12,Carbamidomethyl[C]; | 2.1 | 0.0 | 0.0 | 4.65E-02 |
| P02780 | LVKPYVQDHFTEK | 4,Pro->Ser[P]; | 5.0 | 0.0 | 0.0 | 4.65E-02 |
| P14046 | QWQDVKTENGLKQLSFSLSAEPIQGPYK | 12,AFB1_Dialdehyde[K]; | 2.1 | 0.0 | 0.0 | 4.65E-02 |
| P32038 | LGGQEAMAHAR | 0,Ethylphosphate[AnyN-term]; | 2.1 | 0.0 | 0.0 | 4.65E-02 |
| Q6IE52 | QWQDVKTENGLKQLSFSLSAEPIQGPYK | 12,AFB1_Dialdehyde[K]; | 2.1 | 0.0 | 0.0 | 4.65E-02 |
| P14740 | KYPLLIDVYAGPCSQK | 13,Carbamidomethyl[C]; | 15.4 | 0.0 | 0.0 | 4.65E-02 |
| P07647 | AYDHNNDLMLLHLSK | 5,Deamidated[N]; | 8.1 | 0.0 | 0.0 | 4.68E-02 |
| Q01177 | PWCFTTDPNKR | 3,Carbamidomethyl[C]; | 3.1 | 0.0 | 0.0 | 4.71E-02 |
| P02770 | HPYFYAPELLYYAEKY | 16,Tyr->Arg[Y]; | 3.7 | 0.0 | 0.0 | 4.71E-02 |
| P54311 | ELAGHTGYLSCCR | 11,Carbamidomethyl[C];12,Carbamidomethyl[C]; | 3.7 | 0.0 | 0.0 | 4.71E-02 |
| P02782 | KLSLCLLIMLAVCCYEANASQICELVAHETISFLMK | 5,Cys->Ala[C]; | 3.4 | 0.0 | 0.0 | 4.72E-02 |
| P81828 | QKCQTYPDEICAWVVVTTR | 11,DiDehydro[C]; | 73.3 | 0.0 | 0.0 | 4.81E-02 |
| P02770 | RHPYFYAPELLYYAEK | 13,Tyr->Trp[Y]; | 6.3 | 0.0 | 0.0 | 4.83E-02 |
| P02770 | KAADKDNCFATEGPNLVAR | 8,Cys->Dha[C]; | 3.9 | 0.0 | 0.0 | 4.88E-02 |
| P02770 | FATEGPNLVAR | 0,Diethylphosphate[AnyN-term]; | 2.6 | 0.0 | 0.0 | 4.88E-02 |
| P02780 | VKPYVQDHFTEK | 0,Benzoyl[AnyN-term]; | 2.6 | 0.0 | 0.0 | 4.88E-02 |
| P02782 | QICELVAHETISFLMK | 0,Gln->pyro-Glu[AnyN-termQ];3,Carbamidomethyl[C]; | 1.3 | 0.0 | 0.0 | 4.88E-02 |
| P08723 | FLLEDGEHVTQVSGTR | 0,ICPL_13C(6)2H(4)[AnyN-term]; | 2.6 | 0.0 | 0.0 | 4.88E-02 |
| P10960 | SSLSASCKEVVDSYLPVILDMIK | 7,Nethylmaleimide[C]; | 1.3 | 0.0 | 0.0 | 4.88E-02 |
| P04276 | DLCGQSATQAMDQYTFELSR | 3,Carbamidomethyl[C]; | 16.4 | 0.0 | 0.0 | 4.93E-02 |
| P83121 | HICQTYPDEICAWVVVTTR | 3,HN3_mustard[C]; | 46.1 | 0.0 | 0.0 | 4.95E-02 |
| P07522 | LSVIEMADLDGSKR | 1,Xle->Cys[L]; | 4.6 | 0.0 | 0.0 | 4.95E-02 |
| P07647 | QRAYDHNNDLMLLHLSKPADITGGVK | 0,Gln->pyro-Glu[AnyN-termQ]; | 8.1 | 0.0 | 0.0 | 4.96E-02 |
| P23785 | VHCCPHGASCDLVHTR | 3,Carbamidomethyl[C];4,Carbamidomethyl[C];10,Carbamidomethyl[C]; | 3.3 | 0.0 | 0.0 | 4.97E-02 |
| P02761 | TPEDGEYFVEYDGGNTFTILK | 10,Cation_Na[E]; | 96.0 | 0.0 | 0.0 | 4.98E-02 |
| P02770 | APQVSTPTLVEAAR | 2,Pro->Cys[P]; | 2.0 | 0.0 | 0.0 | 4.98E-02 |
| P28826 | MTDDNGSYLWDRPSK | 5,Deamidated[N]; | 2.0 | 0.0 | 0.0 | 4.98E-02 |
| P00689 | MAVGFMLAHPYGFTR | 1,Oxidation[M]; | 13.1 | 0.2 | 0.0 | 2.37E-02 |
| P02780 | QCFLDQTDK | 3,Phe->Trp[F]; | 10.1 | 0.3 | 0.0 | 4.49E-02 |
| P02770 | RHPDYSVSLLLR | 0,Formyl[AnyN-term]; | 140.4 | 4.8 | 0.0 | 2.94E-02 |
| P36373 | AQHRLVSQSFPHPDYKPFLMR | 5,Xle->Cys[L]; | 17.3 | 0.7 | 0.0 | 2.56E-02 |
| P31211 | DVPLTLTMVHK | 8,Oxidation[M]; | 16.6 | 0.7 | 0.0 | 3.96E-03 |
| P27590 | DWMSIVTPAR | 3,Oxidation[M]; | 194.4 | 14.8 | 0.1 | 5.60E-03 |
| P10960 | SLPCDICK | 4,Carbamidomethyl[C];7,Carbamidomethyl[C]; | 5.9 | 0.5 | 0.1 | 2.69E-02 |
| P22282 | TPGETMYYISLPGSVR | 4,Cation_Na[E]; | 33.7 | 4.0 | 0.1 | 2.80E-02 |
| P28826 | QELSIGTNCDR | 9,Carbamidomethyl[C]; | 4.9 | 0.8 | 0.2 | 4.48E-02 |
| P27590 | CPHTEDTTIQVTENGESSQAR | 1,Carbamidomethyl[C]; | 777.4 | 188.5 | 0.2 | 4.20E-02 |
| Q01177 | NPDGDVNGPWCYTMNPR | 11,Carbamidomethyl[C]; | 12.0 | 3.0 | 0.3 | 2.75E-02 |
| P02770 | SIHTLFGDK | 0,Formyl[AnyN-term]; | 22.6 | 5.8 | 0.3 | 4.45E-02 |
| P05544 | DEELSCSVLELK | 6,Carbamidomethyl[C]; | 48.3 | 14.3 | 0.3 | 4.46E-03 |
| P05545 | DEELSCSVLELK | 6,Carbamidomethyl[C]; | 48.3 | 14.3 | 0.3 | 4.46E-03 |
| P09005 | DEELSCSVLELK | 6,Carbamidomethyl[C]; | 48.3 | 14.3 | 0.3 | 4.46E-03 |
| Q63556 | DEELSCSVLELK | 6,Carbamidomethyl[C]; | 48.3 | 14.3 | 0.3 | 4.46E-03 |
| P27590 | CQLQSLGFMK | 0,Pyro-carbamidomethyl[AnyN-termC]; | 83.9 | 26.3 | 0.3 | 3.27E-02 |
| P02761 | VFMQHIDVLENSLGFK | 11,Deamidated[N]; | 46.7 | 17.2 | 0.4 | 3.27E-02 |
| P18292 | REECSIPVCGQEGR | 4,Carbamidomethyl[C];9,Carbamidomethyl[C]; | 22.7 | 8.5 | 0.4 | 4.57E-02 |
| P02770 | AETFTFHSDICTLPDKEK | 11,Carbamidomethyl[C]; | 1158.0 | 441.7 | 0.4 | 2.78E-02 |
| P27590 | MAETCVPVLR | 1,Oxidation[M];5,Carbamidomethyl[C]; | 145.7 | 74.3 | 0.5 | 5.97E-03 |
| P36953 | LCFFYNK | 2,Carbamidomethyl[C]; | 7.4 | 15.0 | 2.0 | 2.37E-02 |
| P36953 | SLAMVQQECK | 9,Carbamidomethyl[C]; | 5.4 | 12.0 | 2.2 | 1.79E-03 |
| P13635 | DCNKPSPDDDIQDR | 2,Carbamidomethyl[C]; | 7.7 | 19.0 | 2.5 | 1.33E-02 |
| P14046 | ISLCHGNPTFSSETK | 4,Carbamidomethyl[C]; | 15.4 | 39.8 | 2.6 | 3.68E-04 |
| P02764 | QQLELEKETK | 0,Gln->pyro-Glu[AnyN-termQ]; | 16.3 | 42.3 | 2.6 | 5.13E-04 |
| P02770 | QACCDKPVLQK | 4,maleimide[C]; | 4.1 | 11.8 | 2.9 | 1.52E-02 |
| P20760 | ETWQQGNTFTCSVLHEGLHNHHTEK | 11,Carbamidomethyl[C]; | 30.3 | 95.5 | 3.2 | 1.31E-04 |
| P12346 | INHCKFDEFFSQGCAPGYK | 4,Carbamidomethyl[C];14,Carbamidomethyl[C]; | 14.1 | 48.0 | 3.4 | 2.42E-02 |
| Q01177 | TPENFPCK | 7,Carbamidomethyl[C]; | 10.1 | 34.7 | 3.4 | 3.28E-05 |
| P02770 | LPCVEDYLSAILNR | 3,Crotonaldehyde[C]; | 20.6 | 76.8 | 3.7 | 2.84E-05 |
| P02770 | RHPDYSVSLLLR | 8,Formyl[S](Ser->Asp[S]); | 7.3 | 29.2 | 4.0 | 5.97E-03 |
| P36953 | VYKDLLEDCCNRK | 9,Carbamidomethyl[C];10,Carbamidomethyl[C]; | 2.1 | 8.7 | 4.0 | 3.75E-03 |
| P02770 | APQVSTPTLVEAAR | 0,ISD_z+2_ion[AnyN-term]; | 16.9 | 75.2 | 4.5 | 1.91E-04 |
| P12346 | TSYQDCIK | 6,Carbamidomethyl[C]; | 6.4 | 32.2 | 5.0 | 4.23E-04 |
| P02770 | LPCVEDYLSAILNR | 3,CarbamidomethylDTT[C]; | 3.0 | 15.2 | 5.1 | 2.19E-03 |
| P20059 | DYFISCPGR | 6,Carbamidomethyl[C]; | 14.1 | 79.0 | 5.6 | 2.26E-03 |
| P02770 | RHPYFYAPELLYYAEK | 11,Xle->His[L]; | 0.9 | 4.8 | 5.6 | 3.99E-02 |
| Q03626 | SSGSLFNNAMK | 10,Oxidation[M]; | 0.3 | 1.7 | 5.8 | 2.38E-02 |
| Q6IE52 | SSGSLFNNAMK | 10,Oxidation[M]; | 0.3 | 1.7 | 5.8 | 2.38E-02 |
| P05371 | TQQYNELLHSLQSK | 0,Succinyl_2H(4)[AnyN-term]; | 2.1 | 13.3 | 6.2 | 2.52E-02 |
| P06866 | MPICLPSKDYVAPGR | 4,Hydroxycinnamyl[C]; | 0.9 | 5.3 | 6.2 | 3.17E-02 |
| P02770 | DDNPNLPPFQRPEAEAMCTSFQENPTSFLGHYLHEVAR | 18,Carbamidomethyl[C]; | 46.1 | 309.2 | 6.7 | 1.12E-03 |
| P98158 | CADGSSCINSR | 1,Carbamidomethyl[C];7,Carbamidomethyl[C]; | 0.7 | 4.8 | 6.8 | 3.15E-04 |
| P02770 | TSFLGHYLHEVAR | 0,Acetyl_2H(3)[AnyN-term]; | 4.4 | 30.0 | 6.8 | 4.85E-02 |
| P02770 | PCVEDYLSAILNR | 2,MTSL[C]; | 1.0 | 8.3 | 8.3 | 4.60E-02 |
| P02770 | LVQEVTDFAK | 7,Cation_Na[D]; | 8.6 | 71.5 | 8.3 | 1.48E-06 |
| P07647 | NLVKDEPFAQR | 0,Ethylphosphate[AnyN-term]; | 0.9 | 7.3 | 8.6 | 2.21E-02 |
| P02770 | LPCVEDYLSAILNR | 3,Propionamide[C]; | 0.6 | 5.3 | 9.3 | 7.61E-03 |
| P02770 | FLGHYLHEVAR | 2,Xle->Cys[L]; | 0.4 | 4.0 | 9.3 | 4.70E-02 |
| Q01177 | CQSWVSMTPHSHSK | 1,Carbamidomethyl[C]; | 2.9 | 27.2 | 9.5 | 3.59E-03 |
| P01048 | EIPVDSPELK | 0,C+12[AnyN-term]; | 1.3 | 12.3 | 9.6 | 6.04E-05 |
| P02770 | AENCDKSIHTLFGDK | 4,DiDehydro[C]; | 0.4 | 4.2 | 9.7 | 3.87E-02 |
| P02761 | EKIEENGSMR | 6,Deamidated[N]; | 30.0 | 314.5 | 10.5 | 6.70E-03 |
| P02770 | SFQENPTSFLGHYLHEVAR | 1,PhosphoHexNAc[S]; | 1.6 | 20.3 | 12.9 | 7.61E-04 |
| P02770 | LRDNYGELADCCAK | 11,Diethylphosphate[C]; | 1.3 | 17.8 | 13.9 | 1.08E-03 |
| P20760 | ETWQQGNTFTCSVLHEGLHNHHTEK | 11,Carbamidomethyl[C];20,Deamidated[N]; | 1.6 | 22.0 | 14.0 | 4.05E-04 |
| P36373 | NNLLEDEPFAQHR | 3,Xle->Cys[L]; | 0.9 | 13.5 | 15.8 | 3.86E-03 |
| P36374 | NNLLEDEPFAQHR | 3,Xle->Cys[L]; | 0.9 | 13.5 | 15.8 | 3.86E-03 |
| P20760 | GPQVYTMAPPK | 2,Pro->Cys[P]; | 0.7 | 11.7 | 16.3 | 7.19E-03 |
| P02770 | RHPYFYAPELLYYAEK | 9,Cation_Ca[II][E]; | 2.1 | 36.3 | 17.0 | 5.35E-05 |
| P02770 | NPNLPPFQRPEAEAMCTSFQENPTSFLGHYLHEVAR | 16,Carbamidomethyl[C]; | 0.4 | 8.0 | 18.7 | 6.62E-03 |
| P02770 | CPYEEHIK | 1,CarbamidomethylDTT[C]; | 0.9 | 16.8 | 19.6 | 1.57E-05 |
| P02770 | CENQATISSK | 1,ICAT-H[C]; | 0.4 | 8.7 | 20.2 | 4.50E-02 |
| P09006 | DSLRPSMIDELYLPK | 7,Oxidation[M]; | 1.6 | 33.0 | 21.0 | 1.03E-03 |
| P07647 | VSQSFQHPDYIPVFMR | 0,Iodoacetanilide[AnyN-term]; | 0.4 | 10.0 | 23.3 | 3.33E-02 |
| P02761 | LCEAHGITR | 2,CarbamidomethylDTT[C]; | 3.4 | 88.8 | 25.9 | 1.31E-02 |
| Q9EQV9 | FIKPTCAEALAAVSK | 6,Carbamidomethyl[C]; | 0.4 | 13.0 | 30.3 | 1.72E-02 |
| P23680 | ALNYEINGYVVIKPR | 7,Deamidated[N]; | 0.9 | 26.8 | 31.3 | 1.84E-04 |
| P17475 | EDAQETDTSQQDQSPTYR | 5,Cation_Ca[II][E]; | 0.1 | 4.5 | 31.5 | 7.72E-03 |
| Q9QX79 | KIHSMCPDCPHPVDLSAPSVLEAATESLAK | 5,Oxidation[M];6,Carbamidomethyl[C];9,Carbamidomethyl[C]; | 2.3 | 73.8 | 32.3 | 5.36E-03 |
| P02770 | ENPTSFLGHYLHEVAR | 1,Glu->His[E]; | 0.4 | 14.3 | 33.4 | 1.32E-02 |
| P12346 | DGGGDVAFVK | 5,Cation_Na[D]; | 0.1 | 4.8 | 33.8 | 7.92E-03 |
| P02770 | CDKPVLQK | 1,Carbamidomethyl[C]; | 0.1 | 5.0 | 35.0 | 5.04E-03 |
| P02770 | NPTSFLGHYLHEVAR | 0,Succinyl_2H(4)[AnyN-term]; | 0.1 | 5.2 | 36.2 | 1.16E-02 |
| P06866 | PICLPSKDYVAPGR | 3,Carbamidomethyl[C]; | 1.3 | 49.2 | 38.2 | 8.85E-03 |
| P17475 | GTEAAGATVVEAVPMSLPPQVK | 2,Dehydrated[T]; | 0.4 | 22.8 | 53.3 | 7.03E-03 |
| P05545 | IAELFSELDER | 3,Glu->His[E]; | 0.4 | 23.0 | 53.7 | 3.66E-02 |
| Q03626 | DMGLTAFTNLK | 2,Oxidation[M]; | 0.4 | 27.0 | 63.0 | 2.41E-04 |
| Q6IE52 | DMGLTAFTNLK | 2,Oxidation[M]; | 0.4 | 27.0 | 63.0 | 2.41E-04 |
| P12346 | TAGWNIPMGLLFSR | 0,C+12[AnyN-term]; | 0.4 | 32.3 | 75.4 | 9.84E-04 |
| Q03626 | GGEFEMMPLGVNK | 6,Oxidation[M]; | 0.3 | 23.7 | 82.8 | 3.74E-04 |
| P22283 | LVNCPFEEQTEQLKR | 4,Carbamidomethyl[C]; | 0.3 | 51.2 | 179.1 | 6.93E-04 |
| P02770 | DNCFATEGPNLVAR | 2,Hex(3)[N]; | 0.0 | 83.7 | #DIV/0! | 5.89E-07 |
| P02770 | AADKDNCFATEGPNLVAR | 7,Ub-Br2[C]; | 0.0 | 43.8 | #DIV/0! | 7.96E-07 |
| P01048 | FSVATQICNITPGK | 8,Carbamidomethyl[C]; | 0.0 | 40.0 | #DIV/0! | 1.86E-06 |
| P08932 | FSVATQICNITPGK | 8,Carbamidomethyl[C]; | 0.0 | 40.0 | #DIV/0! | 1.86E-06 |
| P06866 | YVMLPVADQEKCELHYEK | 12,Carbamidomethyl[C]; | 0.0 | 57.2 | #DIV/0! | 2.79E-06 |
| P04276 | YCSSQIDAEMR | 2,Carbamidomethyl[C]; | 0.0 | 33.5 | #DIV/0! | 3.59E-06 |
| P13635 | EMGPTYADPVCLSK | 11,Carbamidomethyl[C]; | 0.0 | 39.2 | #DIV/0! | 3.96E-06 |
| P06866 | GAVSPVGVQPILNK | 2,Carbonyl[A]; | 0.0 | 40.5 | #DIV/0! | 6.81E-06 |
| P02770 | RHPDYSVSLLLR | 1,Arg->Tyr[R]; | 0.0 | 48.2 | #DIV/0! | 6.89E-06 |
| P05544 | MQQVESSLQPETLKK | 5,Cation_Na[E]; | 0.0 | 31.2 | #DIV/0! | 9.43E-06 |
| P05545 | MQQVESSLQPETLKK | 5,Cation_Na[E]; | 0.0 | 31.2 | #DIV/0! | 9.43E-06 |
| P09005 | MQQVESSLQPETLKK | 5,Cation_Na[E]; | 0.0 | 31.2 | #DIV/0! | 9.43E-06 |
| P12346 | DFQLFGSPLGK | 1,Cation_Na[D]; | 0.0 | 15.0 | #DIV/0! | 1.11E-05 |
| P15083 | NKETCDVIINTLGK | 5,Carbamidomethyl[C]; | 0.0 | 13.5 | #DIV/0! | 1.39E-05 |
| P02770 | TEDQLKTVMGDFAQFVDK | 0,Phenylisocyanate_2H(5)[AnyN-term]; | 0.0 | 9.3 | #DIV/0! | 1.39E-05 |
| P02770 | QRFPNAEFAEITK | 0,Pyridylacetyl[AnyN-term](Phenylisocyanate[AnyN-term]); | 0.0 | 13.8 | #DIV/0! | 1.82E-05 |
| P36953 | IANDAIQDMLCDMK | 11,Carbamidomethyl[C]; | 0.0 | 18.5 | #DIV/0! | 1.89E-05 |
| P02770 | QTALAELVKHKPK | 0,Carbamyl[AnyN-term]; | 0.0 | 24.8 | #DIV/0! | 1.89E-05 |
| P02770 | QTALAELVK | 0,Gln->pyro-Glu[AnyN-termQ]; | 0.0 | 43.2 | #DIV/0! | 1.91E-05 |
| P05545 | VLDVDETGTEGAAATAVTAALK | 0,Propyl_2H(6)[AnyN-term]; | 0.0 | 25.5 | #DIV/0! | 2.05E-05 |
| P12346 | TCHLAQAPNHVVVSR | 1,Thr->Cys[T]; | 0.0 | 8.2 | #DIV/0! | 2.35E-05 |
| P17475 | VFNNDADLSGITEDAPLK | 7,Cation_Na[D]; | 0.0 | 30.8 | #DIV/0! | 2.39E-05 |
| P02764 | EFQTTDDQCVYNFTHLGVQR | 9,Carbamidomethyl[C]; | 0.0 | 34.3 | #DIV/0! | 3.12E-05 |
| P02770 | MSQRFPNAEFAEITK | 0,Phenylisocyanate_2H(5)[AnyN-term]; | 0.0 | 287.3 | #DIV/0! | 4.19E-05 |
| P12346 | TCHLAQAPNHVVVSR | 2,CarbamidomethylDTT[C]; | 0.0 | 39.7 | #DIV/0! | 4.64E-05 |
| P02761 | VFMQHIDVLENSLGFK | 3,Oxidation[M];11,Deamidated[N]; | 0.0 | 22.5 | #DIV/0! | 4.75E-05 |
| P09006 | MQQVEASLQPETLRR | 1,Met->His[M]; | 0.0 | 18.8 | #DIV/0! | 4.78E-05 |
| P07314 | NIDQVVTAGLK | 0,Methyl[AnyN-term]; | 0.0 | 14.8 | #DIV/0! | 5.35E-05 |
| P02770 | RHPYFYAPELLYYAEK | 12,Dehydrated[Y]; | 0.0 | 72.3 | #DIV/0! | 5.41E-05 |
| P02761 | LNGDWFSIVVASNKR | 0,Methyl_2H(2)[AnyN-term]; | 0.0 | 40.5 | #DIV/0! | 5.66E-05 |
| P02770 | RHPDYSVSLLLR | 5,Dehydrated[Y]; | 0.0 | 191.7 | #DIV/0! | 6.28E-05 |
| P12346 | DQYELLCLDNTR | 7,Carbamidomethyl[C]; | 0.0 | 62.5 | #DIV/0! | 8.42E-05 |
| P02761 | VFMQHIDVLENSLGFK | 3,Oxidation[M]; | 0.0 | 970.7 | #DIV/0! | 8.46E-05 |
| P20759 | DVLTITLTPK | 1,Dehydrated[D]; | 0.0 | 25.8 | #DIV/0! | 1.01E-04 |
| P20760 | DVLTITLTPK | 1,Dehydrated[D]; | 0.0 | 25.8 | #DIV/0! | 1.01E-04 |
| P20759 | NTPPTMDTDGSYFLYSK | 6,Oxidation[M]; | 0.0 | 186.8 | #DIV/0! | 1.13E-04 |
| P20760 | NTPPTMDTDGSYFLYSK | 6,Oxidation[M]; | 0.0 | 186.8 | #DIV/0! | 1.13E-04 |
| P12346 | WCAVSEHENTK | 2,Carbamidomethyl[C]; | 0.0 | 83.7 | #DIV/0! | 1.16E-04 |
| Q64240 | TIAACNLPIVQGPCR | 5,Carbamidomethyl[C];14,Carbamidomethyl[C]; | 0.0 | 38.0 | #DIV/0! | 1.17E-04 |
| P12346 | KTSYQDCIK | 7,Carbamidomethyl[C]; | 0.0 | 8.7 | #DIV/0! | 1.18E-04 |
| P02770 | AADKDNCFATEGPNLVAR | 7,Carbamidomethyl[C]; | 0.0 | 605.7 | #DIV/0! | 1.19E-04 |
| P20762 | VVSTLHIQHQDWMSGK | 3,Methylamine[S]; | 0.0 | 7.7 | #DIV/0! | 1.22E-04 |
| P02770 | LLECADDRAELAK | 4,Carbamidomethyl[C]; | 0.0 | 27.7 | #DIV/0! | 1.25E-04 |
| P02770 | AADKDNCFATEGPNLVAR | 6,Deamidated[N];7,Carbamidomethyl[C]; | 0.0 | 59.7 | #DIV/0! | 1.27E-04 |
| Q01177 | TAVTAAGTPCQEWAAQEPHSHR | 10,Carbamidomethyl[C]; | 0.0 | 50.5 | #DIV/0! | 1.41E-04 |
| P12346 | WCAVSEHENTK | 2,Carbamidomethyl[C];9,Deamidated[N]; | 0.0 | 8.0 | #DIV/0! | 1.46E-04 |
| P15684 | QVTPLFAYFK | 0,Gln->pyro-Glu[AnyN-termQ]; | 0.0 | 7.2 | #DIV/0! | 1.56E-04 |
| P02770 | CCAEGDPPACYGTVLAEFQPLVEEPK | 1,NEMsulfur[C]; | 0.0 | 47.2 | #DIV/0! | 1.56E-04 |
| P02770 | LRDNYGELADCCAK | 11,DTT[C]; | 0.0 | 18.7 | #DIV/0! | 1.59E-04 |
| P02770 | LQACCDKPVLQK | 4,maleimide[C]; | 0.0 | 16.7 | #DIV/0! | 1.71E-04 |
| P20760 | CKVNSGAFPAPIEK | 0,Pyro-carbamidomethyl[AnyN-termC]; | 0.0 | 45.7 | #DIV/0! | 1.73E-04 |
| P02770 | KDNCFATEGPNLVAR | 4,HexNAc[C]; | 0.0 | 22.8 | #DIV/0! | 1.83E-04 |
| P02770 | TFHSDICTLPDKEK | 7,Acetyl[C]; | 0.0 | 7.5 | #DIV/0! | 1.87E-04 |
| P20059 | NGTAHGNSTHPMHSRCNADPGLSALLSDHR | 16,Cys->Arg[C]; | 0.0 | 12.8 | #DIV/0! | 2.01E-04 |
| P02770 | LQACCDKPVLQK | 4,Carbamidomethyl[C];5,Carbamidomethyl[C]; | 0.0 | 283.5 | #DIV/0! | 2.02E-04 |
| P09006 | AVLDVAETGTEAAAATGVK | 7,Cation_Na[E]; | 0.0 | 28.5 | #DIV/0! | 2.06E-04 |
| Q62930 | AIEDYVNEFSAR | 0,C+12[AnyN-term]; | 0.0 | 18.7 | #DIV/0! | 2.07E-04 |
| P02770 | HPYFYAPELLYYAEK | 0,TMAB_2H(9)[AnyN-term]; | 0.0 | 3.8 | #DIV/0! | 2.13E-04 |
| P02770 | PDYSVSLLLR | 1,Pro->Cys[P]; | 0.0 | 27.8 | #DIV/0! | 2.13E-04 |
| P02770 | QEPERNECFLQHKDDNPNLPPFQRPEAEAMCTSFQENPTSFLGHYLHEVAR | 8,GG[C](Dicarbamidomethyl[C]); | 0.0 | 81.8 | #DIV/0! | 2.16E-04 |
| P02770 | CPYEEHIKLVQEVTDFAK | 0,Succinyl_2H(4)[AnyN-term]; | 0.0 | 10.0 | #DIV/0! | 2.31E-04 |
| P12346 | SSTKDLLFR | 0,Carbamyl[AnyN-term]; | 0.0 | 26.7 | #DIV/0! | 2.43E-04 |
| P02770 | KQTALAELVKHKPK | 10,Carbamyl[K]; | 0.0 | 7.7 | #DIV/0! | 2.69E-04 |
| P02770 | EFKAETFTFHSDICTLPDK | 14,Carbamidomethyl[C]; | 0.0 | 13.3 | #DIV/0! | 2.78E-04 |
| P02770 | VFLGTFLYEYSR | 0,ICPL_13C(6)[AnyN-term]; | 0.0 | 22.7 | #DIV/0! | 2.79E-04 |
| P02770 | LRDNYGELADCCAK | 11,Hydroxycinnamyl[C]; | 0.0 | 12.3 | #DIV/0! | 2.95E-04 |
| P12346 | EYLGAEYLQAVGNIRK | 0,Glu->pyro-Glu+Methyl[AnyN-termE]; | 0.0 | 5.8 | #DIV/0! | 2.95E-04 |
| P04276 | EVVSLTEECCAEGADPNCYDTR | 9,Carbamidomethyl[C];10,Carbamidomethyl[C];18,Carbamidomethyl[C]; | 0.0 | 45.2 | #DIV/0! | 3.26E-04 |
| P02770 | GLVLIAFSQYLQK | 14,Cation_Na[AnyC-term]; | 0.0 | 9.0 | #DIV/0! | 3.29E-04 |
| Q64240 | AFAELWAFDAAQGK | 1,Carbonyl[A]; | 0.0 | 5.3 | #DIV/0! | 3.36E-04 |
| Q9R0T4 | DTANWLEINPETGVISTR | 1,Cation_Na[D]; | 0.0 | 8.2 | #DIV/0! | 3.44E-04 |
| P20059 | ECQSEGVLFFQGNRK | 0,TMAB_2H(9)[AnyN-term]; | 0.0 | 12.8 | #DIV/0! | 3.51E-04 |
| Q01177 | MRDVILFEK | 1,Oxidation[M]; | 0.0 | 14.5 | #DIV/0! | 3.53E-04 |
| P01048 | KTEEDLCVGCFQPIPMDSSDLKPVLK | 7,Carbamidomethyl[C];10,Carbamidomethyl[C]; | 0.0 | 48.8 | #DIV/0! | 3.54E-04 |
| P08932 | KTEEDLCVGCFQPIPMDSSDLKPVLK | 7,Carbamidomethyl[C];10,Carbamidomethyl[C]; | 0.0 | 48.8 | #DIV/0! | 3.54E-04 |
| P02770 | CFATEGPNLVAR | 0,ICPL_13C(6)[AnyN-term]; | 0.0 | 10.3 | #DIV/0! | 3.91E-04 |
| P14046 | SGVKEEHSFTVMEFVLPR | 0,Succinyl_2H(4)[AnyN-term]; | 0.0 | 19.3 | #DIV/0! | 3.94E-04 |
| Q03626 | SGVKEEHSFTVMEFVLPR | 0,Succinyl_2H(4)[AnyN-term]; | 0.0 | 19.3 | #DIV/0! | 3.94E-04 |
| Q6IE52 | SGVKEEHSFTVMEFVLPR | 0,Succinyl_2H(4)[AnyN-term]; | 0.0 | 19.3 | #DIV/0! | 3.94E-04 |
| P02770 | AADKDNCFATEGPNLVAR | 7,NEM_2H(5)+H2O[C]; | 0.0 | 60.0 | #DIV/0! | 3.96E-04 |
| P20759 | CSVLHEGLHNHHTEK | 1,cysTMT[C]; | 0.0 | 49.7 | #DIV/0! | 4.01E-04 |
| P20760 | CSVLHEGLHNHHTEK | 1,cysTMT[C]; | 0.0 | 49.7 | #DIV/0! | 4.01E-04 |
| P02770 | TFTFHSDICTLPDKEK | 9,Carbamidomethyl[C]; | 0.0 | 66.3 | #DIV/0! | 4.05E-04 |
| P02770 | CVEDYLSAILNR | 1,2-dimethylsuccinyl[C]; | 0.0 | 26.8 | #DIV/0! | 4.77E-04 |
| P02770 | PQVSTPTLVEAAR | 0,Propionyl_13C(3)[AnyN-term]; | 0.0 | 19.8 | #DIV/0! | 4.83E-04 |
| P02770 | QEPERNECFLQHKDDNPNLPPFQRPEAEAMCTSFQENPTSFLGHYLHEVAR | 31,GG[C](Dicarbamidomethyl[C]); | 0.0 | 115.7 | #DIV/0! | 5.00E-04 |
| P20760 | CKVNSGAFPAPIEK | 1,Carbamyl[C]; | 0.0 | 15.5 | #DIV/0! | 5.02E-04 |
| P12346 | ASDSSINWNNLK | 0,C+12[AnyN-term]; | 0.0 | 29.5 | #DIV/0! | 5.32E-04 |
| P13635 | KALYSEYTDGTFTK | 0,Methyl_2H(2)[AnyN-term]; | 0.0 | 9.8 | #DIV/0! | 5.32E-04 |
| P00787 | EQWSNCPTIAQIR | 6,Carbamidomethyl[C]; | 0.0 | 19.0 | #DIV/0! | 5.70E-04 |
| P12346 | SHAVVARNGDGKEDLIWEILK | 0,NIC[AnyN-term](ICPL[AnyN-term]); | 0.0 | 6.0 | #DIV/0! | 5.73E-04 |
| P27590 | WRSTDYGAGYSCDSDMHGWYR | 6,Dehydrated[Y]; | 0.0 | 11.2 | #DIV/0! | 5.84E-04 |
| P02770 | RHPDYSVSLLLR | 3,Pro->Cys[P]; | 0.0 | 42.2 | #DIV/0! | 6.64E-04 |
| P02770 | LPCVEDYLSAILNR | 3,monomethylphosphothione[C]; | 0.0 | 23.2 | #DIV/0! | 6.69E-04 |
| P02770 | AADKDNCFATEGPNLVAR | 7,Cys->Arg[C]; | 0.0 | 15.3 | #DIV/0! | 7.16E-04 |
| P02770 | RHPDYSVSLLLR | 4,Dehydrated[D]; | 0.0 | 8.2 | #DIV/0! | 7.22E-04 |
| P14046 | VYHKEESSCIHSSCTAER | 0,Methyl_2H(2)[AnyN-term];9,Carbamidomethyl[C];14,Carbamidomethyl[C]; | 0.0 | 6.8 | #DIV/0! | 7.92E-04 |
| Q03626 | VYHKEESSCIHSSCTAER | 0,Methyl_2H(2)[AnyN-term];9,Carbamidomethyl[C];14,Carbamidomethyl[C]; | 0.0 | 6.8 | #DIV/0! | 7.92E-04 |
| P02770 | QEPERNECFLQHK | 0,Gln->pyro-Glu[AnyN-termQ];8,Carbamidomethyl[C]; | 0.0 | 17.0 | #DIV/0! | 9.06E-04 |
| P15083 | ETCDVIINTLGK | 3,Carbamidomethyl[C]; | 0.0 | 18.8 | #DIV/0! | 9.14E-04 |
| P02770 | LVQEVTDFAK | 1,Xle->Cys[L]; | 0.0 | 42.0 | #DIV/0! | 9.21E-04 |
| P20760 | CKVNSGAFPAPIEK | 1,Acetyl[C]; | 0.0 | 23.3 | #DIV/0! | 9.42E-04 |
| P02761 | CEAHGITR | 0,Pyro-carbamidomethyl[AnyN-termC]; | 0.0 | 11.8 | #DIV/0! | 9.44E-04 |
| P02770 | LQACCDKPVLQK | 5,maleimide[C]; | 0.0 | 17.2 | #DIV/0! | 9.45E-04 |
| P12346 | SHAVVARNGDGKEDLIWEILK | 10,Cation_Ag[D]; | 0.0 | 17.8 | #DIV/0! | 9.84E-04 |
| P02770 | QENPTSFLGHYLHEVAR | 0,Propyl[AnyN-term]; | 0.0 | 16.8 | #DIV/0! | 1.01E-03 |
| P20759 | SNSMVTLGCLVK | 4,Oxidation[M];9,Carbamidomethyl[C]; | 0.0 | 21.2 | #DIV/0! | 1.08E-03 |
| P20760 | SNSMVTLGCLVK | 4,Oxidation[M];9,Carbamidomethyl[C]; | 0.0 | 21.2 | #DIV/0! | 1.08E-03 |
| P14046 | MLIYTILPDGEVIADSVK | 3,Xle->Ser[I]; | 0.0 | 1.5 | #DIV/0! | 1.11E-03 |
| P20059 | ELGSPPGISLDTIDAAFSCPGSSK | 19,Carbamidomethyl[C]; | 0.0 | 24.0 | #DIV/0! | 1.11E-03 |
| Q6IE52 | MLIYTILPDGEVIADSVK | 3,Xle->Ser[I]; | 0.0 | 1.5 | #DIV/0! | 1.11E-03 |
| P17475 | VFNNDADLSGITEDAPLK | 1,Val->Met[V]; | 0.0 | 14.8 | #DIV/0! | 1.12E-03 |
| P12346 | CALSHQER | 0,Biotin[AnyN-term]; | 0.0 | 6.3 | #DIV/0! | 1.17E-03 |
| P20759 | MNGQPQENYK | 1,Oxidation[M]; | 0.0 | 18.3 | #DIV/0! | 1.18E-03 |
| P20760 | MNGQPQENYK | 1,Oxidation[M]; | 0.0 | 18.3 | #DIV/0! | 1.18E-03 |
| P02770 | CTSFQENPTSFLGHYLHEVAR | 0,Methyl_2H(2)[AnyN-term];7,Deamidated[N]; | 0.0 | 4.7 | #DIV/0! | 1.26E-03 |
| P02782 | CVDQMSNGDR | 1,Carbamidomethyl[C];5,Oxidation[M]; | 0.0 | 15.3 | #DIV/0! | 1.29E-03 |
| P02770 | DQLKTVMGDFAQFVDK | 5,HexNAc(1)NeuAc(1)[T]; | 0.0 | 9.7 | #DIV/0! | 1.33E-03 |
| P02770 | FKDLGEQHFK | 0,Methyl[AnyN-term]; | 0.0 | 20.8 | #DIV/0! | 1.39E-03 |
| P05544 | STMEEILEGLK | 0,Ethylphosphate[AnyN-term]; | 0.0 | 7.5 | #DIV/0! | 1.40E-03 |
| P02770 | PTSFLGHYLHEVAR | 0,Methyl[AnyN-term]; | 0.0 | 17.8 | #DIV/0! | 1.41E-03 |
| P02770 | RMSQRFPNAEFAEITK | 4,Gln->Pro[Q]; | 0.0 | 17.2 | #DIV/0! | 1.42E-03 |
| P07632 | SLSGEHSIIGR | 0,ISD_z+2_ion[AnyN-term]; | 0.0 | 1.3 | #DIV/0! | 1.46E-03 |
| P17475 | LAEDAQETDTSQQDQSPTYR | 3,phosphoRibosyl[E]; | 0.0 | 6.7 | #DIV/0! | 1.46E-03 |
| P24090 | CNLLAEK | 1,Carbamidomethyl[C]; | 0.0 | 24.2 | #DIV/0! | 1.51E-03 |
| P02770 | SIHTLFGDKLCAIPK | 11,Carbamidomethyl[C]; | 0.0 | 108.2 | #DIV/0! | 1.52E-03 |
| P02761 | DLSSDIKEKFAK | 0,Carbamyl[AnyN-term]; | 0.0 | 13.5 | #DIV/0! | 1.57E-03 |
| P02770 | TEDQLKTVMGDFAQFVDK | 0,Methyl_2H(2)[AnyN-term]; | 0.0 | 13.7 | #DIV/0! | 1.63E-03 |
| P01026 | SPMYSIITPNVLR | 0,C+12[AnyN-term]; | 0.0 | 7.2 | #DIV/0! | 1.66E-03 |
| P08932 | EGAQEMDCNDETVFQAVDTALK | 8,IBTP[C]; | 0.0 | 8.5 | #DIV/0! | 1.67E-03 |
| P17475 | EDAQETDTSQQDQSPTYR | 1,Cation_Fe[III][E]; | 0.0 | 21.5 | #DIV/0! | 1.68E-03 |
| P02770 | INKECCHGDLLECADDRAELAK | 13,CarbamidomethylDTT[C]; | 0.0 | 10.0 | #DIV/0! | 1.68E-03 |
| P02770 | HPYFYAPELLYYAEK | 3,Nitro[Y]; | 0.0 | 11.3 | #DIV/0! | 1.71E-03 |
| P02770 | AMCTSFQENPTSFLGHYLHEVAR | 3,Dipyrrolylmethanemethyl[C]; | 0.0 | 44.3 | #DIV/0! | 1.73E-03 |
| P02770 | RHPYFYAPELLYYAEK | 13,Dehydrated[Y]; | 0.0 | 6.2 | #DIV/0! | 1.74E-03 |
| P12346 | EGVCPEGSIDSAPVK | 4,Cys->Arg[C]; | 0.0 | 23.5 | #DIV/0! | 1.77E-03 |
| P02770 | FTFHSDICTLPDKEK | 8,Carbamidomethyl[C]; | 0.0 | 50.5 | #DIV/0! | 1.77E-03 |
| P02770 | KAETFTFHSDICTLPDKEK | 12,IBTP[C]; | 0.0 | 45.3 | #DIV/0! | 1.80E-03 |
| P09006 | RPSMIDELYLPK | 3,Pent(1)HexNAc(1)[S]; | 0.0 | 6.8 | #DIV/0! | 1.84E-03 |
| P02770 | LRDNYGELADCCAK | 14,Xlink_BS2G[96][K]; | 0.0 | 14.0 | #DIV/0! | 1.86E-03 |
| P02770 | PTSFLGHYLHEVAR | 0,C+12[AnyN-term]; | 0.0 | 9.3 | #DIV/0! | 1.89E-03 |
| P02770 | FPNAEFAEITK | 0,Formyl[AnyN-term]; | 0.0 | 13.5 | #DIV/0! | 1.89E-03 |
| P02770 | DNPNLPPFQRPEAEAMCTSFQENPTSFLGHYLHEVAR | 17,Cys->Arg[C]; | 0.0 | 2.8 | #DIV/0! | 1.94E-03 |
| P02770 | CCAEGDPPACYGTVLAEFQPLVEEPK | 2,4-ONE[C]; | 0.0 | 2.8 | #DIV/0! | 1.94E-03 |
| Q03626 | HAEEHHTAYAVYSLSK | 11,Val->Xle[V]; | 0.0 | 2.8 | #DIV/0! | 1.94E-03 |
| P09006 | AVLDVAETGTEAAAATGVK | 8,Dehydrated[T]; | 0.0 | 10.0 | #DIV/0! | 1.98E-03 |
| P02770 | CFATEGPNLVAR | 0,Methyl_2H(2)[AnyN-term]; | 0.0 | 42.3 | #DIV/0! | 2.00E-03 |
| P02770 | LRDNYGELADCCAK | 12,maleimide[C]; | 0.0 | 8.7 | #DIV/0! | 2.03E-03 |
| P04276 | VCQELSTLGKDDFR | 2,Carbamidomethyl[C]; | 0.0 | 54.7 | #DIV/0! | 2.07E-03 |
| P02770 | TSFLGHYLHEVAR | 0,Ethylphosphate[AnyN-term]; | 0.0 | 8.2 | #DIV/0! | 2.10E-03 |
| P02770 | TCVADENAENCDKSIHTLFGDK | 2,Carbamidomethyl[C];10,Deamidated[N];11,Carbamidomethyl[C]; | 0.0 | 25.2 | #DIV/0! | 2.15E-03 |
| P20759 | NTPPTMDTDGSYFLYSK | 0,Methyl[AnyN-term]; | 0.0 | 14.0 | #DIV/0! | 2.18E-03 |
| P20760 | NTPPTMDTDGSYFLYSK | 0,Methyl[AnyN-term]; | 0.0 | 14.0 | #DIV/0! | 2.18E-03 |
| P81827 | CESLDSTGLCR | 1,Carbamidomethyl[C];10,Carbamidomethyl[C]; | 0.0 | 468.3 | #DIV/0! | 2.23E-03 |
| P12346 | VCPEGSIDSAPVK | 0,Unknown_302[AnyN-term]; | 0.0 | 9.2 | #DIV/0! | 2.27E-03 |
| P22057 | GHDTVQPNFQQDK | 0,GIST-Quat[AnyN-term]; | 0.0 | 10.5 | #DIV/0! | 2.30E-03 |
| P14841 | EADASEEGVQRALDFAVSEYNK | 5,Thiophospho[S]; | 0.0 | 4.3 | #DIV/0! | 2.32E-03 |
| P01026 | LVAYYTLIGANGQR | 11,Deamidated[N]; | 0.0 | 25.3 | #DIV/0! | 2.36E-03 |
| P02770 | MGDFAQFVDK | 0,C+12[AnyN-term]; | 0.0 | 13.0 | #DIV/0! | 2.38E-03 |
| P02770 | CFATEGPNLVAR | 2,Fluoro[F]; | 0.0 | 13.2 | #DIV/0! | 2.41E-03 |
| Q68FP1 | AQHVQVEEGSEPDGFWEALGGK | 13,Asp->Cys[D]; | 0.0 | 10.0 | #DIV/0! | 2.53E-03 |
| P02770 | HPYFYAPELLYYAEK | 0,Diethylphosphate[AnyN-term]; | 0.0 | 21.0 | #DIV/0! | 2.59E-03 |
| P02770 | NCFATEGPNLVAR | 0,Piperidine[AnyN-term]; | 0.0 | 5.5 | #DIV/0! | 2.62E-03 |
| P02761 | MQHIDVLENSLGFK | 0,Unknown_248[AnyN-term]; | 0.0 | 15.8 | #DIV/0! | 2.77E-03 |
| P02761 | GNLDVAKLNGDWFSIVVASNKR | 9,Deamidated[N]; | 0.0 | 44.2 | #DIV/0! | 2.79E-03 |
| P02770 | CCAEGDPPACYGTVLAEFQPLVEEPK | 10,4-ONE[C]; | 0.0 | 34.0 | #DIV/0! | 2.83E-03 |
| P02770 | RHPDYSVSLLLR | 12,Arg->Tyr[R]; | 0.0 | 16.2 | #DIV/0! | 2.89E-03 |
| P02770 | LPCVEDYLSAILNR | 3,GG[C](Dicarbamidomethyl[C]); | 0.0 | 9.8 | #DIV/0! | 2.90E-03 |
| P14046 | PGPCGSEVATVPETGRK | 4,EHD-diphenylpentanone[C]; | 0.0 | 15.0 | #DIV/0! | 2.90E-03 |
| P02770 | SFQENPTSFLGHYLHEVAR | 0,Phenylisocyanate_2H(5)[AnyN-term]; | 0.0 | 2.2 | #DIV/0! | 2.95E-03 |
| P02770 | GLVLIAFSQYLQK | 0,ISD_z+2_ion[AnyN-term]; | 0.0 | 10.3 | #DIV/0! | 3.03E-03 |
| P02770 | ADKDNCFATEGPNLVAR | 6,Dimethylphosphothione[C]; | 0.0 | 26.8 | #DIV/0! | 3.05E-03 |
| P02770 | RHPDYSVSLLLR | 0,ISD_z+2_ion[AnyN-term]; | 0.0 | 36.8 | #DIV/0! | 3.07E-03 |
| P98158 | CAADGSPPLLLLPENVR | 1,Carbamidomethyl[C]; | 0.0 | 12.5 | #DIV/0! | 3.14E-03 |
| P02770 | YMCENQATISSK | 2,Oxidation[M]; | 0.0 | 15.3 | #DIV/0! | 3.14E-03 |
| P02770 | FQENPTSFLGHYLHEVAR | 0,Methyl[AnyN-term]; | 0.0 | 8.8 | #DIV/0! | 3.15E-03 |
| P02770 | TVMGDFAQFVDK | 2,Val->Thr[V]; | 0.0 | 22.5 | #DIV/0! | 3.33E-03 |
| P12346 | KGTDFQLNQLQGK | 14,Cation_Na[AnyC-term]; | 0.0 | 5.5 | #DIV/0! | 3.45E-03 |
| P12346 | HTTIFEVLPQK | 3,Dehydrated[T]; | 0.0 | 15.2 | #DIV/0! | 3.51E-03 |
| P02770 | HPYFYAPELLYYAEK | 0,Iodoacetanilide_13C(6)[AnyN-term]; | 0.0 | 16.2 | #DIV/0! | 3.65E-03 |
| P04276 | DYEKDKVCQELSTLGKDDFR | 2,Isopropylphospho[Y]; | 0.0 | 8.3 | #DIV/0! | 3.90E-03 |
| P06866 | GAVSPVGVQPILNK | 3,Val->Asn[V]; | 0.0 | 6.5 | #DIV/0! | 3.90E-03 |
| P02770 | PPFQRPEAEAMCTSFQENPTSFLGHYLHEVAR | 12,NEMsulfur[C]; | 0.0 | 27.2 | #DIV/0! | 3.90E-03 |
| P02761 | FAKLCEAHGITR | 5,GG[C](Dicarbamidomethyl[C]); | 0.0 | 7.7 | #DIV/0! | 3.96E-03 |
| P20759 | MNGQPQENYK | 1,Oxidation[M];2,Deamidated[N]; | 0.0 | 15.3 | #DIV/0! | 4.02E-03 |
| P20760 | MNGQPQENYK | 1,Oxidation[M];2,Deamidated[N]; | 0.0 | 15.3 | #DIV/0! | 4.02E-03 |
| P02770 | NLPPFQRPEAEAMCTSFQENPTSFLGHYLHEVAR | 14,MDCC[C]; | 0.0 | 0.8 | #DIV/0! | 4.10E-03 |
| P02770 | NCFATEGPNLVAR | 2,Cys->SecNEM_2H(5)[C]; | 0.0 | 0.8 | #DIV/0! | 4.10E-03 |
| P02770 | QTALAELVK | 0,Unknown_302[AnyN-term]; | 0.0 | 2.5 | #DIV/0! | 4.10E-03 |
| P12346 | GTTYEEYLGAEYLQAVGNIR | 4,Phosphoadenosine[Y]; | 0.0 | 10.0 | #DIV/0! | 4.10E-03 |
| P12346 | VAQEHFGKGK | 0,Carbamyl[AnyN-term]; | 0.0 | 1.7 | #DIV/0! | 4.10E-03 |
| Q03626 | ALGCLEASWETIEQGR | 4,Carbamidomethyl[C]; | 0.0 | 5.8 | #DIV/0! | 4.10E-03 |
| Q6IE52 | ALGCLEASWETIEQGR | 4,Carbamidomethyl[C]; | 0.0 | 5.8 | #DIV/0! | 4.10E-03 |
| P01048 | TELTADCETK | 7,Carbamidomethyl[C]; | 0.0 | 26.2 | #DIV/0! | 4.12E-03 |
| P08932 | TELTADCETK | 7,Carbamidomethyl[C]; | 0.0 | 26.2 | #DIV/0! | 4.12E-03 |
| P02770 | PTSFLGHYLHEVAR | 0,CLIP_TRAQ_2[AnyN-term]; | 0.0 | 22.3 | #DIV/0! | 4.15E-03 |
| P02770 | SFQENPTSFLGHYLHEVAR | 0,Biotin[AnyN-term]; | 0.0 | 10.7 | #DIV/0! | 4.16E-03 |
| P17475 | EDAQETDTSQQDQSPTYR | 7,Cation_Ca[II][D]; | 0.0 | 7.8 | #DIV/0! | 4.25E-03 |
| Q63041 | PLSLCALTAVDQSVLLLKPEAK | 5,Delta_Hg(1)[C]; | 0.0 | 5.7 | #DIV/0! | 4.29E-03 |
| P05545 | YQAEAFVADFK | 2,Gln->His[Q]; | 0.0 | 5.2 | #DIV/0! | 4.33E-03 |
| P02770 | LRDNYGELADCCAK | 11,Cys->Dha[C]; | 0.0 | 7.3 | #DIV/0! | 4.49E-03 |
| P02770 | FQENPTSFLGHYLHEVAR | 5,Pro->Cys[P]; | 0.0 | 38.0 | #DIV/0! | 4.50E-03 |
| P02770 | KQTALAELVK | 0,Carbamyl[AnyN-term]; | 0.0 | 9.7 | #DIV/0! | 4.52E-03 |
| P20760 | TCVVVDISQNDPEVR | 2,Hydroxycinnamyl[C]; | 0.0 | 11.2 | #DIV/0! | 4.66E-03 |
| P02770 | HPYFYAPELLYYAEK | 8,Cation_Ca[II][E]; | 0.0 | 3.2 | #DIV/0! | 4.71E-03 |
| P27590 | TACAHWSDHCCLWSTEIQVK | 3,Carbamidomethyl[C];10,Carbamidomethyl[C];11,Carbamidomethyl[C]; | 0.0 | 24.8 | #DIV/0! | 5.09E-03 |
| P02780 | IRGTINSTVTLHDYMK | 0,Unknown_302[AnyN-term]; | 0.0 | 8.2 | #DIV/0! | 5.12E-03 |
| Q9JHB9 | IRGTINSTVTLHDYMK | 0,Unknown_302[AnyN-term]; | 0.0 | 8.2 | #DIV/0! | 5.12E-03 |
| P12346 | TCHLAQAPNHVVVSR | 0,Propyl[AnyN-term]; | 0.0 | 6.7 | #DIV/0! | 5.15E-03 |
| P02770 | EAEAMCTSFQENPTSFLGHYLHEVAR | 6,NEIAA_2H(5)[C]; | 0.0 | 10.0 | #DIV/0! | 5.19E-03 |
| P02770 | YMCENQATISSK | 3,Cys->Arg[C]; | 0.0 | 70.2 | #DIV/0! | 5.21E-03 |
| P02770 | TSFLGHYLHEVAR | 0,mTRAQ_13C(6)15N(2)[AnyN-term]; | 0.0 | 2.8 | #DIV/0! | 5.27E-03 |
| P07632 | ISLSGEHSIIGR | 0,Phenylisocyanate_2H(5)[AnyN-term]; | 0.0 | 9.0 | #DIV/0! | 5.34E-03 |
| P20059 | GPDSVFLIK | 0,Methyl[AnyN-term]; | 0.0 | 13.5 | #DIV/0! | 5.34E-03 |
| P20059 | FNPVTGEVPPR | 0,Methyl_2H(2)[AnyN-term];2,Deamidated[N]; | 0.0 | 22.2 | #DIV/0! | 5.36E-03 |
| P01048 | CQALDMMISRPPGFSPFR | 2,Gln->Tyr[Q]; | 0.0 | 7.5 | #DIV/0! | 5.50E-03 |
| P08932 | CQALDMMISRPPGFSPFR | 2,Gln->Tyr[Q]; | 0.0 | 7.5 | #DIV/0! | 5.50E-03 |
| P02770 | RHPYFYAPELLYYAEK | 6,Dehydrated[Y]; | 0.0 | 12.7 | #DIV/0! | 5.52E-03 |
| P02761 | CRELYLVAYK | 0,Pyro-carbamidomethyl[AnyN-termC]; | 0.0 | 22.2 | #DIV/0! | 5.57E-03 |
| P02770 | HPDYSVSLLLR | 0,Hex[AnyN-term]; | 0.0 | 16.2 | #DIV/0! | 5.59E-03 |
| P04276 | DYEKDKVCQELSTLGKDDFR | 0,CAF[AnyN-term]; | 0.0 | 11.0 | #DIV/0! | 5.73E-03 |
| P83121 | CISFDSTGFCYVGR | 1,Carbamidomethyl[C];10,Carbamidomethyl[C]; | 0.0 | 44.8 | #DIV/0! | 5.74E-03 |
| P15083 | WSNDGCHILPSHDEGAR | 6,Carbamidomethyl[C]; | 0.0 | 9.2 | #DIV/0! | 5.81E-03 |
| P24090 | QQAEHAVEGDCDFHILK | 11,Ub-Br2[C]; | 0.0 | 20.7 | #DIV/0! | 5.88E-03 |
| P12346 | CHLAQAPNHVVVSR | 0,Succinyl_2H(4)[AnyN-term]; | 0.0 | 3.5 | #DIV/0! | 5.93E-03 |
| P02770 | KTVMGDFAQFVDK | 4,EHD-diphenylpentanone[M]; | 0.0 | 13.2 | #DIV/0! | 5.99E-03 |
| Q62930 | VMLTTTFLDDVK | 0,Carbamyl[AnyN-term]; | 0.0 | 5.5 | #DIV/0! | 6.02E-03 |
| Q64230 | QQCLQFFYK | 3,Carbamidomethyl[C]; | 0.0 | 14.7 | #DIV/0! | 6.02E-03 |
| P02761 | LNGDWFSIVVASNKR | 5,Delta_H(-4)O(3)[W]; | 0.0 | 22.2 | #DIV/0! | 6.06E-03 |
| P02770 | SFLGHYLHEVAR | 1,Didehydro[S]; | 0.0 | 11.7 | #DIV/0! | 6.12E-03 |
| P02761 | LCEAHGITR | 2,Trioxidation[C]; | 0.0 | 28.0 | #DIV/0! | 6.16E-03 |
| P02770 | CTSFQENPTSFLGHYLHEVAR | 1,Carbamidomethyl[C]; | 0.0 | 18.8 | #DIV/0! | 6.23E-03 |
| P20059 | GAQATWAELSWPHEK | 0,ICPL_2H(4)[AnyN-term]; | 0.0 | 25.5 | #DIV/0! | 6.26E-03 |
| P02770 | SQRFPNAEFAEITK | 5,Pro->Lys[P]; | 0.0 | 6.7 | #DIV/0! | 6.30E-03 |
| P12346 | LLEACTFHKS | 5,Carbamidomethyl[C]; | 0.0 | 29.3 | #DIV/0! | 6.36E-03 |
| P02770 | PTSFLGHYLHEVAR | 0,iTRAQ4plex114[AnyN-term]; | 0.0 | 13.8 | #DIV/0! | 6.37E-03 |
| P02761 | NGDWFSIVVASNKR | 0,C+12[AnyN-term]; | 0.0 | 16.2 | #DIV/0! | 6.52E-03 |
| P15083 | QSSVSCDQSSQIVSMTLNPVKK | 6,Acetyl[C]; | 0.0 | 4.0 | #DIV/0! | 6.57E-03 |
| P02770 | PEAQRLPCVEDYLSAILNR | 8,iodoTMT[C]; | 0.0 | 5.5 | #DIV/0! | 6.60E-03 |
| P17475 | NNYHSEAFSVNFADSEEAKK | 14,Cation_Na[D]; | 0.0 | 30.7 | #DIV/0! | 6.61E-03 |
| P02770 | AADKDNCFATEGPNLVAR | 5,Asp->Cys[D]; | 0.0 | 7.3 | #DIV/0! | 6.68E-03 |
| P02770 | KQTALAELVK | 0,Hex[AnyN-term]; | 0.0 | 16.5 | #DIV/0! | 6.74E-03 |
| P01048 | PMDSSDLKPVLKHAVEHFNNNTK | 22,FMN[T]; | 0.0 | 2.5 | #DIV/0! | 6.75E-03 |
| P08932 | PMDSSDLKPVLKHAVEHFNNNTK | 22,FMN[T]; | 0.0 | 2.5 | #DIV/0! | 6.75E-03 |
| P02770 | CTSFQENPTSFLGHYLHEVAR | 7,Asn->His[N]; | 0.0 | 9.5 | #DIV/0! | 6.78E-03 |
| P14046 | PRKDPPPKDPVIETIR | 0,glycidamide[AnyN-term]; | 0.0 | 15.0 | #DIV/0! | 6.83E-03 |
| Q03626 | PRKDPPPKDPVIETIR | 0,glycidamide[AnyN-term]; | 0.0 | 15.0 | #DIV/0! | 6.83E-03 |
| P02770 | CFATEGPNLVAR | 1,NEM_2H(5)+H2O[C]; | 0.0 | 13.5 | #DIV/0! | 6.87E-03 |
| P02770 | ADDRAELAKYMCENQATISSK | 11,Oxidation[M]; | 0.0 | 3.8 | #DIV/0! | 7.06E-03 |
| P02770 | CTSFQENPTSFLGHYLHEVAR | 1,Propionamide_2H(3)[C]; | 0.0 | 1.5 | #DIV/0! | 7.08E-03 |
| P02770 | LVQEVTDFAK | 0,Carbamyl[AnyN-term]; | 0.0 | 1.5 | #DIV/0! | 7.08E-03 |
| P02770 | YMCENQATISSK | 0,ISD_z+2_ion[AnyN-term]; | 0.0 | 1.5 | #DIV/0! | 7.08E-03 |
| P14740 | CLSCDLNPER | 1,Carbamidomethyl[C];4,Carbamidomethyl[C]; | 0.0 | 7.5 | #DIV/0! | 7.08E-03 |
| P02761 | LNGDWFSIVVASNK | 4,Cation_Na[D]; | 0.0 | 30.3 | #DIV/0! | 7.08E-03 |
| P17475 | NNYHSEAFSVNFADSEEAKK | 6,Cation_Na[E]; | 0.0 | 12.5 | #DIV/0! | 7.10E-03 |
| P14046 | HTSSWLVTPK | 2,Dehydrated[T]; | 0.0 | 4.3 | #DIV/0! | 7.14E-03 |
| Q03626 | HTSSWLVTPK | 2,Dehydrated[T]; | 0.0 | 4.3 | #DIV/0! | 7.14E-03 |
| Q6IE52 | HTSSWLVTPK | 2,Dehydrated[T]; | 0.0 | 4.3 | #DIV/0! | 7.14E-03 |
| P17475 | ISSNLADFAFSLYR | 4,Asn->Met[N]; | 0.0 | 7.0 | #DIV/0! | 7.42E-03 |
| P12346 | VTEFATCHLAQAPNHVVVSR | 8,Phosphoguanosine[H]; | 0.0 | 10.8 | #DIV/0! | 7.42E-03 |
| Q01177 | WEYCNLKR | 4,Carbamidomethyl[C]; | 0.0 | 10.8 | #DIV/0! | 7.42E-03 |
| P02761 | LNGDWFSIVVASNKR | 2,Deamidated_18O(1)[N](Delta_H(1)N(-1)18O(1)[N]); | 0.0 | 24.8 | #DIV/0! | 7.44E-03 |
| P02770 | DKDNCFATEGPNLVAR | 5,NEM_2H(5)[C]; | 0.0 | 12.8 | #DIV/0! | 7.46E-03 |
| P14046 | VTASPQSLCGLR | 9,Cys->Dha[C]; | 0.0 | 4.0 | #DIV/0! | 7.74E-03 |
| Q03626 | VTASPQSLCGLR | 9,Cys->Dha[C]; | 0.0 | 4.0 | #DIV/0! | 7.74E-03 |
| Q6IE52 | VTASPQSLCGLR | 9,Cys->Dha[C]; | 0.0 | 4.0 | #DIV/0! | 7.74E-03 |
| P20760 | EEMTQSQVSITCMVK | 12,Carbamidomethyl[C]; | 0.0 | 21.2 | #DIV/0! | 7.80E-03 |
| P20059 | NPLPAAHETVAK | 0,Carbamyl[AnyN-term]; | 0.0 | 13.3 | #DIV/0! | 7.87E-03 |
| P02770 | PEAEAMCTSFQENPTSFLGHYLHEVAR | 7,Cys->Asn[C]; | 0.0 | 6.5 | #DIV/0! | 7.93E-03 |
| P02770 | DNIPADLPSIAADFVEDK | 0,Ethylphosphate[AnyN-term]; | 0.0 | 7.7 | #DIV/0! | 8.04E-03 |
| P27590 | ERDWMSIVTPAR | 0,Amidine[AnyN-term]; | 0.0 | 5.7 | #DIV/0! | 8.09E-03 |
| P12788 | NKDTLDNDIMLIK | 4,Thr->Ala[T]; | 0.0 | 5.7 | #DIV/0! | 8.09E-03 |
| P27590 | DWMSIVTPAR | 1,Cation_Na[D]; | 0.0 | 19.0 | #DIV/0! | 8.10E-03 |
| P02770 | PNAEFAEITK | 0,iTRAQ4plex114[AnyN-term]; | 0.0 | 30.3 | #DIV/0! | 8.50E-03 |
| P17475 | HSEAFSVNFADSEEAKK | 0,iTRAQ4plex114[AnyN-term]; | 0.0 | 17.0 | #DIV/0! | 8.58E-03 |
| P20059 | NPLPAAHETVAK | 1,Oxidation[N]; | 0.0 | 2.3 | #DIV/0! | 8.63E-03 |
| P02650 | SKMEEQTQQIR | 3,Oxidation[M]; | 0.0 | 2.3 | #DIV/0! | 8.63E-03 |
| P24090 | VLHAQCHSTPDSAEDVRK | 6,Hydroxycinnamyl[C]; | 0.0 | 2.3 | #DIV/0! | 8.63E-03 |
| P02770 | CYGTVLAEFQPLVEEPK | 1,Cys->Met[C]; | 0.0 | 10.3 | #DIV/0! | 8.65E-03 |
| P20759 | NTPPTMDTDGSYFLYSK | 1,Oxidation[N]; | 0.0 | 25.5 | #DIV/0! | 8.73E-03 |
| P20760 | NTPPTMDTDGSYFLYSK | 1,Oxidation[N]; | 0.0 | 25.5 | #DIV/0! | 8.73E-03 |
| P02770 | TCVADENAENCDKSIHTLFGDK | 2,Carbamidomethyl[C];11,Carbamidomethyl[C]; | 0.0 | 130.7 | #DIV/0! | 8.75E-03 |
| P12346 | DVAFVKHTTIFEVLPQK | 6,Lys->Thr[K]; | 0.0 | 5.8 | #DIV/0! | 8.78E-03 |
| Q9QX79 | SIQHLPEQEEPEDSKGK | 0,Acetyl_2H(3)[AnyN-term]; | 0.0 | 5.8 | #DIV/0! | 8.78E-03 |
| P02770 | AELAKYMCENQATISSK | 8,cGMP+RMP-loss[C]; | 0.0 | 12.3 | #DIV/0! | 8.80E-03 |
| P02770 | FATEGPNLVAR | 0,Pyridylacetyl[AnyN-term](Phenylisocyanate[AnyN-term]); | 0.0 | 11.8 | #DIV/0! | 8.83E-03 |
| P02770 | QEPERNECFLQHKDDNPNLPPFQRPEAEAMCTSFQENPTSFLGHYLHEVAR | 27,Ala->Trp[A]; | 0.0 | 10.5 | #DIV/0! | 8.86E-03 |
| P02761 | LCEAHGITR | 2,GG[C](Dicarbamidomethyl[C]); | 0.0 | 3.3 | #DIV/0! | 8.89E-03 |
| P13635 | EYTDDSFTNRK | 0,ISD_z+2_ion[AnyN-term]; | 0.0 | 6.7 | #DIV/0! | 8.89E-03 |
| P12346 | AIRNQREGVCPEGSIDSAPVK | 10,IGBP_13C(2)[C]; | 0.0 | 9.8 | #DIV/0! | 8.95E-03 |
| P20767 | GAPISQGVDTANPTK | 5,AEBS[S]; | 0.0 | 4.8 | #DIV/0! | 8.97E-03 |
| P20059 | SGAQATWAELSWPHEK | 1,Didehydro[S]; | 0.0 | 5.2 | #DIV/0! | 9.04E-03 |
| P27590 | TDYGAGYSCDSDMHGWYR | 3,DNCB_hapten[Y]; | 0.0 | 19.7 | #DIV/0! | 9.06E-03 |
| P02770 | HPYFYAPELLYYAEK | 0,Carbamyl[AnyN-term]; | 0.0 | 6.2 | #DIV/0! | 9.07E-03 |
| P02770 | INKECCHGDLLECADDRAELAK | 5,CarbamidomethylDTT[C]; | 0.0 | 25.8 | #DIV/0! | 9.16E-03 |
| P12346 | CLRDGGGDVAFVK | 1,Pyridylethyl[C]; | 0.0 | 3.0 | #DIV/0! | 9.28E-03 |
| P02770 | AADKDNCFATEGPNLVAR | 7,Dehydro[C]; | 0.0 | 13.2 | #DIV/0! | 9.32E-03 |
| O70417 | AYLISNTPVDGGFNYIQTR | 10,Cation_Na[D]; | 0.0 | 9.3 | #DIV/0! | 9.34E-03 |
| P02770 | AFKAWAVAR | 0,Formyl[AnyN-term]; | 0.0 | 6.5 | #DIV/0! | 9.38E-03 |
| P05545 | AVLDVDETGTEGAAATAVTAALK | 7,Cation_Na[E]; | 0.0 | 10.3 | #DIV/0! | 9.43E-03 |
| P02770 | SQCLAEIEHDNIPADLPSIAADFVEDK | 3,GG[C](Dicarbamidomethyl[C]);11,Deamidated[N]; | 0.0 | 7.7 | #DIV/0! | 9.43E-03 |
| P17475 | VEAVPMSLPPQVK | 0,Formyl[AnyN-term]; | 0.0 | 13.2 | #DIV/0! | 9.44E-03 |
| P02770 | CKAADKDNCFATEGPNLVAR | 9,Cys->Met[C]; | 0.0 | 4.7 | #DIV/0! | 9.58E-03 |
| P17475 | SVNFADSEEAKK | 1,Glucuronyl[S]; | 0.0 | 7.8 | #DIV/0! | 9.58E-03 |
| P02770 | RHPYFYAPELLYYAEK | 15,Cation_Ca[II][E]; | 0.0 | 12.0 | #DIV/0! | 9.87E-03 |
| P02761 | RGNLDVAKLNGDWFSIVVASNKR | 10,Asn->His[N]; | 0.0 | 4.8 | #DIV/0! | 9.87E-03 |
| P02770 | VFLGTFLYEYSR | 0,ICPL_2H(4)[AnyN-term]; | 0.0 | 9.5 | #DIV/0! | 9.88E-03 |
| P02770 | HIKLVQEVTDFAK | 0,3sulfo[AnyN-term]; | 0.0 | 20.5 | #DIV/0! | 9.90E-03 |
| P12346 | CLRDGGGDVAFVK | 4,Cation_Ag[D]; | 0.0 | 15.7 | #DIV/0! | 9.93E-03 |
| P02770 | RPCFSALTVDETYVPK | 3,CarbamidomethylDTT[C]; | 0.0 | 15.5 | #DIV/0! | 1.02E-02 |
| P02770 | FKAETFTFHSDICTLPDKEK | 13,phenylsulfonylethyl[C]; | 0.0 | 6.2 | #DIV/0! | 1.02E-02 |
| P02770 | EAHKSEIAHR | 0,3sulfo[AnyN-term]; | 0.0 | 4.0 | #DIV/0! | 1.03E-02 |
| P02770 | FQENPTSFLGHYLHEVAR | 2,Xlink_DFDNB[Q]; | 0.0 | 2.7 | #DIV/0! | 1.03E-02 |
| P12346 | VSTVLTAQK | 6,Dehydrated[T]; | 0.0 | 2.7 | #DIV/0! | 1.03E-02 |
| P20059 | PDSVFLIKEDK | 0,Amidine[AnyN-term]; | 0.0 | 1.3 | #DIV/0! | 1.03E-02 |
| P27590 | ISLSKCQLQSLGFMK | 6,NEMsulfur[C]; | 0.0 | 8.5 | #DIV/0! | 1.03E-02 |
| P05545 | AVLDVDETGTEGAAATAVTAALK | 6,Cation_Na[D]; | 0.0 | 7.0 | #DIV/0! | 1.04E-02 |
| P17475 | DADFHVDK | 1,Dehydrated[D]; | 0.0 | 8.7 | #DIV/0! | 1.04E-02 |
| Q9QX79 | SAPSVLEAATESLAK | 0,Succinyl[AnyN-term]; | 0.0 | 2.2 | #DIV/0! | 1.04E-02 |
| P01048 | NCNDETVFQAVDTALKK | 4,Asp->Pro[D]; | 0.0 | 3.7 | #DIV/0! | 1.04E-02 |
| P02770 | YNEVLTQCCTESDKAACLTPK | 2,Deamidated[N];8,GG[C](Dicarbamidomethyl[C]);17,Carbamidomethyl[C]; | 0.0 | 16.8 | #DIV/0! | 1.05E-02 |
| P02770 | RPCFSALTVDETYVPK | 3,Cys->Arg[C]; | 0.0 | 30.7 | #DIV/0! | 1.07E-02 |
| P02770 | HPYFYAPELLYYAEK | 13,Ala->Asp[A]; | 0.0 | 1.7 | #DIV/0! | 1.08E-02 |
| P05544 | DLEERTSMVLVNYLLFK | 0,GIST-Quat[AnyN-term]; | 0.0 | 1.7 | #DIV/0! | 1.08E-02 |
| P05545 | IFSQQADLSR | 0,ICPL_2H(4)[AnyN-term]; | 0.0 | 1.7 | #DIV/0! | 1.08E-02 |
| P09006 | LYQAEAFTADFQQSR | 4,Ala->Asp[A]; | 0.0 | 9.2 | #DIV/0! | 1.10E-02 |
| P17475 | VFNNDADLSGITEDAPLK | 0,Dimethyl_2H(4)[AnyN-term]; | 0.0 | 22.5 | #DIV/0! | 1.10E-02 |
| P02761 | TKDLSSDIK | 0,Carbamyl[AnyN-term]; | 0.0 | 16.0 | #DIV/0! | 1.11E-02 |
| P02770 | FATEGPNLVAR | 0,SPITC[AnyN-term]; | 0.0 | 11.2 | #DIV/0! | 1.11E-02 |
| P02770 | TSFLGHYLHEVAR | 1,Palmitoyl[T]; | 0.0 | 18.5 | #DIV/0! | 1.13E-02 |
| P01835 | RDGVLDSVTDQDSK | 0,Iodoacetanilide[AnyN-term]; | 0.0 | 10.2 | #DIV/0! | 1.15E-02 |
| P01836 | RDGVLDSVTDQDSK | 0,Iodoacetanilide[AnyN-term]; | 0.0 | 10.2 | #DIV/0! | 1.15E-02 |
| P02770 | LRDNYGELADCCAK | 12,Hydroxycinnamyl[C]; | 0.0 | 4.3 | #DIV/0! | 1.16E-02 |
| P02770 | HPYFYAPELLYYAEK | 0,Methylthio[AnyN-term]; | 0.0 | 4.7 | #DIV/0! | 1.16E-02 |
| P20059 | ECQSEGVLFFQGNRK | 2,Diethylphosphate[C]; | 0.0 | 8.2 | #DIV/0! | 1.16E-02 |
| Q9QX79 | TAPTSSPSITAPR | 0,NIC[AnyN-term](ICPL[AnyN-term]); | 0.0 | 7.7 | #DIV/0! | 1.17E-02 |
| P08426 | IRHPSYNANTFDNDIMLIK | 0,Acetyl_2H(3)[AnyN-term]; | 0.0 | 2.0 | #DIV/0! | 1.17E-02 |
| P12346 | NGDGKEDLIWEILK | 10,Delta_H(-4)O(3)[W]; | 0.0 | 2.0 | #DIV/0! | 1.17E-02 |
| P14046 | VYHKEESSCIHSSCTAER | 9,Cys->Dha[C]; | 0.0 | 1.0 | #DIV/0! | 1.17E-02 |
| P17475 | FLEEVKNNYHSEAFSVNFADSEEAKK | 7,Asn->His[N]; | 0.0 | 1.0 | #DIV/0! | 1.17E-02 |
| Q03626 | VYHKEESSCIHSSCTAER | 9,Cys->Dha[C]; | 0.0 | 1.0 | #DIV/0! | 1.17E-02 |
| P05545 | INDYVSNQTQGKIAELFSELDER | 0,ISD_z+2_ion[AnyN-term]; | 0.0 | 13.8 | #DIV/0! | 1.17E-02 |
| P01048 | YNAELESGNQFVLYR | 2,dHex[N]; | 0.0 | 8.3 | #DIV/0! | 1.18E-02 |
| Q68FP1 | TPSAAYLWVGTGASDAEK | 1,Thr->Pro[T]; | 0.0 | 8.0 | #DIV/0! | 1.21E-02 |
| P02761 | EKIEENGSMR | 1,Cation_Ca[II][E]; | 0.0 | 1.8 | #DIV/0! | 1.21E-02 |
| P21704 | ILSRYDIAVVQEVR | 0,CAF[AnyN-term]; | 0.0 | 1.8 | #DIV/0! | 1.21E-02 |
| Q03626 | AEEHHTAYAVYSLSK | 0,AccQTag[AnyN-term]; | 0.0 | 1.8 | #DIV/0! | 1.21E-02 |
| Q6IRK9 | LQQDGLENVHLEQVR | 0,Propyl[AnyN-term]; | 0.0 | 3.7 | #DIV/0! | 1.21E-02 |
| P02770 | DVFLGTFLYEYSR | 12,Formyl[S](Ser->Asp[S]); | 0.0 | 11.5 | #DIV/0! | 1.22E-02 |
| P02761 | VFMQHIDVLENSLGFK | 6,Xle->Trp[I]; | 0.0 | 34.8 | #DIV/0! | 1.22E-02 |
| P02761 | FMQHIDVLENSLGFK | 0,Carbamyl[AnyN-term]; | 0.0 | 8.7 | #DIV/0! | 1.25E-02 |
| P02770 | QLKTVMGDFAQFVDK | 0,Delta_H(2)C(2)[AnyN-term]; | 0.0 | 8.5 | #DIV/0! | 1.25E-02 |
| P02770 | FAQFVDKCCKAADKDNCFATEGPNLVAR | 0,AccQTag[AnyN-term]; | 0.0 | 6.2 | #DIV/0! | 1.26E-02 |
| P02761 | CEAHGITR | 1,Glutathione[C]; | 0.0 | 3.2 | #DIV/0! | 1.26E-02 |
| P02770 | LECADDRAELAK | 3,Carbamidomethyl[C]; | 0.0 | 6.3 | #DIV/0! | 1.26E-02 |
| P14740 | YYQLGCR | 6,Carbamidomethyl[C]; | 0.0 | 3.2 | #DIV/0! | 1.26E-02 |
| P02761 | LNGDWFSIVVASNK | 4,Cation_Fe[III][D]; | 0.0 | 5.0 | #DIV/0! | 1.27E-02 |
| P12346 | TTIFEVLPQK | 1,Decanoyl[T]; | 0.0 | 25.5 | #DIV/0! | 1.27E-02 |
| P02761 | KLCEAHGITR | 0,Methyl_2H(2)[AnyN-term]; | 0.0 | 1.2 | #DIV/0! | 1.27E-02 |
| P02770 | TSFLGHYLHEVAR | 1,monomethylphosphothione[T]; | 0.0 | 3.5 | #DIV/0! | 1.27E-02 |
| P02770 | YMCENQATISSK | 0,Methyl_2H(3)13C(1)[AnyN-term]; | 0.0 | 1.2 | #DIV/0! | 1.27E-02 |
| P12346 | KPVDQYEDCYLAR | 9,Propionamide[C]; | 0.0 | 1.2 | #DIV/0! | 1.27E-02 |
| P20761 | PIQHQDWMSGKEFK | 14,Xlink_DTSSP[88][K]; | 0.0 | 1.2 | #DIV/0! | 1.27E-02 |
| P20059 | THPMHSRCNADPGLSALLSDHR | 8,Acetyl[C]; | 0.0 | 15.7 | #DIV/0! | 1.28E-02 |
| P27274 | QVYQQCWR | 6,Carbamidomethyl[C]; | 0.0 | 10.3 | #DIV/0! | 1.28E-02 |
| P02761 | VFMQHIDVLENSLGFK | 0,C+12[AnyN-term]; | 0.0 | 2.8 | #DIV/0! | 1.29E-02 |
| Q6IG01 | FLEQQNQVLQTK | 2,Xle->Cys[L]; | 0.0 | 2.8 | #DIV/0! | 1.29E-02 |
| Q6IG03 | FLEQQNQVLQTK | 2,Xle->Cys[L]; | 0.0 | 2.8 | #DIV/0! | 1.29E-02 |
| Q6IMF3 | FLEQQNQVLQTK | 2,Xle->Cys[L]; | 0.0 | 2.8 | #DIV/0! | 1.29E-02 |
| P02770 | LRDNYGELADCCAK | 12,NO_SMX_SMCT[C]; | 0.0 | 9.5 | #DIV/0! | 1.29E-02 |
| P02770 | PEAEAMCTSFQENPTSFLGHYLHEVAR | 8,Thr->Pro[T]; | 0.0 | 3.3 | #DIV/0! | 1.29E-02 |
| P02770 | TSFQENPTSFLGHYLHEVAR | 5,Glu->Thr[E]; | 0.0 | 3.3 | #DIV/0! | 1.29E-02 |
| P81827 | PDEICAWVVVTTR | 5,Malonyl[C]; | 0.0 | 17.3 | #DIV/0! | 1.31E-02 |
| P81828 | PDEICAWVVVTTR | 5,Malonyl[C]; | 0.0 | 17.3 | #DIV/0! | 1.31E-02 |
| P83121 | PDEICAWVVVTTR | 5,Malonyl[C]; | 0.0 | 17.3 | #DIV/0! | 1.31E-02 |
| P12346 | QAPNHVVVSR | 0,Cytopiloyne+water[AnyN-term]; | 0.0 | 7.0 | #DIV/0! | 1.32E-02 |
| P00787 | ILRGENHCGIESEIVAGIPR | 8,Carbamidomethyl[C]; | 0.0 | 12.3 | #DIV/0! | 1.32E-02 |
| P02770 | SFLGHYLHEVAR | 0,ICPL_2H(4)[AnyN-term]; | 0.0 | 10.3 | #DIV/0! | 1.35E-02 |
| P02770 | ENPTSFLGHYLHEVAR | 2,Asn->His[N]; | 0.0 | 2.5 | #DIV/0! | 1.36E-02 |
| P17475 | TDTSQQDQSPTYRK | 0,Methyl_2H(2)[AnyN-term]; | 0.0 | 2.5 | #DIV/0! | 1.36E-02 |
| P02770 | FQENPTSFLGHYLHEVAR | 0,Diisopropylphosphate[AnyN-term]; | 0.0 | 6.0 | #DIV/0! | 1.37E-02 |
| P02770 | HPYFYAPELLYYAEKYNEVLTQCCTESDK | 17,Deamidated[N];23,Carbamidomethyl[C];24,Carbamidomethyl[C]; | 0.0 | 9.2 | #DIV/0! | 1.38E-02 |
| P02770 | SFLGHYLHEVAR | 0,Biotin_Thermo-21328[AnyN-term]; | 0.0 | 3.7 | #DIV/0! | 1.39E-02 |
| P02770 | TVLAEFQPLVEEPK | 1,Methylphosphonate[T]; | 0.0 | 3.7 | #DIV/0! | 1.39E-02 |
| P02770 | APQVSTPTLVEAAR | 1,Ala->Asp[A]; | 0.0 | 14.2 | #DIV/0! | 1.42E-02 |
| P02770 | QRLPCVEDYLSAILNR | 0,Unknown_302[AnyN-term]; | 0.0 | 7.3 | #DIV/0! | 1.43E-02 |
| P06866 | LQTEGDGIYTLNSEK | 4,Cation_Fe[III][E]; | 0.0 | 7.3 | #DIV/0! | 1.43E-02 |
| P02761 | LNGDWFSIVVASNKR | 1,Oxidation[L]; | 0.0 | 13.3 | #DIV/0! | 1.45E-02 |
| P12346 | CSTSRLLEACTFHK | 0,Methyl_2H(2)[AnyN-term];10,Carbamidomethyl[C]; | 0.0 | 6.7 | #DIV/0! | 1.45E-02 |
| P02770 | CFATEGPNLVAR | 0,Methyl_2H(3)13C(1)[AnyN-term]; | 0.0 | 5.0 | #DIV/0! | 1.46E-02 |
| P02770 | HPDYSVSLLLR | 0,mTRAQ[AnyN-term]; | 0.0 | 146.2 | #DIV/0! | 1.46E-02 |
| P07632 | PVQGVIHFEQK | 0,SPITC_13C(6)[AnyN-term]; | 0.0 | 7.3 | #DIV/0! | 1.48E-02 |
| P20760 | CKVNSGAFPAPIEK | 2,Xlink_DST[114][K]; | 0.0 | 7.5 | #DIV/0! | 1.49E-02 |
| P12346 | CLRDGGGDVAFVK | 1,Carbamidomethyl[C]; | 0.0 | 43.3 | #DIV/0! | 1.51E-02 |
| Q9QX79 | LPQKNTAPTSSPSITAPR | 4,Xlink_DTSSP[174][K]; | 0.0 | 22.2 | #DIV/0! | 1.52E-02 |
| P02770 | CTSFQENPTSFLGHYLHEVAR | 1,Carbamidomethyl[C];7,Deamidated[N]; | 0.0 | 7.8 | #DIV/0! | 1.53E-02 |
| P05545 | ALYQAEAFVADFK | 3,Difuran[Y]; | 0.0 | 8.7 | #DIV/0! | 1.55E-02 |
| P12346 | PVTEFATCHLAQAPNHVVVSR | 0,Unknown_302[AnyN-term]; | 0.0 | 2.2 | #DIV/0! | 1.55E-02 |
| P12346 | GTTYEEYLGAEYLQAVGNIR | 7,Phosphoadenosine[Y]; | 0.0 | 5.0 | #DIV/0! | 1.56E-02 |
| P02770 | PYFYAPELLYYAEK | 0,Phenylisocyanate_2H(5)[AnyN-term]; | 0.0 | 3.7 | #DIV/0! | 1.57E-02 |
| P02770 | TALAELVKHKPK | 0,Unknown_302[AnyN-term]; | 0.0 | 2.8 | #DIV/0! | 1.59E-02 |
| P02770 | SFLGHYLHEVAR | 1,NeuAc[S]; | 0.0 | 10.8 | #DIV/0! | 1.61E-02 |
| P83121 | ALQCFRCISFDSTGFCYVGR | 0,Phenylisocyanate_2H(5)[AnyN-term];16,Carbamidomethyl[C]; | 0.0 | 9.0 | #DIV/0! | 1.63E-02 |
| P02770 | QLKTVMGDFAQFVDK | 1,Ethyl+Deamidated[Q]; | 0.0 | 5.8 | #DIV/0! | 1.63E-02 |
| P02770 | LRDNYGELADCCAK | 11,NO_SMX_SMCT[C]; | 0.0 | 6.2 | #DIV/0! | 1.65E-02 |
| P04276 | FPSSTFEQVSQLVK | 2,Pro->Cys[P]; | 0.0 | 11.3 | #DIV/0! | 1.65E-02 |
| P02770 | CAEGDPPACYGTVLAEFQPLVEEPK | 0,Phosphogluconoylation[AnyN-term]; | 0.0 | 6.5 | #DIV/0! | 1.67E-02 |
| P20761 | IEKNYKNTEPVMDSDGSFFMYSK | 4,Hep[N]; | 0.0 | 3.5 | #DIV/0! | 1.67E-02 |
| P02761 | LNGDWFSIVVASNKR | 1,Xle->Glu[L]; | 0.0 | 8.2 | #DIV/0! | 1.68E-02 |
| P02770 | LTVDETYVPK | 0,Tween20[AnyN-term]; | 0.0 | 11.2 | #DIV/0! | 1.68E-02 |
| P17475 | LSQAVHKAVLTLDER | 0,Carbamyl[AnyN-term]; | 0.0 | 6.3 | #DIV/0! | 1.70E-02 |
| Q64240 | AFAELWAFDAAQGK | 0,Methyl[AnyN-term]; | 0.0 | 7.2 | #DIV/0! | 1.70E-02 |
| P02770 | YMCENQATISSK | 0,Methyl_2H(2)[AnyN-term];2,Oxidation[M];3,Carbamidomethyl[C]; | 0.0 | 6.0 | #DIV/0! | 1.72E-02 |
| P02770 | TSFLGHYLHEVAR | 0,Succinyl_2H(4)[AnyN-term]; | 0.0 | 1.5 | #DIV/0! | 1.72E-02 |
| P05544 | IAELFSDLEER | 3,Glu->His[E]; | 0.0 | 1.5 | #DIV/0! | 1.72E-02 |
| P09006 | AVLDVAETGTEAAAATGVK | 10,Dehydrated[T]; | 0.0 | 6.0 | #DIV/0! | 1.72E-02 |
| P20760 | FSWFIDDVEVHTAQTHAPEK | 12,Dehydrated[T]; | 0.0 | 3.0 | #DIV/0! | 1.72E-02 |
| Q01177 | TAVTAAGTPCQEWAAQEPHSHR | 10,Acetyl[C]; | 0.0 | 4.5 | #DIV/0! | 1.72E-02 |
| P09006 | AVLDVAETGTEAAAATGVK | 4,Cation_Na[D]; | 0.0 | 2.3 | #DIV/0! | 1.73E-02 |
| P01048 | HLGQSLNCNANVYMRPWENK | 7,Hydroxymethyl[N]; | 0.0 | 2.3 | #DIV/0! | 1.73E-02 |
| P02770 | GDFAQFVDKCCKAADKDNCFATEGPNLVAR | 11,Cys->Thr[C]; | 0.0 | 2.3 | #DIV/0! | 1.73E-02 |
| P08932 | HLGQSLNCNANVYMRPWENK | 7,Hydroxymethyl[N]; | 0.0 | 2.3 | #DIV/0! | 1.73E-02 |
| P14046 | HTSSWLVTPK | 8,Dehydrated[T]; | 0.0 | 2.3 | #DIV/0! | 1.73E-02 |
| Q03626 | HTSSWLVTPK | 8,Dehydrated[T]; | 0.0 | 2.3 | #DIV/0! | 1.73E-02 |
| Q6IE52 | HTSSWLVTPK | 8,Dehydrated[T]; | 0.0 | 2.3 | #DIV/0! | 1.73E-02 |
| P02770 | LRDNYGELADCCAK | 12,DTT[C]; | 0.0 | 9.7 | #DIV/0! | 1.74E-02 |
| P12346 | NGDGKEDLIWEILK | 6,Cation_Ca[II][E]; | 0.0 | 3.3 | #DIV/0! | 1.74E-02 |
| P02770 | LRDNYGELADCCAKQEPER | 11,Ub-amide[C]; | 0.0 | 4.2 | #DIV/0! | 1.75E-02 |
| P02770 | PEAEAMCTSFQENPTSFLGHYLHEVAR | 6,Met->Aha[M]; | 0.0 | 5.7 | #DIV/0! | 1.76E-02 |
| P02770 | AMCTSFQENPTSFLGHYLHEVAR | 3,NEIAA_2H(5)[C]; | 0.0 | 17.7 | #DIV/0! | 1.78E-02 |
| P02770 | TSFQENPTSFLGHYLHEVAR | 0,CLIP_TRAQ_4[AnyN-term]; | 0.0 | 5.8 | #DIV/0! | 1.79E-02 |
| P02761 | NLDVAKLNGDWFSIVVASNKR | 0,ICPL_2H(4)[AnyN-term]; | 0.0 | 6.2 | #DIV/0! | 1.79E-02 |
| P02770 | SQCLAEIEHDNIPADLPSIAADFVEDK | 3,2-succinyl[C]; | 0.0 | 6.5 | #DIV/0! | 1.79E-02 |
| P05545 | ELDERTSMVLVNYLLFK | 0,GIST-Quat[AnyN-term]; | 0.0 | 2.0 | #DIV/0! | 1.80E-02 |
| P12346 | GTDFQLNQLQGK | 0,ICPL_13C(6)[AnyN-term]; | 0.0 | 2.0 | #DIV/0! | 1.80E-02 |
| P13635 | KLFPGQQYLYVLR | 1,ZQG[K]; | 0.0 | 2.0 | #DIV/0! | 1.80E-02 |
| P20759 | NTPPTMDTDGSYFLYSK | 0,Methyl_2H(2)[AnyN-term]; | 0.0 | 36.8 | #DIV/0! | 1.85E-02 |
| P20760 | NTPPTMDTDGSYFLYSK | 0,Methyl_2H(2)[AnyN-term]; | 0.0 | 36.8 | #DIV/0! | 1.85E-02 |
| P01048 | YNAELESGNQFVLYR | 0,mTRAQ_13C(6)15N(2)[AnyN-term]; | 0.0 | 4.7 | #DIV/0! | 1.85E-02 |
| P20759 | TGSEVSSVFIFPPK | 0,ICPL_2H(4)[AnyN-term]; | 0.0 | 20.8 | #DIV/0! | 1.86E-02 |
| P20760 | TGSEVSSVFIFPPK | 0,ICPL_2H(4)[AnyN-term]; | 0.0 | 20.8 | #DIV/0! | 1.86E-02 |
| P27590 | INFECSYPLDMK | 5,Carbamyl[C]; | 0.0 | 6.2 | #DIV/0! | 1.86E-02 |
| P02770 | LLECADDRAELAK | 3,Glu->Trp[E]; | 0.0 | 6.7 | #DIV/0! | 1.86E-02 |
| P02770 | VQEVTDFAK | 0,NIC[AnyN-term](ICPL[AnyN-term]); | 0.0 | 16.3 | #DIV/0! | 1.87E-02 |
| P02770 | ATEDQLKTVMGDFAQFVDK | 6,Xle->Gly[L]; | 0.0 | 6.0 | #DIV/0! | 1.87E-02 |
| P07522 | YVCNCVIGYIGER | 3,Carbamidomethyl[C];5,Carbamidomethyl[C]; | 0.0 | 27.2 | #DIV/0! | 1.88E-02 |
| P05545 | EAKKFINDYVSNQTQGKIAELFSELDER | 9,Trioxidation[Y]; | 0.0 | 17.5 | #DIV/0! | 1.91E-02 |
| P02761 | LNGDWFSIVVASNKR | 5,Nitro[W]; | 0.0 | 6.8 | #DIV/0! | 1.91E-02 |
| P02770 | FQENPTSFLGHYLHEVAR | 4,HexN[N]; | 0.0 | 12.8 | #DIV/0! | 1.92E-02 |
| P12346 | TCHLAQAPNHVVVSR | 2,NEIAA_2H(5)[C]; | 0.0 | 18.0 | #DIV/0! | 1.92E-02 |
| P02770 | EFKAETFTFHSDICTLPDKEK | 12,Cation_Ca[II][D]; | 0.0 | 2.8 | #DIV/0! | 1.93E-02 |
| P02770 | RMSQRFPNAEFAEITK | 6,Phe->Asn[F]; | 0.0 | 9.3 | #DIV/0! | 1.95E-02 |
| P12346 | LAQAPNHVVVSR | 0,ESP_2H(10)[AnyN-term]; | 0.0 | 6.3 | #DIV/0! | 1.97E-02 |
| P02764 | DESEIVFVDWTKDK | 3,Hex(1)Pent(2)Me(1)[S]; | 0.0 | 1.8 | #DIV/0! | 1.97E-02 |
| P12346 | ADRDQYELLCLDNTR | 10,Crotonaldehyde[C]; | 0.0 | 1.8 | #DIV/0! | 1.97E-02 |
| P26051 | FIEGHVVIPR | 0,Biotin[AnyN-term]; | 0.0 | 3.7 | #DIV/0! | 1.97E-02 |
| P07522 | VNLDPASVPPR | 1,Val->Thr[V]; | 0.0 | 10.7 | #DIV/0! | 1.97E-02 |
| P02770 | CVADENAENCDKSIHTLFGDK | 10,NO_SMX_SMCT[C]; | 0.0 | 9.5 | #DIV/0! | 1.98E-02 |
| P02770 | YNEVLTQCCTESDKAACLTPK | 2,Deamidated[N];9,GG[C](Dicarbamidomethyl[C]);17,Carbamidomethyl[C]; | 0.0 | 12.3 | #DIV/0! | 1.98E-02 |
| P01026 | GIPIVTSPYQIHFTK | 0,Piperidine[AnyN-term]; | 0.0 | 1.7 | #DIV/0! | 1.99E-02 |
| P07522 | VVHLHAQPGTENR | 12,Ammonia-loss[N]; | 0.0 | 1.7 | #DIV/0! | 1.99E-02 |
| P08426 | IRHPSYNANTFDNDIMLIK | 0,Carbamyl[AnyN-term]; | 0.0 | 1.7 | #DIV/0! | 1.99E-02 |
| P12346 | VSTVLTAQK | 2,Hex(1)HexNAc(1)Phos(1)[S]; | 0.0 | 12.2 | #DIV/0! | 1.99E-02 |
| P02770 | YMCENQATISSK | 2,Oxidation[M];3,GG[C](Dicarbamidomethyl[C]); | 0.0 | 4.5 | #DIV/0! | 1.99E-02 |
| P12346 | CLRDGGGDVAFVK | 1,Cys->CamSec[C]; | 0.0 | 3.8 | #DIV/0! | 1.99E-02 |
| P12346 | SKDFQLFGSPLGKDLLFK | 0,Carbamyl[AnyN-term]; | 0.0 | 3.2 | #DIV/0! | 2.04E-02 |
| P02770 | QCLAEIEHDNIPADLPSIAADFVEDK | 2,HexNAc[C]; | 0.0 | 4.8 | #DIV/0! | 2.06E-02 |
| P12346 | DLKQEDFQLLCPDGTKKPVTEFATCHLAQAPNHVVVSR | 25,DTT[C]; | 0.0 | 4.8 | #DIV/0! | 2.06E-02 |
| Q63041 | AEQGAYLGPLPYK | 0,ISD_z+2_ion[AnyN-term]; | 0.0 | 5.5 | #DIV/0! | 2.06E-02 |
| P12346 | GAFKCLRDGGGDVAFVK | 5,CarboxymethylDTT[C]; | 0.0 | 4.2 | #DIV/0! | 2.07E-02 |
| P02761 | RGNLDVAKLNGDWFSIVVASNKR | 5,Cation_Na[D]; | 0.0 | 6.2 | #DIV/0! | 2.08E-02 |
| P27590 | ACAHWSDHCCLWSTEIQVK | 6,Hex(1)Pent(1)[S]; | 0.0 | 2.7 | #DIV/0! | 2.10E-02 |
| P02770 | CYGTVLAEFQPLVEEPK | 1,GG[C](Dicarbamidomethyl[C]); | 0.0 | 7.8 | #DIV/0! | 2.10E-02 |
| P13635 | MYYSGVDLTK | 1,Met->Aha[M]; | 0.0 | 5.7 | #DIV/0! | 2.10E-02 |
| P02780 | SGSGCSILDEVIR | 5,Carbamidomethyl[C]; | 0.0 | 134.0 | #DIV/0! | 2.11E-02 |
| Q9JHB9 | SGSGCSILDEVIR | 5,Carbamidomethyl[C]; | 0.0 | 134.0 | #DIV/0! | 2.11E-02 |
| P02770 | TVLAEFQPLVEEPK | 1,FMN[T]; | 0.0 | 41.5 | #DIV/0! | 2.12E-02 |
| P02761 | LNGDWFSIVVASNKR | 0,Methyl_2H(2)[AnyN-term];2,Deamidated[N]; | 0.0 | 3.5 | #DIV/0! | 2.12E-02 |
| P02770 | LPCVEDYLSAILNR | 2,Pro->Tyr[P]; | 0.0 | 3.5 | #DIV/0! | 2.12E-02 |
| P07522 | DQILAGDNADLSKEVASLDNSPK | 12,Hex(2)[S]; | 0.0 | 8.2 | #DIV/0! | 2.13E-02 |
| P02770 | ENPTSFLGHYLHEVAR | 4,PET[T]; | 0.0 | 6.2 | #DIV/0! | 2.15E-02 |
| P02770 | PNAEFAEITK | 0,DiLeu4plex115[AnyN-term]; | 0.0 | 5.5 | #DIV/0! | 2.16E-02 |
| P02770 | RFPNAEFAEITK | 0,CLIP_TRAQ_4[AnyN-term]; | 0.0 | 27.3 | #DIV/0! | 2.16E-02 |
| P02770 | ADKDNCFATEGPNLVAR | 6,DHP[C]; | 0.0 | 27.3 | #DIV/0! | 2.18E-02 |
| P27274 | CCQADLCNK | 1,Carbamidomethyl[C];2,Carbamidomethyl[C];7,Carbamidomethyl[C]; | 0.0 | 3.8 | #DIV/0! | 2.18E-02 |
| P02770 | DDNPNLPPFQRPEAEAMCTSFQENPTSFLGHYLHEVAR | 5,Deamidated[N];18,Carbamidomethyl[C]; | 0.0 | 5.0 | #DIV/0! | 2.21E-02 |
| P02770 | GDFAQFVDK | 0,CLIP_TRAQ_2[AnyN-term]; | 0.0 | 12.7 | #DIV/0! | 2.21E-02 |
| P02770 | MSQRFPNAEFAEITK | 2,O-Isopropylmethylphosphonate[S]; | 0.0 | 5.2 | #DIV/0! | 2.22E-02 |
| P12346 | KPVTEFATCHLAQAPNHVVVSR | 1,Xlink_DTSSP[174][K]; | 0.0 | 4.7 | #DIV/0! | 2.24E-02 |
| P02770 | MSQRFPNAEFAEITK | 0,CLIP_TRAQ_3[AnyN-term]; | 0.0 | 14.8 | #DIV/0! | 2.24E-02 |
| Q64240 | EYCGVPGDGYEELTR | 3,Cys->Trp[C]; | 0.0 | 4.5 | #DIV/0! | 2.27E-02 |
| P02770 | APQVSTPTLVEAARNLGRVGTKCCTLPEAQRLPCVEDYLSAILNR | 34,Iodoacetanilide[C]; | 0.0 | 7.8 | #DIV/0! | 2.28E-02 |
| P02770 | HPDYSVSLLLR | 0,Iodoacetanilide_13C(6)[AnyN-term]; | 0.0 | 27.7 | #DIV/0! | 2.28E-02 |
| P27274 | CYNCLDPVSSCK | 1,Carbamidomethyl[C];4,Carbamidomethyl[C];11,Carbamidomethyl[C]; | 0.0 | 14.7 | #DIV/0! | 2.29E-02 |
| P36953 | FQELGKDALQR | 0,Pyridylacetyl[AnyN-term](Phenylisocyanate[AnyN-term]); | 0.0 | 10.3 | #DIV/0! | 2.30E-02 |
| P02781 | EEFDAPPEAVEANLK | 0,NP40[AnyN-term]; | 0.0 | 11.7 | #DIV/0! | 2.30E-02 |
| P09006 | MQQVEASLQPETLRR | 1,Met->Asp[M]; | 0.0 | 8.2 | #DIV/0! | 2.30E-02 |
| Q01177 | LSRPATITDNVIPACLPSPNYVVADR | 9,Propargylamine[D]; | 0.0 | 7.5 | #DIV/0! | 2.31E-02 |
| P01048 | KYNAELESGNQFVLYR | 5,Cation_Na[E]; | 0.0 | 4.0 | #DIV/0! | 2.32E-02 |
| P05545 | NDYVSNQTQGKIAELFSELDER | 1,Tris[N]; | 0.0 | 4.0 | #DIV/0! | 2.32E-02 |
| P12346 | EYLGAEYLQAVGNIR | 1,Cation_Na[E]; | 0.0 | 4.0 | #DIV/0! | 2.32E-02 |
| P02770 | RHPYFYAPELLYYAEKYNEV | 17,Fluorescein-tyramine[Y]; | 0.0 | 5.2 | #DIV/0! | 2.32E-02 |
| P22282 | RTPGETMYYISLPGSVR | 7,Oxidation[M]; | 0.0 | 7.3 | #DIV/0! | 2.34E-02 |
| P20761 | AQTQPREEQYNSTFRVVSALPIQHQDWMSGK | 5,Pro->Cys[P]; | 0.0 | 8.2 | #DIV/0! | 2.34E-02 |
| P48199 | SFSIFSYATK | 2,Phe->His[F]; | 0.0 | 3.5 | #DIV/0! | 2.35E-02 |
| P17475 | FLEEVKNNYHSEAFSVNFADSEEAKK | 4,Cation_Na[E]; | 0.0 | 20.5 | #DIV/0! | 2.36E-02 |
| P04276 | TKMPNASPEELADMVAK | 2,Lys->Asp[K]; | 0.0 | 5.8 | #DIV/0! | 2.36E-02 |
| P02770 | VMGDFAQFVDK | 0,Succinyl_2H(4)[AnyN-term]; | 0.0 | 29.7 | #DIV/0! | 2.38E-02 |
| P12346 | GYYAVAVVK | 10,Cation_Na[AnyC-term]; | 0.0 | 5.3 | #DIV/0! | 2.40E-02 |
| P02770 | RLPCVEDYLSAILNR | 0,Biotin_Thermo-21328[AnyN-term]; | 0.0 | 7.2 | #DIV/0! | 2.41E-02 |
| P02770 | PTSFLGHYLHEVAR | 0,mTRAQ[AnyN-term]; | 0.0 | 10.8 | #DIV/0! | 2.41E-02 |
| P06866 | QWVNPAAGDKLPK | 1,Gln->Pro[Q]; | 0.0 | 7.8 | #DIV/0! | 2.42E-02 |
| P02770 | RPCFSALTVDETYVPKEFK | 3,Phospho[C]; | 0.0 | 5.5 | #DIV/0! | 2.44E-02 |
| P01026 | VRVELLHNPAFCSMATAK | 12,Carbamidomethyl[C]; | 0.0 | 9.2 | #DIV/0! | 2.45E-02 |
| Q63041 | DLNDNDAYSVFQSIGLK | 0,Iodoacetanilide_13C(6)[AnyN-term]; | 0.0 | 2.7 | #DIV/0! | 2.50E-02 |
| P02761 | GNLDVAKLNGDWFSIVVASNKR | 20,Deamidated[N]; | 0.0 | 1.3 | #DIV/0! | 2.50E-02 |
| P02761 | KFAKLCEAHGITR | 0,SPITC_13C(6)[AnyN-term]; | 0.0 | 1.3 | #DIV/0! | 2.50E-02 |
| P02761 | LCEAHGITR | 0,Acetyl_2H(3)[AnyN-term]; | 0.0 | 0.7 | #DIV/0! | 2.50E-02 |
| P02770 | NCFATEGPNLVAR | 2,DiDehydro[C]; | 0.0 | 2.0 | #DIV/0! | 2.50E-02 |
| P02770 | PCVEDYLSAILNR | 0,Biotin[AnyN-term]; | 0.0 | 1.3 | #DIV/0! | 2.50E-02 |
| P02770 | TCVADENAENCDKSIHTLFGDK | 2,DNCB_hapten[C]; | 0.0 | 0.7 | #DIV/0! | 2.50E-02 |
| P05544 | VFSQQADLSR | 1,Val->Asn[V]; | 0.0 | 2.7 | #DIV/0! | 2.50E-02 |
| P05544 | LSLGAKDSTMEEILEGLK | 8,N-dimethylphosphate[S]; | 0.0 | 0.7 | #DIV/0! | 2.50E-02 |
| P09005 | VFSQQADLSR | 1,Val->Asn[V]; | 0.0 | 2.7 | #DIV/0! | 2.50E-02 |
| P14841 | SQTNLTNCPFHDQPHLMR | 4,Deamidated[N];8,Carbamidomethyl[C]; | 0.0 | 0.7 | #DIV/0! | 2.50E-02 |
| P15684 | KVVATTQMQAADAR | 8,Oxidation[M]; | 0.0 | 1.3 | #DIV/0! | 2.50E-02 |
| P17475 | ISSNLADFAFSLYR | 4,Deamidated[N]; | 0.0 | 0.7 | #DIV/0! | 2.50E-02 |
| P22282 | LDNCPFEEQTEQLKR | 4,Methylthio[C]; | 0.0 | 1.3 | #DIV/0! | 2.50E-02 |
| P22283 | LVNCPFEEQTEQLKR | 3,Deamidated[N];4,Carbamidomethyl[C]; | 0.0 | 2.0 | #DIV/0! | 2.50E-02 |
| Q63556 | VFSQQADLSR | 1,Val->Asn[V]; | 0.0 | 2.7 | #DIV/0! | 2.50E-02 |
| P02770 | PEAEAMCTSFQENPTSFLGHYLHEVAR | 4,Glu->His[E]; | 0.0 | 9.8 | #DIV/0! | 2.51E-02 |
| P02770 | QKAPQVSTPTLVEAAR | 2,Phosphoguanosine[K]; | 0.0 | 23.0 | #DIV/0! | 2.52E-02 |
| P02761 | NGDWFSIVVASNKR | 0,Phenylisocyanate_2H(5)[AnyN-term]; | 0.0 | 5.5 | #DIV/0! | 2.54E-02 |
| P02770 | RHPYFYAPELLYYAEK | 4,Dehydrated[Y]; | 0.0 | 13.8 | #DIV/0! | 2.54E-02 |
| P02770 | NECFLQHKDDNPNLPPFQRPEAEAMCTSFQENPTSFLGHYLHEVAR | 3,maleimide[C]; | 0.0 | 5.0 | #DIV/0! | 2.56E-02 |
| Q9QX79 | SYFVEYLIKESPCTQSQDSCSLQASDSEPVGLCQGSLIK | 33,Carbamidomethyl[C]; | 0.0 | 5.0 | #DIV/0! | 2.56E-02 |
| P02770 | NECFLQHKDDNPNLPPFQRPEAEAMCTSFQENPTSFLGHYLHEVAR | 26,maleimide[C]; | 0.0 | 11.0 | #DIV/0! | 2.56E-02 |
| P02770 | CCAEGDPPACYGTVLAEFQPLVEEPK | 10,NEMsulfur[C]; | 0.0 | 10.2 | #DIV/0! | 2.57E-02 |
| P17475 | FNNDADLSGITEDAPLK | 8,Pentose[S]; | 0.0 | 3.8 | #DIV/0! | 2.58E-02 |
| P20759 | WQQGNTFTCSVLHEGLHNHHTEK | 5,Asn->His[N]; | 0.0 | 5.3 | #DIV/0! | 2.61E-02 |
| P20760 | WQQGNTFTCSVLHEGLHNHHTEK | 5,Asn->His[N]; | 0.0 | 5.3 | #DIV/0! | 2.61E-02 |
| P02770 | CTSFQENPTSFLGHYLHEVAR | 4,Fluoro[F]; | 0.0 | 3.7 | #DIV/0! | 2.61E-02 |
| P02770 | LDAVKEKALVAAVR | 5,Malonyl[K]; | 0.0 | 11.3 | #DIV/0! | 2.62E-02 |
| P02782 | EELKKELEMYNAPPAAVEAK | 11,Hep[N]; | 0.0 | 11.3 | #DIV/0! | 2.62E-02 |
| P02761 | LCEAHGITR | 2,2-dimethylsuccinyl[C]; | 0.0 | 9.8 | #DIV/0! | 2.64E-02 |
| Q9QX79 | NTAPTSSPSITAPR | 2,Hex(1)HexNAc(1)[T]; | 0.0 | 4.7 | #DIV/0! | 2.64E-02 |
| P02770 | LRDNYGELADCCAK | 12,Diethylphosphate[C]; | 0.0 | 2.8 | #DIV/0! | 2.64E-02 |
| P02770 | FQENPTSFLGHYLHEVAR | 0,Methyl_2H(2)[AnyN-term]; | 0.0 | 2.8 | #DIV/0! | 2.64E-02 |
| P04276 | DKVCQELSTLGKDDFR | 2,Xlink_BS2G[96][K]; | 0.0 | 2.8 | #DIV/0! | 2.64E-02 |
| P02770 | CENQATISSK | 1,Carbamidomethyl[C];3,Deamidated[N]; | 0.0 | 3.0 | #DIV/0! | 2.67E-02 |
| P12346 | CPEGSIDSAPVK | 0,NHS-LC-Biotin[AnyN-term]; | 0.0 | 3.0 | #DIV/0! | 2.67E-02 |
| P20759 | SEVSSVFIFPPK | 1,Pent(2)[S]; | 0.0 | 3.0 | #DIV/0! | 2.67E-02 |
| P20760 | SEVSSVFIFPPK | 1,Pent(2)[S]; | 0.0 | 3.0 | #DIV/0! | 2.67E-02 |
| Q68FP1 | AQHVQVEEGSEPDGFWEALGGK | 11,Glu->Asp[E]; | 0.0 | 2.5 | #DIV/0! | 2.68E-02 |
| P20059 | ECHRGECQSEGVLFFQGNR | 2,MDCC[C]; | 0.0 | 6.2 | #DIV/0! | 2.69E-02 |
| P02761 | TDYDRYVMFHLINFK | 8,Oxidation[M]; | 0.0 | 4.0 | #DIV/0! | 2.69E-02 |
| P02770 | AADKDNCFATEGPNLVAR | 7,Trioxidation[C]; | 0.0 | 4.0 | #DIV/0! | 2.69E-02 |
| P02770 | YTQKAPQVSTPTLVEAAR | 4,Dimethyl_2H(4)13C(2)[K]; | 0.0 | 20.3 | #DIV/0! | 2.74E-02 |
| P02770 | EAHKSEIAHR | 0,Unknown_250[AnyN-term]; | 0.0 | 2.2 | #DIV/0! | 2.74E-02 |
| P27590 | CQLQSLGFMK | 1,Propionamide_2H(3)[C]; | 0.0 | 2.2 | #DIV/0! | 2.74E-02 |
| P36373 | CSSNNYQVWLGRNNLLEDEPFAQHR | 1,Diethylphosphothione[C]; | 0.0 | 13.0 | #DIV/0! | 2.74E-02 |
| P02770 | MCENQATISSK | 1,AHA-SS_CAM[M]; | 0.0 | 46.0 | #DIV/0! | 2.76E-02 |
| Q6IE52 | FGNSKYMVLVPSQLYTETPEK | 6,Fluorescein-tyramine[Y]; | 0.0 | 3.3 | #DIV/0! | 2.77E-02 |
| P01835 | DSTYSMSSTLSLTK | 1,Asp->Pro[D]; | 0.0 | 2.3 | #DIV/0! | 2.78E-02 |
| P01836 | DSTYSMSSTLSLTK | 1,Asp->Pro[D]; | 0.0 | 2.3 | #DIV/0! | 2.78E-02 |
| Q03626 | HAEEHHTAYAVYSLSK | 16,Lys->CamCys[K]; | 0.0 | 2.3 | #DIV/0! | 2.78E-02 |
| P02770 | FQENPTSFLGHYLHEVAR | 2,Gln->Phe[Q]; | 0.0 | 4.8 | #DIV/0! | 2.81E-02 |
| P12346 | CPDGTKKPVTEFATCHLAQAPNHVVVSR | 1,Delta_Hg(1)[C]; | 0.0 | 4.8 | #DIV/0! | 2.81E-02 |
| P12346 | FKCLRDGGGDVAFVK | 0,NHS-LC-Biotin[AnyN-term]; | 0.0 | 4.8 | #DIV/0! | 2.81E-02 |
| Q01177 | EQQCVIMAENSK | 0,Acetyl_2H(3)[AnyN-term]; | 0.0 | 5.3 | #DIV/0! | 2.82E-02 |
| P04276 | MPNASPEELADMVAK | 3,Deamidated[N]; | 0.0 | 18.0 | #DIV/0! | 2.83E-02 |
| P02761 | VFMQHIDVLENSLGFK | 0,Methyl[AnyN-term]; | 0.0 | 14.5 | #DIV/0! | 2.84E-02 |
| P02770 | TSFLGHYLHEVAR | 2,NeuAc[S]; | 0.0 | 1.8 | #DIV/0! | 2.84E-02 |
| P12346 | AQAPNHVVVSR | 0,BITC[AnyN-term]; | 0.0 | 1.8 | #DIV/0! | 2.84E-02 |
| P02770 | PEAEAMCTSFQENPTSFLGHYLHEVAR | 2,Glu->His[E]; | 0.0 | 6.3 | #DIV/0! | 2.84E-02 |
| P12346 | GTTYEEYLGAEYLQAVGNIR | 3,Phosphoadenosine[T]; | 0.0 | 4.5 | #DIV/0! | 2.86E-02 |
| P01835 | DSTYSMSSTLSLTK | 6,Oxidation[M]; | 0.0 | 143.8 | #DIV/0! | 2.86E-02 |
| P01836 | DSTYSMSSTLSLTK | 6,Oxidation[M]; | 0.0 | 143.8 | #DIV/0! | 2.86E-02 |
| P02770 | ENPTSFLGHYLHEVAR | 6,Phe->Pro[F]; | 0.0 | 13.0 | #DIV/0! | 2.91E-02 |
| P02770 | QENPTSFLGHYLHEVAR | 2,Carboxy[E]; | 0.0 | 6.3 | #DIV/0! | 2.92E-02 |
| P02770 | KPKATEDQLKTVMGDFAQFVDK | 1,Xlink_BS2G[96][K]; | 0.0 | 3.2 | #DIV/0! | 2.92E-02 |
| P02770 | LPCVEDYLSAILNR | 3,Cys->Tyr[C]; | 0.0 | 14.2 | #DIV/0! | 2.93E-02 |
| P01048 | KTEEDLCVGCFQPIPMDSSDLKPVLK | 10,DNCB_hapten[C]; | 0.0 | 8.5 | #DIV/0! | 2.94E-02 |
| P08932 | KTEEDLCVGCFQPIPMDSSDLKPVLK | 10,DNCB_hapten[C]; | 0.0 | 8.5 | #DIV/0! | 2.94E-02 |
| P02761 | NIIDLTKTDR | 0,CAF[AnyN-term]; | 0.0 | 5.5 | #DIV/0! | 2.94E-02 |
| P02770 | QENPTSFLGHYLHEVAR | 5,HexNAc(1)NeuAc(1)[T]; | 0.0 | 7.8 | #DIV/0! | 2.97E-02 |
| P26644 | VCPFAGILENGVVR | 2,Carbamidomethyl[C]; | 0.0 | 22.3 | #DIV/0! | 2.99E-02 |
| P02770 | ENPTSFLGHYLHEVAR | 2,Hex(3)[N]; | 0.0 | 3.0 | #DIV/0! | 3.01E-02 |
| P02770 | KAADKDNCFATEGPNLVAR | 8,DHP[C]; | 0.0 | 3.0 | #DIV/0! | 3.01E-02 |
| P09006 | EKAKALYQAEAFTADFQQSR | 2,Xlink_DSS[155][K]; | 0.0 | 1.5 | #DIV/0! | 3.01E-02 |
| P14046 | LMAYAFALAGNQEK | 0,ICPL_13C(6)[AnyN-term]; | 0.0 | 1.5 | #DIV/0! | 3.01E-02 |
| P17475 | VFNNDADLSGITEDAPLK | 5,Cation_Na[D]; | 0.0 | 1.5 | #DIV/0! | 3.01E-02 |
| P20759 | TGSEVSSVFIFPPK | 0,iTRAQ4plex114[AnyN-term]; | 0.0 | 4.5 | #DIV/0! | 3.01E-02 |
| P20760 | TGSEVSSVFIFPPK | 0,iTRAQ4plex114[AnyN-term]; | 0.0 | 4.5 | #DIV/0! | 3.01E-02 |
| P21704 | SRYDIAVVQEVR | 3,Biotin-tyramide[Y]; | 0.0 | 3.0 | #DIV/0! | 3.01E-02 |
| Q03626 | LMAYAFALAGNQEK | 0,ICPL_13C(6)[AnyN-term]; | 0.0 | 1.5 | #DIV/0! | 3.01E-02 |
| P01048 | YNAELESGNQFVLYR | 0,Diethylphosphate[AnyN-term]; | 0.0 | 6.5 | #DIV/0! | 3.03E-02 |
| P06866 | KPPEIANGYVEHLVR | 7,Deamidated[N]; | 0.0 | 6.5 | #DIV/0! | 3.03E-02 |
| P12346 | TEFATCHLAQAPNHVVVSR | 0,Biotin_Thermo-21328[AnyN-term]; | 0.0 | 5.5 | #DIV/0! | 3.04E-02 |
| P02761 | KLCEAHGITR | 0,ISD_z+2_ion[AnyN-term]; | 0.0 | 1.7 | #DIV/0! | 3.05E-02 |
| P02770 | FSALTVDETYVPK | 0,Diethylphosphate[AnyN-term]; | 0.0 | 1.7 | #DIV/0! | 3.05E-02 |
| P04276 | YCSSQIDAEMRDILQS | 2,NEIAA_2H(5)[C]; | 0.0 | 1.7 | #DIV/0! | 3.05E-02 |
| P14046 | NLHPLNELFPLAYIEDPK | 0,Methyl_2H(2)[AnyN-term]; | 0.0 | 1.7 | #DIV/0! | 3.05E-02 |
| Q03626 | NLHPLNELFPLAYIEDPK | 0,Methyl_2H(2)[AnyN-term]; | 0.0 | 1.7 | #DIV/0! | 3.05E-02 |
| P01048 | CFQPIPMDSSDLKPVLK | 1,Dimethylphosphothione[C]; | 0.0 | 8.8 | #DIV/0! | 3.06E-02 |
| P08932 | CFQPIPMDSSDLKPVLK | 1,Dimethylphosphothione[C]; | 0.0 | 8.8 | #DIV/0! | 3.06E-02 |
| P02770 | SFLGHYLHEVAR | 1,monomethylphosphothione[S]; | 0.0 | 7.5 | #DIV/0! | 3.07E-02 |
| Q99PS8 | AENGDFASFR | 3,Deamidated[N]; | 0.0 | 13.3 | #DIV/0! | 3.07E-02 |
| Q03626 | GDPIPNEQVFIK | 3,Pro->Cys[P]; | 0.0 | 3.7 | #DIV/0! | 3.07E-02 |
| Q6IE52 | GDPIPNEQVFIK | 3,Pro->Cys[P]; | 0.0 | 3.7 | #DIV/0! | 3.07E-02 |
| P20759 | SSVFIFPPK | 1,Hex(1)HexA(1)[S]; | 0.0 | 8.7 | #DIV/0! | 3.07E-02 |
| P20760 | SSVFIFPPK | 1,Hex(1)HexA(1)[S]; | 0.0 | 8.7 | #DIV/0! | 3.07E-02 |
| P20059 | FNPVTGEVPPR | 2,Asn->Met[N]; | 0.0 | 35.7 | #DIV/0! | 3.09E-02 |
| P02770 | SFQENPTSFLGHYLHEVAR | 5,Asn->His[N]; | 0.0 | 3.5 | #DIV/0! | 3.09E-02 |
| P02770 | PTSFLGHYLHEVAR | 2,Thr->Asn[T]; | 0.0 | 3.5 | #DIV/0! | 3.09E-02 |
| P02770 | CCSGSLVER | 0,Methyl_2H(2)[AnyN-term];1,Carbamidomethyl[C];2,Carbamidomethyl[C]; | 0.0 | 3.5 | #DIV/0! | 3.09E-02 |
| P02770 | AHRFKDLGEQHFK | 3,2HPG[R]; | 0.0 | 6.8 | #DIV/0! | 3.12E-02 |
| P20762 | VVSTLHIQHQDWMSGK | 14,Methylamine[S]; | 0.0 | 5.2 | #DIV/0! | 3.12E-02 |
| P02770 | APQVSTPTLVEAARNLGRVGTKCCTLPEAQRLPCVEDYLSAILNR | 34,HCysteinyl[C]; | 0.0 | 6.3 | #DIV/0! | 3.15E-02 |
| P02770 | CTSFQENPTSFLGHYLHEVAR | 0,Methyl_2H(2)[AnyN-term];1,Carbamidomethyl[C]; | 0.0 | 8.7 | #DIV/0! | 3.16E-02 |
| P02767 | PFASGKTAESGELHGLTTDEK | 6,AEBS[K]; | 0.0 | 4.5 | #DIV/0! | 3.16E-02 |
| P07314 | IDQVVTAGLK | 0,Phenylisocyanate_2H(5)[AnyN-term]; | 0.0 | 4.5 | #DIV/0! | 3.16E-02 |
| P09006 | PDQGKMQQVEASLQPETLR | 0,Unknown_302[AnyN-term]; | 0.0 | 14.3 | #DIV/0! | 3.17E-02 |
| P02770 | QLKTVMGDFAQFVDK | 0,Formyl[AnyN-term]; | 0.0 | 23.2 | #DIV/0! | 3.17E-02 |
| P02770 | AKYMCENQATISSK | 5,DiDehydro[C]; | 0.0 | 8.0 | #DIV/0! | 3.17E-02 |
| P02770 | FQENPTSFLGHYLHEVAR | 3,UgiJoullieProGlyProGly[E]; | 0.0 | 2.5 | #DIV/0! | 3.18E-02 |
| P02770 | CCTLPEAQRLPCVEDYLSAILNR | 1,CarbamidomethylDTT[C]; | 0.0 | 2.5 | #DIV/0! | 3.18E-02 |
| P02761 | NIIDLTKTDR | 0,NIC[AnyN-term](ICPL[AnyN-term]); | 0.0 | 9.3 | #DIV/0! | 3.21E-02 |
| P12346 | GAFQCLVEKGDVAFVK | 5,Cys->Xle[C]; | 0.0 | 4.8 | #DIV/0! | 3.21E-02 |
| P12346 | HTTIFEVLPQKADRDQYELLCLDNTR | 17,sulfo+amino[Y]; | 0.0 | 9.7 | #DIV/0! | 3.21E-02 |
| P02770 | LTVDETYVPKEFK | 2,Thr->Ala[T]; | 0.0 | 12.5 | #DIV/0! | 3.22E-02 |
| P17475 | VFNNDADLSGITEDAPLK | 0,Formyl[AnyN-term]; | 0.0 | 3.2 | #DIV/0! | 3.23E-02 |
| P02770 | DNPNLPPFQRPEAEAMCTSFQENPTSFLGHYLHEVAR | 17,Carbamidomethyl[C]; | 0.0 | 5.5 | #DIV/0! | 3.25E-02 |
| P02770 | GDFAQFVDK | 0,DiLeu4plex115[AnyN-term]; | 0.0 | 10.5 | #DIV/0! | 3.27E-02 |
| P02761 | LCEAHGITR | 2,Nethylmaleimide+water[C]; | 0.0 | 2.0 | #DIV/0! | 3.27E-02 |
| P02770 | FQENPTSFLGHYLHEVAR | 7,HexNAc(1)dHex(1)[S]; | 0.0 | 2.0 | #DIV/0! | 3.27E-02 |
| P02770 | LRDNYGELADCCAK | 11,Diethylphosphothione[C]; | 0.0 | 2.0 | #DIV/0! | 3.27E-02 |
| P02770 | MCENQATISSK | 0,Unknown_248[AnyN-term]; | 0.0 | 4.0 | #DIV/0! | 3.27E-02 |
| P14046 | IKLPTYCNTDYDMVPLAVPAVALDSSTDR | 7,Carbamidomethyl[C]; | 0.0 | 2.0 | #DIV/0! | 3.27E-02 |
| P20059 | VTSVDAAFR | 0,Delta_H(2)C(2)[AnyN-term]; | 0.0 | 2.0 | #DIV/0! | 3.27E-02 |
| P27590 | ERDWMSIVTPAR | 0,Carbamyl[AnyN-term]; | 0.0 | 4.2 | #DIV/0! | 3.28E-02 |
| P12346 | DLKQEDFQLLCPDGTKKPVTEFATCHLAQAPNHVVVSR | 25,Crotonaldehyde[C]; | 0.0 | 5.0 | #DIV/0! | 3.30E-02 |
| P02770 | CTSFQENPTSFLGHYLHEVAR | 1,Dipyridyl[C]; | 0.0 | 4.5 | #DIV/0! | 3.32E-02 |
| P14046 | ALMAYAFALAGNQEK | 3,Met->His[M]; | 0.0 | 7.8 | #DIV/0! | 3.33E-02 |
| P20059 | THPMHSRCNADPGLSALLSDHR | 0,Amidine[AnyN-term]; | 0.0 | 3.3 | #DIV/0! | 3.34E-02 |
| P00758 | NDLMLLHLSQPADITDGVK | 0,Carboxymethyl[AnyN-term]; | 0.0 | 1.2 | #DIV/0! | 3.35E-02 |
| P02770 | CCAEGDPPACYGTVLAEFQPLVEEPK | 6,UgiJoullieProGly[D]; | 0.0 | 3.5 | #DIV/0! | 3.35E-02 |
| P02770 | CCAEGDPPACYGTVLAEFQPLVEEPK | 1,4-ONE[C]; | 0.0 | 1.2 | #DIV/0! | 3.35E-02 |
| P02770 | QLKTVMGDFAQFVDK | 4,Dehydrated[T]; | 0.0 | 3.5 | #DIV/0! | 3.35E-02 |
| P02770 | MCENQATISSK | 2,Cys->Arg[C]; | 0.0 | 2.3 | #DIV/0! | 3.35E-02 |
| P17475 | FNNDADLSGITEDAPLK | 0,GIST-Quat_2H(3)[AnyN-term]; | 0.0 | 3.5 | #DIV/0! | 3.35E-02 |
| P20759 | NTPPTMDTDGSYFLYSK | 2,Thr->Xle[T]; | 0.0 | 1.2 | #DIV/0! | 3.35E-02 |
| P20760 | NTPPTMDTDGSYFLYSK | 2,Thr->Xle[T]; | 0.0 | 1.2 | #DIV/0! | 3.35E-02 |
| P36373 | NDLMLLHLSQPADITDGVK | 0,Carboxymethyl[AnyN-term]; | 0.0 | 1.2 | #DIV/0! | 3.35E-02 |
| P83121 | HICQTYPDEICAWVVVTTR | 11,DTT[C]; | 0.0 | 1.2 | #DIV/0! | 3.35E-02 |
| Q01177 | KLYDYCNIPLCASLSSFECGKPQVEPK | 7,Deamidated[N];11,GG[C](Dicarbamidomethyl[C]);19,Carbamidomethyl[C]; | 0.0 | 1.2 | #DIV/0! | 3.35E-02 |
| P02770 | PACYGTVLAEFQPLVEEPK | 3,IBTP[C]; | 0.0 | 3.0 | #DIV/0! | 3.36E-02 |
| P13635 | TYEWTVPK | 1,Hex(1)HexNAc(1)[T]; | 0.0 | 6.0 | #DIV/0! | 3.36E-02 |
| P14046 | EDNSIHWERPQKPTK | 3,Asn->Lys[N]; | 0.0 | 2.8 | #DIV/0! | 3.42E-02 |
| Q03626 | EDNSIHWERPQKPTK | 3,Asn->Lys[N]; | 0.0 | 2.8 | #DIV/0! | 3.42E-02 |
| P02770 | GDFAQFVDKCCKAADKDNCFATEGPNLVAR | 10,Cys->Thr[C]; | 0.0 | 3.8 | #DIV/0! | 3.42E-02 |
| P20760 | FSWFIDDVEVHTAQTHAPEK | 4,Phe->Glu[F]; | 0.0 | 14.3 | #DIV/0! | 3.45E-02 |
| P12346 | LCPDGTKKPVTEFATCHLAQAPNHVVVSR | 16,GG[C](Dicarbamidomethyl[C]); | 0.0 | 4.0 | #DIV/0! | 3.47E-02 |
| P02770 | PTSFLGHYLHEVAR | 2,O-Methylphosphate[T]; | 0.0 | 7.3 | #DIV/0! | 3.48E-02 |
| P02770 | ENPTSFLGHYLHEVAR | 0,AccQTag[AnyN-term]; | 0.0 | 7.0 | #DIV/0! | 3.48E-02 |
| P02770 | YTQKAPQVSTPTLVEAAR | 0,Galactosyl[AnyN-term]; | 0.0 | 6.5 | #DIV/0! | 3.48E-02 |
| P02770 | CCAEGDPPACYGTVLAEFQPLVEEPK | 2,NEMsulfur[C]; | 0.0 | 33.8 | #DIV/0! | 3.48E-02 |
| P02770 | PQVSTPTLVEAAR | 1,Pro->Arg[P]; | 0.0 | 11.8 | #DIV/0! | 3.51E-02 |
| P02770 | LRDNYGELADCCAK | 5,Dehydrated[Y];11,Carbamidomethyl[C];12,Carbamidomethyl[C]; | 0.0 | 16.3 | #DIV/0! | 3.51E-02 |
| P08426 | IIRHPSYNANTFDNDIMLIK | 3,Arg->Ser[R]; | 0.0 | 15.2 | #DIV/0! | 3.52E-02 |
| Q01177 | NPDNDEQGPWCYTTDPDQR | 11,Carbamidomethyl[C]; | 0.0 | 14.8 | #DIV/0! | 3.52E-02 |
| P02770 | PEAEAMCTSFQENPTSFLGHYLHEVAR | 0,C+12[AnyN-term]; | 0.0 | 8.7 | #DIV/0! | 3.54E-02 |
| P04276 | KMPNASPEELADMVAK | 0,ICPL_2H(4)[AnyN-term]; | 0.0 | 3.2 | #DIV/0! | 3.55E-02 |
| P13635 | FTKENEGTYYGPDGR | 0,ISD_z+2_ion[AnyN-term]; | 0.0 | 3.2 | #DIV/0! | 3.55E-02 |
| P02770 | DDRAELAKYMCENQATISSK | 8,Xlink_DTSSP[88][K]; | 0.0 | 3.5 | #DIV/0! | 3.62E-02 |
| Q01177 | WSDTSPHVPK | 0,ISD_z+2_ion[AnyN-term]; | 0.0 | 3.5 | #DIV/0! | 3.62E-02 |
| P02770 | PEAEAMCTSFQENPTSFLGHYLHEVAR | 12,Glu->His[E]; | 0.0 | 4.8 | #DIV/0! | 3.62E-02 |
| P20759 | SNSMVTLGCLVK | 2,Deamidated[N];4,Oxidation[M];9,Carbamidomethyl[C]; | 0.0 | 6.0 | #DIV/0! | 3.63E-02 |
| P20760 | SNSMVTLGCLVK | 2,Deamidated[N];4,Oxidation[M];9,Carbamidomethyl[C]; | 0.0 | 6.0 | #DIV/0! | 3.63E-02 |
| P02770 | NPTSFLGHYLHEVAR | 1,Hydroxymethyl[N]; | 0.0 | 6.5 | #DIV/0! | 3.64E-02 |
| P01026 | VELKPGDNLNVNFHLR | 0,Methyl[AnyN-term]; | 0.0 | 4.5 | #DIV/0! | 3.64E-02 |
| P48199 | FVLSPEQINAVYVGRVFSPNVLNWR | 12,DNCB_hapten[Y]; | 0.0 | 4.2 | #DIV/0! | 3.65E-02 |
| P02770 | SFLGHYLHEVAR | 1,Ethanedithiol[S]; | 0.0 | 10.2 | #DIV/0! | 3.66E-02 |
| P20059 | CNADPGLSALLSDHR | 1,Cys->His[C]; | 0.0 | 8.5 | #DIV/0! | 3.69E-02 |
| P02770 | QENPTSFLGHYLHEVAR | 5,BEMAD_ST_2H(6)[T]; | 0.0 | 2.5 | #DIV/0! | 3.69E-02 |
| P02770 | GDPPACYGTVLAEFQPLVEEPK | 6,Methylthio[C]; | 0.0 | 2.5 | #DIV/0! | 3.69E-02 |
| P26644 | CPFPSRPDNGFVNYPAKPVLSYK | 1,Carbamidomethyl[C];9,Deamidated[N]; | 0.0 | 19.8 | #DIV/0! | 3.70E-02 |
| P02770 | DLPSIAADFVEDK | 1,SulfanilicAcid[D]; | 0.0 | 8.5 | #DIV/0! | 3.73E-02 |
| P05544 | VVHKAVLDVDETGTEATAATGVATVIR | 14,PhosphoHexNAc[T]; | 0.0 | 1.8 | #DIV/0! | 3.79E-02 |
| P09006 | QQVEASLQPETLR | 0,CAF[AnyN-term]; | 0.0 | 5.5 | #DIV/0! | 3.79E-02 |
| P12346 | HLAQAPNHVVVSR | 0,TNBS[AnyN-term]; | 0.0 | 1.8 | #DIV/0! | 3.79E-02 |
| P02770 | QLKTVMGDFAQFVDK | 0,Dimethyl[AnyN-term](Ethyl[AnyN-term]); | 0.0 | 6.3 | #DIV/0! | 3.79E-02 |
| P02770 | CVADENAENCDKSIHTLFGDK | 1,NO_SMX_SMCT[C]; | 0.0 | 9.7 | #DIV/0! | 3.80E-02 |
| P02770 | LVERRPCFSALTVDETYVPK | 7,HexNAc[C]; | 0.0 | 24.7 | #DIV/0! | 3.80E-02 |
| P07522 | GSMLKPSSLVVVHPLAK | 6,Pro->Cys[P]; | 0.0 | 10.3 | #DIV/0! | 3.81E-02 |
| P24090 | QYGFCK | 5,Carbamidomethyl[C]; | 0.0 | 7.0 | #DIV/0! | 3.82E-02 |
| P02770 | RPCFSALTVDETYVPK | 2,Pro->Tyr[P]; | 0.0 | 5.7 | #DIV/0! | 3.82E-02 |
| P02770 | GLVLIAFSQYLQKCPYEEHIK | 13,Succinyl[K]; | 0.0 | 2.7 | #DIV/0! | 3.83E-02 |
| P02770 | QLKTVMGDFAQFVDK | 0,LG-pyrrole[AnyN-term]; | 0.0 | 5.3 | #DIV/0! | 3.83E-02 |
| P02770 | LQACCDKPVLQK | 5,Diethylphosphate[C]; | 0.0 | 4.2 | #DIV/0! | 3.84E-02 |
| P02761 | PEDGEYFVEYDGGNTFTILK | 9,Unknown_216[E]; | 0.0 | 6.7 | #DIV/0! | 3.85E-02 |
| P02761 | FMQHIDVLENSLGFK | 0,AccQTag[AnyN-term]; | 0.0 | 3.8 | #DIV/0! | 3.87E-02 |
| P02761 | LCEAHGITR | 2,HexNAc[C]; | 0.0 | 3.8 | #DIV/0! | 3.87E-02 |
| P12346 | CPEGSIDSAPVK | 1,Carbamidomethyl[C]; | 0.0 | 12.2 | #DIV/0! | 3.87E-02 |
| P09006 | LDVAETGTEAAAATGVK | 0,Tween20[AnyN-term]; | 0.0 | 6.5 | #DIV/0! | 3.87E-02 |
| P01048 | HLGQSLNCNANVYMRPWENK | 9,Hydroxymethyl[N]; | 0.0 | 3.2 | #DIV/0! | 3.88E-02 |
| P05544 | PDQGKMQQVESSLQPETLKK | 0,Cytopiloyne+water[AnyN-term]; | 0.0 | 3.2 | #DIV/0! | 3.88E-02 |
| P05545 | PDQGKMQQVESSLQPETLKK | 0,Cytopiloyne+water[AnyN-term]; | 0.0 | 3.2 | #DIV/0! | 3.88E-02 |
| P08932 | HLGQSLNCNANVYMRPWENK | 9,Hydroxymethyl[N]; | 0.0 | 3.2 | #DIV/0! | 3.88E-02 |
| P09005 | PDQGKMQQVESSLQPETLKK | 0,Cytopiloyne+water[AnyN-term]; | 0.0 | 3.2 | #DIV/0! | 3.88E-02 |
| P02770 | AEAMCTSFQENPTSFLGHYLHEVAR | 0,AccQTag[AnyN-term]; | 0.0 | 4.0 | #DIV/0! | 3.88E-02 |
| P02770 | LQACCDKPVLQK | 4,Diethylphosphate[C]; | 0.0 | 4.0 | #DIV/0! | 3.88E-02 |
| P22282 | QLKREICYFVVYPDYIEQNIHAVR | 7,ICAT-H_13C(6)[C]; | 0.0 | 3.5 | #DIV/0! | 3.88E-02 |
| P07522 | LFWTDTGMSPR | 8,Oxidation[M]; | 0.0 | 13.7 | #DIV/0! | 3.89E-02 |
| P02770 | TSFQENPTSFLGHYLHEVAR | 1,PET[T]; | 0.0 | 4.8 | #DIV/0! | 3.90E-02 |
| P20759 | MNGQPQENYK | 0,ISD_z+2_ion[AnyN-term]; | 0.0 | 6.0 | #DIV/0! | 3.90E-02 |
| P20760 | MNGQPQENYK | 0,ISD_z+2_ion[AnyN-term]; | 0.0 | 6.0 | #DIV/0! | 3.90E-02 |
| P02770 | YGTVLAEFQPLVEEPK | 0,Carbamidomethyl[AnyN-term]; | 0.0 | 4.3 | #DIV/0! | 3.93E-02 |
| P02770 | VMGDFAQFVDK | 3,Gly->Pro[G]; | 0.0 | 2.3 | #DIV/0! | 3.95E-02 |
| P02770 | YMCENQATISSK | 3,Malonyl[C]; | 0.0 | 2.3 | #DIV/0! | 3.95E-02 |
| P12346 | FATCHLAQAPNHVVVSR | 4,Acetyl[C]; | 0.0 | 2.3 | #DIV/0! | 3.95E-02 |
| P36953 | ADWCQPLK | 4,Carbamidomethyl[C]; | 0.0 | 7.0 | #DIV/0! | 3.95E-02 |
| P20759 | NTPPTMDTDGSYFLYSK | 1,Methyl+Deamidated[N](Asn->Glu[N]); | 0.0 | 9.0 | #DIV/0! | 4.00E-02 |
| P20760 | NTPPTMDTDGSYFLYSK | 1,Methyl+Deamidated[N](Asn->Glu[N]); | 0.0 | 9.0 | #DIV/0! | 4.00E-02 |
| P02770 | PFQRPEAEAMCTSFQENPTSFLGHYLHEVAR | 11,Acetyl[C]; | 0.0 | 9.3 | #DIV/0! | 4.03E-02 |
| P02770 | SFLGHYLHEVAR | 0,Ethylphosphate[AnyN-term]; | 0.0 | 4.2 | #DIV/0! | 4.03E-02 |
| P98158 | TCKVTGSENPLLVVASR | 3,Dap-DSP[K]; | 0.0 | 3.7 | #DIV/0! | 4.03E-02 |
| P02770 | RHPDYSVSLLLR | 0,CLIP_TRAQ_4[AnyN-term]; | 0.0 | 16.0 | #DIV/0! | 4.03E-02 |
| Q03626 | QQNSHGGFSSTQDTVVALDALSK | 3,Asn->Met[N]; | 0.0 | 5.3 | #DIV/0! | 4.06E-02 |
| Q6IE52 | QQNSHGGFSSTQDTVVALDALSK | 3,Asn->Met[N]; | 0.0 | 5.3 | #DIV/0! | 4.06E-02 |
| P02770 | ALTVDETYVPK | 0,Dimethyl[AnyN-term](Ethyl[AnyN-term]); | 0.0 | 12.5 | #DIV/0! | 4.07E-02 |
| P01026 | VHQFFNVGLIQPGSVK | 6,Ammonia-loss[N]; | 0.0 | 1.0 | #DIV/0! | 4.09E-02 |
| P02770 | ELAKYMCENQATISSKLQACCDKPVLQKSQCLAEIEHDNIPADLPSIAADFVEDK | 20,Trioxidation[C]; | 0.0 | 1.0 | #DIV/0! | 4.09E-02 |
| P02770 | FSALTVDETYVPK | 4,Xle->Cys[L]; | 0.0 | 1.0 | #DIV/0! | 4.09E-02 |
| P02770 | TSFQENPTSFLGHYLHEVAR | 8,PhosphoHex[T]; | 0.0 | 1.0 | #DIV/0! | 4.09E-02 |
| P02770 | VTKCCSGSLVER | 4,NEMsulfur[C]; | 0.0 | 1.0 | #DIV/0! | 4.09E-02 |
| P02770 | PYFYAPELLYYAEK | 0,GIST-Quat[AnyN-term]; | 0.0 | 1.0 | #DIV/0! | 4.09E-02 |
| P12346 | KGTDFQLNQLQGK | 10,Xle->Pro[L]; | 0.0 | 2.0 | #DIV/0! | 4.09E-02 |
| P20059 | GAQATWAELSWPHEK | 0,Ethylphosphate[AnyN-term]; | 0.0 | 1.0 | #DIV/0! | 4.09E-02 |
| P27590 | HTEDTTIQVTENGESSQAR | 0,Biotin_Thermo-21328[AnyN-term]; | 0.0 | 1.0 | #DIV/0! | 4.09E-02 |
| P36953 | WCQPLKEDLQNKR | 2,ICAT-C_13C(9)[C]; | 0.0 | 1.0 | #DIV/0! | 4.09E-02 |
| Q9QX79 | PSVLEAATESLAK | 2,Hex(2)[S]; | 0.0 | 2.0 | #DIV/0! | 4.09E-02 |
| P08932 | AQEEGAQEMDCNDETVFQAVDTALK | 10,Asp->Cys[D]; | 0.0 | 2.2 | #DIV/0! | 4.11E-02 |
| P12346 | ATCHLAQAPNHVVVSR | 2,HexNAc(1)NeuAc(1)[T]; | 0.0 | 2.2 | #DIV/0! | 4.11E-02 |
| P12346 | DLLFKDSAFGLLR | 0,Carbamyl[AnyN-term]; | 0.0 | 2.2 | #DIV/0! | 4.11E-02 |
| P20760 | VEVHTAQTHAPEK | 0,SPITC_13C(6)[AnyN-term]; | 0.0 | 2.2 | #DIV/0! | 4.11E-02 |
| P23680 | SSPSIVLGQEQDTYGGGFDK | 3,Pro->Tyr[P]; | 0.0 | 5.5 | #DIV/0! | 4.11E-02 |
| P02761 | NGDWFSIVVASNKR | 0,NDA[AnyN-term]; | 0.0 | 7.8 | #DIV/0! | 4.12E-02 |
| P01835 | DSTYSMSSTLSLTK | 3,Dehydrated[T]; | 0.0 | 3.5 | #DIV/0! | 4.15E-02 |
| P01836 | DSTYSMSSTLSLTK | 3,Dehydrated[T]; | 0.0 | 3.5 | #DIV/0! | 4.15E-02 |
| P02761 | LNGDWFSIVVASNKR | 0,Methyl[AnyN-term]; | 0.0 | 9.0 | #DIV/0! | 4.17E-02 |
| P02770 | RHPYFYAPELLYYAEK | 5,Phe->Glu[F]; | 0.0 | 12.5 | #DIV/0! | 4.18E-02 |
| P02770 | CTSFQENPTSFLGHYLHEVAR | 1,BHT[C]; | 0.0 | 6.5 | #DIV/0! | 4.18E-02 |
| P10959 | LKGEEVAFWTELLAK | 5,Cation_Na[E]; | 0.0 | 3.2 | #DIV/0! | 4.21E-02 |
| P02761 | LDVAKLNGDWFSIVVASNK | 5,PyridoxalPhosphate[K]; | 0.0 | 0.8 | #DIV/0! | 4.22E-02 |
| P02770 | QEPERNECFLQHKDDNPNLPPFQRPEAEAMCTSFQENPTSFLGHYLHEVAR | 29,Ala->Trp[A]; | 0.0 | 0.8 | #DIV/0! | 4.22E-02 |
| P02770 | YFYAPELLYYAEK | 0,Phenylisocyanate_2H(5)[AnyN-term]; | 0.0 | 0.8 | #DIV/0! | 4.22E-02 |
| P02770 | TSFQENPTSFLGHYLHEVAR | 1,PhosphoHex[T]; | 0.0 | 0.8 | #DIV/0! | 4.22E-02 |
| P02770 | DDRAELAKYMCENQATISSK | 0,glycidamide[AnyN-term]; | 0.0 | 1.7 | #DIV/0! | 4.22E-02 |
| P02770 | YMCENQATISSK | 0,glycidamide[AnyN-term]; | 0.0 | 0.8 | #DIV/0! | 4.22E-02 |
| P02770 | HPDYSVSLLLR | 2,Dioxidation[P](Pro->Glu[P]); | 0.0 | 0.8 | #DIV/0! | 4.22E-02 |
| P05545 | KIFSQQADLSR | 3,Phe->Gln[F]; | 0.0 | 0.8 | #DIV/0! | 4.22E-02 |
| P07647 | AYDHNNDLMLLHLSK | 13,Xle->Pro[L]; | 0.0 | 0.8 | #DIV/0! | 4.22E-02 |
| P12346 | WCAVSEHENTK | 2,HMVK[C]; | 0.0 | 0.8 | #DIV/0! | 4.22E-02 |
| P12346 | WCAVSEHENTK | 0,glycidamide[AnyN-term]; | 0.0 | 0.8 | #DIV/0! | 4.22E-02 |
| P12346 | NGDGKEDLIWEILK | 7,Cation_Ca[II][D]; | 0.0 | 0.8 | #DIV/0! | 4.22E-02 |
| P12346 | FATCHLAQAPNHVVVSR | 4,Carbamyl[C]; | 0.0 | 1.7 | #DIV/0! | 4.22E-02 |
| P12346 | VCPEGSIDSAPVK | 2,NO_SMX_SMCT[C]; | 0.0 | 0.8 | #DIV/0! | 4.22E-02 |
| P12346 | LCDLCIGPAKCAPNNREGYNGYTGAFQCLVEKGDVAFVK | 2,Diethylphosphate[C]; | 0.0 | 0.8 | #DIV/0! | 4.22E-02 |
| P12346 | AFQCLVEKGDVAFVK | 4,SulfurDioxide[C]; | 0.0 | 0.8 | #DIV/0! | 4.22E-02 |
| P14046 | DYDMVPLAVPAVALDSSTDR | 4,Oxidation[M]; | 0.0 | 0.8 | #DIV/0! | 4.22E-02 |
| P14046 | QPAPSPEDLALSMGTIK | 5,PhosphoHexNAc[S]; | 0.0 | 0.8 | #DIV/0! | 4.22E-02 |
| P15684 | YLSYTLNPDYIR | 2,Xle->Pro[L]; | 0.0 | 1.7 | #DIV/0! | 4.22E-02 |
| P17475 | FSVNFADSEEAKK | 4,Asn->Phe[N]; | 0.0 | 2.5 | #DIV/0! | 4.22E-02 |
| P20059 | STHPMHSRCNADPGLSALLSDHR | 9,Cys->Gly[C]; | 0.0 | 1.7 | #DIV/0! | 4.22E-02 |
| P20759 | DVLTITLTPK | 6,Dehydrated[T]; | 0.0 | 1.7 | #DIV/0! | 4.22E-02 |
| P20760 | DVLTITLTPK | 6,Dehydrated[T]; | 0.0 | 1.7 | #DIV/0! | 4.22E-02 |
| P60711 | DDDIAALVVDNGSGMCK | 0,Acetyl[ProteinN-term];16,Carbamidomethyl[C]; | 0.0 | 1.7 | #DIV/0! | 4.22E-02 |
| Q01177 | VIPACLPSPNYVVADR | 0,PEITC[AnyN-term]; | 0.0 | 0.8 | #DIV/0! | 4.22E-02 |
| Q03626 | QPAPSPEDLALSMGTIK | 5,PhosphoHexNAc[S]; | 0.0 | 0.8 | #DIV/0! | 4.22E-02 |
| Q62867 | ARVVPIRLDLNDAQYETLFR | 0,Methyl[AnyN-term]; | 0.0 | 0.8 | #DIV/0! | 4.22E-02 |
| Q6IG01 | FLEQQNQVLQTK | 2,Xle->Pro[L]; | 0.0 | 1.7 | #DIV/0! | 4.22E-02 |
| Q6IG03 | FLEQQNQVLQTK | 2,Xle->Pro[L]; | 0.0 | 1.7 | #DIV/0! | 4.22E-02 |
| Q6IMF3 | FLEQQNQVLQTK | 2,Xle->Pro[L]; | 0.0 | 1.7 | #DIV/0! | 4.22E-02 |
| Q03626 | SFVYTKTLMAYAFALAGNQEK | 8,trifluoro[L]; | 0.0 | 2.8 | #DIV/0! | 4.24E-02 |
| P02770 | TSFQENPTSFLGHYLHEVAR | 0,Unknown_248[AnyN-term]; | 0.0 | 6.5 | #DIV/0! | 4.26E-02 |
| P20759 | VSSVFIFPPKPK | 3,Sulfo[S]; | 0.0 | 6.0 | #DIV/0! | 4.27E-02 |
| P02767 | TAESGELHGLTTDEKFTEGVYR | 1,Thr->Asn[T]; | 0.0 | 8.5 | #DIV/0! | 4.28E-02 |
| P12346 | VKHTTIFEVLPQKADR | 2,Phosphoguanosine[K]; | 0.0 | 4.5 | #DIV/0! | 4.29E-02 |
| P02770 | CTSFQENPTSFLGHYLHEVAR | 7,Asn->Met[N]; | 0.0 | 1.3 | #DIV/0! | 4.29E-02 |
| P02770 | LCVLHEKTPVSEK | 2,Malonyl[C]; | 0.0 | 1.3 | #DIV/0! | 4.29E-02 |
| P02770 | FQENPTSFLGHYLHEVAR | 0,Unknown_210[AnyN-term]; | 0.0 | 1.3 | #DIV/0! | 4.29E-02 |
| P05371 | SLLNSLEEAKK | 0,NIC[AnyN-term](ICPL[AnyN-term]); | 0.0 | 1.3 | #DIV/0! | 4.29E-02 |
| P05544 | TAATGVATVIR | 1,Hex(1)HexNAc(1)Phos(1)[T]; | 0.0 | 1.3 | #DIV/0! | 4.29E-02 |
| P07151 | PNIEMSDLSFSK | 0,Unknown_234[AnyN-term]; | 0.0 | 2.7 | #DIV/0! | 4.29E-02 |
| P09005 | TAATGVATVIR | 1,Hex(1)HexNAc(1)Phos(1)[T]; | 0.0 | 1.3 | #DIV/0! | 4.29E-02 |
| P09006 | EVFSTQADLSGITGDKDLMVSQVVHK | 4,azole[S]; | 0.0 | 4.0 | #DIV/0! | 4.29E-02 |
| Q03626 | SIDTSSISGYSLNIK | 0,GIST-Quat_2H(9)[AnyN-term]; | 0.0 | 1.3 | #DIV/0! | 4.29E-02 |
| Q62930 | DHIIDDVISFIR | 1,Asp->Pro[D]; | 0.0 | 1.3 | #DIV/0! | 4.29E-02 |
| Q6IE52 | SIDTSSISGYSLNIK | 0,GIST-Quat_2H(9)[AnyN-term]; | 0.0 | 1.3 | #DIV/0! | 4.29E-02 |
| P12346 | LPEGTTYEEYLGAEYLQAVGNIR | 5,dHex(1)Hex(2)[T]; | 0.0 | 13.2 | #DIV/0! | 4.33E-02 |
| Q9QX79 | TVTVSCEFFESQDQVPGGENPADTQDAK | 6,Carbamidomethyl[C]; | 0.0 | 19.3 | #DIV/0! | 4.34E-02 |
| P05545 | DYVSNQTQGKIAELFSELDER | 10,BHT[K]; | 0.0 | 5.2 | #DIV/0! | 4.34E-02 |
| P10959 | TGDEHSRGNWAHLDQLAALR | 0,C8-QAT[AnyN-term]; | 0.0 | 15.5 | #DIV/0! | 4.34E-02 |
| P20059 | SGAQATWAELSWPHEKVDGALCLEK | 22,Carbamidomethyl[C]; | 0.0 | 35.3 | #DIV/0! | 4.37E-02 |
| P02770 | NPTSFLGHYLHEVAR | 1,Ethyl+Deamidated[N]; | 0.0 | 5.8 | #DIV/0! | 4.42E-02 |
| P17475 | YHSEAFSVNFADSEEAK | 0,NP40[AnyN-term]; | 0.0 | 5.7 | #DIV/0! | 4.45E-02 |
| P02761 | DVAKLNGDWFSIVVASNKR | 0,Diisopropylphosphate[AnyN-term]; | 0.0 | 8.0 | #DIV/0! | 4.45E-02 |
| P02761 | GDWFSIVVASNKR | 0,Phenylisocyanate_2H(5)[AnyN-term]; | 0.0 | 1.5 | #DIV/0! | 4.46E-02 |
| P02761 | TDYDRYVMFHLINFK | 0,Methyl_2H(2)[AnyN-term]; | 0.0 | 3.0 | #DIV/0! | 4.46E-02 |
| P02761 | MQHIDVLENSLGFK | 0,Triton[AnyN-term]; | 0.0 | 1.5 | #DIV/0! | 4.46E-02 |
| P02767 | EPFASGKTAESGELHGLTTDEK | 7,Xlink_DST[56][K]; | 0.0 | 1.5 | #DIV/0! | 4.46E-02 |
| P02770 | TVMGDFAQFVDK | 2,Val->Met[V]; | 0.0 | 3.0 | #DIV/0! | 4.46E-02 |
| P02770 | MCTSFQENPTSFLGHYLHEVAR | 6,Gln->Ala[Q]; | 0.0 | 1.5 | #DIV/0! | 4.46E-02 |
| P02770 | YMCENQATISSK | 1,Dehydrated[Y]; | 0.0 | 1.5 | #DIV/0! | 4.46E-02 |
| P02770 | QEPERNECFLQHKDDNPNLPPFQRPEAEAMCTSFQENPTSFLGHYLHEVAR | 0,ICPL_13C(6)2H(4)[AnyN-term]; | 0.0 | 1.5 | #DIV/0! | 4.46E-02 |
| P10959 | EAELLELTVK | 0,NHS-LC-Biotin[AnyN-term]; | 0.0 | 1.5 | #DIV/0! | 4.46E-02 |
| P12346 | WCAVSEHENTK | 2,Cys->Gln[C]; | 0.0 | 1.5 | #DIV/0! | 4.46E-02 |
| P12346 | PDGTKKPVTEFATCHLAQAPNHVVVSR | 9,PhosphoCytidine[T]; | 0.0 | 1.5 | #DIV/0! | 4.46E-02 |
| P17475 | VEAVPMSLPPQVK | 1,Val->Gln[V]; | 0.0 | 1.5 | #DIV/0! | 4.46E-02 |
| P20759 | PCICTGSEVSSVFIFPPK | 0,Methyl_2H(2)[AnyN-term];2,Carbamidomethyl[C];4,Carbamidomethyl[C]; | 0.0 | 1.5 | #DIV/0! | 4.46E-02 |
| P12346 | KPVTEFATCHLAQAPNHVVVSR | 8,Thr->Arg[T]; | 0.0 | 7.3 | #DIV/0! | 4.47E-02 |
| P02770 | AADKDNCFATEGPNLVAR | 7,DiDehydro[C]; | 0.0 | 18.2 | #DIV/0! | 4.49E-02 |
| P02770 | VADENAENCDKSIHTLFGDK | 9,Glutathione[C]; | 0.0 | 5.0 | #DIV/0! | 4.49E-02 |
| P02761 | LNGDWFSIVVASNKR | 2,Asn->Met[N]; | 0.0 | 17.0 | #DIV/0! | 4.49E-02 |
| P12346 | NQREGVCPEGSIDSAPVK | 7,4-ONE[C]; | 0.0 | 4.0 | #DIV/0! | 4.51E-02 |
| P02770 | NPTSFLGHYLHEVAR | 3,Methylphosphonate[T]; | 0.0 | 3.7 | #DIV/0! | 4.53E-02 |
| P02770 | NCFATEGPNLVAR | 1,Deamidated_18O(1)[N](Delta_H(1)N(-1)18O(1)[N]); | 0.0 | 3.7 | #DIV/0! | 4.53E-02 |
| P02770 | MCTSFQENPTSFLGHYLHEVAR | 0,Hex[AnyN-term]; | 0.0 | 5.3 | #DIV/0! | 4.53E-02 |
| sP02761 | LNGDWFSIVVASNKR | 0,C+12[AnyN-term]; | 0.0 | 5.7 | #DIV/0! | 4.55E-02 |
| P12346 | PDGTKKPVTEFATCHLAQAPNHVVVSR | 13,PhosphoCytidine[T]; | 0.0 | 19.5 | #DIV/0! | 4.59E-02 |
| P24090 | HAFSPVASVESASGEVLHSPK | 21,Lys->AminoadipicAcid[K]; | 0.0 | 5.8 | #DIV/0! | 4.62E-02 |
| P04276 | VCQELSTLGKDDFR | 0,NHS-LC-Biotin[AnyN-term]; | 0.0 | 4.5 | #DIV/0! | 4.63E-02 |
| P02770 | FKDLGEQHFK | 2,Lys->AminoadipicAcid[K]; | 0.0 | 11.5 | #DIV/0! | 4.63E-02 |
| P60711 | VTHTVPIYEGYALPHAILR | 0,Phosphogluconoylation[AnyN-term]; | 0.0 | 4.3 | #DIV/0! | 4.64E-02 |
| P63259 | VTHTVPIYEGYALPHAILR | 0,Phosphogluconoylation[AnyN-term]; | 0.0 | 4.3 | #DIV/0! | 4.64E-02 |
| P12346 | GVCPEGSIDSAPVK | 7,Glucuronyl[S]; | 0.0 | 5.3 | #DIV/0! | 4.65E-02 |
| P02770 | NPTSFLGHYLHEVAR | 4,azole[S]; | 0.0 | 2.8 | #DIV/0! | 4.66E-02 |
| P05545 | AVLDVDETGTEGAAATAVTAALK | 11,Cation_Na[E]; | 0.0 | 2.8 | #DIV/0! | 4.66E-02 |
| P12346 | SSTKDLLFRDDTK | 0,Carbamyl[AnyN-term]; | 0.0 | 11.7 | #DIV/0! | 4.72E-02 |
| P02770 | FLGHYLHEVAR | 0,Hex[AnyN-term]; | 0.0 | 4.0 | #DIV/0! | 4.72E-02 |
| P05545 | QPEDQAEINTGSALFIDK | 0,NDA[AnyN-term]; | 0.0 | 14.2 | #DIV/0! | 4.72E-02 |
| P20759 | VSSVFIFPPKPK | 2,Sulfo[S]; | 0.0 | 6.0 | #DIV/0! | 4.74E-02 |
| P02761 | EKIEENGSMRVFMQHIDVLENSLGFK | 12,Phe->CamCys[F]; | 0.0 | 3.8 | #DIV/0! | 4.78E-02 |
| P02761 | DYDRYVMFHLINFK | 2,Sulfo[Y]; | 0.0 | 1.8 | #DIV/0! | 4.78E-02 |
| P08723 | LCLTSLSFTTNK | 2,Thiadiazole[C]; | 0.0 | 1.8 | #DIV/0! | 4.78E-02 |
| P20059 | PVTSVDAAFR | 0,ICPL_2H(4)[AnyN-term]; | 0.0 | 1.8 | #DIV/0! | 4.78E-02 |
| Q03626 | SSGSLFNNAMK | 0,C+12[AnyN-term]; | 0.0 | 1.8 | #DIV/0! | 4.78E-02 |
| Q6IE52 | SSGSLFNNAMK | 0,C+12[AnyN-term]; | 0.0 | 1.8 | #DIV/0! | 4.78E-02 |
| Q9R0T4 | DNAPIFNPSTYQGQVLENEVGAR | 2,PhosphoHex(2)[N]; | 0.0 | 1.8 | #DIV/0! | 4.78E-02 |
| P12346 | DLKQEDFQLLCPDGTKKPVTEFATCHLAQAPNHVVVSR | 11,DTT[C]; | 0.0 | 11.3 | #DIV/0! | 4.82E-02 |
| P06866 | ATDLKDWVQETMAKN | 12,Oxidation[M]; | 0.0 | 16.0 | #DIV/0! | 4.82E-02 |
| P02761 | KLCEAHGITR | 0,Piperidine[AnyN-term]; | 0.0 | 2.2 | #DIV/0! | 4.82E-02 |
| P12346 | DFQLFGSPLGK | 1,Asp->Cys[D]; | 0.0 | 4.3 | #DIV/0! | 4.82E-02 |
| P22057 | KFITFSKDQGLTEEDIVFLPQPDK | 0,GIST-Quat_2H(3)[AnyN-term]; | 0.0 | 2.2 | #DIV/0! | 4.82E-02 |
| P36953 | ADTTYALPSVSALVSALR | 0,ISD_z+2_ion[AnyN-term]; | 0.0 | 2.2 | #DIV/0! | 4.82E-02 |
| Q62930 | AIEDYVNEFSAR | 0,Methyl[AnyN-term]; | 0.0 | 2.2 | #DIV/0! | 4.82E-02 |
| P02761 | GDWFSIVVASNKR | 0,Hex[AnyN-term]; | 0.0 | 3.3 | #DIV/0! | 4.83E-02 |
| P02770 | SQCLAEIEHDNIPADLPSIAADFVEDK | 3,GG[C](Dicarbamidomethyl[C]); | 0.0 | 3.0 | #DIV/0! | 4.84E-02 |
| P02770 | FKDLGEQHFK | 1,Phe->CamCys[F]; | 0.0 | 3.0 | #DIV/0! | 4.84E-02 |
| P08723 | KEFLLEDGEHVTQVSGTR | 0,3sulfo[AnyN-term]; | 0.0 | 8.8 | #DIV/0! | 4.84E-02 |
| P02770 | CTSFQENPTSFLGHYLHEVAR | 0,Brij58[AnyN-term]; | 0.0 | 7.0 | #DIV/0! | 4.84E-02 |
| P02770 | FQENPTSFLGHYLHEVAR | 6,dHex(1)Hex(1)[T]; | 0.0 | 9.8 | #DIV/0! | 4.86E-02 |
| P02770 | DRAELAKYMCENQATISSK | 8,Dehydrated[Y]; | 0.0 | 6.2 | #DIV/0! | 4.92E-02 |
| P12346 | TEFATCHLAQAPNHVVVSR | 6,GG[C](Dicarbamidomethyl[C]); | 0.0 | 6.2 | #DIV/0! | 4.92E-02 |
| P02770 | IAFSQYLQK | 0,Carbamyl[AnyN-term]; | 0.0 | 15.2 | #DIV/0! | 4.93E-02 |
| P01048 | QEEGAQELNCNDETVFQAVDTALK | 10,Propionamide_2H(3)[C]; | 0.0 | 2.0 | #DIV/0! | 4.93E-02 |
| P01048 | KDGAETLYSFK | 0,Sulfo-NHS-LC-LC-Biotin[AnyN-term]; | 0.0 | 2.0 | #DIV/0! | 4.93E-02 |
| P02770 | TVMGDFAQFVDK | 1,Thr->Cys[T]; | 0.0 | 4.0 | #DIV/0! | 4.93E-02 |
| P08932 | KDGAETLYSFK | 0,Sulfo-NHS-LC-LC-Biotin[AnyN-term]; | 0.0 | 2.0 | #DIV/0! | 4.93E-02 |
| P36953 | KDDRPEDLSLR | 1,Hex(2)[K]; | 0.0 | 2.0 | #DIV/0! | 4.93E-02 |
| Q63621 | DSCFNSPMR | 3,Carbamidomethyl[C]; | 0.0 | 2.0 | #DIV/0! | 4.93E-02 |
| P13635 | EFTDSTFREQVK | 0,Methyl_2H(2)[AnyN-term]; | 0.0 | 6.3 | #DIV/0! | 4.97E-02 |
| P02764 | DVGMDESEIVFVDWTKDK | 2,Val->His[V]; | 0.0 | 3.5 | #DIV/0! | 4.98E-02 |

Appendix Table 3 Differential modifications screened by FC ≥ 1.5 or ≤ 0.67 with P < 0.05 in the 14–15-month-old group

| Uniprot ID | Peptide | Modification | Control group | Experimental group | Flod change | P value |
| --- | --- | --- | --- | --- | --- | --- |
| P83121 | CQTYPDEICAWVVVTTR | 1,Carbamidomethyl[C];9,Carbamidomethyl[C]; | 451.0 | 0.0 | 0.0 | 7.66E-07 |
| P81827 | CQTYPDEICAWVVVTTR | 1,Carbamidomethyl[C];9,Carbamidomethyl[C]; | 451.0 | 0.0 | 0.0 | 7.66E-07 |
| P81828 | CQTYPDEICAWVVVTTR | 1,Carbamidomethyl[C];9,Carbamidomethyl[C]; | 451.0 | 0.0 | 0.0 | 7.66E-07 |
| P02770 | ADKDNCFATEGPNLVAR | 6,DNCB_hapten[C]; | 86.9 | 0.0 | 0.0 | 4.96E-06 |
| P83121 | CQTYPDEICAWVVVTTR | 9,maleimide[C]; | 131.7 | 0.0 | 0.0 | 8.37E-06 |
| P81827 | CQTYPDEICAWVVVTTR | 9,maleimide[C]; | 131.7 | 0.0 | 0.0 | 8.37E-06 |
| P81828 | CQTYPDEICAWVVVTTR | 9,maleimide[C]; | 131.7 | 0.0 | 0.0 | 8.37E-06 |
| P05545 | MQQVESSLQPETLKK | 1,Oxidation[M]; | 43.9 | 0.0 | 0.0 | 1.66E-05 |
| P05544 | MQQVESSLQPETLKK | 1,Oxidation[M]; | 43.9 | 0.0 | 0.0 | 1.66E-05 |
| P09005 | MQQVESSLQPETLKK | 1,Oxidation[M]; | 43.9 | 0.0 | 0.0 | 1.66E-05 |
| P02770 | FPNAEFAEITK | 3,Deamidated[N]; | 172.0 | 0.0 | 0.0 | 2.64E-05 |
| P02770 | ADKDNCFATEGPNLVAR | 6,Nmethylmaleimide[C]; | 54.4 | 0.0 | 0.0 | 3.70E-05 |
| P02770 | YMCENQATISSK | 2,Oxidation[M];3,Carbamidomethyl[C]; | 145.4 | 0.0 | 0.0 | 3.94E-05 |
| Q01177 | CEGETDFICR | 1,Carbamidomethyl[C];9,Carbamidomethyl[C]; | 14.3 | 0.0 | 0.0 | 4.02E-05 |
| P02770 | AADKDNCFATEGPNLVAR | 8,Phe->Trp[F]; | 120.9 | 0.0 | 0.0 | 4.31E-05 |
| Q63678 | CLAYDFYPQR | 1,Carbamidomethyl[C]; | 10.3 | 0.0 | 0.0 | 6.97E-05 |
| P36953 | TINPTVDHCCR | 9,Carbamidomethyl[C];10,Carbamidomethyl[C]; | 6.6 | 0.0 | 0.0 | 7.34E-05 |
| P20760 | SNSMVTLGCLVK | 2,Deamidated[N];9,Carbamidomethyl[C]; | 57.6 | 0.0 | 0.0 | 1.51E-04 |
| P20759 | SNSMVTLGCLVK | 2,Deamidated[N];9,Carbamidomethyl[C]; | 57.6 | 0.0 | 0.0 | 1.51E-04 |
| P05545 | TEGAAATAVTAALK | 0,NDA[AnyN-term]; | 298.0 | 0.0 | 0.0 | 2.54E-04 |
| Q01177 | KQLAAGSIADCLAK | 11,Carbamidomethyl[C]; | 9.7 | 0.0 | 0.0 | 3.02E-04 |
| P48199 | VFSPNVLNWR | 8,Deamidated[N]; | 20.4 | 0.0 | 0.0 | 4.63E-04 |
| P02770 | APQVSTPTLVEAAR | 5,Formyl[S](Ser->Asp[S]); | 16.1 | 0.0 | 0.0 | 5.55E-04 |
| P12346 | ASDSSINWNNLK | 7,Deamidated[N]; | 21.9 | 0.0 | 0.0 | 6.17E-04 |
| P14046 | VTASPQSLCGLR | 9,Carbamidomethyl[C]; | 16.1 | 0.0 | 0.0 | 6.17E-04 |
| Q03626 | VTASPQSLCGLR | 9,Carbamidomethyl[C]; | 16.1 | 0.0 | 0.0 | 6.17E-04 |
| Q6IE52 | VTASPQSLCGLR | 9,Carbamidomethyl[C]; | 16.1 | 0.0 | 0.0 | 6.17E-04 |
| P02770 | DKDNCFATEGPNLVAR | 5,PropylNAGthiazoline[C]; | 47.9 | 0.0 | 0.0 | 6.84E-04 |
| P20760 | SNSMVTLGCLVK | 9,Carbamidomethyl[C]; | 56.0 | 0.0 | 0.0 | 6.86E-04 |
| P20759 | SNSMVTLGCLVK | 9,Carbamidomethyl[C]; | 56.0 | 0.0 | 0.0 | 6.86E-04 |
| P20767 | ANGAPISQGVDTANPTK | 2,Deamidated[N]; | 23.6 | 0.0 | 0.0 | 7.58E-04 |
| P20760 | CKVNSGAFPAPIEK | 1,Carbamidomethyl[C];4,Deamidated[N]; | 14.7 | 0.0 | 0.0 | 8.19E-04 |
| P02770 | FPNAEFAEITK | 8,Cation_Na[E]; | 55.9 | 0.0 | 0.0 | 8.43E-04 |
| P20059 | CNADPGLSALLSDHR | 1,Carbamidomethyl[C]; | 14.9 | 0.0 | 0.0 | 9.19E-04 |
| P12346 | ASDSSINWNNLK | 9,Deamidated[N]; | 89.3 | 0.0 | 0.0 | 1.19E-03 |
| P02770 | FVEDKEVCK | 8,Carbamidomethyl[C]; | 2.4 | 0.0 | 0.0 | 1.30E-03 |
| P04642 | VIGSGCNLDSAR | 6,Carbamidomethyl[C]; | 4.7 | 0.0 | 0.0 | 1.34E-03 |
| P19629 | VIGSGCNLDSAR | 6,Carbamidomethyl[C]; | 4.7 | 0.0 | 0.0 | 1.34E-03 |
| P42123 | VIGSGCNLDSAR | 6,Carbamidomethyl[C]; | 4.7 | 0.0 | 0.0 | 1.34E-03 |
| P02770 | HPDYSVSLLLR | 0,BITC[AnyN-term]; | 16.6 | 0.0 | 0.0 | 1.52E-03 |
| P12346 | DLKQEDFQLLCPDGTK | 11,Carbamidomethyl[C]; | 26.7 | 0.0 | 0.0 | 1.55E-03 |
| Q63621 | GCTEIVNFHNVQPK | 2,Carbamidomethyl[C]; | 4.6 | 0.0 | 0.0 | 1.61E-03 |
| P02770 | HPDYSVSLLLR | 0,C+12[AnyN-term]; | 97.1 | 0.0 | 0.0 | 1.62E-03 |
| P02770 | TVMGDFAQFVDK | 3,Oxidation[M]; | 383.3 | 0.0 | 0.0 | 1.90E-03 |
| P20760 | CKVNSGAFPAPIEK | 1,Carbamidomethyl[C]; | 12.6 | 0.0 | 0.0 | 1.94E-03 |
| P12346 | DCTGNFCLFR | 2,Carbamidomethyl[C];7,Carbamidomethyl[C]; | 55.6 | 0.0 | 0.0 | 1.99E-03 |
| Q01177 | YSPSTHPSEGLEENYCR | 16,Carbamidomethyl[C]; | 6.3 | 0.0 | 0.0 | 2.02E-03 |
| P81827 | ESLDSTGLCR | 9,Carbamidomethyl[C]; | 6.7 | 0.0 | 0.0 | 2.12E-03 |
| P02770 | PPACYGTVLAEFQPLVEEPK | 4,Carbamidomethyl[C]; | 190.6 | 0.0 | 0.0 | 2.28E-03 |
| P02770 | CCAEGDPPACYGTVLAEFQPLVEEPK | 1,CarbamidomethylDTT[C]; | 30.9 | 0.0 | 0.0 | 2.43E-03 |
| Q5FVF9 | QPCLGSDPGCPQDGR | 3,Carbamidomethyl[C];10,Carbamidomethyl[C]; | 7.7 | 0.0 | 0.0 | 2.54E-03 |
| P12346 | EGVCPEGSIDSAPVK | 4,Carbamidomethyl[C]; | 51.1 | 0.0 | 0.0 | 2.55E-03 |
| P12788 | HPEYNKDTLDNDIMLIK | 6,TMT2plex[K]; | 16.9 | 0.0 | 0.0 | 2.65E-03 |
| P48199 | VFSPNVLNWR | 5,Deamidated[N]; | 9.3 | 0.0 | 0.0 | 2.74E-03 |
| P02770 | CCSGSLVER | 1,Carbamidomethyl[C];2,Carbamidomethyl[C]; | 341.3 | 0.0 | 0.0 | 2.91E-03 |
| P02770 | YNEVLTQCCTESDK | 2,Deamidated[N];8,Carbamidomethyl[C];9,Carbamidomethyl[C]; | 8.9 | 0.0 | 0.0 | 2.99E-03 |
| P17475 | AFHHLLQTLNRPDSELQLNTGNGLFVNK | 22,Deamidated[N]; | 36.1 | 0.0 | 0.0 | 3.22E-03 |
| P12346 | KPVTEFATCHLAQAPNHVVVSR | 9,Carbamidomethyl[C];16,Deamidated[N]; | 34.1 | 0.0 | 0.0 | 3.50E-03 |
| Q63041 | ATPLSLCALTAVDQSVLLLKPEAK | 7,Carbamidomethyl[C]; | 3.3 | 0.0 | 0.0 | 3.69E-03 |
| P15083 | KNKETCDVIINTLGK | 6,Carbamidomethyl[C]; | 2.4 | 0.0 | 0.0 | 3.70E-03 |
| P10960 | SLPCDICK | 4,Carbamidomethyl[C];7,Carbamidomethyl[C]; | 7.9 | 0.0 | 0.0 | 3.72E-03 |
| P12346 | LLEACTFHK | 5,Carbamidomethyl[C]; | 22.1 | 0.0 | 0.0 | 3.73E-03 |
| Q9R0T4 | DWVIPPINCPENQK | 9,Carbamidomethyl[C]; | 39.0 | 0.0 | 0.0 | 3.76E-03 |
| P01836 | DSKDSTYSMSSTLSLTK | 0,Propionyl_13C(3)[AnyN-term]; | 25.7 | 0.0 | 0.0 | 3.78E-03 |
| P01835 | DSKDSTYSMSSTLSLTK | 0,Propionyl_13C(3)[AnyN-term]; | 25.7 | 0.0 | 0.0 | 3.78E-03 |
| P02770 | APQVSTPTLVEAAR | 6,Formyl[T](Thr->Glu[T]); | 10.7 | 0.0 | 0.0 | 3.91E-03 |
| P05544 | WKDSLIPR | 0,Formyl[AnyN-term]; | 7.0 | 0.0 | 0.0 | 3.95E-03 |
| P36953 | RPCFEHLK | 3,Carbamidomethyl[C]; | 4.7 | 0.0 | 0.0 | 3.96E-03 |
| P02770 | CCAEGDPPACYGTVLAEFQPLVEEPK | 2,CarbamidomethylDTT[C]; | 19.0 | 0.0 | 0.0 | 3.99E-03 |
| P81828 | DSTGFCHVGR | 6,Carbamidomethyl[C]; | 54.3 | 0.0 | 0.0 | 4.03E-03 |
| P02770 | CFATEGPNLVAR | 1,Carbamidomethyl[C]; | 17.4 | 0.0 | 0.0 | 4.04E-03 |
| P20767 | NSFTCQVTHEGNTVEK | 5,Carbamidomethyl[C]; | 15.1 | 0.0 | 0.0 | 4.09E-03 |
| P12346 | KPVTEFATCHLAQAPNHVVVSR | 9,Carbamidomethyl[C]; | 171.3 | 0.0 | 0.0 | 4.14E-03 |
| P02770 | QEPERNECFLQHK | 8,Carbamidomethyl[C]; | 26.9 | 0.0 | 0.0 | 4.24E-03 |
| P02770 | CCTLPEAQR | 1,Carbamidomethyl[C];2,Carbamidomethyl[C]; | 82.0 | 0.0 | 0.0 | 4.42E-03 |
| P01836 | SKDSTYSMSSTLSLTK | 0,Galactosyl[AnyN-term]; | 3.0 | 0.0 | 0.0 | 4.84E-03 |
| P01835 | SKDSTYSMSSTLSLTK | 0,Galactosyl[AnyN-term]; | 3.0 | 0.0 | 0.0 | 4.84E-03 |
| P02770 | LGEYGFQNAVLVR | 8,Ammonia-loss[N]; | 7.6 | 0.0 | 0.0 | 4.87E-03 |
| P36953 | ADWCQPLKEDLQNKR | 4,Carbamidomethyl[C]; | 12.1 | 0.0 | 0.0 | 4.88E-03 |
| P02770 | EYSRRHPDYSVSLLLR | 2,Tyr->Cys[Y]; | 10.6 | 0.0 | 0.0 | 4.90E-03 |
| P13635 | DIASGLIGPLILCK | 13,Carbamidomethyl[C]; | 27.1 | 0.0 | 0.0 | 5.13E-03 |
| P02770 | AWAVARMSQRFPNAEFAEITK | 0,Pyridylacetyl[AnyN-term](Phenylisocyanate[AnyN-term]); | 16.4 | 0.0 | 0.0 | 5.18E-03 |
| P02770 | TCVADENAENCDKSIHTLFGDK | 2,Carbamidomethyl[C]; | 6.1 | 0.0 | 0.0 | 5.25E-03 |
| Q99PS8 | TSEVVIGQCK | 9,Carbamidomethyl[C]; | 5.7 | 0.0 | 0.0 | 5.30E-03 |
| P05545 | TSMVLVNYLLFK | 3,Oxidation[M]; | 16.9 | 0.0 | 0.0 | 5.31E-03 |
| P05544 | TSMVLVNYLLFK | 3,Oxidation[M]; | 16.9 | 0.0 | 0.0 | 5.31E-03 |
| P02770 | SFLGHYLHEVAR | 0,iTRAQ8plex[AnyN-term]; | 19.4 | 0.0 | 0.0 | 5.35E-03 |
| P02770 | APQVSTPTLVEAAR | 0,Formyl[AnyN-term]; | 10.1 | 0.0 | 0.0 | 5.57E-03 |
| P21704 | SDHYPVEVTLR | 1,HexNAc[S]; | 11.7 | 0.0 | 0.0 | 5.71E-03 |
| P05544 | DYVSNQTQGKIAELFSDLEER | 0,NDA[AnyN-term]; | 17.1 | 0.0 | 0.0 | 5.75E-03 |
| P81828 | TSFDSTGFCHVGR | 9,Carbamidomethyl[C]; | 69.3 | 0.0 | 0.0 | 5.93E-03 |
| P02770 | LAEIEHDNIPADLPSIAADFVEDK | 3,Biotin-PEO-Amine[E]; | 25.1 | 0.0 | 0.0 | 5.95E-03 |
| P12346 | YLGHSYVTAIR | 0,NIC[AnyN-term](ICPL[AnyN-term]); | 17.6 | 0.0 | 0.0 | 6.03E-03 |
| P02770 | TPTLVEAAR | 0,C+12[AnyN-term]; | 21.0 | 0.0 | 0.0 | 6.15E-03 |
| P12346 | CAVSEHENTK | 1,Carbamidomethyl[C]; | 10.3 | 0.0 | 0.0 | 6.31E-03 |
| P02770 | CFATEGPNLVAR | 1,Cys->Gly[C]; | 31.3 | 0.0 | 0.0 | 6.48E-03 |
| P02770 | FQRPEAEAMCTSFQENPTSFLGHYLHEVAR | 9,Met->His[M]; | 255.7 | 0.0 | 0.0 | 6.58E-03 |
| P04276 | QLTSFIEK | 0,Gln->pyro-Glu[AnyN-termQ]; | 8.1 | 0.0 | 0.0 | 6.82E-03 |
| P12346 | NGDGKEDLIWEILK | 1,Methyl+Deamidated[N](Asn->Glu[N]); | 14.6 | 0.0 | 0.0 | 6.90E-03 |
| Q9JJ40 | GSNGYGFYLR | 3,Deamidated[N]; | 17.0 | 0.0 | 0.0 | 7.15E-03 |
| P02770 | FATEGPNLVAR | 0,ICPL_13C(6)2H(4)[AnyN-term]; | 2.7 | 0.0 | 0.0 | 7.19E-03 |
| P20059 | CNADPGLSALLSDHR | 1,Carbamidomethyl[C];2,Deamidated[N]; | 28.1 | 0.0 | 0.0 | 7.51E-03 |
| P05544 | AVLDVDETGTEATAATGVATVIR | 7,Glu->Asn[E]; | 130.9 | 0.0 | 0.0 | 7.51E-03 |
| P05544 | AVLDVDETGTEATAATGVATVIR | 11,Glu->Asn[E]; | 24.7 | 0.0 | 0.0 | 7.59E-03 |
| P01048 | QEEGAQELNCNDETVFQAVDTALK | 0,Gln->pyro-Glu[AnyN-termQ];10,Carbamidomethyl[C]; | 29.6 | 0.0 | 0.0 | 7.64E-03 |
| P98158 | CQTTNICVPR | 1,Carbamidomethyl[C];7,Carbamidomethyl[C]; | 4.1 | 0.0 | 0.0 | 7.78E-03 |
| P02770 | CCTLPEAQR | 1,maleimide[C]; | 21.1 | 0.0 | 0.0 | 7.85E-03 |
| P05545 | IKDLTTPYIRDEELSCSVLELK | 16,Carbamidomethyl[C]; | 1067.0 | 0.0 | 0.0 | 7.87E-03 |
| P02770 | LRDNYGELADCCAK | 12,Carbamidomethyl[C]; | 10.3 | 0.0 | 0.0 | 8.06E-03 |
| P01836 | DSKDSTYSMSSTLSLTK | 3,Methyl_2H(3)+Acetyl_2H(3)[K]; | 21.4 | 0.0 | 0.0 | 8.27E-03 |
| P01835 | DSKDSTYSMSSTLSLTK | 3,Methyl_2H(3)+Acetyl_2H(3)[K]; | 21.4 | 0.0 | 0.0 | 8.27E-03 |
| P02770 | ALTVDETYVPK | 0,C+12[AnyN-term]; | 19.3 | 0.0 | 0.0 | 8.70E-03 |
| P02770 | CKAADKDNCFATEGPNLVAR | 1,Trioxidation[C]; | 2.7 | 0.0 | 0.0 | 8.97E-03 |
| P12346 | CAVSEHENTK | 0,Biotin[AnyN-term]; | 8.6 | 0.0 | 0.0 | 8.97E-03 |
| P02770 | CPYEEHIK | 1,Trioxidation[C]; | 10.6 | 0.0 | 0.0 | 9.01E-03 |
| Q01177 | SLDGYVSTQGASLHSLTK | 0,Pyridylacetyl[AnyN-term](Phenylisocyanate[AnyN-term]); | 5.0 | 0.0 | 0.0 | 9.32E-03 |
| Q64240 | CIQFIYGGCK | 1,Carbamidomethyl[C];9,Carbamidomethyl[C]; | 13.3 | 0.0 | 0.0 | 9.52E-03 |
| P15083 | DDNGRFSVLITGLR | 3,Deamidated[N]; | 8.6 | 0.0 | 0.0 | 9.54E-03 |
| Q63678 | AYLEEECPTMLKK | 7,Carbamidomethyl[C]; | 3.9 | 0.0 | 0.0 | 9.71E-03 |
| P27590 | GERDWMSIVTPAR | 0,Formyl[AnyN-term]; | 6.3 | 0.0 | 0.0 | 9.76E-03 |
| P13635 | DLYSGLIGPLIVCR | 13,Carbamidomethyl[C]; | 31.7 | 0.0 | 0.0 | 9.80E-03 |
| P83121 | PDEICAWVVVTTR | 5,Carbamidomethyl[C]; | 110.6 | 0.0 | 0.0 | 9.91E-03 |
| P81827 | PDEICAWVVVTTR | 5,Carbamidomethyl[C]; | 110.6 | 0.0 | 0.0 | 9.91E-03 |
| P81828 | PDEICAWVVVTTR | 5,Carbamidomethyl[C]; | 110.6 | 0.0 | 0.0 | 9.91E-03 |
| P02770 | DNCFATEGPNLVAR | 3,Atto495Maleimide[C]; | 5.1 | 0.0 | 0.0 | 1.00E-02 |
| P81828 | LALQCFRCTSFDSTGFCHVGR | 8,Carbamidomethyl[C];17,Carbamidomethyl[C]; | 25.0 | 0.0 | 0.0 | 1.03E-02 |
| P02770 | HPDYSVSLLLR | 0,Formyl[AnyN-term]; | 16.0 | 0.0 | 0.0 | 1.04E-02 |
| P81828 | ALQCFRCTSFDSTGFCHVGR | 4,Carbamidomethyl[C];7,GG[C](Dicarbamidomethyl[C]);16,Carbamidomethyl[C]; | 6.9 | 0.0 | 0.0 | 1.05E-02 |
| Q63257 | VLGDPTCFSDYIR | 7,Carbamidomethyl[C]; | 38.7 | 0.0 | 0.0 | 1.06E-02 |
| P12346 | HQTVLENTNGK | 0,Formyl[AnyN-term];9,Deamidated[N]; | 1.9 | 0.0 | 0.0 | 1.07E-02 |
| P02782 | VAHETISFLMK | 0,Propionyl[AnyN-term]; | 12.1 | 0.0 | 0.0 | 1.07E-02 |
| P02782 | ICELVAHETISFLMK | 2,BMP-piperidinol[C]; | 7.3 | 0.0 | 0.0 | 1.07E-02 |
| P08932 | EDFPFLREDCVPLPYGDHGECR | 10,Carbamidomethyl[C];21,Carbamidomethyl[C]; | 6.3 | 0.0 | 0.0 | 1.09E-02 |
| P12346 | GTDFQLNQLQGK | 7,Deamidated[N]; | 22.9 | 0.0 | 0.0 | 1.09E-02 |
| P02770 | YMCENQATISSK | 3,Delta_S(-1)Se(1)[C]; | 7.1 | 0.0 | 0.0 | 1.10E-02 |
| P15083 | SSVTFECDLGR | 7,Carbamidomethyl[C]; | 127.3 | 0.0 | 0.0 | 1.10E-02 |
| P05545 | TGTEGAAATAVTAALK | 4,Hydroxamic_acid[E]; | 20.7 | 0.0 | 0.0 | 1.11E-02 |
| P12346 | DSAFGLLR | 1,Dehydrated[D]; | 13.7 | 0.0 | 0.0 | 1.12E-02 |
| Q03626 | IMQWQDIKTENGLK | 11,Deamidated[N]; | 6.7 | 0.0 | 0.0 | 1.13E-02 |
| P20059 | NPLPAAHETVAK | 1,Deamidated[N]; | 2.1 | 0.0 | 0.0 | 1.13E-02 |
| P12346 | WCAVSEHENTK | 2,Crotonaldehyde[C]; | 2.6 | 0.0 | 0.0 | 1.17E-02 |
| P12346 | ASDSSINWNNLK | 10,Deamidated[N]; | 20.0 | 0.0 | 0.0 | 1.20E-02 |
| P81828 | LALQCFRCTSFDSTGFCHVGR | 5,Carbamidomethyl[C];17,Carbamidomethyl[C]; | 10.4 | 0.0 | 0.0 | 1.23E-02 |
| P24090 | VGQPGDAGAAGPVAPLCPGR | 17,Carbamidomethyl[C]; | 8.3 | 0.0 | 0.0 | 1.24E-02 |
| Q6P7A9 | AVPTQCDVTPNSR | 6,Carbamidomethyl[C]; | 3.4 | 0.0 | 0.0 | 1.24E-02 |
| P01835 | ADYESHNLYTCEVVHK | 0,Formyl[AnyN-term];11,Carbamidomethyl[C]; | 3.6 | 0.0 | 0.0 | 1.28E-02 |
| P07522 | ADLGGMDVK | 6,Oxidation[M]; | 2.7 | 0.0 | 0.0 | 1.30E-02 |
| O70244 | FCTSLSTPPLK | 2,Carbamidomethyl[C]; | 11.1 | 0.0 | 0.0 | 1.30E-02 |
| P07522 | HDLLTMAIFGDK | 6,Oxidation[M]; | 9.3 | 0.0 | 0.0 | 1.32E-02 |
| P36953 | LCFFYNKK | 2,Carbamidomethyl[C]; | 7.6 | 0.0 | 0.0 | 1.33E-02 |
| P02770 | CFATEGPNLVAR | 1,biotinAcrolein298[C]; | 4.7 | 0.0 | 0.0 | 1.33E-02 |
| P27590 | INFECSYPLDMK | 5,Carbamidomethyl[C]; | 25.1 | 0.0 | 0.0 | 1.35E-02 |
| P02761 | NGDWFSIVVASNKR | 0,Pyridylacetyl[AnyN-term](Phenylisocyanate[AnyN-term]); | 2.9 | 0.0 | 0.0 | 1.37E-02 |
| P36953 | SLAMVQQECK | 4,Oxidation[M];9,Carbamidomethyl[C]; | 5.3 | 0.0 | 0.0 | 1.39E-02 |
| Q01460 | EDCPCSEASLCRPIR | 3,Carbamidomethyl[C];5,Carbamidomethyl[C];11,Carbamidomethyl[C]; | 5.4 | 0.0 | 0.0 | 1.41E-02 |
| P02780 | LVKPYVQDHFTEK | 1,Xle->Cys[L]; | 7.7 | 0.0 | 0.0 | 1.44E-02 |
| P02770 | ALTVDETYVPK | 1,Carbonyl[A]; | 81.6 | 0.0 | 0.0 | 1.47E-02 |
| P20767 | NSFTCQVTHEGNTVEK | 5,Carbamidomethyl[C];12,Deamidated[N]; | 6.0 | 0.0 | 0.0 | 1.48E-02 |
| P20760 | KETWQQGNTFTCSVLHEGLHNHHTEK | 12,Carbamidomethyl[C]; | 28.1 | 0.0 | 0.0 | 1.50E-02 |
| P02770 | EDQLKTVMGDFAQFVDK | 0,Propionyl[AnyN-term]; | 7.0 | 0.0 | 0.0 | 1.52E-02 |
| P29598 | RPWCYVQIGLK | 4,Carbamidomethyl[C]; | 13.7 | 0.0 | 0.0 | 1.52E-02 |
| P12346 | HAVVARNGDGKEDLIWEILK | 1,Diethylphosphothione[H]; | 8.6 | 0.0 | 0.0 | 1.53E-02 |
| P02770 | KQTALAELVK | 0,ISD_z+2_ion[AnyN-term]; | 3.7 | 0.0 | 0.0 | 1.53E-02 |
| P02770 | KDNCFATEGPNLVAR | 4,lapachenole[C]; | 1.9 | 0.0 | 0.0 | 1.53E-02 |
| P02770 | CFATEGPNLVAR | 1,BHT[C]; | 1.9 | 0.0 | 0.0 | 1.53E-02 |
| P81828 | QKCQTYPDEICAWVVVTTR | 11,DiDehydro[C]; | 132.0 | 0.0 | 0.0 | 1.53E-02 |
| P01048 | DTALKKYNAELESGNQFVLYR | 0,Dansyl[AnyN-term]; | 3.6 | 0.0 | 0.0 | 1.54E-02 |
| P05545 | DETGTEGAAATAVTAALK | 3,HexNAc(1)NeuAc(1)[T]; | 9.0 | 0.0 | 0.0 | 1.57E-02 |
| P05545 | IAELFSELDER | 0,Formyl[AnyN-term]; | 4.3 | 0.0 | 0.0 | 1.57E-02 |
| P02770 | QENPTSFLGHYLHEVAR | 6,Malonyl[S]; | 8.4 | 0.0 | 0.0 | 1.61E-02 |
| P28826 | QELSIGTNCDR | 9,Carbamidomethyl[C]; | 7.3 | 0.0 | 0.0 | 1.62E-02 |
| P10959 | GNWAHLDQLAALR | 2,Ammonia-loss[N]; | 7.9 | 0.0 | 0.0 | 1.62E-02 |
| P02770 | LRDNYGELADCCAK | 11,Carbamidomethyl[C]; | 4.9 | 0.0 | 0.0 | 1.64E-02 |
| P08932 | QALDMMISRPPGFSPFR | 0,Diethylphosphate[AnyN-term]; | 5.4 | 0.0 | 0.0 | 1.64E-02 |
| P01048 | QALDMMISRPPGFSPFR | 0,Diethylphosphate[AnyN-term]; | 5.4 | 0.0 | 0.0 | 1.64E-02 |
| P09006 | MQQVEASLQPETLR | 1,Oxidation[M]; | 6.4 | 0.0 | 0.0 | 1.65E-02 |
| P02770 | YEKLGEYGFQNAVLVR | 3,Xlink_BuUrBu[317][K]; | 1.7 | 0.0 | 0.0 | 1.67E-02 |
| P12346 | WCAVSEHENTK | 2,Cys->Dha[C]; | 1.7 | 0.0 | 0.0 | 1.67E-02 |
| P02770 | LQACCDKPVLQK | 4,DiDehydro[C]; | 17.9 | 0.0 | 0.0 | 1.69E-02 |
| P05545 | AVLDVDETGTEGAAATAVTAALK | 8,Didehydro[T]; | 4.7 | 0.0 | 0.0 | 1.71E-02 |
| P29598 | KPSSTVDQQGFQCGQK | 13,Carbamidomethyl[C]; | 3.1 | 0.0 | 0.0 | 1.71E-02 |
| P02770 | RLPCVEDYLSAILNR | 4,Carbamidomethyl[C]; | 30.4 | 0.0 | 0.0 | 1.73E-02 |
| P02770 | VGTKCCTLPEAQR | 5,Carbamidomethyl[C];6,Carbamidomethyl[C]; | 10.9 | 0.0 | 0.0 | 1.73E-02 |
| P20761 | APFVCSVVHEGLHNHHVEK | 5,Carbamidomethyl[C]; | 34.6 | 0.0 | 0.0 | 1.73E-02 |
| P02770 | SFQENPTSFLGHYLHEVAR | 0,PyMIC[AnyN-term]; | 13.1 | 0.0 | 0.0 | 1.74E-02 |
| P02770 | DKDNCFATEGPNLVAR | 2,CresylSaligeninPhosphate[K]; | 11.9 | 0.0 | 0.0 | 1.75E-02 |
| P05545 | IFSQQADLSR | 3,Formyl[S](Ser->Asp[S]); | 2.0 | 0.0 | 0.0 | 1.77E-02 |
| P02782 | QICELVAHETISFLMK | 3,ethylsulfonylethyl[C](BEMAD_C[C]); | 2.4 | 0.0 | 0.0 | 1.79E-02 |
| P04276 | VCQELSTLGK | 2,Carbamidomethyl[C]; | 4.7 | 0.0 | 0.0 | 1.79E-02 |
| P98158 | HQCLCEEGYILER | 3,Carbamidomethyl[C];5,Carbamidomethyl[C]; | 2.9 | 0.0 | 0.0 | 1.79E-02 |
| P05545 | TGTEGAAATAVTAALK | 1,Oxidation[T]; | 6.3 | 0.0 | 0.0 | 1.80E-02 |
| P02770 | AMCTSFQENPTSFLGHYLHEVAR | 3,4-ONE+Delta_H(-2)O(-1)[C]; | 8.9 | 0.0 | 0.0 | 1.85E-02 |
| P05545 | AVLDVDETGTEGAAATAVTAALK | 11,Glu->Gln[E]; | 13.9 | 0.0 | 0.0 | 1.86E-02 |
| P07151 | KIPNIEMSDLSFSK | 7,Oxidation[M]; | 3.1 | 0.0 | 0.0 | 1.89E-02 |
| P02770 | YNEVLTQCCTESDK | 8,Carbamidomethyl[C];9,Carbamidomethyl[C]; | 527.7 | 0.0 | 0.0 | 1.90E-02 |
| P05545 | LDVDETGTEGAAATAVTAALK | 0,Tween20[AnyN-term]; | 8.3 | 0.0 | 0.0 | 1.91E-02 |
| P02770 | QENPTSFLGHYLHEVAR | 3,Deamidated[N]; | 5.6 | 0.0 | 0.0 | 1.93E-02 |
| P02770 | TVMGDFAQFVDK | 1,Didehydro[T]; | 23.9 | 0.0 | 0.0 | 1.94E-02 |
| P14740 | KYPLLIDVYAGPCSQK | 13,Carbamidomethyl[C]; | 29.9 | 0.0 | 0.0 | 1.94E-02 |
| P02770 | CCTLPEAQR | 2,maleimide[C]; | 27.6 | 0.0 | 0.0 | 1.94E-02 |
| P13635 | GQFPLSIQPMGVR | 10,Oxidation[M]; | 19.0 | 0.0 | 0.0 | 1.96E-02 |
| P08426 | LNSPATLNSR | 2,Asn->Ser[N]; | 3.3 | 0.0 | 0.0 | 1.96E-02 |
| P02770 | LGEYGFQNAVLVR | 8,Deamidated[N]; | 4.3 | 0.0 | 0.0 | 1.97E-02 |
| P02770 | ENPTSFLGHYLHEVAR | 0,Glu->pyro-Glu[AnyN-termE]; | 12.3 | 0.0 | 0.0 | 1.97E-02 |
| P36373 | NNLLEDEPFAQHR | 0,Formyl[AnyN-term]; | 3.9 | 0.0 | 0.0 | 1.97E-02 |
| P05544 | DSTMEEILEGLK | 0,C+12[AnyN-term]; | 11.6 | 0.0 | 0.0 | 1.97E-02 |
| P36374 | NNLLEDEPFAQHR | 0,Formyl[AnyN-term]; | 3.9 | 0.0 | 0.0 | 1.97E-02 |
| P02770 | FQENPTSFLGHYLHEVAR | 0,CLIP_TRAQ_3[AnyN-term]; | 8.1 | 0.0 | 0.0 | 2.01E-02 |
| P02770 | YNEVLTQCCTESDKAACLTPK | 0,Formyl[AnyN-term];8,GG[C](Dicarbamidomethyl[C]);17,Carbamidomethyl[C]; | 8.1 | 0.0 | 0.0 | 2.01E-02 |
| P02770 | AACLTPKLDAVK | 3,Carbamidomethyl[C]; | 2.9 | 0.0 | 0.0 | 2.02E-02 |
| P17475 | GTEAAGATVVEAVPMSLPPQVK | 15,Oxidation[M]; | 34.4 | 0.0 | 0.0 | 2.02E-02 |
| P02770 | CFATEGPNLVAR | 1,maleimide[C]; | 4.7 | 0.0 | 0.0 | 2.04E-02 |
| P83121 | CQTYPDEICAWVVVTTR | 1,maleimide[C]; | 65.4 | 0.0 | 0.0 | 2.04E-02 |
| P81827 | CQTYPDEICAWVVVTTR | 1,maleimide[C]; | 65.4 | 0.0 | 0.0 | 2.04E-02 |
| P81828 | CQTYPDEICAWVVVTTR | 1,maleimide[C]; | 65.4 | 0.0 | 0.0 | 2.04E-02 |
| P02770 | SFQENPTSFLGHYLHEVAR | 4,Cation_Li[E]; | 6.4 | 0.0 | 0.0 | 2.04E-02 |
| P02782 | EMYNAPPAAVEAK | 0,Unknown_250[AnyN-term]; | 6.4 | 0.0 | 0.0 | 2.04E-02 |
| P12346 | EYLGAEYLQAVGNIR | 3,Xle->Met[L]; | 1.9 | 0.0 | 0.0 | 2.05E-02 |
| P01836 | RDGVLDSVTDQDSK | 12,Asp->Thr[D]; | 1.9 | 0.0 | 0.0 | 2.05E-02 |
| P01835 | RDGVLDSVTDQDSK | 12,Asp->Thr[D]; | 1.9 | 0.0 | 0.0 | 2.05E-02 |
| Q64240 | QGPCRAFAELWAFDAAQGK | 4,BHT[C]; | 7.7 | 0.0 | 0.0 | 2.07E-02 |
| P02770 | FPNAEFAEITK | 3,Ammonia-loss[N]; | 15.3 | 0.0 | 0.0 | 2.07E-02 |
| P20059 | CNADPGLSALLSDHR | 0,Pyro-carbamidomethyl[AnyN-termC]; | 54.1 | 0.0 | 0.0 | 2.08E-02 |
| Q9ESG3 | ELCRPDAENAFK | 3,Carbamidomethyl[C]; | 7.3 | 0.0 | 0.0 | 2.10E-02 |
| P13432 | NPPTQLHSTEQANTK | 7,His->Xle[H]; | 10.4 | 0.0 | 0.0 | 2.13E-02 |
| P02770 | PCVEDYLSAILNR | 0,Formyl[AnyN-term]; | 3.4 | 0.0 | 0.0 | 2.14E-02 |
| P02761 | VFMQHIDVLENSLGFK | 0,Formyl[AnyN-term]; | 9.4 | 0.0 | 0.0 | 2.15E-02 |
| P14046 | GMYESLPVVAVK | 2,Oxidation[M]; | 4.3 | 0.0 | 0.0 | 2.17E-02 |
| Q04807 | LTMLSQAVQK | 3,Oxidation[M]; | 6.6 | 0.0 | 0.0 | 2.18E-02 |
| P28826 | LSQVLSGPENDHSNMGQCK | 18,Carbamidomethyl[C]; | 11.0 | 0.0 | 0.0 | 2.20E-02 |
| P02770 | TCVADENAENCDK | 11,Carbamidomethyl[C]; | 11.4 | 0.0 | 0.0 | 2.21E-02 |
| P02770 | HPYFYAPELLYYAEK | 0,Formyl[AnyN-term]; | 23.3 | 0.0 | 0.0 | 2.21E-02 |
| P12346 | YLGAEYLQAVGNIR | 0,BITC[AnyN-term]; | 2.3 | 0.0 | 0.0 | 2.22E-02 |
| P02770 | ECCHGDLLECADDRAELAK | 2,Carbamidomethyl[C];3,Carbamidomethyl[C]; | 33.4 | 0.0 | 0.0 | 2.22E-02 |
| P02770 | LYEKLGEYGFQNAVLVR | 4,Lys->Asn[K]; | 7.1 | 0.0 | 0.0 | 2.22E-02 |
| P02770 | RHPDYSVSLLLR | 11,Xle->Cys[L]; | 10.0 | 0.0 | 0.0 | 2.24E-02 |
| P02770 | TSFLGHYLHEVAR | 1,Oxidation[T]; | 7.3 | 0.0 | 0.0 | 2.25E-02 |
| Q03626 | SSGSLFNNAMK | 7,Deamidated[N]; | 3.6 | 0.0 | 0.0 | 2.26E-02 |
| Q6IE52 | SSGSLFNNAMK | 7,Deamidated[N]; | 3.6 | 0.0 | 0.0 | 2.26E-02 |
| P02770 | RHPYFYAPELLYYAEK | 6,Acetyl[Y]; | 2.1 | 0.0 | 0.0 | 2.28E-02 |
| P02770 | AADKDNCFATEGPNLVAR | 5,Dehydrated[D]; | 2.1 | 0.0 | 0.0 | 2.28E-02 |
| P05545 | QVVHKAVLDVDETGTEGAAATAVTAALK | 5,Xlink_DST[132][K]; | 2.1 | 0.0 | 0.0 | 2.28E-02 |
| P10959 | TEAELLELTVK | 0,Unknown_250[AnyN-term]; | 5.0 | 0.0 | 0.0 | 2.30E-02 |
| P05544 | IAELFSDLEER | 0,C+12[AnyN-term]; | 8.1 | 0.0 | 0.0 | 2.30E-02 |
| P01836 | KDSTYSMSSTLSLTK | 4,Pent(2)[T]; | 5.1 | 0.0 | 0.0 | 2.30E-02 |
| P01835 | KDSTYSMSSTLSLTK | 4,Pent(2)[T]; | 5.1 | 0.0 | 0.0 | 2.30E-02 |
| P36373 | MLLHLSQPADITDGVK | 0,Galactosyl[AnyN-term]; | 5.7 | 0.0 | 0.0 | 2.31E-02 |
| P00758 | MLLHLSQPADITDGVK | 0,Galactosyl[AnyN-term]; | 5.7 | 0.0 | 0.0 | 2.31E-02 |
| P02770 | QRPEAEAMCTSFQENPTSFLGHYLHEVAR | 9,Carbamidomethyl[C]; | 3.3 | 0.0 | 0.0 | 2.31E-02 |
| P08932 | FQPIPMDSSDLKPVLK | 0,AccQTag[AnyN-term]; | 6.1 | 0.0 | 0.0 | 2.32E-02 |
| P01048 | FQPIPMDSSDLKPVLK | 0,AccQTag[AnyN-term]; | 6.1 | 0.0 | 0.0 | 2.32E-02 |
| P98158 | IESAWMNGEHR | 7,Deamidated[N]; | 3.6 | 0.0 | 0.0 | 2.42E-02 |
| P02770 | AADKDNCFATEGPNLVAR | 0,Ethylphosphate[AnyN-term]; | 5.9 | 0.0 | 0.0 | 2.42E-02 |
| P02770 | NCFATEGPNLVAR | 0,NIC[AnyN-term](ICPL[AnyN-term]); | 5.6 | 0.0 | 0.0 | 2.43E-02 |
| P02770 | LGEYGFQNAVLVR | 0,Formyl[AnyN-term]; | 49.9 | 0.0 | 0.0 | 2.43E-02 |
| P07522 | NLDPASVPPR | 0,Ethylphosphate[AnyN-term]; | 9.6 | 0.0 | 0.0 | 2.47E-02 |
| P02770 | SFQENPTSFLGHYLHEVAR | 2,Phe->Trp[F]; | 3.7 | 0.0 | 0.0 | 2.48E-02 |
| P02770 | MCTSFQENPTSFLGHYLHEVAR | 2,Myristoyl[C]; | 1.6 | 0.0 | 0.0 | 2.48E-02 |
| P02770 | LPEAQRLPCVEDYLSAILNR | 0,TNBS[AnyN-term]; | 1.4 | 0.0 | 0.0 | 2.49E-02 |
| P14659 | LLQDFFNGK | 7,Deamidated[N]; | 13.6 | 0.0 | 0.0 | 2.51E-02 |
| P63018 | LLQDFFNGK | 7,Deamidated[N]; | 13.6 | 0.0 | 0.0 | 2.51E-02 |
| Q9QX79 | IFYETVHGQCK | 10,Carbamidomethyl[C]; | 4.9 | 0.0 | 0.0 | 2.52E-02 |
| P12346 | TAGWNIPMGLLFSR | 0,ISD_z+2_ion[AnyN-term]; | 2.6 | 0.0 | 0.0 | 2.53E-02 |
| P07314 | TPALCEVFCR | 5,Carbamidomethyl[C];9,Carbamidomethyl[C]; | 20.6 | 0.0 | 0.0 | 2.55E-02 |
| P02770 | CFATEGPNLVAR | 1,Thiazolidine[C]; | 20.1 | 0.0 | 0.0 | 2.56E-02 |
| P13635 | TFCSEPEKVDKDNEDFQESNR | 3,Carbamidomethyl[C]; | 3.6 | 0.0 | 0.0 | 2.57E-02 |
| P02770 | FKAETFTFHSDICTLPDKEK | 13,CarbamidomethylDTT[C]; | 7.4 | 0.0 | 0.0 | 2.59E-02 |
| P02770 | ENPTSFLGHYLHEVAR | 0,Unknown_216[AnyN-term]; | 78.0 | 0.0 | 0.0 | 2.60E-02 |
| P12346 | VAQEHFGK | 0,Formyl[AnyN-term]; | 2.3 | 0.0 | 0.0 | 2.60E-02 |
| P20059 | YYCFQGNKFLR | 3,Carbamidomethyl[C]; | 4.9 | 0.0 | 0.0 | 2.60E-02 |
| Q9JJ19 | IVEVNGVCMEGK | 5,Deamidated[N];8,Carbamidomethyl[C];9,Oxidation[M]; | 3.0 | 0.0 | 0.0 | 2.63E-02 |
| P19468 | GYVSDIDCR | 8,Carbamidomethyl[C]; | 3.0 | 0.0 | 0.0 | 2.63E-02 |
| P05545 | PDKNVVFSPLSISAALAILSLGAK | 0,TNBS[AnyN-term]; | 8.1 | 0.0 | 0.0 | 2.67E-02 |
| P02770 | ADKDNCFATEGPNLVAR | 6,PEITC[C]; | 3.1 | 0.0 | 0.0 | 2.68E-02 |
| P02770 | HPDYSVSLLLR | 1,Aspartylurea[H]; | 3.1 | 0.0 | 0.0 | 2.68E-02 |
| Q5XI43 | IDDLEPADEGIYSCHLHHHYCGLHER | 14,Carbamidomethyl[C];21,Carbamidomethyl[C]; | 9.4 | 0.0 | 0.0 | 2.71E-02 |
| P02770 | SIHTLFGDKLCAIPKLR | 11,Carbamidomethyl[C]; | 2.9 | 0.0 | 0.0 | 2.74E-02 |
| P12346 | HQTVLENTNGKNTAAWAK | 7,Deamidated[N];12,Deamidated[N]; | 2.9 | 0.0 | 0.0 | 2.74E-02 |
| P02770 | AETFTFHSDICTLPDKEK | 8,Carbonyl[S]; | 2.4 | 0.0 | 0.0 | 2.78E-02 |
| P22282 | LDNCPFEEQTEQLKR | 1,trifluoro[L]; | 2.4 | 0.0 | 0.0 | 2.78E-02 |
| P08721 | SQESDEAIKVIPVAQR | 4,Phospho[S]; | 2.4 | 0.0 | 0.0 | 2.78E-02 |
| P02770 | DQLKTVMGDFAQFVDK | 2,spermine[Q]; | 4.6 | 0.0 | 0.0 | 2.80E-02 |
| P17475 | FPKLSISGTYNLK | 0,3sulfo[AnyN-term]; | 17.9 | 0.0 | 0.0 | 2.82E-02 |
| P12346 | HQTVLENTNGKNTAAWAK | 9,Deamidated[N];12,Deamidated[N]; | 7.7 | 0.0 | 0.0 | 2.84E-02 |
| P02770 | QLKTVMGDFAQFVDK | 4,PhosphoCytidine[T]; | 9.3 | 0.0 | 0.0 | 2.89E-02 |
| P02770 | LQACCDKPVLQK | 5,Carbamidomethyl[C]; | 5.6 | 0.0 | 0.0 | 2.89E-02 |
| P02770 | RHPYFYAPELLYYAEK | 9,Cation_Na[E]; | 5.4 | 0.0 | 0.0 | 2.89E-02 |
| P50123 | MNFAEEEPSKK | 0,Acetyl[ProteinN-term]; | 3.6 | 0.0 | 0.0 | 2.90E-02 |
| O70244 | NGDSPGSPVIGR | 1,Deamidated[N]; | 4.6 | 0.0 | 0.0 | 2.90E-02 |
| P02782 | CELVAHETISFLMK | 1,Cytopiloyne[C]; | 33.9 | 0.0 | 0.0 | 2.97E-02 |
| P02770 | VGTKCCTLPEAQR | 0,Formyl[AnyN-term];5,Carbamidomethyl[C];6,Carbamidomethyl[C]; | 1.7 | 0.0 | 0.0 | 3.00E-02 |
| P05545 | NPDKNVVFSPLSISAALAILSLGAK | 11,Xle->Val[L]; | 5.1 | 0.0 | 0.0 | 3.00E-02 |
| P01835 | ADYESHNLYTCEVVHK | 7,Deamidated[N];11,Carbamidomethyl[C]; | 1.7 | 0.0 | 0.0 | 3.00E-02 |
| P07522 | RLFWTDTGMSPR | 9,Oxidation[M]; | 2.3 | 0.0 | 0.0 | 3.00E-02 |
| P15684 | VVATTQMQAADAR | 7,Oxidation[M]; | 1.7 | 0.0 | 0.0 | 3.00E-02 |
| P09006 | LDVAETGTEAAAATGVK | 6,Hep[T]; | 8.4 | 0.0 | 0.0 | 3.01E-02 |
| P02770 | MSQRFPNAEFAEITK | 2,O-Et-N-diMePhospho[S]; | 10.4 | 0.0 | 0.0 | 3.03E-02 |
| P02770 | RAELAKYMCENQATISSK | 9,NEIAA_2H(5)[C]; | 4.7 | 0.0 | 0.0 | 3.04E-02 |
| P07522 | HDLLTMAIFGDK | 5,Dehydrated[T]; | 4.9 | 0.0 | 0.0 | 3.05E-02 |
| P12346 | LPEGTTYEEYLGAEYLQAVGNIR | 9,Glu->Asn[E]; | 7.7 | 0.0 | 0.0 | 3.06E-02 |
| P05544 | AVLDVDETGTEATAATGVATVIR | 13,Dehydrated[T]; | 12.0 | 0.0 | 0.0 | 3.06E-02 |
| P02761 | LCEAHGITR | 0,Formyl[AnyN-term];2,Carbamidomethyl[C]; | 10.7 | 0.0 | 0.0 | 3.06E-02 |
| P05544 | MPKFSISTDYSLK | 0,SPITC[AnyN-term]; | 3.6 | 0.0 | 0.0 | 3.06E-02 |
| Q9EQV6 | LLNGRPPLGFLNPR | 3,Deamidated[N]; | 3.3 | 0.0 | 0.0 | 3.08E-02 |
| P10959 | NGNPNGEGLPHWPK | 3,Deamidated[N];5,Deamidated[N]; | 3.0 | 0.0 | 0.0 | 3.09E-02 |
| P09006 | DVAETGTEAAAATGVK | 0,Unknown_306[AnyN-term]; | 10.0 | 0.0 | 0.0 | 3.10E-02 |
| P12346 | HQTVLENTNGK | 0,Formyl[AnyN-term]; | 2.7 | 0.0 | 0.0 | 3.11E-02 |
| P02770 | AADKDNCFATEGPNLVAR | 10,Hex(1)NeuGc(1)[T]; | 3.4 | 0.0 | 0.0 | 3.18E-02 |
| P02770 | PACYGTVLAEFQPLVEEPK | 4,Tyr->Gly[Y]; | 3.4 | 0.0 | 0.0 | 3.18E-02 |
| P02770 | CENQATISSK | 1,BADGE[C]; | 1.9 | 0.0 | 0.0 | 3.24E-02 |
| P02770 | AADKDNCFATEGPNLVAR | 6,Methyl+Deamidated[N](Asn->Glu[N]); | 3.7 | 0.0 | 0.0 | 3.24E-02 |
| P17475 | MQHLEQTLTK | 0,Formyl[AnyN-term];1,Oxidation[M]; | 1.9 | 0.0 | 0.0 | 3.24E-02 |
| P14046 | MVLVPSQLYTETPEK | 6,Glucuronyl[S]; | 1.9 | 0.0 | 0.0 | 3.24E-02 |
| Q03626 | MVLVPSQLYTETPEK | 6,Glucuronyl[S]; | 1.9 | 0.0 | 0.0 | 3.24E-02 |
| Q6IE52 | MVLVPSQLYTETPEK | 6,Glucuronyl[S]; | 1.9 | 0.0 | 0.0 | 3.24E-02 |
| P02770 | MCENQATISSK | 2,Carbamidomethyl[C]; | 5.3 | 0.0 | 0.0 | 3.26E-02 |
| P81828 | QCFRCTSFDSTGFCHVGR | 0,Gln->pyro-Glu[AnyN-termQ];2,GG[C](Dicarbamidomethyl[C]);14,Carbamidomethyl[C]; | 2.0 | 0.0 | 0.0 | 3.27E-02 |
| P60711 | IEHGIVTNWDDMEK | 0,LG-anhyropyrrole[AnyN-term]; | 2.0 | 0.0 | 0.0 | 3.27E-02 |
| P63259 | IEHGIVTNWDDMEK | 0,LG-anhyropyrrole[AnyN-term]; | 2.0 | 0.0 | 0.0 | 3.27E-02 |
| P02782 | QICELVAHETISFLMK | 0,PyMIC[AnyN-term]; | 5.0 | 0.0 | 0.0 | 3.29E-02 |
| P02770 | MGDFAQFVDK | 1,Oxidation[M]; | 4.3 | 0.0 | 0.0 | 3.29E-02 |
| P12346 | NGDGKEDLIWEILK | 1,Ammonia-loss[N]; | 36.3 | 0.0 | 0.0 | 3.31E-02 |
| P02770 | AMCTSFQENPTSFLGHYLHEVAR | 3,Carbamidomethyl[C]; | 26.9 | 0.0 | 0.0 | 3.31E-02 |
| P02770 | CFATEGPNLVAR | 1,DeStreak[C]; | 1.6 | 0.0 | 0.0 | 3.33E-02 |
| P05545 | AVLDVDETGTEGAAATAVTAALK | 10,Thr->Pro[T]; | 1.6 | 0.0 | 0.0 | 3.33E-02 |
| P05544 | IAELFSDLEER | 0,Formyl[AnyN-term]; | 1.6 | 0.0 | 0.0 | 3.33E-02 |
| P02770 | DQLKTVMGDFAQFVDK | 0,Triton[AnyN-term]; | 8.3 | 0.0 | 0.0 | 3.33E-02 |
| P05545 | IFSQQADLSR | 0,Formyl[AnyN-term]; | 12.7 | 0.0 | 0.0 | 3.34E-02 |
| P02770 | LGEYGFQNAVLVR | 4,Dehydrated[Y]; | 21.4 | 0.0 | 0.0 | 3.36E-02 |
| P12346 | HQTVLENTNGKNTAAWAK | 9,Ammonia-loss[N]; | 5.7 | 0.0 | 0.0 | 3.40E-02 |
| P12788 | KDTLDNDIMLIK | 0,NIC[AnyN-term](ICPL[AnyN-term]); | 2.7 | 0.0 | 0.0 | 3.41E-02 |
| P12346 | HQTVLENTNGKNTAAWAK | 7,Deamidated[N];9,Deamidated[N]; | 6.6 | 0.0 | 0.0 | 3.43E-02 |
| P98158 | GSYECFCVDGFK | 5,Carbamidomethyl[C];7,Carbamidomethyl[C]; | 8.6 | 0.0 | 0.0 | 3.44E-02 |
| P02770 | RPCFSALTVDETYVPK | 3,NEMsulfurWater[C]; | 5.9 | 0.0 | 0.0 | 3.47E-02 |
| P22282 | LDNCPFEEQTEQLKR | 4,Carbamidomethyl[C]; | 66.7 | 0.0 | 0.0 | 3.48E-02 |
| P05544 | AELFSDLEER | 0,Phenylisocyanate_2H(5)[AnyN-term]; | 1.3 | 0.0 | 0.0 | 3.49E-02 |
| P02770 | LQACCDKPVLQK | 5,BEMAD_C_2H(6)[C]; | 7.3 | 0.0 | 0.0 | 3.51E-02 |
| P10960 | GCSFLPDPYQK | 2,Carbamidomethyl[C]; | 13.6 | 0.0 | 0.0 | 3.51E-02 |
| P02770 | SFQENPTSFLGHYLHEVAR | 1,O-Et-N-diMePhospho[S]; | 1.4 | 0.0 | 0.0 | 3.53E-02 |
| P02770 | NPTSFLGHYLHEVAR | 5,Phe->Arg[F]; | 1.4 | 0.0 | 0.0 | 3.53E-02 |
| P02770 | VGTKCCTLPEAQR | 5,GG[C](Dicarbamidomethyl[C]);6,Carbamidomethyl[C]; | 1.4 | 0.0 | 0.0 | 3.53E-02 |
| Q9JJ19 | LVEVNGENVEK | 5,Deamidated[N]; | 1.4 | 0.0 | 0.0 | 3.53E-02 |
| P81828 | LQCFRCTSFDSTGFCHVGR | 3,Carbamidomethyl[C];15,Carbamidomethyl[C]; | 3.7 | 0.0 | 0.0 | 3.55E-02 |
| P02770 | YMCENQATISSK | 1,sulfo+amino[Y]; | 14.0 | 0.0 | 0.0 | 3.60E-02 |
| P12346 | LPEGTTYEEYLGAEYLQAVGNIR | 10,Dehydrated[Y]; | 126.6 | 0.0 | 0.0 | 3.61E-02 |
| P02770 | PCVEDYLSAILNR | 2,Carbamidomethyl[C]; | 14.6 | 0.0 | 0.0 | 3.63E-02 |
| P02770 | ENPTSFLGHYLHEVAR | 0,Carbamyl[AnyN-term]; | 2.1 | 0.0 | 0.0 | 3.66E-02 |
| P02770 | DNCFATEGPNLVAR | 0,Amidine[AnyN-term]; | 2.1 | 0.0 | 0.0 | 3.66E-02 |
| P02770 | AADKDNCFATEGPNLVAR | 0,NIC[AnyN-term](ICPL[AnyN-term]); | 2.1 | 0.0 | 0.0 | 3.66E-02 |
| P17475 | SSNLADFAFSLYR | 1,Isopropylphospho[S]; | 2.1 | 0.0 | 0.0 | 3.66E-02 |
| P14046 | ALMAYAFALAGNQEK | 0,C+12[AnyN-term]; | 2.1 | 0.0 | 0.0 | 3.66E-02 |
| P14046 | AHHTAYAVYSLSK | 0,ESP_2H(10)[AnyN-term]; | 2.1 | 0.0 | 0.0 | 3.66E-02 |
| Q6IE52 | AHHTAYAVYSLSK | 0,ESP_2H(10)[AnyN-term]; | 2.1 | 0.0 | 0.0 | 3.66E-02 |
| P02764 | KPDLSPELR | 0,Formyl[AnyN-term]; | 2.1 | 0.0 | 0.0 | 3.66E-02 |
| Q63041 | QNNICFDNWWVDEYHTQADHSAAR | 5,Carbamidomethyl[C]; | 2.1 | 0.0 | 0.0 | 3.66E-02 |
| Q64319 | KQCYFHQFLK | 3,Carbamidomethyl[C]; | 4.3 | 0.0 | 0.0 | 3.66E-02 |
| P12346 | EGVCPEGSIDSAPVK | 0,Formyl[AnyN-term];4,Carbamidomethyl[C]; | 5.3 | 0.0 | 0.0 | 3.66E-02 |
| P05545 | AVLDVDETGTEGAAATAVTAALK | 10,Didehydro[T]; | 11.0 | 0.0 | 0.0 | 3.66E-02 |
| P02770 | YMCENQATISSK | 0,Acetyl_2H(3)[AnyN-term]; | 9.0 | 0.0 | 0.0 | 3.66E-02 |
| P02770 | NPTSFLGHYLHEVAR | 4,Phosphopantetheine[S]; | 12.9 | 0.0 | 0.0 | 3.67E-02 |
| Q64240 | AFAELWAFDAAQGK | 4,Glu->Gln[E]; | 4.0 | 0.0 | 0.0 | 3.68E-02 |
| P02770 | AADKDNCFATEGPNLVAR | 7,Cys->Gly[C]; | 3.3 | 0.0 | 0.0 | 3.68E-02 |
| P83121 | CQTYPDEICAWVVVTTR | 0,NDA[AnyN-term];1,Carbamidomethyl[C];9,Carbamidomethyl[C]; | 133.9 | 0.0 | 0.0 | 3.72E-02 |
| P81827 | CQTYPDEICAWVVVTTR | 0,NDA[AnyN-term];1,Carbamidomethyl[C];9,Carbamidomethyl[C]; | 133.9 | 0.0 | 0.0 | 3.72E-02 |
| P81828 | CQTYPDEICAWVVVTTR | 0,NDA[AnyN-term];1,Carbamidomethyl[C];9,Carbamidomethyl[C]; | 133.9 | 0.0 | 0.0 | 3.72E-02 |
| P05544 | GTEATAATGVATVIR | 1,Gly->Met[G]; | 46.0 | 0.0 | 0.0 | 3.72E-02 |
| P09005 | GTEATAATGVATVIR | 1,Gly->Met[G]; | 46.0 | 0.0 | 0.0 | 3.72E-02 |
| O70244 | QCMNDFLEVR | 2,Carbamidomethyl[C]; | 4.7 | 0.0 | 0.0 | 3.73E-02 |
| P02770 | DKCCKAADKDNCFATEGPNLVAR | 12,Dioxidation[C]; | 1.7 | 0.0 | 0.0 | 3.75E-02 |
| P08649 | EPFLSCCK | 6,Carbamidomethyl[C];7,Carbamidomethyl[C]; | 1.7 | 0.0 | 0.0 | 3.75E-02 |
| Q01177 | CQEWAAQEPHSHR | 1,Dioxidation[C]; | 3.1 | 0.0 | 0.0 | 3.78E-02 |
| P15083 | CHILPSHDEGAR | 1,Cys->Gly[C]; | 3.1 | 0.0 | 0.0 | 3.78E-02 |
| P15684 | SFPCFDEPAMK | 4,Carbamidomethyl[C]; | 7.0 | 0.0 | 0.0 | 3.78E-02 |
| P02770 | MCTSFQENPTSFLGHYLHEVAR | 4,Ser->His[S]; | 11.3 | 0.0 | 0.0 | 3.78E-02 |
| P02770 | EDQLKTVMGDFAQFVDK | 0,Diethyl[AnyN-term]; | 4.9 | 0.0 | 0.0 | 3.79E-02 |
| P02770 | YSRRHPDYSVSLLLR | 1,PhosphoCytidine[Y]; | 31.7 | 0.0 | 0.0 | 3.81E-02 |
| P02770 | DNCFATEGPNLVAR | 0,Piperidine[AnyN-term]; | 4.0 | 0.0 | 0.0 | 3.82E-02 |
| Q62930 | SCADALGDR | 2,Carbamidomethyl[C]; | 2.0 | 0.0 | 0.0 | 3.82E-02 |
| Q63041 | GGENCIKAEDITHNGIVYTPK | 5,Carbamidomethyl[C];14,Deamidated[N]; | 3.0 | 0.0 | 0.0 | 3.82E-02 |
| P08649 | VPQPACREPFLSCCK | 6,Carbamidomethyl[C];13,Carbamidomethyl[C];14,Carbamidomethyl[C]; | 3.0 | 0.0 | 0.0 | 3.82E-02 |
| P05545 | MQQVESSLQPETLKK | 0,ISD_z+2_ion[AnyN-term]; | 2.6 | 0.0 | 0.0 | 3.83E-02 |
| P05544 | MQQVESSLQPETLKK | 0,ISD_z+2_ion[AnyN-term]; | 2.6 | 0.0 | 0.0 | 3.83E-02 |
| P09005 | MQQVESSLQPETLKK | 0,ISD_z+2_ion[AnyN-term]; | 2.6 | 0.0 | 0.0 | 3.83E-02 |
| P01836 | GVLDSVTDQDSKDSTYSMSSTLSLTK | 4,Cation_Ag[D]; | 6.1 | 0.0 | 0.0 | 3.84E-02 |
| P01835 | GVLDSVTDQDSKDSTYSMSSTLSLTK | 4,Cation_Ag[D]; | 6.1 | 0.0 | 0.0 | 3.84E-02 |
| P20767 | NSFTCQVTHEGNTVEK | 1,Deamidated[N];5,Carbamidomethyl[C]; | 10.6 | 0.0 | 0.0 | 3.88E-02 |
| P81828 | QKCQTYPDEICAWVVVTTR | 3,BEMAD_C_2H(6)[C]; | 12.7 | 0.0 | 0.0 | 3.90E-02 |
| P02770 | CPYEEHIK | 0,Formyl[AnyN-term];1,Carbamidomethyl[C]; | 7.1 | 0.0 | 0.0 | 3.93E-02 |
| Q01177 | TPANFPDAGLEMNYCRNPDNDQR | 15,Carbamidomethyl[C]; | 7.6 | 0.0 | 0.0 | 3.94E-02 |
| P02770 | TCVADENAENCDK | 2,Carbamidomethyl[C]; | 4.6 | 0.0 | 0.0 | 3.95E-02 |
| P02770 | FATEGPNLVAR | 0,Galactosyl[AnyN-term]; | 4.6 | 0.0 | 0.0 | 3.95E-02 |
| Q03336 | CLLFVDIPSK | 1,Carbamidomethyl[C]; | 16.7 | 0.0 | 0.0 | 3.96E-02 |
| P02770 | KQTALAELVK | 3,Dehydrated[T]; | 5.3 | 0.0 | 0.0 | 3.98E-02 |
| Q01177 | YILQGVTSWGLGCARPNKPGVYVR | 13,Carbamidomethyl[C]; | 8.4 | 0.0 | 0.0 | 4.01E-02 |
| P02770 | FATEGPNLVAR | 0,NDA[AnyN-term]; | 2.7 | 0.0 | 0.0 | 4.01E-02 |
| P98158 | NPCASASCSHLCLLSAQAPR | 3,Carbamidomethyl[C];8,Carbamidomethyl[C];12,Carbamidomethyl[C]; | 9.3 | 0.0 | 0.0 | 4.06E-02 |
| P02770 | FQENPTSFLGHYLHEVAR | 0,DiART6plex118[AnyN-term]; | 3.0 | 0.0 | 0.0 | 4.08E-02 |
| P15083 | YWCHWEADENGR | 3,Carbamidomethyl[C]; | 3.0 | 0.0 | 0.0 | 4.08E-02 |
| P19468 | LGCPGFTLPEHRPNPEEGGASK | 3,Carbamidomethyl[C]; | 6.1 | 0.0 | 0.0 | 4.08E-02 |
| P02761 | LDVAKLNGDWFSIVVASNKR | 12,Hex(1)HexA(1)[S]; | 2.9 | 0.0 | 0.0 | 4.09E-02 |
| P02764 | IFAHLIVLK | 0,Formyl[AnyN-term]; | 5.4 | 0.0 | 0.0 | 4.09E-02 |
| P12788 | HPEYNKDTLDNDIMLIK | 6,TMT[K]; | 3.6 | 0.0 | 0.0 | 4.11E-02 |
| P02770 | TVMGDFAQFVDK | 3,Met->Lys[M]; | 5.9 | 0.0 | 0.0 | 4.12E-02 |
| P12346 | KSCHTGVDRTAGWNIPMGLLFSR | 3,Carbamidomethyl[C]; | 14.0 | 0.0 | 0.0 | 4.14E-02 |
| P20760 | MNGQPQENYK | 2,Deamidated[N]; | 7.4 | 0.0 | 0.0 | 4.17E-02 |
| P20759 | MNGQPQENYK | 2,Deamidated[N]; | 7.4 | 0.0 | 0.0 | 4.17E-02 |
| P02770 | TVMGDFAQFVDK | 1,Thr->Val[T]; | 9.6 | 0.0 | 0.0 | 4.17E-02 |
| P12346 | NGDGKEDLIWEILK | 0,C+12[AnyN-term]; | 25.9 | 0.0 | 0.0 | 4.20E-02 |
| P02770 | RPCFSALTVDETYVPK | 0,Formyl[AnyN-term];3,Carbamidomethyl[C]; | 66.9 | 0.0 | 0.0 | 4.21E-02 |
| P02770 | ECCHGDLLECADDRAELAK | 2,Carbamidomethyl[C];10,Carbamidomethyl[C]; | 40.4 | 0.0 | 0.0 | 4.22E-02 |
| P08932 | KTEEDLCVGCFQPIPMDSSDLKPVLK | 10,NIPCAM[C]; | 1.6 | 0.0 | 0.0 | 4.24E-02 |
| P01048 | KTEEDLCVGCFQPIPMDSSDLKPVLK | 10,NIPCAM[C]; | 1.6 | 0.0 | 0.0 | 4.24E-02 |
| P17475 | ILYFPKLSISGTYNLK | 6,Phosphoguanosine[K]; | 3.1 | 0.0 | 0.0 | 4.24E-02 |
| P08932 | EALGHSIAQLNAQHNHIFYFK | 0,Glu->pyro-Glu[AnyN-termE]; | 9.1 | 0.0 | 0.0 | 4.25E-02 |
| P01048 | EALGHSIAQLNAQHNHIFYFK | 0,Glu->pyro-Glu[AnyN-termE]; | 9.1 | 0.0 | 0.0 | 4.25E-02 |
| P98158 | GFTCACPDDFQTVQLR | 4,Carbamidomethyl[C];6,Carbamidomethyl[C]; | 4.3 | 0.0 | 0.0 | 4.27E-02 |
| P02770 | TSFLGHYLHEVAR | 1,Thr->Phe[T]; | 2.4 | 0.0 | 0.0 | 4.27E-02 |
| P05545 | IAELFSELDER | 0,C+12[AnyN-term]; | 9.0 | 0.0 | 0.0 | 4.28E-02 |
| P08932 | CQALDMMISRPPGFSPFR | 0,Propyl_2H(6)[AnyN-term]; | 2.3 | 0.0 | 0.0 | 4.28E-02 |
| P01048 | CQALDMMISRPPGFSPFR | 0,Propyl_2H(6)[AnyN-term]; | 2.3 | 0.0 | 0.0 | 4.28E-02 |
| P02770 | HPDYSVSLLLR | 1,Diethylphosphothione[H]; | 2.3 | 0.0 | 0.0 | 4.28E-02 |
| Q9WUC4 | PKHEFSVDMTCGGCAEAVSR | 11,Carbamidomethyl[C];14,Carbamidomethyl[C]; | 3.6 | 0.0 | 0.0 | 4.29E-02 |
| P02770 | LRDNYGELADCCAK | 0,Formyl[AnyN-term];11,Carbamidomethyl[C];12,Carbamidomethyl[C]; | 24.4 | 0.0 | 0.0 | 4.32E-02 |
| P02770 | LPCVEDYLSAILNRLCVLHEK | 3,Carbamidomethyl[C];16,Carbamidomethyl[C]; | 9.6 | 0.0 | 0.0 | 4.32E-02 |
| Q01177 | CTTPPPPPGPTYQCLK | 1,Carbamidomethyl[C];14,Carbamidomethyl[C]; | 16.0 | 0.0 | 0.0 | 4.32E-02 |
| P98158 | DGRTCKVTGSENPLLVVASR | 5,dichlorination[C]; | 2.7 | 0.0 | 0.0 | 4.32E-02 |
| P02770 | LYEYSRRHPDYSVSLLLR | 0,BITC[AnyN-term]; | 3.4 | 0.0 | 0.0 | 4.33E-02 |
| P02770 | TVMGDFAQFVDK | 1,Thr->Pro[T]; | 22.7 | 0.0 | 0.0 | 4.33E-02 |
| P02770 | DNCFATEGPNLVAR | 3,SulfoGMBS[C]; | 65.0 | 0.0 | 0.0 | 4.35E-02 |
| P02770 | PTSFLGHYLHEVAR | 1,OxProBiotin[P]; | 12.0 | 0.0 | 0.0 | 4.36E-02 |
| P02770 | YEYSRRHPDYSVSLLLR | 5,FTC[R]; | 2.9 | 0.0 | 0.0 | 4.37E-02 |
| P02770 | KYEATLEK | 0,C+12[AnyN-term]; | 19.4 | 0.0 | 0.0 | 4.38E-02 |
| P02770 | TFHSDICTLPDKEK | 7,Cys->Dha[C]; | 4.0 | 0.0 | 0.0 | 4.40E-02 |
| P02770 | TSFLGHYLHEVAR | 4,Xle->Thr[L]; | 11.3 | 0.0 | 0.0 | 4.43E-02 |
| P02770 | NCFATEGPNLVAR | 2,Carbamidomethyl[C]; | 29.7 | 0.0 | 0.0 | 4.43E-02 |
| P02770 | FATEGPNLVAR | 1,Bromo[F]; | 6.1 | 0.0 | 0.0 | 4.45E-02 |
| P02770 | TVMGDFAQFVDK | 3,Carbamyl[M]; | 13.7 | 0.0 | 0.0 | 4.46E-02 |
| P07522 | VNLDPASVPPR | 0,C+12[AnyN-term]; | 8.1 | 0.0 | 0.0 | 4.46E-02 |
| P12346 | GWNIPMGLLFSR | 0,3sulfo[AnyN-term]; | 3.1 | 0.0 | 0.0 | 4.47E-02 |
| P01836 | RDGVLDSVTDQDSK | 12,Dehydrated[D]; | 12.6 | 0.0 | 0.0 | 4.49E-02 |
| P01835 | RDGVLDSVTDQDSK | 12,Dehydrated[D]; | 12.6 | 0.0 | 0.0 | 4.49E-02 |
| P14046 | ETGLMAFTNLK | 1,Cation_Li[E]; | 10.1 | 0.0 | 0.0 | 4.49E-02 |
| P02770 | DKDNCFATEGPNLVAR | 5,HN2_mustard[C]; | 2.6 | 0.0 | 0.0 | 4.53E-02 |
| P02770 | GEYGFQNAVLVR | 3,Tyr->Cys[Y]; | 2.6 | 0.0 | 0.0 | 4.53E-02 |
| P02770 | AELVKHKPKATEDQLKTVMGDFAQFVDK | 0,TNBS[AnyN-term]; | 2.6 | 0.0 | 0.0 | 4.53E-02 |
| P08932 | CQALDMMISRPPGFSPFR | 1,Trioxidation[C]; | 2.6 | 0.0 | 0.0 | 4.53E-02 |
| P01048 | CQALDMMISRPPGFSPFR | 1,Trioxidation[C]; | 2.6 | 0.0 | 0.0 | 4.53E-02 |
| P12346 | DGGGDVAFVK | 1,Dehydrated[D]; | 2.6 | 0.0 | 0.0 | 4.53E-02 |
| Q01177 | NLEENYCR | 7,Carbamidomethyl[C]; | 2.6 | 0.0 | 0.0 | 4.53E-02 |
| Q6IRK9 | QQDGLENVHLEQVR | 0,3sulfo[AnyN-term]; | 2.6 | 0.0 | 0.0 | 4.53E-02 |
| P05544 | ATAATGVATVIR | 1,Dap-DSP[A]; | 4.4 | 0.0 | 0.0 | 4.53E-02 |
| P09005 | ATAATGVATVIR | 1,Dap-DSP[A]; | 4.4 | 0.0 | 0.0 | 4.53E-02 |
| P02770 | TVMGDFAQFVDK | 5,Cation_Ca[II][D]; | 41.3 | 0.0 | 0.0 | 4.53E-02 |
| P02770 | EYSRRHPDYSVSLLLR | 4,Arg->Cys[R]; | 1.9 | 0.0 | 0.0 | 4.54E-02 |
| P20760 | KETWQQGNTFTCSVLHEGLHNHHTEK | 12,Carbamidomethyl[C];21,Deamidated[N]; | 1.9 | 0.0 | 0.0 | 4.54E-02 |
| P17475 | KISSNLADFAFSLYR | 5,Deamidated[N]; | 1.9 | 0.0 | 0.0 | 4.54E-02 |
| P09005 | LDVAETGTEATAATGVATVIR | 5,GlycerylPE[E]; | 1.9 | 0.0 | 0.0 | 4.54E-02 |
| P05371 | LTQQYNELLHSLQSK | 0,ISD_z+2_ion[AnyN-term]; | 1.9 | 0.0 | 0.0 | 4.54E-02 |
| P02770 | ADKDNCFATEGPNLVAR | 0,Formyl[AnyN-term]; | 3.9 | 0.0 | 0.0 | 4.56E-02 |
| P02770 | HKPKATEDQLKTVMGDFAQFVDK | 0,Propionamide[AnyN-term]; | 7.3 | 0.0 | 0.0 | 4.58E-02 |
| P02770 | AADKDNCFATEGPNLVAR | 7,Pyridylethyl[C]; | 1.4 | 0.0 | 0.0 | 4.65E-02 |
| P02770 | PTSFLGHYLHEVAR | 2,Oxidation[T]; | 1.4 | 0.0 | 0.0 | 4.65E-02 |
| P02770 | MCTSFQENPTSFLGHYLHEVAR | 0,Phenylisocyanate_2H(5)[AnyN-term]; | 4.3 | 0.0 | 0.0 | 4.65E-02 |
| P02770 | NPTSFLGHYLHEVAR | 3,Cresylphosphate[T]; | 1.4 | 0.0 | 0.0 | 4.65E-02 |
| P02770 | PYFYAPELLYYAEK | 3,Phe->Gly[F]; | 1.4 | 0.0 | 0.0 | 4.65E-02 |
| P02770 | AADKDNCFATEGPNLVAR | 4,AFB1_Dialdehyde[K]; | 1.4 | 0.0 | 0.0 | 4.65E-02 |
| P02770 | EKALVAAVR | 0,C+12[AnyN-term]; | 1.4 | 0.0 | 0.0 | 4.65E-02 |
| P05544 | AVLDVDETGTEATAATGVATVIR | 6,Dehydrated[D]; | 1.4 | 0.0 | 0.0 | 4.65E-02 |
| P05544 | STMEEILEGLK | 0,Phenylisocyanate_2H(5)[AnyN-term]; | 3.6 | 0.0 | 0.0 | 4.65E-02 |
| P81827 | CESLDSTGLCR | 1,GG[C](Dicarbamidomethyl[C]);10,Carbamidomethyl[C]; | 1.4 | 0.0 | 0.0 | 4.65E-02 |
| Q9JJ19 | IVEVNGVCMEGK | 8,Carbamidomethyl[C]; | 1.4 | 0.0 | 0.0 | 4.65E-02 |
| P02780 | PICCYASGSGCSILDEVIR | 3,Carbamidomethyl[C];11,Carbamidomethyl[C]; | 2.4 | 0.0 | 0.0 | 4.67E-02 |
| Q9JHB9 | PICCYASGSGCSILDEVIR | 3,Carbamidomethyl[C];11,Carbamidomethyl[C]; | 2.4 | 0.0 | 0.0 | 4.67E-02 |
| P05544 | TEATAATGVATVIR | 1,Pentose[T]; | 10.4 | 0.0 | 0.0 | 4.68E-02 |
| P09005 | TEATAATGVATVIR | 1,Pentose[T]; | 10.4 | 0.0 | 0.0 | 4.68E-02 |
| P02770 | TVMGDFAQFVDK | 3,Met->Gln[M]; | 6.4 | 0.0 | 0.0 | 4.71E-02 |
| P02770 | FQENPTSFLGHYLHEVAR | 0,NDA[AnyN-term]; | 5.3 | 0.0 | 0.0 | 4.72E-02 |
| P36373 | KPGDDHSNDLMLLHLSQPADITDGVK | 23,Dehydrated[D]; | 1.1 | 0.0 | 0.0 | 4.72E-02 |
| P02770 | CFSALTVDETYVPK | 0,ESP[AnyN-term]; | 11.9 | 0.0 | 0.0 | 4.77E-02 |
| P02770 | LQACCDKPVLQK | 4,Carbamidomethyl[C]; | 8.4 | 0.0 | 0.0 | 4.79E-02 |
| P01836 | SKDSTYSMSSTLSLTK | 1,Glucuronyl[S]; | 7.0 | 0.0 | 0.0 | 4.81E-02 |
| P01835 | SKDSTYSMSSTLSLTK | 1,Glucuronyl[S]; | 7.0 | 0.0 | 0.0 | 4.81E-02 |
| P02770 | RHPDYSVSLLLR | 2,His->Glu[H]; | 3.0 | 0.0 | 0.0 | 4.85E-02 |
| P02761 | GNLDVAKLNGDWFSIVVASNKR | 7,3-phosphoglyceryl[K]; | 13.1 | 0.0 | 0.0 | 4.86E-02 |
| P02770 | ENPTSFLGHYLHEVAR | 1,phosphoRibosyl[E]; | 5.9 | 0.0 | 0.0 | 4.87E-02 |
| P83121 | EICAWVVVTTR | 3,biotinAcrolein298[C]; | 98.3 | 0.0 | 0.0 | 4.87E-02 |
| P81827 | EICAWVVVTTR | 3,biotinAcrolein298[C]; | 98.3 | 0.0 | 0.0 | 4.87E-02 |
| P81828 | EICAWVVVTTR | 3,biotinAcrolein298[C]; | 98.3 | 0.0 | 0.0 | 4.87E-02 |
| P02770 | AADKDNCFATEGPNLVAR | 0,C+12[AnyN-term]; | 1.3 | 0.0 | 0.0 | 4.88E-02 |
| P02770 | QLKTVMGDFAQFVDK | 4,Pent(1)HexNAc(1)[T]; | 1.3 | 0.0 | 0.0 | 4.88E-02 |
| P02770 | LPCVEDYLSAILNR | 1,Xle->Asp[L]; | 1.3 | 0.0 | 0.0 | 4.88E-02 |
| P02770 | TVMGDFAQFVDKCCK | 0,Formyl[AnyN-term];3,Oxidation[M];13,GG[C](Dicarbamidomethyl[C]); | 1.3 | 0.0 | 0.0 | 4.88E-02 |
| P02761 | VFMQHIDVLENSLGFK | 12,Formyl[S](Ser->Asp[S]); | 3.9 | 0.0 | 0.0 | 4.88E-02 |
| P12346 | AVVARNGDGKEDLIWEILK | 10,Glucosylgalactosyl[K]; | 1.3 | 0.0 | 0.0 | 4.88E-02 |
| P12346 | LPEGTTYEEYLGAEYLQAVGNIR | 8,Cation_Ca[II][E]; | 1.3 | 0.0 | 0.0 | 4.88E-02 |
| P05545 | NLHVSQVVHK | 0,C+12[AnyN-term]; | 1.3 | 0.0 | 0.0 | 4.88E-02 |
| P02782 | QICELVAHETISFLMK | 3,4-ONE+Delta_H(-2)O(-1)[C]; | 1.3 | 0.0 | 0.0 | 4.88E-02 |
| P81828 | RCTSFDSTGFCHVGR | 2,Carbamidomethyl[C];11,Carbamidomethyl[C]; | 2.6 | 0.0 | 0.0 | 4.88E-02 |
| P07314 | QPLSSMCPSIIVDKDGK | 0,Gln->pyro-Glu[AnyN-termQ];6,Oxidation[M];7,Carbamidomethyl[C]; | 1.3 | 0.0 | 0.0 | 4.88E-02 |
| Q64319 | THAVSLEK | 1,Hex(1)NeuGc(1)[T]; | 1.3 | 0.0 | 0.0 | 4.88E-02 |
| P50123 | GCQIYLENFK | 2,Carbamidomethyl[C]; | 9.3 | 0.0 | 0.0 | 4.93E-02 |
| P01836 | DGVLDSVTDQDSKDSTYSMSSTLSLTK | 7,Val->Ser[V]; | 6.1 | 0.0 | 0.0 | 4.94E-02 |
| P01835 | DGVLDSVTDQDSKDSTYSMSSTLSLTK | 7,Val->Ser[V]; | 6.1 | 0.0 | 0.0 | 4.94E-02 |
| P12346 | EGVCPEGSIDSAPVK | 5,Pro->Trp[P]; | 5.4 | 0.0 | 0.0 | 4.96E-02 |
| P13635 | DIASGLIGPLILCKK | 13,Carbamidomethyl[C]; | 5.4 | 0.0 | 0.0 | 4.96E-02 |
| P27590 | GYRGDGWYCECSPGFCEPGLDCLPQGPSGK | 11,GG[C](Dicarbamidomethyl[C]);16,Carbamidomethyl[C];22,Carbamidomethyl[C]; | 2.7 | 0.0 | 0.0 | 4.96E-02 |
| P02770 | CFATEGPNLVAR | 1,Dioxidation[C]; | 61.9 | 0.0 | 0.0 | 4.97E-02 |
| P02770 | TNCELYEK | 0,Formyl[AnyN-term];3,Carbamidomethyl[C]; | 3.3 | 0.0 | 0.0 | 4.97E-02 |
| P02770 | PPFQRPEAEAMCTSFQENPTSFLGHYLHEVAR | 11,Met->Phe[M]; | 4.0 | 0.0 | 0.0 | 4.98E-02 |
| P02770 | KDNCFATEGPNLVAR | 1,HCysThiolactone[K]; | 2.0 | 0.0 | 0.0 | 4.98E-02 |
| P02761 | DVAKLNGDWFSIVVASNKR | 11,Hex(1)NeuAc(1)[S]; | 2.0 | 0.0 | 0.0 | 4.98E-02 |
| P17475 | KVINDYVEK | 0,Formyl[AnyN-term]; | 2.0 | 0.0 | 0.0 | 4.98E-02 |
| P05544 | DLYVSQVVHKAVLDVDETGTEATAATGVATVIR | 10,AFB1_Dialdehyde[K]; | 2.0 | 0.0 | 0.0 | 4.98E-02 |
| P05544 | LEERTSMVLVNYLLFK | 0,3sulfo[AnyN-term]; | 48.3 | 0.2 | 0.0 | 1.59E-02 |
| P31211 | DVPLTLTMVHK | 8,Oxidation[M]; | 17.4 | 0.2 | 0.0 | 3.97E-02 |
| P13635 | IYHSHVDAPKDIASGLIGPLILCK | 23,Carbamidomethyl[C]; | 8.3 | 0.2 | 0.0 | 4.95E-02 |
| P98158 | CACDPEYTLESDGR | 1,Carbamidomethyl[C];3,Carbamidomethyl[C]; | 14.7 | 0.3 | 0.0 | 2.23E-02 |
| P02770 | CCTLPEAQRLPCVEDYLSAILNR | 1,Carbamidomethyl[C];2,Carbamidomethyl[C];12,Carbamidomethyl[C]; | 190.4 | 6.0 | 0.0 | 4.22E-02 |
| P02770 | YMCENQATISSK | 3,Trioxidation[C]; | 28.9 | 1.0 | 0.0 | 3.93E-04 |
| P01048 | ELESGNQFVLYR | 0,ESP[AnyN-term]; | 18.9 | 0.8 | 0.0 | 4.11E-02 |
| P81828 | STGFCHVGR | 5,Carbamidomethyl[C]; | 33.0 | 1.7 | 0.1 | 3.01E-02 |
| P12346 | DLFWKGDKDCTGNFCLFR | 10,Carbamidomethyl[C];13,Deamidated[N];15,Carbamidomethyl[C]; | 13.1 | 1.0 | 0.1 | 4.95E-02 |
| P02770 | RHPDYSVSLLLR | 0,Formyl[AnyN-term]; | 282.7 | 22.2 | 0.1 | 2.95E-04 |
| P02770 | EFKAETFTFHSDICTLPDKEK | 14,Carbamidomethyl[C]; | 743.1 | 70.2 | 0.1 | 2.47E-02 |
| P02770 | CENQATISSK | 1,Carbamidomethyl[C]; | 40.0 | 4.0 | 0.1 | 5.70E-06 |
| P02770 | INKECCHGDLLECADDRAELAK | 5,Carbamidomethyl[C];6,Carbamidomethyl[C];13,Carbamidomethyl[C]; | 740.9 | 84.8 | 0.1 | 1.70E-02 |
| P02770 | AELAKYMCENQATISSK | 0,Unknown_248[AnyN-term]; | 2.4 | 0.3 | 0.1 | 3.68E-02 |
| P10960 | LGPGVSDICK | 9,Carbamidomethyl[C]; | 9.6 | 1.3 | 0.1 | 3.49E-02 |
| P27590 | CQLQSLGFMK | 0,Pyro-carbamidomethyl[AnyN-termC]; | 85.6 | 13.7 | 0.2 | 1.73E-02 |
| P02770 | FKDLGEQHFK | 3,Dehydrated[D]; | 8.0 | 1.5 | 0.2 | 2.46E-02 |
| P02761 | LNGDWFSIVVASNKR | 2,Deamidated[N]; | 313.4 | 63.5 | 0.2 | 3.11E-02 |
| P05545 | DEELSCSVLELK | 6,Carbamidomethyl[C]; | 74.1 | 18.2 | 0.2 | 5.24E-04 |
| P05544 | DEELSCSVLELK | 6,Carbamidomethyl[C]; | 74.1 | 18.2 | 0.2 | 5.24E-04 |
| P09005 | DEELSCSVLELK | 6,Carbamidomethyl[C]; | 74.1 | 18.2 | 0.2 | 5.24E-04 |
| Q63556 | DEELSCSVLELK | 6,Carbamidomethyl[C]; | 74.1 | 18.2 | 0.2 | 5.24E-04 |
| P02770 | AETFTFHSDICTLPDKEK | 11,Carbamidomethyl[C]; | 1719.0 | 451.7 | 0.3 | 5.86E-05 |
| P20059 | YYCFQGNK | 3,Carbamidomethyl[C]; | 14.9 | 22.8 | 1.5 | 2.72E-02 |
| P02770 | LPCVEDYLSAILNR | 3,Crotonaldehyde[C]; | 32.7 | 57.3 | 1.8 | 2.25E-02 |
| P20760 | ETWQQGNTFTCSVLHEGLHNHHTEK | 11,Carbamidomethyl[C]; | 49.7 | 91.5 | 1.8 | 5.00E-02 |
| P32038 | ILGGQEAMAHAR | 8,Oxidation[M]; | 2.9 | 5.8 | 2.0 | 1.80E-02 |
| P02764 | QQLELEKETK | 0,Gln->pyro-Glu[AnyN-termQ]; | 15.4 | 31.7 | 2.1 | 1.59E-03 |
| P13635 | DCNKPSPDDDIQDR | 2,Carbamidomethyl[C]; | 7.1 | 16.5 | 2.3 | 9.57E-05 |
| P14841 | SQTNLTNCPFHDQPHLMR | 8,Carbamidomethyl[C]; | 6.3 | 14.8 | 2.4 | 1.52E-02 |
| P20059 | GATYAFSGSHYWR | 3,Dehydrated[T]; | 2.7 | 6.7 | 2.5 | 1.47E-02 |
| P02770 | QEPERNECFLQHK | 6,Deamidated[N];8,Carbamidomethyl[C]; | 21.9 | 55.7 | 2.5 | 5.81E-03 |
| P02770 | NPNLPPFQRPEAEAMCTSFQENPTSFLGHYLHEVAR | 16,Carbamidomethyl[C]; | 6.6 | 18.0 | 2.7 | 2.95E-03 |
| P12346 | INHCKFDEFFSQGCAPGYK | 4,Carbamidomethyl[C];14,Carbamidomethyl[C]; | 16.4 | 46.0 | 2.8 | 3.17E-03 |
| Q99PS8 | HPPVFGLCR | 8,Carbamidomethyl[C]; | 4.1 | 11.8 | 2.9 | 4.51E-03 |
| Q01177 | NPDGDVNGPWCYTMNPR | 11,Carbamidomethyl[C]; | 6.9 | 20.5 | 3.0 | 1.69E-05 |
| Q01177 | TPENFPCK | 7,Carbamidomethyl[C]; | 9.6 | 33.5 | 3.5 | 2.16E-04 |
| P36953 | VYKDLLEDCCNRK | 9,Carbamidomethyl[C];10,Carbamidomethyl[C]; | 2.4 | 8.8 | 3.6 | 3.28E-05 |
| P20760 | ETWQQGNTFTCSVLHEGLHNHHTEK | 7,Deamidated[N];11,Carbamidomethyl[C]; | 1.0 | 3.7 | 3.7 | 4.85E-02 |
| P02770 | LPCVEDYLSAILNR | 3,CarbamidomethylDTT[C]; | 3.4 | 13.0 | 3.8 | 1.45E-04 |
| P02770 | FKDLGEQHFK | 0,Carbamyl[AnyN-term]; | 6.6 | 25.7 | 3.9 | 3.38E-04 |
| P02770 | LPCVEDYLSAILNR | 3,monomethylphosphothione[C]; | 5.0 | 19.8 | 4.0 | 4.99E-03 |
| P02770 | LVQEVTDFAK | 7,Cation_Na[D]; | 15.7 | 63.7 | 4.1 | 7.22E-05 |
| P02770 | ENPTSFLGHYLHEVAR | 0,Phenylisocyanate_2H(5)[AnyN-term]; | 1.6 | 6.5 | 4.1 | 4.10E-03 |
| P01048 | EIPVDSPELK | 0,C+12[AnyN-term]; | 3.0 | 12.7 | 4.2 | 5.03E-03 |
| P20760 | ETWQQGNTFTCSVLHEGLHNHHTEK | 11,Carbamidomethyl[C];20,Deamidated[N]; | 6.1 | 31.7 | 5.2 | 1.95E-03 |
| P24090 | QQAEHAVEGDCDFHILK | 0,Gln->pyro-Glu[AnyN-termQ];11,Carbamidomethyl[C]; | 3.0 | 15.7 | 5.2 | 4.83E-04 |
| P12346 | CALSHQER | 0,Biotin[AnyN-term]; | 0.9 | 4.5 | 5.3 | 1.79E-03 |
| P12346 | TSYQDCIK | 6,Carbamidomethyl[C]; | 5.6 | 30.3 | 5.4 | 8.25E-03 |
| P02680 | DNCCILDER | 3,Carbamidomethyl[C];4,Carbamidomethyl[C]; | 2.7 | 15.5 | 5.7 | 6.81E-03 |
| P20059 | DYFISCPGR | 6,Carbamidomethyl[C]; | 18.1 | 106.2 | 5.9 | 1.09E-03 |
| P02761 | LCEAHGITR | 2,CarbamidomethylDTT[C]; | 2.1 | 12.8 | 6.0 | 2.00E-03 |
| P06866 | PICLPSKDYVAPGR | 3,Carbamidomethyl[C]; | 4.4 | 29.5 | 6.7 | 1.74E-02 |
| P02770 | RHPDYSVSLLLR | 8,Formyl[S](Ser->Asp[S]); | 9.0 | 61.3 | 6.8 | 1.48E-05 |
| P02770 | HSDICTLPDKEK | 5,Carbamidomethyl[C]; | 0.9 | 6.2 | 7.2 | 1.19E-03 |
| P09006 | DSLRPSMIDELYLPK | 7,Oxidation[M]; | 6.0 | 44.2 | 7.4 | 1.14E-05 |
| P08932 | CQALDMMISRPPGFSPFR | 2,Gln->Tyr[Q]; | 0.4 | 3.2 | 7.4 | 4.77E-02 |
| P01048 | CQALDMMISRPPGFSPFR | 2,Gln->Tyr[Q]; | 0.4 | 3.2 | 7.4 | 4.77E-02 |
| Q01177 | TAVTAAGTPCQEWAAQEPHSHR | 10,Acetyl[C]; | 0.7 | 5.7 | 7.9 | 3.67E-02 |
| P24090 | ELACDDPETEHVALIAVDYLNK | 4,Carbamidomethyl[C]; | 4.9 | 39.0 | 8.0 | 8.23E-04 |
| P02770 | TSFLGHYLHEVAR | 1,dHex[T]; | 0.7 | 5.8 | 8.2 | 4.17E-02 |
| P01026 | LEESCTRFYHPEKDDGMLSK | 7,Arg->Met[R]; | 0.4 | 3.7 | 8.6 | 2.03E-02 |
| P01026 | VELLHNPAFCSMATAK | 10,Carbamidomethyl[C]; | 3.3 | 28.5 | 8.7 | 1.07E-06 |
| Q01177 | CQSWVSMTPHSHSK | 1,Carbamidomethyl[C]; | 3.0 | 26.5 | 8.8 | 4.81E-05 |
| P02770 | DDNPNLPPFQRPEAEAMCTSFQENPTSFLGHYLHEVAR | 18,Carbamidomethyl[C]; | 39.0 | 364.2 | 9.3 | 1.31E-09 |
| P02770 | RHPYFYAPELLYYAEK | 15,Cation_Ca[II][E]; | 0.6 | 5.7 | 9.9 | 8.14E-03 |
| P02770 | LPEAQRLPCVEDYLSAILNR | 0,SPITC[AnyN-term]; | 0.4 | 4.5 | 10.5 | 4.98E-02 |
| P13635 | MYYSGVDLTK | 1,Oxidation[M]; | 1.0 | 10.7 | 10.7 | 3.25E-03 |
| P02770 | LRDNYGELADCCAK | 11,Diethylphosphate[C]; | 0.7 | 7.7 | 10.7 | 9.58E-04 |
| P02770 | AKYMCENQATISSK | 5,DiDehydro[C]; | 0.4 | 4.7 | 10.9 | 1.81E-02 |
| P02770 | ENPTSFLGHYLHEVAR | 1,Glu->His[E]; | 3.3 | 39.2 | 11.9 | 1.04E-03 |
| P17475 | GTEAAGATVVEAVPMSLPPQVK | 2,Dehydrated[T]; | 0.9 | 10.3 | 12.1 | 7.96E-03 |
| P02680 | IHDTTGKDCQDIANK | 9,Carbamidomethyl[C]; | 0.3 | 3.5 | 12.3 | 2.15E-02 |
| P02770 | CFATEGPNLVAR | 1,NEM_2H(5)+H2O[C]; | 1.4 | 19.0 | 13.3 | 1.07E-05 |
| P02770 | DKDNCFATEGPNLVAR | 5,NEIAA_2H(5)[C]; | 0.6 | 8.2 | 14.3 | 3.08E-02 |
| P36953 | ADWCQPLK | 4,Carbamidomethyl[C]; | 0.9 | 13.7 | 15.9 | 1.90E-04 |
| P02770 | PTSFLGHYLHEVAR | 0,C+12[AnyN-term]; | 0.6 | 9.3 | 16.3 | 1.29E-02 |
| P02770 | TCVADENAENCDKSIHTLFGDK | 2,Pyridylethyl[C]; | 0.1 | 2.3 | 16.3 | 3.39E-02 |
| P05371 | TQQYNELLHSLQSK | 0,Succinyl_2H(4)[AnyN-term]; | 0.3 | 5.3 | 18.7 | 3.36E-02 |
| P20059 | NPLPAAHETVAK | 1,Ammonia-loss[N]; | 0.1 | 2.8 | 19.8 | 1.76E-03 |
| P02770 | QKCPYEEHIK | 3,Carbamidomethyl[C]; | 0.1 | 2.8 | 19.8 | 5.71E-03 |
| P02782 | ICELVAHETISFLMK | 2,Carbamidomethyl[C]; | 0.1 | 2.8 | 19.8 | 4.02E-02 |
| P36374 | VGSTCLTSGWGSITPLK | 5,Carbamidomethyl[C]; | 0.7 | 15.5 | 21.7 | 2.49E-02 |
| P12346 | TAGWNIPMGLLFSR | 0,C+12[AnyN-term]; | 1.0 | 24.5 | 24.5 | 2.21E-04 |
| P02770 | PTSFLGHYLHEVAR | 0,Methyl[AnyN-term]; | 1.4 | 38.2 | 26.7 | 9.20E-06 |
| P09006 | MQQVEASLQPETLRR | 1,Met->His[M]; | 0.6 | 16.7 | 29.2 | 1.37E-04 |
| P02770 | QENPTSFLGHYLHEVAR | 0,TMAB_2H(9)[AnyN-term]; | 0.1 | 4.5 | 31.5 | 1.77E-02 |
| P02770 | CPYEEHIK | 1,CarbamidomethylDTT[C]; | 0.6 | 19.7 | 34.4 | 1.40E-05 |
| P12346 | QAPNHVVVSR | 0,Cytopiloyne+water[AnyN-term]; | 0.1 | 9.7 | 67.7 | 4.07E-03 |
| P17475 | EDAQETDTSQQDQSPTYR | 7,Cation_Ca[II][D]; | 0.1 | 13.0 | 91.0 | 2.28E-04 |
| P12346 | HTTIFEVLPQK | 6,Cation_Na[E]; | 0.1 | 13.2 | 92.2 | 1.33E-02 |
| P20760 | MNGQPQENYK | 1,Oxidation[M];2,Deamidated[N]; | 0.3 | 31.5 | 110.3 | 4.99E-07 |
| P20759 | MNGQPQENYK | 1,Oxidation[M];2,Deamidated[N]; | 0.3 | 31.5 | 110.3 | 4.99E-07 |
| Q03626 | DMGLTAFTNLK | 2,Oxidation[M]; | 0.1 | 16.3 | 114.3 | 7.99E-03 |
| Q6IE52 | DMGLTAFTNLK | 2,Oxidation[M]; | 0.1 | 16.3 | 114.3 | 7.99E-03 |
| Q9QX79 | KIHSMCPDCPHPVDLSAPSVLEAATESLAK | 5,Oxidation[M];6,Carbamidomethyl[C];9,Carbamidomethyl[C]; | 0.6 | 81.0 | 141.8 | 4.27E-04 |
| P02770 | YTQKAPQVSTPTLVEAAR | 0,Galactosyl[AnyN-term]; | 0.1 | 20.5 | 143.5 | 3.16E-03 |
| P17475 | VEAVPMSLPPQVK | 0,Formyl[AnyN-term]; | 0.1 | 20.8 | 145.8 | 4.93E-03 |
| P02770 | DNCFATEGPNLVAR | 2,Hex(3)[N]; | 0.1 | 75.0 | 525.0 | 3.31E-07 |
| Q9EQV6 | VFNNDADLSGITEDAPLK | 7,Cation_Na[D]; | 0.0 | 35.2 | #DIV/0! | 1.66E-08 |
| Q9EQV6 | EFKAETFTFHSDICTLPDK | 14,Carbamidomethyl[C]; | 0.0 | 23.0 | #DIV/0! | 5.25E-08 |
| Q9EQV6 | PTSFLGHYLHEVAR | 0,CLIP_TRAQ_2[AnyN-term]; | 0.0 | 32.3 | #DIV/0! | 7.04E-08 |
| P06238 | EMGPTYADPVCLSK | 11,Carbamidomethyl[C]; | 0.0 | 42.5 | #DIV/0! | 1.79E-07 |
| Q9EQV6 | DFQLFGSPLGK | 1,Cation_Na[D]; | 0.0 | 16.0 | #DIV/0! | 3.21E-07 |
| Q9EQV6 | AADKDNCFATEGPNLVAR | 7,Carbamidomethyl[C]; | 0.0 | 420.0 | #DIV/0! | 4.75E-07 |
| Q9EQV6 | EEMTQSQVSITCMVK | 12,Carbamidomethyl[C]; | 0.0 | 28.0 | #DIV/0! | 6.24E-07 |
| Q9EQV6 | QTALAELVK | 0,Gln->pyro-Glu[AnyN-termQ]; | 0.0 | 38.5 | #DIV/0! | 7.75E-07 |
| Q9EQV6 | MNGQPQENYK | 1,Oxidation[M]; | 0.0 | 27.2 | #DIV/0! | 9.58E-07 |
| P06238 | MNGQPQENYK | 1,Oxidation[M]; | 0.0 | 27.2 | #DIV/0! | 9.58E-07 |
| Q9EQV6 | FPSSTFEQVSQLVK | 2,Pro->Cys[P]; | 0.0 | 17.0 | #DIV/0! | 1.33E-06 |
| Q9EQV6 | KQTALAELVKHKPK | 10,Carbamyl[K]; | 0.0 | 10.7 | #DIV/0! | 1.80E-06 |
| Q9EQV6 | VFNNDADLSGITEDAPLK | 0,Dimethyl_2H(4)[AnyN-term]; | 0.0 | 65.0 | #DIV/0! | 2.03E-06 |
| Q9EQV6 | RHPDYSVSLLLR | 5,Dehydrated[Y]; | 0.0 | 235.0 | #DIV/0! | 2.72E-06 |
| Q9EQV6 | FPNAEFAEITK | 0,Formyl[AnyN-term]; | 0.0 | 18.7 | #DIV/0! | 2.74E-06 |
| Q9EQV6 | YMCENQATISSK | 3,Cys->Arg[C]; | 0.0 | 91.7 | #DIV/0! | 4.00E-06 |
| Q9EQV6 | MSQRFPNAEFAEITK | 0,Phenylisocyanate_2H(5)[AnyN-term]; | 0.0 | 315.8 | #DIV/0! | 4.17E-06 |
| Q9EQV6 | TCHLAQAPNHVVVSR | 2,CarbamidomethylDTT[C]; | 0.0 | 38.7 | #DIV/0! | 4.28E-06 |
| Q9EQV6 | RMSQRFPNAEFAEITK | 4,Gln->Pro[Q]; | 0.0 | 34.0 | #DIV/0! | 5.99E-06 |
| Q9EQV6 | RHPDYSVSLLLR | 1,Arg->Tyr[R]; | 0.0 | 73.3 | #DIV/0! | 6.06E-06 |
| P02767 | TAVTAAGTPCQEWAAQEPHSHR | 10,Carbamidomethyl[C]; | 0.0 | 44.3 | #DIV/0! | 6.16E-06 |
| Q9EQV6 | QKAPQVSTPTLVEAAR | 2,Phosphoguanosine[K]; | 0.0 | 44.5 | #DIV/0! | 6.68E-06 |
| P02767 | LVAYYTLIGANGQR | 11,Deamidated[N]; | 0.0 | 37.8 | #DIV/0! | 8.72E-06 |
| Q9EQV6 | NTPPTMDTDGSYFLYSK | 6,Oxidation[M]; | 0.0 | 207.8 | #DIV/0! | 9.84E-06 |
| P06238 | NTPPTMDTDGSYFLYSK | 6,Oxidation[M]; | 0.0 | 207.8 | #DIV/0! | 9.84E-06 |
| Q9EQV6 | EVVSLTEECCAEGADPNCYDTR | 9,Carbamidomethyl[C];10,Carbamidomethyl[C];18,Carbamidomethyl[C]; | 0.0 | 50.8 | #DIV/0! | 1.14E-05 |
| Q9EQV6 | ENPTSFLGHYLHEVAR | 2,Asn->His[N]; | 0.0 | 21.5 | #DIV/0! | 1.26E-05 |
| Q9EQV6 | ELYLVAYK | 1,Cation_Na[E]; | 0.0 | 5.7 | #DIV/0! | 1.29E-05 |
| P06238 | SQSQVMHHICSK | 10,Carbamidomethyl[C]; | 0.0 | 2.8 | #DIV/0! | 1.29E-05 |
| Q9EQV6 | ATCHLAQAPNHVVVSR | 2,CresylSaligeninPhosphate[T]; | 0.0 | 17.3 | #DIV/0! | 1.30E-05 |
| Q9EQV6 | LQACCDKPVLQK | 4,Carbamidomethyl[C];5,Carbamidomethyl[C]; | 0.0 | 220.8 | #DIV/0! | 1.36E-05 |
| P06238 | KALYSEYTDGTFTK | 0,Methyl_2H(2)[AnyN-term]; | 0.0 | 5.2 | #DIV/0! | 1.36E-05 |
| Q9WUW3 | TIAACNLPIVQGPCR | 5,Carbamidomethyl[C];14,Carbamidomethyl[C]; | 0.0 | 29.2 | #DIV/0! | 1.38E-05 |
| P22283 | EFQTTDDQCVYNFTHLGVQR | 9,Carbamidomethyl[C]; | 0.0 | 26.3 | #DIV/0! | 1.44E-05 |
| Q9EQV6 | AADKDNCFATEGPNLVAR | 6,Deamidated[N];7,Carbamidomethyl[C]; | 0.0 | 75.5 | #DIV/0! | 1.48E-05 |
| Q9EQV6 | DQYELLCLDNTR | 7,Carbamidomethyl[C]; | 0.0 | 86.2 | #DIV/0! | 1.71E-05 |
| Q9EQV6 | CTSFQENPTSFLGHYLHEVAR | 1,Carbamidomethyl[C]; | 0.0 | 30.3 | #DIV/0! | 1.72E-05 |
| Q9EQV6 | VFNNDADLSGITEDAPLK | 9,Formyl[S](Ser->Asp[S]); | 0.0 | 11.8 | #DIV/0! | 1.85E-05 |
| P02767 | ELGSPPGISLDTIDAAFSCPGSSK | 19,Carbamidomethyl[C]; | 0.0 | 36.2 | #DIV/0! | 2.27E-05 |
| Q9EQV6 | WCAVSEHENTK | 2,Carbamidomethyl[C];9,Deamidated[N]; | 0.0 | 8.5 | #DIV/0! | 2.30E-05 |
| P02767 | GPDSVFLIK | 0,Methyl[AnyN-term]; | 0.0 | 29.8 | #DIV/0! | 2.39E-05 |
| P06238 | IANDAIQDMLCDMK | 11,Carbamidomethyl[C]; | 0.0 | 25.0 | #DIV/0! | 3.37E-05 |
| P02767 | AVLDVAETGTEAAAATGVK | 7,Cation_Na[E]; | 0.0 | 38.0 | #DIV/0! | 3.38E-05 |
| Q9EQV6 | YNEVLTQCCTESDKAACLTPK | 2,Deamidated[N];8,GG[C](Dicarbamidomethyl[C]);17,Carbamidomethyl[C]; | 0.0 | 17.2 | #DIV/0! | 3.67E-05 |
| Q9EQV6 | LVQEVTDFAK | 1,Xle->Cys[L]; | 0.0 | 52.2 | #DIV/0! | 3.80E-05 |
| Q9EQV6 | ADRDQYELLCLDNTR | 6,sulfo+amino[Y]; | 0.0 | 12.3 | #DIV/0! | 4.11E-05 |
| P02767 | QQAEHAVEGDCDFHILK | 11,Ub-Br2[C]; | 0.0 | 21.2 | #DIV/0! | 4.14E-05 |
| P02767 | VRVELLHNPAFCSMATAK | 12,Carbamidomethyl[C]; | 0.0 | 12.2 | #DIV/0! | 4.18E-05 |
| Q9EQV6 | FSVATQICNITPGK | 8,Carbamidomethyl[C]; | 0.0 | 63.3 | #DIV/0! | 4.19E-05 |
| Q9EQV6 | FSVATQICNITPGK | 8,Carbamidomethyl[C]; | 0.0 | 63.3 | #DIV/0! | 4.19E-05 |
| P06238 | TYEWTVPK | 1,Hex(1)HexNAc(1)[T]; | 0.0 | 12.5 | #DIV/0! | 4.70E-05 |
| Q9EQV6 | AADKDNCFATEGPNLVAR | 7,NEM_2H(5)+H2O[C]; | 0.0 | 99.0 | #DIV/0! | 4.94E-05 |
| Q9EQV6 | FNNDADLSGITEDAPLK | 0,GIST-Quat_2H(3)[AnyN-term]; | 0.0 | 11.2 | #DIV/0! | 5.18E-05 |
| Q9EQV6 | SHAVVARNGDGKEDLIWEILK | 10,Cation_Ag[D]; | 0.0 | 23.5 | #DIV/0! | 5.23E-05 |
| P02767 | MRDVILFEK | 1,Oxidation[M]; | 0.0 | 19.7 | #DIV/0! | 5.27E-05 |
| Q9EQV6 | WCAVSEHENTK | 2,Carbamidomethyl[C]; | 0.0 | 84.0 | #DIV/0! | 5.32E-05 |
| Q9EQV6 | QEPERNECFLQHK | 0,Gln->pyro-Glu[AnyN-termQ];8,Carbamidomethyl[C]; | 0.0 | 38.3 | #DIV/0! | 5.58E-05 |
| Q9EQV6 | TELTADCETK | 7,Carbamidomethyl[C]; | 0.0 | 34.2 | #DIV/0! | 6.25E-05 |
| Q9EQV6 | TELTADCETK | 7,Carbamidomethyl[C]; | 0.0 | 34.2 | #DIV/0! | 6.25E-05 |
| Q9EQV6 | AFKAWAVAR | 0,Formyl[AnyN-term]; | 0.0 | 24.2 | #DIV/0! | 6.42E-05 |
| Q9EQV6 | HPYFYAPELLYYAEK | 0,Iodoacetanilide_13C(6)[AnyN-term]; | 0.0 | 15.7 | #DIV/0! | 6.50E-05 |
| Q9EQV6 | TFTFHSDICTLPDKEK | 9,Carbamidomethyl[C]; | 0.0 | 64.7 | #DIV/0! | 7.21E-05 |
| Q9EQV6 | QEPERNECFLQHKDDNPNLPPFQRPEAEAMCTSFQENPTSFLGHYLHEVAR | 31,GG[C](Dicarbamidomethyl[C]); | 0.0 | 166.8 | #DIV/0! | 7.39E-05 |
| P06238 | ADDKLFPGQQYLYVLR | 3,Cation_Na[D]; | 0.0 | 11.3 | #DIV/0! | 7.46E-05 |
| Q9EQV6 | SALTVDETYVPK | 0,CAF[AnyN-term]; | 0.0 | 10.5 | #DIV/0! | 7.49E-05 |
| Q9EQV6 | YCSSQIDAEMR | 2,Carbamidomethyl[C]; | 0.0 | 48.8 | #DIV/0! | 7.58E-05 |
| Q9EQV6 | FLGHYLHEVAR | 0,Hex[AnyN-term]; | 0.0 | 24.2 | #DIV/0! | 7.84E-05 |
| Q9EQV6 | HTTIFEVLPQK | 3,Dehydrated[T]; | 0.0 | 33.5 | #DIV/0! | 8.63E-05 |
| Q9EQV6 | YNAELESGNQFVLYR | 0,Methyl[AnyN-term]; | 0.0 | 21.0 | #DIV/0! | 9.57E-05 |
| Q9EQV6 | PEAEAMCTSFQENPTSFLGHYLHEVAR | 12,Glu->His[E]; | 0.0 | 7.5 | #DIV/0! | 9.99E-05 |
| P02767 | MQQVESSLQPETLKK | 5,Cation_Na[E]; | 0.0 | 28.5 | #DIV/0! | 1.03E-04 |
| P06911 | MQQVESSLQPETLKK | 5,Cation_Na[E]; | 0.0 | 28.5 | #DIV/0! | 1.03E-04 |
| P06911 | MQQVESSLQPETLKK | 5,Cation_Na[E]; | 0.0 | 28.5 | #DIV/0! | 1.03E-04 |
| Q9EQV6 | LAQAPNHVVVSR | 0,ESP_2H(10)[AnyN-term]; | 0.0 | 25.8 | #DIV/0! | 1.10E-04 |
| Q9EQV6 | TSFLGHYLHEVAR | 1,Palmitoyl[T]; | 0.0 | 12.2 | #DIV/0! | 1.24E-04 |
| Q9EQV6 | CFATEGPNLVAR | 0,Methyl_2H(2)[AnyN-term]; | 0.0 | 36.2 | #DIV/0! | 1.25E-04 |
| Q9EQV6 | CTSFQENPTSFLGHYLHEVAR | 7,Asn->His[N]; | 0.0 | 13.7 | #DIV/0! | 1.39E-04 |
| Q9EQV6 | CVEDYLSAILNR | 1,2-dimethylsuccinyl[C]; | 0.0 | 23.8 | #DIV/0! | 1.53E-04 |
| Q9EQV6 | EGYNGYTGAFQCLVEK | 4,Deamidated[N];6,Dehydrated[Y];12,Carbamidomethyl[C]; | 0.0 | 16.3 | #DIV/0! | 1.67E-04 |
| Q9EQV6 | MQHIDVLENSLGFK | 0,Unknown_248[AnyN-term]; | 0.0 | 39.5 | #DIV/0! | 1.69E-04 |
| Q9EQV6 | FDEFFSQGCAPGYK | 3,Cation_Na[E];9,Carbamidomethyl[C]; | 0.0 | 118.2 | #DIV/0! | 1.69E-04 |
| P02767 | SPMYSIITPNVLR | 0,C+12[AnyN-term]; | 0.0 | 12.5 | #DIV/0! | 1.76E-04 |
| Q9EQV6 | VFLGTFLYEYSR | 0,ICPL_13C(6)[AnyN-term]; | 0.0 | 18.2 | #DIV/0! | 1.87E-04 |
| Q9EQV6 | EGVCPEGSIDSAPVK | 4,Cys->Arg[C]; | 0.0 | 18.7 | #DIV/0! | 1.90E-04 |
| P06238 | GGEFEMMPLGVNK | 6,Oxidation[M]; | 0.0 | 24.8 | #DIV/0! | 1.94E-04 |
| Q9EQV6 | LRDNYGELADCCAK | 11,DTT[C]; | 0.0 | 19.3 | #DIV/0! | 1.98E-04 |
| Q9EQV6 | NTAPTSSPSITAPR | 2,Hex(1)HexNAc(1)[T]; | 0.0 | 7.3 | #DIV/0! | 2.03E-04 |
| Q9EQV6 | CKVNSGAFPAPIEK | 0,Pyro-carbamidomethyl[AnyN-termC]; | 0.0 | 43.0 | #DIV/0! | 2.27E-04 |
| Q9EQV6 | LQACCDKPVLQK | 5,maleimide[C]; | 0.0 | 16.2 | #DIV/0! | 2.30E-04 |
| Q9EQV6 | NTPPTMDTDGSYFLYSK | 1,Deamidated[N]; | 0.0 | 6.7 | #DIV/0! | 2.39E-04 |
| P06238 | NTPPTMDTDGSYFLYSK | 1,Deamidated[N]; | 0.0 | 6.7 | #DIV/0! | 2.39E-04 |
| Q9EQV6 | LRDNYGELADCCAK | 12,DTT[C]; | 0.0 | 10.3 | #DIV/0! | 2.46E-04 |
| Q9EQV6 | PTSFLGHYLHEVAR | 0,iTRAQ4plex114[AnyN-term]; | 0.0 | 16.7 | #DIV/0! | 2.60E-04 |
| Q9EQV6 | LLECADDRAELAK | 4,Carbamidomethyl[C]; | 0.0 | 33.7 | #DIV/0! | 2.63E-04 |
| Q9EQV6 | LCVLHEKTPVSEK | 2,Carbamidomethyl[C]; | 0.0 | 16.2 | #DIV/0! | 2.69E-04 |
| Q9EQV6 | EAMCTSFQENPTSFLGHYLHEVAR | 4,Palmitoyl[C]; | 0.0 | 9.2 | #DIV/0! | 2.77E-04 |
| Q9EQV6 | QEPERNECFLQHKDDNPNLPPFQRPEAEAMCTSFQENPTSFLGHYLHEVAR | 8,GG[C](Dicarbamidomethyl[C]); | 0.0 | 80.0 | #DIV/0! | 2.85E-04 |
| Q9EQV6 | PERNECFLQHKDDNPNLPPFQRPEAEAMCTSFQENPTSFLGHYLHEVAR | 29,4AcAllylGal[C]; | 0.0 | 8.2 | #DIV/0! | 2.86E-04 |
| Q9EQV6 | APQVSTPTLVEAARNLGRVGTKCCTLPEAQRLPCVEDYLSAILNR | 34,Iodoacetanilide[C]; | 0.0 | 15.5 | #DIV/0! | 2.87E-04 |
| P06238 | DKDNEDFQESNR | 0,Biotin[AnyN-term]; | 0.0 | 5.5 | #DIV/0! | 3.01E-04 |
| Q9EQV6 | FTFHSDICTLPDKEK | 8,Carbamidomethyl[C]; | 0.0 | 24.8 | #DIV/0! | 3.02E-04 |
| Q9EQV6 | KYNAELESGNQFVLYR | 5,Cation_Na[E]; | 0.0 | 16.8 | #DIV/0! | 3.08E-04 |
| Q9EQV6 | LPQKNTAPTSSPSITAPR | 4,Xlink_DTSSP[174][K]; | 0.0 | 17.8 | #DIV/0! | 3.12E-04 |
| Q9EQV6 | RHPYFYAPELLYYAEK | 12,Dehydrated[Y]; | 0.0 | 35.5 | #DIV/0! | 3.16E-04 |
| Q9EQV6 | SIHTLFGDKLCAIPK | 11,Carbamidomethyl[C]; | 0.0 | 80.5 | #DIV/0! | 3.21E-04 |
| P02767 | DSTYSMSSTLSLTK | 6,Oxidation[M]; | 0.0 | 135.2 | #DIV/0! | 3.24E-04 |
| P06238 | DSTYSMSSTLSLTK | 6,Oxidation[M]; | 0.0 | 135.2 | #DIV/0! | 3.24E-04 |
| Q9EQV6 | CSVLHEGLHNHHTEK | 1,cysTMT[C]; | 0.0 | 37.7 | #DIV/0! | 3.31E-04 |
| P06238 | CSVLHEGLHNHHTEK | 1,cysTMT[C]; | 0.0 | 37.7 | #DIV/0! | 3.31E-04 |
| Q9EQV6 | VFMQHIDVLENSLGFK | 3,Oxidation[M]; | 0.0 | 145.0 | #DIV/0! | 3.38E-04 |
| Q9EQV6 | RHPDYSVSLLLR | 12,Arg->Tyr[R]; | 0.0 | 12.3 | #DIV/0! | 3.46E-04 |
| Q9EQV6 | TNCELYEK | 3,Carbamidomethyl[C];4,Cation_Na[E]; | 0.0 | 15.3 | #DIV/0! | 3.68E-04 |
| P06238 | PGPCGSEVATVPETGRK | 4,EHD-diphenylpentanone[C]; | 0.0 | 13.5 | #DIV/0! | 3.69E-04 |
| P02767 | WEYCNLKR | 4,Carbamidomethyl[C]; | 0.0 | 16.3 | #DIV/0! | 3.75E-04 |
| P06238 | AAPQLPMEELVSLSK | 9,Cation_Na[E]; | 0.0 | 7.5 | #DIV/0! | 3.76E-04 |
| Q9EQV6 | LLEACTFHKS | 5,Carbamidomethyl[C]; | 0.0 | 24.3 | #DIV/0! | 3.83E-04 |
| Q9WUW3 | SFECLHPEIK | 4,Carbamidomethyl[C]; | 0.0 | 31.0 | #DIV/0! | 4.09E-04 |
| Q9EQV6 | CTSFQENPTSFLGHYLHEVAR | 1,Carbamidomethyl[C];7,Deamidated[N]; | 0.0 | 9.2 | #DIV/0! | 4.21E-04 |
| Q9EQV6 | TCVADENAENCDKSIHTLFGDK | 2,Carbamidomethyl[C];10,Deamidated[N];11,Carbamidomethyl[C]; | 0.0 | 42.2 | #DIV/0! | 4.25E-04 |
| Q9EQV6 | CLRDGGGDVAFVK | 1,Carbamidomethyl[C]; | 0.0 | 35.2 | #DIV/0! | 4.26E-04 |
| P02767 | MQQVEASLQPETLRR | 1,Met->Asp[M]; | 0.0 | 8.7 | #DIV/0! | 4.33E-04 |
| Q9EQV6 | SFQENPTSFLGHYLHEVAR | 5,Asn->His[N]; | 0.0 | 7.2 | #DIV/0! | 4.35E-04 |
| Q9EQV6 | QTALAELVKHKPK | 0,Carbamyl[AnyN-term]; | 0.0 | 36.8 | #DIV/0! | 4.42E-04 |
| Q9EQV6 | TVLAEFQPLVEEPK | 1,FMN[T]; | 0.0 | 70.2 | #DIV/0! | 4.42E-04 |
| Q9EQV6 | HSEAFSVNFADSEEAKK | 0,iTRAQ4plex114[AnyN-term]; | 0.0 | 25.7 | #DIV/0! | 4.44E-04 |
| P02767 | YVMLPVADQEKCELHYEK | 12,Carbamidomethyl[C]; | 0.0 | 34.5 | #DIV/0! | 4.48E-04 |
| Q9EQV6 | TGSEVSSVFIFPPK | 0,iTRAQ4plex114[AnyN-term]; | 0.0 | 7.5 | #DIV/0! | 4.56E-04 |
| P06238 | TGSEVSSVFIFPPK | 0,iTRAQ4plex114[AnyN-term]; | 0.0 | 7.5 | #DIV/0! | 4.56E-04 |
| P02767 | SGAQATWAELSWPHEK | 9,Cation_Na[E]; | 0.0 | 48.0 | #DIV/0! | 4.68E-04 |
| Q9EQV6 | HPYFYAPELLYYAEK | 8,Cation_Ca[II][E]; | 0.0 | 14.7 | #DIV/0! | 4.76E-04 |
| Q9EQV6 | GTEAAGATVVEAVPMSLPPQVK | 0,ISD_z+2_ion[AnyN-term]; | 0.0 | 19.3 | #DIV/0! | 4.94E-04 |
| Q9EQV6 | FSWFIDDVEVHTAQTHAPEK | 12,Dehydrated[T]; | 0.0 | 7.8 | #DIV/0! | 4.96E-04 |
| P02767 | AVLDVAETGTEAAAATGVK | 8,Dehydrated[T]; | 0.0 | 20.7 | #DIV/0! | 4.99E-04 |
| Q9EQV6 | QLKTVMGDFAQFVDK | 0,Dimethyl[AnyN-term](Ethyl[AnyN-term]); | 0.0 | 8.3 | #DIV/0! | 5.21E-04 |
| Q9EQV6 | NGDGKEDLIWEILK | 6,Cation_Ca[II][E]; | 0.0 | 1.7 | #DIV/0! | 5.21E-04 |
| Q9EQV6 | CLRDGGGDVAFVK | 4,Cation_Ag[D]; | 0.0 | 18.2 | #DIV/0! | 5.24E-04 |
| Q9EQV6 | VCQELSTLGKDDFR | 2,Carbamidomethyl[C]; | 0.0 | 87.5 | #DIV/0! | 5.27E-04 |
| P18292 | ETFMDCLEGR | 6,Carbamidomethyl[C]; | 0.0 | 21.2 | #DIV/0! | 5.43E-04 |
| Q9EQV6 | YNEVLTQCCTESDK | 3,Cation_Na[E];8,Carbamidomethyl[C];9,Carbamidomethyl[C]; | 0.0 | 83.2 | #DIV/0! | 5.47E-04 |
| Q9EQV6 | YGTVLAEFQPLVEEPK | 0,Carbamidomethyl[AnyN-term]; | 0.0 | 42.2 | #DIV/0! | 5.52E-04 |
| P06238 | FTKENEGTYYGPDGR | 0,ISD_z+2_ion[AnyN-term]; | 0.0 | 6.2 | #DIV/0! | 5.61E-04 |
| Q9EQV6 | LRDNYGELADCCAK | 11,Hydroxycinnamyl[C]; | 0.0 | 11.5 | #DIV/0! | 5.61E-04 |
| Q9EQV6 | KTSYQDCIK | 7,CarbamidomethylDTT[C]; | 0.0 | 6.0 | #DIV/0! | 5.73E-04 |
| P26644 | IEKNYKNTEPVMDSDGSFFMYSK | 7,Hep[N]; | 0.0 | 13.2 | #DIV/0! | 5.76E-04 |
| Q9EQV6 | AADKDNCFATEGPNLVAR | 7,DiDehydro[C]; | 0.0 | 26.7 | #DIV/0! | 5.80E-04 |
| P06238 | ALMAYAFALAGNQEK | 3,Met->His[M]; | 0.0 | 9.5 | #DIV/0! | 5.84E-04 |
| Q9EQV6 | PDYSVSLLLR | 1,Pro->Cys[P]; | 0.0 | 33.8 | #DIV/0! | 5.89E-04 |
| Q9EQV6 | KQTALAELVK | 0,Hex[AnyN-term]; | 0.0 | 17.3 | #DIV/0! | 6.01E-04 |
| Q9EQV6 | DYEKDKVCQELSTLGKDDFR | 2,Isopropylphospho[Y]; | 0.0 | 13.8 | #DIV/0! | 6.18E-04 |
| P23680 | SSPSIVLGQEQDTYGGGFDK | 3,Pro->Tyr[P]; | 0.0 | 18.0 | #DIV/0! | 6.24E-04 |
| Q9EQV6 | AMFHVNKPR | 2,Oxidation[M]; | 0.0 | 21.3 | #DIV/0! | 6.32E-04 |
| Q9EQV6 | DYEKDKVCQELSTLGKDDFR | 0,CAF[AnyN-term]; | 0.0 | 12.3 | #DIV/0! | 6.33E-04 |
| Q9EQV6 | LNGDWFSIVVASNK | 4,Cation_Na[D]; | 0.0 | 8.0 | #DIV/0! | 6.60E-04 |
| Q9EQV6 | GDGKEDLIWEILK | 0,Phenylisocyanate_2H(5)[AnyN-term]; | 0.0 | 6.3 | #DIV/0! | 6.62E-04 |
| P02767 | GAVSPVGVQPILNK | 2,Carbonyl[A]; | 0.0 | 39.5 | #DIV/0! | 6.69E-04 |
| P04639 | AIEDYVNEFSAR | 0,C+12[AnyN-term]; | 0.0 | 21.7 | #DIV/0! | 6.72E-04 |
| P20762 | LSVDTDSWMR | 9,Oxidation[M]; | 0.0 | 10.8 | #DIV/0! | 6.72E-04 |
| Q9EQV6 | FATEGPNLVAR | 0,Pyridylacetyl[AnyN-term](Phenylisocyanate[AnyN-term]); | 0.0 | 8.3 | #DIV/0! | 6.79E-04 |
| Q9EQV6 | KTEEDLCVGCFQPIPMDSSDLKPVLK | 7,Carbamidomethyl[C];10,Carbamidomethyl[C]; | 0.0 | 44.3 | #DIV/0! | 6.80E-04 |
| Q9EQV6 | KTEEDLCVGCFQPIPMDSSDLKPVLK | 7,Carbamidomethyl[C];10,Carbamidomethyl[C]; | 0.0 | 44.3 | #DIV/0! | 6.80E-04 |
| Q9EQV6 | FQENPTSFLGHYLHEVAR | 5,Pro->Cys[P]; | 0.0 | 94.7 | #DIV/0! | 7.00E-04 |
| P02767 | WKIDGSEQRDGVLDSVTDQDSKDSTYSMSSTLSLTK | 1,Trp->Cys[W]; | 0.0 | 6.2 | #DIV/0! | 7.09E-04 |
| Q9EQV6 | CFATEGPNLVAR | 0,ICPL_13C(6)[AnyN-term]; | 0.0 | 27.7 | #DIV/0! | 7.25E-04 |
| Q9EQV6 | SFLGHYLHEVAR | 0,Ethylphosphate[AnyN-term]; | 0.0 | 5.2 | #DIV/0! | 7.33E-04 |
| P02767 | FLRFNPVTGEVPPR | 0,TNBS[AnyN-term]; | 0.0 | 5.2 | #DIV/0! | 7.33E-04 |
| Q9EQV6 | AEAMCTSFQENPTSFLGHYLHEVAR | 5,BHT[C]; | 0.0 | 40.7 | #DIV/0! | 7.37E-04 |
| Q9EQV6 | KTSYQDCIK | 7,Carbamidomethyl[C]; | 0.0 | 9.2 | #DIV/0! | 7.38E-04 |
| Q9EQV6 | KQTALAELVK | 0,Carbamyl[AnyN-term]; | 0.0 | 10.0 | #DIV/0! | 7.46E-04 |
| Q9EQV6 | LRDNYGELADCCAK | 7,Cation_Na[E];11,Carbamidomethyl[C];12,Carbamidomethyl[C]; | 0.0 | 18.3 | #DIV/0! | 7.56E-04 |
| P02767 | AVLDVAETGTEAAAATGVK | 7,Cation_Na[E];11,Cation_Na[E]; | 0.0 | 15.2 | #DIV/0! | 8.01E-04 |
| Q9EQV6 | RHPYFYAPELLYYAEK | 4,Dehydrated[Y]; | 0.0 | 36.3 | #DIV/0! | 8.23E-04 |
| Q9EQV6 | NNYHSEAFSVNFADSEEAKK | 14,Cation_Na[D]; | 0.0 | 24.8 | #DIV/0! | 8.36E-04 |
| Q9EQV6 | FSWFIDDVEVHTAQTHAPEK | 7,Cation_Fe[III][D]; | 0.0 | 13.5 | #DIV/0! | 8.41E-04 |
| P02767 | GECQSEGVLFFQGNR | 3,Carbamidomethyl[C];6,Cation_Na[E]; | 0.0 | 14.3 | #DIV/0! | 8.44E-04 |
| P26644 | CPFPSRPDNGFVNYPAKPVLSYK | 1,Carbamidomethyl[C];9,Deamidated[N]; | 0.0 | 30.5 | #DIV/0! | 8.66E-04 |
| Q9EQV6 | DYRLYLGHSYVTAIRNQREGVCPEGSIDSAPVK | 0,Unknown_234[AnyN-term]; | 0.0 | 2.2 | #DIV/0! | 8.87E-04 |
| Q9EQV6 | DNIPADLPSIAADFVEDK | 0,Ethylphosphate[AnyN-term]; | 0.0 | 14.8 | #DIV/0! | 8.88E-04 |
| Q9EQV6 | DNCFATEGPNLVAR | 3,Iodoacetanilide[C]; | 0.0 | 1.2 | #DIV/0! | 9.17E-04 |
| P02767 | CNLLAEK | 1,Carbamidomethyl[C]; | 0.0 | 26.2 | #DIV/0! | 9.25E-04 |
| Q9EQV6 | ENPTSFLGHYLHEVAR | 6,Phe->Pro[F]; | 0.0 | 16.0 | #DIV/0! | 9.61E-04 |
| Q9EQV6 | EGAQEMDCNDETVFQAVDTALK | 8,IBTP[C]; | 0.0 | 8.0 | #DIV/0! | 9.61E-04 |
| Q9EQV6 | VFNNDADLSGITEDAPLK | 0,Formyl[AnyN-term]; | 0.0 | 15.0 | #DIV/0! | 9.83E-04 |
| Q9EQV6 | PNAEFAEITK | 0,iTRAQ4plex114[AnyN-term]; | 0.0 | 59.8 | #DIV/0! | 9.94E-04 |
| Q9EQV6 | YMCENQATISSK | 2,Oxidation[M]; | 0.0 | 17.7 | #DIV/0! | 1.03E-03 |
| Q9EQV6 | DVLTITLTPK | 6,Dehydrated[T]; | 0.0 | 7.0 | #DIV/0! | 1.06E-03 |
| P06238 | DVLTITLTPK | 6,Dehydrated[T]; | 0.0 | 7.0 | #DIV/0! | 1.06E-03 |
| Q9EQV6 | TSFQENPTSFLGHYLHEVAR | 6,Hex(3)[N]; | 0.0 | 4.5 | #DIV/0! | 1.11E-03 |
| P06238 | AENGDFASFR | 3,Deamidated[N]; | 0.0 | 29.2 | #DIV/0! | 1.12E-03 |
| P06238 | QSGVKEEHSFTVMEFVLPR | 2,Ser->Gly[S]; | 0.0 | 15.2 | #DIV/0! | 1.14E-03 |
| P06238 | QSGVKEEHSFTVMEFVLPR | 2,Ser->Gly[S]; | 0.0 | 15.2 | #DIV/0! | 1.14E-03 |
| P06238 | QSGVKEEHSFTVMEFVLPR | 2,Ser->Gly[S]; | 0.0 | 15.2 | #DIV/0! | 1.14E-03 |
| Q9EQV6 | FQENPTSFLGHYLHEVAR | 4,Hep[N]; | 0.0 | 10.7 | #DIV/0! | 1.16E-03 |
| P06238 | LVCQDPCNVYETLTEYWR | 0,Phenylisocyanate_2H(5)[AnyN-term]; | 0.0 | 9.8 | #DIV/0! | 1.21E-03 |
| Q9EQV6 | LQACCDKPVLQK | 4,maleimide[C]; | 0.0 | 15.0 | #DIV/0! | 1.22E-03 |
| Q9EQV6 | TCVADENAENCDKSIHTLFGDK | 2,Carbamidomethyl[C];11,Carbamidomethyl[C]; | 0.0 | 109.0 | #DIV/0! | 1.22E-03 |
| Q9EQV6 | CPEGSIDSAPVK | 1,Carbamidomethyl[C]; | 0.0 | 18.2 | #DIV/0! | 1.24E-03 |
| Q9EQV6 | FQENPTSFLGHYLHEVAR | 4,HexN[N]; | 0.0 | 15.2 | #DIV/0! | 1.28E-03 |
| P02767 | LYQAEAFTADFQQSR | 4,Ala->Asp[A]; | 0.0 | 17.3 | #DIV/0! | 1.29E-03 |
| P22057 | KFITFSKDQGLTEEDIVFLPQPDK | 0,GIST-Quat_2H(3)[AnyN-term]; | 0.0 | 5.5 | #DIV/0! | 1.29E-03 |
| Q9EQV6 | YNEVLTQCCTESDKAACLTPK | 2,Deamidated[N];9,GG[C](Dicarbamidomethyl[C]);17,Carbamidomethyl[C]; | 0.0 | 14.5 | #DIV/0! | 1.29E-03 |
| P02767 | AVLDVAETGTEAAAATGVK | 10,Dehydrated[T]; | 0.0 | 4.8 | #DIV/0! | 1.33E-03 |
| Q9EQV6 | DLKQEDFQLLCPDGTK | 12,Pro->His[P]; | 0.0 | 3.8 | #DIV/0! | 1.40E-03 |
| Q63416 | HPEYNKDTLDNDIMLIK | 5,Hep[N]; | 0.0 | 3.8 | #DIV/0! | 1.40E-03 |
| Q9EQV6 | NTPPTMDTDGSYFLYSK | 0,Methyl[AnyN-term]; | 0.0 | 29.2 | #DIV/0! | 1.44E-03 |
| P06238 | NTPPTMDTDGSYFLYSK | 0,Methyl[AnyN-term]; | 0.0 | 29.2 | #DIV/0! | 1.44E-03 |
| P02767 | DGVLDSVTDQDSK | 5,Dehydrated[D]; | 0.0 | 1.3 | #DIV/0! | 1.46E-03 |
| P06238 | DGVLDSVTDQDSK | 5,Dehydrated[D]; | 0.0 | 1.3 | #DIV/0! | 1.46E-03 |
| Q9EQV6 | AADKDNCFATEGPNLVAR | 6,Asn->Tyr[N]; | 0.0 | 50.2 | #DIV/0! | 1.51E-03 |
| Q9EQV6 | CENQATISSK | 1,MDCC[C]; | 0.0 | 11.2 | #DIV/0! | 1.58E-03 |
| Q9EQV6 | SQYLQKCPYEEHIKLVQEVTDFAK | 7,monomethylphosphothione[C]; | 0.0 | 5.2 | #DIV/0! | 1.59E-03 |
| Q9EQV6 | RHPDYSVSLLLR | 3,Pro->Cys[P]; | 0.0 | 25.2 | #DIV/0! | 1.60E-03 |
| Q9EQV6 | NNYHSEAFSVNFADSEEAK | 4,His->Met[H]; | 0.0 | 11.3 | #DIV/0! | 1.63E-03 |
| Q9EQV6 | QLKTVMGDFAQFVDK | 0,Formyl[AnyN-term]; | 0.0 | 5.0 | #DIV/0! | 1.68E-03 |
| P26644 | VPELLGGPSVFIFPPKPK | 3,phosphoRibosyl[E]; | 0.0 | 13.5 | #DIV/0! | 1.73E-03 |
| Q9EQV6 | QRFPNAEFAEITK | 0,Pyridylacetyl[AnyN-term](Phenylisocyanate[AnyN-term]); | 0.0 | 6.2 | #DIV/0! | 1.74E-03 |
| Q9EQV6 | FATEGPNLVAR | 0,SPITC[AnyN-term]; | 0.0 | 19.3 | #DIV/0! | 1.74E-03 |
| Q9EQV6 | NNYHSEAFSVNFADSEEAKK | 6,Cation_Na[E]; | 0.0 | 31.5 | #DIV/0! | 1.76E-03 |
| Q9EQV6 | RPCFSALTVDETYVPK | 3,CarbamidomethylDTT[C]; | 0.0 | 12.8 | #DIV/0! | 1.77E-03 |
| Q9EQV6 | FSWFIDDVEVHTAQTHAPEK | 6,Cation_Ca[II][D]; | 0.0 | 11.3 | #DIV/0! | 1.78E-03 |
| Q9EQV6 | VFNNDADLSGITEDAPLK | 1,Val->Met[V]; | 0.0 | 10.5 | #DIV/0! | 1.83E-03 |
| Q9EQV6 | MCTSFQENPTSFLGHYLHEVAR | 0,Hex[AnyN-term]; | 0.0 | 6.3 | #DIV/0! | 1.83E-03 |
| P06238 | TFCSEPEKVDKDNEDFQESNR | 3,Carboxymethyl[C]; | 0.0 | 3.3 | #DIV/0! | 1.88E-03 |
| Q9EQV6 | EFKAETFTFHSDICTLPDK | 1,Cation_Na[E];14,Carbamidomethyl[C]; | 0.0 | 10.2 | #DIV/0! | 1.88E-03 |
| Q9EQV6 | ENPTSFLGHYLHEVAR | 4,PET[T]; | 0.0 | 6.5 | #DIV/0! | 1.88E-03 |
| P08934 | NKETCDVIINTLGK | 5,Carbamidomethyl[C]; | 0.0 | 10.8 | #DIV/0! | 1.89E-03 |
| Q9EQV6 | CTSFQENPTSFLGHYLHEVAR | 4,Fluoro[F]; | 0.0 | 3.7 | #DIV/0! | 1.89E-03 |
| Q9EQV6 | NQREGVCPEGSIDSAPVK | 7,4-ONE[C]; | 0.0 | 3.7 | #DIV/0! | 1.89E-03 |
| Q9EQV6 | ELPEHTLK | 4,Cation_Na[E]; | 0.0 | 1.8 | #DIV/0! | 1.89E-03 |
| P02767 | AVLDVDETGTEGAAATAVTAALK | 7,Cation_Na[E]; | 0.0 | 12.0 | #DIV/0! | 1.92E-03 |
| Q9EQV6 | LRDNYGELADCCAK | 5,Dehydrated[Y];11,Carbamidomethyl[C];12,Carbamidomethyl[C]; | 0.0 | 37.3 | #DIV/0! | 1.95E-03 |
| P06238 | SGSGCSILDEVIR | 5,Carbamidomethyl[C]; | 0.0 | 92.7 | #DIV/0! | 1.96E-03 |
| P06238 | SGSGCSILDEVIR | 5,Carbamidomethyl[C]; | 0.0 | 92.7 | #DIV/0! | 1.96E-03 |
| Q9EQV6 | PTSFLGHYLHEVAR | 2,O-Methylphosphate[T]; | 0.0 | 10.5 | #DIV/0! | 2.01E-03 |
| Q9EQV6 | DNCFATEGPNLVAR | 3,NEM_2H(5)[C]; | 0.0 | 7.7 | #DIV/0! | 2.05E-03 |
| Q9EQV6 | KTEEDLCVGCFQPIPMDSSDLKPVLK | 10,DNCB_hapten[C]; | 0.0 | 11.0 | #DIV/0! | 2.06E-03 |
| Q9EQV6 | KTEEDLCVGCFQPIPMDSSDLKPVLK | 10,DNCB_hapten[C]; | 0.0 | 11.0 | #DIV/0! | 2.06E-03 |
| Q9EQV6 | CFSALTVDETYVPK | 1,SulfurDioxide[C]; | 0.0 | 5.0 | #DIV/0! | 2.08E-03 |
| Q9EQV6 | GDGKEDLIWEILK | 0,Pyridylacetyl[AnyN-term](Phenylisocyanate[AnyN-term]); | 0.0 | 2.5 | #DIV/0! | 2.08E-03 |
| Q9EQV6 | KLVEKFLEEVKNNYHSEAFSVNFADSEEAKK | 20,Pentose[S]; | 0.0 | 2.5 | #DIV/0! | 2.08E-03 |
| P06238 | AAPQLPMEELVSLSK | 8,Cation_Na[E]; | 0.0 | 2.5 | #DIV/0! | 2.08E-03 |
| Q9EQV6 | SNSMVTLGCLVK | 4,Oxidation[M];9,Carbamidomethyl[C]; | 0.0 | 42.2 | #DIV/0! | 2.09E-03 |
| P06238 | SNSMVTLGCLVK | 4,Oxidation[M];9,Carbamidomethyl[C]; | 0.0 | 42.2 | #DIV/0! | 2.09E-03 |
| Q9EQV6 | HPYFYAPELLYYAEK | 0,Methylthio[AnyN-term]; | 0.0 | 14.3 | #DIV/0! | 2.10E-03 |
| Q9EQV6 | RHPDYSVSLLLR | 0,ISD_z+2_ion[AnyN-term]; | 0.0 | 20.3 | #DIV/0! | 2.13E-03 |
| Q9EQV6 | VAHPASSTKVDK | 0,Carbamidomethyl[AnyN-term]; | 0.0 | 3.0 | #DIV/0! | 2.13E-03 |
| P06238 | VAHPASSTKVDK | 0,Carbamidomethyl[AnyN-term]; | 0.0 | 3.0 | #DIV/0! | 2.13E-03 |
| P26644 | VAHPASSTKVDK | 0,Carbamidomethyl[AnyN-term]; | 0.0 | 3.0 | #DIV/0! | 2.13E-03 |
| Q9EQV6 | ALTVDETYVPK | 0,Dimethyl[AnyN-term](Ethyl[AnyN-term]); | 0.0 | 55.0 | #DIV/0! | 2.18E-03 |
| Q9EQV6 | NTPPTMDTDGSYFLYSK | 0,Methyl_2H(2)[AnyN-term]; | 0.0 | 36.7 | #DIV/0! | 2.24E-03 |
| P06238 | NTPPTMDTDGSYFLYSK | 0,Methyl_2H(2)[AnyN-term]; | 0.0 | 36.7 | #DIV/0! | 2.24E-03 |
| P08934 | ETCDVIINTLGK | 3,Carbamidomethyl[C]; | 0.0 | 15.0 | #DIV/0! | 2.33E-03 |
| Q9EQV6 | LPCVEDYLSAILNR | 3,Carbamidomethyl[C];5,Cation_Na[E];7,Dehydrated[Y]; | 0.0 | 10.3 | #DIV/0! | 2.36E-03 |
| Q9EQV6 | TVTVSCEFFESQDQVPGGENPADTQDAK | 6,Carbamidomethyl[C]; | 0.0 | 23.5 | #DIV/0! | 2.42E-03 |
| P02767 | VLDVDETGTEGAAATAVTAALK | 0,Propyl_2H(6)[AnyN-term]; | 0.0 | 14.0 | #DIV/0! | 2.46E-03 |
| Q9EQV6 | MNGQPQENYK | 0,ISD_z+2_ion[AnyN-term]; | 0.0 | 15.2 | #DIV/0! | 2.48E-03 |
| P02767 | NPLPAAHETVAK | 0,Carbamyl[AnyN-term]; | 0.0 | 15.2 | #DIV/0! | 2.48E-03 |
| P06238 | MNGQPQENYK | 0,ISD_z+2_ion[AnyN-term]; | 0.0 | 15.2 | #DIV/0! | 2.48E-03 |
| Q9EQV6 | SYFVEYLIKESPCTQSQDSCSLQASDSEPVGLCQGSLIK | 33,Carbamidomethyl[C]; | 0.0 | 7.8 | #DIV/0! | 2.52E-03 |
| P06238 | PCICTGSEVSSVFIFPPK | 5,HexNAc(1)NeuAc(1)[T]; | 0.0 | 7.7 | #DIV/0! | 2.62E-03 |
| P02767 | LFQEEFPGIPYPPDAAVECHR | 11,Dehydrated[Y];19,Carbamidomethyl[C]; | 0.0 | 17.0 | #DIV/0! | 2.66E-03 |
| Q9EQV6 | KDDNPNLPPFQRPEAEAMCTSFQENPTSFLGHYLHEVAR | 19,Carbamidomethyl[C]; | 0.0 | 10.3 | #DIV/0! | 2.68E-03 |
| P55314 | YVCNCVIGYIGER | 3,Carbamidomethyl[C];5,Carbamidomethyl[C]; | 0.0 | 26.5 | #DIV/0! | 2.76E-03 |
| P01681 | DVQMTQSPSYLAASPGESVSISCK | 4,Oxidation[M];17,Cation_Na[E];23,Carbamidomethyl[C]; | 0.0 | 8.8 | #DIV/0! | 2.83E-03 |
| Q9EQV6 | CTSFQENPTSFLGHYLHEVAR | 1,Carbamidomethyl[C];6,Cation_Na[E]; | 0.0 | 18.0 | #DIV/0! | 2.84E-03 |
| P02767 | LQTEGDGIYTLNSEK | 4,Glu->His[E]; | 0.0 | 7.0 | #DIV/0! | 2.89E-03 |
| Q9EQV6 | ENPTSFLGHYLHEVAR | 2,Hex(3)[N]; | 0.0 | 15.5 | #DIV/0! | 2.90E-03 |
| Q9EQV6 | SFQENPTSFLGHYLHEVAR | 1,N-dimethylphosphate[S]; | 0.0 | 7.5 | #DIV/0! | 2.90E-03 |
| Q9EQV6 | DNPNLPPFQRPEAEAMCTSFQENPTSFLGHYLHEVAR | 17,Carbamidomethyl[C]; | 0.0 | 12.0 | #DIV/0! | 3.11E-03 |
| Q9EQV6 | CDKPVLQK | 1,Carbamidomethyl[C]; | 0.0 | 4.8 | #DIV/0! | 3.16E-03 |
| Q9EQV6 | ADKDNCFATEGPNLVAR | 6,DHP[C]; | 0.0 | 16.5 | #DIV/0! | 3.16E-03 |
| Q9EQV6 | MCENQATISSK | 0,Unknown_248[AnyN-term]; | 0.0 | 6.2 | #DIV/0! | 3.23E-03 |
| P02767 | ILGSDVQQIAVTK | 2,Xle->Cys[L]; | 0.0 | 6.2 | #DIV/0! | 3.23E-03 |
| Q9EQV6 | DAEEAATGECTTTLGK | 3,Cation_Na[E];10,Carbamidomethyl[C]; | 0.0 | 6.3 | #DIV/0! | 3.27E-03 |
| Q9EQV6 | DAEEAATGECTTTLGK | 3,Cation_Na[E];10,Carbamidomethyl[C]; | 0.0 | 6.3 | #DIV/0! | 3.27E-03 |
| Q9EQV6 | GSIQHLPEQEEPEDSKGK | 2,Dehydrated[S]; | 0.0 | 9.7 | #DIV/0! | 3.30E-03 |
| Q9EQV6 | YGTVLAEFQPLVEEPK | 0,Carboxymethyl[AnyN-term]; | 0.0 | 33.8 | #DIV/0! | 3.31E-03 |
| P22283 | LVNCPFEEQTEQLKR | 4,Carbamidomethyl[C]; | 0.0 | 34.2 | #DIV/0! | 3.33E-03 |
| Q9EQV6 | VFNNDADLSGITEDAPLK | 1,Val->Glu[V]; | 0.0 | 8.2 | #DIV/0! | 3.35E-03 |
| Q9EQV6 | HPDYSVSLLLR | 0,Iodoacetanilide_13C(6)[AnyN-term]; | 0.0 | 34.7 | #DIV/0! | 3.39E-03 |
| Q9EQV6 | CTSFQENPTSFLGHYLHEVAR | 1,2-dimethylsuccinyl[C]; | 0.0 | 12.3 | #DIV/0! | 3.40E-03 |
| P02767 | KPPEIANGYVEHLVR | 4,Glu->Lys[E]; | 0.0 | 5.7 | #DIV/0! | 3.41E-03 |
| Q9WUW3 | AFAELWAFDAAQGK | 0,Methyl[AnyN-term]; | 0.0 | 5.5 | #DIV/0! | 3.45E-03 |
| Q9EQV6 | MGDFAQFVDK | 0,C+12[AnyN-term]; | 0.0 | 22.8 | #DIV/0! | 3.47E-03 |
| P06238 | EDNSIHWERPQKPTK | 3,Asn->Lys[N]; | 0.0 | 10.2 | #DIV/0! | 3.48E-03 |
| P06238 | EDNSIHWERPQKPTK | 3,Asn->Lys[N]; | 0.0 | 10.2 | #DIV/0! | 3.48E-03 |
| Q9EQV6 | SFQENPTSFLGHYLHEVAR | 0,NDA[AnyN-term]; | 0.0 | 9.3 | #DIV/0! | 3.52E-03 |
| Q9EQV6 | RHPYFYAPELLYYAEK | 6,Dehydrated[Y]; | 0.0 | 5.0 | #DIV/0! | 3.54E-03 |
| Q9EQV6 | GLVLIAFSQYLQK | 0,ISD_z+2_ion[AnyN-term]; | 0.0 | 8.5 | #DIV/0! | 3.59E-03 |
| P02767 | SDVDEDIIPEEDIISR | 0,Dimethyl_2H(4)[AnyN-term]; | 0.0 | 4.3 | #DIV/0! | 3.65E-03 |
| P02767 | MHSRCNADPGLSALLSDHR | 5,IED-Biotin[C]; | 0.0 | 3.8 | #DIV/0! | 3.72E-03 |
| Q9EQV6 | FSALTVDETYVPK | 0,Diethylphosphate[AnyN-term]; | 0.0 | 4.8 | #DIV/0! | 3.74E-03 |
| Q9EQV6 | SVNFADSEEAKK | 1,Glucuronyl[S]; | 0.0 | 14.5 | #DIV/0! | 3.80E-03 |
| Q9EQV6 | LRDNYGELADCCAK | 12,Hydroxycinnamyl[C]; | 0.0 | 6.8 | #DIV/0! | 3.92E-03 |
| P02767 | RPSMIDELYLPK | 3,Pent(1)HexNAc(1)[S]; | 0.0 | 7.7 | #DIV/0! | 3.96E-03 |
| Q9EQV6 | RHPYFYAPELLYYAEK | 13,Dehydrated[Y]; | 0.0 | 8.5 | #DIV/0! | 3.98E-03 |
| P02767 | DSCVGTLVVK | 3,Carbamidomethyl[C]; | 0.0 | 9.0 | #DIV/0! | 4.00E-03 |
| Q9EQV6 | LNCNDETVFQAVDTALKK | 5,Asp->Gly[D]; | 0.0 | 9.8 | #DIV/0! | 4.01E-03 |
| Q9EQV6 | TSFQENPTSFLGHYLHEVAR | 7,Pro->Cys[P]; | 0.0 | 3.3 | #DIV/0! | 4.10E-03 |
| Q9EQV6 | TVMGDFAQFVDK | 0,Ethylphosphate[AnyN-term]; | 0.0 | 3.3 | #DIV/0! | 4.10E-03 |
| Q9EQV6 | SIHTLFGDKLCAIPK | 11,Nethylmaleimide[C]; | 0.0 | 0.8 | #DIV/0! | 4.10E-03 |
| P06238 | FTKENEGTYYGPDGR | 5,Ammonia-loss[N]; | 0.0 | 0.8 | #DIV/0! | 4.10E-03 |
| P06911 | STMEEILEGLK | 0,Ethylphosphate[AnyN-term]; | 0.0 | 7.3 | #DIV/0! | 4.21E-03 |
| Q9EQV6 | YLGHSYVTAIR | 0,Methyl[AnyN-term]; | 0.0 | 6.8 | #DIV/0! | 4.23E-03 |
| Q9EQV6 | CPYEEHIK | 1,Carboxymethyl[C]; | 0.0 | 8.0 | #DIV/0! | 4.24E-03 |
| Q9EQV6 | PQVSTPTLVEAAR | 1,Pro->Arg[P]; | 0.0 | 21.5 | #DIV/0! | 4.34E-03 |
| P02767 | CNADPGLSALLSDHR | 0,Piperidine[AnyN-term]; | 0.0 | 8.5 | #DIV/0! | 4.39E-03 |
| P02767 | FNPVTGEVPPR | 0,Methyl[AnyN-term]; | 0.0 | 20.5 | #DIV/0! | 4.41E-03 |
| Q9EQV6 | DFQLFGSPLGK | 0,ISD_z+2_ion[AnyN-term]; | 0.0 | 5.3 | #DIV/0! | 4.42E-03 |
| Q9EQV6 | NCFATEGPNLVAR | 2,Cys->Dha[C]; | 0.0 | 23.2 | #DIV/0! | 4.51E-03 |
| P02767 | KLYDYCNIPLCASLSSFECGKPQVEPK | 6,GG[C](Dicarbamidomethyl[C]);7,Deamidated[N];19,Carbamidomethyl[C]; | 0.0 | 4.7 | #DIV/0! | 4.52E-03 |
| P02781 | EEFDAPPEAVEANLK | 0,NP40[AnyN-term]; | 0.0 | 28.5 | #DIV/0! | 4.54E-03 |
| P55314 | CECICPAGFQGVACEVTNRK | 3,GG[C](Dicarbamidomethyl[C]);5,Carbamidomethyl[C];14,Carbamidomethyl[C]; | 0.0 | 11.2 | #DIV/0! | 4.54E-03 |
| P02767 | EVFSTQADLSGITGDKDLMVSQVVHK | 0,C+12[AnyN-term]; | 0.0 | 4.5 | #DIV/0! | 4.55E-03 |
| Q9EQV6 | YNAELESGNQFVLYR | 0,mTRAQ_13C(6)15N(2)[AnyN-term]; | 0.0 | 10.2 | #DIV/0! | 4.60E-03 |
| Q9EQV6 | LCEAHGITR | 2,2-dimethylsuccinyl[C]; | 0.0 | 6.5 | #DIV/0! | 4.60E-03 |
| Q9EQV6 | KPVTEFATCHLAQAPNHVVVSR | 8,Thr->Arg[T]; | 0.0 | 22.7 | #DIV/0! | 4.61E-03 |
| Q9EQV6 | SFLGHYLHEVAR | 0,dNIC[AnyN-term]; | 0.0 | 9.7 | #DIV/0! | 4.69E-03 |
| P06238 | SGVKEEHSFTVMEFVLPR | 0,Succinyl_2H(4)[AnyN-term]; | 0.0 | 15.5 | #DIV/0! | 4.71E-03 |
| P06238 | SGVKEEHSFTVMEFVLPR | 0,Succinyl_2H(4)[AnyN-term]; | 0.0 | 15.5 | #DIV/0! | 4.71E-03 |
| P06238 | SGVKEEHSFTVMEFVLPR | 0,Succinyl_2H(4)[AnyN-term]; | 0.0 | 15.5 | #DIV/0! | 4.71E-03 |
| P02767 | TAHGNSTHPMHSRCNADPGLSALLSDHR | 0,Brij58[AnyN-term]; | 0.0 | 3.8 | #DIV/0! | 4.72E-03 |
| Q9EQV6 | MPNASPEELADMVAK | 1,Oxidation[M];12,Oxidation[M]; | 0.0 | 30.2 | #DIV/0! | 4.73E-03 |
| Q9EQV6 | PVTEFATCHLAQAPNHVVVSR | 8,Carbamidomethyl[C]; | 0.0 | 20.0 | #DIV/0! | 4.87E-03 |
| P02767 | NPDNDEQGPWCYTTDPDQR | 11,Carbamidomethyl[C]; | 0.0 | 19.3 | #DIV/0! | 4.90E-03 |
| P02767 | DGVLDSVTDQDSK | 1,Dehydrated[D]; | 0.0 | 4.8 | #DIV/0! | 5.03E-03 |
| P06238 | DGVLDSVTDQDSK | 1,Dehydrated[D]; | 0.0 | 4.8 | #DIV/0! | 5.03E-03 |
| Q9EQV6 | AETFTFHSDICTLPDK | 11,IED-Biotin[C]; | 0.0 | 20.5 | #DIV/0! | 5.08E-03 |
| Q9EQV6 | KATEDQLKTVMGDFAQFVDK | 3,Hex(1)HexNAc(1)Phos(1)[T]; | 0.0 | 12.5 | #DIV/0! | 5.09E-03 |
| P02770 | KEAYYLLEPAYFYPHR | 13,Dehydrated[Y]; | 0.0 | 28.0 | #DIV/0! | 5.16E-03 |
| Q9EQV6 | EAHKSEIAHR | 0,Unknown_250[AnyN-term]; | 0.0 | 4.3 | #DIV/0! | 5.26E-03 |
| P02767 | HAFSPVASVESASGEVLHSPK | 21,Lys->AminoadipicAcid[K]; | 0.0 | 10.0 | #DIV/0! | 5.35E-03 |
| Q9EQV6 | GLVLIAFSQYLQK | 14,Cation_Na[AnyC-term]; | 0.0 | 6.5 | #DIV/0! | 5.36E-03 |
| Q9EQV6 | RPCFSALTVDETYVPKEFK | 3,Carbamidomethyl[C];17,Cation_Na[E]; | 0.0 | 13.5 | #DIV/0! | 5.43E-03 |
| Q9EQV6 | ASDSSINWNNLK | 0,C+12[AnyN-term]; | 0.0 | 35.2 | #DIV/0! | 5.44E-03 |
| Q9EQV6 | YMCENQATISSK | 1,Dehydrated[Y]; | 0.0 | 5.5 | #DIV/0! | 5.45E-03 |
| Q9EQV6 | DQLKTVMGDFAQFVDK | 5,HexNAc(1)NeuAc(1)[T]; | 0.0 | 4.0 | #DIV/0! | 5.48E-03 |
| P07632 | DTANWLEINPETGVISTR | 1,Cation_Na[D]; | 0.0 | 5.3 | #DIV/0! | 5.54E-03 |
| P06238 | MFTTAPENVDKEDEDFQESNK | 1,Met->Asp[M]; | 0.0 | 7.3 | #DIV/0! | 5.70E-03 |
| Q9EQV6 | MNGQPQENYK | 7,Cation_Na[E]; | 0.0 | 4.8 | #DIV/0! | 5.73E-03 |
| P06238 | MNGQPQENYK | 7,Cation_Na[E]; | 0.0 | 4.8 | #DIV/0! | 5.73E-03 |
| Q9EQV6 | FKDLGEQHFK | 2,Lys->AminoadipicAcid[K]; | 0.0 | 9.3 | #DIV/0! | 5.82E-03 |
| P02767 | DTFQSEFYSGK | 6,Cation_Na[E]; | 0.0 | 12.5 | #DIV/0! | 5.84E-03 |
| P04639 | VMLTTTFLDDVK | 0,Carbamyl[AnyN-term]; | 0.0 | 3.5 | #DIV/0! | 5.93E-03 |
| P00762 | ANTFDNDIMLIK | 0,dNIC[AnyN-term]; | 0.0 | 9.0 | #DIV/0! | 5.96E-03 |
| Q9EQV6 | NPTSFLGHYLHEVAR | 5,Bromo[F]; | 0.0 | 11.8 | #DIV/0! | 5.98E-03 |
| P02767 | GECQSEGVLFFQGNRK | 2,Cation_Na[E];3,Carbamidomethyl[C]; | 0.0 | 13.3 | #DIV/0! | 6.00E-03 |
| P20762 | VVSTLHIQHQDWMSGK | 14,Methylamine[S]; | 0.0 | 3.7 | #DIV/0! | 6.02E-03 |
| Q9EQV6 | CTSFQENPTSFLGHYLHEVAR | 6,Cation_Na[E];7,Deamidated[N]; | 0.0 | 9.3 | #DIV/0! | 6.02E-03 |
| Q9EQV6 | SFQENPTSFLGHYLHEVAR | 0,ESP_2H(10)[AnyN-term]; | 0.0 | 18.3 | #DIV/0! | 6.07E-03 |
| Q9EQV6 | QEPERNECFLQHK | 0,Gln->pyro-Glu[AnyN-termQ];4,Cation_Na[E];8,Carbamidomethyl[C]; | 0.0 | 5.2 | #DIV/0! | 6.16E-03 |
| Q9EQV6 | NTPPTMDTDGSYFLYSK | 2,Oxidation[T]; | 0.0 | 2.2 | #DIV/0! | 6.17E-03 |
| P06238 | NTPPTMDTDGSYFLYSK | 2,Oxidation[T]; | 0.0 | 2.2 | #DIV/0! | 6.17E-03 |
| Q9EQV6 | ATEDQLKTVMGDFAQFVDK | 0,Diethylphosphate[AnyN-term]; | 0.0 | 4.5 | #DIV/0! | 6.18E-03 |
| Q9EQV6 | GSIQHLPEQEEPEDSKGK | 0,Dimethyl_2H(4)[AnyN-term]; | 0.0 | 11.3 | #DIV/0! | 6.20E-03 |
| Q9EQV6 | ACYGTVLAEFQPLVEEPK | 2,NEMsulfur[C]; | 0.0 | 17.7 | #DIV/0! | 6.26E-03 |
| P26644 | CPVPELLGGPSVFIFPPKPK | 0,C+12[AnyN-term]; | 0.0 | 14.2 | #DIV/0! | 6.38E-03 |
| Q9EQV6 | DENAENCDKSIHTLFGDK | 7,4AcAllylGal[C]; | 0.0 | 3.2 | #DIV/0! | 6.38E-03 |
| Q9EQV6 | GTTYEEYLGAEYLQAVGNIR | 2,Phosphoadenosine[T]; | 0.0 | 7.7 | #DIV/0! | 6.44E-03 |
| P02767 | AKALYQAEAFTADFQQSR | 0,Carbamyl[AnyN-term]; | 0.0 | 15.7 | #DIV/0! | 6.46E-03 |
| Q9EQV6 | FSWFIDDVEVHTAQTHAPEK | 4,Phe->Glu[F]; | 0.0 | 12.7 | #DIV/0! | 6.49E-03 |
| Q9EQV9 | FIKPTCAEALAAVSK | 6,Carbamidomethyl[C]; | 0.0 | 13.2 | #DIV/0! | 6.50E-03 |
| P02767 | EFQEKAKALYQAEAFTADFQQSR | 0,Ethylphosphate[AnyN-term]; | 0.0 | 7.2 | #DIV/0! | 6.53E-03 |
| Q9EQV6 | HPYFYAPELLYYAEK | 0,Carbamyl[AnyN-term]; | 0.0 | 5.5 | #DIV/0! | 6.60E-03 |
| Q9EQV6 | LCEAHGITR | 2,Trioxidation[C]; | 0.0 | 5.5 | #DIV/0! | 6.60E-03 |
| P02767 | TAVTAAGTPCQEWAAQEPHSHR | 10,monomethylphosphothione[C]; | 0.0 | 10.3 | #DIV/0! | 6.67E-03 |
| Q9EQV6 | FQENPTSFLGHYLHEVAR | 6,dHex(1)Hex(1)[T]; | 0.0 | 25.5 | #DIV/0! | 6.71E-03 |
| Q9EQV6 | PCVEDYLSAILNR | 0,iTRAQ4plex114[AnyN-term]; | 0.0 | 5.0 | #DIV/0! | 6.75E-03 |
| Q9EQV6 | YNAELESGNQFVLYR | 4,Methyl_2H(3)[E]; | 0.0 | 5.0 | #DIV/0! | 6.75E-03 |
| Q9EQV6 | DETVFQAVDTALKK | 0,LG-anhydrolactam[AnyN-term]; | 0.0 | 11.0 | #DIV/0! | 6.75E-03 |
| Q9EQV6 | DETVFQAVDTALKK | 0,LG-anhydrolactam[AnyN-term]; | 0.0 | 11.0 | #DIV/0! | 6.75E-03 |
| P02767 | CNADPGLSALLSDHR | 1,Cys->His[C]; | 0.0 | 11.3 | #DIV/0! | 6.76E-03 |
| Q9EQV6 | NPTSFLGHYLHEVAR | 0,Dimethyl[AnyN-term](Ethyl[AnyN-term]); | 0.0 | 8.8 | #DIV/0! | 6.78E-03 |
| Q9EQV6 | CKVNSGAFPAPIEK | 1,Acetyl[C]; | 0.0 | 9.5 | #DIV/0! | 6.78E-03 |
| Q9EQV6 | CTSFQENPTSFLGHYLHEVAR | 6,Cation_Na[E]; | 0.0 | 16.2 | #DIV/0! | 6.80E-03 |
| Q9EQV6 | TSFLGHYLHEVAR | 2,MeMePhosphorothioate[S]; | 0.0 | 11.0 | #DIV/0! | 7.06E-03 |
| Q9EQV6 | KGTDFQLNQLQGK | 14,Cation_Na[AnyC-term]; | 0.0 | 3.8 | #DIV/0! | 7.06E-03 |
| Q9EQV6 | ADRDQYELLCLDNTR | 10,Crotonaldehyde[C]; | 0.0 | 3.8 | #DIV/0! | 7.06E-03 |
| Q9EQV6 | CKVNSGAFPAPIEK | 1,monomethylphosphothione[C]; | 0.0 | 14.3 | #DIV/0! | 7.07E-03 |
| Q9EQV6 | NLPPFQRPEAEAMCTSFQENPTSFLGHYLHEVAR | 14,MDCC[C]; | 0.0 | 3.0 | #DIV/0! | 7.08E-03 |
| Q9EQV6 | YYAVAVVKASDSSINWNNLK | 10,HexNAc(2)Sulf(1)[S]; | 0.0 | 1.5 | #DIV/0! | 7.08E-03 |
| Q9EQV6 | KMPNASPEELADMVAK | 3,Pro->Trp[P]; | 0.0 | 1.5 | #DIV/0! | 7.08E-03 |
| Q9EQV6 | HEGLHNHHTEK | 0,Dansyl[AnyN-term]; | 0.0 | 1.5 | #DIV/0! | 7.08E-03 |
| P02767 | RDGVLDSVTDQDSKDSTYSMSSTLSLTK | 10,Cation_Ca[II][D]; | 0.0 | 1.5 | #DIV/0! | 7.08E-03 |
| P06238 | HEGLHNHHTEK | 0,Dansyl[AnyN-term]; | 0.0 | 1.5 | #DIV/0! | 7.08E-03 |
| P06238 | RDGVLDSVTDQDSKDSTYSMSSTLSLTK | 10,Cation_Ca[II][D]; | 0.0 | 1.5 | #DIV/0! | 7.08E-03 |
| P07632 | PVQGVIHFEQK | 0,SPITC_13C(6)[AnyN-term]; | 0.0 | 3.3 | #DIV/0! | 7.12E-03 |
| Q9EQV6 | HPDYSVSLLLR | 0,Hex[AnyN-term]; | 0.0 | 14.0 | #DIV/0! | 7.13E-03 |
| Q9EQV6 | FQENPTSFLGHYLHEVAR | 0,Diisopropylphosphate[AnyN-term]; | 0.0 | 8.0 | #DIV/0! | 7.14E-03 |
| P02767 | VHQFFNVGLIQPGSVK | 6,Ammonia-loss[N]; | 0.0 | 10.3 | #DIV/0! | 7.19E-03 |
| P02767 | PDSVFLIKEDK | 0,Amidine[AnyN-term]; | 0.0 | 5.5 | #DIV/0! | 7.22E-03 |
| Q9EQV6 | LRDNYGELADCCAK | 11,Cys->Dha[C]; | 0.0 | 10.2 | #DIV/0! | 7.26E-03 |
| P02770 | SGLYFIRPLK | 0,CAF[AnyN-term]; | 0.0 | 7.8 | #DIV/0! | 7.29E-03 |
| P06238 | CVDQMSNGDR | 1,Carbamidomethyl[C];5,Oxidation[M]; | 0.0 | 25.5 | #DIV/0! | 7.34E-03 |
| P06238 | VSSVFIFPPKPK | 3,Sulfo[S]; | 0.0 | 2.7 | #DIV/0! | 7.44E-03 |
| Q9EQV6 | ACYGTVLAEFQPLVEEPK | 1,Ala->Asp[A]; | 0.0 | 27.5 | #DIV/0! | 7.54E-03 |
| Q68FP1 | GSEPDGFWEALGGK | 2,Dimethylphosphothione[S]; | 0.0 | 6.0 | #DIV/0! | 7.61E-03 |
| P55314 | CSDILNEIKDR | 1,Carbamidomethyl[C]; | 0.0 | 8.3 | #DIV/0! | 7.81E-03 |
| Q9EQV6 | DLKQEDFQLLCPDGTKK | 11,Carbamidomethyl[C]; | 0.0 | 9.8 | #DIV/0! | 7.91E-03 |
| Q9EQV6 | RPCFSALTVDETYVPKEFK | 3,Carbamidomethyl[C];11,Cation_Na[E]; | 0.0 | 25.5 | #DIV/0! | 8.08E-03 |
| Q9EQV6 | AADKDNCFATEGPNLVAR | 7,Dehydro[C]; | 0.0 | 12.5 | #DIV/0! | 8.09E-03 |
| P20762 | VVSTLHIQHQDWMSGK | 3,Methylamine[S]; | 0.0 | 7.3 | #DIV/0! | 8.09E-03 |
| Q9EQV6 | NPTSFLGHYLHEVAR | 0,Delta_H(2)C(2)[AnyN-term]; | 0.0 | 4.8 | #DIV/0! | 8.11E-03 |
| Q9EQV6 | HAVVARNGDGKEDLIWEILK | 7,Hep[N]; | 0.0 | 9.7 | #DIV/0! | 8.11E-03 |
| Q9EQV6 | FLGHYLHEVAR | 0,CarboxymethylDMAP[AnyN-term]; | 0.0 | 8.0 | #DIV/0! | 8.36E-03 |
| P55314 | CECICPAGFQGVACEVTNRK | 1,GG[C](Dicarbamidomethyl[C]);5,Carbamidomethyl[C];14,Carbamidomethyl[C]; | 0.0 | 8.2 | #DIV/0! | 8.40E-03 |
| Q9EQV6 | KPVTEFATCHLAQAPNHVVVSR | 9,monomethylphosphothione[C]; | 0.0 | 10.8 | #DIV/0! | 8.42E-03 |
| Q9EQV6 | YNEVLTQCCTESDKAACLTPK | 3,Cation_Na[E];8,GG[C](Dicarbamidomethyl[C]);17,Carbamidomethyl[C]; | 0.0 | 27.0 | #DIV/0! | 8.49E-03 |
| Q9EQV6 | PVTEFATCHLAQAPNHVVVSR | 0,Unknown_302[AnyN-term]; | 0.0 | 2.3 | #DIV/0! | 8.63E-03 |
| Q6IG01 | NKYEDEINKR | 0,Formyl[AnyN-term]; | 0.0 | 2.3 | #DIV/0! | 8.63E-03 |
| Q6P6Q2 | NKYEDEINKR | 0,Formyl[AnyN-term]; | 0.0 | 2.3 | #DIV/0! | 8.63E-03 |
| Q10758 | NKYEDEINKR | 0,Formyl[AnyN-term]; | 0.0 | 2.3 | #DIV/0! | 8.63E-03 |
| P26644 | VCPFAGILENGVVR | 2,Carbamidomethyl[C]; | 0.0 | 16.2 | #DIV/0! | 8.81E-03 |
| P02767 | ECQSEGVLFFQGNRK | 0,TMAB_2H(9)[AnyN-term]; | 0.0 | 10.7 | #DIV/0! | 8.83E-03 |
| P04639 | GAPISQGVDTANPTK | 5,AEBS[S]; | 0.0 | 8.5 | #DIV/0! | 8.86E-03 |
| Q9EQV6 | SSTKDLLFRDDTK | 0,Carbamyl[AnyN-term]; | 0.0 | 17.8 | #DIV/0! | 8.87E-03 |
| P02767 | QQVEASLQPETLR | 0,CAF[AnyN-term]; | 0.0 | 14.3 | #DIV/0! | 8.88E-03 |
| P02767 | ECHRGECQSEGVLFFQGNR | 7,MDCC[C]; | 0.0 | 14.3 | #DIV/0! | 8.88E-03 |
| Q9EQV6 | FYAPELLYYAEK | 2,Didehydro[Y]; | 0.0 | 8.3 | #DIV/0! | 8.96E-03 |
| Q9EQV6 | CHLAQAPNHVVVSR | 1,Palmitoyl[C]; | 0.0 | 17.3 | #DIV/0! | 8.98E-03 |
| Q9EQV6 | TCVVVDISQNDPEVR | 2,Hydroxycinnamyl[C]; | 0.0 | 12.5 | #DIV/0! | 9.00E-03 |
| Q9EQV6 | KQTALAELVKHKPK | 1,Malonyl[K]; | 0.0 | 3.5 | #DIV/0! | 9.05E-03 |
| P16391 | EGPEYWEQQTR | 8,Deamidated[Q]; | 0.0 | 3.5 | #DIV/0! | 9.05E-03 |
| Q6IMF3 | FLEQQNQVLQTK | 2,Xle->Pro[L]; | 0.0 | 6.2 | #DIV/0! | 9.07E-03 |
| Q6IG03 | FLEQQNQVLQTK | 2,Xle->Pro[L]; | 0.0 | 6.2 | #DIV/0! | 9.07E-03 |
| Q6IG01 | FLEQQNQVLQTK | 2,Xle->Pro[L]; | 0.0 | 6.2 | #DIV/0! | 9.07E-03 |
| Q9EQV6 | MSQRFPNAEFAEITK | 2,O-Isopropylmethylphosphonate[S]; | 0.0 | 18.3 | #DIV/0! | 9.09E-03 |
| P22283 | LVNCPFEEQTEQLKR | 3,Deamidated[N];4,Carbamidomethyl[C]; | 0.0 | 4.3 | #DIV/0! | 9.26E-03 |
| Q9EQV6 | PERNECFLQHKDDNPNLPPFQRPEAEAMCTSFQENPTSFLGHYLHEVAR | 6,4AcAllylGal[C]; | 0.0 | 3.0 | #DIV/0! | 9.28E-03 |
| Q9EQV6 | DLKQEDFQLLCPDGTKKPVTEFATCHLAQAPNHVVVSR | 25,DTT[C]; | 0.0 | 6.5 | #DIV/0! | 9.38E-03 |
| Q9EQV6 | ECCHGDLLECADDRAELAK | 1,Cation_Na[E];2,GG[C](Dicarbamidomethyl[C]);10,Carbamidomethyl[C]; | 0.0 | 12.5 | #DIV/0! | 9.40E-03 |
| Q9EQV6 | CKAADKDNCFATEGPNLVAR | 0,Formyl[AnyN-term]; | 0.0 | 11.3 | #DIV/0! | 9.69E-03 |
| Q9EQV6 | DLKQEDFQLLCPDGTKKPVTEFATCHLAQAPNHVVVSR | 11,DTT[C]; | 0.0 | 10.7 | #DIV/0! | 9.75E-03 |
| P07647 | VGSICLASGWGMTNPSEMK | 5,Carbamidomethyl[C];12,Oxidation[M]; | 0.0 | 12.8 | #DIV/0! | 9.80E-03 |
| Q9EQV6 | SFLGHYLHEVAR | 1,Didehydro[S]; | 0.0 | 5.2 | #DIV/0! | 9.83E-03 |
| Q9EQV6 | PEAEAMCTSFQENPTSFLGHYLHEVAR | 6,Met->Aha[M]; | 0.0 | 16.7 | #DIV/0! | 9.87E-03 |
| P00762 | LGEHNINVLEGDEQFINAAK | 10,Cation_Na[E];13,Cation_Na[E]; | 0.0 | 5.5 | #DIV/0! | 9.89E-03 |
| Q9EQV6 | AEAMCTSFQENPTSFLGHYLHEVAR | 0,Brij35[AnyN-term]; | 0.0 | 12.3 | #DIV/0! | 9.90E-03 |
| Q9EQV6 | LDAVKEKALVAAVR | 5,Malonyl[K]; | 0.0 | 10.7 | #DIV/0! | 9.94E-03 |
| P06238 | LVKPYVQDHFTEK | 1,Carbonyl[L]; | 0.0 | 7.3 | #DIV/0! | 1.00E-02 |
| Q9EQV6 | MSQRFPNAEFAEITK | 0,Phenylisocyanate_2H(5)[AnyN-term];7,Deamidated[N]; | 0.0 | 16.2 | #DIV/0! | 1.00E-02 |
| Q9EQV6 | NCFATEGPNLVAR | 1,Deamidated[N]; | 0.0 | 2.8 | #DIV/0! | 1.00E-02 |
| Q9EQV6 | LDAVKEKALVAAVR | 0,glycidamide[AnyN-term]; | 0.0 | 17.3 | #DIV/0! | 1.00E-02 |
| Q9EQV6 | RHPYFYAPELLYYAEK | 8,Pro->Lys[P]; | 0.0 | 14.0 | #DIV/0! | 1.01E-02 |
| Q9EQV6 | HQTVLENTNGKNTAAWAK | 11,Carbamidomethyl[K](Gly[K]); | 0.0 | 32.5 | #DIV/0! | 1.01E-02 |
| P06238 | ALGCLEASWETIEQGR | 4,Carbamidomethyl[C]; | 0.0 | 6.8 | #DIV/0! | 1.02E-02 |
| P06238 | ALGCLEASWETIEQGR | 4,Carbamidomethyl[C]; | 0.0 | 6.8 | #DIV/0! | 1.02E-02 |
| P02767 | GAQATWAELSWPHEK | 0,dNIC[AnyN-term]; | 0.0 | 13.5 | #DIV/0! | 1.02E-02 |
| Q9EQV6 | EAEAMCTSFQENPTSFLGHYLHEVAR | 0,Biotin_Thermo-21330[AnyN-term]; | 0.0 | 9.2 | #DIV/0! | 1.02E-02 |
| Q9EQV6 | VFMQHIDVLENSLGFK | 1,Val->Asn[V]; | 0.0 | 1.3 | #DIV/0! | 1.03E-02 |
| Q9EQV6 | NDADLSGITEDAPLK | 0,ICPL_13C(6)2H(4)[AnyN-term]; | 0.0 | 5.3 | #DIV/0! | 1.03E-02 |
| P06238 | EESSCIHSSCTAER | 2,Cation_Na[E];5,Carbamidomethyl[C];10,Carbamidomethyl[C]; | 0.0 | 1.3 | #DIV/0! | 1.03E-02 |
| P06238 | EESSCIHSSCTAER | 2,Cation_Na[E];5,Carbamidomethyl[C];10,Carbamidomethyl[C]; | 0.0 | 1.3 | #DIV/0! | 1.03E-02 |
| P04639 | EKGVELSDVKR | 1,Glu->His[E]; | 0.0 | 1.3 | #DIV/0! | 1.03E-02 |
| Q63416 | YETHGDVFIKPQLWPLTDCCES | 19,Hydroxycinnamyl[C]; | 0.0 | 1.3 | #DIV/0! | 1.03E-02 |
| P06238 | ICTGSEVSSVFIFPPKPK | 0,SPITC[AnyN-term]; | 0.0 | 3.2 | #DIV/0! | 1.04E-02 |
| Q9EQV6 | EGAQEMDCNDETVFQAVDTALKK | 0,Triton[AnyN-term]; | 0.0 | 3.7 | #DIV/0! | 1.04E-02 |
| Q9EQV6 | FTFHSDICTLPDKEK | 8,GG[C](Dicarbamidomethyl[C]); | 0.0 | 5.2 | #DIV/0! | 1.06E-02 |
| Q9EQV6 | TVMGDFAQFVDK | 1,Thr->Cys[T]; | 0.0 | 16.7 | #DIV/0! | 1.07E-02 |
| Q9EQV6 | AADKDNCFATEGPNLVAR | 7,Ub-Br2[C]; | 0.0 | 35.8 | #DIV/0! | 1.08E-02 |
| P02767 | RDGVLDSVTDQDSKDSTYSMSSTLSLTK | 12,Cation_Ca[II][D]; | 0.0 | 1.7 | #DIV/0! | 1.08E-02 |
| P06238 | RDGVLDSVTDQDSKDSTYSMSSTLSLTK | 12,Cation_Ca[II][D]; | 0.0 | 1.7 | #DIV/0! | 1.08E-02 |
| Q9EQV6 | FAKLCEAHGITR | 5,GG[C](Dicarbamidomethyl[C]); | 0.0 | 8.0 | #DIV/0! | 1.10E-02 |
| Q9EQV6 | VQEVTDFAK | 0,Succinyl_2H(4)[AnyN-term]; | 0.0 | 10.3 | #DIV/0! | 1.11E-02 |
| Q9EQV6 | SFLGHYLHEVAR | 0,Biotin_Thermo-21328[AnyN-term]; | 0.0 | 7.7 | #DIV/0! | 1.13E-02 |
| P00762 | LGEHNINVLEGDEQFINAAK | 12,Dehydrated[D]; | 0.0 | 3.8 | #DIV/0! | 1.13E-02 |
| P02767 | HNLYTCEVVHK | 6,Carbamidomethyl[C];7,Cation_Na[E]; | 0.0 | 5.8 | #DIV/0! | 1.13E-02 |
| P06238 | HNLYTCEVVHK | 6,Carbamidomethyl[C];7,Cation_Na[E]; | 0.0 | 5.8 | #DIV/0! | 1.13E-02 |
| Q9EQV6 | CFATEGPNLVAR | 2,Fluoro[F]; | 0.0 | 6.5 | #DIV/0! | 1.15E-02 |
| P06238 | NVDKEDEDFQESNK | 1,Asn->Gly[N]; | 0.0 | 11.2 | #DIV/0! | 1.15E-02 |
| P26644 | VPELLGGPSVFIFPPKPK | 0,TNBS[AnyN-term]; | 0.0 | 4.3 | #DIV/0! | 1.16E-02 |
| Q9EQV6 | LVQEVTDFAK | 1,Xle->Thr[L]; | 0.0 | 9.8 | #DIV/0! | 1.17E-02 |
| Q9EQV6 | HQTVLENTNGKNTAAWAK | 6,Glu->Pro[E]; | 0.0 | 3.0 | #DIV/0! | 1.17E-02 |
| P06238 | SQICELVAHETISFLMK | 4,Carbamidomethyl[C];5,Cation_Na[E];16,Oxidation[M]; | 0.0 | 13.8 | #DIV/0! | 1.17E-02 |
| Q9EQV6 | FPSSTFEQVSQLVK | 0,Dicarbamidomethyl[AnyN-term]; | 0.0 | 28.3 | #DIV/0! | 1.18E-02 |
| P02767 | FNPVTGEVPPR | 0,Methyl_2H(3)13C(1)[AnyN-term]; | 0.0 | 4.2 | #DIV/0! | 1.18E-02 |
| Q9EQV6 | NPTSFLGHYLHEVAR | 1,Hydroxymethyl[N]; | 0.0 | 19.7 | #DIV/0! | 1.19E-02 |
| Q9EQV6 | YNEVLTQCCTESDKAACLTPK | 3,Cation_Na[E];9,GG[C](Dicarbamidomethyl[C]);17,Carbamidomethyl[C]; | 0.0 | 28.8 | #DIV/0! | 1.19E-02 |
| Q9EQV6 | LCMAALSHQPQEFPAYVEPTNDEICEAFR | 2,Carbamidomethyl[C];3,Oxidation[M];25,Carbamidomethyl[C]; | 0.0 | 1.8 | #DIV/0! | 1.21E-02 |
| P06238 | ASQICELVAHETISFLMK | 4,Xle->Met[I]; | 0.0 | 7.2 | #DIV/0! | 1.22E-02 |
| Q9EQV6 | SQCLAEIEHDNIPADLPSIAADFVEDK | 3,GG[C](Dicarbamidomethyl[C]);11,Deamidated[N]; | 0.0 | 15.8 | #DIV/0! | 1.23E-02 |
| P02767 | DSTYSMSSTLSLTK | 0,ISD_z+2_ion[AnyN-term]; | 0.0 | 12.0 | #DIV/0! | 1.25E-02 |
| P06238 | DSTYSMSSTLSLTK | 0,ISD_z+2_ion[AnyN-term]; | 0.0 | 12.0 | #DIV/0! | 1.25E-02 |
| Q68FP1 | VMPKAGALNSNDAFVLK | 0,SPITC_13C(6)[AnyN-term]; | 0.0 | 12.0 | #DIV/0! | 1.25E-02 |
| Q9EQV6 | TSFQENPTSFLGHYLHEVAR | 0,Unknown_248[AnyN-term]; | 0.0 | 6.3 | #DIV/0! | 1.26E-02 |
| Q9EQV6 | FLEEVKNNYHSEAFSVNFADSEEAKK | 4,Cation_Na[E];7,Deamidated[N]; | 0.0 | 3.2 | #DIV/0! | 1.26E-02 |
| Q9EQV6 | FQENPTSFLGHYLHEVAR | 4,Asn->Tyr[N]; | 0.0 | 3.5 | #DIV/0! | 1.27E-02 |
| Q9EQV6 | APQVSTPTLVEAARNLGRVGTKCCTLPEAQRLPCVEDYLSAILNR | 23,Iodoacetanilide[C]; | 0.0 | 1.2 | #DIV/0! | 1.27E-02 |
| Q9EQV6 | LRDNYGELADCCAK | 12,NO_SMX_SIMD[C]; | 0.0 | 7.0 | #DIV/0! | 1.27E-02 |
| Q9EQV6 | DGAETLYSFK | 0,Iodoacetanilide[AnyN-term]; | 0.0 | 1.2 | #DIV/0! | 1.27E-02 |
| Q9EQV6 | DGAETLYSFK | 0,Iodoacetanilide[AnyN-term]; | 0.0 | 1.2 | #DIV/0! | 1.27E-02 |
| Q9EQV6 | HLAQAPNHVVVSR | 0,Propionyl_13C(3)[AnyN-term]; | 0.0 | 2.3 | #DIV/0! | 1.27E-02 |
| Q9EQV6 | YHSEAFSVNFADSEEAKK | 1,Tyr->Pro[Y]; | 0.0 | 2.3 | #DIV/0! | 1.27E-02 |
| P02767 | NADPGLSALLSDHR | 0,CAF[AnyN-term]; | 0.0 | 1.2 | #DIV/0! | 1.27E-02 |
| P06238 | HAEEHHTAYAVYSLSK | 16,Lys->CamCys[K]; | 0.0 | 4.7 | #DIV/0! | 1.27E-02 |
| Q63416 | EKIIRHPEYNKDTLDNDIMLIK | 13,dHex(1)Hex(1)[T]; | 0.0 | 1.2 | #DIV/0! | 1.27E-02 |
| Q9EQV6 | FKDLGEQHFK | 1,Phe->Lys[F]; | 0.0 | 2.8 | #DIV/0! | 1.29E-02 |
| P27274 | QVYQQCWR | 6,Carbamidomethyl[C]; | 0.0 | 5.7 | #DIV/0! | 1.29E-02 |
| P06238 | TFCSEPEKVDKDNEDFQESNR | 3,Delta_H(6)C(3)O(1)[C]; | 0.0 | 3.8 | #DIV/0! | 1.29E-02 |
| P06238 | SAYINTKDSPVLVDSFEDSEPYR | 3,AEBS[Y]; | 0.0 | 3.8 | #DIV/0! | 1.29E-02 |
| Q9EQV6 | PSVLEAATESLAK | 2,Hex(2)[S]; | 0.0 | 7.8 | #DIV/0! | 1.29E-02 |
| P02767 | WFWDFATR | 0,Methyl[AnyN-term]; | 0.0 | 3.3 | #DIV/0! | 1.29E-02 |
| Q9EQV6 | HPYFYAPELLYYAEK | 0,Diethylphosphate[AnyN-term]; | 0.0 | 16.0 | #DIV/0! | 1.30E-02 |
| P06911 | KLYSRSLDDNGEALYNFR | 3,sulfo+amino[Y]; | 0.0 | 9.7 | #DIV/0! | 1.30E-02 |
| Q9EQV6 | KMPNASPEELADMVAK | 1,Xlink_DTSSP[88][K]; | 0.0 | 16.8 | #DIV/0! | 1.31E-02 |
| Q9EQV6 | TCVADENAENCDKSIHTLFGDK | 2,Cys->CamSec[C]; | 0.0 | 4.0 | #DIV/0! | 1.32E-02 |
| Q9EQV6 | PTSFLGHYLHEVAR | 1,Pro->His[P]; | 0.0 | 12.2 | #DIV/0! | 1.32E-02 |
| Q9EQV6 | CADYSENTFTEYKK | 0,Sulfo-NHS-LC-LC-Biotin[AnyN-term]; | 0.0 | 13.0 | #DIV/0! | 1.33E-02 |
| Q9EQV6 | DGGGDVAFVK | 5,Cation_Na[D]; | 0.0 | 7.3 | #DIV/0! | 1.34E-02 |
| Q9EQV6 | QEPERNECFLQHKDDNPNLPPFQRPEAEAMCTSFQENPTSFLGHYLHEVAR | 27,Ala->Trp[A]; | 0.0 | 9.2 | #DIV/0! | 1.35E-02 |
| Q9EQV6 | AADKDNCFATEGPNLVAR | 7,Carboxymethyl[C]; | 0.0 | 2.7 | #DIV/0! | 1.36E-02 |
| Q9EQV6 | DKDNCFATEGPNLVAR | 5,NEM_2H(5)[C]; | 0.0 | 7.2 | #DIV/0! | 1.36E-02 |
| Q9EQV6 | CYGTVLAEFQPLVEEPK | 0,Formyl[AnyN-term]; | 0.0 | 2.5 | #DIV/0! | 1.36E-02 |
| Q9EQV6 | AETFTFHSDICTLPDKEKQIK | 11,BEMAD_C_2H(6)[C]; | 0.0 | 3.7 | #DIV/0! | 1.39E-02 |
| Q9WUW3 | EYCGVPGDGYEELTR | 1,Cation_Na[E];3,Carbamidomethyl[C]; | 0.0 | 15.3 | #DIV/0! | 1.39E-02 |
| Q9EQV6 | GEQHFKGLVLIAFSQYLQK | 8,Xle->Cys[L]; | 0.0 | 4.3 | #DIV/0! | 1.41E-02 |
| Q9EQV6 | CFATEGPNLVAR | 1,2-succinyl[C]; | 0.0 | 4.3 | #DIV/0! | 1.41E-02 |
| Q9EQV6 | PDGTKKPVTEFATCHLAQAPNHVVVSR | 13,PhosphoCytidine[T]; | 0.0 | 22.3 | #DIV/0! | 1.43E-02 |
| Q9EQV6 | NPTSFLGHYLHEVAR | 3,Ethanedithiol[T]; | 0.0 | 15.5 | #DIV/0! | 1.44E-02 |
| P06238 | ETFTYEWTVPK | 6,Cation_Na[E]; | 0.0 | 5.7 | #DIV/0! | 1.44E-02 |
| Q9EQV6 | DNPNLPPFQRPEAEAMCTSFQENPTSFLGHYLHEVAR | 17,Delta_H(6)C(3)O(1)[C]; | 0.0 | 3.0 | #DIV/0! | 1.44E-02 |
| Q9EQV6 | IQHLPEQEEPEDSKGK | 0,GIST-Quat_2H(3)[AnyN-term]; | 0.0 | 3.0 | #DIV/0! | 1.44E-02 |
| Q9EQV6 | CFATEGPNLVAR | 1,IED-Biotin[C]; | 0.0 | 13.5 | #DIV/0! | 1.44E-02 |
| Q9EQV6 | KAETFTFHSDICTLPDKEK | 12,IBTP[C]; | 0.0 | 26.3 | #DIV/0! | 1.45E-02 |
| Q9EQV6 | NPTSFLGHYLHEVAR | 4,Sulfo[S]; | 0.0 | 7.7 | #DIV/0! | 1.45E-02 |
| Q9EQV6 | DAEEAATGECTTTLGK | 4,Cation_Na[E];10,Carbamidomethyl[C]; | 0.0 | 3.5 | #DIV/0! | 1.47E-02 |
| Q9EQV6 | DAEEAATGECTTTLGK | 4,Cation_Na[E];10,Carbamidomethyl[C]; | 0.0 | 3.5 | #DIV/0! | 1.47E-02 |
| P02767 | LSRPATITDNVIPACLPSPNYVVADR | 9,Propargylamine[D]; | 0.0 | 3.5 | #DIV/0! | 1.47E-02 |
| Q9EQV6 | RHPYFYAPELLYYAEK | 5,Phe->Glu[F]; | 0.0 | 15.5 | #DIV/0! | 1.47E-02 |
| Q9EQV6 | TCHLAQAPNHVVVSR | 1,Thr->Cys[T]; | 0.0 | 12.8 | #DIV/0! | 1.47E-02 |
| Q9EQV6 | LVQEVTDFAK | 4,Cation_Na[E]; | 0.0 | 34.2 | #DIV/0! | 1.49E-02 |
| Q9EQV6 | AETFTFHSDICTLPDKEK | 2,Cation_Na[E];11,Carbamidomethyl[C]; | 0.0 | 13.0 | #DIV/0! | 1.50E-02 |
| Q9EQV6 | VCPEGSIDSAPVK | 2,lapachenole[C]; | 0.0 | 5.7 | #DIV/0! | 1.51E-02 |
| Q9EQV6 | YNAELESGNQFVLYR | 0,Diethylphosphate[AnyN-term]; | 0.0 | 7.3 | #DIV/0! | 1.53E-02 |
| P02767 | QYGFCK | 5,Carbamidomethyl[C]; | 0.0 | 5.8 | #DIV/0! | 1.56E-02 |
| Q9EQV6 | CPDGTKKPVTEFATCHLAQAPNHVVVSR | 1,Delta_Hg(1)[C]; | 0.0 | 6.3 | #DIV/0! | 1.57E-02 |
| P02767 | SGAQATWAELSWPHEKVDGALCLEK | 22,Carbamidomethyl[C]; | 0.0 | 34.8 | #DIV/0! | 1.58E-02 |
| Q9EQV6 | PCFSALTVDETYVPK | 3,Phe->Asp[F]; | 0.0 | 8.5 | #DIV/0! | 1.59E-02 |
| Q9EQV6 | ATEDQLKTVMGDFAQFVDK | 0,CAF[AnyN-term]; | 0.0 | 2.8 | #DIV/0! | 1.59E-02 |
| Q9EQV6 | TCVADENAENCDKSIHTLFGDK | 2,DiDehydro[C]; | 0.0 | 2.8 | #DIV/0! | 1.59E-02 |
| Q9EQV6 | DVEVHTAQTHAPEK | 0,Propyl[AnyN-term]; | 0.0 | 2.8 | #DIV/0! | 1.59E-02 |
| P26644 | CPTCPTCHKCPVPELLGGPSVFIFPPKPK | 1,Iodoacetanilide[C]; | 0.0 | 2.8 | #DIV/0! | 1.59E-02 |
| Q9EQV6 | VFLGTFLYEYSR | 0,dNIC[AnyN-term]; | 0.0 | 4.8 | #DIV/0! | 1.60E-02 |
| P02767 | CRQFYKLQTEGDGIYTLNSEK | 0,Methyl_2H(2)[AnyN-term]; | 0.0 | 4.2 | #DIV/0! | 1.60E-02 |
| Q63416 | KIIRHPEYNKDTLDNDIMLIK | 12,FMN[T]; | 0.0 | 4.2 | #DIV/0! | 1.60E-02 |
| P02767 | ALYQAEAFTADFQQSR | 2,Xle->Ser[L]; | 0.0 | 4.7 | #DIV/0! | 1.61E-02 |
| Q9EQV6 | YNAELESGNQFVLYR | 5,Xle->Glu[L]; | 0.0 | 9.7 | #DIV/0! | 1.62E-02 |
| Q9EQV6 | CPYEEHIKLVQEVTDFAK | 0,Succinyl_2H(4)[AnyN-term]; | 0.0 | 4.0 | #DIV/0! | 1.63E-02 |
| Q9EQV6 | VLAEFQPLVEEPK | 0,Cytopiloyne+water[AnyN-term]; | 0.0 | 8.0 | #DIV/0! | 1.63E-02 |
| Q9EQV6 | TALAELVKHKPK | 0,Unknown_302[AnyN-term]; | 0.0 | 4.0 | #DIV/0! | 1.63E-02 |
| Q9EQV6 | LAEDAQETDTSQQDQSPTYR | 3,phosphoRibosyl[E]; | 0.0 | 4.0 | #DIV/0! | 1.63E-02 |
| P08723 | KEFLLEDGEHVTQVSGTR | 0,3sulfo[AnyN-term]; | 0.0 | 11.7 | #DIV/0! | 1.63E-02 |
| P06238 | EYTDDSFTNRK | 0,ISD_z+2_ion[AnyN-term]; | 0.0 | 6.3 | #DIV/0! | 1.63E-02 |
| Q9EQV6 | NNYHSEAFSVNFADSEEAKK | 16,Cation_Na[E]; | 0.0 | 8.8 | #DIV/0! | 1.66E-02 |
| Q9EQV6 | MPNASPEELADMVAK | 3,Deamidated[N]; | 0.0 | 29.2 | #DIV/0! | 1.68E-02 |
| Q9EQV6 | EGYNGYTGAFQCLVEK | 3,Dehydrated[Y];4,Deamidated[N];12,Carbamidomethyl[C]; | 0.0 | 22.5 | #DIV/0! | 1.69E-02 |
| Q9EQV6 | TKDLSSDIK | 0,Carbamyl[AnyN-term]; | 0.0 | 7.8 | #DIV/0! | 1.70E-02 |
| P02767 | VVPEGMR | 6,Oxidation[M]; | 0.0 | 5.2 | #DIV/0! | 1.70E-02 |
| P26644 | IEKNYKNTEPVMDSDGSFFMYSK | 4,Hep[N]; | 0.0 | 2.7 | #DIV/0! | 1.71E-02 |
| P02767 | GAVSPVGVQPILNK | 0,Carbamyl[AnyN-term]; | 0.0 | 8.7 | #DIV/0! | 1.72E-02 |
| Q9EQV6 | SQRFPNAEFAEITK | 0,Dimethyl_2H(4)[AnyN-term]; | 0.0 | 4.5 | #DIV/0! | 1.72E-02 |
| Q9EQV6 | KCCTLPEAQR | 3,GG[C](Dicarbamidomethyl[C]); | 0.0 | 1.5 | #DIV/0! | 1.72E-02 |
| Q9EQV6 | YLQKCPYEEHIKLVQEVTDFAK | 5,IED-Biotin[C]; | 0.0 | 1.5 | #DIV/0! | 1.72E-02 |
| Q9EQV6 | VFMQHIDVLENSLGFK | 0,Methyl[AnyN-term]; | 0.0 | 3.0 | #DIV/0! | 1.72E-02 |
| Q9EQV6 | VFNNDADLSGITEDAPLK | 5,Cation_Na[D]; | 0.0 | 1.5 | #DIV/0! | 1.72E-02 |
| P02767 | DGVLDSVTDQDSK | 0,Methyl_2H(2)[AnyN-term]; | 0.0 | 1.5 | #DIV/0! | 1.72E-02 |
| P06238 | DGVLDSVTDQDSK | 0,Methyl_2H(2)[AnyN-term]; | 0.0 | 1.5 | #DIV/0! | 1.72E-02 |
| P02767 | IFSQQADLSR | 0,dNIC[AnyN-term]; | 0.0 | 2.3 | #DIV/0! | 1.73E-02 |
| Q9WUW3 | AFAELWAFDAAQGK | 0,C+12[AnyN-term]; | 0.0 | 2.3 | #DIV/0! | 1.73E-02 |
| Q9EQV6 | TSFLGHYLHEVAR | 0,Ethylphosphate[AnyN-term]; | 0.0 | 3.3 | #DIV/0! | 1.74E-02 |
| Q9EQV6 | RRHPYFYAPELLYYAEK | 6,Phe->Trp[F]; | 0.0 | 3.3 | #DIV/0! | 1.74E-02 |
| P02767 | TAESGELHGLTTDEKFTEGVYR | 2,Carbonyl[A]; | 0.0 | 4.2 | #DIV/0! | 1.75E-02 |
| Q9EQV6 | LRDNYGELADCCAK | 12,DiDehydro[C]; | 0.0 | 7.7 | #DIV/0! | 1.76E-02 |
| Q9EQV6 | GDFAQFVDK | 0,CLIP_TRAQ_2[AnyN-term]; | 0.0 | 3.2 | #DIV/0! | 1.76E-02 |
| Q9EQV6 | QNDPEVRFSWFIDDVEVHTAQTHAPEK | 0,C+12[AnyN-term]; | 0.0 | 3.2 | #DIV/0! | 1.76E-02 |
| P06238 | MLIYTILPDGEVIADSVK | 3,Xle->Ser[I]; | 0.0 | 3.2 | #DIV/0! | 1.76E-02 |
| P06238 | MLIYTILPDGEVIADSVK | 3,Xle->Ser[I]; | 0.0 | 3.2 | #DIV/0! | 1.76E-02 |
| Q9EQV6 | DNPNLPPFQRPEAEAMCTSFQENPTSFLGHYLHEVAR | 17,Carboxymethyl[C]; | 0.0 | 2.5 | #DIV/0! | 1.77E-02 |
| P23680 | LKDSFLADVK | 1,Xle->Asp[L]; | 0.0 | 2.5 | #DIV/0! | 1.77E-02 |
| Q9EQV6 | MCTSFQENPTSFLGHYLHEVAR | 0,CarboxymethylDMAP[AnyN-term]; | 0.0 | 7.0 | #DIV/0! | 1.78E-02 |
| Q9EQV6 | RLPCVEDYLSAILNR | 0,Biotin_Thermo-21328[AnyN-term]; | 0.0 | 7.0 | #DIV/0! | 1.78E-02 |
| Q9EQV6 | APQVSTPTLVEAAR | 1,Ala->Asp[A]; | 0.0 | 26.3 | #DIV/0! | 1.79E-02 |
| Q9EQV6 | CTSFQENPTSFLGHYLHEVAR | 1,Propionamide_2H(3)[C]; | 0.0 | 2.0 | #DIV/0! | 1.80E-02 |
| Q9EQV6 | PEAEAMCTSFQENPTSFLGHYLHEVAR | 6,Oxidation[M]; | 0.0 | 5.3 | #DIV/0! | 1.81E-02 |
| Q9EQV6 | VFNNDADLSGITEDAPLKLSQAVHK | 0,Carbamyl[AnyN-term]; | 0.0 | 5.3 | #DIV/0! | 1.81E-02 |
| Q9EQV6 | EQHFKGLVLIAFSQYLQK | 0,Methylthio[AnyN-term]; | 0.0 | 4.3 | #DIV/0! | 1.82E-02 |
| Q9EQV6 | HLAQAPNHVVVSR | 0,TNBS[AnyN-term]; | 0.0 | 7.7 | #DIV/0! | 1.85E-02 |
| Q9EQV6 | DNCFATEGPNLVAR | 3,MDCC[C]; | 0.0 | 13.0 | #DIV/0! | 1.86E-02 |
| Q9EQV6 | CLRDGGGDVAFVK | 1,Cys->CamSec[C]; | 0.0 | 5.8 | #DIV/0! | 1.87E-02 |
| Q9EQV6 | YMCENQATISSK | 2,Oxidation[M];3,GG[C](Dicarbamidomethyl[C]); | 0.0 | 6.0 | #DIV/0! | 1.87E-02 |
| Q9EQV6 | DDNPNLPPFQRPEAEAMCTSFQENPTSFLGHYLHEVAR | 5,Deamidated[N];18,Carbamidomethyl[C]; | 0.0 | 9.2 | #DIV/0! | 1.87E-02 |
| P04639 | DHIIDDVISFIR | 1,Asp->Pro[D]; | 0.0 | 5.0 | #DIV/0! | 1.88E-02 |
| Q9EQV6 | TCHLAQAPNHVVVSR | 0,Propyl[AnyN-term]; | 0.0 | 10.7 | #DIV/0! | 1.88E-02 |
| Q9EQV6 | SDSSINWNNLKGK | 0,Ethylphosphate[AnyN-term]; | 0.0 | 5.5 | #DIV/0! | 1.88E-02 |
| Q9EQV6 | FKDLGEQHFK | 0,Methyl[AnyN-term]; | 0.0 | 10.8 | #DIV/0! | 1.88E-02 |
| P02767 | LQVLAEFQEK | 9,Cation_Na[E]; | 0.0 | 3.5 | #DIV/0! | 1.89E-02 |
| P02770 | LQVLAEFQEK | 9,Cation_Na[E]; | 0.0 | 3.5 | #DIV/0! | 1.89E-02 |
| Q9EQV6 | YHSEAFSVNFADSEEAK | 0,NP40[AnyN-term]; | 0.0 | 7.7 | #DIV/0! | 1.90E-02 |
| P02767 | QWVNPAAGDKLPK | 1,Gln->Pro[Q]; | 0.0 | 5.2 | #DIV/0! | 1.90E-02 |
| Q9EQV6 | DDNPNLPPFQRPEAEAMCTSFQENPTSFLGHYLHEVAR | 3,Deamidated[N];18,Carbamidomethyl[C]; | 0.0 | 9.3 | #DIV/0! | 1.91E-02 |
| P06238 | QCSHALSPTNCDASKPLAEK | 10,Deamidated[N];11,Carbamidomethyl[C]; | 0.0 | 2.8 | #DIV/0! | 1.93E-02 |
| P07632 | ISLSGEHSIIGR | 0,Phenylisocyanate_2H(5)[AnyN-term]; | 0.0 | 4.8 | #DIV/0! | 1.94E-02 |
| Q9EQV6 | CYGTVLAEFQPLVEEPK | 1,Cys->Met[C]; | 0.0 | 39.8 | #DIV/0! | 1.96E-02 |
| Q9EQV6 | VEVHTAQTHAPEK | 0,SPITC_13C(6)[AnyN-term]; | 0.0 | 8.2 | #DIV/0! | 1.96E-02 |
| Q9EQV6 | GLVLIAFSQYLQK | 10,Dehydrated[Y]; | 0.0 | 7.2 | #DIV/0! | 1.97E-02 |
| Q9EQV6 | QEPERNECFLQHKDDNPNLPPFQRPEAEAMCTSFQENPTSFLGHYLHEVAR | 0,ICPL_13C(6)2H(4)[AnyN-term]; | 0.0 | 1.8 | #DIV/0! | 1.97E-02 |
| Q9EQV6 | REAHKSEIAHR | 0,Ethylphosphate[AnyN-term]; | 0.0 | 1.8 | #DIV/0! | 1.97E-02 |
| Q9EQV6 | KKPVTEFATCHLAQAPNHVVVSR | 11,His->Trp[H]; | 0.0 | 1.8 | #DIV/0! | 1.97E-02 |
| P06238 | DTTYALPSVSALVSALR | 1,Cation_Fe[III][D]; | 0.0 | 1.8 | #DIV/0! | 1.97E-02 |
| P08934 | ESNAELTADCETKR | 10,Carbamidomethyl[C]; | 0.0 | 1.8 | #DIV/0! | 1.97E-02 |
| Q63416 | HPEYNKDTLDNDIMLIK | 0,Unknown_248[AnyN-term]; | 0.0 | 1.8 | #DIV/0! | 1.97E-02 |
| Q9EQV6 | SFQENPTSFLGHYLHEVAR | 0,Phenylisocyanate_2H(5)[AnyN-term]; | 0.0 | 4.7 | #DIV/0! | 1.98E-02 |
| Q9EQV6 | CPDGTKKPVTEFATCHLAQAPNHVVVSR | 15,Delta_Hg(1)[C]; | 0.0 | 4.7 | #DIV/0! | 1.98E-02 |
| Q9EQV6 | TVMGDFAQFVDK | 2,Val->Glu[V]; | 0.0 | 13.0 | #DIV/0! | 1.99E-02 |
| Q9EQV6 | YMCENQATISSK | 0,Phenylisocyanate_2H(5)[AnyN-term];2,Oxidation[M];3,Carbamidomethyl[C]; | 0.0 | 1.7 | #DIV/0! | 1.99E-02 |
| Q9EQV6 | QENPTSFLGHYLHEVAR | 5,Diethylphosphothione[T]; | 0.0 | 3.3 | #DIV/0! | 1.99E-02 |
| Q9EQV6 | CPYEEHIK | 1,maleimide[C]; | 0.0 | 3.3 | #DIV/0! | 1.99E-02 |
| Q9EQV6 | RGNLDVAKLNGDWFSIVVASNKR | 10,Asn->His[N]; | 0.0 | 1.7 | #DIV/0! | 1.99E-02 |
| Q9EQV6 | EKIEENGSMR | 2,Lys->AminoadipicAcid[K]; | 0.0 | 1.7 | #DIV/0! | 1.99E-02 |
| Q9EQV6 | NCNDETVFQAVDTALKK | 4,Asp->Pro[D]; | 0.0 | 1.7 | #DIV/0! | 1.99E-02 |
| Q9EQV6 | GKEDLIWEILK | 0,Hex[AnyN-term]; | 0.0 | 5.0 | #DIV/0! | 1.99E-02 |
| P06238 | FTKENEGTYYGPDGR | 1,Phe->Met[F]; | 0.0 | 3.3 | #DIV/0! | 1.99E-02 |
| P01015 | STLLGAEANLGK | 0,Phenylisocyanate_2H(5)[AnyN-term]; | 0.0 | 1.7 | #DIV/0! | 1.99E-02 |
| Q9EQV6 | HIKLVQEVTDFAK | 0,3sulfo[AnyN-term]; | 0.0 | 3.8 | #DIV/0! | 1.99E-02 |
| P02767 | GECQSEGVLFFQGNR | 3,CarbamidomethylDTT[C]; | 0.0 | 5.3 | #DIV/0! | 2.00E-02 |
| Q9EQV6 | TCVADENAENCDKSIHTLFGDK | 2,Carbamidomethyl[C];11,GG[C](Dicarbamidomethyl[C]); | 0.0 | 7.8 | #DIV/0! | 2.01E-02 |
| Q9EQV6 | HPDYSVSLLLR | 0,mTRAQ[AnyN-term]; | 0.0 | 59.0 | #DIV/0! | 2.02E-02 |
| P02767 | FLRFNPVTGEVPPR | 3,phosphoRibosyl[R]; | 0.0 | 13.8 | #DIV/0! | 2.03E-02 |
| Q9EQV6 | FLGHYLHEVAR | 1,Phe->Met[F]; | 0.0 | 3.2 | #DIV/0! | 2.04E-02 |
| Q9EQV6 | PPFQRPEAEAMCTSFQENPTSFLGHYLHEVAR | 12,NEMsulfur[C]; | 0.0 | 4.8 | #DIV/0! | 2.06E-02 |
| Q9EQV6 | FSVNFADSEEAKK | 0,Piperidine[AnyN-term]; | 0.0 | 5.5 | #DIV/0! | 2.06E-02 |
| Q9EQV6 | DVLTITLTPK | 1,Dehydrated[D]; | 0.0 | 12.2 | #DIV/0! | 2.08E-02 |
| P06238 | DVLTITLTPK | 1,Dehydrated[D]; | 0.0 | 12.2 | #DIV/0! | 2.08E-02 |
| Q9EQV6 | ATEDQLKTVMGDFAQFVDKCCK | 20,Carbamidomethyl[C];21,Carbamidomethyl[C]; | 0.0 | 2.7 | #DIV/0! | 2.10E-02 |
| P02767 | RDGVLDSVTDQDSKDSTYSMSSTLSLTK | 6,Cation_Ca[II][D]; | 0.0 | 2.7 | #DIV/0! | 2.10E-02 |
| P06238 | RDGVLDSVTDQDSKDSTYSMSSTLSLTK | 6,Cation_Ca[II][D]; | 0.0 | 2.7 | #DIV/0! | 2.10E-02 |
| Q9EQV6 | DDNPNLPPFQRPEAEAMCTSFQENPTSFLGHYLHEVAR | 10,Gln->Ala[Q]; | 0.0 | 5.0 | #DIV/0! | 2.10E-02 |
| Q9EQV6 | FQENPTSFLGHYLHEVAR | 5,Pro->Phe[P]; | 0.0 | 8.3 | #DIV/0! | 2.11E-02 |
| Q9EQV6 | LLECADDRAELAK | 4,HNE+Delta_H(2)[C]; | 0.0 | 3.5 | #DIV/0! | 2.12E-02 |
| P02767 | DSTYSMSSTLSLTK | 4,Dehydrated[Y]; | 0.0 | 3.5 | #DIV/0! | 2.12E-02 |
| P06238 | DSTYSMSSTLSLTK | 4,Dehydrated[Y]; | 0.0 | 3.5 | #DIV/0! | 2.12E-02 |
| Q9EQV6 | PEAEAMCTSFQENPTSFLGHYLHEVAR | 2,Glu->His[E]; | 0.0 | 11.5 | #DIV/0! | 2.14E-02 |
| Q9EQV6 | VFNNDADLSGITEDAPLK | 0,Dimethyl[AnyN-term](Ethyl[AnyN-term]); | 0.0 | 4.0 | #DIV/0! | 2.14E-02 |
| Q9EQV6 | CTSFQENPTSFLGHYLHEVAR | 1,Hydroxycinnamyl[C]; | 0.0 | 10.3 | #DIV/0! | 2.19E-02 |
| Q9EQV6 | CCAEGDPPACYGTVLAEFQPLVEEPK | 10,4-ONE[C]; | 0.0 | 22.8 | #DIV/0! | 2.21E-02 |
| Q9EQV6 | PYFYAPELLYYAEK | 0,GIST-Quat[AnyN-term]; | 0.0 | 2.5 | #DIV/0! | 2.21E-02 |
| Q9EQV6 | PTSFLGHYLHEVAR | 0,mTRAQ[AnyN-term]; | 0.0 | 7.7 | #DIV/0! | 2.23E-02 |
| Q9EQV6 | PNAEFAEITK | 0,mTRAQ[AnyN-term]; | 0.0 | 2.3 | #DIV/0! | 2.24E-02 |
| P06238 | VYHKEESSCIHSSCTAER | 2,Dehydrated[Y];9,Carbamidomethyl[C];14,Carbamidomethyl[C]; | 0.0 | 3.3 | #DIV/0! | 2.24E-02 |
| P06238 | VYHKEESSCIHSSCTAER | 2,Dehydrated[Y];9,Carbamidomethyl[C];14,Carbamidomethyl[C]; | 0.0 | 3.3 | #DIV/0! | 2.24E-02 |
| Q9EQV6 | SKDFQLFGSPLGKDLLFK | 0,Carbamyl[AnyN-term]; | 0.0 | 6.0 | #DIV/0! | 2.26E-02 |
| Q9EQV6 | GNTFTCSVLHEGLHNHHTEK | 6,MM-diphenylpentanone[C]; | 0.0 | 4.5 | #DIV/0! | 2.27E-02 |
| P06238 | GNTFTCSVLHEGLHNHHTEK | 6,MM-diphenylpentanone[C]; | 0.0 | 4.5 | #DIV/0! | 2.27E-02 |
| Q9EQV6 | GTTYEEYLGAEYLQAVGNIR | 3,Phosphoadenosine[T]; | 0.0 | 8.7 | #DIV/0! | 2.27E-02 |
| Q9EQV6 | EGVCPEGSIDSAPVK | 1,Cation_Na[E];4,Carbamidomethyl[C]; | 0.0 | 2.8 | #DIV/0! | 2.28E-02 |
| Q9EQV6 | EDAQETDTSQQDQSPTYR | 2,Cation_Fe[III][D]; | 0.0 | 14.2 | #DIV/0! | 2.28E-02 |
| Q9EQV6 | HPYFYAPELLYYAEK | 3,Nitro[Y]; | 0.0 | 6.7 | #DIV/0! | 2.37E-02 |
| Q9EQV6 | TTIFEVLPQK | 1,Decanoyl[T]; | 0.0 | 27.0 | #DIV/0! | 2.38E-02 |
| Q9EQV6 | SFLGHYLHEVAR | 1,monomethylphosphothione[S]; | 0.0 | 3.7 | #DIV/0! | 2.39E-02 |
| P02767 | PFASGKTAESGELHGLTTDEK | 6,AEBS[K]; | 0.0 | 4.5 | #DIV/0! | 2.41E-02 |
| Q9EQV9 | DSCFNSPMR | 3,Carbamidomethyl[C]; | 0.0 | 4.5 | #DIV/0! | 2.41E-02 |
| Q9EQV6 | DQLKTVMGDFAQFVDK | 0,dNIC[AnyN-term]; | 0.0 | 4.3 | #DIV/0! | 2.42E-02 |
| Q9EQV6 | TEFATCHLAQAPNHVVVSR | 0,Biotin_Thermo-21328[AnyN-term]; | 0.0 | 7.7 | #DIV/0! | 2.43E-02 |
| Q9EQV6 | HLAQAPNHVVVSR | 0,Unknown_248[AnyN-term]; | 0.0 | 5.0 | #DIV/0! | 2.44E-02 |
| Q9EQV6 | AHRFKDLGEQHFK | 3,2HPG[R]; | 0.0 | 7.5 | #DIV/0! | 2.47E-02 |
| Q9EQV6 | SQAVTCNVAHPASSTK | 6,Propionamide[C]; | 0.0 | 1.3 | #DIV/0! | 2.50E-02 |
| Q9EQV6 | PEAQRLPCVEDYLSAILNR | 5,Arg->Tyr[R]; | 0.0 | 1.3 | #DIV/0! | 2.50E-02 |
| Q9EQV6 | PYFYAPELLYYAEK | 0,Phenylisocyanate_2H(5)[AnyN-term]; | 0.0 | 0.7 | #DIV/0! | 2.50E-02 |
| Q9EQV6 | HPYFYAPELLYYAEK | 3,Tyr->His[Y]; | 0.0 | 2.0 | #DIV/0! | 2.50E-02 |
| Q9EQV6 | CYGTVLAEFQPLVEEPK | 0,Delta_H(2)C(2)[AnyN-term]; | 0.0 | 0.7 | #DIV/0! | 2.50E-02 |
| Q9EQV6 | KCCAEGDPPACYGTVLAEFQPLVEEPK | 0,Hex[AnyN-term]; | 0.0 | 2.7 | #DIV/0! | 2.50E-02 |
| Q9EQV6 | PEAQRLPCVEDYLSAILNR | 2,iodoTMT[E]; | 0.0 | 0.7 | #DIV/0! | 2.50E-02 |
| Q9EQV6 | QEEGAQELNCNDETVFQAVDTALK | 10,Propionamide_2H(3)[C]; | 0.0 | 0.7 | #DIV/0! | 2.50E-02 |
| Q9EQV6 | SKDFQLFGSPLGKDLLFK | 9,Carbamyl[S]; | 0.0 | 0.7 | #DIV/0! | 2.50E-02 |
| Q9EQV6 | VSTVLTAQK | 2,Dehydrated[S]; | 0.0 | 0.7 | #DIV/0! | 2.50E-02 |
| Q9EQV6 | GNTFTCSVLHEGLHNHHTEK | 6,EHD-diphenylpentanone[C]; | 0.0 | 0.7 | #DIV/0! | 2.50E-02 |
| Q9EQV6 | MNGQPQENYK | 1,Oxidation[M];8,Deamidated[N]; | 0.0 | 0.7 | #DIV/0! | 2.50E-02 |
| Q9EQV6 | FLEEVKNNYHSEAFSVNFADSEEAKK | 7,Asn->His[N]; | 0.0 | 2.7 | #DIV/0! | 2.50E-02 |
| Q9EQV6 | EDAQETDTSQQDQSPTYR | 5,Cation_Ca[II][E]; | 0.0 | 2.7 | #DIV/0! | 2.50E-02 |
| P02767 | PATITDNVIPACLPSPNYVVADR | 0,Biotin_Thermo-21328[AnyN-term]; | 0.0 | 0.7 | #DIV/0! | 2.50E-02 |
| P02767 | QSWVSMTPHSHSK | 0,Tween20[AnyN-term]; | 0.0 | 0.7 | #DIV/0! | 2.50E-02 |
| P02767 | HSRCNADPGLSALLSDHR | 4,SulfoGMBS[C]; | 0.0 | 5.3 | #DIV/0! | 2.50E-02 |
| P02767 | LISERWKNPVTSVDAAFR | 0,Diisopropylphosphate[AnyN-term]; | 0.0 | 0.7 | #DIV/0! | 2.50E-02 |
| P02767 | RDGVLDSVTDQDSKDSTYSMSSTLSLTK | 2,Cation_Ca[II][D]; | 0.0 | 2.0 | #DIV/0! | 2.50E-02 |
| P06238 | GNTFTCSVLHEGLHNHHTEK | 6,EHD-diphenylpentanone[C]; | 0.0 | 0.7 | #DIV/0! | 2.50E-02 |
| P06238 | MNGQPQENYK | 1,Oxidation[M];8,Deamidated[N]; | 0.0 | 0.7 | #DIV/0! | 2.50E-02 |
| P06238 | RDGVLDSVTDQDSKDSTYSMSSTLSLTK | 2,Cation_Ca[II][D]; | 0.0 | 2.0 | #DIV/0! | 2.50E-02 |
| P04639 | LNLNLLDNWDTLGSTVGR | 0,BITC[AnyN-term]; | 0.0 | 0.7 | #DIV/0! | 2.50E-02 |
| P06238 | PRKDPPPKDPVIETIR | 0,glycidamide[AnyN-term]; | 0.0 | 18.2 | #DIV/0! | 2.51E-02 |
| P06238 | PRKDPPPKDPVIETIR | 0,glycidamide[AnyN-term]; | 0.0 | 18.2 | #DIV/0! | 2.51E-02 |
| Q9EQV6 | EAEAMCTSFQENPTSFLGHYLHEVAR | 6,NEIAA_2H(5)[C]; | 0.0 | 5.0 | #DIV/0! | 2.56E-02 |
| Q9EQV6 | DVAFVKHTTIFEVLPQK | 6,Lys->Thr[K]; | 0.0 | 4.2 | #DIV/0! | 2.57E-02 |
| Q9EQV6 | LKTVMGDFAQFVDK | 2,OxLysBiotin[K]; | 0.0 | 4.2 | #DIV/0! | 2.57E-02 |
| Q9EQV6 | PTSFLGHYLHEVAR | 0,Unknown_250[AnyN-term]; | 0.0 | 3.8 | #DIV/0! | 2.58E-02 |
| Q9EQV6 | VCPEGSIDSAPVK | 0,Unknown_302[AnyN-term]; | 0.0 | 4.3 | #DIV/0! | 2.58E-02 |
| P06238 | TACAHWSDHCCLWSTEIQVK | 3,Carbamidomethyl[C];10,GG[C](Dicarbamidomethyl[C]);16,Cation_Na[E]; | 0.0 | 6.0 | #DIV/0! | 2.59E-02 |
| Q9EQV6 | GLVLIAFSQYLQKCPYEEHIK | 13,Succinyl[K]; | 0.0 | 3.5 | #DIV/0! | 2.59E-02 |
| Q9EQV6 | VTEFATCHLAQAPNHVVVSR | 8,Phosphoguanosine[H]; | 0.0 | 6.5 | #DIV/0! | 2.60E-02 |
| Q9EQV6 | PPTMDTDGSYFLYSK | 13,Didehydro[Y]; | 0.0 | 3.2 | #DIV/0! | 2.61E-02 |
| P06238 | PPTMDTDGSYFLYSK | 13,Didehydro[Y]; | 0.0 | 3.2 | #DIV/0! | 2.61E-02 |
| P06238 | ANEPSPGEGDSNCVTR | 0,ISD_z+2_ion[AnyN-term];13,Carbamidomethyl[C]; | 0.0 | 3.2 | #DIV/0! | 2.61E-02 |
| Q9EQV6 | TSFQENPTSFLGHYLHEVAR | 0,Hex[AnyN-term]; | 0.0 | 7.8 | #DIV/0! | 2.62E-02 |
| Q9EQV6 | DDNPNLPPFQRPEAEAMCTSFQENPTSFLGHYLHEVAR | 18,Carbamidomethyl[C];24,Deamidated[N]; | 0.0 | 5.8 | #DIV/0! | 2.62E-02 |
| Q9EQV6 | EDQLKTVMGDFAQFVDK | 6,Hex(1)HexNAc(1)[T]; | 0.0 | 4.7 | #DIV/0! | 2.64E-02 |
| Q9EQV6 | NECFLQHKDDNPNLPPFQRPEAEAMCTSFQENPTSFLGHYLHEVAR | 3,maleimide[C]; | 0.0 | 6.0 | #DIV/0! | 2.67E-02 |
| Q9EQV6 | RLPCVEDYLSAILNR | 4,Delta_H(6)C(3)O(1)[C]; | 0.0 | 3.0 | #DIV/0! | 2.67E-02 |
| Q9EQV6 | CNANVYMRPWENK | 0,15N-oxobutanoic[AnyN-termC]; | 0.0 | 3.0 | #DIV/0! | 2.67E-02 |
| Q9EQV6 | CNANVYMRPWENK | 0,15N-oxobutanoic[AnyN-termC]; | 0.0 | 3.0 | #DIV/0! | 2.67E-02 |
| Q9EQV6 | FATCHLAQAPNHVVVSR | 1,Phe->His[F]; | 0.0 | 3.0 | #DIV/0! | 2.67E-02 |
| P08934 | CNANVYMRPWENK | 0,15N-oxobutanoic[AnyN-termC]; | 0.0 | 3.0 | #DIV/0! | 2.67E-02 |
| Q9EQV6 | CKVNSGAFPAPIEK | 0,Carbamyl[AnyN-term]; | 0.0 | 17.5 | #DIV/0! | 2.69E-02 |
| P06238 | TACAHWSDHCCLWSTEIQVK | 3,Carbamidomethyl[C];11,GG[C](Dicarbamidomethyl[C]);16,Cation_Na[E]; | 0.0 | 9.8 | #DIV/0! | 2.70E-02 |
| Q9EQV6 | FSWFIDDVEVHTAQTHAPEK | 6,Cation_Fe[III][D]; | 0.0 | 6.7 | #DIV/0! | 2.71E-02 |
| Q9EQV6 | KMPNASPEELADMVAK | 0,dNIC[AnyN-term]; | 0.0 | 4.5 | #DIV/0! | 2.71E-02 |
| P27274 | CYNCLDPVSSCK | 1,Carbamidomethyl[C];4,Carbamidomethyl[C];11,Carbamidomethyl[C]; | 0.0 | 5.7 | #DIV/0! | 2.74E-02 |
| P00787 | EQWSNCPTIAQIR | 6,Carbamidomethyl[C]; | 0.0 | 10.3 | #DIV/0! | 2.74E-02 |
| Q9EQV6 | TGSEVSSVFIFPPK | 0,CLIP_TRAQ_2[AnyN-term]; | 0.0 | 2.2 | #DIV/0! | 2.74E-02 |
| P02767 | AVLDVDETGTEGAAATAVTAALK | 11,Cation_Na[E]; | 0.0 | 2.2 | #DIV/0! | 2.74E-02 |
| P02767 | SMEEILEGLK | 0,Galactosyl[AnyN-term]; | 0.0 | 2.2 | #DIV/0! | 2.74E-02 |
| P06238 | TGSEVSSVFIFPPK | 0,CLIP_TRAQ_2[AnyN-term]; | 0.0 | 2.2 | #DIV/0! | 2.74E-02 |
| P22057 | GHDTVQPNFQQDK | 0,GIST-Quat[AnyN-term]; | 0.0 | 2.2 | #DIV/0! | 2.74E-02 |
| Q63416 | ATPANLEEAR | 0,C+12[AnyN-term]; | 0.0 | 4.3 | #DIV/0! | 2.74E-02 |
| Q9EQV6 | TSFLGHYLHEVAR | 1,monomethylphosphothione[T]; | 0.0 | 3.3 | #DIV/0! | 2.77E-02 |
| Q9EQV6 | NNYHSEAFSVNFADSEEAKK | 14,Asp->His[D]; | 0.0 | 3.8 | #DIV/0! | 2.78E-02 |
| P26644 | AQTQPREEQYNSTFRVVSALPIQHQDWMSGK | 5,Pro->Cys[P]; | 0.0 | 7.2 | #DIV/0! | 2.82E-02 |
| Q9EQV6 | TVMGDFAQFVDK | 0,Piperidine[AnyN-term]; | 0.0 | 1.8 | #DIV/0! | 2.84E-02 |
| Q9EQV6 | RPEAEAMCTSFQENPTSFLGHYLHEVAR | 0,NP40[AnyN-term]; | 0.0 | 1.8 | #DIV/0! | 2.84E-02 |
| Q9EQV6 | MSQRFPNAEFAEITK | 2,Isopropylphospho[S]; | 0.0 | 1.8 | #DIV/0! | 2.84E-02 |
| P02767 | SGELHGLTTDEKFTEGVYR | 0,LG-anhydrolactam[AnyN-term]; | 0.0 | 1.8 | #DIV/0! | 2.84E-02 |
| Q9EQV6 | RHPDYSVSLLLR | 0,CLIP_TRAQ_4[AnyN-term]; | 0.0 | 4.5 | #DIV/0! | 2.86E-02 |
| Q9EQV6 | MSQRFPNAEFAEITK | 4,Trimethyl_2H(9)[R]; | 0.0 | 8.3 | #DIV/0! | 2.87E-02 |
| P06238 | INFECSYPLDMK | 0,ISD_z+2_ion[AnyN-term];2,Deamidated[N];5,Carbamidomethyl[C]; | 0.0 | 4.0 | #DIV/0! | 2.88E-02 |
| P26644 | QPREEQYNSTFRVVSALPIQHQDWMSGK | 10,PhosphoCytidine[T]; | 0.0 | 6.5 | #DIV/0! | 2.89E-02 |
| P06238 | SVSSAYINTK | 0,C+12[AnyN-term]; | 0.0 | 7.5 | #DIV/0! | 2.90E-02 |
| Q9EQV6 | GDWFSIVVASNKR | 0,CarboxymethylDMAP[AnyN-term]; | 0.0 | 3.2 | #DIV/0! | 2.92E-02 |
| Q9EQV6 | LRDNYGELADCCAKQEPER | 11,Ub-amide[C]; | 0.0 | 2.7 | #DIV/0! | 2.93E-02 |
| Q9EQV6 | SSTKDLLFR | 0,Carbamyl[AnyN-term]; | 0.0 | 22.2 | #DIV/0! | 2.93E-02 |
| P02767 | NPVTGEVPPR | 0,Diisopropylphosphate[AnyN-term]; | 0.0 | 7.7 | #DIV/0! | 2.94E-02 |
| Q9EQV6 | GKSKDFQLFGSPLGK | 0,3sulfo[AnyN-term]; | 0.0 | 6.7 | #DIV/0! | 2.98E-02 |
| P02767 | TADGSWEPFASGK | 0,Triton[AnyN-term]; | 0.0 | 6.3 | #DIV/0! | 3.00E-02 |
| P02770 | NLQVLAEFQEK | 0,Biotin_Thermo-21330[AnyN-term]; | 0.0 | 5.2 | #DIV/0! | 3.00E-02 |
| Q9EQV6 | CTSFQENPTSFLGHYLHEVAR | 1,Carboxymethyl[C]; | 0.0 | 3.0 | #DIV/0! | 3.01E-02 |
| Q9EQV6 | FKDLGEQHFK | 1,Phe->Gln[F]; | 0.0 | 1.5 | #DIV/0! | 3.01E-02 |
| Q9EQV6 | QHLPEQEEPEDSK | 2,BDMAPP[H]; | 0.0 | 1.5 | #DIV/0! | 3.01E-02 |
| Q6IMF3 | FLEQQNQVLQTK | 1,Phe->Met[F]; | 0.0 | 4.5 | #DIV/0! | 3.01E-02 |
| Q6IG03 | FLEQQNQVLQTK | 1,Phe->Met[F]; | 0.0 | 4.5 | #DIV/0! | 3.01E-02 |
| Q6IG01 | FLEQQNQVLQTK | 1,Phe->Met[F]; | 0.0 | 4.5 | #DIV/0! | 3.01E-02 |
| P00762 | LGEHNINVLEGDEQFINAAK | 3,Cation_Na[E];10,Cation_Na[E]; | 0.0 | 1.5 | #DIV/0! | 3.01E-02 |
| Q9EQV6 | TCVADENAENCDKSIHTLFGDK | 2,GG[C](Dicarbamidomethyl[C]);11,Carbamidomethyl[C]; | 0.0 | 7.7 | #DIV/0! | 3.05E-02 |
| Q9EQV6 | MGDFAQFVDK | 0,Methyl[AnyN-term]; | 0.0 | 1.7 | #DIV/0! | 3.05E-02 |
| Q9EQV6 | CKVNSGAFPAPIEK | 0,ISD_z+2_ion[AnyN-term];1,Carbamidomethyl[C]; | 0.0 | 7.3 | #DIV/0! | 3.07E-02 |
| P06238 | DWMSIVTPAR | 1,Cation_Na[D]; | 0.0 | 3.5 | #DIV/0! | 3.09E-02 |
| P00762 | HPSYNANTFDNDIMLIK | 0,LG-anhydrolactam[AnyN-term]; | 0.0 | 4.2 | #DIV/0! | 3.10E-02 |
| Q9EQV6 | CRELYLVAYK | 0,Pyro-carbamidomethyl[AnyN-termC]; | 0.0 | 14.7 | #DIV/0! | 3.13E-02 |
| Q9EQV6 | GVCPEGSIDSAPVK | 7,Glucuronyl[S]; | 0.0 | 10.2 | #DIV/0! | 3.17E-02 |
| Q9EQV6 | CCAEGDPPACYGTVLAEFQPLVEEPK | 1,NEMsulfur[C]; | 0.0 | 12.2 | #DIV/0! | 3.17E-02 |
| Q9EQV6 | CVLHEKTPVSEK | 1,Delta_Hg(1)[C]; | 0.0 | 2.5 | #DIV/0! | 3.18E-02 |
| P02767 | SSPTVFRLLWESGSLLR | 0,Biotin_Thermo-21328[AnyN-term]; | 0.0 | 7.5 | #DIV/0! | 3.18E-02 |
| P02767 | NGTAHGNSTHPMHSRCNADPGLSALLSDHR | 16,Cys->Arg[C]; | 0.0 | 7.3 | #DIV/0! | 3.19E-02 |
| Q9EQV6 | YMCENQATISSK | 3,MM-diphenylpentanone[C]; | 0.0 | 6.7 | #DIV/0! | 3.19E-02 |
| Q9EQV6 | LVERRPCFSALTVDETYVPK | 7,HexNAc[C]; | 0.0 | 19.3 | #DIV/0! | 3.20E-02 |
| Q9EQV6 | PQVSTPTLVEAAR | 0,Propionyl_13C(3)[AnyN-term]; | 0.0 | 12.0 | #DIV/0! | 3.21E-02 |
| Q9EQV6 | EAEAMCTSFQENPTSFLGHYLHEVAR | 0,Ethylphosphate[AnyN-term]; | 0.0 | 3.2 | #DIV/0! | 3.23E-02 |
| Q9EQV6 | AMCTSFQENPTSFLGHYLHEVAR | 3,Dipyrrolylmethanemethyl[C]; | 0.0 | 40.7 | #DIV/0! | 3.26E-02 |
| Q9EQV6 | KQEPERNECFLQHKDDNPNLPPFQRPEAEAMCTSFQENPTSFLGHYLHEVAR | 0,ISD_z+2_ion[AnyN-term]; | 0.0 | 2.0 | #DIV/0! | 3.27E-02 |
| Q9EQV6 | REAHKSEIAHR | 1,Arg->Ser[R]; | 0.0 | 4.0 | #DIV/0! | 3.27E-02 |
| P06238 | KTPEEILTEK | 0,Cytopiloyne+water[AnyN-term]; | 0.0 | 2.0 | #DIV/0! | 3.27E-02 |
| P00689 | GAGGASILTFWDAR | 0,DiART6plex118[AnyN-term]; | 0.0 | 2.0 | #DIV/0! | 3.27E-02 |
| P02767 | LDVAETGTEAAAATGVK | 0,Tween20[AnyN-term]; | 0.0 | 14.3 | #DIV/0! | 3.29E-02 |
| Q9EQV6 | FQENPTSFLGHYLHEVAR | 4,HexNAc(1)dHex(1)[N]; | 0.0 | 11.3 | #DIV/0! | 3.29E-02 |
| Q9EQV6 | TCVADENAENCDKSIHTLFGDK | 11,Cys->CamSec[C]; | 0.0 | 3.7 | #DIV/0! | 3.30E-02 |
| Q9EQV6 | PQTHYLAVAVVK | 3,Glucuronyl[T]; | 0.0 | 6.3 | #DIV/0! | 3.31E-02 |
| P02767 | GRGHGKLRNGTAHGNSTHPMHSRCNADPGLSALLSDHR | 24,Delta_Se(1)[C]; | 0.0 | 5.7 | #DIV/0! | 3.32E-02 |
| Q9EQV6 | VSTVLTAQK | 2,Hex(1)HexNAc(1)Phos(1)[S]; | 0.0 | 4.5 | #DIV/0! | 3.32E-02 |
| Q9EQV6 | QEPERNECFLQHK | 0,Gln->pyro-Glu[AnyN-termQ];2,Cation_Na[E];8,Carbamidomethyl[C]; | 0.0 | 1.2 | #DIV/0! | 3.35E-02 |
| Q9EQV6 | CFSALTVDETYVPKEFK | 0,Cytopiloyne+water[AnyN-term]; | 0.0 | 1.2 | #DIV/0! | 3.35E-02 |
| Q9EQV6 | SFLGHYLHEVAR | 0,mTRAQ[AnyN-term]; | 0.0 | 2.3 | #DIV/0! | 3.35E-02 |
| Q9EQV6 | CPYEEHIK | 1,NEM_2H(5)+H2O[C]; | 0.0 | 1.2 | #DIV/0! | 3.35E-02 |
| Q9EQV6 | EAMCTSFQENPTSFLGHYLHEVAR | 12,dHex(1)Hex(1)[T]; | 0.0 | 1.2 | #DIV/0! | 3.35E-02 |
| Q9EQV6 | TVMGDFAQFVDK | 1,Thr->Ala[T]; | 0.0 | 1.2 | #DIV/0! | 3.35E-02 |
| Q9EQV6 | GCFQPIPMDSSDLKPVLK | 2,SulfurDioxide[C]; | 0.0 | 1.2 | #DIV/0! | 3.35E-02 |
| Q9EQV6 | GCFQPIPMDSSDLKPVLK | 2,SulfurDioxide[C]; | 0.0 | 1.2 | #DIV/0! | 3.35E-02 |
| Q9EQV6 | FATCHLAQAPNHVVVSR | 4,Bromobimane[C]; | 0.0 | 2.3 | #DIV/0! | 3.35E-02 |
| Q9EQV6 | HQTVLENTNGK | 11,Xlink_DSSO[86][K]; | 0.0 | 1.2 | #DIV/0! | 3.35E-02 |
| P02767 | GIPIVTSPYQIHFTK | 0,Piperidine[AnyN-term]; | 0.0 | 1.2 | #DIV/0! | 3.35E-02 |
| P06238 | SGSLLNNAMK | 3,Sulfo[S]; | 0.0 | 3.5 | #DIV/0! | 3.35E-02 |
| P06238 | MLIYTILPDGEVIADSVK | 1,Oxidation[M]; | 0.0 | 2.3 | #DIV/0! | 3.35E-02 |
| P06238 | SGSLLNNAMK | 3,Sulfo[S]; | 0.0 | 3.5 | #DIV/0! | 3.35E-02 |
| P06238 | MLIYTILPDGEVIADSVK | 1,Oxidation[M]; | 0.0 | 2.3 | #DIV/0! | 3.35E-02 |
| P08934 | SGSLLNNAMK | 3,Sulfo[S]; | 0.0 | 3.5 | #DIV/0! | 3.35E-02 |
| P01015 | PFTPAIFPR | 0,Cytopiloyne+water[AnyN-term]; | 0.0 | 1.2 | #DIV/0! | 3.35E-02 |
| Q9EQV6 | QENPTSFLGHYLHEVAR | 5,Decanoyl[T]; | 0.0 | 5.2 | #DIV/0! | 3.36E-02 |
| P06238 | SGSGCSILDEVIR | 5,Carboxymethyl[C]; | 0.0 | 3.0 | #DIV/0! | 3.36E-02 |
| P06238 | SGSGCSILDEVIR | 5,Carboxymethyl[C]; | 0.0 | 3.0 | #DIV/0! | 3.36E-02 |
| P04639 | KNFRTENYEEQFEMFK | 1,Retinylidene[K]; | 0.0 | 3.0 | #DIV/0! | 3.36E-02 |
| P08934 | YDGEDFIEFNK | 0,Biotin[AnyN-term]; | 0.0 | 3.0 | #DIV/0! | 3.36E-02 |
| Q9EQV6 | AEAMCTSFQENPTSFLGHYLHEVAR | 5,lapachenole[C]; | 0.0 | 2.7 | #DIV/0! | 3.37E-02 |
| Q9EQV6 | TEFATCHLAQAPNHVVVSR | 1,Hex(1)HexNAc(1)Phos(1)[T]; | 0.0 | 6.7 | #DIV/0! | 3.41E-02 |
| P06238 | SGSLLNNAMK | 1,Sulfo[S]; | 0.0 | 3.8 | #DIV/0! | 3.42E-02 |
| P06238 | SGSLLNNAMK | 1,Sulfo[S]; | 0.0 | 3.8 | #DIV/0! | 3.42E-02 |
| P08934 | SGSLLNNAMK | 1,Sulfo[S]; | 0.0 | 3.8 | #DIV/0! | 3.42E-02 |
| Q9EQV6 | NYAEAKDVFLGTFLYEYSR | 6,PS_Hapten[K]; | 0.0 | 39.2 | #DIV/0! | 3.44E-02 |
| Q9EQV6 | PEAEAMCTSFQENPTSFLGHYLHEVAR | 0,C+12[AnyN-term]; | 0.0 | 5.8 | #DIV/0! | 3.45E-02 |
| Q9EQV6 | VSTVLTAQK | 6,Dehydrated[T]; | 0.0 | 4.2 | #DIV/0! | 3.46E-02 |
| Q9EQV6 | TAPTSSPSITAPR | 0,NIC[AnyN-term](ICPL[AnyN-term]); | 0.0 | 4.2 | #DIV/0! | 3.46E-02 |
| P06238 | SEEELKKELEMYNAPPAAVEAK | 10,Cation_Na[E];11,Oxidation[M]; | 0.0 | 14.5 | #DIV/0! | 3.54E-02 |
| Q9EQV6 | NCFATEGPNLVAR | 1,Deamidated_18O(1)[N](Delta_H(1)N(-1)18O(1)[N]); | 0.0 | 3.2 | #DIV/0! | 3.55E-02 |
| P02767 | LQGTPVAQMAEDAVDGER | 4,Pent(2)[T]; | 0.0 | 5.0 | #DIV/0! | 3.56E-02 |
| Q9EQV6 | KCCAEGDPPACYGTVLAEFQPLVEEPK | 0,Formyl[AnyN-term]; | 0.0 | 4.3 | #DIV/0! | 3.58E-02 |
| Q9EQV6 | FMQHIDVLENSLGFK | 0,Carbamyl[AnyN-term]; | 0.0 | 3.5 | #DIV/0! | 3.62E-02 |
| Q9EQV6 | GTTYEEYLGAEYLQAVGNIR | 7,Phosphoadenosine[Y]; | 0.0 | 3.5 | #DIV/0! | 3.62E-02 |
| P02767 | IPIVTSPYQIHFTK | 0,SMA[AnyN-term]; | 0.0 | 7.0 | #DIV/0! | 3.62E-02 |
| P02767 | KPDGVFQEDGPVIHQEMIGGFR | 0,Pyridylacetyl[AnyN-term](Phenylisocyanate[AnyN-term]); | 0.0 | 8.7 | #DIV/0! | 3.62E-02 |
| P02767 | GECQSEGVLFFQGNR | 3,Cys->Dha[C]; | 0.0 | 5.8 | #DIV/0! | 3.64E-02 |
| Q9EQV6 | YMCENQATISSK | 0,glycidamide[AnyN-term]; | 0.0 | 6.5 | #DIV/0! | 3.64E-02 |
| Q9EQV6 | RMSQRFPNAEFAEITK | 6,Phe->Asn[F]; | 0.0 | 7.7 | #DIV/0! | 3.64E-02 |
| Q9EQV6 | KPVTEFATCHLAQAPNHVVVSR | 9,GG[C](Dicarbamidomethyl[C]); | 0.0 | 3.8 | #DIV/0! | 3.64E-02 |
| Q9EQV6 | LPCVEDYLSAILNR | 3,GG[C](Dicarbamidomethyl[C]); | 0.0 | 13.7 | #DIV/0! | 3.65E-02 |
| Q9EQV6 | ARMSQRFPNAEFAEITK | 1,Ala->Asp[A]; | 0.0 | 14.7 | #DIV/0! | 3.65E-02 |
| Q9EQV6 | GNLDVAKLNGDWFSIVVASNKR | 9,Deamidated[N]; | 0.0 | 9.7 | #DIV/0! | 3.66E-02 |
| P02767 | AFSPVASVESASGEVLHSPK | 3,Diethylphosphothione[S]; | 0.0 | 6.8 | #DIV/0! | 3.66E-02 |
| Q9EQV6 | TVMGDFAQFVDK | 0,ISD_z+2_ion[AnyN-term];3,Oxidation[M]; | 0.0 | 27.3 | #DIV/0! | 3.69E-02 |
| Q9EQV6 | GTTYEEYLGAEYLQAVGNIR | 4,Phosphoadenosine[Y]; | 0.0 | 8.5 | #DIV/0! | 3.69E-02 |
| Q9EQV6 | VDETYVPKEFK | 0,3sulfo[AnyN-term]; | 0.0 | 2.5 | #DIV/0! | 3.69E-02 |
| Q9EQV6 | CVADENAENCDKSIHTLFGDK | 1,HexNAc[C]; | 0.0 | 2.5 | #DIV/0! | 3.69E-02 |
| Q9EQV6 | RLRTKMPNASPEELADMVAK | 6,AHA-SS[M]; | 0.0 | 2.5 | #DIV/0! | 3.69E-02 |
| Q9EQV6 | CKVNSGAFPAPIEK | 2,Xlink_DST[114][K]; | 0.0 | 7.7 | #DIV/0! | 3.70E-02 |
| Q9EQV6 | AMCTSFQENPTSFLGHYLHEVAR | 3,NEIAA_2H(5)[C]; | 0.0 | 10.7 | #DIV/0! | 3.71E-02 |
| Q9EQV6 | MCTSFQENPTSFLGHYLHEVAR | 2,Cys->Xle[C]; | 0.0 | 1.8 | #DIV/0! | 3.79E-02 |
| Q9EQV6 | DNCFATEGPNLVAR | 3,4AcAllylGal[C]; | 0.0 | 3.7 | #DIV/0! | 3.79E-02 |
| Q9EQV6 | RLPCVEDYLSAILNR | 4,GG[C](Dicarbamidomethyl[C]); | 0.0 | 1.8 | #DIV/0! | 3.79E-02 |
| P06238 | VHLSFSPSQSLPASQTHMR | 4,Dehydrated[S]; | 0.0 | 1.8 | #DIV/0! | 3.79E-02 |
| P06238 | VHLSFSPSQSLPASQTHMR | 4,Dehydrated[S]; | 0.0 | 1.8 | #DIV/0! | 3.79E-02 |
| P06238 | KFTKAAPQLPMEELVSLSK | 3,Phospho[T]; | 0.0 | 3.7 | #DIV/0! | 3.79E-02 |
| P06238 | VHLSFSPSQSLPASQTHMR | 4,Dehydrated[S]; | 0.0 | 1.8 | #DIV/0! | 3.79E-02 |
| Q9EQV6 | AADKDNCFATEGPNLVAR | 5,Cation_Fe[III][D]; | 0.0 | 6.5 | #DIV/0! | 3.79E-02 |
| Q9EQV6 | HTAQTHAPEK | 0,Propyl_2H(6)[AnyN-term]; | 0.0 | 4.7 | #DIV/0! | 3.80E-02 |
| Q9EQV6 | SDSSINWNNLKGK | 1,N-dimethylphosphate[S]; | 0.0 | 7.0 | #DIV/0! | 3.82E-02 |
| P06238 | QVATVPETGRK | 0,dNIC[AnyN-term]; | 0.0 | 2.8 | #DIV/0! | 3.82E-02 |
| P02767 | LSQPEDQAEINTGSALFIDK | 3,Gln->Cys[Q]; | 0.0 | 15.3 | #DIV/0! | 3.83E-02 |
| Q9EQV6 | FKDLGEQHFK | 0,C+12[AnyN-term]; | 0.0 | 2.7 | #DIV/0! | 3.83E-02 |
| Q9EQV6 | DDNPNLPPFQRPEAEAMCTSFQENPTSFLGHYLHEVAR | 18,Carboxymethyl[C]; | 0.0 | 2.7 | #DIV/0! | 3.83E-02 |
| Q9EQV6 | FQENPTSFLGHYLHEVAR | 2,Xlink_DFDNB[Q]; | 0.0 | 3.5 | #DIV/0! | 3.88E-02 |
| P02767 | ECHRGECQSEGVLFFQGNR | 2,MDCC[C]; | 0.0 | 3.5 | #DIV/0! | 3.88E-02 |
| P02767 | ECQSEGVLFFQGNR | 2,maleimide[C]; | 0.0 | 5.5 | #DIV/0! | 3.89E-02 |
| P02767 | AAVECHRGECQSEGVLFFQGNR | 6,Diphthamide[H]; | 0.0 | 5.7 | #DIV/0! | 3.93E-02 |
| Q9EQV6 | VTCVVVDISQNDPEVR | 3,Methylthio[C]; | 0.0 | 5.8 | #DIV/0! | 3.93E-02 |
| Q9EQV6 | QENPTSFLGHYLHEVAR | 0,Propyl[AnyN-term]; | 0.0 | 8.3 | #DIV/0! | 3.93E-02 |
| Q9EQV6 | CCKAADKDNCFATEGPNLVAR | 10,HN3_mustard[C]; | 0.0 | 2.3 | #DIV/0! | 3.95E-02 |
| Q9EQV6 | CHLAQAPNHVVVSR | 0,iTRAQ4plex114[AnyN-term]; | 0.0 | 4.7 | #DIV/0! | 3.95E-02 |
| P06238 | MYYSGVDLTK | 1,Met->Aha[M]; | 0.0 | 4.7 | #DIV/0! | 3.95E-02 |
| P04639 | LTYYRKPWNVAFLAYETK | 3,sulfo+amino[Y]; | 0.0 | 2.3 | #DIV/0! | 3.95E-02 |
| P27274 | CCQADLCNK | 1,Carbamidomethyl[C];2,Carbamidomethyl[C];7,Carbamidomethyl[C]; | 0.0 | 2.3 | #DIV/0! | 3.95E-02 |
| P02767 | PDQGKMQQVESSLQPETLKK | 0,Cytopiloyne+water[AnyN-term]; | 0.0 | 5.5 | #DIV/0! | 4.00E-02 |
| P06911 | PDQGKMQQVESSLQPETLKK | 0,Cytopiloyne+water[AnyN-term]; | 0.0 | 5.5 | #DIV/0! | 4.00E-02 |
| P06911 | PDQGKMQQVESSLQPETLKK | 0,Cytopiloyne+water[AnyN-term]; | 0.0 | 5.5 | #DIV/0! | 4.00E-02 |
| P00762 | LGEHNINVLEGDEQFINAAK | 7,Ammonia-loss[N]; | 0.0 | 10.2 | #DIV/0! | 4.01E-02 |
| Q9EQV6 | LAELVKHKPK | 0,Carbamyl[AnyN-term]; | 0.0 | 4.2 | #DIV/0! | 4.03E-02 |
| Q9EQV6 | TEFATCHLAQAPNHVVVSR | 6,GG[C](Dicarbamidomethyl[C]); | 0.0 | 3.7 | #DIV/0! | 4.03E-02 |
| Q9EQV6 | MSQRFPNAEFAEITK | 0,CLIP_TRAQ_3[AnyN-term]; | 0.0 | 13.7 | #DIV/0! | 4.07E-02 |
| Q9EQV6 | DVFLGTFLYEYSR | 12,Formyl[S](Ser->Asp[S]); | 0.0 | 1.0 | #DIV/0! | 4.09E-02 |
| Q9EQV6 | DQLKTVMGDFAQFVDK | 5,FMN[T]; | 0.0 | 1.0 | #DIV/0! | 4.09E-02 |
| Q9EQV6 | PEAEAMCTSFQENPTSFLGHYLHEVAR | 9,Methylamine[S]; | 0.0 | 2.0 | #DIV/0! | 4.09E-02 |
| Q9EQV6 | KCPYEEHIK | 2,Carbamidomethyl[C]; | 0.0 | 2.0 | #DIV/0! | 4.09E-02 |
| Q9EQV6 | QEPERNECFLQHKDDNPNLPPFQRPEAEAMCTSFQENPTSFLGHYLHEVAR | 29,Ala->Trp[A]; | 0.0 | 1.0 | #DIV/0! | 4.09E-02 |
| Q9EQV6 | PPFQRPEAEAMCTSFQENPTSFLGHYLHEVAR | 18,HexNAc(1)dHex(2)[N]; | 0.0 | 1.0 | #DIV/0! | 4.09E-02 |
| Q9EQV6 | TCVADENAENCDKSIHTLFGDK | 11,DiDehydro[C]; | 0.0 | 1.0 | #DIV/0! | 4.09E-02 |
| Q9EQV6 | DKCCKAADKDNCFATEGPNLVAR | 0,Phenylisocyanate_2H(5)[AnyN-term]; | 0.0 | 1.0 | #DIV/0! | 4.09E-02 |
| Q9EQV6 | FATEGPNLVAR | 0,Unknown_250[AnyN-term]; | 0.0 | 1.0 | #DIV/0! | 4.09E-02 |
| Q9EQV6 | TVFQAVDTALKK | 0,Succinyl_2H(4)[AnyN-term]; | 0.0 | 1.0 | #DIV/0! | 4.09E-02 |
| Q9EQV6 | TVFQAVDTALKK | 0,Succinyl_2H(4)[AnyN-term]; | 0.0 | 1.0 | #DIV/0! | 4.09E-02 |
| Q9EQV6 | ASDSSINWNNLK | 0,Methyl[AnyN-term]; | 0.0 | 1.0 | #DIV/0! | 4.09E-02 |
| Q9EQV6 | DLLFKDSAFGLLR | 5,Lys->His[K]; | 0.0 | 1.0 | #DIV/0! | 4.09E-02 |
| Q9EQV6 | DGGGDVAFVK | 0,Biotin_Thermo-21330[AnyN-term]; | 0.0 | 2.0 | #DIV/0! | 4.09E-02 |
| Q9EQV6 | NNYHSEAFSVNFADSEEAKK | 17,Cation_Na[E]; | 0.0 | 1.0 | #DIV/0! | 4.09E-02 |
| P02767 | YEYCNIPECEEECMYCSGEK | 9,GG[C](Dicarbamidomethyl[C]);13,Carbamidomethyl[C];16,Carbamidomethyl[C]; | 0.0 | 1.0 | #DIV/0! | 4.09E-02 |
| P02767 | WSDTSPHVPK | 0,ISD_z+2_ion[AnyN-term]; | 0.0 | 2.0 | #DIV/0! | 4.09E-02 |
| Q9EQV6 | RPCFSALTVDETYVPKEFK | 3,Propionamide_2H(3)[C]; | 0.0 | 4.7 | #DIV/0! | 4.10E-02 |
| Q9EQV6 | PYFYAPELLYYAEK | 0,Delta_H(2)C(2)[AnyN-term]; | 0.0 | 2.2 | #DIV/0! | 4.11E-02 |
| Q9EQV6 | EGYNGYTGAFQCLVEK | 3,Dehydrated[Y];12,Carbamidomethyl[C]; | 0.0 | 2.2 | #DIV/0! | 4.11E-02 |
| Q9EQV6 | YYAVAVVKASDSSINWNNLK | 12,HexNAc(2)Sulf(1)[S]; | 0.0 | 2.2 | #DIV/0! | 4.11E-02 |
| P06238 | EEHHTAYAVYSLSK | 0,Iodoacetanilide[AnyN-term]; | 0.0 | 2.2 | #DIV/0! | 4.11E-02 |
| Q9EQV6 | LLECADDRAELAK | 3,Glu->Trp[E]; | 0.0 | 12.5 | #DIV/0! | 4.11E-02 |
| P06238 | QLKREICYFVVYPDYIEQNIHAVR | 7,ICAT-H_13C(6)[C]; | 0.0 | 4.5 | #DIV/0! | 4.13E-02 |
| P06238 | GDPIPNEQVFIK | 3,Pro->Cys[P]; | 0.0 | 4.5 | #DIV/0! | 4.13E-02 |
| P06238 | GDPIPNEQVFIK | 3,Pro->Cys[P]; | 0.0 | 4.5 | #DIV/0! | 4.13E-02 |
| Q9EQV6 | DVEVHTAQTHAPEK | 0,Ethylphosphate[AnyN-term]; | 0.0 | 5.7 | #DIV/0! | 4.13E-02 |
| Q9EQV6 | RPCFSALTVDETYVPK | 3,Cys->Arg[C]; | 0.0 | 15.5 | #DIV/0! | 4.14E-02 |
| Q9EQV6 | NPTSFLGHYLHEVAR | 3,Sulfo[T]; | 0.0 | 9.7 | #DIV/0! | 4.15E-02 |
| Q9EQV6 | SFLGHYLHEVAR | 1,NeuAc[S]; | 0.0 | 3.5 | #DIV/0! | 4.15E-02 |
| Q9EQV6 | KDNCFATEGPNLVAR | 4,HexNAc[C]; | 0.0 | 12.7 | #DIV/0! | 4.19E-02 |
| Q9EQV6 | CTSFQENPTSFLGHYLHEVAR | 1,MDCC[C]; | 0.0 | 0.8 | #DIV/0! | 4.22E-02 |
| Q9EQV6 | AEGDPPACYGTVLAEFQPLVEEPK | 11,Hex(1)HexNAc(1)[T]; | 0.0 | 0.8 | #DIV/0! | 4.22E-02 |
| Q9EQV6 | TSFQENPTSFLGHYLHEVAR | 6,HexN[N]; | 0.0 | 0.8 | #DIV/0! | 4.22E-02 |
| Q9EQV6 | ENPTSFLGHYLHEVAR | 0,AccQTag[AnyN-term]; | 0.0 | 0.8 | #DIV/0! | 4.22E-02 |
| Q9EQV6 | SFQENPTSFLGHYLHEVAR | 1,Glucuronyl[S]; | 0.0 | 0.8 | #DIV/0! | 4.22E-02 |
| Q9EQV6 | CTSFQENPTSFLGHYLHEVAR | 0,iTRAQ4plex114[AnyN-term]; | 0.0 | 0.8 | #DIV/0! | 4.22E-02 |
| Q9EQV6 | TSFQENPTSFLGHYLHEVAR | 0,Pyridylacetyl[AnyN-term](Phenylisocyanate[AnyN-term]); | 0.0 | 0.8 | #DIV/0! | 4.22E-02 |
| Q9EQV6 | CEAHGITR | 0,Pyro-carbamidomethyl[AnyN-termC]; | 0.0 | 1.7 | #DIV/0! | 4.22E-02 |
| Q9EQV6 | LNGDWFSIVVASNK | 1,Xle->Cys[L]; | 0.0 | 0.8 | #DIV/0! | 4.22E-02 |
| Q9EQV6 | KFAKLCEAHGITR | 0,SPITC_13C(6)[AnyN-term]; | 0.0 | 0.8 | #DIV/0! | 4.22E-02 |
| Q9EQV6 | QEDFQLLCPDGTK | 2,Cation_Na[E];8,Carbamidomethyl[C]; | 0.0 | 5.8 | #DIV/0! | 4.22E-02 |
| Q9EQV6 | CAVSEHENTK | 1,2-dimethylsuccinyl[C]; | 0.0 | 0.8 | #DIV/0! | 4.22E-02 |
| Q9EQV6 | PTQPFFGYVGAFKCLRDGGGDVAFVK | 0,Ethylphosphate[AnyN-term]; | 0.0 | 0.8 | #DIV/0! | 4.22E-02 |
| Q9EQV6 | VFNNDADLSGITEDAPLKLSQAVHK | 6,Ala->Asp[A]; | 0.0 | 0.8 | #DIV/0! | 4.22E-02 |
| P02767 | VEGDCDFHILKQDGQFR | 2,Cation_Na[E];5,Carbamidomethyl[C]; | 0.0 | 1.7 | #DIV/0! | 4.22E-02 |
| P02767 | ALYQAEAFVADFK | 2,Xle->Val[L]; | 0.0 | 0.8 | #DIV/0! | 4.22E-02 |
| P02767 | ILPDQGKMQQVESSLQPETLKK | 4,SulfanilicAcid[D]; | 0.0 | 0.8 | #DIV/0! | 4.22E-02 |
| P02767 | TYSMSSTLSLTK | 0,3sulfo[AnyN-term]; | 0.0 | 0.8 | #DIV/0! | 4.22E-02 |
| P02767 | STYSMSSTLSLTK | 3,sulfo+amino[Y]; | 0.0 | 0.8 | #DIV/0! | 4.22E-02 |
| P06238 | SEEELKKELEMYNAPPAAVEAK | 8,Cation_Na[E];11,Oxidation[M]; | 0.0 | 0.8 | #DIV/0! | 4.22E-02 |
| P06238 | HRFLVNLVKWMPEITDEER | 9,Xlink_DTSSP[174][K]; | 0.0 | 0.8 | #DIV/0! | 4.22E-02 |
| P06238 | TYSMSSTLSLTK | 0,3sulfo[AnyN-term]; | 0.0 | 0.8 | #DIV/0! | 4.22E-02 |
| P06238 | STYSMSSTLSLTK | 3,sulfo+amino[Y]; | 0.0 | 0.8 | #DIV/0! | 4.22E-02 |
| P06911 | ILPDQGKMQQVESSLQPETLKK | 4,SulfanilicAcid[D]; | 0.0 | 0.8 | #DIV/0! | 4.22E-02 |
| P06911 | ILPDQGKMQQVESSLQPETLKK | 4,SulfanilicAcid[D]; | 0.0 | 0.8 | #DIV/0! | 4.22E-02 |
| P04639 | VKDSGRDYVSQFESSTLGKQLNLNLLDNWDTLGSTVGR | 19,Malonyl[K]; | 0.0 | 0.8 | #DIV/0! | 4.22E-02 |
| P08934 | SQTNLTNCPFHDQPHLMR | 4,Deamidated[N];8,Carbamidomethyl[C]; | 0.0 | 0.8 | #DIV/0! | 4.22E-02 |
| P55314 | CECICPAGFQGVACEVTNRK | 1,GG[C](Dicarbamidomethyl[C]);3,Carbamidomethyl[C];14,Carbamidomethyl[C]; | 0.0 | 0.8 | #DIV/0! | 4.22E-02 |
| Q9EQV6 | FATEGPNLVAR | 3,NeuAc[T]; | 0.0 | 2.8 | #DIV/0! | 4.24E-02 |
| Q9EQV6 | NTAPTSSPSITAPR | 2,HexNAc[T]; | 0.0 | 9.0 | #DIV/0! | 4.25E-02 |
| Q9EQV6 | LRDNYGELADCCAK | 11,DiDehydro[C]; | 0.0 | 4.7 | #DIV/0! | 4.26E-02 |
| P08934 | WSNDGCHILPSHDEGAR | 6,Carbamidomethyl[C]; | 0.0 | 4.3 | #DIV/0! | 4.28E-02 |
| Q9EQV6 | TSFQENPTSFLGHYLHEVAR | 1,PET[T]; | 0.0 | 6.7 | #DIV/0! | 4.29E-02 |
| Q9EQV6 | QENPTSFLGHYLHEVAR | 6,Glycerophospho[S]; | 0.0 | 1.3 | #DIV/0! | 4.29E-02 |
| Q9EQV6 | TSFQENPTSFLGHYLHEVAR | 6,Deamidated[N]; | 0.0 | 1.3 | #DIV/0! | 4.29E-02 |
| Q9EQV6 | NNYHSEAFSVNFADSEEAKK | 12,Fluoro[F]; | 0.0 | 1.3 | #DIV/0! | 4.29E-02 |
| P02767 | DTLPHEDQGKGR | 1,Cation_Ca[II][D]; | 0.0 | 2.7 | #DIV/0! | 4.29E-02 |
| P02767 | AADLSDQVPDTDSETRILLQGTPVAQMAEDAVDGER | 13,HexNAc(2)Sulf(1)[S]; | 0.0 | 1.3 | #DIV/0! | 4.29E-02 |
| P02767 | AVLDVAETGTEAAAATGVK | 6,Ala->Asp[A]; | 0.0 | 2.7 | #DIV/0! | 4.29E-02 |
| P06238 | SFVYTKALMAYAFALAGNQEK | 0,CAF[AnyN-term]; | 0.0 | 1.3 | #DIV/0! | 4.29E-02 |
| Q9WUW3 | TIAACNLPIVQGPCR | 5,Carbamidomethyl[C];6,Deamidated[N];14,Carbamidomethyl[C]; | 0.0 | 1.3 | #DIV/0! | 4.29E-02 |
| Q9EQV6 | ADKDNCFATEGPNLVAR | 6,Dimethylphosphothione[C]; | 0.0 | 8.7 | #DIV/0! | 4.33E-02 |
| P06238 | EICYFVVYPDYIEQNIHAVR | 3,Ethanolyl[C]; | 0.0 | 7.3 | #DIV/0! | 4.34E-02 |
| P02767 | VSTQGASLHSLTK | 4,Gln->Lys[Q]; | 0.0 | 5.3 | #DIV/0! | 4.41E-02 |
| Q9EQV6 | ASDSSINWNNLK | 1,Carbonyl[A]; | 0.0 | 6.3 | #DIV/0! | 4.46E-02 |
| Q9EQV6 | FKDLGEQHFK | 1,Phe->Glu[F]; | 0.0 | 1.5 | #DIV/0! | 4.46E-02 |
| Q9EQV6 | NCDKSIHTLFGDK | 2,NEMsulfur[C]; | 0.0 | 1.5 | #DIV/0! | 4.46E-02 |
| Q9EQV6 | PMDSSDLKPVLKHAVEHFNNNTK | 22,FMN[T]; | 0.0 | 1.5 | #DIV/0! | 4.46E-02 |
| Q9EQV6 | YNAELESGNQFVLYR | 2,dHex[N]; | 0.0 | 1.5 | #DIV/0! | 4.46E-02 |
| Q9EQV6 | PMDSSDLKPVLKHAVEHFNNNTK | 22,FMN[T]; | 0.0 | 1.5 | #DIV/0! | 4.46E-02 |
| Q9EQV6 | KLPQKNTAPTSSPSITAPR | 0,Carbamyl[AnyN-term]; | 0.0 | 1.5 | #DIV/0! | 4.46E-02 |
| Q9EQV6 | TDTSQQDQSPTYRK | 0,Methyl_2H(2)[AnyN-term]; | 0.0 | 1.5 | #DIV/0! | 4.46E-02 |
| P06238 | TLMAYAFALAGNQEK | 5,Tyr->His[Y]; | 0.0 | 1.5 | #DIV/0! | 4.46E-02 |
| P06238 | CQLQSLGFMK | 1,Propionamide_2H(3)[C]; | 0.0 | 1.5 | #DIV/0! | 4.46E-02 |
| P06911 | STMEEILEGLK | 0,ICPL_13C(6)[AnyN-term]; | 0.0 | 1.5 | #DIV/0! | 4.46E-02 |
| P26644 | PIQHQDWMSGKEFK | 14,Xlink_DTSSP[88][K]; | 0.0 | 3.0 | #DIV/0! | 4.46E-02 |
| P20762 | VPGCSGTSGSLVTLGCLVK | 0,ISD_z+2_ion[AnyN-term];4,Carbamidomethyl[C];16,Carbamidomethyl[C]; | 0.0 | 1.5 | #DIV/0! | 4.46E-02 |
| P02767 | VELGNDATDIEDDSCPKPPEIANGYVEHLVR | 15,Carbamidomethyl[C];23,Deamidated[N];25,Dehydrated[Y]; | 0.0 | 4.0 | #DIV/0! | 4.51E-02 |
| P06238 | ANKDDRPEDLSLR | 0,Iodoacetanilide_13C(6)[AnyN-term]; | 0.0 | 3.7 | #DIV/0! | 4.53E-02 |
| Q9EQV6 | CCAEGDPPACYGTVLAEFQPLVEEPK | 10,NEMsulfur[C]; | 0.0 | 33.8 | #DIV/0! | 4.55E-02 |
| Q9EQV6 | SFQENPTSFLGHYLHEVAR | 0,SPITC_13C(6)[AnyN-term]; | 0.0 | 6.0 | #DIV/0! | 4.55E-02 |
| Q9EQV6 | TVMGDFAQFVDK | 2,Val->Cys[V]; | 0.0 | 8.7 | #DIV/0! | 4.55E-02 |
| Q9EQV6 | NGDGKEDLIWEILK | 7,Cation_Ca[II][D]; | 0.0 | 2.3 | #DIV/0! | 4.57E-02 |
| Q9EQV6 | NAELESGNQFVLYR | 0,Galactosyl[AnyN-term]; | 0.0 | 6.8 | #DIV/0! | 4.66E-02 |
| Q9EQV6 | NECFLQHKDDNPNLPPFQRPEAEAMCTSFQENPTSFLGHYLHEVAR | 18,Gln->Phe[Q]; | 0.0 | 3.5 | #DIV/0! | 4.70E-02 |
| Q9EQV6 | VEYDGGNTFTILK | 0,DiART6plex118[AnyN-term]; | 0.0 | 3.5 | #DIV/0! | 4.70E-02 |
| Q9EQV6 | PNASPEELADMVAK | 4,Pentose[S]; | 0.0 | 3.5 | #DIV/0! | 4.70E-02 |
| Q9EQV6 | TVDETYVPKEFK | 1,Phospho[T]; | 0.0 | 4.7 | #DIV/0! | 4.72E-02 |
| Q9EQV6 | DLKQEDFQLLCPDGTKKPVTEFATCHLAQAPNHVVVSR | 11,Diethylphosphothione[C]; | 0.0 | 8.3 | #DIV/0! | 4.75E-02 |
| Q9EQV6 | KLVVLPFPGK | 0,Carbamyl[AnyN-term]; | 0.0 | 4.8 | #DIV/0! | 4.76E-02 |
| Q9EQV6 | CTSFQENPTSFLGHYLHEVAR | 9,Hex(1)Pent(1)[T]; | 0.0 | 2.7 | #DIV/0! | 4.77E-02 |
| Q9EQV6 | PTSFLGHYLHEVAR | 2,Hex(1)HexNAc(1)[T]; | 0.0 | 2.7 | #DIV/0! | 4.77E-02 |
| Q9EQV6 | GDPPACYGTVLAEFQPLVEEPK | 6,Methylthio[C]; | 0.0 | 2.7 | #DIV/0! | 4.77E-02 |
| Q9EQV6 | KPKATEDQLKTVMGDFAQFVDK | 1,Xlink_BS2G[96][K]; | 0.0 | 2.7 | #DIV/0! | 4.77E-02 |
| P06238 | DGPCGTVLR | 0,ISD_z+2_ion[AnyN-term];4,Carbamidomethyl[C]; | 0.0 | 2.7 | #DIV/0! | 4.77E-02 |
| Q9EQV6 | LECADDRAELAK | 3,Carbamidomethyl[C]; | 0.0 | 1.8 | #DIV/0! | 4.78E-02 |
| Q9EQV6 | RMSQRFPNAEFAEITK | 3,Ser->Gly[S]; | 0.0 | 3.7 | #DIV/0! | 4.78E-02 |
| Q9EQV6 | NGDGKEDLIWEILK | 0,Carbamyl[AnyN-term]; | 0.0 | 3.7 | #DIV/0! | 4.78E-02 |
| Q9EQV6 | GNTFTCSVLHEGLHNHHTEK | 6,Cys->Pro[C]; | 0.0 | 1.8 | #DIV/0! | 4.78E-02 |
| P06238 | GNTFTCSVLHEGLHNHHTEK | 6,Cys->Pro[C]; | 0.0 | 1.8 | #DIV/0! | 4.78E-02 |
| P06238 | AEEHHTAYAVYSLSK | 0,glycidamide[AnyN-term]; | 0.0 | 1.8 | #DIV/0! | 4.78E-02 |
| P00762 | IIRHPSYNANTFDNDIMLIK | 3,Arg->Ser[R]; | 0.0 | 1.8 | #DIV/0! | 4.78E-02 |
| P02767 | STYSMSSTLSLTK | 1,Thiophospho[S]; | 0.0 | 4.5 | #DIV/0! | 4.79E-02 |
| P06238 | STYSMSSTLSLTK | 1,Thiophospho[S]; | 0.0 | 4.5 | #DIV/0! | 4.79E-02 |
| P02767 | LQTEGDGIYTLNSEK | 6,Cation_Fe[III][D]; | 0.0 | 6.3 | #DIV/0! | 4.80E-02 |
| Q9EQV6 | NPTSFLGHYLHEVAR | 3,Phospho[T]; | 0.0 | 2.2 | #DIV/0! | 4.82E-02 |
| Q9EQV6 | ASDSSINWNNLKGK | 3,Cation_Ca[II][D]; | 0.0 | 4.3 | #DIV/0! | 4.82E-02 |
| Q9EQV6 | CAEGDPPACYGTVLAEFQPLVEEPK | 0,Phosphogluconoylation[AnyN-term]; | 0.0 | 3.0 | #DIV/0! | 4.84E-02 |
| Q9EQV6 | QIKKQTALAELVK | 3,Diethylphosphothione[K]; | 0.0 | 3.0 | #DIV/0! | 4.84E-02 |
| Q9EQV6 | DKAACLTPKLDAVKEKALVAAVR | 2,Lys->Ala[K]; | 0.0 | 4.7 | #DIV/0! | 4.88E-02 |
| Q9EQV6 | ADKDNCFATEGPNLVAR | 0,GIST-Quat_2H(3)[AnyN-term]; | 0.0 | 2.0 | #DIV/0! | 4.93E-02 |
| P06238 | MLIYTILPDGEVIADSVK | 2,Xle->Ser[L]; | 0.0 | 2.0 | #DIV/0! | 4.93E-02 |
| P06238 | MLIYTILPDGEVIADSVK | 2,Xle->Ser[L]; | 0.0 | 2.0 | #DIV/0! | 4.93E-02 |
| P02767 | PFASGKTAESGELHGLTTDEK | 4,AEBS[S]; | 0.0 | 5.2 | #DIV/0! | 4.98E-02 |

Appendix Table 4 The types of differential post-translational modifications in the 1-month-old group, 8-month-old group, and 14-15-month-old group

| 1-month-old group | 8-month-old group | 14-15-month-old group |
| --- | --- | --- |
| 2-succinyl | 2-dimethylsuccinyl | 15N-oxobutanoic |
| 3-phosphoglyceryl | 2HPG | 2-dimethylsuccinyl |
| 3sulfo | 2-succinyl | 2HPG |
| 4AcAllylGal | 3sulfo | 2-succinyl |
| 4-ONE | 4-ONE | 3-phosphoglyceryl |
| AccQTag | 4-ONE+Delta_H(-2)O(-1) | 3sulfo |
| Acetyl | AccQTag | 4AcAllylGal |
| Acetyl_2H(3) | Acetyl | 4-ONE |
| Acetyldeoxyhypusine | Acetyl_2H(3) | 4-ONE+Delta_H(-2)O(-1) |
| Acetylhypusine | AEBS | AccQTag |
| AEBS | AEC-MAEC_2H(4) | Acetyl |
| AEC-MAEC_2H(4) | AFB1_Dialdehyde | Acetyl_2H(3) |
| Ala->His | AHA-SS_CAM | AEBS |
| Amidine | Ala->Asp | AFB1_Dialdehyde |
| Ammonia-loss | Ala->Trp | AHA-SS |
| Ammonium | Amidine | Ala->Asp |
| Arg->Glu | Ammonia-loss | Ala->Trp |
| Arg->GluSA | Arg->Gly | Amidine |
| Arg->Lys | Arg->Ser | Ammonia-loss |
| Arg->Ser | Arg->Tyr | Arg->Cys |
| Asn->Cys | Asn->Cys | Arg->Met |
| Asn->His | Asn->His | Arg->Ser |
| Asn->Lys | Asn->Lys | Arg->Tyr |
| Asn->Met | Asn->Met | Asn->Gly |
| Asn->Phe | Asn->Phe | Asn->His |
| Asn->Thr | Asn->Xle | Asn->Lys |
| Asn->Tyr | Asp->Cys | Asn->Ser |
| Asp->Gln | Asp->Gln | Asn->Tyr |
| Asp->Lys | Asp->Lys | Asp->Gly |
| Aspartylurea | Asp->Pro | Asp->His |
| AzidoF | azole | Asp->Pro |
| azole | BEMAD_ST_2H(6) | Asp->Thr |
| Bacillosamine | Benzoyl | Aspartylurea |
| BEMAD_ST_2H(6) | BHT | Atto495Maleimide |
| BHT | BHTOH | BADGE |
| BHTOH | Biotin | BDMAPP |
| Biotin_Thermo-21328 | Biotin_Thermo-21328 | BEMAD_C_2H(6) |
| Biotin_Thermo-21330 | Biotin-tyramide | BHT |
| BITC | BITC | Biotin |
| Brij35 | Brij58 | Biotin_Thermo-21328 |
| Brij58 | C+12 | Biotin_Thermo-21330 |
| Bromobimane | C8-QAT | biotinAcrolein298 |
| C+12 | CAF | Biotin-PEO-Amine |
| C8-QAT | Carbamidomethyl | BITC |
| CAF | CarbamidomethylDTT | BMP-piperidinol |
| Carbamidomethyl | Carbamyl | Brij35 |
| CarbamidomethylDTT | Carbonyl | Brij58 |
| Carbamyl | Carboxy | Bromo |
| Carbonyl | Carboxymethyl | Bromobimane |
| Carboxy | CarboxymethylDTT | C+12 |
| Carboxyethyl | Cation_Ag | CAF |
| Carboxymethyl | Cation_Ca | Carbamidomethyl |
| Carboxymethyl_13C(2) | Cation_Fe | CarbamidomethylDTT |
| CarboxymethylDMAP | Cation_Na | Carbamyl |
| CarboxymethylDTT | cGMP+RMP-loss | Carbonyl |
| Cation_Ag | CLIP_TRAQ_2 | Carboxymethyl |
| Cation_Al | CLIP_TRAQ_3 | CarboxymethylDMAP |
| Cation_Ca | CLIP_TRAQ_4 | Cation_Ag |
| Cation_Fe | Crotonaldehyde | Cation_Ca |
| Cation_Li | Cys->Ala | Cation_Fe |
| Cation_Na | Cys->Arg | Cation_Li |
| CLIP_TRAQ_2 | Cys->Asn | Cation_Na |
| CLIP_TRAQ_4 | Cys->CamSec | CLIP_TRAQ_2 |
| CresylSaligeninPhosphate | Cys->Dha | CLIP_TRAQ_3 |
| Crotonaldehyde | Cys->Gln | CLIP_TRAQ_4 |
| Cys->Ala | Cys->Gly | Cresylphosphate |
| Cys->Dha | Cys->His | CresylSaligeninPhosphate |
| Cys->Glu | Cys->Met | Crotonaldehyde |
| Cys->Met | Cys->SecNEM_2H(5) | Cys->Arg |
| Cys->SecNEM_2H(5) | Cys->Thr | Cys->CamSec |
| Cys->Trp | Cys->Trp | Cys->Dha |
| Cys->Val | Cys->Tyr | Cys->Gly |
| Cytopiloyne+water | Cys->Xle | Cys->His |
| DAET | cysTMT | Cys->Met |
| Dansyl | Cytopiloyne | Cys->Pro |
| Dap-DSP | Cytopiloyne+water | Cys->Xle |
| Deamidated | Dap-DSP | cysTMT |
| Deamidated_18O(1) | Deamidated | Cytopiloyne |
| Decanoyl | Deamidated_18O(1) | Cytopiloyne+water |
| Dehydrated | Decanoyl | Dansyl |
| Delta_H(2)C(2) | Dehydrated | Dap-DSP |
| Delta_H(-4)O(3) | Dehydro | Deamidated |
| Delta_O(4) | Delta_H(2)C(2) | Deamidated_18O(1) |
| Deoxy | Delta_H(4)C(3)O(1) | Decanoyl |
| dHex(1)Hex(2) | Delta_H(-4)O(3) | Dehydrated |
| DHP | Delta_Hg(1) | Dehydro |
| DiART6plex115 | dHex | Delta_H(2)C(2) |
| DiART6plex118 | dHex(1)Hex(1) | Delta_H(6)C(3)O(1) |
| Dicarbamidomethyl | dHex(1)Hex(2) | Delta_Hg(1) |
| Didehydro | DHP | Delta_S(-1)Se(1) |
| Diethyl | Didehydro | Delta_Se(1) |
| Diethylphosphate | Diethylphosphate | DeStreak |
| Diisopropylphosphate | Diethylphosphothione | dHex |
| DiLeu4plex118 | Difuran | dHex(1)Hex(1) |
| Dimethyl_2H(4) | Diisopropylphosphate | DHP |
| Dimethyl_2H(6)13C(2) | DiLeu4plex115 | DiART6plex118 |
| Dimethylphosphothione | Dimethyl | Dicarbamidomethyl |
| Dioxidation | Dimethyl_2H(4) | dichlorination |
| Dipyridyl | Dimethyl_2H(4)13C(2) | DiDehydro |
| DNCB_hapten | Dimethylphosphothione | Diethyl |
| dNIC | Dioxidation | Diethylphosphate |
| DTT | Dipyridyl | Diethylphosphothione |
| EQAT_2H(5) | Dipyrrolylmethanemethyl | Diisopropylphosphate |
| ESP | DNCB_hapten | Dimethyl |
| Ethanolamine | DTT | Dimethyl_2H(4) |
| Ethanolyl | EHD-diphenylpentanone | Dimethylphosphothione |
| Ethylphosphate | ESP_2H(10) | Dioxidation |
| ethylsulfonylethyl | Ethanedithiol | Diphthamide |
| Fluoro | Ethyl+Deamidated | Dipyrrolylmethanemethyl |
| Formyl | Ethylphosphate | DNCB_hapten |
| Formylasparagine | Fluorescein-tyramine | dNIC |
| Furan | Fluoro | DTT |
| Galactosyl | FMN | EHD-diphenylpentanone |
| GG | Formyl | ESP |
| GIST-Quat | Galactosyl | ESP_2H(10) |
| GIST-Quat_2H(3) | GG | Ethanedithiol |
| GIST-Quat_2H(9) | GIST-Quat | Ethanolyl |
| Gln->Cys | GIST-Quat_2H(3) | Ethylphosphate |
| Gln->His | GIST-Quat_2H(9) | ethylsulfonylethyl |
| Gln->Pro | Gln->Ala | Fluoro |
| Gln->pyro-Glu | Gln->His | FMN |
| Gln->Thr | Gln->Phe | Formyl |
| Gln->Val | Gln->Pro | FTC |
| Glu->His | Gln->pyro-Glu | Galactosyl |
| Glu->Met | Gln->Tyr | GG |
| Glu->pyro-Glu | Glu->Asn | GIST-Quat |
| Glu->pyro-Glu+Methyl | Glu->Asp | GIST-Quat_2H(3) |
| Glu->Xle | Glu->Gln | Gln->Ala |
| Glucosylgalactosyl | Glu->His | Gln->Cys |
| Glucuronyl | Glu->pyro-Glu | Gln->Lys |
| Glutathione | Glu->pyro-Glu+Methyl | Gln->Phe |
| Gly->Arg | Glu->Thr | Gln->Pro |
| Gly->Met | Glu->Trp | Gln->pyro-Glu |
| GlycerylPE | Glucuronyl | Gln->Tyr |
| glycidamide | Glutathione | Glu->Asn |
| Guanidinyl | Gly->Pro | Glu->Gln |
| HCysThiolactone | glycidamide | Glu->His |
| Hep | Glycosyl | Glu->Lys |
| Hex | HCysteinyl | Glu->Pro |
| Hex(1)HexA(1) | Hep | Glu->pyro-Glu |
| Hex(1)HexNAc(1)Phos(1) | Hex | Glu->Trp |
| Hex(1)NeuAc(1) | Hex(1)HexA(1) | Glucosylgalactosyl |
| Hex(1)Pent(2) | Hex(1)HexNAc(1) | Glucuronyl |
| Hex(2) | Hex(1)HexNAc(1)Phos(1) | Gly->Met |
| HexN | Hex(1)Pent(1) | Glycerophospho |
| HexNAc(1)dHex(1) | Hex(1)Pent(2)Me(1) | GlycerylPE |
| HexNAc(1)dHex(2) | Hex(2) | glycidamide |
| HexNAc(1)NeuAc(1) | Hex(3) | HCysThiolactone |
| HexNAc(2) | HexN | Hep |
| His->Glu | HexNAc | Hex |
| His->Met | HexNAc(1)dHex(1) | Hex(1)HexA(1) |
| His->Phe | HexNAc(1)NeuAc(1) | Hex(1)HexNAc(1) |
| His->Ser | His->Xle | Hex(1)HexNAc(1)Phos(1) |
| His->Thr | HMVK | Hex(1)NeuAc(1) |
| His->Xle | HN3_mustard | Hex(1)NeuGc(1) |
| HNE | HNE-Delta_H(2)O | Hex(1)Pent(1) |
| HNE+Delta_H(2) | Hydroxamic_acid | Hex(2) |
| HNE-Delta_H(2)O | Hydroxycinnamyl | Hex(3) |
| Hydroxamic_acid | Hydroxymethyl | HexN |
| IBTP | IBTP | HexNAc |
| ICAT-C | ICAT-C_13C(9) | HexNAc(1)dHex(1) |
| ICPL_13C(6) | ICAT-H | HexNAc(1)dHex(2) |
| ICPL_13C(6)2H(4) | ICAT-H_13C(6) | HexNAc(1)NeuAc(1) |
| IDEnT | ICPL_13C(6) | HexNAc(2)Sulf(1) |
| Iminobiotin | ICPL_13C(6)2H(4) | His->Glu |
| Iodoacetanilide | ICPL_2H(4) | His->Met |
| Iodoacetanilide_13C(6) | IGBP_13C(2) | His->Trp |
| iodoTMT | Iodoacetanilide | His->Xle |
| ISD_z+2_ion | Iodoacetanilide_13C(6) | HN2_mustard |
| LG-anhyropyrrole | iodoTMT | HN3_mustard |
| LG-pyrrole | ISD_z+2_ion | HNE+Delta_H(2) |
| Lys->AminoadipicAcid | Isopropylphospho | Hydroxamic_acid |
| Lys->Asn | iTRAQ4plex114 | Hydroxycinnamyl |
| Lys->Asp | LG-pyrrole | Hydroxymethyl |
| Lys->CamCys | Lys->AminoadipicAcid | IBTP |
| Lys->Cys | Lys->Asp | ICAT-H_13C(6) |
| Lys->His | Lys->CamCys | ICPL_13C(6) |
| Lys->Thr | Lys->Thr | ICPL_13C(6)2H(4) |
| Lys->Xle | maleimide | IED-Biotin |
| maleimide | Malonyl | Iodoacetanilide |
| MDCC | MDCC | Iodoacetanilide_13C(6) |
| Met->AspSA | Met->Aha | iodoTMT |
| Met->Cys | Met->Asp | ISD_z+2_ion |
| Met->Gln | Met->His | Isopropylphospho |
| Met->His | Methyl | iTRAQ4plex114 |
| Met->Thr | Methyl_2H(2) | iTRAQ8plex |
| Methyl | Methyl_2H(3)13C(1) | lapachenole |
| Methyl_2H(2) | Methyl+Deamidated | LG-anhydrolactam |
| Methyl_2H(3)+Acetyl_2H(3) | Methylamine | LG-anhyropyrrole |
| Methyl_2H(3)13C(1) | Methylphosphonate | Lys->Ala |
| Methyl+Deamidated | Methylthio | Lys->AminoadipicAcid |
| Methylphosphonate | monomethylphosphothione | Lys->Asn |
| Methylthio | mTRAQ | Lys->CamCys |
| monomethylphosphothione | mTRAQ_13C(6)15N(2) | Lys->His |
| mTRAQ | MTSL | Lys->Thr |
| mTRAQ_13C(6)15N(2) | NDA | maleimide |
| MurNAc | N-dimethylphosphate | Malonyl |
| NDA | NEIAA_2H(5) | MDCC |
| NEM_2H(5)+H2O | NEM_2H(5) | MeMePhosphorothioate |
| NEMsulfur | NEM_2H(5)+H2O | Met->Aha |
| Nethylmaleimide | NEMsulfur | Met->Asp |
| NeuAc | Nethylmaleimide | Met->Gln |
| NHS-fluorescein | Nethylmaleimide+water | Met->His |
| NHS-LC-Biotin | NeuAc | Met->Lys |
| NIC | NHS-LC-Biotin | Met->Phe |
| NIPCAM | NIC | Methyl |
| Nitro | Nitro | Methyl_2H(2) |
| Nitrosyl | Nmethylmaleimide | Methyl_2H(3) |
| NP40 | NO_SMX_SMCT | Methyl_2H(3)+Acetyl_2H(3) |
| O-Et-N-diMePhospho | NP40 | Methyl_2H(3)13C(1) |
| Oxidation | O-Isopropylmethylphosphonate | Methyl+Deamidated |
| OxProBiotinRed | O-Methylphosphate | Methylamine |
| PEITC | Oxidation | Methylthio |
| Pent(1)HexNAc(1) | Palmitoyl | MM-diphenylpentanone |
| Pent(2) | PEITC | monomethylphosphothione |
| Pentose | Pent(1)HexNAc(1) | mTRAQ |
| PET | Pent(2) | mTRAQ_13C(6)15N(2) |
| Phe->Arg | Pentose | Myristoyl |
| Phe->Asp | PET | NDA |
| Phe->CamCys | Phe->Asn | N-dimethylphosphate |
| Phe->His | Phe->CamCys | NEIAA_2H(5) |
| Phe->Lys | Phe->Gln | NEM_2H(5) |
| Phe->Met | Phe->Glu | NEM_2H(5)+H2O |
| Phe->Trp | Phe->His | NEMsulfur |
| Phenylisocyanate_2H(5) | Phe->Pro | NEMsulfurWater |
| Phosphoadenosine | Phe->Trp | Nethylmaleimide |
| PhosphoCytidine | Phenylisocyanate_2H(5) | NeuAc |
| Phosphogluconoylation | phenylsulfonylethyl | NIC |
| PhosphoHex | Phospho | NIPCAM |
| PhosphoHex(2) | Phosphoadenosine | Nitro |
| PhosphoHexNAc | PhosphoCytidine | Nmethylmaleimide |
| Phosphopantetheine | Phosphogluconoylation | NO_SMX_SIMD |
| Phosphopropargyl | Phosphoguanosine | NP40 |
| phosphoRibosyl | PhosphoHex | O-Et-N-diMePhospho |
| PhosphoUridine | PhosphoHex(2) | O-Isopropylmethylphosphonate |
| Piperidine | PhosphoHexNAc | O-Methylphosphate |
| Pro->Cys | phosphoRibosyl | Oxidation |
| Pro->Gly | Piperidine | OxLysBiotin |
| Pro->Phe | Pro->Arg | OxProBiotin |
| Pro->Ser | Pro->Cys | Palmitoyl |
| Pro->Tyr | Pro->Lys | PEITC |
| Propargylamine | Pro->Ser | Pent(1)HexNAc(1) |
| Propionamide | Pro->Thr | Pent(2) |
| Propionamide_2H(3) | Pro->Tyr | Pentose |
| Propionyl_13C(3) | Propargylamine | PET |
| Propyl | Propionamide | Phe->Arg |
| Propyl_2H(6) | Propionamide_2H(3) | Phe->Asn |
| PyMIC | Propionyl | Phe->Asp |
| PyridoxalPhosphate | Propionyl_13C(3) | Phe->Gln |
| Pyridylacetyl | Propyl | Phe->Glu |
| Pyro-carbamidomethyl | Propyl_2H(6) | Phe->Gly |
| pyrophospho | PyMIC | Phe->His |
| QAT | PyridoxalPhosphate | Phe->Lys |
| Ser->Cys | Pyridylacetyl | Phe->Met |
| Ser->Gly | Pyridylethyl | Phe->Pro |
| Ser->Pro | Pyro-carbamidomethyl | Phe->Trp |
| SMA | SPITC | Phenylisocyanate_2H(5) |
| SPITC | SPITC_13C(6) | Phospho |
| SPITC_13C(6) | Succinyl | Phosphoadenosine |
| Succinyl | Succinyl_2H(4) | PhosphoCytidine |
| Succinyl_2H(4) | SulfanilicAcid | Phosphogluconoylation |
| SulfanilicAcid | Sulfo | Phosphoguanosine |
| Sulfo | sulfo+amino | Phosphopantetheine |
| SulfoGMBS | Sulfo-NHS-LC-LC-Biotin | phosphoRibosyl |
| SulfurDioxide | SulfurDioxide | Piperidine |
| Thiadiazole | Thiadiazole | Pro->Arg |
| Thr->Ala | Thiophospho | Pro->Cys |
| Thr->Asn | Thr->Ala | Pro->His |
| Thr->Pro | Thr->Arg | Pro->Lys |
| Thr->Tyr | Thr->Asn | Pro->Phe |
| Thr->Xle | Thr->Asp | Pro->Trp |
| TMAB_2H(9) | Thr->Cys | Pro->Tyr |
| TMT | Thr->Pro | Propargylamine |
| TMT2plex | Thr->Xle | Propionamide |
| TNBS | TMAB_2H(9) | Propionamide_2H(3) |
| trifluoro | TMT | Propionyl |
| Trioxidation | TMT2plex | Propionyl_13C(3) |
| Triton | TNBS | Propyl |
| Trp->Kynurenin | trifluoro | Propyl_2H(6) |
| Tween20 | Trioxidation | PropylNAGthiazoline |
| Tyr->Arg | Tris | PS_Hapten |
| Tyr->Xle | Triton | PyMIC |
| Ub-Br2 | Tween20 | Pyridylacetyl |
| Unknown_162 | Tyr->Arg | Pyridylethyl |
| Unknown_210 | Tyr->Trp | Pyro-carbamidomethyl |
| Unknown_216 | Ub-amide | Retinylidene |
| Unknown_234 | Ub-Br2 | Ser->Gly |
| Unknown_250 | UgiJoullieProGly | Ser->His |
| Unknown_302 | UgiJoullieProGlyProGly | SMA |
| Unknown_306 | Unknown_210 | spermine |
| Unknown_420 | Unknown_216 | SPITC |
| Val->Arg | Unknown_234 | SPITC_13C(6) |
| Val->Asn | Unknown_248 | Succinyl |
| Val->Cys | Unknown_250 | Succinyl_2H(4) |
| Val->Glu | Unknown_302 | SulfanilicAcid |
| Val->His | Val->Asn | Sulfo |
| Val->Pro | Val->Gln | sulfo+amino |
| Val->Ser | Val->His | SulfoGMBS |
| Val->Thr | Val->Met | Sulfo-NHS-LC-LC-Biotin |
| Val->Tyr | Val->Thr | SulfurDioxide |
| Xle->Ala | Val->Xle | Thiazolidine |
| Xle->Arg | Xle->Cys | Thiophospho |
| Xle->Asp | Xle->Glu | Thr->Ala |
| Xle->Cys | Xle->Gly | Thr->Arg |
| Xle->Gln | Xle->His | Thr->Cys |
| Xle->Glu | Xle->Lys | Thr->Phe |
| Xle->Met | Xle->Pro | Thr->Pro |
| Xle->Phe | Xle->Ser | Thr->Val |
| Xle->Pro | Xle->Thr | TMAB_2H(9) |
| Xle->Ser | Xle->Trp | TMT |
| Xle->Thr | Xlink_BS2G | TMT2plex |
| Xle->Trp | Xlink_DFDNB | TNBS |
| Xle->Tyr | Xlink_DSS | trifluoro |
| Xlink_BS2G | Xlink_DST | Trimethyl_2H(9) |
| Xlink_DFDNB | Xlink_DTSSP | Trioxidation |
| Xlink_DSSO | ZQG | Triton |
| Xlink_DTSSP |  | Trp->Cys |
| Xlink_EGS |  | Tween20 |
|  |  | Tyr->Cys |
|  |  | Tyr->Gly |
|  |  | Tyr->His |
|  |  | Tyr->Pro |
|  |  | Ub-amide |
|  |  | Ub-Br2 |
|  |  | Unknown_216 |
|  |  | Unknown_234 |
|  |  | Unknown_248 |
|  |  | Unknown_250 |
|  |  | Unknown_302 |
|  |  | Unknown_306 |
|  |  | Val->Asn |
|  |  | Val->Cys |
|  |  | Val->Glu |
|  |  | Val->Met |
|  |  | Val->Ser |
|  |  | Xle->Asp |
|  |  | Xle->Cys |
|  |  | Xle->Glu |
|  |  | Xle->Met |
|  |  | Xle->Pro |
|  |  | Xle->Ser |
|  |  | Xle->Thr |
|  |  | Xle->Val |
|  |  | Xlink_BS2G |
|  |  | Xlink_BuUrBu |
|  |  | Xlink_DFDNB |
|  |  | Xlink_DSSO |
|  |  | Xlink_DST |
|  |  | Xlink_DTSSP |

Appendix Table 5 The fold changes of the shared differential post-translational modifications among the 1-month-old, 8-month-old, and 14–15-month-old groups

| Protein Names | Peptide | Modification | 1-month-old | 8-month-old | 14–15-month-old |
| --- | --- | --- | --- | --- | --- |
| CD59 glycoprotein | CYNCLDPVSSCK | 1,Carbamidomethyl[C];4,Carbamidomethyl[C];11,Carbamidomethyl[C]; | #DIV/0! | #DIV/0! | #DIV/0! |
| CD59 glycoprotein | CCQADLCNK | 1,Carbamidomethyl[C];2,Carbamidomethyl[C];7,Carbamidomethyl[C]; | #DIV/0! | #DIV/0! | #DIV/0! |
| CD59 glycoprotein | QVYQQCWR | 6,Carbamidomethyl[C]; | #DIV/0! | #DIV/0! | #DIV/0! |
| Beta-2-glycoprotein 1 | VCPFAGILENGVVR | 2,Carbamidomethyl[C]; | #DIV/0! | #DIV/0! | #DIV/0! |
| Cystatin-related protein 2 | LVNCPFEEQTEQLKR | 3,Deamidated[N];4,Carbamidomethyl[C]; | #DIV/0! | #DIV/0! | #DIV/0! |
| Prostaglandin-H2 D-isomerase | GHDTVQPNFQQDK | 0,GIST-Quat[AnyN-term]; | #DIV/0! | #DIV/0! | #DIV/0! |
| Superoxide dismutase [Cu-Zn] | PVQGVIHFEQK | 0,SPITC_13C(6)[AnyN-term]; | #DIV/0! | #DIV/0! | #DIV/0! |
| Albumin | YTQKAPQVSTPTLVEAAR | 0,Galactosyl[AnyN-term]; | #DIV/0! | #DIV/0! | 143.50 |
| Cathepsin B | EQWSNCPTIAQIR | 6,Carbamidomethyl[C]; | #DIV/0! | #DIV/0! | #DIV/0! |
| Cystatin-related protein 2 | LVNCPFEEQTEQLKR | 4,Carbamidomethyl[C]; | 134.361111111112 | 179.083333333333 | #DIV/0! |
| Major urinary protein | LCEAHGITR | 2,CarbamidomethylDTT[C]; | 72.98 | 25.91 | 5.99 |
| Afamin | ADWCQPLK | 4,Carbamidomethyl[C]; | 7.78 | #DIV/0! | 15.94 |
| Albumin | DDNPNLPPFQRPEAEAMCTSFQENPTSFLGHYLHEVAR | 18,Carbamidomethyl[C]; | 4.63 | 6.70 | 9.34 |
| Alpha-1-acid glycoprotein | QQLELEKETK | 0,Gln->pyro-Glu[AnyN-termQ]; | 3.52 | 2.60 | 2.05 |
| Clusterin | TQQYNELLHSLQSK | 0,Succinyl_2H(4)[AnyN-term]; | 2.84 | 6.22 | 18.67 |
| Plasminogen | TPENFPCK | 7,Carbamidomethyl[C]; | 1.99 | 3.42 | 3.50 |
| Prosaposin | SLPCDICK | 4,Carbamidomethyl[C];7,Carbamidomethyl[C]; | 1.71 | 0.09 | 0.00 |
| Hemopexin | DYFISCPGR | 6,Carbamidomethyl[C]; | 1.71 | 5.59 | 5.85 |
| Albumin | AETFTFHSDICTLPDKEK | 11,Carbamidomethyl[C]; | 0.26 | 0.38 | 0.26 |
| Serotransferrin | TSYQDCIK | 6,Carbamidomethyl[C]; | 0.02 | 5.00 | 5.44 |
| Serine protease inhibitor A3M | DEELSCSVLELK | 6,Carbamidomethyl[C]; | 0.02 | 0.30 | 0.25 |
| Serine protease inhibitor 2.1 | DEELSCSVLELK | 6,Carbamidomethyl[C]; | 0.02 | 0.30 | 0.25 |
| Serine protease inhibitor A3K | DEELSCSVLELK | 6,Carbamidomethyl[C]; | 0.02 | 0.30 | 0.25 |
| Serine protease inhibitor A3L | DEELSCSVLELK | 6,Carbamidomethyl[C]; | 0.02 | 0.30 | 0.25 |
| Cadherin-1 | DWVIPPINCPENQK | 9,Carbamidomethyl[C]; | 0.00 | 0.00 | 0.00 |
| Na | IVEVNGVCMEGK | 5,Deamidated[N];8,Carbamidomethyl[C];9,Oxidation[M]; | 0.00 | 0.00 | 0.00 |
| Lysosomal alpha-glucosidase | AVPTQCDVTPNSR | 6,Carbamidomethyl[C]; | 0.00 | 0.00 | 0.00 |
| Murinoglobulin-2 | VTASPQSLCGLR | 9,Carbamidomethyl[C]; | 0.00 | 0.00 | 0.00 |
| Interleukin-4 receptor subunit alpha | VLGDPTCFSDYIR | 7,Carbamidomethyl[C]; | 0.00 | 0.00 | 0.00 |
| Alpha-1-macroglobulin | ATPLSLCALTAVDQSVLLLKPEAK | 7,Carbamidomethyl[C]; | 0.00 | 0.00 | 0.00 |
| Matrix remodeling-associated protein 8 | IDDLEPADEGIYSCHLHHHYCGLHER | 14,Carbamidomethyl[C];21,Carbamidomethyl[C]; | 0.00 | 0.00 | 0.00 |
| Glycosylation-dependent cell adhesion molecule 1 | LTMLSQAVQK | 3,Oxidation[M]; | 0.00 | 0.00 | 0.00 |
| Murinoglobulin-1 | VTASPQSLCGLR | 9,Carbamidomethyl[C]; | 0.00 | 0.00 | 0.00 |
| Di-N-acetylchitobiase | EDCPCSEASLCRPIR | 3,Carbamidomethyl[C];5,Carbamidomethyl[C];11,Carbamidomethyl[C]; | 0.00 | 0.00 | 0.00 |
| Plasminogen | KQLAAGSIADCLAK | 11,Carbamidomethyl[C]; | 0.00 | 0.00 | 0.00 |
| Plasminogen | YSPSTHPSEGLEENYCR | 16,Carbamidomethyl[C]; | 0.00 | 0.00 | 0.00 |
| Plasminogen | CEGETDFICR | 1,Carbamidomethyl[C];9,Carbamidomethyl[C]; | 0.00 | 0.00 | 0.00 |
| Plasminogen | CTTPPPPPGPTYQCLK | 1,Carbamidomethyl[C];14,Carbamidomethyl[C]; | 0.00 | 0.00 | 0.00 |
| Low-density lipoprotein receptor-related protein 2 | NPCASASCSHLCLLSAQAPR | 3,Carbamidomethyl[C];8,Carbamidomethyl[C];12,Carbamidomethyl[C]; | 0.00 | 0.00 | 0.00 |
| Low-density lipoprotein receptor-related protein 2 | GFTCACPDDFQTVQLR | 4,Carbamidomethyl[C];6,Carbamidomethyl[C]; | 0.00 | 0.00 | 0.00 |
| Low-density lipoprotein receptor-related protein 2 | GSYECFCVDGFK | 5,Carbamidomethyl[C];7,Carbamidomethyl[C]; | 0.00 | 0.00 | 0.00 |
| Urinary protein 3 | CQTYPDEICAWVVVTTR | 1,Carbamidomethyl[C];9,Carbamidomethyl[C]; | 0.00 | 0.00 | 0.00 |
| Urinary protein 2 | CQTYPDEICAWVVVTTR | 1,Carbamidomethyl[C];9,Carbamidomethyl[C]; | 0.00 | 0.00 | 0.00 |
| Urinary protein 2 | TSFDSTGFCHVGR | 9,Carbamidomethyl[C]; | 0.00 | 0.00 | 0.00 |
| Urinary protein 2 | LALQCFRCTSFDSTGFCHVGR | 5,Carbamidomethyl[C];17,Carbamidomethyl[C]; | 0.00 | 0.00 | 0.00 |
| Urinary protein 2 | DSTGFCHVGR | 6,Carbamidomethyl[C]; | 0.00 | 0.00 | 0.00 |
| Urinary protein 2 | LALQCFRCTSFDSTGFCHVGR | 8,Carbamidomethyl[C];17,Carbamidomethyl[C]; | 0.00 | 0.00 | 0.00 |
| Urinary protein 1 | CQTYPDEICAWVVVTTR | 1,Carbamidomethyl[C];9,Carbamidomethyl[C]; | 0.00 | 0.00 | 0.00 |
| Heat shock cognate 71 kDa protein | LLQDFFNGK | 7,Deamidated[N]; | 0.00 | 0.00 | 0.00 |
| Afamin | TINPTVDHCCR | 9,Carbamidomethyl[C];10,Carbamidomethyl[C]; | 0.00 | 0.00 | 0.00 |
| Afamin | SLAMVQQECK | 4,Oxidation[M];9,Carbamidomethyl[C]; | 0.00 | 0.00 | 0.00 |
| Meprin A subunit beta | LSQVLSGPENDHSNMGQCK | 18,Carbamidomethyl[C]; | 0.00 | 0.00 | 0.00 |
| Cystatin-related protein 1 | LDNCPFEEQTEQLKR | 4,Carbamidomethyl[C]; | 0.00 | 0.00 | 0.00 |
| Ig lambda-2 chain C region | NSFTCQVTHEGNTVEK | 5,Carbamidomethyl[C]; | 0.00 | 0.00 | 0.00 |
| Ig lambda-2 chain C region | NSFTCQVTHEGNTVEK | 1,Deamidated[N];5,Carbamidomethyl[C]; | 0.00 | 0.00 | 0.00 |
| Ig lambda-2 chain C region | ANGAPISQGVDTANPTK | 2,Deamidated[N]; | 0.00 | 0.00 | 0.00 |
| Ig lambda-2 chain C region | NSFTCQVTHEGNTVEK | 5,Carbamidomethyl[C];12,Deamidated[N]; | 0.00 | 0.00 | 0.00 |
| Ig gamma-2A chain C region | SNSMVTLGCLVK | 2,Deamidated[N];9,Carbamidomethyl[C]; | 0.00 | 0.00 | 0.00 |
| Ig gamma-2A chain C region | SNSMVTLGCLVK | 9,Carbamidomethyl[C]; | 0.00 | 0.00 | 0.00 |
| Ig gamma-2A chain C region | KETWQQGNTFTCSVLHEGLHNHHTEK | 12,Carbamidomethyl[C]; | 0.00 | 0.00 | 0.00 |
| Ig gamma-1 chain C region | SNSMVTLGCLVK | 2,Deamidated[N];9,Carbamidomethyl[C]; | 0.00 | 0.00 | 0.00 |
| Ig gamma-1 chain C region | SNSMVTLGCLVK | 9,Carbamidomethyl[C]; | 0.00 | 0.00 | 0.00 |
| Hemopexin | CNADPGLSALLSDHR | 0,Pyro-carbamidomethyl[AnyN-termC]; | 0.00 | 0.00 | 0.00 |
| Aminopeptidase N | VVATTQMQAADAR | 7,Oxidation[M]; | 0.00 | 0.00 | 0.00 |
| Polymeric immunoglobulin receptor | SSVTFECDLGR | 7,Carbamidomethyl[C]; | 0.00 | 0.00 | 0.00 |
| Polymeric immunoglobulin receptor | YWCHWEADENGR | 3,Carbamidomethyl[C]; | 0.00 | 0.00 | 0.00 |
| Dipeptidyl peptidase 4 | KYPLLIDVYAGPCSQK | 13,Carbamidomethyl[C]; | 0.00 | 0.00 | 0.00 |
| Heat shock-related 70 kDa protein 2 | LLQDFFNGK | 7,Deamidated[N]; | 0.00 | 0.00 | 0.00 |
| Alpha-1-inhibitor 3 | VTASPQSLCGLR | 9,Carbamidomethyl[C]; | 0.00 | 0.00 | 0.00 |
| Ceruloplasmin | DLYSGLIGPLIVCR | 13,Carbamidomethyl[C]; | 0.00 | 0.00 | 0.00 |
| Ceruloplasmin | DIASGLIGPLILCK | 13,Carbamidomethyl[C]; | 0.00 | 0.00 | 0.00 |
| Trypsin-4 | HPEYNKDTLDNDIMLIK | 6,TMT2plex[K]; | 0.00 | 0.00 | 0.00 |
| Trypsin-4 | HPEYNKDTLDNDIMLIK | 6,TMT[K]; | 0.00 | 0.00 | 0.00 |
| Serotransferrin | KPVTEFATCHLAQAPNHVVVSR | 9,Carbamidomethyl[C]; | 0.00 | 0.00 | 0.00 |
| Serotransferrin | EGVCPEGSIDSAPVK | 4,Carbamidomethyl[C]; | 0.00 | 0.00 | 0.00 |
| Serotransferrin | DLKQEDFQLLCPDGTK | 11,Carbamidomethyl[C]; | 0.00 | 0.00 | 0.00 |
| Serotransferrin | ASDSSINWNNLK | 9,Deamidated[N]; | 0.00 | 0.00 | 0.00 |
| Serotransferrin | LLEACTFHK | 5,Carbamidomethyl[C]; | 0.00 | 0.00 | 0.00 |
| Serotransferrin | DCTGNFCLFR | 2,Carbamidomethyl[C];7,Carbamidomethyl[C]; | 0.00 | 0.00 | 0.00 |
| Prosaposin | GCSFLPDPYQK | 2,Carbamidomethyl[C]; | 0.00 | 0.00 | 0.00 |
| Serine protease inhibitor 2.1 | MQQVESSLQPETLKK | 0,ISD_z+2_ion[AnyN-term]; | 0.00 | 0.00 | 0.00 |
| Serine protease inhibitor 2.1 | MQQVESSLQPETLKK | 1,Oxidation[M]; | 0.00 | 0.00 | 0.00 |
| Pro-epidermal growth factor | HDLLTMAIFGDK | 5,Dehydrated[T]; | 0.00 | 0.00 | 0.00 |
| Pro-epidermal growth factor | ADLGGMDVK | 6,Oxidation[M]; | 0.00 | 0.00 | 0.00 |
| Pro-epidermal growth factor | HDLLTMAIFGDK | 6,Oxidation[M]; | 0.00 | 0.00 | 0.00 |
| Glutathione hydrolase 1 proenzyme | TPALCEVFCR | 5,Carbamidomethyl[C];9,Carbamidomethyl[C]; | 0.00 | 0.00 | 0.00 |
| Serine protease inhibitor A3K | IKDLTTPYIRDEELSCSVLELK | 16,Carbamidomethyl[C]; | 0.00 | 0.00 | 0.00 |
| Serine protease inhibitor A3K | TEGAAATAVTAALK | 0,NDA[AnyN-term]; | 0.00 | 0.00 | 0.00 |
| Serine protease inhibitor A3K | TSMVLVNYLLFK | 3,Oxidation[M]; | 0.00 | 0.00 | 0.00 |
| Serine protease inhibitor A3K | MQQVESSLQPETLKK | 0,ISD_z+2_ion[AnyN-term]; | 0.00 | 0.00 | 0.00 |
| Serine protease inhibitor A3K | IAELFSELDER | 0,C+12[AnyN-term]; | 0.00 | 0.00 | 0.00 |
| Serine protease inhibitor A3K | MQQVESSLQPETLKK | 1,Oxidation[M]; | 0.00 | 0.00 | 0.00 |
| Serine protease inhibitor A3L | TSMVLVNYLLFK | 3,Oxidation[M]; | 0.00 | 0.00 | 0.00 |
| Serine protease inhibitor A3L | MQQVESSLQPETLKK | 0,ISD_z+2_ion[AnyN-term]; | 0.00 | 0.00 | 0.00 |
| Serine protease inhibitor A3L | MQQVESSLQPETLKK | 1,Oxidation[M]; | 0.00 | 0.00 | 0.00 |
| Albumin | PPACYGTVLAEFQPLVEEPK | 4,Carbamidomethyl[C]; | 0.00 | 0.00 | 0.00 |
| Albumin | YNEVLTQCCTESDK | 8,Carbamidomethyl[C];9,Carbamidomethyl[C]; | 0.00 | 0.00 | 0.00 |
| Albumin | FPNAEFAEITK | 3,Deamidated[N]; | 0.00 | 0.00 | 0.00 |
| Albumin | QEPERNECFLQHK | 8,Carbamidomethyl[C]; | 0.00 | 0.00 | 0.00 |
| Albumin | YNEVLTQCCTESDK | 2,Deamidated[N];8,Carbamidomethyl[C];9,Carbamidomethyl[C]; | 0.00 | 0.00 | 0.00 |
| Albumin | AADKDNCFATEGPNLVAR | 8,Phe->Trp[F]; | 0.00 | 0.00 | 0.00 |
| Albumin | CCTLPEAQR | 1,Carbamidomethyl[C];2,Carbamidomethyl[C]; | 0.00 | 0.00 | 0.00 |
| Albumin | CCSGSLVER | 1,Carbamidomethyl[C];2,Carbamidomethyl[C]; | 0.00 | 0.00 | 0.00 |
| Albumin | CCAEGDPPACYGTVLAEFQPLVEEPK | 1,CarbamidomethylDTT[C]; | 0.00 | 0.00 | 0.00 |
| Albumin | CCAEGDPPACYGTVLAEFQPLVEEPK | 2,CarbamidomethylDTT[C]; | 0.00 | 0.00 | 0.00 |
| Albumin | ADKDNCFATEGPNLVAR | 6,DNCB_hapten[C]; | 0.00 | 0.00 | 0.00 |
| Major urinary protein | VFMQHIDVLENSLGFK | 12,Formyl[S](Ser->Asp[S]); | 0.00 | 0.00 | 0.00 |
| Ig kappa chain C region, A allele | SKDSTYSMSSTLSLTK | 1,Glucuronyl[S]; | 0.00 | 0.00 | 0.00 |
| Ig kappa chain C region, B allele | SKDSTYSMSSTLSLTK | 1,Glucuronyl[S]; | 0.00 | 0.00 | 0.00 |

Appendix Table 6 The biological process (BP) enrichment analysis of proteins with differential modifications in the 1-month-old group

| Pathways | P-Value | Count |
| --- | --- | --- |
| zymogen activation | 8.60E-24 | 18 |
| proteolysis | 8.70E-20 | 39 |
| regulation of systemic arterial blood pressure | 6.70E-17 | 13 |
| acute-phase response | 7.80E-15 | 13 |
| positive regulation of gene expression | 9.30E-10 | 27 |
| protein refolding | 5.80E-09 | 8 |
| response to xenobiotic stimulus | 1.60E-08 | 24 |
| substantia nigra development | 3.40E-07 | 8 |
| response to ethanol | 5.60E-07 | 15 |
| juxtaglomerular apparatus development | 1.30E-06 | 5 |
| chaperone cofactor-dependent protein refolding | 1.60E-06 | 7 |
| positive regulation of hepatic stellate cell contraction | 4.20E-06 | 4 |
| response to hypoxia | 5.20E-06 | 15 |
| response to odorant | 7.00E-06 | 5 |
| negative regulation of blood coagulation | 9.70E-06 | 5 |
| positive regulation of hepatic stellate cell migration | 1.00E-05 | 4 |
| mesenchyme migration | 1.00E-05 | 4 |
| protein catabolic process | 1.00E-05 | 10 |
| cellular response to type II interferon | 1.80E-05 | 9 |
| regulation of synaptic vesicle endocytosis | 2.00E-05 | 6 |
| regulation of blood pressure | 2.20E-05 | 8 |
| response to nutrient | 3.00E-05 | 10 |
| positive regulation of ERK1 and ERK2 cascade | 3.00E-05 | 12 |
| response to organic cyclic compound | 3.50E-05 | 14 |
| glomerular mesangial cell development | 3.60E-05 | 4 |
| vascular associated smooth muscle contraction | 3.60E-05 | 4 |
| response to mechanical stimulus | 4.60E-05 | 9 |
| proteolysis involved in protein catabolic process | 5.20E-05 | 7 |
| canonical glycolysis | 5.50E-05 | 5 |
| positive regulation of phagocytosis | 5.70E-05 | 7 |
| positive regulation of T cell mediated cytotoxicity | 6.20E-05 | 7 |
| regulation of transmembrane transporter activity | 1.20E-04 | 4 |
| carbohydrate metabolic process | 1.40E-04 | 8 |
| gluconeogenesis | 1.50E-04 | 6 |
| lactate metabolic process | 1.60E-04 | 4 |
| binding of sperm to zona pellucida | 2.00E-04 | 6 |
| response to lead ion | 2.20E-04 | 6 |
| glutathione metabolic process | 2.40E-04 | 6 |
| positive regulation of hepatic stellate cell activation | 2.80E-04 | 4 |
| response to activity | 2.80E-04 | 8 |
| postsynaptic actin cytoskeleton organization | 3.00E-04 | 5 |
| negative regulation of activation of membrane attack complex | 3.10E-04 | 3 |
| cellular response to cytochalasin B | 3.10E-04 | 3 |
| establishment of protein localization to mitochondrial membrane | 3.10E-04 | 3 |
| regulation of amyloid-beta clearance | 3.10E-04 | 3 |
| blood coagulation | 3.20E-04 | 7 |
| negative regulation of proteolysis | 3.40E-04 | 5 |
| leukocyte cell-cell adhesion | 3.40E-04 | 5 |
| response to calcium ion | 3.80E-04 | 7 |
| ERK1 and ERK2 cascade | 4.20E-04 | 6 |
| fibrinolysis | 4.30E-04 | 4 |
| cellular response to glucose stimulus | 4.80E-04 | 8 |
| response to bacterium | 4.90E-04 | 10 |
| receptor-mediated endocytosis | 6.10E-04 | 6 |
| embryo implantation | 6.10E-04 | 6 |
| negative regulation of calcium ion export across plasma membrane | 6.10E-04 | 3 |
| regulation of iron ion transport | 6.10E-04 | 3 |
| negative regulation of high voltage-gated calcium channel activity | 6.10E-04 | 3 |
| glyceraldehyde-3-phosphate metabolic process | 6.10E-04 | 3 |
| protein targeting to lysosome involved in chaperone-mediated autophagy | 6.10E-04 | 3 |
| pyruvate metabolic process | 6.40E-04 | 4 |
| amyloid-beta clearance | 6.40E-04 | 4 |
| peptide catabolic process | 7.60E-04 | 4 |
| regulation of immune system process | 7.90E-04 | 5 |
| negative regulation of endopeptidase activity | 8.70E-04 | 5 |
| cellular response to hepatocyte growth factor stimulus | 9.00E-04 | 4 |
| response to nutrient levels | 9.30E-04 | 9 |
| epithelial cell differentiation | 9.70E-04 | 7 |
| response to lipopolysaccharide | 1.00E-03 | 11 |
| regulation of norepinephrine uptake | 1.00E-03 | 3 |
| aflatoxin catabolic process | 1.00E-03 | 3 |
| positive regulation of antibacterial peptide production | 1.00E-03 | 3 |
| pyruvate catabolic process | 1.00E-03 | 3 |
| organelle localization by membrane tethering | 1.00E-03 | 3 |
| methylglyoxal biosynthetic process | 1.00E-03 | 3 |
| regulation of store-operated calcium channel activity | 1.00E-03 | 3 |
| response to heat | 1.00E-03 | 5 |
| lysosomal transport | 1.20E-03 | 4 |
| gene expression | 1.30E-03 | 11 |
| antimicrobial humoral immune response mediated by antimicrobial peptide | 1.30E-03 | 7 |
| glycolytic process | 1.40E-03 | 6 |
| negative regulation of apoptotic process | 1.50E-03 | 16 |
| regulation of transepithelial transport | 1.50E-03 | 3 |
| presynaptic endocytosis | 1.50E-03 | 3 |
| membrane to membrane docking | 1.50E-03 | 3 |
| glyceraldehyde-3-phosphate biosynthetic process | 1.50E-03 | 3 |
| regulation of high voltage-gated calcium channel activity | 1.50E-03 | 3 |
| intracellular iron ion homeostasis | 1.50E-03 | 6 |
| positive regulation of fibroblast proliferation | 1.80E-03 | 6 |
| cellular response to cAMP | 1.80E-03 | 6 |
| positive regulation of cytokine production | 2.00E-03 | 6 |
| vitamin D metabolic process | 2.10E-03 | 3 |
| protein localization to adherens junction | 2.10E-03 | 3 |
| negative regulation of fibrinolysis | 2.10E-03 | 3 |
| regulation of cardiac muscle cell action potential | 2.10E-03 | 3 |
| positive regulation of oligodendrocyte progenitor proliferation | 2.10E-03 | 3 |
| response to estradiol | 2.20E-03 | 9 |
| cellular response to amyloid-beta | 2.40E-03 | 5 |
| positive regulation of receptor-mediated endocytosis | 2.60E-03 | 4 |
| positive regulation of superoxide anion generation | 2.60E-03 | 4 |
| positive regulation of ryanodine-sensitive calcium-release channel activity | 2.80E-03 | 3 |
| protein localization to bicellular tight junction | 2.80E-03 | 3 |
| autophagic cell death | 2.80E-03 | 3 |
| NAD metabolic process | 2.80E-03 | 3 |
| skeletal muscle tissue development | 2.90E-03 | 6 |
| animal organ regeneration | 2.90E-03 | 6 |
| positive regulation of nitric oxide biosynthetic process | 2.90E-03 | 5 |
| antibacterial humoral response | 3.10E-03 | 5 |
| positive regulation of cell population proliferation | 3.10E-03 | 15 |
| response to copper ion | 3.10E-03 | 4 |
| negative regulation of ryanodine-sensitive calcium-release channel activity | 3.60E-03 | 3 |
| glycerol catabolic process | 3.60E-03 | 3 |
| morphogenesis of a polarized epithelium | 3.60E-03 | 3 |
| autophagosome membrane docking | 3.60E-03 | 3 |
| cellular response to iron ion | 4.40E-03 | 3 |
| mitochondrion-endoplasmic reticulum membrane tethering | 4.40E-03 | 3 |
| negative regulation of inclusion body assembly | 4.40E-03 | 3 |
| protein import | 4.40E-03 | 3 |
| positive regulation of protein localization to early endosome | 4.40E-03 | 3 |
| positive regulation of long-term synaptic potentiation | 4.60E-03 | 4 |
| kidney development | 4.70E-03 | 8 |
| positive regulation of endocytosis | 5.00E-03 | 4 |
| vasodilation | 5.10E-03 | 5 |
| axo-dendritic transport | 5.40E-03 | 3 |
| regulation of ryanodine-sensitive calcium-release channel activity | 5.40E-03 | 3 |
| glutathione biosynthetic process | 5.40E-03 | 3 |
| tissue regeneration | 5.40E-03 | 4 |
| positive regulation of interleukin-8 production | 5.40E-03 | 5 |
| ATP metabolic process | 6.30E-03 | 4 |
| calcineurin-mediated signaling | 6.40E-03 | 3 |
| cellular response to follicle-stimulating hormone stimulus | 6.80E-03 | 4 |
| protein maturation | 6.90E-03 | 5 |
| negative regulation of neuron apoptotic process | 7.20E-03 | 8 |
| establishment of protein localization to membrane | 7.50E-03 | 3 |
| negative regulation of lipid biosynthetic process | 7.50E-03 | 3 |
| detection of calcium ion | 7.50E-03 | 3 |
| cellular oxidant detoxification | 7.60E-03 | 5 |
| protein stabilization | 7.70E-03 | 8 |
| chaperone-mediated protein folding | 7.80E-03 | 4 |
| response to oxidative stress | 7.90E-03 | 7 |
| retina development in camera-type eye | 8.30E-03 | 6 |
| lysosomal protein catabolic process | 8.70E-03 | 3 |
| negative regulation of CD4-positive, alpha-beta T cell proliferation | 8.70E-03 | 3 |
| positive regulation of tumor necrosis factor-mediated signaling pathway | 8.70E-03 | 3 |
| adherens junction assembly | 8.70E-03 | 3 |
| cellular response to unfolded protein | 8.70E-03 | 3 |
| cell adhesion | 8.70E-03 | 11 |
| actin cytoskeleton organization | 8.90E-03 | 8 |
| regulation of heart rate | 9.00E-03 | 4 |
| cellular response to cadmium ion | 9.00E-03 | 4 |
| positive regulation of proteasomal ubiquitin-dependent protein catabolic process | 9.50E-03 | 5 |
| positive regulation of tumor necrosis factor production | 9.80E-03 | 6 |
| glutamate metabolic process | 1.00E-02 | 3 |
| positive regulation of cell migration | 1.00E-02 | 9 |
| cochlea development | 1.00E-02 | 4 |
| glucose metabolic process | 1.10E-02 | 5 |
| G2/M transition of mitotic cell cycle | 1.10E-02 | 4 |
| wound healing, spreading of cells | 1.10E-02 | 3 |
| apical protein localization | 1.10E-02 | 3 |
| cellular response to thyroid hormone stimulus | 1.10E-02 | 3 |
| thyroid hormone generation | 1.10E-02 | 3 |
| response to corticosterone | 1.20E-02 | 4 |
| response to peptide hormone | 1.20E-02 | 6 |
| apoptotic process | 1.20E-02 | 12 |
| response to glucocorticoid | 1.20E-02 | 6 |
| negative regulation of neuron projection development | 1.30E-02 | 5 |
| negative regulation of mitochondrial outer membrane permeabilization involved in apoptotic signaling pathway | 1.30E-02 | 3 |
| digestion | 1.30E-02 | 3 |
| regulation of cyclin-dependent protein serine/threonine kinase activity | 1.30E-02 | 3 |
| regulation of calcium-mediated signaling | 1.30E-02 | 3 |
| regulation of cell shape | 1.30E-02 | 6 |
| neuron apoptotic process | 1.30E-02 | 6 |
| positive regulation of axon extension | 1.30E-02 | 4 |
| iron ion transport | 1.40E-02 | 4 |
| establishment of epithelial cell apical/basal polarity | 1.40E-02 | 3 |
| antigen processing and presentation of peptide antigen via MHC class I | 1.40E-02 | 3 |
| positive regulation of nitric-oxide synthase activity | 1.60E-02 | 3 |
| positive regulation of G protein-coupled receptor signaling pathway | 1.60E-02 | 3 |
| negative regulation of cell-substrate adhesion | 1.60E-02 | 3 |
| muscle contraction | 1.60E-02 | 4 |
| cellular response to electrical stimulus | 1.70E-02 | 3 |
| response to zinc ion | 1.80E-02 | 4 |
| protein folding | 1.90E-02 | 6 |
| response to virus | 1.90E-02 | 5 |
| T cell mediated cytotoxicity | 1.90E-02 | 3 |
| protein processing | 2.00E-02 | 5 |
| positive regulation of phosphatidylinositol 3-kinase/protein kinase B signal transduction | 2.00E-02 | 7 |
| endocytic hemoglobin import into cell | 2.00E-02 | 2 |
| chemoattraction of axon | 2.00E-02 | 2 |
| protein transmembrane import into intracellular organelle | 2.00E-02 | 2 |
| chaperone-mediated autophagy translocation complex disassembly | 2.00E-02 | 2 |
| positive regulation of endoribonuclease activity | 2.00E-02 | 2 |
| positive regulation of neurofibrillary tangle assembly | 2.00E-02 | 2 |
| slow axonal transport | 2.00E-02 | 2 |
| positive regulation of protein refolding | 2.00E-02 | 2 |
| positive regulation of T cell migration | 2.10E-02 | 3 |
| chaperone-mediated protein complex assembly | 2.10E-02 | 3 |
| amyloid fibril formation | 2.10E-02 | 3 |
| symbiont entry into host cell | 2.10E-02 | 4 |
| female pregnancy | 2.20E-02 | 6 |
| regulation of release of sequestered calcium ion into cytosol by sarcoplasmic reticulum | 2.30E-02 | 3 |
| inflammatory response | 2.30E-02 | 9 |
| response to amphetamine | 2.30E-02 | 4 |
| antigen processing and presentation of exogenous peptide antigen via MHC class II | 2.50E-02 | 3 |
| regulation of cardiac muscle contraction | 2.50E-02 | 3 |
| xenobiotic catabolic process | 2.70E-02 | 3 |
| actin filament organization | 2.70E-02 | 6 |
| cellular response to heat | 2.80E-02 | 4 |
| androgen receptor signaling pathway | 2.90E-02 | 3 |
| positive regulation of MAPK cascade | 2.90E-02 | 7 |
| cysteine metabolic process | 3.00E-02 | 2 |
| peptidyl-cysteine S-trans-nitrosylation | 3.00E-02 | 2 |
| regulation of amyloid fibril formation | 3.00E-02 | 2 |
| cellular response to thyroxine stimulus | 3.00E-02 | 2 |
| angiotensin-mediated vasoconstriction involved in regulation of systemic arterial blood pressure | 3.00E-02 | 2 |
| positive regulation of lysosomal membrane permeability | 3.00E-02 | 2 |
| blood vessel diameter maintenance | 3.10E-02 | 3 |
| response to glucose | 3.10E-02 | 5 |
| response to starvation | 3.20E-02 | 4 |
| neuromuscular process controlling balance | 3.20E-02 | 4 |
| bicellular tight junction assembly | 3.20E-02 | 4 |
| cellular response to transforming growth factor beta stimulus | 3.30E-02 | 5 |
| immune response | 3.30E-02 | 9 |
| angiogenesis | 3.40E-02 | 7 |
| protein localization to plasma membrane | 3.50E-02 | 6 |
| muscle cell cellular homeostasis | 3.50E-02 | 3 |
| positive regulation of bone resorption | 3.50E-02 | 3 |
| signal transduction | 3.70E-02 | 17 |
| positive regulation of cytosolic calcium ion concentration | 3.90E-02 | 6 |
| metanephric distal convoluted tubule development | 4.00E-02 | 2 |
| positive regulation of lipoprotein transport | 4.00E-02 | 2 |
| response to nitrosative stress | 4.00E-02 | 2 |
| glutathione derivative biosynthetic process | 4.00E-02 | 2 |
| micturition | 4.00E-02 | 2 |
| Factor XII activation | 4.00E-02 | 2 |
| late endosomal microautophagy | 4.00E-02 | 2 |
| negative regulation of extracellular matrix disassembly | 4.00E-02 | 2 |
| fusion of virus membrane with host plasma membrane | 4.00E-02 | 2 |
| glucose catabolic process to lactate via pyruvate | 4.00E-02 | 2 |
| intestinal D-glucose absorption | 4.00E-02 | 2 |
| positive regulation of lysosomal protein catabolic process | 4.00E-02 | 2 |
| astrocyte activation involved in immune response | 4.00E-02 | 2 |
| insulin catabolic process | 4.00E-02 | 2 |
| regulation of protein localization to plasma membrane | 4.20E-02 | 3 |
| positive regulation of NF-kappaB transcription factor activity | 4.50E-02 | 5 |
| collagen catabolic process | 4.70E-02 | 3 |

Appendix Table 7 The biological process (BP) enrichment analysis of proteins with differential modifications in the 8-month-old group

| Pathways | P-Value | Count |
| --- | --- | --- |
| acute-phase response | 5.80E-14 | 11 |
| zymogen activation | 2.50E-12 | 10 |
| proteolysis | 1.00E-11 | 23 |
| protein refolding | 1.50E-10 | 8 |
| regulation of systemic arterial blood pressure | 2.30E-08 | 7 |
| chaperone cofactor-dependent protein refolding | 7.40E-08 | 7 |
| complement activation, classical pathway | 7.40E-08 | 7 |
| response to ethanol | 6.10E-07 | 12 |
| negative regulation of blood coagulation | 1.20E-06 | 5 |
| response to xenobiotic stimulus | 1.60E-06 | 16 |
| regulation of synaptic vesicle endocytosis | 1.60E-06 | 6 |
| negative regulation of fibrinolysis | 7.50E-06 | 4 |
| substantia nigra development | 7.60E-06 | 6 |
| response to estradiol | 9.60E-06 | 10 |
| retina development in camera-type eye | 9.60E-06 | 8 |
| blood coagulation | 1.80E-05 | 7 |
| positive regulation of receptor-mediated endocytosis | 2.00E-05 | 5 |
| antibacterial humoral response | 2.90E-05 | 6 |
| negative regulation of proteolysis | 4.60E-05 | 5 |
| positive regulation of phagocytosis | 5.00E-05 | 6 |
| positive regulation of T cell mediated cytotoxicity | 5.40E-05 | 6 |
| embryo implantation | 5.40E-05 | 6 |
| fibrinolysis | 9.40E-05 | 4 |
| negative regulation of activation of membrane attack complex | 1.10E-04 | 3 |
| cellular response to cytochalasin B | 1.10E-04 | 3 |
| establishment of protein localization to mitochondrial membrane | 1.10E-04 | 3 |
| response to mechanical stimulus | 1.20E-04 | 7 |
| amyloid-beta clearance | 1.40E-04 | 4 |
| intracellular iron ion homeostasis | 1.40E-04 | 6 |
| negative regulation of high voltage-gated calcium channel activity | 2.20E-04 | 3 |
| regulation of iron ion transport | 2.20E-04 | 3 |
| negative regulation of calcium ion export across plasma membrane | 2.20E-04 | 3 |
| protein targeting to lysosome involved in chaperone-mediated autophagy | 2.20E-04 | 3 |
| lysosomal transport | 2.70E-04 | 4 |
| response to lead ion | 3.20E-04 | 5 |
| organelle localization by membrane tethering | 3.60E-04 | 3 |
| regulation of norepinephrine uptake | 3.60E-04 | 3 |
| regulation of store-operated calcium channel activity | 3.60E-04 | 3 |
| positive regulation of gene expression | 3.70E-04 | 13 |
| response to bacterium | 4.30E-04 | 8 |
| regulation of high voltage-gated calcium channel activity | 5.40E-04 | 3 |
| presynaptic endocytosis | 5.40E-04 | 3 |
| regulation of transepithelial transport | 5.40E-04 | 3 |
| cellular response to type II interferon | 5.80E-04 | 6 |
| response to hypoxia | 7.30E-04 | 9 |
| receptor-mediated endocytosis | 7.40E-04 | 5 |
| regulation of cardiac muscle cell action potential | 7.50E-04 | 3 |
| protein localization to adherens junction | 7.50E-04 | 3 |
| positive regulation of oligodendrocyte progenitor proliferation | 7.50E-04 | 3 |
| positive regulation of ryanodine-sensitive calcium-release channel activity | 1.00E-03 | 3 |
| morphogenesis of a polarized epithelium | 1.30E-03 | 3 |
| negative regulation of ryanodine-sensitive calcium-release channel activity | 1.30E-03 | 3 |
| autophagosome membrane docking | 1.30E-03 | 3 |
| kidney development | 1.40E-03 | 7 |
| ATP metabolic process | 1.50E-03 | 4 |
| cellular response to iron ion | 1.60E-03 | 3 |
| protein import | 1.60E-03 | 3 |
| negative regulation of inclusion body assembly | 1.60E-03 | 3 |
| mitochondrion-endoplasmic reticulum membrane tethering | 1.60E-03 | 3 |
| regulation of transmembrane transporter activity | 1.60E-03 | 3 |
| chaperone-mediated protein folding | 1.80E-03 | 4 |
| regulation of ryanodine-sensitive calcium-release channel activity | 1.90E-03 | 3 |
| regulation of immune system process | 2.00E-03 | 4 |
| cellular response to cadmium ion | 2.10E-03 | 4 |
| response to lipopolysaccharide | 2.30E-03 | 8 |
| calcineurin-mediated signaling | 2.30E-03 | 3 |
| complement activation, alternative pathway | 2.30E-03 | 3 |
| response to heat | 2.40E-03 | 4 |
| animal organ regeneration | 2.70E-03 | 5 |
| response to odorant | 2.70E-03 | 3 |
| establishment of protein localization to membrane | 2.70E-03 | 3 |
| detection of calcium ion | 2.70E-03 | 3 |
| detection of chemical stimulus involved in sensory perception of bitter taste | 2.90E-03 | 4 |
| adherens junction assembly | 3.20E-03 | 3 |
| cellular response to unfolded protein | 3.20E-03 | 3 |
| iron ion transport | 3.30E-03 | 4 |
| response to organic cyclic compound | 4.00E-03 | 8 |
| binding of sperm to zona pellucida | 4.10E-03 | 4 |
| apical protein localization | 4.10E-03 | 3 |
| innate immune response | 4.30E-03 | 9 |
| inflammatory response | 4.70E-03 | 8 |
| regulation of calcium-mediated signaling | 4.70E-03 | 3 |
| regulation of cyclin-dependent protein serine/threonine kinase activity | 4.70E-03 | 3 |
| carbohydrate metabolic process | 4.70E-03 | 5 |
| response to amphetamine | 5.80E-03 | 4 |
| positive regulation of nitric-oxide synthase activity | 5.80E-03 | 3 |
| positive regulation of G protein-coupled receptor signaling pathway | 5.80E-03 | 3 |
| cellular response to electrical stimulus | 6.50E-03 | 3 |
| cellular response to heat | 6.90E-03 | 4 |
| chaperone-mediated protein complex assembly | 7.80E-03 | 3 |
| locomotory exploration behavior | 7.80E-03 | 3 |
| vasodilation | 8.20E-03 | 4 |
| regulation of release of sequestered calcium ion into cytosol by sarcoplasmic reticulum | 8.50E-03 | 3 |
| positive regulation of interleukin-8 production | 8.50E-03 | 4 |
| establishment of localization in cell | 9.20E-03 | 6 |
| regulation of cardiac muscle contraction | 9.20E-03 | 3 |
| response to magnesium ion | 9.20E-03 | 3 |
| protein catabolic process | 9.30E-03 | 5 |
| androgen receptor signaling pathway | 1.10E-02 | 3 |
| cellular oxidant detoxification | 1.10E-02 | 4 |
| negative regulation of inflammatory response | 1.20E-02 | 5 |
| positive regulation of endoribonuclease activity | 1.20E-02 | 2 |
| positive regulation of protein refolding | 1.20E-02 | 2 |
| protein transmembrane import into intracellular organelle | 1.20E-02 | 2 |
| endocytic hemoglobin import into cell | 1.20E-02 | 2 |
| slow axonal transport | 1.20E-02 | 2 |
| positive regulation of neurofibrillary tangle assembly | 1.20E-02 | 2 |
| chaperone-mediated autophagy translocation complex disassembly | 1.20E-02 | 2 |
| protein folding | 1.30E-02 | 5 |
| positive regulation of proteasomal ubiquitin-dependent protein catabolic process | 1.30E-02 | 4 |
| muscle cell cellular homeostasis | 1.30E-02 | 3 |
| positive regulation of bone resorption | 1.30E-02 | 3 |
| response to copper ion | 1.30E-02 | 3 |
| signal transduction | 1.40E-02 | 13 |
| positive regulation of endothelial cell proliferation | 1.50E-02 | 4 |
| cellular response to cAMP | 1.50E-02 | 4 |
| regulation of protein localization to plasma membrane | 1.60E-02 | 3 |
| postsynaptic actin cytoskeleton organization | 1.60E-02 | 3 |
| positive regulation of immunoglobulin production | 1.70E-02 | 3 |
| positive regulation of endocytosis | 1.80E-02 | 3 |
| complement-dependent cytotoxicity | 1.80E-02 | 2 |
| positive regulation of lysosomal membrane permeability | 1.80E-02 | 2 |
| regulation of amyloid-beta clearance | 1.80E-02 | 2 |
| response to calcium ion | 1.90E-02 | 4 |
| tissue regeneration | 1.90E-02 | 3 |
| skeletal muscle tissue development | 2.00E-02 | 4 |
| positive regulation of phosphorylation | 2.10E-02 | 3 |
| estrous cycle | 2.30E-02 | 3 |
| protein processing | 2.40E-02 | 4 |
| negative regulation of extracellular matrix disassembly | 2.40E-02 | 2 |
| positive regulation of lipoprotein transport | 2.40E-02 | 2 |
| Factor XII activation | 2.40E-02 | 2 |
| late endosomal microautophagy | 2.40E-02 | 2 |
| micturition | 2.40E-02 | 2 |
| response to tumor necrosis factor | 2.50E-02 | 3 |
| response to unfolded protein | 2.60E-02 | 3 |
| cell motility | 2.60E-02 | 3 |
| response to cytokine | 2.60E-02 | 4 |
| regulation of heart rate | 2.70E-02 | 3 |
| negative regulation of endopeptidase activity | 2.70E-02 | 3 |
| regulation of cytokinesis | 2.80E-02 | 3 |
| establishment or maintenance of cell polarity | 2.90E-02 | 3 |
| negative regulation of supramolecular fiber organization | 3.00E-02 | 2 |
| response to triglyceride | 3.00E-02 | 2 |
| positive regulation of DNA binding | 3.00E-02 | 2 |
| positive regulation of type IIa hypersensitivity | 3.00E-02 | 2 |
| positive regulation of antibacterial peptide production | 3.00E-02 | 2 |
| response to nickel cation | 3.00E-02 | 2 |
| cellular heat acclimation | 3.00E-02 | 2 |
| defense response | 3.10E-02 | 3 |
| G2/M transition of mitotic cell cycle | 3.10E-02 | 3 |
| negative regulation of endothelial cell proliferation | 3.10E-02 | 3 |
| regulation of neuron projection development | 3.10E-02 | 3 |
| response to nutrient levels | 3.20E-02 | 5 |
| positive regulation of double-strand break repair via homologous recombination | 3.20E-02 | 3 |
| response to corticosterone | 3.20E-02 | 3 |
| positive regulation of cell migration | 3.20E-02 | 6 |
| negative regulation of apoptotic process | 3.20E-02 | 9 |
| cellular response to interferon-beta | 3.60E-02 | 3 |
| cell adhesion | 3.70E-02 | 7 |
| cellular response to hydrogen peroxide | 3.90E-02 | 4 |
| gene expression | 3.90E-02 | 6 |
| clathrin coat disassembly | 4.20E-02 | 2 |
| synaptic vesicle uncoating | 4.20E-02 | 2 |
| positive regulation of nucleotide-binding oligomerization domain containing 2 signaling pathway | 4.20E-02 | 2 |
| hemoglobin metabolic process | 4.20E-02 | 2 |
| vitamin D metabolic process | 4.20E-02 | 2 |
| negative regulation of cell growth | 4.50E-02 | 4 |
| response to immobilization stress | 4.50E-02 | 3 |
| positive regulation of vasoconstriction | 4.50E-02 | 3 |
| positive regulation of NF-kappaB transcription factor activity | 4.60E-02 | 4 |
| protein stabilization | 4.60E-02 | 5 |
| regulation of synaptic vesicle exocytosis | 4.70E-02 | 3 |
| lysosome organization | 4.70E-02 | 3 |
| regulation of cell cycle | 4.70E-02 | 5 |
| chaperone-mediated autophagy | 4.80E-02 | 2 |
| positive regulation of microtubule nucleation | 4.80E-02 | 2 |
| response to thyroxine | 4.80E-02 | 2 |
| response to glucocorticoid | 4.80E-02 | 4 |
| positive regulation of nitric oxide biosynthetic process | 4.80E-02 | 3 |
| negative regulation of cell adhesion | 4.80E-02 | 3 |
| positive regulation of MAPK cascade | 4.90E-02 | 5 |

Appendix Table 8 The biological process (BP) enrichment analysis of proteins with differential modifications in the 14-15-month-old group

| Pathways | P-Value | Count |
| --- | --- | --- |
| acute-phase response | 1.00E-15 | 12 |
| proteolysis | 1.90E-13 | 25 |
| complement activation, classical pathway | 7.30E-13 | 10 |
| zymogen activation | 2.70E-12 | 10 |
| fibrinolysis | 2.30E-08 | 6 |
| regulation of systemic arterial blood pressure | 2.40E-08 | 7 |
| response to nutrient | 4.50E-07 | 10 |
| negative regulation of blood coagulation | 1.30E-06 | 5 |
| blood coagulation | 1.30E-06 | 8 |
| negative regulation of proteolysis | 1.60E-06 | 6 |
| response to xenobiotic stimulus | 1.70E-06 | 16 |
| positive regulation of phagocytosis | 3.10E-06 | 7 |
| negative regulation of fibrinolysis | 7.70E-06 | 4 |
| response to estradiol | 1.00E-05 | 10 |
| complement activation | 1.80E-05 | 5 |
| positive regulation of receptor-mediated endocytosis | 2.10E-05 | 5 |
| regulation of transmembrane transporter activity | 2.60E-05 | 4 |
| positive regulation of bone resorption | 2.80E-05 | 5 |
| antibacterial humoral response | 3.00E-05 | 6 |
| lactate metabolic process | 3.60E-05 | 4 |
| intracellular phosphate ion homeostasis | 4.70E-05 | 4 |
| complement activation, alternative pathway | 4.70E-05 | 4 |
| embryo implantation | 5.60E-05 | 6 |
| cellular response to cytochalasin B | 1.10E-04 | 3 |
| negative regulation of activation of membrane attack complex | 1.10E-04 | 3 |
| response to mechanical stimulus | 1.30E-04 | 7 |
| pyruvate metabolic process | 1.40E-04 | 4 |
| intracellular iron ion homeostasis | 1.50E-04 | 6 |
| response to hypoxia | 1.50E-04 | 10 |
| peptide catabolic process | 1.70E-04 | 4 |
| protein targeting to lysosome involved in chaperone-mediated autophagy | 2.20E-04 | 3 |
| Factor XII activation | 2.20E-04 | 3 |
| regulation of iron ion transport | 2.20E-04 | 3 |
| animal organ regeneration | 3.00E-04 | 6 |
| response to lead ion | 3.30E-04 | 5 |
| regulation of norepinephrine uptake | 3.70E-04 | 3 |
| pyruvate catabolic process | 3.70E-04 | 3 |
| protein refolding | 5.20E-04 | 4 |
| collecting duct development | 5.50E-04 | 3 |
| regulation of transepithelial transport | 5.50E-04 | 3 |
| intracellular sodium ion homeostasis | 7.30E-04 | 4 |
| positive regulation of T cell mediated cytotoxicity | 7.60E-04 | 5 |
| positive regulation of oligodendrocyte progenitor proliferation | 7.60E-04 | 3 |
| protein localization to adherens junction | 7.60E-04 | 3 |
| vasodilation | 8.00E-04 | 5 |
| response to organic cyclic compound | 9.60E-04 | 9 |
| NAD metabolic process | 1.00E-03 | 3 |
| morphogenesis of a polarized epithelium | 1.30E-03 | 3 |
| kidney development | 1.50E-03 | 7 |
| cellular response to interleukin-6 | 1.60E-03 | 4 |
| negative regulation of lymphocyte proliferation | 1.60E-03 | 3 |
| cellular response to iron ion | 1.60E-03 | 3 |
| positive regulation of endothelial cell proliferation | 1.80E-03 | 5 |
| positive regulation of fibroblast proliferation | 1.80E-03 | 5 |
| chaperone-mediated protein folding | 1.90E-03 | 4 |
| cellular response to cadmium ion | 2.20E-03 | 4 |
| response to bacterium | 2.50E-03 | 7 |
| positive regulation of MAPK cascade | 2.70E-03 | 7 |
| skeletal muscle tissue development | 2.80E-03 | 5 |
| response to odorant | 2.80E-03 | 3 |
| plasminogen activation | 2.80E-03 | 3 |
| adherens junction assembly | 3.20E-03 | 3 |
| iron ion transport | 3.40E-03 | 4 |
| apical protein localization | 4.20E-03 | 3 |
| intracellular chloride ion homeostasis | 4.20E-03 | 3 |
| innate immune response | 4.50E-03 | 9 |
| amyloid-beta clearance | 4.80E-03 | 3 |
| digestion | 4.80E-03 | 3 |
| regulation of cyclin-dependent protein serine/threonine kinase activity | 4.80E-03 | 3 |
| inflammatory response | 4.90E-03 | 8 |
| negative regulation of cell adhesion | 5.40E-03 | 4 |
| response to estrogen | 5.80E-03 | 5 |
| positive regulation of G protein-coupled receptor signaling pathway | 5.90E-03 | 3 |
| ERK1 and ERK2 cascade | 6.50E-03 | 4 |
| cellular response to electrical stimulus | 6.60E-03 | 3 |
| retina development in camera-type eye | 6.70E-03 | 5 |
| keratinization | 7.10E-03 | 4 |
| cellular response to fluid shear stress | 7.20E-03 | 3 |
| cellular response to testosterone stimulus | 7.20E-03 | 3 |
| protein catabolic process | 9.60E-03 | 5 |
| intermediate filament organization | 9.80E-03 | 4 |
| response to lipopolysaccharide | 1.00E-02 | 7 |
| positive regulation of superoxide anion generation | 1.20E-02 | 3 |
| chaperone-mediated autophagy translocation complex disassembly | 1.20E-02 | 2 |
| slow axonal transport | 1.20E-02 | 2 |
| negative regulation of collateral sprouting of intact axon in response to injury | 1.20E-02 | 2 |
| protein transmembrane import into intracellular organelle | 1.20E-02 | 2 |
| positive regulation of protein refolding | 1.20E-02 | 2 |
| endocytic hemoglobin import into cell | 1.20E-02 | 2 |
| positive regulation of gene expression | 1.30E-02 | 10 |
| response to hydrogen peroxide | 1.30E-02 | 4 |
| muscle cell cellular homeostasis | 1.40E-02 | 3 |
| regulation of blood pressure | 1.40E-02 | 4 |
| postsynaptic actin cytoskeleton organization | 1.60E-02 | 3 |
| regulation of protein localization to plasma membrane | 1.60E-02 | 3 |
| regulation of synaptic vesicle endocytosis | 1.70E-02 | 3 |
| complement-dependent cytotoxicity | 1.80E-02 | 2 |
| positive regulation of lysosomal membrane permeability | 1.80E-02 | 2 |
| tissue regeneration | 1.90E-02 | 3 |
| response to vitamin D | 2.00E-02 | 3 |
| chaperone cofactor-dependent protein refolding | 2.00E-02 | 3 |
| positive regulation of phosphorylation | 2.20E-02 | 3 |
| liver development | 2.20E-02 | 5 |
| cellular response to follicle-stimulating hormone stimulus | 2.30E-02 | 3 |
| response to ethanol | 2.30E-02 | 6 |
| estrous cycle | 2.40E-02 | 3 |
| protein processing | 2.40E-02 | 4 |
| glucose catabolic process to lactate via pyruvate | 2.40E-02 | 2 |
| vitamin transport | 2.40E-02 | 2 |
| negative regulation of extracellular matrix disassembly | 2.40E-02 | 2 |
| micturition | 2.40E-02 | 2 |
| late endosomal microautophagy | 2.40E-02 | 2 |
| cell motility | 2.60E-02 | 3 |
| negative regulation of endopeptidase activity | 2.70E-02 | 3 |
| platelet activation | 2.70E-02 | 3 |
| establishment or maintenance of cell polarity | 3.00E-02 | 3 |
| response to heat | 3.00E-02 | 3 |
| response to triglyceride | 3.00E-02 | 2 |
| response to nickel cation | 3.00E-02 | 2 |
| negative regulation of supramolecular fiber organization | 3.00E-02 | 2 |
| positive regulation of antibacterial peptide production | 3.00E-02 | 2 |
| positive regulation of type IIa hypersensitivity | 3.00E-02 | 2 |
| androgen catabolic process | 3.00E-02 | 2 |
| carbohydrate metabolic process | 3.10E-02 | 4 |
| positive regulation of double-strand break repair via homologous recombination | 3.20E-02 | 3 |
| calcium ion homeostasis | 3.20E-02 | 3 |
| detection of chemical stimulus involved in sensory perception of bitter taste | 3.40E-02 | 3 |
| positive regulation of ATPase-coupled calcium transmembrane transporter activity | 3.60E-02 | 2 |
| positive regulation of estradiol secretion | 3.60E-02 | 2 |
| positive regulation of hydrolase activity | 3.60E-02 | 2 |
| intracellular calcium ion homeostasis | 3.70E-02 | 4 |
| cellular response to hydrogen peroxide | 4.00E-02 | 4 |
| spermatogenesis | 4.00E-02 | 8 |
| gene expression | 4.10E-02 | 6 |
| response to activity | 4.10E-02 | 4 |
| urate biosynthetic process | 4.20E-02 | 2 |
| positive regulation of apoptotic cell clearance | 4.20E-02 | 2 |
| regulation of systemic arterial blood pressure by renin-angiotensin | 4.20E-02 | 2 |
| clathrin coat disassembly | 4.20E-02 | 2 |
| positive regulation of extracellular matrix constituent secretion | 4.20E-02 | 2 |
| synaptic vesicle uncoating | 4.20E-02 | 2 |
| glucocorticoid metabolic process | 4.20E-02 | 2 |
| response to macrophage colony-stimulating factor | 4.20E-02 | 2 |
| hemoglobin metabolic process | 4.20E-02 | 2 |
| vitamin D metabolic process | 4.20E-02 | 2 |
| platelet aggregation | 4.30E-02 | 3 |
| positive regulation of vasoconstriction | 4.60E-02 | 3 |
| response to immobilization stress | 4.60E-02 | 3 |
| negative regulation of angiogenesis | 4.70E-02 | 4 |
| negative regulation of T cell proliferation | 4.80E-02 | 3 |
| positive regulation of CoA-transferase activity | 4.80E-02 | 2 |
| chaperone-mediated autophagy | 4.80E-02 | 2 |
| regulation of cell cycle | 4.80E-02 | 5 |
| positive regulation of nitric oxide biosynthetic process | 4.90E-02 | 3 |
| response to glucocorticoid | 4.90E-02 | 4 |
